# SAP30BP interacts with RBM17/SPF45 to promote splicing in a subset of human short introns

**DOI:** 10.1101/2022.12.30.522300

**Authors:** Kazuhiro Fukumura, Luca Sperotto, Stefanie Seuß, Hyun-Seo Kang, Rei Yoshimoto, Michael Sattler, Akila Mayeda

## Abstract

Human pre-mRNA splicing requires the removal of introns with highly variable lengths, from tens to over a million nucleotides. Therefore, mechanisms of intron recognition and splicing are likely not universal. Recently, we reported that splicing in a subset of human short introns with truncated polypyrimidine tracts depends on RBM17 (SPF45), instead of the canonical splicing factor U2AF heterodimer. Here, we demonstrate that SAP30BP, a factor previously implicated in transcriptional control, is an essential splicing cofactor for RBM17. *In vitro* binding and NMR analyses demonstrate that a U2AF-homology motif (UHM) in RBM17 binds directly to a newly identified UHM-ligand motif (ULM) in SAP30BP. We show that this RBM17–SAP30BP interaction is required to specifically recruit RBM17 to phosphorylated SF3B1 (SF3b155), a U2 snRNP component in active spliceosomes. We propose a unique mechanism for splicing in a subset of short introns, in which SAP30BP guides RBM17 in the assembly of active spliceosomes.

**Graphical abstract:** 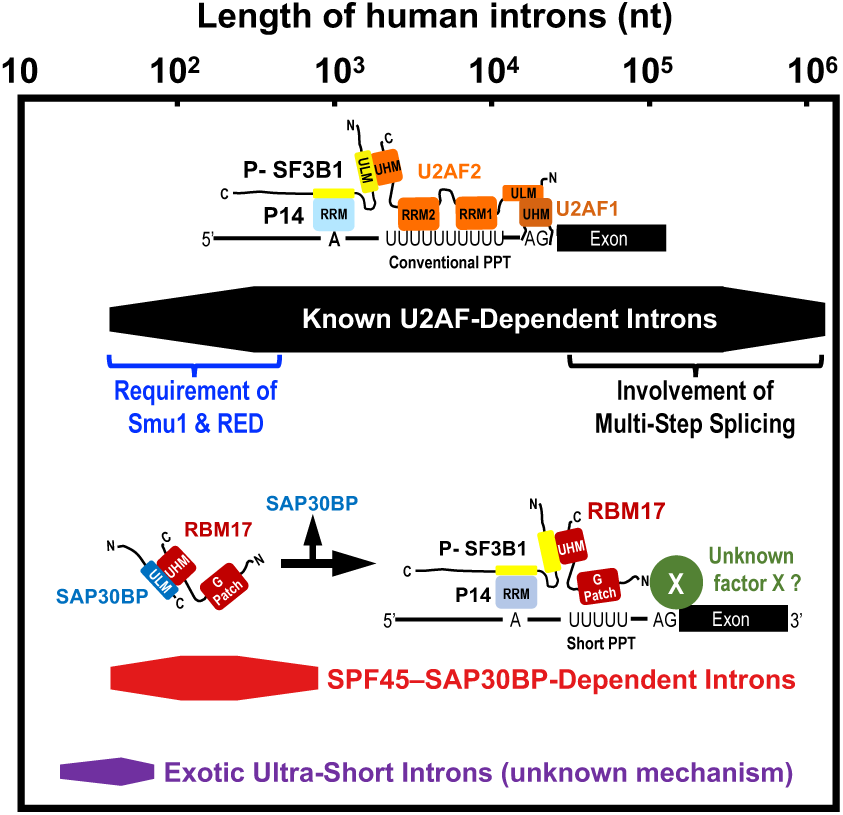

**In brief:** Fukumura et al. discover a general splicing mechanism in a subset of human short introns with truncated polypyrimidine tracts. This splicing reaction is mediated by intermediary RBM17–SAP30BP complex, instead of the known U2AF heterodimer. SAP30BP binding to RBM17 may support RBM17 association with active phosphorylated SF3B1 in U2 snRNP.

**Highlights:**

- RBM17 (SPF45) is a splicing factor required for a subset of human short introns
- SAP30BP is an essential cofactor, which interacts with RBM17 *via* UHM–ULM binding
- RBM17 forms a weak complex with SAP30BP before its binding with SF3B1 in U2 snRNP
- RBM17–SAP30BP complex supports RBM17 to be recruited to active phosphorylated SF3B1

## INTRODUCTION

Human pre-mRNA introns vary extensively in length, ranging from 30 nucleotides (nt) to 1,160,411 nt^1^. On the other hand, the known molecular mechanism of pre-mRNA splicing was biochemically established using ideal model pre-mRNAs with a single relatively short intron. The common model substrates are human β-globin pre-mRNA with 158-nt intron and adenovirus 2 major late (AdML) pre-mRNA with 231-nt intron, which are efficiently spliced *in vitro* and *in cellulo* (protocols in Ref.^2,3^).

Therefore, the known essential *cis*-acting splicing signals and the binding *trans*-acting splicing factors in the early spliceosomal complex were analyzed with such ideal model pre-mRNAs; i.e., the 5′ splice site, the branch-site (BS) sequence, and the poly-pyrimidine tract (PPT) followed by the 3′ splice site that are bound by U1 snRNP, U2 snRNP and U2AF2–U2AF1 (U2AF^65^–U2AF^35^), respectively (reviewed in Ref.^4,5^). This aspect was realized by our recent discovery; i.e., a subset of human short introns with truncated PPTs are spliced out by RBM17 (SPF45), but not by the known U2AF heterodimer^6^. Interestingly, the human splicing factors Smu1 and RED are involved in splicing targeting another distinct subset of short introns in which distances between the 5′ splice site and branch site are sufficiently short^7^.

In the RBM17-dependent splicing mechanism, the U2AF-homology motif (UHM) of RBM17 competes out that of U2AF2 for binding to the N-terminal UHM-ligand motif (ULM) of the U2 snRNP component, SF3B1 (SF3b155; Figure 1A), whose C-terminal HEAT domain provides the binding pocket for BS adenosine and SF3B1 contacts the intron region upstream and downstream of the BS^8-10^. Our nuclear magnetic resonance (NMR) spectroscopy data revealed that neither the UHM domain nor the G-patch motif of RBM17 binds the truncated PPT *in vitro*, while both of these are critical for RBM17 to be localized on the truncated PPT and for eventual RBM17-dependent splicing *in cellulo*^6^. Therefore, we postulate that the recruitment of RBM17 in the functional position on short introns requires an additional protein factor (‘Unknown factor X’ in Figure 1A).

**Figure 1.**
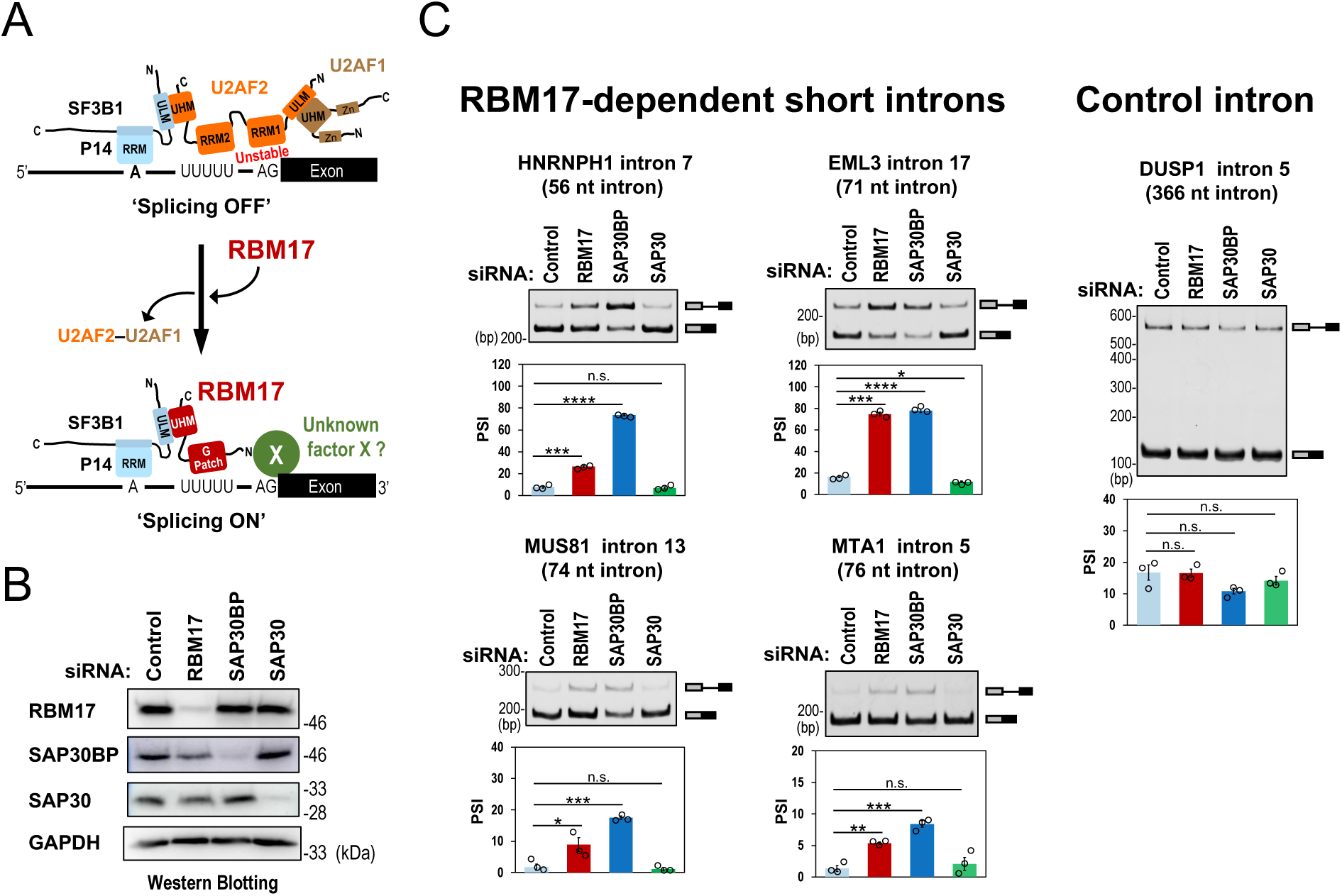
SAP30BP is additionally required for RBM17-dependent splicing of pre-mRNAs with short introns. **(A)** The model of RBM17-dependent splicing on a short intron with a truncated PPT (indicated by UUUUU)^6^. The associated splicing factors with the domain structures and the target sequences of pre-mRNAs are represented schematically. ‘Unknown factor X’ was postulated in this study. **(B)** The depletion of RBM17, SAP30BP, SAP30 proteins by siRNA-knockdown in HeLa cells. The depleted proteins were checked by Western blotting with indicated antibodies. **(C)** *In cellulo* splicing assays of the pre-mRNAs including indicated introns. After the indicated siRNA transfection in HeLa cells, endogenous splicing of the four RBM17-dependent short introns and one RBM17-independent control intron were analyzed by RT–PCR. Means ± SEM are given for three independent experiments and Welch’s *t*-test values were calculated (\**P* < 0.05, \*\**P* < 0.01, \*\*\**P* < 0.001, \*\*\*\**P* < 0.0001, ^n.s.^ *P* > 0.05).

Intriguingly, three previous studies have implicated SAP30BP as a promising potential RBM17-interacting cofactor: (i) SAP30BP (HCNGP) was detected as a non-snRNP protein in human spliceosomal A complexes^11^, (ii) the fruit fly homologs of SAP30BP and RBM17 were detected in spliceosomal complexes formed on a short intron (62-nt) but not on a long intron (147-nt)^12^, and (iii) yeast two-hybrid protein interaction assays showed that RBM17 can interact with SAP30BP (Table S3 in Ref.^13^). Here we demonstrate that SAP30BP is a functional partner in RBM17-dependent splicing. Interestingly, the binding of SAP30BP to RBM17 is important prior to the RBM17 interaction with phosphorylated SF3B1, a component of active U2 snRNP.

## RESULTS

### siRNA-knockdown of SAP30BP represses endogenous RBM17-dependent splicing

To test the functional role of SAP30BP in RBM17-dependent splicing, we performed siRNA-mediated depletions of SAP30B, RBM17 (positive control), and SAP30 (negative control) in HeLa cells (Figure 1B). Total RNAs were then analyzed by RT– PCR to examine RBM17-dependent splicing activity of the endogenous pre-mRNAs targeting the 56-nt (intron 7 of *HNRNPH1*), the 71-nt (intron 17 of *EML3*), the 74-nt (intron 13 of *MUS81*), and the 76-nt (intron 5 of *MTA1*); which were all previously identified as RBM17-dependently spliced pre-mRNAs^6^. These short introns were effectively repressed not only in RBM17-depleted cells but also in SAP30BP-depleted cells, whereas such splicing repression was not observed with the control pre-mRNA (366-nt intron 5 of *DUSP1*; Figure 1C). These data suggest that SAP30BP plays a key role in RBM17-dependent splicing but not in U2AF-dependent conventional splicing.

### SAP30BP and RBM17 are critical to splice out many short introns

To test whether SAP30BP is generally required for RBM17-dependent splicing, we performed whole-transcriptome sequencing (RNA-Seq) analysis in either RBM17-deficient or SAP30BP-deficient HeLa cells. The sequencing reads were mapped to the human genome reference sequence, and we identified total 2077 events of intron retention (i.e., splicing inhibition) in which 326 events overlapped with knockdown of RBM17 and SAP30BP; namely, more than half of the retained introns in RBM17-deficient cells were shared with those in SAP30BP-deficient cells (Figure 2A; see Supplementary Table S1 for the list of all 326 introns). Notably, the length distribution of these retained-introns in RBM17-depleted and SAP30BP-depleted cells is strongly biased towards shorter lengths compared to all introns in general (Figure 2B).

**Figure 2.**
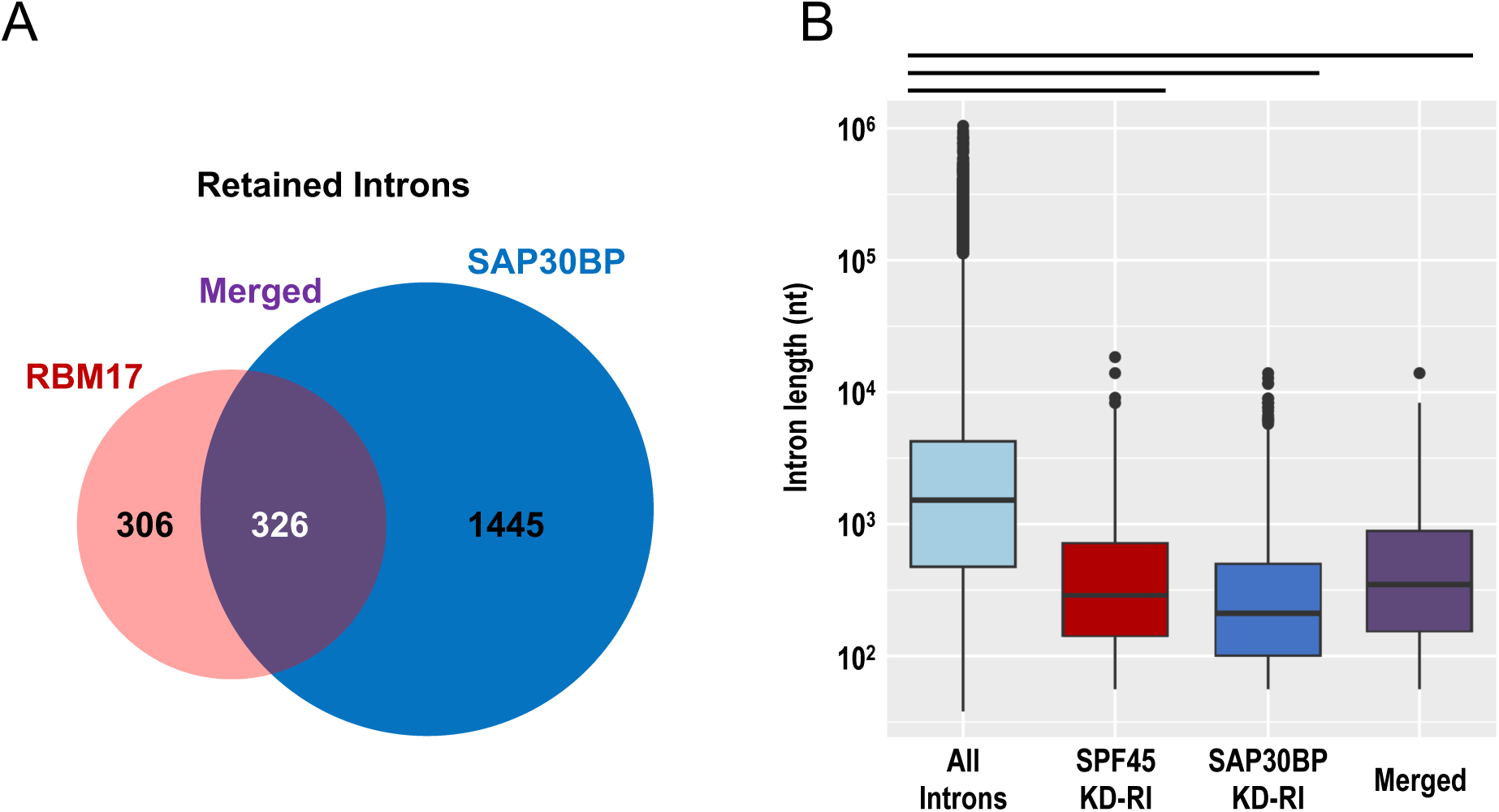
RBM17 and SAP30BP are generally required for the splicing of pre-mRNAs including short introns. **(A)** Venn diagram for significantly retained introns upon siRNA-knockdown of RBM17 and SAP30BP in HeLa cells. **(B)** Box-plots for the intron-length distributions of all introns in human RefGene and the retained introns (RI) in RBM17- and SAP30BP-knockdown (KD) HeLa cells. Box-plots indicate the summary of the dataset; median (middle line), 25–75th percentile (box), and <5TH and >95th percentile (whiskers), together with outliers (single points). Wilcoxon rank-sum test was used to calculate the statistical significance (indicated by three bars; all *p* < 2.2 × 10^-16^).

We compared 82 intron retention events in U2AF2-deficient HeLa cells (using RNA-Seq data^14^) with 1771 intron retention events in SAP30BP-deficient HeLa cells (Figure 2A). Then we found only 5 shared intron retentions, indicating that U2AF-dependent introns and SAP30BP-dependent introns are separate sets of introns.

Together, these results demonstrate that both RBM17 and SAP30BP ubiquitously function for efficient splicing of a distinct population of pre-mRNAs with short introns.

### RBM17 and SAP30BP function together in RBM17-dependent splicing

We recently demonstrated that a truncated PPT is a responsive *cis*-element for RBM17-dependent splicing. Therefore, we examined whether the SAP30BP-dependency is also related to the truncated PPT.

First, we confirmed that RBM17-dependent splicing, as well as SAP30BP-dependent splicing, is recapitulated in ectopically expressed mini-genes including short introns as in the corresponding endogenous genes (Figure 1C). The splicing in HNRNPH1 (56-nt intron) and EML3 (71-nt intron) pre-mRNAs was repressed by depletion of either RBM17 or SAP30BP, whereas splicing of the control adenovirus 2 major late (AdML) pre-mRNA (231-nt intron) was unaffected (Supplementary Figure S1A).

Next, we proceeded to analyze the role of the PPT using ectopically expressed chimeric mini-genes. Regarding the RBM17-dependency on the chimeric HNRNPH1 pre-mRNAs (the intronic fragments were replaced with those from control AdML pre-mRNA), we observed results consistent with our recent report^6^; i.e., (i) the replacement of the truncated 13-nt PPT of HNRNPH1 with the conventional 25-nt PPT of AdML switched the pre-mRNA to RBM17-independent splicing (AdML PPT25; Supplementary Figure S1B), (ii) the truncation of this AdML PPT to 13-nt regained RBM17-dependency (AdML PPT13), and (iii) the expansion of the upstream intron region between the 5′ splice site and the branch site with a short HNRNPH1 PPT retained RBM17 dependency (AdML 5′MT). Remarkably, knockdown of SAP30BP exactly follows the above RBM17-dependency (Supplementary Figure S1B). All these results were also recapitulated with another chimeric EML3 pre-mRNA with truncated 12-nt PPT (Supplementary Figure S1C). Our global PPT length analysis of the retained introns in SAP30BP-knockdown cells showed similar PPT length distribution as those in RBM17-knockdown cells (Supplementary Figure S2).

We conclude that RBM17 and SAP30BP act together in RBM17-dependent splicing, specifically for short introns with truncated PPTs.

### RBM17 binds to SAP30BP through UHM–ULM interaction

The cooperation of RBM17 and SAP30BP in splicing of pre-mRNAs with short introns suggest a direct interaction of these two factors *in cellulo*. To test this hypothesis, HeLa cells were co-transfected with Myc-SAP30BP and FLAG-RBM17 expression plasmids, immunoprecipitated with anti-Flag antibody, and the co-precipitated proteins were detected by anti-Flag or anti-Myc antibody. The immunoblot showed that RBM17 bound to SAP30BP and SF3B1 (positive control), but not to SAP30 (negative control), suggesting the formation of a RBM17–SAP30BP complex (Figure 3A).

**Figure 3.**
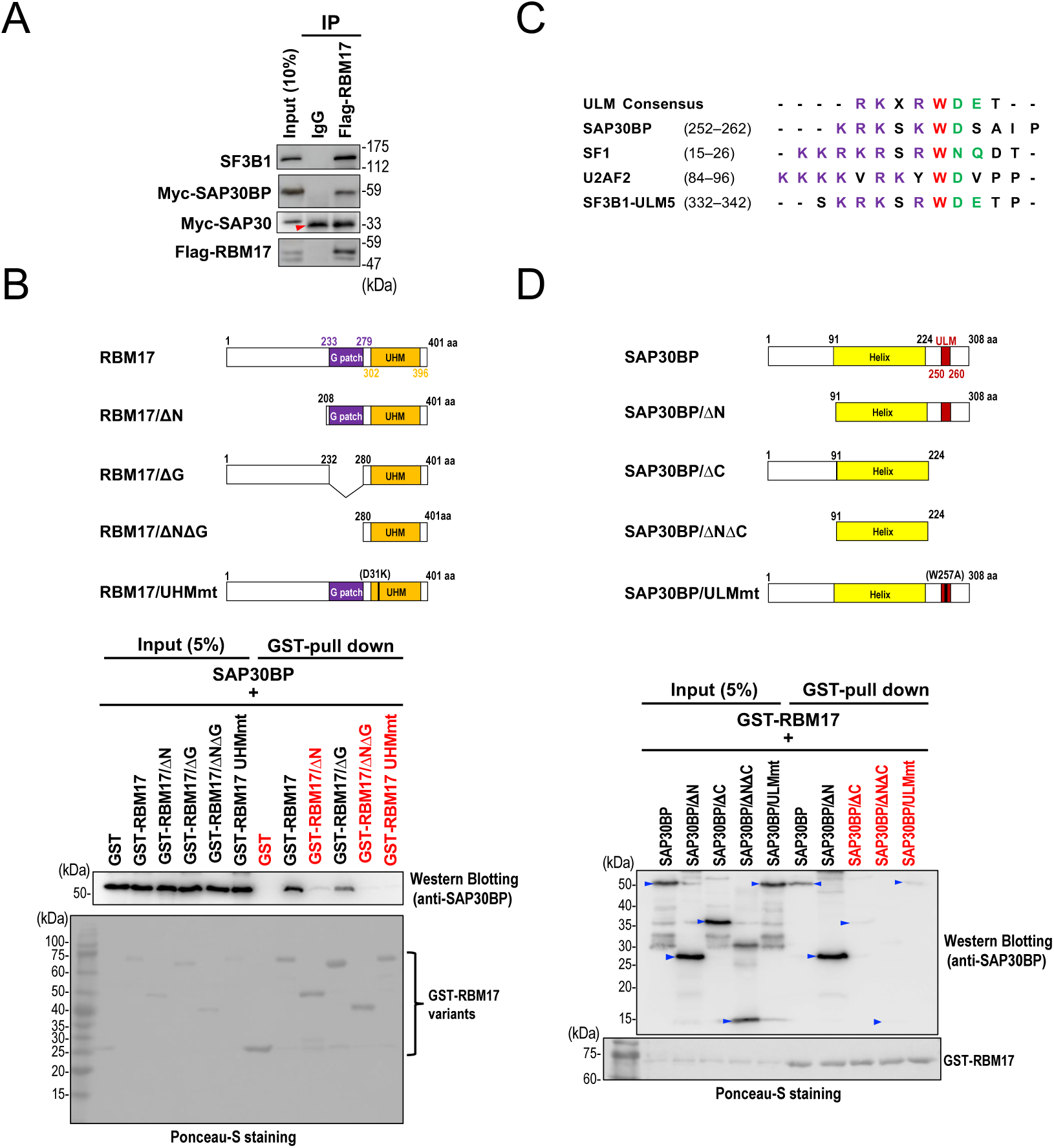
UHM in RBM17 and newly found ULM in SAP30BP are critical for the binding between RBM17 and SAP30BP. **(A)** Co-immunoprecipitation of RBM17 with SAP30BP and SF3B1. HeLa cells were co-transfected with Myc-SAP30BP, Myc-SAP30 (negative control) and Flag-RBM17 plasmids, these transfected cell extracts were immunoprecipitated using mouse IgG or anti-Flag antibody, and the immunoprecipitated proteins were analyzed by Western blotting. The red arrow head indicates IgG light chain of anti-Flag antibody. **(B)** *In vitro* GST-pulldown assay (after RNase A-treatment) to detect the direct binding between indicated RBM17 variant proteins and SAP30BP protein. Red-colored variants indicate the diminished protein binding with SAP30BP. Recombinant SAP30BP protein associated with these GST-fused proteins was detected by Western blotting using anti-SAP30BP antibody. The same membrane was also stained with ponceau S to evaluate the loading amount and the comparable precipitation of the GST or GST fusion proteins. **(C)** Alignment of ULM sequences in SAP30BP, SF1, U2AF2 and SF3B1 (ULM5) with the ULM consensus sequence. Conserved amino acid residues are denoted by colors: PURPLE–basic residues preceding the conserved tryptophan (W); RED–conserved W; GREEN–acidic and N/Q-type residues following the conserved W. **(D)** *In vitro* GST-pulldown assay (with RNase A-treatment) to detect the direct binding between indicated SAP30BP variants (indicated by blue arrow heads) and RBM17. Red-colored variants indicate the diminished protein binding with RBM17. See (B) for the methods.

Next, we investigated which regions in RBM17 and SAP30BP are required for complex formation. We prepared *E. coli* recombinant glutathione S-transferase (GST)-fusion proteins of RBM17, SAP30BP, and various deletion and mutation constructs (Figures 3B, 3D; upper schemata). Our GST pull-down assays revealed that deletion of the N-terminal region, including or excluding the G-patch motif (RBM17/ΔNΔG, RBM17ΔN), greatly reduced binding to SAP30BP (lower panel). While deletion of the G-patch motif alone (RBM17/ΔG) did not affect the SAP30BP interaction. Deletion of either half of the N-terminal domain (RBM17/ΔN1, RBM17/ΔN2) strongly reduced binding to SAP30BP indicating that the complete N-terminal domain contributes to the SAP30BP interaction (Supplementary Figure S3). However, the complete or middle part of the N-terminal domain alone (RBM17/N, RBM17/N3) showed no binding to SAP30BP (Supplementary Figure S3). The C-terminal region of RBM17 harbors the UHM domain, and a D319K mutation in the UHM (RBM17/UHMmt), which inhibits ULM binding^6^, greatly reduced the SAP30BP interaction. These data suggest that a ULM–UHM interaction is important for the complex formation. Together, we conclude that the RBM17–UHM is essential to interact with SAP30BP and that the N-terminal domain of RBM17 might additionally contribute to SAP30BP binding.

Consistent with the RBM17–UHM-mediated binding of SAP30BP, we indeed identified a ULM sequence in the C-terminal region of SAP30BP, which could mediate the interaction with the RBM17–UHM (Figure 3C; Figure 3D, upper schemata). Deletion of the C-terminal region of SAP30BP alone or both N- and C-terminal regions (SAP30BP/ΔC, SAP30BP/ΔNΔC) abrogated binding to RBM17, while deletion of the N-terminal domain alone (SAP30BP/ΔN) had no effect on the RBM17 interaction (lower panel). A W257A mutation in the ULM of SAP30BP (SAP30BP/ULMmt), which impairs UHM binding^15^, disrupted binding to RBM17 (Figure 3D, lower panel), confirming that the UHM–ULM interaction is critical for the RBM17–SAP30BP complex.

### NMR analysis confirmed the RBM17–SAP30BP interaction

To obtain a residue-level resolution mapping of the RBM17–SAP30BP interaction, we performed NMR titration experiments by testing the binding of ^15^N-labeled RBM17– UHM (RBM17/ΔNΔG) to the C-terminal SAP30BP fragment that harbors the ULM (SAP30BP/ΔNΔH, Figure 4A). Significant amide chemical shift changes confirm the UHM–ULM interaction *via* the RBM17–UHM domain (Figure 4B, Supplementary Figure S4). The most affected area includes the two α-helices and residues close to the characteristic Arg-Tyr-Phe (RYF) motif (important for ULM binding) of the UHM, indicating a canonical UHM–ULM interaction (Figure 4B, D). Isothermal titration calorimetry (ITC) shows a binding affinity with a dissociation constant *K*_d_ = 34.4 µM (Figure 4C). Of note, this is somewhat lower than other reported UHM–ULM interactions^16^. In particular, the affinity is about one order of magnitude lower than the interaction of the RBM17–UHM domain with SF3B1–ULM5^16^. Nevertheless, we could verify the binding between endogenous RBM17 and SAP30BP *in cellulo* (Supplementary Figure S5).

**Figure 4.**
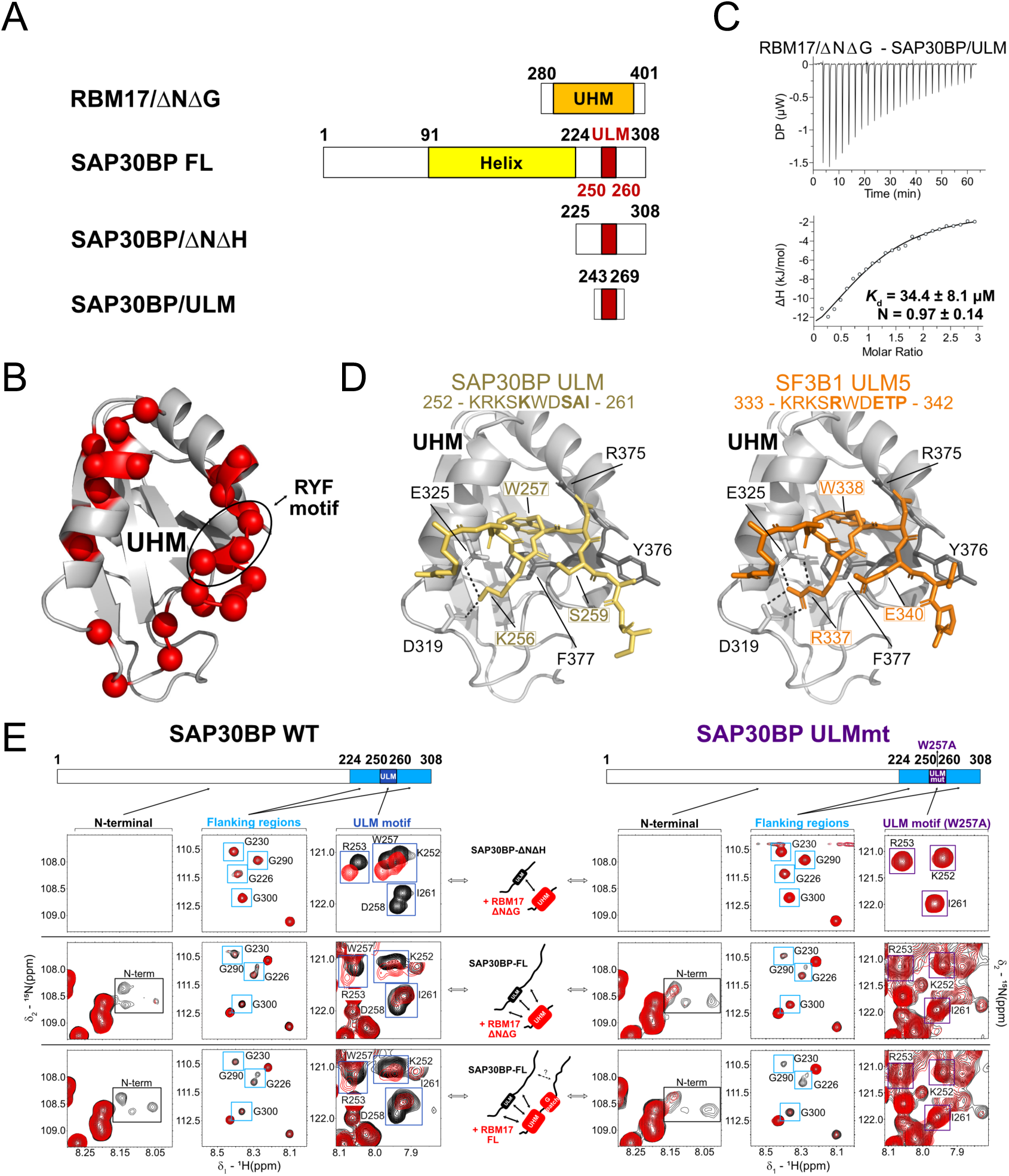
The RBM17–SAP30BP interaction was confirmed by NMR and isothermal titration calorimetry (ITC) **(A)** RBM17 and SAP30BP variant constructs used for NMR and ITC experiments. **(B)** NMR chemical shift perturbation (>0.65 ppm) of RBM17/ΔNΔG, upon adding 6-fold molar excess of SAP30BP/ΔNΔH, is highlighted on RBM17–UHM structure (red spheres). The position of the Arg-Tyr-Phe (RYF)-motif is indicated. **(C)** ITC binding analysis (isotherm) of RBM17/ΔNΔG and SAP30BP/ULM. **(D)** Crystal structure of SF3B1–ULM5 bound to RBM17–UHM (right, PDB accession code 2PEH) and homology model of the SAP30BP–ULM peptide bound to the RBM17–UHM (left). Differences in amino acid sequences are shown in bold. The RYF motif (R375-Y376-F377) of RBM17–UHM and other key interface residues are labelled. **(E)** Comparison of ^1^H-^15^N HSQC NMR spectra of SAP30BP (ΔNΔH or full-length, in black) with (left) or without (right) mutation in ULM upon addition of RBM17 (ΔNΔG or full-length, in red). Constructs are schematically indicated (center). Representative NMR signals belonging to the N-terminal (black square), flanking (cyan square), and ULM (blue square for wild-type, violet square for ULMmt) regions in SAP30BP are displayed.

To analyze which differences in the ULMs of SF3B1 and SAB30BP could explain the reduced affinity, we performed homology modeling based on the known crystal structure of RBM17–UHM bound to SF3B1–ULM5^16^. Considering the sequence similarity of the two ULM sequences, we replaced the ligand by the SAP30BP–ULM (Figure 4D). Analysis of the model complex with the RBM17–UHM and SF3B1– ULM5 indicates that key interactions are preserved, however, with a few exceptions. For example, Arg337 (preceding the important Trp in the SF3B1–ULM) is substituted by Lys256 in SAP30BP (R→K in Figure 3C). While this maintains electrostatic interactions with Asp319 and Glu325 in the RBM17–UHM, hydrogen bond interactions are likely reduced (Figure 4D). Also, the presence of Ser259 in SAP30BP instead of Gln340 in SF3B1 (E→S in Figure 3C) removes a negative charge that interacts with a positively charged region on the surface of the UHM. Taken together, the amino acid differences may rationalize the lower binding affinity of the SAP30BP–ULM. This is consistent with our observation that mechanistically the SF3B1–ULM likely displaces SAP30BP for the binding to the eventual RBM17–UHM.

The GST pull-down assays suggest a contribution of the N-terminal region of RBM17 for the RBM17–SAP30BP interaction (Figure 3B). We therefore performed NMR titration experiments with full-length proteins to identify additional contacts between the two proteins involving their N-terminal regions. To this end, we monitored NMR spectra of three different ^15^N-labeled SAP30BP constructs upon titration with unlabeled RBM17 variants; namely, SAP30BP/ΔNΔH with RBM17/ΔNΔG, full-length SAP30BP with RBM17/ΔNΔG, and full-length SAP30BP with full-length RBM17 (Figure 4E, left). NMR chemical shift changes of the SAP30BP amide signals were monitored using ^1^H-^15^N correlation experiments. The experiments with the full-length proteins confirmed the expected spectral changes for residues in the ULM sequence. Spectral changes were not observed for residues in the ULM anymore upon mutation of the key Trp residue in the ULM sequence that is critical for the UHM–ULM interaction (Figure 4E, right). However, additional spectral changes were observed for residues flanking the ULM sequence and in the N-terminal region of SAP30BP upon binding to RBM17 (Figure 4E, left). These spectral changes were still observable when mutating the Trp in the ULM (Figure 4E, right), confirming additional RBM17–SAP30BP interactions outside the ULM sequence.

These results, together with the data from the GST pull-down assays, demonstrate that the N-terminal regions of RBM17 and SAP30BP also contribute to the interaction between the two proteins.

### RBM17–SAP30BP interaction plays a role to recruit phosphorylated SF3B1

Lastly, we investigated the functional role of the SAP30BP in the context of the following RBM17 binding to SF3B1. Since RBM17 has only one UHM, the RBM17– SAP30BP interaction has to be released prior to the RBM17 interaction with SF3B1 (Figure 1A). This scenario is supported by the stronger binding affinity of RBM17 with SF3B1 compared to that with SAP30BP (see above section). This raises the question: what is the role of the RBM17–SAP30BP interaction? Previously, it was shown that SF3B1 is phosphorylated in the active spliceosome *in vitro*^17-19^. SF3B1 phosphorylated at Thr313, mostly localizes in the chromatin fraction, whereas free unphosphorylated SF3B1 is predominantly found in the nucleoplasm^20^. We therefore postulate that SAP30BP prevents RBM17 binding to inactive unphosphorylated SF3B1 and promotes its binding to active phosphorylated SF3B1.

To examine this hypothesis, we took advantage of an anti-phospho-SF3B1 antibody in our GST pull-down assay (Figure 5A). We found that the depletion of SAP30BP markedly decreased RBM17 binding to phospho-SF3B1 (lanes 4 and 8), indicating that RBM17 alone binds to the excess of unphosphorylated SF3B1, which results in a decrease of the RBM17 association with active phospho-SF3B1. Since other phosphorylation sites are known in the SF3B1–ULM region, we performed this GST pull-down assay with another anti-phosphoSF3B1 antibody(Thr211) (Supplementary Figure S6). We obtained essentially same data as in Figure 5A. These results suggest that a SAP30BP interaction with RBM17 might shield the RBM17–UHM binding surface to prevent binding with inactive free SF3B1–ULM (Figure 6).

**Figure 5.**
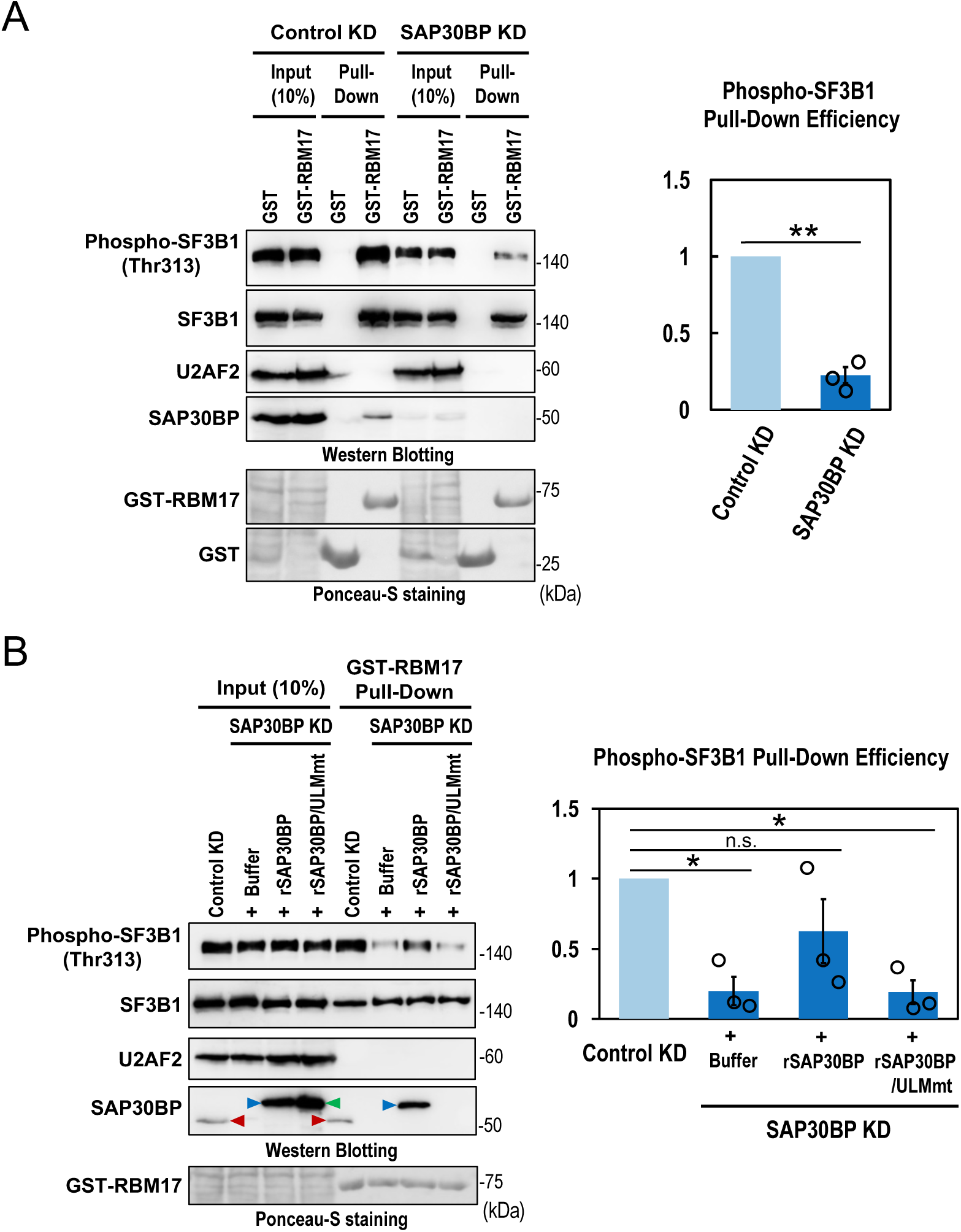
Binding of RBM17 to SAP30BP is critical for RBM17 to be recruited in active phospho-SF3B1. **(A)** Binding of RBM17 to total SF3B1 and phosphorylated SF3B1 under SAP30bp-depletion. GST pull-down assay with whole cell extracts prepared from SAP30BP-knockdown HeLa cells. Proteins associated with GST-RBM17 protein were detected by Western blotting using the indicated antibodies. Membrane was also stained with Ponceau S for loading and precipitation controls. The bar graph shows quantification of the bands on Western blots. Phospho-SF3B1 scanned data were normalized to SF3B1 (Phospho-SF3B1/SF3B1) and plotted. Means ± SEM are given for three independent experiments and Welch’s *t*-test values were calculated (\*\**P* < 0.01). **(B)** Rescue experiments of diminished RBM17–phospho-SF3B1 binding under SAP30BP-depletion. The same reaction mixtures in panel (A) were supplemented with recombinant SAP30BP or SAP30BP/ULMmt proteins and the associated proteins with GST-RBM17 were analyzed by Western blotting with indicated antibodies. See (A) for plotted scanned data and the statistical analysis (\**P* < 0.05, ^n.s.^ *P* > 0.05).

**Figure 6.**
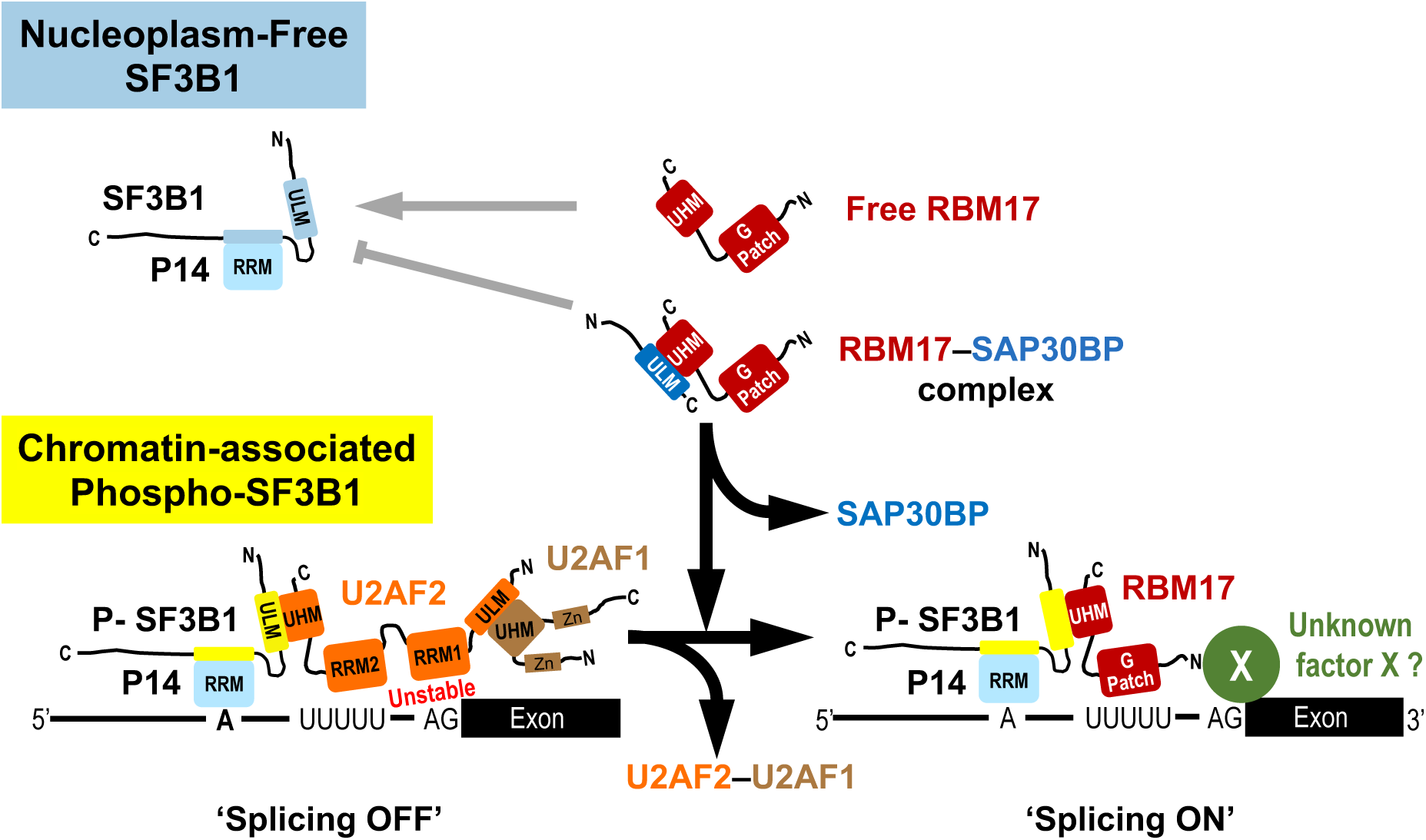
The model of RBM17–SAP30BP complex-dependent splicing on a short intron with a truncated PPT. We assume that RBM17 alone can bind with free unphosphorylated SF3B1 in the nucleoplasm, while intermediary RBM17–SAP30BP complex prevents this non-functional binding, which allows functional RBM17 binding to active phosphorylated SF3B1 on pre-mRNA. The associated splicing factors with the domain structures and the target sites of pre-mRNAs are represented schematically. The bold black arrows indicate validated pathways, whereas the narrow gray arrows indicate speculative pathways based on our GST pull-down assays (Figure 5).

To confirm the functional role of an intermediary RBM17–SAP30BP complex, we performed rescue experiments *in vitro* with SAP30BP-depleted HeLa cell extracts supplemented by two recombinant proteins: rSAP30BP and rSAP30BP/ULMmt (Figure 5B). rSAP30BP rescued the binding to phospho-SF3B1 in SAP30BP-depleted extracts, however the rSAP30BP/ULMmt, which hardly bind to RBM17 (Figure 3D), did not (Figure 5B, ‘Pull-Down’ panel; compare with control ‘Buffer’ lane). We conclude that the intermediary RBM17–SAP30BP complex is important for RBM17 to be recruited to active phosphorylated SF3B1 (Figure 6).

## DISCUSSION

SAP30BP was first identified and cloned as HTRP (human transcription regulator protein), which is a cellular co-repressor induced by HSV1 infection and it interacts with SAP30 associated with histone deacetylase complex (HDAC)^21^. The transcriptional co-repressors, SAP30BP and HDAC, interact with BAHCC1 to control the modified histone H3 (H3K27me3)-mediated gene repression^22^. Here we identified SAP30BP as a functional cofactor of RBM17-dependent splicing in short introns. Therefore SAP30BP adds to the growing list of factors with dual roles in transcription and pre-mRNA splicing. Consistent with our results, a recent study revealing an important role of murine Sap30bp in promoting splicing of short introns and suggested the functional association of Sap30bp in the control of nucleocytoplasmic transport of transcripts associated with the nervous system development (J. Roth, A.-C. Gingras and B.J. Blencowe, personal communication). Another study also observed that knockdown or degradation of human SAP30BP induces many intron retention events and SAP30BP was identified as a CDK11 activator that regulates splicing globally (C. Wang and H. Cheng, personal communication).

### A discovered important UHM–ULM interaction in RBM17-dependent splicing

UHM–ULM-mediated protein–protein interactions; e.g., U2AF2–SF1, U2AF2–SF3B1, and U2AF1–U2AF2, play essential roles for spliceosome assembly in constitutive splicing^23-25^ (reviewed in Ref.^26^). UHM–ULM interactions are also involved in alternative splicing regulation; e.g., RBM17–SF3B1^16^, PUF60–SF3B1^27^ and RBM39 (CAPERα)–SF3B1^28^. Recently, we demonstrated that the SF3B1–RBM17 complex (*via* ULM–UHM binding) replaces the SF3B1–U2AF complex (*via* ULM–UHM binding) on the truncated PPT of a subset of short introns^6^.

Here, we discovered a so far unknown ULM–UHM interaction in atypical constitutive splicing targeting short introns. We show that the RBM17–UHM binds to a newly identified ULM in SAP30BP prior to the recruitment of RBM17 to active SF3B1 on pre-mRNA. The fact that the UHM–ULM interaction in the RBM17–SAP30BP complex is significantly weaker than other UHM–ULM interactions, as demonstrated by our ITC data (Figure 4C), is consistent with a release of SAP30BP from RBM17 upon SF3B1 binding. The SAP30BP–ULM may thus occupy and shield the RBM17–UHM to avoid undesired interactions with other ULMs. And finally, SAP30BP–ULM is replaced by SF3B1–ULM, which has an order of magnitude stronger binding affinity.

The weak and reversible, albeit functional and critical, ULM–UHM interaction between RBM17 and SAP30BP thus provides a unique splicing reaction not known previously.

### A conceivable role of RBM17–SAP30BP binding prior to interaction with U2 snRNP

Why does RBM17 need to bind SAP30BP before its interaction with SF3B1 in U2 snRNP? Several data provide clues to address this question. Previous *in vitro* spliceosome analyses demonstrated that SF3B1 is phosphorylated during spliceosome activation, typcally from B to B^act^ complex^17-19,29,30^. We therefore paid attention to the phosphorylation status of SF3B1 that interacts with RBM17. Our *in vitro* binding assay revealed that RBM17 can bind to SF3B1 anyway, but without SAP30BP, RBM17 does not bind well to active phosphorylated SF3B1. Importantly, most phosphorylated SF3B1 is present in spliceosomes detected in the chromatin fraction, whereas free unphosphorylated SF3B1 predominantly exists in the nucleoplasmic fraction^20^. We thus hypothesize that SAP30BP might shield and sequester the UHM domain of RBM17 to prevent its association with free unphosphorylated SF3B1 in the nucleoplasm. And thereby it will ensure that RBM17 can be recruited to active phosphorylated SF3B1 on pre-mRNA (Figure 6).

The composition of spliceosomes and their assembly in RBM17-dependent splicing has not been studied, and thus it is unclear whether the U2 snRNP (SF3B1)–RBM17 complex formation occurs in A complex or the following B complex, as the U2 snRNP (SF3B1)–U2AF interaction occurs in the A complex, and afterwards RBM17 replaces U2AF (Figure 6). In either case, our model suggests that the phosphorylation of SF3B1 for spliceosome activation, in RBM17-dependent splicing occurs somewhat earlier than in conventional U2AF-dependent splicing. The exact timing of SF3B1 phosphorylation during the RBM17-dependent splicing process remains to be determined. It will also be interesting to investigate if and how phosphorylation of SAP30BP may affect its structural interactions with RBM17 and other binding partners in this process.

### 3′ splice site recognition in RBM17-dependent splicing

We have not yet identified the responsible factor of the 3′ splice site recognition in the early spliceosomes of RBM17-dependent short introns, such as U2AF1 in U2AF-dependent conventional introns (‘Unknown factor X’ in Figure 6). An involvement of U2AF1 was ruled out by the observation of active RBM17-dependent splicing upon co-depletion of U2AF2 and U2AF1^6^. Since the RBM17 G-patch motif is critical for RBM17-dependent splicing^6^, we speculate that such a ‘Factor X’, if any, interacts with RBM17 through the G-patch motif to recognize the 3′ splice site. Our ongoing studies aim to identify the putative ‘Factor X’.

The point mutation in the 3′ splice site ‘AG’ has no effects on U2AF-dependent splicing reaction until the first step of splicing in C complex (Ref.^31^ and references therein). Therefore in RBM17-dependent splicing, it is also possible that 3′ splice site recognition may also occur later in the splicing process. We have not yet identified the recognition timing of the 3′ splice site ‘AG’ during the RBM17-dependent splicing cycle.

Previous studies uncovered diversity in recognition of the 3′ splice site. Interestingly, it was reported that excess levels of the SR protein, SRSF2 (SC35), can functionally substitute for U2AF2 in a substrate-specific manner^32^, although the mechanism of this U2AF-independent splicing remains unknown. Indeed, genome-wide analysis of U2AF–RNA interactions suggests that ∼12% of functional 3′ splice sites do not require U2AF2 binding or even its function for their recognition^14^. We are convinced that the 3′ splice sites of the RBM17-dependent introns must be one category of such U2AF2-independent 3′ splice sites.

### Limitations of the study

We proposed the model of RBM17–SAP30BP complex-dependent splicing acting in a subset of human introns with truncated PPT (Figure 6). We have not yet obtained definitive evidence to prove this model, and therefore, we cannot rule out the possibility of alternative models. We do not yet know whether SAP30BP directly promotes association of RBM17 with phosphorylated SF3B1.

Our data (Figure 2A) apparently indicated that there is a SAP30BP-dependent but not RBM17-dependent splicing. Since UHM–ULM interactions play an important role in splicing (reviewed in Ref.^26^), SAP30BP-ULM interactions with UHM of other splicing factors are also possible. Indeed, SAP30BP interactions with splicing regulators PUF60 and RBM39, as well as RBM17, were detected by yeast two-hybrid screening^13^.

## EXPERIMENTAL PROCEDURES

### Construction of expression plasmids

The mini-gene plasmids (pcDNA3-AdML, pcDNA3-HNRNPH1 and pcDNA3-EML3), their chimeric variants (pcDNA3-HNRNPH1/AdML-PPT25, pcDNA3-HNRNPH1/AdML-PPT13, pcDNA3-HNRNPH1/5′AdML, pcDNA3-EML3/AdML-PPT25, pcDNA3-EML3/AdML-PPT13 and pcDNA3-EML3/5′AdML), and the RBM17 expression plasmids (pcDNA3-Flag-RBM17, pGEX6p2-RBM17, pGEX6p2-RBM17/ΔG and pGEX6p2-RBM17/UHMmt) were constructed previously^6^.

To construct the deleted variants (pGEX6p2-RBM17/ΔN, pGEX6p2-RBM17/ΔN1, pGEX6p2-RBM17/ΔN2, pGEX6p2-RBM17/ΔNΔG pGEX6p2-RBM17/N and pGEX6p2-RBM17/N3), the corresponding ORF regions were PCR-amplified from the parent plasmid pcDNA3-Flag-RBM17 with specific primer sets (Supplementary Table S2) and subcloned into the pGEX6p2 vector (GE Healthcare Life Sciences). To construct the series of SAP30BP and SAP30 expression plasmids (pcDNA3-Myc-SAP30BP, pGEX6p2-SAP30BP, pGEX6p2-SAP30BP/ΔN, pGEX6p2-SAP30BP/ΔC, pGEX6p2-SAP30BP/ΔNΔC and pcDNA3-Myc-SAP30), the corresponding ORF regions were PCR-amplified from HeLa cells cDNA with specific primer sets (Supplementary Table S2) and subcloned into the pcDNA3-Myc or pGEX6p2 vectors (GE Healthcare Life Sciences). Overlap extension PCR was performed to induce the mutation in the ULM of pcDNA3-Myc-SAP30BP (pGEX6p2-SAP30BP/ULMmt).

### Expression and purification of recombinant proteins

Various recombinant RBM17 and SAP30BP proteins were expressed from pETM11 vectors with His6-ProteinA TEV cleavable tag using *E. coli* BL21(DE3) in LB medium or in M9 medium supplemented with ^15^NH_4_Cl or ^15^NH_4_Cl and ^13^C-D-Glucose. Protein expression was induced at OD_600_ around 0.8–1.0 with 1 mM isopropyl-β-D-thiogalactopyranoside (IPTG), followed by overnight expression at 20°C. Cells were resuspended in 50 mM Tris-HCl (pH 8.0) buffer with 500 mM NaCl, 10 mM imidazole and protease inhibitors, and lysed using French press and sonicator. After centrifugation, the cleared lysates were purified on a Ni-NTA resin column. The protein samples were further purified by ion exchange chromatography, according to their pI, on a 5 mL HiTrap SP/Q HP column (Cytiva) with 20 mM sodium phosphate (pH 6.5) and by size-exclusion chromatography on a HiLoad 16/60 Superdex 75 column (GE Healthcare Life Sciences) with 20 mM sodium phosphate (pH 6.5) buffer containing 150 mM NaCl. The tag was cleaved with TEV protease and removed with a Ni-NTA column as the final step for SAP30BP/ULM, or before ion exchange chromatography for all other constructs (Figure 4A).

### Western blotting analyses

Western blotting was performed as previously described^6^. Briefly, protein samples were boiled with ‘NuPAGE LDS Sample Buffer’ (Thermo Fisher Scientific), separated by SDS-polyacrylamide gel electrophoresis (SDS-PAGE). The gel was stained with Ponceau S (Cell Signaling Technology), if necessary, and immuno-reactive protein bands were detected by the ECL system and visualized by imaging analyzer (ImageQuant LAS 500, GE Healthcare Life Sciences).

The following commercially available antibodies were used to detect targeted proteins: anti-RBM17 (1:1500 dilution; HPA037478, Sigma-Aldrich), anti-SF3B1 (1:3000 dilution; D221-3, MBL Life Science), anti-Phospho-SF3B1(Thr313) (1:1500 dilution; D8D8V, Cell Signaling Technology), anti-Phospho-SF3B1(Thr211) (1:1500 dilution; PA5-105427, Thermo Fisher Scientific), anti-U2AF2 (1:1500 dilution; U4758, Sigma-Aldrich), anti-SAP30BP (1:1500 dilution; HPA052943, Sigma-Aldrich), anti-SAP30 (1:1500 dilution; 27679-1-AP, Proteintech), anti-GAPDH (1:3000 dilution; M171-3, MBL Life Science), anti-Flag (1:3000 dilution; anti-DDDDK tag, M185-3L, MBL Life Science) and anti-cMyc 9E10 antibodies (1:1500 dilution; sc-40, Santa Cruz Biotechnology). Immuno-reactive protein bands were detected by the ECL system and visualized by imaging analyzer (ImageQuant LAS 500, GE Healthcare Life Sciences).

### Human cell line culture

HeLa cells (ATCC) were cultured in Dulbecco’s modified Eagle’s medium (Wako) supplemented with 10% fetal bovine serum (Sigma-Aldrich) and 1% penicillin-streptomycin (Nakarai Tesque). Cells were grown at 37°C in 5% CO_2_.

### siRNA knockdown and splicing assays

siRNA-mediated knockdown in HeLa cells was performed as previously described^6^. The siRNAs targeting RBM17, SAP30BP and SAP30 were purchased from Fasmac (Supplementary Table S2 for the sequences). At 72 h post-transfection, each protein depletion was checked by Western blotting with indicated antibodies.

Endogenous splicing products derived from the *HNRNPH1*, *EML3*, *MUS81*, MTA1 and *DUSP1* genes were analyzed by RT–PCR with oligo-dT, random primers and specific primer sets (Supplementary Table S2) as previously described^6^.

To analyze splicing products derived from mini-genes, RBM17- and SAP30BP-depleted HeLa cells were transfected at 48 h post-transfection with 0.5 µg of mini-gene plasmid (Supplementary Table S2) using lipofectamine 2000 reagent (Invitrogen–Thermo Fisher Scientific). These cells were incubated for 24 h prior to the extraction of RNAs described previously^6^. To analyze splicing products from mini-genes, RT–PCR was performed with T7 primer and a specific primer set for each mini-gene (Supplementary Table S2).

All the oligonucleotide primers were purchased from Fasmac (Supplementary Table S2). The PCRs were performed with Blend Taq polymerase (Toyobo Life Science), and the PCR products were analyzed by 6% PAGE and quantified as described previously^6^. All the experiments were independently repeated three times and the means and standard errors of the splicing efficiencies were calculated.

### High-throughput RNA sequencing (RNA-Seq) analyses

Six independent total RNAs were prepared by ‘NucleoSpin RNA’ kit (Macherey-Nagel): these are derived from HeLa cells, treated with two control siRNAs (universal negative control siRNA, Nippon gene), with two RBM17-targeted siRNAs and with two SAP30BP-treated siRNAs. Then enrichment of Poly(A) mRNA was performed with the ‘NEBNext Poly(A) mRNA Magnetic Isolation Module’ (New England Biolabs). RNA libraries were prepared using the ‘NEBNext Ultra Directional RNA Library Prep Kit for Illumina’ (New England Biolabs). These samples were sequenced on the high-throughput sequencing platform (NovaSeq6000, Illumina) using a 150 bp paired-end strategy. cDNA library construction and RNA sequencing were performed by a company (Rhelixa).

The sequencing data was analyzed as previously described^33^. Obtained sequence reads were mapped onto the human genome reference sequences (hg19) using the HISAT2 version 2.2.1^34^; the mapped sequence read were assembled using StringTie version 2.1.7^35^, and the changes of alternative splicing isoforms were analyzed using rMATS version 4.1.1^36^ as previously described^6^. Obtained splicing events were defined as significant changes when the false discovery rate (FDR) was calculated at less than 0.05.

PPT score and length were calculated by SVM-BP finder software^37^ as previously described^33^. The raw data from the RNA-Seq analysis have been deposited in the SRA database (https://www.ncbi.nlm.nih.gov/sra) under accession number GSE220906.

### Immunoprecipitation assays

HeLa cells (in 100-mm dishes) were co-transfected with pcDNA3-Flag-RBM17 and pcDNA3-Myc-SAP30BP (or pcDNA3-Myc-SAP30 as a negative control) using Lipofectamine 2000 reagent (Thermo Fisher Scientific). At 48 h post-transfection, harvested cells were suspended in buffer D [20 mM HEPES-NaOH (pH 7.9), 50 mM KCl, 0.2 mM EDTA, 20% (v/v) glycerol] and sonicated for 20 sec with an ultrasonic disrupter (UR-20P, Tomy Digital Biology). The lysate was centrifuged to remove debris and the supernatant was rocked at 4°C for 2 h with an anti-Flag antibody (anti-DDDDK tag, MBL Life Science) or mouse anti-IgG antibody (Santa Cruz Biotechnology) that was conjugated with ‘Dynabeads Protein A’ (Veritas) in NET2 buffer [50 mM Tris-HCl (pH7.5), 150 mM NaCl, 0.05% Triton-X100]. The beads were washed five times with NET2 buffer, and the associated proteins were detected by Western blotting (described above) with anti-SF3B1, anti-cMyc, and anti-Flag antibodies.

For immunoprecipitation experiments with endogenous proteins, HeLa cell lysate was rocked at 4°C for 16 h with anti-SAP30BP antibody (HPA052943, Sigma-Aldrich) or rabbit anti-IgG antibody (PM035, MBL) that was conjugated with Dynalbeads Protein A (Veritas) in NET2 buffer. The beads were washed five times with NET2 buffer, and the associated proteins were detected by Western blotting (described above) with indicated antibodies.

### Immunofluorescence confocal microscopic assays

HeLa cells were fixed with 3% formaldehyde in phosphate-buffered saline (PBS), permeabilized with 0.1% Triton X-100 in PBS, blocked with 2% bovine serum albumin (BSA) in PBS, and then incubated with the following primary antibodies in 2% BSA in PBS for 0.5 h; anti-SAP30BP (1:500 dilution; OTI4A9, ThermoFisher Scientific), anti-SAP30BP (1:500 dilution; HPA052943, Sigma-Aldrich) aiti-RBM17 (1:500 dilution; HPA037478, Sigma-Aldrich), anti-SF3B1 (1:500 dilution; D221-3, MBL Life Science) and anti-phosph-SF3B1(Thr313) (1:500 dilution; D8D8V, Cell Signaling Technology). After washing three times with PBS, cells were incubated with Alexa Fluor 488 or Alexa Fluor 555 secondary antibodies (1:500 dilution; ThermoFisher Scientific), and then washed three times with PBS. The nucleus in cells were counter-stained with 4’, 6-diamidino-2-phenylindole (DAPI). The images were analyzed by a confocal laser microscope (LSM710, Carl Zeiss). Quantification of Pearson’s correlation coefficient was calculated by Fiji (National Institute of Health).

### GST pull-down assays

Recombinant GST-tagged proteins (GST-RBM17, GST-RBM17/ΔN, GST-RBM17/ΔN1, GST-RBM17/ΔN2, GST-RBM17/N, GST-RBM17/N3, GST-RBM17/ΔG, GST-RBM17/ΔNΔG, GST-RBM17/UHMmt, GST-SAP30BP, GST-SAP30BP/ΔN, GST-SAP30BP/ΔC, GST-SAP30BP/ΔNΔC or GST-SAP30BP/ULMmt) were expressed in *E. coli* ‘BL21-CodonPlus (DE3)-RIPL’ competent cells (Stratagene–Agilent) and purified as previously described^6^. To remove the GST-tag from GST-SAP30BP proteins (GST-SAP30BP, GST-SAP30BP/ΔN, GST-SAP30BP/ΔC, GST-SAP30BP/ΔNΔC or GST-SAP30BP/ULMmt), ‘PreScission Protease’ (GE Healthcare Life Sciences) was used according to the manufacturer’s protocol.

For *in vitro* binding assays with the recombinant proteins (Figure 3, Supplementary Figure S3), GST-RBM17 variant proteins (20 µg) were mixed with untagged SAP30BP variant proteins (1 µg) in NET2 buffer supplemented with 0.5 mM dithiothreitol (DTT) and incubated at 4°C for 1 h. Then 20 µL of Glutathione Sepharose 4B (GE Healthcare Life Sciences) was added and incubated at 4°C for another 2 h. The incubated beads were washed five times with NET2 buffer and boiled with ‘NuPAGE LDS Sample Buffer’ to analyze SAP30BP binding by Western blotting with anti-SAP30BP antibody (described above).

For *in vitro* binding assays in HeLa cell extracts (Figure 5), HeLa cells (in 100-mm dishes) were transfected with 500 pmol SAP30BP siRNA or negative control siRNA (Nippon gene) using Lipofectamine RNAiMAX (Invitrogen–Thermo Fisher Scientific) according to the manufacturer’s protocol. At 72 h post-transfection, transfected HeLa cells were suspended in Buffer D and sonicated for 20 sec twice (described above) and centrifuged to remove debris. Obtained siRNA-treated HeLa whole cell extracts (200 µL) were mixed with recombinant GST-RBM17 proteins (20 µg) and incubated at 30°C for 15 min. The SAP30BP or SAP30BP/ULMmt recombinant proteins (1 µg) were added to the above cell extract mixtures to perform rescue experiments (Figure 5B). After RNase A (10 µg, Nippon Gene) treatment at 30°C for 5 min, NET2 buffer was added to a final volume of 500 µL with 20 µL of Glutathione Sepharose 4B and incubated at 4°C for 3 h. The incubated beads were washed five times with NET2 buffer and boiled with ‘NuPAGE LDS Sample Buffer’ to analyze SF3B1 binding by Western blotting (described above) with indicated antibodies.

### NMR spectroscopy

NMR experiments were recorded at 298 K on 500/600/900-MHz Bruker Avance NMR spectrometers equipped with cryogenic triple resonance gradient probes. Backbone ^1^H, ^15^N and ^13^C resonances were assigned with standard triple resonance experiments (HNCO, HN(CA)CO, HNCACB and CBCA(CO)NH; described in Ref.^38^). NMR spectra were processed by TOPSPIN3.5 (Bruker) or NMRPipe^39^ and analyzed using NMRFAM-SPARKY (T. D. Goddard and D. G. Kneller, SPARKY 3, University of California, San Francisco)^40^. Samples were measured at 0.6–0.8 mM protein concentration for triple resonance experiments and 50–100 μM protein concentration for ^1^H-^15^N HSQC in NMR buffer [20 mM sodium phosphate (pH 6.5), 150 mM NaCl, 2 mM DTT] with 10% D_2_O added as lock signal.

### ITC binding analysis

Binding affinities of RBM17/ΔNΔG with SAP30BP/ULM (Figure 4A) were measured using a MicroCal PEAQ-ITC (Malvern Panalytical) in 20 mM sodium phosphate (pH 6.5), 150 mM NaCl at 25°C. RBM17 protein in the cell (concentration ranges, 60–100 μM) was titrated with SAP30BP/ULM (concentration of 800–900 μM) for the final molar ratios of 1:1.7 to 1:3, adjusted depending on the affinity interaction. The standard deviation was determined for 3 replicates.

### Quantification and statistical analysis

Mean values, SEM values, and Welch’s *t*-test were calculated using Excel (Microsoft). Wilcoxon rank-sum test was performed in R software (The R Foundation for Statistical Computing).

## ACKNOWLEDGMENTS

We thank Drs. B. Blencowe and Hong Cheng for sharing results prior to publication; Dr. A. R. Krainer for critical reading of the manuscript; H. Shirasaki and T. Kanehisa for technical support; and members of our labs for constructive discussions. We thank Sam Asami and Gerd Gemmecker for support with NMR experiments and acknowledge access to NMR measurements at the Bavarian NMR center in Garching.

K.F. was partly supported by Grants-in-Aid for Scientific Research (C) [Grant number: 18K07304] from the Japan Society for the Promotion of Science (JSPS), a Research Grant from the Hori Sciences and Arts Foundation, and a Research Grant from Nitto Foundation, a Research Grant from Takeda Science foundation, and a Research Grant from Aichi Cancer Research Foundation. A.M. was partly supported by Grants-in-Aid for Scientific Research (B) [Grant number: JP16H04705] and for Scientific Research (C) [Grant number: JP21K06024] from JSPS. L.S. and M.S. acknowledge support by the Marie Skłodowska-Curie Innovative Training Network (MSCA-ITN) RNAct research of the European Union’s Horizon 2020 program [Grant number: 813239]. M.S. acknowledges funding by the Deutsche Forschungsgemeinschaft (DFG, German Research Foundation) with grants [SPP1935 project number 273941853 and SFB1035 project number 201302640].

## AUTHOR CONTRIBUTIONS

K.F. and A.M. conceived and designed the experiments; K.F. performed most of the experiments and organized the data; R.Y. performed bioinformatics analyses of the sequencing data; H.-S.K., S.S, L.S. and M.S. performed NMR and ITC experiments and data analysis; A.M. drafted the manuscript and edited the manuscript together with K.F. and M.S. All authors read, corrected and approved the final manuscript. A.M. coordinated and supervised the whole project.

## DECLARATION OF INTERESTS

The authors declare no competing interests.

## Supplementary Information

**Table S1.** Uploaded in a separate Excel File

**Table S2.**
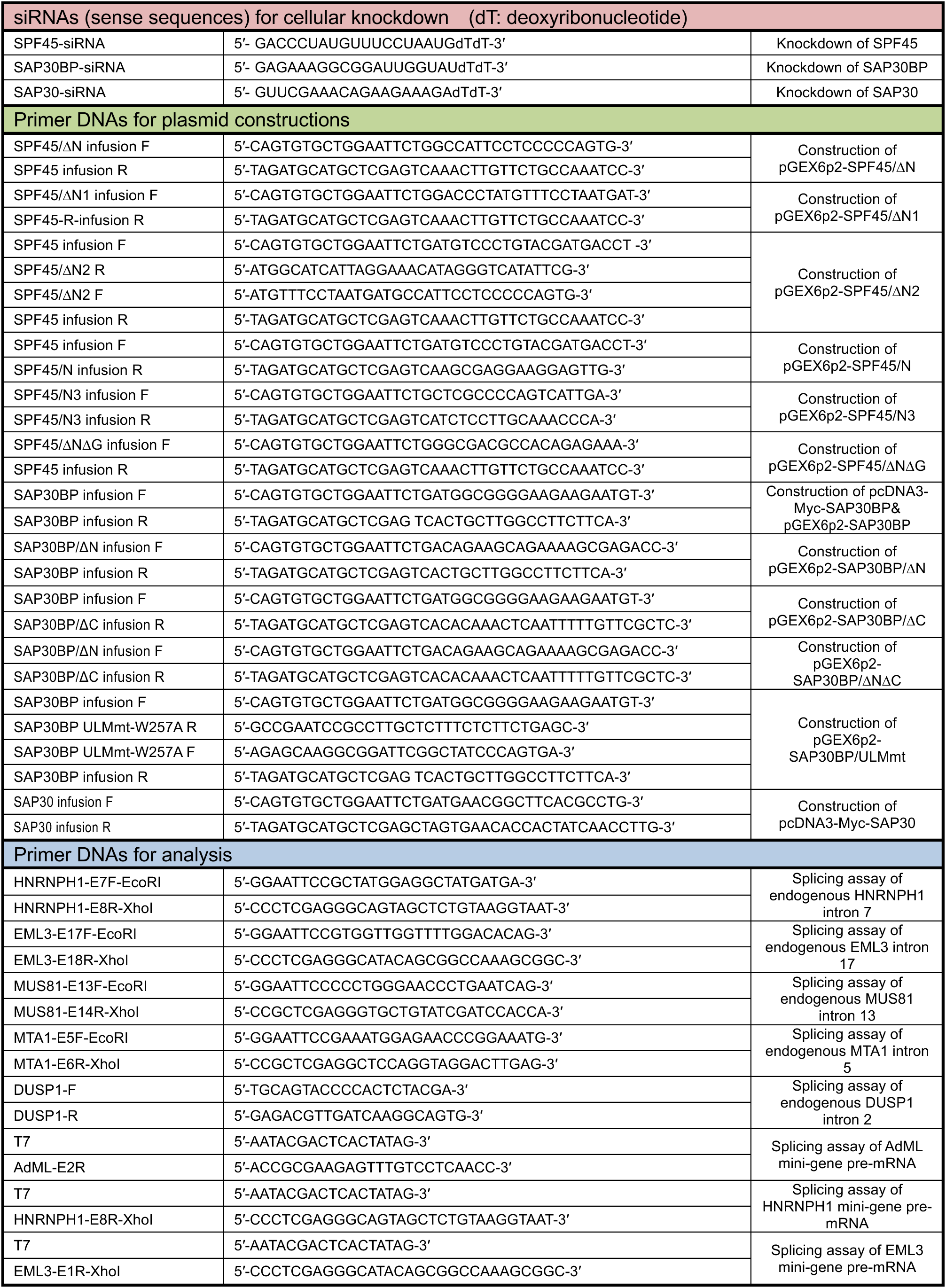
List of synthetic oligonucleotides used in the experiments (Related to STAR⋆Methods)

**Figure S1.**
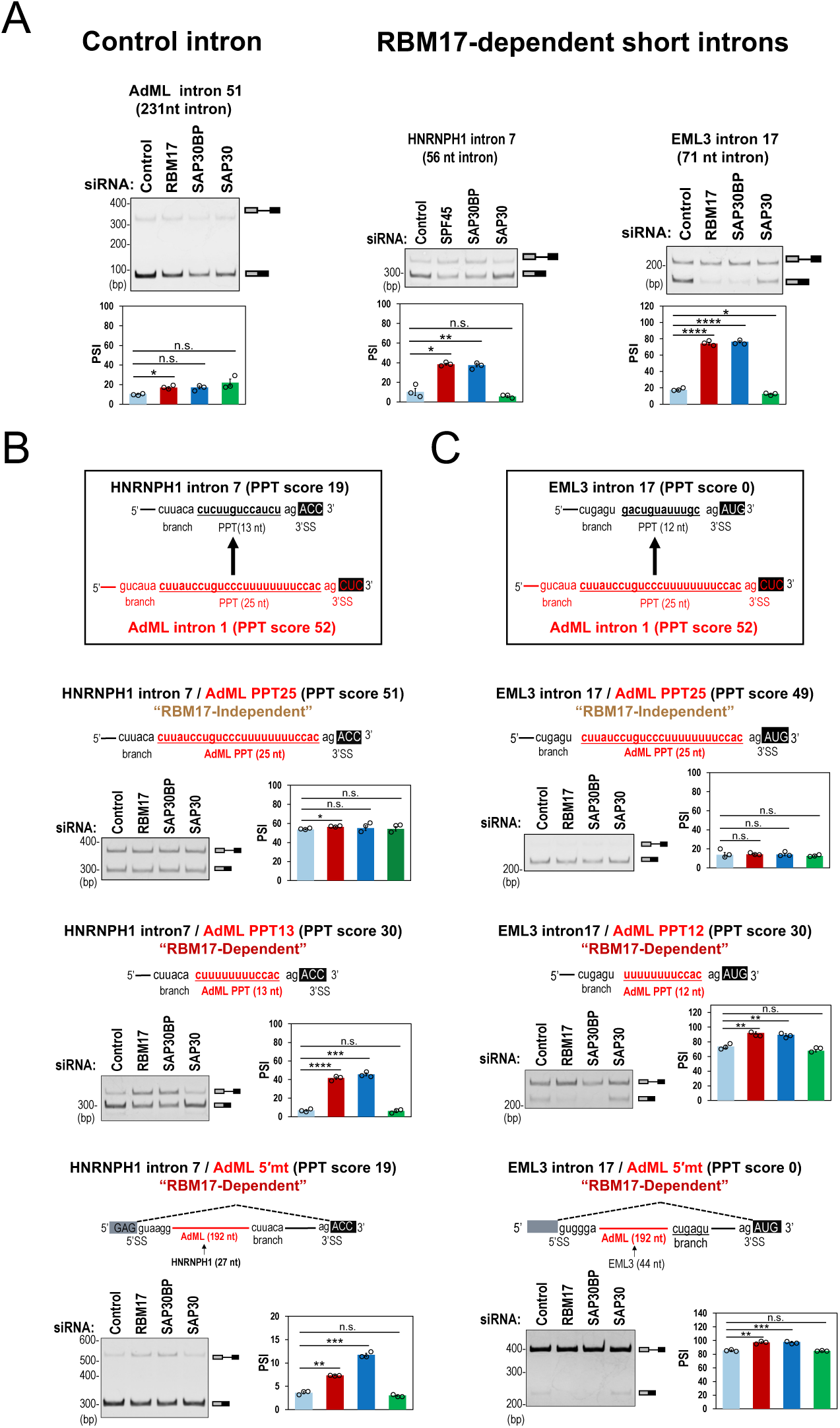
The length of poly-pyrimidine tracts (PPTs) determines the co-dependency of RBM17 and SAP30BP in splicing (Related to Figure 1) **(A)** Selectively repressed splicing of pre-mRNAs with short introns in RBM17- and SAP30BP-depleted cells. Indicated three pre-mRNAs were expressed from mini-genes in HeLa cells treated with RBM17, SAP30BP or SAP30 (negative control) siRNA and their splicing was assayed by RT–PCR. PAGE images and graphs of PSI (percent spliced in) values are shown. Means ± SEM are given for three independent experiments and Welch’s *t*-test values were calculated (\**P* < 0.05, \*\**P* < 0.01, \*\*\**P* < 0.001, \*\*\*\**P* < 0.0001, ^n.s.^ *P* > 0.05). **(B, C)** Selectively repressed splicing of the chimeric HNRNH1 and EML3 pre-mRNAs with truncated PPT in RBM17- and SAP30BP-depleted cells. These chimeric pre-mRNAs are schematically shown (red color indicates AdML derived sequences). The PPT score (see Methods) is one of the criteria to evaluate effective PPTs. See (A) for *in cellulo* splicing assay and the statistical analysis.

**Figure S2.**
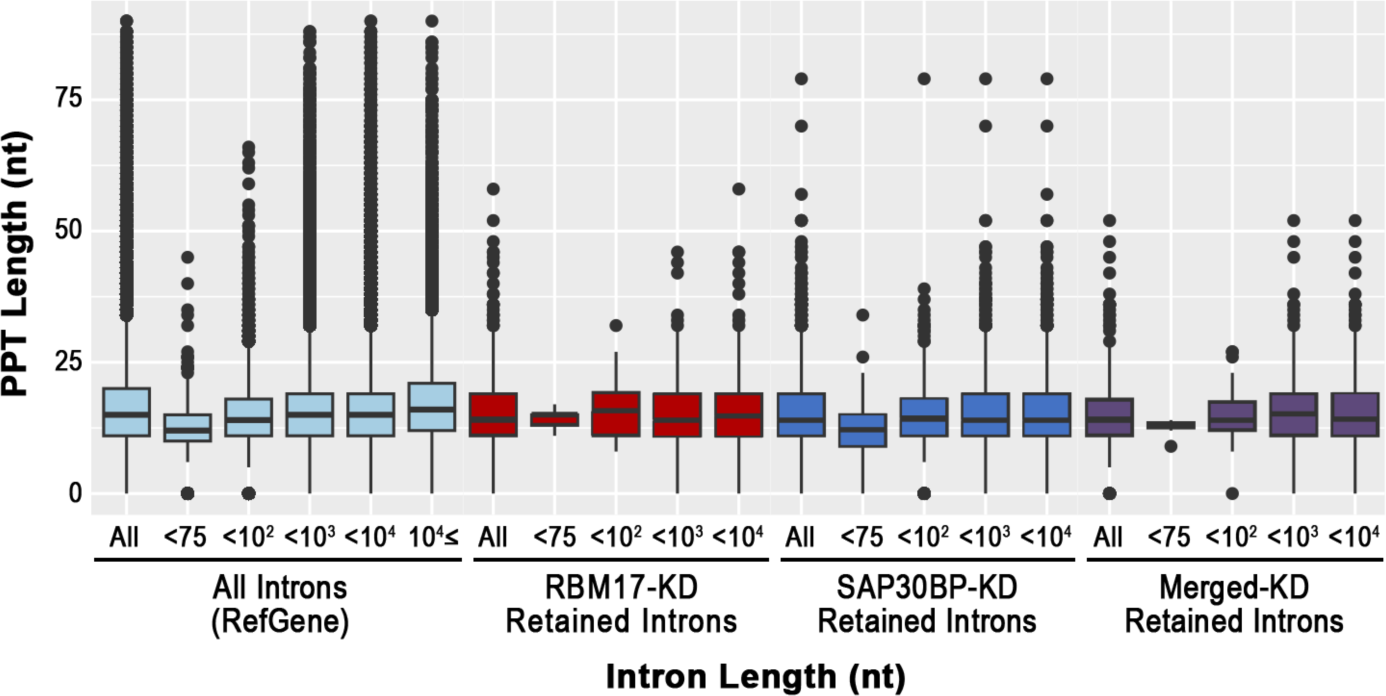
Truncated polypyrimidine tract (PPT) is dominated in introns shorter than 75 nt (Related to Figure 2) The box plots show the similar PPT-length distributions of all introns (in human RefGene) with those of the retained introns in RBM17- and SAP30BP-knockdown HeLa cells, suggesting that shorter introns generally tend to possess the shorter PPTs. The same analysis in RBM17-knockdown HEK293 cells were described previously (see Ref.^6^ in the text), in which PPT lengths of RBM17-dependent short introns were shorter than those of the whole RefGene introns. Box plots show the summary of the data set; median (middle line), 25–75th percentile (box), and <5TH and >95th percentile (whiskers), together with outliers (single points).

**Figure S3.**
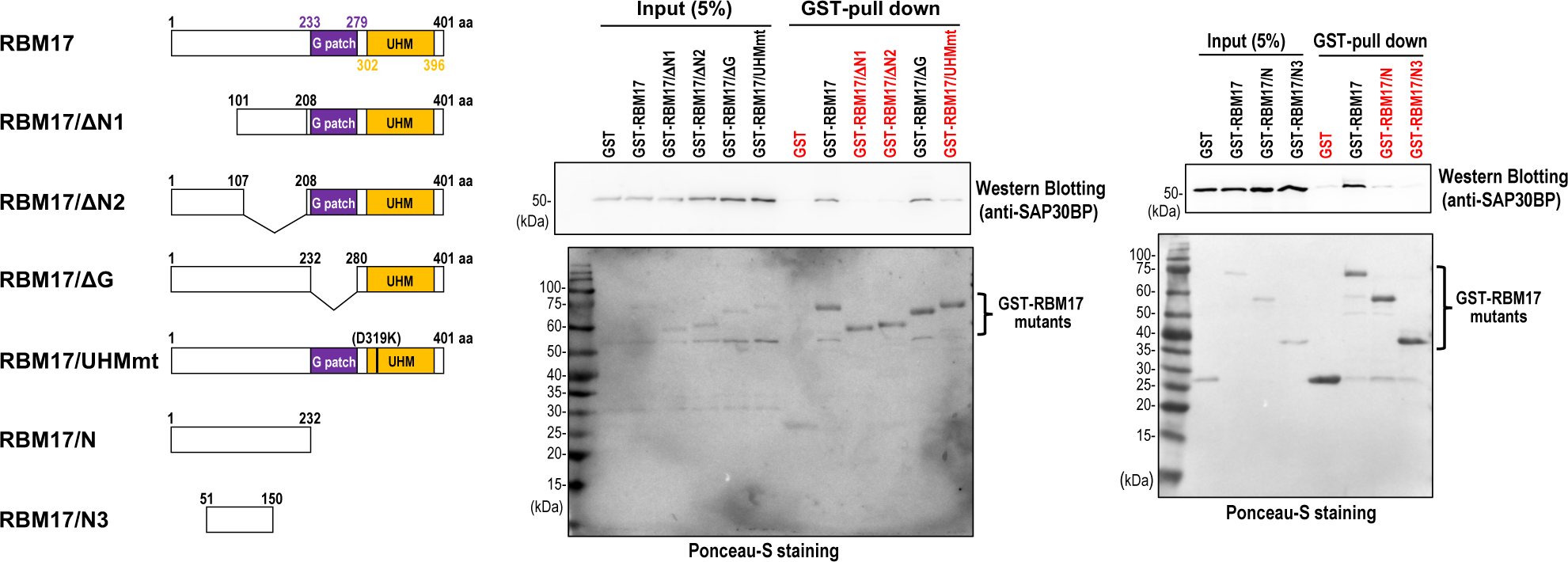
UHM and full N-terminal region in RBM17 are important for the SAP30BP binding (Related to Figure 3) Direct interaction of RBM17 mutants with SAP30BP, as shown by immunoblot analysis of the *in vitro* GST-pulldown assay for purified GST, GST-RBM17/DG, RBM17/UHMmt,GST-RBM17/DN1, GST-RBM17/DN2 or RBM17/N of RBM17/N3 with purified SAP30BP. Membrane was also stained by ponceau S to show the general loading as well as the comparable precipitation of the GST or GST fusion proteins.

**Figure S4.**
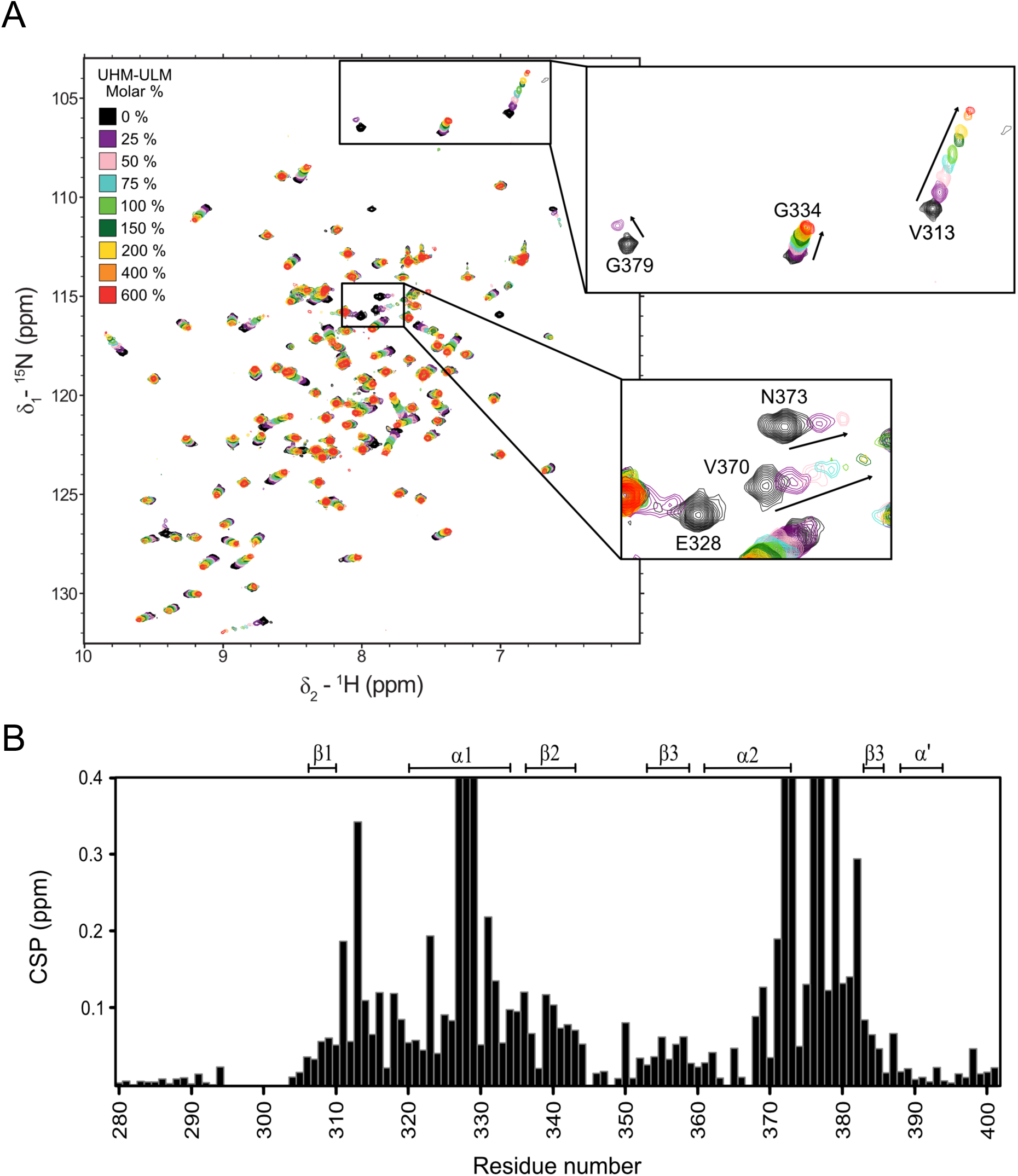
NMR spectral changes reveal the interaction between the RBM17–UHM and SAP30BP–ULM (Related to Figure 4) **(A)** 1H-15N HSQC NMR spectra of RBM17–UHM domain (RBM17/ΔNΔG) upon titration with SAP30BP–ULM domain (SAP30BP/ΔNΔH). **(B)** Bar plot of the calculated chemical shift perturbation observed in the spectrum in panel A. The secondary structure of the RBM17 UHM domain is indicated on top.

**Figure S5.**
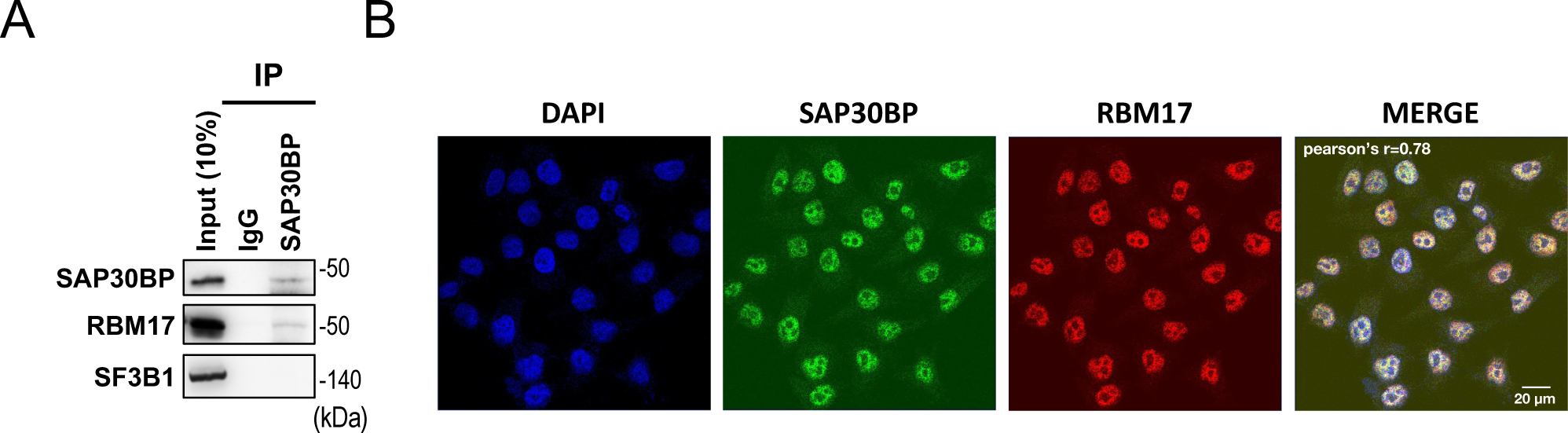
Interaction of the endogenous SAP30BP and RBM17 (Related to Figure 4) **(A)** Co-immunoprecipitation of the endogenous SAP30BP with RBM17. HeLa cell extracts were immunoprecipitated using Rabbit-IgG or anti-SAP30BP antibody, and the immunsoprecipitated proteins were analyzed by Western blotting. **(B)** Subcellular co-localization of SAP30BP and SAP30BP proteins. Co-localizations were examined by immunofluorescence confocal microscopy using indicated antibodies. Pearson’s correlation coefficient is indicated. DAPI was used to stain the nuclear DNA.

**Figure S6.**
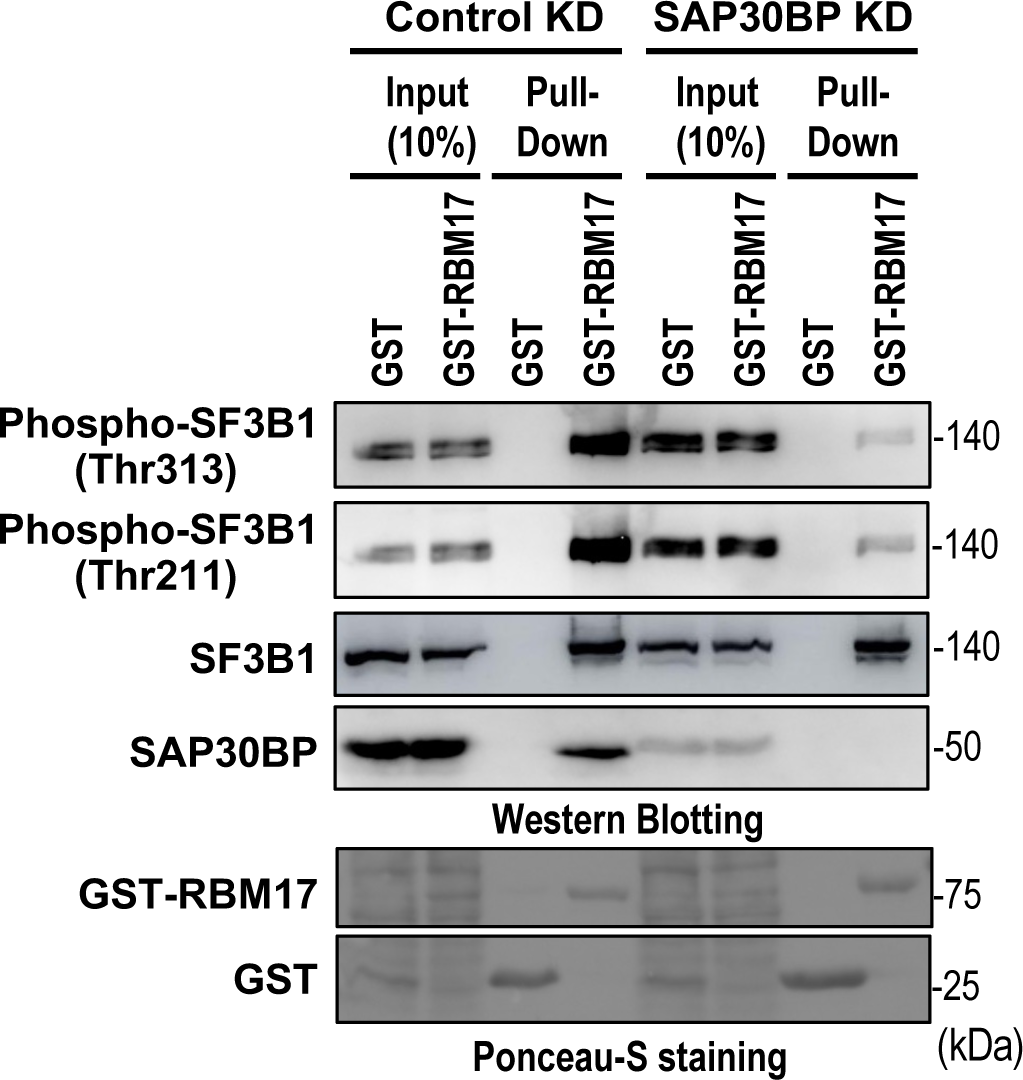
SAP30BP is critical for RBM17 to be recruited in active phospho-SF3B1 (Related to Figure 5) Binding of RBM17 to total SF3B1 and phosphorylated SF3B1under SAP30BP-depletion. GST pull-down assay with whole cell extracts prepared from SAP30BP-knockdown HeLa cells. Proteins associated with GST-RBM17 protein were detected by Western blotting using the indicated antibodies. Active-phosphorylated SF3B1 was detected using antibodies which recognize phosphorylated Thr313 or Thr211.

## Notes

### Competing Interest Statement

The authors have declared no competing interest.

### Summary of Updates

Other statistical analyses (Welch's t-test) were applied in Figures 1C, 5 and Supplementary Figure S1. The statistical analysis (Wilcoxon rank-sum test) was added in Figure 2B. Using confocal fluorescence microscopy, Supplementary FigureS5B was re-analyzed with higher resolution. Accordingly, these figures with the legends were replaced. Minor modifications and corrections were applied in the text and figures.

