## Supplementary Table S1 for "SAP30BP interacts with RBM17/SPF45 to promote splicing in a subset of human short introns" (from another version of this preprint: version 3, 2023-10-04)

### Table S1 (1) The list of retained introns in RBM17-knockdown HeLa cells

See rMATS (http://rnaseq-mats.sourceforge.net/) for the format of this table.

| ID | Gene ID | Chromosome | Strand | RI-Exon Start | RI-Exon End | Upstream ES | Upstream EE | Downstream ES | Intron (nt) | Downstream Exon | ID.1 | IJC SAMPLE 1 | SJC SAMPLE 1 | IJC SAMPLE 2 | SJC SAMPLE 2 | Inc.-Form (nt) | Skip-Form (nt) | P Value | FDR | Inc. Level 1 | Inc. Level 2 | Inc. Level Differ. |
|---|---|---|---|---|---|---|---|---|---|---|---|---|---|---|---|---|---|---|---|---|---|---|
| 352 | XLOC_035171 | chr5 | - | 179044799 | 179045061 | 179044799 | 179044933 | 179044989 | 56 | 179045061 | 352 | 257,227 | 17,401,655 | 778,733 | 14,231,535 | 205 | 149 | 0 | 0 | 0.097,0.091 | 0.284,0.258 | -0.177 |
| 9885 | XLOC_009615 | chr12 | - | 132624967 | 132625280 | 132624967 | 132625100 | 132625164 | 64 | 132625280 | 9885 | 1,1 | 1,3 | 4,2 | 1,0 | 213 | 149 | 9.22E-05 | 0.001935548 | 0.412,0.189 | 0.737,1.0 | -0.568 |
| 4436 | XLOC_041106 | chr7 | + | 56126054 | 56126403 | 56126054 | 56126214 | 56126281 | 67 | 56126403 | 4436 | 11,14 | 5,4 | 26,16 | 1,0 | 216 | 149 | 2.25E-05 | 0.00063373 | 0.603,0.707 | 0.947,1.0 | -0.319 |
| 7506 | XLOC_021015 | chr19 | + | 14674469 | 14674904 | 14674469 | 14674514 | 14674582 | 68 | 14674904 | 7506 | 10,12 | 5,3 | 4,6 | 0,1 | 217 | 149 | 0.002214968 | 0.022451179 | 0.579,0.733 | 1.0,0.805 | -0.247 |
| 3274 | XLOC_018288 | chr17 | + | 79661844 | 79662097 | 79661844 | 79661883 | 79661953 | 70 | 79662097 | 3274 | 254,254 | 394,338 | 501,467 | 501,335 | 219 | 149 | 0.000131947 | 0.002590726 | 0.305,0.338 | 0.405,0.487 | -0.124 |
| 9147 | XLOC_021814 | chr19 | - | 49348626 | 49349103 | 49348626 | 49348703 | 49348773 | 70 | 49349103 | 9147 | 0,0 | 2,6 | 4,5 | 3,1 | 219 | 149 | 2.60E-07 | 1.57E-05 | 0.0,0.0 | 0.476,0.773 | -0.625 |
| 15126 | XLOC_000825 | chr1 | - | 1233392 | 1233801 | 1233392 | 1233493 | 1233564 | 71 | 1233801 | 15126 | 7,6 | 3,3 | 31,51 | 0,2 | 220 | 149 | 0.000124699 | 0.00247131 | 0.612,0.575 | 1.0,0.945 | -0.379 |
| 2483 | XLOC_007374 | chr11 | - | 800761 | 801120 | 800761 | 800912 | 800984 | 72 | 801120 | 2483 | 16,9 | 19,13 | 38,41 | 8,10 | 221 | 149 | 1.93E-06 | 8.40E-05 | 0.362,0.318 | 0.762,0.734 | -0.408 |
| 9037 | XLOC_016435 | chr16 | + | 67201205 | 67201502 | 67201205 | 67201305 | 67201377 | 72 | 67201502 | 9037 | 22,28 | 4,4 | 66,53 | 0,2 | 221 | 149 | 0.000421669 | 0.006381025 | 0.788,0.825 | 1.0,0.947 | -0.167 |
| 10309 | XLOC_013110 | chr14 | + | 50246312 | 50246528 | 50246312 | 50246393 | 50246465 | 72 | 50246528 | 10309 | 64,66 | 179,203 | 123,134 | 138,138 | 221 | 149 | 9.46E-10 | 1.34E-07 | 0.194,0.18 | 0.375,0.396 | -0.199 |
| 234 | XLOC_041771 | chr7 | - | 150773999 | 150774323 | 150773999 | 150774114 | 150774187 | 73 | 150774323 | 234 | 127,112 | 177,191 | 163,213 | 192,165 | 222 | 149 | 0.005520697 | 0.04474263 | 0.325,0.282 | 0.363,0.464 | -0.11 |
| 450 | XLOC_047682 | chrX | + | 44941820 | 44942034 | 44941820 | 44941885 | 44941959 | 74 | 44942034 | 450 | 4,0 | 3,12 | 19,14 | 6,4 | 223 | 149 | 0.000543375 | 0.007718076 | 0.471,0.0 | 0.679,0.7 | -0.454 |
| 14034 | XLOC_023538 | chr2 | + | 74719266 | 74719572 | 74719266 | 74719356 | 74719430 | 74 | 74719572 | 14034 | 20,20 | 41,58 | 53,43 | 52,23 | 223 | 149 | 5.63E-05 | 0.001318181 | 0.246,0.187 | 0.405,0.555 | -0.264 |
| 95 | XLOC_044466 | chr8 | - | 145738024 | 145738521 | 145738024 | 145738154 | 145738229 | 75 | 145738521 | 95 | 97,99 | 272,272 | 229,239 | 234,192 | 224 | 149 | 1.09E-14 | 4.96E-12 | 0.192,0.195 | 0.394,0.453 | -0.23 |
| 3056 | XLOC_045945 | chr9 | + | 79898270 | 79898566 | 79898270 | 79898387 | 79898462 | 75 | 79898566 | 3056 | 0,0 | 22,28 | 6,9 | 54,35 | 224 | 149 | 6.06E-05 | 0.001392389 | 0.0,0.0 | 0.069,0.146 | -0.107 |
| 10769 | XLOC_018709 | chr17 | - | 73127137 | 73127376 | 73127137 | 73127200 | 73127275 | 75 | 73127376 | 10769 | 233,241 | 264,283 | 333,372 | 248,282 | 224 | 149 | 0.001065349 | 0.012897979 | 0.37,0.362 | 0.472,0.467 | -0.104 |
| 2980 | XLOC_024343 | chr2 | - | 190618641 | 190619071 | 190618641 | 190618790 | 190618866 | 76 | 190619071 | 2980 | 3,0 | 32,30 | 8,5 | 28,17 | 225 | 149 | 0.000766191 | 0.010013069 | 0.058,0.0 | 0.159,0.163 | -0.132 |
| 3275 | XLOC_018288 | chr17 | + | 79663636 | 79664028 | 79663636 | 79663777 | 79663853 | 76 | 79664028 | 3275 | 79,51 | 376,389 | 170,195 | 451,484 | 225 | 149 | 1.74E-08 | 1.53E-06 | 0.122,0.08 | 0.2,0.211 | -0.105 |
| 1737 | XLOC_014574 | chr15 | + | 41106140 | 41106921 | 41106140 | 41106268 | 41106345 | 77 | 41106921 | 1737 | 116,149 | 115,111 | 159,193 | 54,47 | 226 | 149 | 1.13E-09 | 1.54E-07 | 0.399,0.469 | 0.66,0.73 | -0.261 |
| 3764 | XLOC_046457 | chr9 | - | 135772570 | 135772928 | 135772570 | 135772732 | 135772809 | 77 | 135772928 | 3764 | 16,3 | 61,57 | 30,17 | 58,57 | 226 | 149 | 0.003204065 | 0.029622336 | 0.147,0.034 | 0.254,0.164 | -0.119 |
| 13152 | XLOC_018790 | chr17 | - | 80018576 | 80018830 | 80018576 | 80018659 | 80018736 | 77 | 80018830 | 13152 | 4,7 | 6,4 | 46,62 | 2,0 | 226 | 149 | 1.59E-08 | 1.42E-06 | 0.305,0.536 | 0.938,1.0 | -0.548 |
| 13190 | XLOC_034753 | chr5 | + | 149747251 | 149747480 | 149747251 | 149747329 | 149747406 | 77 | 149747480 | 13190 | 7,2 | 8,0 | 8,8 | 0,0 | 226 | 149 | 0.000580889 | 0.008090486 | 0.366,1.0 | 1.0,1.0 | -0.317 |
| 14440 | XLOC_018386 | chr17 | - | 7224938 | 7225329 | 7224938 | 7225106 | 7225183 | 77 | 7225329 | 14440 | 20,19 | 104,113 | 59,41 | 103,116 | 226 | 149 | 6.43E-05 | 0.001458729 | 0.113,0.1 | 0.274,0.189 | -0.125 |
| 10097 | XLOC_007211 | chr11 | + | 71940967 | 71941312 | 71940967 | 71941063 | 71941141 | 78 | 71941312 | 10097 | 4,6 | 0,5 | 8,9 | 0,0 | 227 | 149 | 0.00188332 | 0.019935885 | 1.0,0.441 | 1.0,1.0 | -0.279 |
| 698 | XLOC_028549 | chr22 | + | 50636251 | 50636615 | 50636251 | 50636424 | 50636503 | 79 | 50636615 | 698 | 194,175 | 248,232 | 475,413 | 330,405 | 228 | 149 | 0.001751648 | 0.018860619 | 0.338,0.33 | 0.485,0.4 | -0.108 |
| 2892 | XLOC_016217 | chr16 | + | 2284859 | 2285144 | 2284859 | 2285002 | 2285081 | 79 | 2285144 | 2892 | 32,31 | 169,182 | 59,84 | 181,142 | 228 | 149 | 2.04E-05 | 0.000583276 | 0.11,0.1 | 0.176,0.279 | -0.122 |
| 6126 | XLOC_044184 | chr8 | + | 145693063 | 145693370 | 145693063 | 145693162 | 145693241 | 79 | 145693370 | 6126 | 11,22 | 126,91 | 28,33 | 70,74 | 228 | 149 | 8.96E-05 | 0.001902785 | 0.054,0.136 | 0.207,0.226 | -0.121 |
| 14657 | XLOC_006953 | chr11 | + | 828574 | 828963 | 828574 | 828772 | 828851 | 79 | 828963 | 14657 | 0,1 | 20,25 | 4,3 | 7,15 | 228 | 149 | 0.000413862 | 0.006300927 | 0.0,0.025 | 0.272,0.116 | -0.181 |
| 556 | XLOC_030002 | chr3 | - | 123674689 | 123674962 | 123674689 | 123674812 | 123674892 | 80 | 123674962 | 556 | 79,86 | 46,69 | 168,110 | 47,42 | 229 | 149 | 0.000214937 | 0.003835212 | 0.528,0.448 | 0.699,0.63 | -0.176 |
| 1060 | XLOC_021627 | chr19 | - | 19767468 | 19767725 | 19767468 | 19767575 | 19767655 | 80 | 19767725 | 1060 | 109,78 | 66,55 | 197,228 | 91,74 | 229 | 149 | 0.006383917 | 0.04947289 | 0.518,0.48 | 0.585,0.667 | -0.127 |
| 3737 | XLOC_046490 | chr9 | - | 139905424 | 139905752 | 139905424 | 139905557 | 139905637 | 80 | 139905752 | 3737 | 43,65 | 211,175 | 118,106 | 131,146 | 229 | 149 | 2.05E-09 | 2.49E-07 | 0.117,0.195 | 0.37,0.321 | -0.19 |
| 4311 | XLOC_041495 | chr7 | - | 44609407 | 44609832 | 44609407 | 44609502 | 44609582 | 80 | 44609832 | 4311 | 11,13 | 11,8 | 28,18 | 6,0 | 229 | 149 | 0.000477482 | 0.006986662 | 0.394,0.514 | 0.752,1.0 | -0.422 |
| 1779 | XLOC_000300 | chr1 | + | 53343353 | 53344440 | 53343353 | 53343515 | 53343596 | 81 | 53344440 | 1779 | 14,16 | 36,37 | 64,55 | 56,74 | 230 | 149 | 0.00271298 | 0.026145685 | 0.201,0.219 | 0.425,0.325 | -0.165 |
| 4220 | XLOC_045898 | chr9 | + | 35660561 | 35661014 | 35660561 | 35660855 | 35660936 | 81 | 35661014 | 4220 | 250,186 | 69,53 | 519,435 | 64,73 | 230 | 149 | 9.91E-05 | 0.002055803 | 0.701,0.695 | 0.84,0.794 | -0.119 |
| 10098 | XLOC_007211 | chr11 | + | 71940967 | 71941312 | 71940967 | 71941063 | 71941144 | 81 | 71941312 | 10098 | 4,6 | 19,8 | 8,9 | 5,2 | 230 | 149 | 0.002335462 | 0.023398133 | 0.12,0.327 | 0.509,0.745 | -0.403 |
| 12589 | XLOC_043958 | chr8 | + | 21957080 | 21957404 | 21957080 | 21957133 | 21957214 | 81 | 21957404 | 12589 | 0,0 | 2,3 | 4,3 | 3,7 | 230 | 149 | 6.50E-05 | 0.001470498 | 0.0,0.0 | 0.463,0.217 | -0.34 |
| 1451 | XLOC_023617 | chr2 | + | 109367719 | 109368159 | 109367719 | 109367901 | 109367983 | 82 | 109368159 | 1451 | 6,14 | 181,158 | 34,40 | 163,139 | 231 | 149 | 2.85E-08 | 2.40E-06 | 0.021,0.054 | 0.119,0.157 | -0.101 |
| 1774 | XLOC_018149 | chr17 | + | 48452545 | 48453554 | 48452545 | 48453344 | 48453426 | 82 | 48453554 | 1774 | 15,17 | 55,96 | 47,35 | 67,65 | 231 | 149 | 1.68E-05 | 0.000493829 | 0.15,0.103 | 0.312,0.258 | -0.159 |
| 2660 | XLOC_011393 | chr13 | + | 115037658 | 115038417 | 115037658 | 115037956 | 115038038 | 82 | 115038417 | 2660 | 240,189 | 144,141 | 259,274 | 100,110 | 231 | 149 | 0.000417365 | 0.00634637 | 0.518,0.464 | 0.626,0.616 | -0.13 |
| 5705 | XLOC_028552 | chr22 | + | 50960771 | 50961002 | 50960771 | 50960844 | 50960926 | 82 | 50961002 | 5705 | 3,4 | 9,29 | 24,11 | 12,21 | 231 | 149 | 0.001161597 | 0.013714868 | 0.177,0.082 | 0.563,0.253 | -0.278 |
| 5771 | XLOC_041629 | chr7 | - | 100175110 | 100175363 | 100175110 | 100175204 | 100175286 | 82 | 100175363 | 5771 | 13,6 | 3,3 | 1,13 | 0,0 | 231 | 149 | 3.84E-05 | 0.000981851 | 0.737,0.563 | 1.0,1.0 | -0.35 |
| 428 | XLOC_034609 | chr5 | + | 81600433 | 81601300 | 81600433 | 81600669 | 81600752 | 83 | 81601300 | 428 | 7,3 | 3,10 | 3,4 | 0,0 | 232 | 149 | 3.41E-07 | 1.97E-05 | 0.6,0.162 | 1.0,1.0 | -0.619 |
| 873 | XLOC_024154 | chr2 | - | 98428883 | 98429212 | 98428883 | 98429018 | 98429101 | 83 | 98429212 | 873 | 0,0 | 25,37 | 7,8 | 34,32 | 232 | 149 | 4.83E-06 | 0.000179869 | 0.0,0.0 | 0.117,0.138 | -0.128 |
| 7622 | XLOC_030172 | chr3 | - | 196735527 | 196735833 | 196735527 | 196735595 | 196735679 | 84 | 196735833 | 7622 | 0,0 | 12,9 | 2,3 | 5,7 | 233 | 149 | 0.001058031 | 0.012865381 | 0.0,0.0 | 0.204,0.215 | -0.209 |
| 9437 | XLOC_014944 | chr15 | - | 65113365 | 65113596 | 65113365 | 65113480 | 65113565 | 85 | 65113596 | 9437 | 21,20 | 36,41 | 45,57 | 31,29 | 234 | 149 | 1.30E-05 | 0.000409695 | 0.271,0.237 | 0.48,0.556 | -0.264 |
| 10110 | XLOC_007211 | chr11 | + | 71943921 | 71944207 | 71943921 | 71944033 | 71944118 | 85 | 71944207 | 10110 | 2,3 | 10,14 | 11,5 | 3,2 | 234 | 149 | 4.65E-06 | 0.000175024 | 0.113,0.12 | 0.7,0.614 | -0.54 |
| 11276 | XLOC_041357 | chr7 | + | 154795156 | 154798693 | 154795156 | 154795586 | 154795671 | 85 | 154798693 | 11276 | 114,90 | 10,12 | 104,74 | 2,0 | 234 | 149 | 3.83E-06 | 0.000145696 | 0.879,0.827 | 0.971,1.0 | -0.133 |
| 353 | XLOC_035168 | chr5 | - | 178029485 | 178030935 | 178029485 | 178030758 | 178030844 | 86 | 178030935 | 353 | 4,5 | 84,53 | 26,27 | 31,58 | 235 | 149 | 3.66E-10 | 5.56E-08 | 0.029,0.056 | 0.347,0.228 | -0.245 |
| 8631 | XLOC_021878 | chr19 | - | 55606658 | 55606968 | 55606658 | 55606726 | 55606812 | 86 | 55606968 | 8631 | 9,0 | 64,72 | 23,18 | 71,89 | 235 | 149 | 7.99E-05 | 0.001739624 | 0.082,0.0 | 0.17,0.114 | -0.101 |
| 10163 | XLOC_020020 | chr18 | + | 2673015 | 2673362 | 2673015 | 2673193 | 2673279 | 86 | 2673362 | 10163 | 3,0 | 8,5 | 3,8 | 3,5 | 235 | 149 | 0.004449088 | 0.038176369 | 0.192,0.0 | 0.388,0.504 | -0.35 |
| 10550 | XLOC_028486 | chr22 | + | 39066888 | 39067297 | 39066888 | 39066994 | 39067080 | 86 | 39067297 | 10550 | 3,9 | 3,1 | 3,10 | 0,0 | 235 | 149 | 0.001915458 | 0.020204989 | 0.388,0.851 | 1.0,1.0 | -0.381 |
| 13221 | XLOC_016216 | chr16 | + | 2257035 | 2257408 | 2257035 | 2257111 | 2257197 | 86 | 2257408 | 13221 | 1,0 | 13,20 | 6,4 | 16,16 | 235 | 149 | 0.000303979 | 0.004975467 | 0.047,0.0 | 0.192,0.137 | -0.141 |
| 13813 | XLOC_021582 | chr19 | - | 14521027 | 14521417 | 14521027 | 14521146 | 14521232 | 86 | 14521417 | 13813 | 7,12 | 12,26 | 31,40 | 20,21 | 235 | 149 | 0.000224344 | 0.003951507 | 0.27,0.226 | 0.496,0.547 | -0.274 |
| 16260 | XLOC_007253 | chr11 | + | 93526926 | 93527201 | 93526926 | 93527030 | 93527116 | 86 | 93527201 | 16260 | 28,30 | 112,117 | 92,114 | 125,126 | 235 | 149 | 1.18E-09 | 1.59E-07 | 0.137,0.14 | 0.318,0.365 | -0.203 |
| 1193 | XLOC_024434 | chr2 | - | 230656581 | 230656942 | 230656581 | 230656781 | 230656868 | 87 | 230656942 | 1193 | 1,13 | 295,232 | 70,98 | 248,279 | 236 | 149 | 0 | 0 | 0.002,0.034 | 0.151,0.182 | -0.148 |
| 5527 | XLOC_028754 | chr22 | - | 51009586 | 51009968 | 51009586 | 51009716 | 51009803 | 87 | 51009968 | 5527 | 2,3 | 2,3 | 3,1 | 0,0 | 236 | 149 | 1.16E-05 | 0.000372384 | 0.387,0.387 | 1.0,1.0 | -0.613 |
| 9624 | XLOC_047718 | chrX | + | 53113942 | 53114281 | 53113942 | 53114048 | 53114135 | 87 | 53114281 | 9624 | 1,8 | 51,52 | 7,13 | 35,28 | 236 | 149 | 0.00245468 | 0.024291638 | 0.012,0.089 | 0.112,0.227 | -0.119 |
| 11856 | XLOC_015047 | chr15 | - | 89870142 | 89870580 | 89870142 | 89870310 | 89870397 | 87 | 89870580 | 11856 | 13,23 | 6,3 | 35,40 | 2,0 | 236 | 149 | 0.000359977 | 0.005675864 | 0.578,0.829 | 0.917,1.0 | -0.255 |
| 13428 | XLOC_044241 | chr8 | - | 22885843 | 22886159 | 22885843 | 22885954 | 22886041 | 87 | 22886159 | 13428 | 282,270 | 94,96 | 284,322 | 61,67 | 236 | 149 | 0.002479796 | 0.024411711 | 0.654,0.64 | 0.746,0.752 | -0.102 |
| 1203 | XLOC_013333 | chr14 | - | 24709468 | 24710081 | 24709468 | 24709536 | 24709624 | 88 | 24710081 | 1203 | 149,178 | 48,40 | 204,207 | 29,33 | 237 | 149 | 0.005472353 | 0.044494671 | 0.661,0.737 | 0.816,0.798 | -0.108 |
| 2509 | XLOC_007251 | chr11 | + | 93462779 | 93463135 | 93462779 | 93462947 | 93463035 | 88 | 93463135 | 2509 | 11,17 | 132,156 | 42,55 | 149,116 | 237 | 149 | 2.88E-08 | 2.40E-06 | 0.05,0.064 | 0.151,0.23 | -0.134 |
| 4655 | XLOC_038403 | chr6 | - | 111995483 | 111995833 | 111995483 | 111995613 | 111995701 | 88 | 111995833 | 4655 | 1,4 | 2,5 | 7,25 | 1,2 | 237 | 149 | 0.000120851 | 0.002414125 | 0.239,0.335 | 0.815,0.887 | -0.564 |
| 7042 | XLOC_029383 | chr3 | + | 40502898 | 40503152 | 40502898 | 40503010 | 40503098 | 88 | 40503152 | 7042 | 3,8 | 4,1 | 12,6 | 0,1 | 237 | 149 | 0.001727975 | 0.018712887 | 0.32,0.834 | 1.0,0.79 | -0.318 |
| 10878 | XLOC_021489 | chr19 | - | 5616138 | 5616497 | 5616138 | 5616343 | 5616432 | 89 | 5616497 | 10878 | 1,0 | 0,3 | 3,1 | 0,0 | 238 | 149 | 0.000475038 | 0.006971048 | 1.0,0.0 | 1.0,1.0 | -0.5 |
| 14289 | XLOC_000967 | chr1 | - | 27682878 | 27683175 | 27682878 | 27682991 | 27683080 | 89 | 27683175 | 14289 | 0,1 | 37,40 | 10,17 | 26,39 | 238 | 149 | 2.61E-09 | 3.09E-07 | 0.0,0.015 | 0.194,0.214 | -0.197 |
| 1035 | XLOC_023585 | chr2 | + | 97866070 | 97866262 | 97866070 | 97866099 | 97866189 | 90 | 97866262 | 1035 | 1,1 | 36,33 | 2,7 | 16,19 | 239 | 149 | 0.000504944 | 0.007296221 | 0.017,0.019 | 0.072,0.187 | -0.112 |
| 1823 | XLOC_013529 | chr14 | - | 102695132 | 102695994 | 102695132 | 102695703 | 102695793 | 90 | 102695994 | 1823 | 12,17 | 19,18 | 44,40 | 9,12 | 239 | 149 | 3.42E-06 | 0.000132998 | 0.283,0.371 | 0.753,0.675 | -0.387 |

| | | | | | | | | | | | | | | | | | | | | | | |
|---|---|---|---|---|---|---|---|---|---|---|---|---|---|---|---|---|---|---|---|---|---|---|
| 9742 | XLOC_014609 | chr15 | + | 44150628 | 44151021 | 44150628 | 44150778 | 44150868 | 90 | 44151021 | 9742 | 0,0 | 2,2 | 1,4 | 5,9 | 239 | 149 | 0.003598115 | 0.032428309 | 0.0,0.0 | 0.111,0.217 | -0.164 |
| 10085 | XLOC_013517 | chr14 | - | 96798641 | 96798989 | 96798641 | 96798747 | 96798837 | 90 | 96798989 | 10085 | 0,0 | 16,13 | 3,3 | 19,7 | 239 | 149 | 0.002239458 | 0.022608002 | 0.0,0.0 | 0.09,0.211 | -0.15 |
| 12629 | XLOC_007186 | chr11 | + | 67163186 | 67163642 | 67163186 | 67163296 | 67163386 | 90 | 67163642 | 12629 | 3,3 | 5,1 | 5,1 | 0,0 | 239 | 149 | 1.72E-06 | 7.63E-05 | 0.272,0.652 | 1.0,1.0 | -0.538 |
| 14521 | XLOC_009183 | chr12 | + | 132514245 | 132514699 | 132514245 | 132514448 | 132514538 | 90 | 132514699 | 14521 | 1,4 | 6,2 | 6,7 | 0,0 | 239 | 149 | 4.78E-08 | 3.69E-06 | 0.094,0.555 | 1.0,1.0 | -0.675 |
| 14721 | XLOC_007127 | chr11 | + | 63970483 | 63970700 | 63970483 | 63970506 | 63970596 | 90 | 63970700 | 14721 | 5,16 | 5,0 | 3,3 | 0,0 | 239 | 149 | 0.00393877 | 0.03462244 | 0.384,1.0 | 1.0,1.0 | -0.308 |
| 15925 | XLOC_001222 | chr1 | - | 109823391 | 109823873 | 109823391 | 109823668 | 109823758 | 90 | 109823873 | 15925 | 136,111 | 105,123 | 155,136 | 58,48 | 239 | 149 | 5.69E-07 | 3.01E-05 | 0.447,0.36 | 0.625,0.639 | -0.229 |
| 5146 | XLOC_044458 | chr8 | - | 145541757 | 145541978 | 145541757 | 145541832 | 145541923 | 91 | 145541978 | 5146 | 12,27 | 135,174 | 25,48 | 107,101 | 240 | 149 | 4.56E-05 | 0.001125993 | 0.052,0.088 | 0.127,0.228 | -0.107 |
| 8731 | XLOC_024384 | chr2 | - | 208446517 | 208446884 | 208446517 | 208446670 | 208446761 | 91 | 208446884 | 8731 | 25,44 | 6,1 | 31,31 | 0,0 | 240 | 149 | 0.000466042 | 0.006886256 | 0.721,0.965 | 1.0,1.0 | -0.157 |
| 5564 | XLOC_028749 | chr22 | - | 50905766 | 50906040 | 50905766 | 50905877 | 50905969 | 92 | 50906040 | 5564 | 5,1 | 18,7 | 11,10 | 5,14 | 241 | 149 | 0.0038066 | 0.033820951 | 0.147,0.081 | 0.576,0.306 | -0.327 |
| 10713 | XLOC_000603 | chr1 | + | 169819405 | 169819707 | 169819405 | 169819486 | 169819578 | 92 | 169819707 | 10713 | 10,7 | 2,2 | 12,10 | 0,0 | 241 | 149 | 0.001746506 | 0.018818726 | 0.756,0.684 | 1.0,1.0 | -0.28 |
| 1181 | XLOC_020233 | chr18 | - | 47812118 | 47812627 | 47812118 | 47812298 | 47812391 | 93 | 47812627 | 1181 | 39,21 | 181,168 | 46,72 | 123,129 | 242 | 149 | 7.47E-07 | 3.73E-05 | 0.117,0.071 | 0.187,0.256 | -0.128 |
| 11658 | XLOC_041003 | chr7 | + | 6054532 | 6054983 | 6054532 | 6054683 | 6054776 | 93 | 6054983 | 11658 | 1,0 | 1,2 | 12,4 | 5,0 | 242 | 149 | 0.000302775 | 0.00496116 | 0.381,0.0 | 0.596,1.0 | -0.608 |
| 8724 | XLOC_021139 | chr19 | + | 36104769 | 36105036 | 36104769 | 36104802 | 36104896 | 94 | 36105036 | 8724 | 2,5 | 4,1 | 8,7 | 0,0 | 243 | 149 | 2.72E-05 | 0.00074809 | 0.235,0.754 | 1.0,1.0 | -0.506 |
| 9385 | XLOC_026746 | chr20 | + | 31015933 | 31016225 | 31015933 | 31016032 | 31016127 | 95 | 31016225 | 9385 | 31,42 | 4,5 | 39,37 | 2,0 | 244 | 149 | 0.004654716 | 0.039468 | 0.826,0.837 | 0.923,1.0 | -0.13 |
| 9975 | XLOC_007774 | chr11 | - | 134026646 | 134026990 | 134026646 | 134026817 | 134026912 | 95 | 134026990 | 9975 | 35,23 | 13,9 | 25,29 | 2,4 | 244 | 149 | 0.004400346 | 0.037822859 | 0.622,0.609 | 0.884,0.816 | -0.235 |
| 4543 | XLOC_038359 | chr6 | - | 90377732 | 90378162 | 90377732 | 90377870 | 90377966 | 96 | 90378162 | 4543 | 5,6 | 4,2 | 5,1 | 0,0 | 245 | 149 | 5.06E-06 | 0.000186952 | 0.432,0.646 | 1.0,1.0 | -0.461 |
| 9660 | XLOC_014829 | chr15 | - | 34823295 | 34823760 | 34823295 | 34823579 | 34823675 | 96 | 34823760 | 9660 | 0,1 | 4,2 | 6,1 | 4,1 | 245 | 149 | 0.0007008 | 0.009361665 | 0.0,0.233 | 0.477,0.378 | -0.311 |
| 9756 | XLOC_014584 | chr15 | + | 41793383 | 41793757 | 41793383 | 41793564 | 41793660 | 96 | 41793757 | 9756 | 1,1 | 0,4 | 2,6 | 0,0 | 245 | 149 | 0.001886595 | 0.019956511 | 1.0,0.132 | 1.0,1.0 | -0.434 |
| 4232 | XLOC_041774 | chr7 | - | 150938249 | 150938739 | 150938249 | 150938480 | 150938577 | 97 | 150938739 | 4232 | 5,12 | 6,0 | 4,1 | 0,0 | 246 | 149 | 0.002094446 | 0.021519576 | 0.335,1.0 | 1.0,1.0 | -0.333 |
| 4931 | XLOC_034487 | chr5 | + | 1802195 | 1802488 | 1802195 | 1802335 | 1802434 | 99 | 1802488 | 4931 | 6,9 | 3,2 | 8,18 | 0,0 | 248 | 149 | 9.01E-05 | 0.001903505 | 0.546,0.73 | 1.0,1.0 | -0.362 |
| 333 | XLOC_037658 | chr6 | + | 17543067 | 17543374 | 17543067 | 17543191 | 17543291 | 100 | 17543374 | 333 | 2,7 | 58,40 | 20,11 | 38,39 | 249 | 149 | 0.000113119 | 0.002299363 | 0.02,0.095 | 0.24,0.144 | -0.135 |
| 7098 | XLOC_021619 | chr19 | - | 19042139 | 19042434 | 19042139 | 19042217 | 19042317 | 100 | 19042434 | 7098 | 5,2 | 2,2 | 7,12 | 0,0 | 249 | 149 | 0.000236471 | 0.004131238 | 0.599,0.374 | 1.0,1.0 | -0.514 |
| 14150 | XLOC_040978 | chr7 | + | 769073 | 769484 | 769073 | 769198 | 769299 | 101 | 769484 | 14150 | 5,2 | 6,3 | 3,9 | 1,0 | 250 | 149 | 4.76E-05 | 0.001166164 | 0.332,0.284 | 0.641,1.0 | -0.512 |
| 4572 | XLOC_038257 | chr6 | - | 38654667 | 38654906 | 38654667 | 38654750 | 38654852 | 102 | 38654906 | 4572 | 5,3 | 1,3 | 5,5 | 0,0 | 251 | 149 | 0.000518825 | 0.007453832 | 0.748,0.372 | 1.0,1.0 | -0.44 |
| 15689 | XLOC_000426 | chr1 | + | 113217176 | 113217541 | 113217176 | 113217351 | 113217453 | 102 | 113217541 | 15689 | 23,52 | 77,77 | 38,41 | 36,38 | 251 | 149 | 0.002458306 | 0.024295557 | 0.151,0.286 | 0.385,0.39 | -0.169 |
| 5476 | XLOC_029386 | chr3 | + | 42260708 | 42261088 | 42260708 | 42260882 | 42260985 | 103 | 42261088 | 5476 | 3,6 | 7,9 | 11,19 | 6,4 | 252 | 149 | 0.002260134 | 0.022786146 | 0.202,0.283 | 0.52,0.737 | -0.386 |
| 6548 | XLOC_024016 | chr2 | - | 44145165 | 44145537 | 44145165 | 44145290 | 44145394 | 104 | 44145537 | 6548 | 0,1 | 2,0 | 1,2 | 1,0 | 253 | 149 | 0.00294212 | 0.027676905 | 0.0,1.0 | 0.371,1.0 | -0.185 |
| 8819 | XLOC_016786 | chr16 | - | 69402278 | 69402498 | 69402278 | 69402385 | 69402489 | 104 | 69402498 | 8819 | 2,0 | 2,6 | 19,11 | 12,5 | 253 | 149 | 0.001411653 | 0.015917608 | 0.371,0.0 | 0.483,0.564 | -0.338 |
| 10595 | XLOC_021613 | chr19 | - | 18548570 | 18548985 | 18548570 | 18548798 | 18548902 | 104 | 18548985 | 10595 | 9,9 | 76,85 | 23,19 | 58,62 | 253 | 149 | 0.000170605 | 0.003216663 | 0.065,0.059 | 0.189,0.153 | -0.109 |
| 14229 | XLOC_018153 | chr17 | + | 48827861 | 48828213 | 48827861 | 48828002 | 48828107 | 105 | 48828213 | 14229 | 915,899 | 219,188 | 11,811,140 | 58,62 | 254 | 149 | 0 | 0 | 0.71,0.737 | 0.923,0.915 | -0.196 |
| 7976 | XLOC_034626 | chr5 | + | 96092881 | 96093198 | 96092881 | 96093040 | 96093147 | 107 | 96093198 | 7976 | 10,5 | 6,2 | 10,8 | 0,0 | 256 | 149 | 1.19E-06 | 5.62E-05 | 0.492,0.593 | 1.0,1.0 | -0.458 |
| 5266 | XLOC_029841 | chr3 | - | 49153246 | 49153595 | 49153246 | 49153371 | 49153479 | 108 | 49153595 | 5266 | 8,3 | 40,40 | 21,11 | 46,35 | 257 | 149 | 0.002881118 | 0.027376988 | 0.104,0.042 | 0.209,0.154 | -0.108 |
| 730 | XLOC_027777 | chr21 | - | 34954256 | 34954552 | 34954256 | 34954361 | 34954470 | 109 | 34954552 | 730 | 31,40 | 136,125 | 46,55 | 95,86 | 258 | 149 | 0.000481775 | 0.007035783 | 0.116,0.156 | 0.219,0.27 | -0.108 |
| 2476 | XLOC_048000 | chrX | - | 71933650 | 71934208 | 71933650 | 71933789 | 71933898 | 109 | 71934208 | 2476 | 129,94 | 33,19 | 206,176 | 24,16 | 258 | 149 | 0.001135289 | 0.013488949 | 0.693,0.741 | 0.832,0.864 | -0.131 |
| 11082 | XLOC_046244 | chr9 | - | 35812893 | 35813335 | 35812893 | 35813153 | 35813262 | 109 | 35813335 | 11082 | 9,2 | 3,7 | 4,4 | 1,0 | 258 | 149 | 0.00059843 | 0.008296397 | 0.634,0.142 | 0.698,1.0 | -0.461 |
| 15204 | XLOC_023671 | chr2 | + | 135920148 | 135920421 | 135920148 | 135920208 | 135920317 | 109 | 135920421 | 15204 | 9,11 | 3,4 | 15,13 | 0,2 | 258 | 149 | 0.005385784 | 0.043957116 | 0.634,0.614 | 1.0,0.79 | -0.271 |
| 6156 | XLOC_041630 | chr7 | - | 100280930 | 100281255 | 100280930 | 100281061 | 100281172 | 111 | 100281255 | 6156 | 11,8 | 5,1 | 10,9 | 0,1 | 260 | 149 | 0.001744375 | 0.018809235 | 0.558,0.821 | 1.0,0.838 | -0.23 |
| 3991 | XLOC_044466 | chr8 | - | 145737063 | 145737445 | 145737063 | 145737172 | 145737284 | 112 | 145737445 | 3991 | 6,6 | 6,4 | 22,23 | 4,3 | 261 | 149 | 0.002924851 | 0.027569316 | 0.363,0.461 | 0.758,0.814 | -0.374 |
| 7940 | XLOC_018714 | chr17 | - | 73282713 | 73283114 | 73282713 | 73282883 | 73282995 | 112 | 73283114 | 7940 | 12,5 | 6,3 | 16,4 | 1,0 | 261 | 149 | 7.12E-05 | 0.00158088 | 0.533,0.488 | 0.901,1.0 | -0.44 |
| 5240 | XLOC_029868 | chr3 | - | 52005475 | 52005908 | 52005475 | 52005714 | 52005827 | 113 | 52005908 | 5240 | 27,44 | 7,2 | 24,42 | 0,1 | 262 | 149 | 0.000556374 | 0.007858199 | 0.687,0.926 | 1.0,0.96 | -0.173 |
| 12922 | XLOC_018124 | chr17 | + | 45696374 | 45696688 | 45696374 | 45696530 | 45696643 | 113 | 45696688 | 12922 | 64,61 | 21,23 | 46,44 | 0,9 | 262 | 149 | 0.002052757 | 0.021207126 | 0.634,0.601 | 1.0,0.735 | -0.25 |
| 10605 | XLOC_005139 | chr10 | - | 14870143 | 14870481 | 14870143 | 14870271 | 14870385 | 114 | 14870481 | 10605 | 0,2 | 3,1 | 2,1 | 1,0 | 263 | 149 | 0.001285184 | 0.014824947 | 0.0,0.531 | 0.531,1.0 | -0.5 |
| 13452 | XLOC_017993 | chr17 | + | 27896335 | 27896643 | 27896335 | 27896377 | 27896491 | 114 | 27896643 | 13452 | 8,15 | 3,1 | 6,10 | 0,0 | 263 | 149 | 0.005922452 | 0.046986036 | 0.602,0.895 | 1.0,1.0 | -0.252 |
| 15273 | XLOC_007445 | chr11 | - | 18320167 | 18320517 | 18320167 | 18320224 | 18320338 | 114 | 18320517 | 15273 | 4,2 | 4,0 | 4,2 | 0,0 | 263 | 149 | 0.005301958 | 0.043343504 | 0.362,1.0 | 1.0,1.0 | -0.319 |
| 1105 | XLOC_021261 | chr19 | + | 45567269 | 45567888 | 45567269 | 45567485 | 45567600 | 115 | 45567888 | 1105 | 46,56 | 117,123 | 125,103 | 92,131 | 264 | 149 | 3.99E-05 | 0.001015057 | 0.182,0.204 | 0.434,0.307 | -0.177 |
| 1697 | XLOC_001548 | chr1 | - | 228295935 | 228296722 | 228295935 | 228296022 | 228296137 | 115 | 228296722 | 1697 | 43,41 | 6,10 | 61,68 | 0,7 | 264 | 149 | 0.006358932 | 0.049330096 | 0.802,0.698 | 1.0,0.846 | -0.173 |
| 7542 | XLOC_038282 | chr6 | - | 43536217 | 43536474 | 43536217 | 43536332 | 43536447 | 115 | 43536474 | 7542 | 0,2 | 1,2 | 6,1 | 0,0 | 264 | 149 | 0.00010575 | 0.002174812 | 0.0,0.361 | 1.0,1.0 | -0.82 |
| 11805 | XLOC_008857 | chr12 | + | 44164840 | 44165168 | 44164840 | 44164907 | 44165022 | 115 | 44165168 | 11805 | 4,1 | 2,1 | 8,3 | 0,0 | 264 | 149 | 0.006028777 | 0.047645284 | 0.53,0.361 | 1.0,1.0 | -0.554 |
| 2931 | XLOC_023538 | chr2 | + | 74717151 | 74717600 | 74717151 | 74717254 | 74717370 | 116 | 74717600 | 2931 | 25,20 | 66,51 | 48,31 | 51,49 | 265 | 149 | 0.004707318 | 0.03982423 | 0.176,0.181 | 0.346,0.262 | -0.126 |
| 8939 | XLOC_016560 | chr16 | - | 774105 | 774523 | 774105 | 774205 | 774321 | 116 | 774523 | 8939 | 1,1 | 22,28 | 10,18 | 42,42 | 265 | 149 | 1.36E-05 | 0.000419071 | 0.025,0.02 | 0.118,0.194 | -0.134 |
| 13043 | XLOC_041378 | chr7 | - | 1515828 | 1516293 | 1515828 | 1516088 | 1516205 | 117 | 1516293 | 13043 | 0,0 | 7,1 | 1,4 | 0,0 | 266 | 149 | 9.06E-11 | 1.62E-08 | 0.0,0.0 | 1.0,1.0 | -1 |
| 1934 | XLOC_028602 | chr22 | - | 24180897 | 24181318 | 24180897 | 24180963 | 24181081 | 118 | 24181318 | 1934 | 20,11 | 26,27 | 61,33 | 36,15 | 267 | 149 | 0.000167636 | 0.003179792 | 0.3,0.185 | 0.486,0.551 | -0.276 |
| 4037 | XLOC_044443 | chr8 | - | 144921343 | 144921513 | 144921343 | 144921367 | 144921485 | 118 | 144921513 | 4037 | 39,28 | 104,76 | 54,51 | 52,53 | 267 | 149 | 6.26E-06 | 0.000220453 | 0.173,0.171 | 0.367,0.349 | -0.186 |
| 7277 | XLOC_021261 | chr19 | + | 45567269 | 45567888 | 45567269 | 45567482 | 45567600 | 118 | 45567888 | 7277 | 21,27 | 10,7 | 65,49 | 4,7 | 267 | 149 | 0.003716081 | 0.033175498 | 0.54,0.683 | 0.901,0.796 | -0.237 |
| 7992 | XLOC_018673 | chr17 | - | 62499920 | 62500205 | 62499920 | 62499986 | 62500104 | 118 | 62500205 | 7992 | 19,14 | 1,5 | 15,12 | 1,0 | 267 | 149 | 0.003386189 | 0.030888447 | 0.914,0.61 | 0.893,1.0 | -0.184 |
| 10448 | XLOC_011394 | chr13 | + | 115051703 | 115052104 | 115051703 | 115051875 | 115051993 | 118 | 115052104 | 10448 | 88,99 | 204,203 | 114,118 | 133,137 | 267 | 149 | 8.95E-05 | 0.001902785 | 0.194,0.214 | 0.324,0.325 | -0.12 |
| 3538 | XLOC_048105 | chrX | - | 138819889 | 138820228 | 138819889 | 138819955 | 138820074 | 119 | 138820228 | 3538 | 2,12 | 2,3 | 4,2 | 0,0 | 268 | 149 | 5.88E-05 | 0.001360044 | 0.357,0.69 | 1.0,1.0 | -0.477 |
| 4595 | XLOC_038080 | chr6 | - | 3285169 | 3285345 | 3285169 | 3285193 | 3285312 | 119 | 3285345 | 4595 | 1,0 | 0,4 | 1,1 | 0,0 | 268 | 149 | 0.000157837 | 0.003009104 | 1.0,0.0 | 1.0,1.0 | -0.5 |
| 15257 | XLOC_047875 | chrX | + | 152959359 | 152959713 | 152959359 | 152959472 | 152959591 | 119 | 152959713 | 15257 | 21,19 | 10,7 | 27,37 | 3,3 | 268 | 149 | 0.001156733 | 0.013678916 | 0.539,0.601 | 0.833,0.873 | -0.283 |
| 10171 | XLOC_013352 | chr14 | - | 35591207 | 35591618 | 35591207 | 35591307 | 35591427 | 120 | 35591618 | 10171 | 23,26 | 70,61 | 54,51 | 50,50 | 269 | 149 | 9.21E-06 | 0.00030587 | 0.154,0.191 | 0.374,0.361 | -0.195 |
| 14017 | XLOC_000373 | chr1 | + | 93586105 | 93586451 | 93586105 | 93586229 | 93586351 | 122 | 93586451 | 14017 | 23,22 | 5,3 | 5,9 | 1,0 | 271 | 149 | 0.005782123 | 0.046140423 | 0.717,0.801 | 0.733,1.0 | -0.108 |
| 10509 | XLOC_027816 | chr21 | - | 46285309 | 46285565 | 46285309 | 46285362 | 46285485 | 123 | 46285565 | 10509 | 24,45 | 4,3 | 43,31 | 1,0 | 272 | 149 | 0.001217753 | 0.01424373 | 0.767,0.892 | 0.959,1.0 | -0.15 |
| 14549 | XLOC_004988 | chr10 | + | 97816649 | 97817004 | 97816649 | 97816757 | 97816880 | 123 | 97817004 | 14549 | 2,0 | 7,8 | 6,4 | 8,6 | 272 | 149 | 0.002580707 | 0.025125565 | 0.135,0.0 | 0.291,0.268 | -0.212 |
| 1360 | XLOC_018027 | chr17 | + | 34956400 | 34957467 | 34956400 | 34956466 | 34956590 | 124 | 34957467 | 1360 | 3,8 | 59,43 | 6,20 | 31,24 | 273 | 149 | 0.00136034 | 0.015466546 | 0.027,0.092 | 0.096,0.313 | -0.145 |
| 13620 | XLOC_018545 | chr17 | - | 40139608 | 40139928 | 40139608 | 40139712 | 40139836 | 124 | 40139928 | 13620 | 6,8 | 11,6 | 25,8 | 5,1 | 273 | 149 | 0.000463905 | 0.006861421 | 0.229,0.421 | 0.732,0.814 | -0.448 |
| 15690 | XLOC_000426 | chr1 | + | 113217318 | 113217541 | 113217318 | 113217351 | 113217475 | 124 | 113217541 | 15690 | 11,7 | 8,12 | 11,18 | 0,3 | 273 | 149 | 6.49E-05 | 0.001470498 | 0.429,0.241 | 1.0,0.766 | -0.548 |
| 14675 | XLOC_007153 | chr11 | + | 65338052 | 65338441 | 65338052 | 65338231 | 65338356 | 125 | 65338441 | 14675 | 1,0 | 4,8 | 7,2 | 14,6 | 274 | 149 | 0.001765024 | 0.018950384 | 0.12,0.0 | 0.214,0.153 | -0.123 |
| 6714 | XLOC_028458 | chr22 | + | 32012730 | 32013122 | 32012730 | 32012863 | 32012989 | 126 | 32013122 | 6714 | 1,0 | 0,2 | 4,3 | 1,0 | 275 | 149 | 0.001760099 | 0.018924525 | 1.0,0.0 | 0.684,1.0 | -0.342 |
| 7080 | XLOC_023353 | chr2 | + | 271892 | 272305 | 271892 | 272065 | 272191 | 126 | 272305 | 7080 | 453,462 | 64,54 | 743,752 | 38,44 | 275 | 149 | 1.64E-06 | 7.39E-05 | 0.793,0.823 | 0.914,0.903 | -0.101 |
| 7330 | XLOC_017967 | chr17 | + | 20163504 | 20163836 | 20163504 | 20163607 | 20163736 | 129 | 20163836 | 7330 | 0,1 | 1,3 | 4,10 | 2,1 | 278 | 149 | 4.46E-05 | 0.001104034 | 0.0,0.152 | 0.517,0.843 | -0.604 |
| 729 | XLOC_027785 | chr21 | - | 37415698 | 37416254 | 37415698 | 37415949 | 37416079 | 130 | 37416254 | 729 | 13,3 | 10,9 | 25,15 | 4,5 | 279 | 149 | 0.001611861 | 0.017697527 | 0.41,0.151 | 0.769,0.616 | -0.412 |

| | | | | | | | | | | | | | | | | | | | | | | |
|---|---|---|---|---|---|---|---|---|---|---|---|---|---|---|---|---|---|---|---|---|---|---|
| 9268 | XLOC_037650 | chr6 | + | 13601065 | 13601353 | 13601065 | 13601181 | 13601312 | 131 | 13601353 | 9268 | 7,2 | 3,2 | 15,14 | 1,0 | 280 | 149 | 0.000131825 | 0.002590726 | 0.554,0.347 | 0.889,1.0 | -0.494 |
| 9623 | XLOC_047718 | chrX | + | 53113616 | 53114048 | 53113616 | 53113804 | 53113936 | 132 | 53114048 | 9623 | 26,16 | 43,37 | 13,21 | 6,8 | 281 | 149 | 2.77E-05 | 0.000759497 | 0.243,0.187 | 0.535,0.582 | -0.344 |
| 10111 | XLOC_007211 | chr11 | + | 71944484 | 71944787 | 71944484 | 71944566 | 71944698 | 132 | 71944787 | 10111 | 191,215 | 771,776 | 367,385 | 561,590 | 281 | 149 | 1.27E-13 | 4.66E-11 | 0.116,0.128 | 0.258,0.257 | -0.136 |
| 4373 | XLOC_041301 | chr7 | + | 129094261 | 129094700 | 129094261 | 129094368 | 129094501 | 133 | 129094700 | 4373 | 0,0 | 1,1 | 1,3 | 0,0 | 282 | 149 | 7.76E-09 | 7.73E-07 | 0.0,0.0 | 1.0,1.0 | -1 |
| 8112 | XLOC_018519 | chr17 | - | 36962368 | 36962648 | 36962368 | 36962416 | 36962549 | 133 | 36962648 | 8112 | 0,1 | 3,0 | 1,1 | 0,0 | 282 | 149 | 0.000288982 | 0.004779308 | 0.0,1.0 | 1.0,1.0 | -0.5 |
| 10662 | XLOC_000946 | chr1 | - | 24297827 | 24298113 | 24297827 | 24297929 | 24298062 | 133 | 24298113 | 10662 | 3,0 | 13,2 | 2,3 | 0,2 | 282 | 149 | 0.004585551 | 0.039035576 | 0.109,0.0 | 1.0,0.442 | -0.666 |
| 4365 | XLOC_041366 | chr7 | + | 157046360 | 157046836 | 157046360 | 157046513 | 157046647 | 134 | 157046836 | 4365 | 0,2 | 1,4 | 9,2 | 0,0 | 283 | 149 | 5.08E-09 | 5.46E-07 | 0.0,0.208 | 1.0,1.0 | -0.896 |
| 10463 | XLOC_016854 | chr16 | - | 88787997 | 88788474 | 88787997 | 88788081 | 88788215 | 134 | 88788474 | 10463 | 44,54 | 42,29 | 48,56 | 8,17 | 283 | 149 | 0.000746149 | 0.009819404 | 0.355,0.495 | 0.76,0.634 | -0.272 |
| 5672 | XLOC_028587 | chr22 | - | 21983589 | 21984050 | 21983589 | 21983696 | 21983831 | 135 | 21984050 | 5672 | 80,43 | 48,39 | 129,133 | 31,52 | 284 | 149 | 0.000893509 | 0.011303753 | 0.466,0.366 | 0.686,0.573 | -0.213 |
| 12026 | XLOC_000229 | chr1 | + | 40214378 | 40214760 | 40214378 | 40214439 | 40214574 | 135 | 40214760 | 12026 | 1,2 | 6,4 | 6,2 | 1,0 | 284 | 149 | 5.31E-06 | 0.000194341 | 0.08,0.208 | 0.759,1.0 | -0.735 |
| 12027 | XLOC_021106 | chr19 | + | 33073100 | 33073364 | 33073100 | 33073138 | 33073275 | 137 | 33073364 | 12027 | 16,23 | 4,2 | 27,18 | 1,0 | 286 | 149 | 0.001172158 | 0.013817865 | 0.676,0.857 | 0.934,1.0 | -0.201 |
| 7752 | XLOC_034882 | chr5 | - | 40833878 | 40834367 | 40833878 | 40834143 | 40834282 | 139 | 40834367 | 7752 | 8,9 | 9,0 | 30,27 | 0,0 | 288 | 149 | 0.000681711 | 0.009188442 | 0.315,1.0 | 1.0,1.0 | -0.343 |
| 9214 | XLOC_016191 | chr16 | + | 778796 | 779088 | 778796 | 778868 | 779008 | 140 | 779088 | 9214 | 148,171 | 18,20 | 233,239 | 10,8 | 289 | 149 | 0.000115647 | 0.002344423 | 0.809,0.815 | 0.923,0.939 | -0.119 |
| 11335 | XLOC_009489 | chr12 | - | 95450972 | 95451419 | 95450972 | 95451188 | 95451328 | 140 | 95451419 | 11335 | 0,5 | 9,15 | 15,11 | 12,21 | 289 | 149 | 0.005621654 | 0.045340978 | 0.0,0.147 | 0.392,0.213 | -0.229 |
| 12943 | XLOC_028747 | chr22 | - | 50720358 | 50720742 | 50720358 | 50720471 | 50720612 | 141 | 50720742 | 12943 | 6,3 | 2,2 | 1,11 | 0,0 | 290 | 149 | 0.000280981 | 0.004672266 | 0.607,0.435 | 1.0,1.0 | -0.479 |
| 8469 | XLOC_017897 | chr17 | + | 7161906 | 7162262 | 7161906 | 7162002 | 7162144 | 142 | 7162262 | 8469 | 100,112 | 203,208 | 104,136 | 131,136 | 291 | 149 | 0.000467095 | 0.006895041 | 0.201,0.216 | 0.289,0.339 | -0.105 |
| 9071 | XLOC_021326 | chr19 | + | 50387509 | 50387813 | 50387509 | 50387626 | 50387768 | 142 | 50387813 | 9071 | 40,52 | 4,3 | 67,63 | 0,1 | 291 | 149 | 0.001204483 | 0.014110465 | 0.837,0.899 | 1.0,0.97 | -0.117 |
| 11561 | XLOC_029823 | chr3 | - | 48664063 | 48664540 | 48664063 | 48664118 | 48664260 | 142 | 48664540 | 11561 | 1,0 | 0,3 | 2,3 | 0,0 | 291 | 149 | 0.000263737 | 0.004481105 | 1.0,0.0 | 1.0,1.0 | -0.5 |
| 16174 | XLOC_028574 | chr22 | - | 19841965 | 19842363 | 19841965 | 19842062 | 19842204 | 142 | 19842363 | 16174 | 32,40 | 5,3 | 42,60 | 1,0 | 291 | 149 | 0.00023816 | 0.004151107 | 0.766,0.872 | 0.956,1.0 | -0.159 |
| 2080 | XLOC_001214 | chr1 | - | 101360188 | 101361665 | 101360188 | 101360342 | 101360485 | 143 | 101361665 | 2080 | 21,14 | 5,3 | 17,17 | 0,0 | 292 | 149 | 1.09E-05 | 0.000353415 | 0.682,0.704 | 1.0,1.0 | -0.307 |
| 7751 | XLOC_034882 | chr5 | - | 40833878 | 40834367 | 40833878 | 40834139 | 40834282 | 143 | 40834367 | 7751 | 8,9 | 30,24 | 30,27 | 39,26 | 292 | 149 | 0.001284459 | 0.014824947 | 0.12,0.161 | 0.282,0.346 | -0.173 |
| 4345 | XLOC_041420 | chr7 | - | 21945580 | 21946074 | 21945580 | 21945762 | 21945906 | 144 | 21946074 | 4345 | 0,4 | 4,3 | 5,8 | 0,1 | 293 | 149 | 8.42E-06 | 0.000283243 | 0.0,0.404 | 1.0,0.803 | -0.7 |
| 10324 | XLOC_021503 | chr19 | - | 6730997 | 6731245 | 6730997 | 6731065 | 6731209 | 144 | 6731245 | 10324 | 24,43 | 5,1 | 62,61 | 1,0 | 293 | 149 | 0.001838997 | 0.019563081 | 0.709,0.956 | 0.969,1.0 | -0.152 |
| 2576 | XLOC_014712 | chr15 | + | 75646993 | 75648036 | 75646993 | 75647159 | 75647304 | 145 | 75648036 | 2576 | 7,5 | 38,40 | 35,32 | 31,45 | 294 | 149 | 1.35E-07 | 8.84E-06 | 0.085,0.06 | 0.364,0.265 | -0.242 |
| 11474 | XLOC_009359 | chr12 | - | 50537734 | 50538147 | 50537734 | 50537840 | 50537985 | 145 | 50538147 | 11474 | 5,0 | 6,1 | 1,6 | 0,0 | 294 | 149 | 1.65E-08 | 1.46E-06 | 0.297,0.0 | 1.0,1.0 | -0.852 |
| 4853 | XLOC_034773 | chr5 | + | 159848933 | 159849410 | 159848933 | 159849162 | 159849308 | 146 | 159849410 | 4853 | 54,50 | 27,41 | 56,33 | 10,11 | 295 | 149 | 0.003105158 | 0.028939145 | 0.503,0.381 | 0.739,0.602 | -0.228 |
| 6046 | XLOC_026812 | chr20 | + | 44441758 | 44442103 | 44441758 | 44441929 | 44442075 | 146 | 44442103 | 6046 | 25,22 | 20,20 | 56,31 | 5,6 | 295 | 149 | 4.96E-07 | 2.67E-05 | 0.387,0.357 | 0.85,0.723 | -0.414 |
| 4503 | XLOC_047701 | chrX | + | 48433948 | 48434397 | 48433948 | 48434055 | 48434202 | 147 | 48434397 | 4503 | 92,71 | 28,36 | 257,229 | 34,19 | 296 | 149 | 5.80E-07 | 3.05E-05 | 0.623,0.498 | 0.792,0.858 | -0.264 |
| 12217 | XLOC_007722 | chr11 | - | 116732917 | 116733293 | 116732917 | 116733043 | 116733190 | 147 | 116733293 | 12217 | 4,2 | 4,0 | 4,1 | 0,0 | 296 | 149 | 0.004739864 | 0.039905334 | 0.335,1.0 | 1.0,1.0 | -0.333 |
| 9619 | XLOC_014857 | chr15 | - | 41809995 | 41810402 | 41809995 | 41810055 | 41810203 | 148 | 41810402 | 9619 | 2,4 | 43,34 | 35,26 | 38,65 | 297 | 149 | 2.15E-07 | 1.34E-05 | 0.023,0.056 | 0.316,0.167 | -0.202 |
| 11904 | XLOC_008751 | chr12 | + | 6643635 | 6644027 | 6643635 | 6643827 | 6643975 | 148 | 6644027 | 11904 | 103,134 | 128,114 | 201,184 | 135,111 | 297 | 149 | 0.004886059 | 0.040730484 | 0.288,0.371 | 0.428,0.454 | -0.111 |
| 8846 | XLOC_016188 | chr16 | + | 722702 | 723134 | 722702 | 722825 | 722975 | 150 | 723134 | 8846 | 6,5 | 1,9 | 5,10 | 0,0 | 299 | 149 | 2.19E-06 | 9.37E-05 | 0.749,0.217 | 1.0,1.0 | -0.517 |
| 16243 | XLOC_047661 | chrX | + | 21985003 | 21985434 | 21985003 | 21985163 | 21985313 | 150 | 21985434 | 16243 | 6,6 | 90,72 | 7,7 | 30,16 | 299 | 149 | 0.002417291 | 0.024055364 | 0.032,0.04 | 0.104,0.179 | -0.105 |
| 12950 | XLOC_028747 | chr22 | - | 50721432 | 50721874 | 50721432 | 50721612 | 50721765 | 153 | 50721874 | 12950 | 20,24 | 1,5 | 16,44 | 0,0 | 302 | 149 | 0.000144326 | 0.002801237 | 0.908,0.703 | 1.0,1.0 | -0.195 |
| 7728 | XLOC_028532 | chr22 | + | 45717927 | 45718307 | 45717927 | 45718096 | 45718251 | 155 | 45718307 | 7728 | 0,2 | 5,6 | 3,2 | 2,0 | 304 | 149 | 0.002370221 | 0.023705363 | 0.0,0.14 | 0.424,1.0 | -0.642 |
| 13929 | XLOC_007065 | chr11 | + | 45948865 | 45949360 | 45948865 | 45949144 | 45949299 | 155 | 45949360 | 13929 | 5,5 | 8,0 | 6,5 | 0,0 | 304 | 149 | 0.001141569 | 0.01355286 | 0.234,1.0 | 1.0,1.0 | -0.383 |
| 14149 | XLOC_029987 | chr3 | - | 121386907 | 121387275 | 121386907 | 121387005 | 121387160 | 155 | 121387275 | 14149 | 0,0 | 10,4 | 3,2 | 1,2 | 304 | 149 | 0.000128343 | 0.002532667 | 0.0,0.0 | 0.595,0.329 | -0.462 |
| 8788 | XLOC_016862 | chr16 | - | 89808927 | 89809346 | 89808927 | 89809051 | 89809207 | 156 | 89809346 | 8788 | 10,9 | 1,4 | 9,11 | 1,0 | 305 | 149 | 0.002221287 | 0.022500071 | 0.83,0.524 | 0.815,1.0 | -0.23 |
| 12347 | XLOC_043957 | chr8 | + | 21937922 | 21938136 | 21937922 | 21937971 | 21938128 | 157 | 21938136 | 12347 | 32,23 | 12,5 | 10,8 | 0,0 | 306 | 149 | 7.80E-06 | 0.00026501 | 0.565,0.691 | 1.0,1.0 | -0.372 |
| 7536 | XLOC_047975 | chrX | - | 57004724 | 57005093 | 57004724 | 57004854 | 57005012 | 158 | 57005093 | 7536 | 0,0 | 3,1 | 9,4 | 2,0 | 307 | 149 | 1.79E-08 | 1.57E-06 | 0.0,0.0 | 0.686,1.0 | -0.843 |
| 15064 | XLOC_018794 | chr17 | - | 80203851 | 80204341 | 80203851 | 80203915 | 80204073 | 158 | 80204341 | 15064 | 105,139 | 83,87 | 194,181 | 37,46 | 307 | 149 | 2.71E-10 | 4.24E-08 | 0.38,0.437 | 0.718,0.656 | -0.279 |
| 16215 | XLOC_000284 | chr1 | + | 46739032 | 46739419 | 46739032 | 46739137 | 46739295 | 158 | 46739419 | 16215 | 47,49 | 94,96 | 88,89 | 98,91 | 307 | 149 | 0.00102962 | 0.012560859 | 0.195,0.199 | 0.304,0.322 | -0.116 |
| 8695 | XLOC_047679 | chrX | + | 41064555 | 41064936 | 41064555 | 41064746 | 41064905 | 159 | 41064936 | 8695 | 0,1 | 2,1 | 1,1 | 0,0 | 308 | 149 | 1.07E-06 | 5.13E-05 | 0.0,0.326 | 1.0,1.0 | -0.837 |
| 8792 | XLOC_016862 | chr16 | - | 89851261 | 89851567 | 89851261 | 89851372 | 89851531 | 159 | 89851567 | 8792 | 0,7 | 4,1 | 5,4 | 0,0 | 308 | 149 | 1.15E-05 | 0.000371005 | 0.0,0.772 | 1.0,1.0 | -0.614 |
| 10092 | XLOC_007211 | chr11 | + | 71939770 | 71940080 | 71939770 | 71939891 | 71940051 | 160 | 71940080 | 10092 | 87,107 | 74,67 | 130,132 | 63,42 | 309 | 149 | 0.004293973 | 0.037094418 | 0.362,0.435 | 0.499,0.602 | -0.152 |
| 5512 | XLOC_032741 | chr4 | - | 122591108 | 122591377 | 122591108 | 122591167 | 122591328 | 161 | 122591377 | 5512 | 6,3 | 1,5 | 8,10 | 0,0 | 310 | 149 | 6.17E-06 | 0.000218375 | 0.743,0.224 | 1.0,1.0 | -0.516 |
| 7929 | XLOC_005345 | chr10 | - | 101945270 | 101945645 | 101945270 | 101945394 | 101945555 | 161 | 101945645 | 7929 | 2,4 | 2,4 | 8,11 | 1,0 | 310 | 149 | 1.65E-05 | 0.000487948 | 0.325,0.325 | 0.794,1.0 | -0.572 |
| 10020 | XLOC_007716 | chr11 | - | 113678672 | 113679159 | 113678672 | 113678858 | 113679019 | 161 | 113679159 | 10020 | 0,1 | 1,3 | 6,3 | 0,0 | 310 | 149 | 4.38E-09 | 4.84E-07 | 0.0,0.138 | 1.0,1.0 | -0.931 |
| 4583 | XLOC_038232 | chr6 | - | 33545311 | 33545685 | 33545311 | 33545412 | 33545574 | 162 | 33545685 | 4583 | 0,0 | 1,6 | 4,2 | 2,0 | 311 | 149 | 5.82E-06 | 0.000210474 | 0.0,0.0 | 0.489,1.0 | -0.744 |
| 12844 | XLOC_006942 | chr11 | + | 380100 | 380445 | 380100 | 380129 | 380291 | 162 | 380445 | 12844 | 27,17 | 1,4 | 21,28 | 0,0 | 311 | 149 | 0.001470696 | 0.016435519 | 0.928,0.671 | 1.0,1.0 | -0.2 |
| 3639 | XLOC_047816 | chrX | + | 110970583 | 110970998 | 110970583 | 110970673 | 110970841 | 168 | 110970998 | 3639 | 0,2 | 26,21 | 12,6 | 27,37 | 317 | 149 | 0.000216868 | 0.003855951 | 0.0,0.043 | 0.173,0.071 | -0.101 |
| 12612 | XLOC_007214 | chr11 | + | 73074791 | 73075284 | 73074791 | 73075047 | 73075216 | 169 | 73075284 | 12612 | 13,13 | 3,1 | 10,7 | 0,0 | 318 | 149 | 0.00446648 | 0.038260136 | 0.67,0.859 | 1.0,1.0 | -0.236 |
| 15968 | XLOC_014828 | chr15 | - | 34653369 | 34653798 | 34653369 | 34653430 | 34653600 | 170 | 34653798 | 15968 | 66,74 | 6,14 | 93,80 | 5,2 | 319 | 149 | 0.00304983 | 0.028511836 | 0.837,0.712 | 0.897,0.949 | -0.149 |
| 3177 | XLOC_026791 | chr20 | + | 39985679 | 39986087 | 39985679 | 39985828 | 39986000 | 172 | 39986087 | 3177 | 28,38 | 29,19 | 61,50 | 2,7 | 321 | 149 | 2.56E-07 | 1.56E-05 | 0.309,0.481 | 0.934,0.768 | -0.456 |
| 2554 | XLOC_007119 | chr11 | + | 62556494 | 62557875 | 62556494 | 62556898 | 62557072 | 174 | 62557875 | 2554 | 21,20 | 82,106 | 46,40 | 64,83 | 323 | 149 | 2.56E-05 | 0.000704955 | 0.106,0.08 | 0.249,0.182 | -0.122 |
| 4856 | XLOC_034759 | chr5 | + | 151179459 | 151179907 | 151179459 | 151179604 | 151179778 | 174 | 151179907 | 4856 | 11,6 | 0,5 | 9,7 | 0,0 | 323 | 149 | 0.002465711 | 0.024333733 | 1.0,0.356 | 1.0,1.0 | -0.322 |
| 5180 | XLOC_030129 | chr3 | - | 184069813 | 184070117 | 184069813 | 184069905 | 184070080 | 175 | 184070117 | 5180 | 1,2 | 26,29 | 13,9 | 29,31 | 324 | 149 | 6.31E-05 | 0.001436213 | 0.017,0.031 | 0.171,0.118 | -0.121 |
| 10765 | XLOC_018709 | chr17 | - | 73126847 | 73127193 | 73126847 | 73126962 | 73127137 | 175 | 73127193 | 10765 | 72,81 | 8,2 | 71,103 | 0,1 | 324 | 149 | 0.000262886 | 0.004478289 | 0.805,0.949 | 1.0,0.979 | -0.113 |
| 12227 | XLOC_021877 | chr19 | - | 55558472 | 55558856 | 55558472 | 55558579 | 55558754 | 175 | 55558856 | 12227 | 25,31 | 4,4 | 53,42 | 0,0 | 324 | 149 | 1.50E-05 | 0.000450719 | 0.742,0.781 | 1.0,1.0 | -0.238 |
| 3898 | XLOC_046087 | chr9 | + | 131831006 | 131831408 | 131831006 | 131831178 | 131831354 | 176 | 131831408 | 3898 | 4,8 | 4,0 | 2,3 | 0,0 | 325 | 149 | 0.006130751 | 0.048130878 | 0.314,1.0 | 1.0,1.0 | -0.343 |
| 10093 | XLOC_007211 | chr11 | + | 71939770 | 71940080 | 71939770 | 71939891 | 71940067 | 176 | 71940080 | 10093 | 61,76 | 46,72 | 116,112 | 32,37 | 325 | 149 | 4.90E-07 | 2.67E-05 | 0.378,0.326 | 0.624,0.581 | -0.251 |
| 1453 | XLOC_009585 | chr12 | - | 121864460 | 121868272 | 121864460 | 121867966 | 121868144 | 178 | 121868272 | 1453 | 137,116 | 9,14 | 176,195 | 1,5 | 327 | 149 | 2.00E-05 | 0.000573554 | 0.874,0.791 | 0.988,0.947 | -0.135 |
| 13358 | XLOC_044359 | chr8 | - | 93904216 | 93904520 | 93904216 | 93904302 | 93904481 | 179 | 93904520 | 13358 | 0,4 | 2,1 | 1,1 | 0,0 | 328 | 149 | 0.000428216 | 0.006454289 | 0.0,0.645 | 1.0,1.0 | -0.677 |
| 3549 | XLOC_048083 | chrX | - | 130928021 | 130928494 | 130928021 | 130928143 | 130928324 | 181 | 130928494 | 3549 | 1,0 | 8,3 | 23,19 | 16,15 | 330 | 149 | 2.89E-06 | 0.000116877 | 0.053,0.0 | 0.394,0.364 | -0.352 |
| 6598 | XLOC_023898 | chr2 | + | 234356503 | 234356827 | 234356503 | 234356551 | 234356732 | 181 | 234356827 | 6598 | 5,14 | 67,68 | 18,40 | 36,17 | 330 | 149 | 1.41E-05 | 0.000429121 | 0.033,0.085 | 0.184,0.515 | -0.291 |
| 11178 | XLOC_007597 | chr11 | - | 66134581 | 66135019 | 66134581 | 66134753 | 66134934 | 181 | 66135019 | 11178 | 34,52 | 7,0 | 47,55 | 0,0 | 330 | 149 | 0.000533202 | 0.007602299 | 0.687,1.0 | 1.0,1.0 | -0.156 |
| 15296 | XLOC_001334 | chr1 | - | 153913379 | 153913870 | 153913379 | 153913524 | 153913706 | 182 | 153913870 | 15296 | 32,33 | 66,60 | 67,73 | 65,46 | 331 | 149 | 0.000114746 | 0.002329292 | 0.179,0.198 | 0.317,0.417 | -0.178 |
| 4935 | XLOC_034481 | chr5 | + | 647919 | 648338 | 647919 | 648031 | 648214 | 183 | 648338 | 4935 | 12,10 | 18,17 | 13,10 | 2,5 | 332 | 149 | 0.00263591 | 0.025596748 | 0.23,0.209 | 0.745,0.473 | -0.389 |
| 6108 | XLOC_043952 | chr8 | + | 19675385 | 19675853 | 19675385 | 19675602 | 19675785 | 183 | 19675853 | 6108 | 8,6 | 3,1 | 15,9 | 0,1 | 332 | 149 | 0.003252071 | 0.029937366 | 0.545,0.729 | 1.0,0.802 | -0.264 |
| 12050 | XLOC_038395 | chr6 | - | 109766973 | 109767615 | 109766973 | 109767155 | 109767338 | 183 | 109767615 | 12050 | 0,1 | 3,1 | 2,1 | 0,0 | 332 | 149 | 4.59E-09 | 5.04E-07 | 0.0,0.31 | 1.0,1.0 | -0.845 |
| 6115 | XLOC_026682 | chr20 | + | 2636783 | 2637195 | 2636783 | 2636860 | 2637046 | 186 | 2637195 | 6115 | 159,136 | 157,144 | 313,362 | 196,202 | 335 | 149 | 0.000128469 | 0.002532667 | 0.311,0.296 | 0.415,0.444 | -0.126 |

| | | | | | | | | | | | | | | | | | | | | | | |
|---|---|---|---|---|---|---|---|---|---|---|---|---|---|---|---|---|---|---|---|---|---|---|
| 11732 | XLOC_008937 | chr12 | + | 56334035 | 56334506 | 56334035 | 56334217 | 56334407 | 190 | 56334506 | 11732 | 9,10 | 79,72 | 15,18 | 30,47 | 339 | 149 | 0.000173689 | 0.003261717 | 0.048,0.058 | 0.18,0.144 | -0.109 |
| 3960 | XLOC_045975 | chr9 | + | 97842975 | 97843454 | 97842975 | 97843062 | 97843253 | 191 | 97843454 | 3960 | 11,17 | 2,2 | 5,5 | 0,0 | 340 | 149 | 0.003682435 | 0.032990374 | 0.707,0.788 | 1.0,1.0 | -0.252 |
| 7420 | XLOC_020958 | chr19 | + | 9939002 | 9939348 | 9939002 | 9939069 | 9939260 | 191 | 9939348 | 7420 | 15,29 | 1,3 | 29,19 | 0,0 | 340 | 149 | 0.00566201 | 0.045599975 | 0.868,0.809 | 1.0,1.0 | -0.161 |
| 7640 | XLOC_020063 | chr18 | + | 21526107 | 21526540 | 21526107 | 21526248 | 21526440 | 192 | 21526540 | 7640 | 5,0 | 1,2 | 2,1 | 0,0 | 341 | 149 | 0.000604824 | 0.008361919 | 0.686,0.0 | 1.0,1.0 | -0.657 |
| 6520 | XLOC_024051 | chr2 | - | 61719710 | 61720188 | 61719710 | 61719856 | 61720049 | 193 | 61720188 | 6520 | 5,3 | 10,5 | 2,13 | 1,0 | 342 | 149 | 3.69E-05 | 0.000954575 | 0.179,0.207 | 0.466,1.0 | -0.54 |
| 7421 | XLOC_020958 | chr19 | + | 9939002 | 9939348 | 9939002 | 9939069 | 9939262 | 193 | 9939348 | 7421 | 15,29 | 1,11 | 29,19 | 1,0 | 342 | 149 | 0.000149057 | 0.002863501 | 0.867,0.535 | 0.927,1.0 | -0.262 |
| 7500 | XLOC_016577 | chr16 | - | 2162788 | 2163285 | 2162788 | 2162964 | 2163157 | 193 | 2163285 | 7500 | 8,2 | 21,13 | 18,12 | 13,18 | 342 | 149 | 0.005425628 | 0.044253352 | 0.142,0.063 | 0.376,0.225 | -0.198 |
| 10492 | XLOC_044467 | chr8 | - | 145753407 | 145753819 | 145753407 | 145753456 | 145753651 | 195 | 145753819 | 10492 | 38,38 | 4,3 | 68,68 | 0,1 | 344 | 149 | 0.000391229 | 0.006029578 | 0.804,0.846 | 1.0,0.967 | -0.159 |
| 674 | XLOC_028745 | chr22 | - | 50689199 | 50691909 | 50689199 | 50689334 | 50689530 | 196 | 50691909 | 674 | 83,84 | 21,13 | 129,150 | 11,11 | 345 | 149 | 0.002510367 | 0.024615994 | 0.631,0.736 | 0.835,0.855 | -0.161 |
| 9808 | XLOC_018520 | chr17 | - | 37009274 | 37009967 | 37009274 | 37009358 | 37009554 | 196 | 37009967 | 9808 | 23,24 | 4,2 | 62,45 | 0,1 | 345 | 149 | 0.000498082 | 0.007224832 | 0.713,0.838 | 1.0,0.951 | -0.2 |
| 7501 | XLOC_016577 | chr16 | - | 2162788 | 2163285 | 2162788 | 2162964 | 2163161 | 197 | 2163285 | 7501 | 8,3 | 53,45 | 22,16 | 58,38 | 346 | 149 | 0.000215923 | 0.003848235 | 0.061,0.028 | 0.14,0.153 | -0.102 |
| 11093 | XLOC_013429 | chr14 | - | 68059031 | 68059435 | 68059031 | 68059153 | 68059351 | 198 | 68059435 | 11093 | 0,0 | 2,4 | 11,2 | 6,5 | 347 | 149 | 4.07E-05 | 0.001029275 | 0.0,0.0 | 0.44,0.147 | -0.293 |
| 2130 | XLOC_007597 | chr11 | - | 66133596 | 66134035 | 66133596 | 66133702 | 66133901 | 199 | 66134035 | 2130 | 78,77 | 106,130 | 122,106 | 102,75 | 348 | 149 | 0.000104229 | 0.002150628 | 0.24,0.202 | 0.339,0.377 | -0.137 |
| 9973 | XLOC_007774 | chr11 | - | 134026646 | 134026990 | 134026646 | 134026713 | 134026912 | 199 | 134026990 | 9973 | 35,25 | 0,5 | 25,30 | 0,0 | 348 | 149 | 0.003992231 | 0.03495032 | 1.0,0.682 | 1.0,1.0 | -0.159 |
| 13771 | XLOC_004802 | chr10 | + | 28822926 | 28823275 | 28822926 | 28822963 | 28823162 | 199 | 28823275 | 13771 | 52,40 | 7,2 | 57,62 | 2,0 | 348 | 149 | 0.002057542 | 0.021227397 | 0.761,0.895 | 0.924,1.0 | -0.134 |
| 10304 | XLOC_029885 | chr3 | - | 53853215 | 53853701 | 53853215 | 53853358 | 53853558 | 200 | 53853701 | 10304 | 2,4 | 3,4 | 13,14 | 3,1 | 349 | 149 | 0.001903209 | 0.020118107 | 0.222,0.299 | 0.649,0.857 | -0.492 |
| 458 | XLOC_041145 | chr7 | + | 75044869 | 75045469 | 75044869 | 75045056 | 75045258 | 202 | 75045469 | 458 | 208,241 | 33,44 | 336,358 | 14,18 | 351 | 149 | 9.26E-09 | 8.71E-07 | 0.728,0.699 | 0.911,0.894 | -0.189 |
| 4296 | XLOC_041531 | chr7 | - | 72419430 | 72420026 | 72419430 | 72419641 | 72419843 | 202 | 72420026 | 4296 | 48,61 | 36,24 | 73,71 | 8,4 | 351 | 149 | 1.24E-07 | 8.29E-06 | 0.361,0.519 | 0.795,0.883 | -0.399 |
| 445 | XLOC_034514 | chr5 | + | 35709049 | 35710594 | 35709049 | 35709223 | 35709426 | 203 | 35710594 | 445 | 17,23 | 2,4 | 12,18 | 0,0 | 352 | 149 | 0.000178437 | 0.003309556 | 0.783,0.709 | 1.0,1.0 | -0.254 |
| 2115 | XLOC_044420 | chr8 | - | 134487961 | 134488843 | 134487961 | 134488317 | 134488520 | 203 | 134488843 | 2115 | 19,14 | 43,39 | 97,66 | 101,67 | 352 | 149 | 0.000218134 | 0.003873853 | 0.158,0.132 | 0.289,0.294 | -0.146 |
| 2199 | XLOC_009410 | chr12 | - | 56720096 | 56720743 | 56720096 | 56720193 | 56720400 | 207 | 56720743 | 2199 | 27,38 | 92,80 | 60,63 | 71,84 | 356 | 149 | 0.000380221 | 0.005888823 | 0.109,0.166 | 0.261,0.239 | -0.112 |
| 6371 | XLOC_024443 | chr2 | - | 233349908 | 233350372 | 233349908 | 233349977 | 233350184 | 207 | 233350372 | 6371 | 20,14 | 20,30 | 106,92 | 23,14 | 356 | 149 | 7.95E-11 | 1.50E-08 | 0.295,0.163 | 0.659,0.733 | -0.467 |
| 15218 | XLOC_005275 | chr10 | - | 79579078 | 79579776 | 79579078 | 79579445 | 79579652 | 207 | 79579776 | 15218 | 4,1 | 20,20 | 15,7 | 19,3 | 356 | 149 | 7.90E-05 | 0.001723812 | 0.077,0.02 | 0.248,0.494 | -0.323 |
| 15749 | XLOC_007017 | chr11 | + | 18418026 | 18418515 | 18418026 | 18418157 | 18418365 | 208 | 18418515 | 15749 | 14,17 | 13,11 | 16,8 | 3,1 | 357 | 149 | 0.003150057 | 0.029295865 | 0.31,0.392 | 0.69,0.77 | -0.379 |
| 457 | XLOC_041145 | chr7 | + | 75044869 | 75045469 | 75044869 | 75045048 | 75045258 | 210 | 75045469 | 457 | 211,241 | 53,34 | 340,367 | 19,12 | 359 | 149 | 6.88E-09 | 7.04E-07 | 0.623,0.746 | 0.881,0.927 | -0.22 |
| 12541 | XLOC_000835 | chr1 | - | 1647784 | 1648250 | 1647784 | 1647917 | 1648127 | 210 | 1648250 | 12541 | 8,4 | 3,1 | 7,11 | 1,0 | 359 | 149 | 0.002159978 | 0.022033115 | 0.525,0.624 | 0.744,1.0 | -0.297 |
| 13305 | XLOC_021144 | chr19 | + | 36215539 | 36215988 | 36215539 | 36215679 | 36215889 | 210 | 36215988 | 13305 | 4,0 | 5,2 | 3,3 | 0,1 | 359 | 149 | 0.000322581 | 0.00520071 | 0.249,0.0 | 1.0,0.555 | -0.653 |
| 9872 | XLOC_018149 | chr17 | + | 48456139 | 48456585 | 48456139 | 48456256 | 48456467 | 211 | 48456585 | 9872 | 62,59 | 101,98 | 91,102 | 75,77 | 360 | 149 | 6.95E-05 | 0.001561627 | 0.203,0.199 | 0.334,0.354 | -0.143 |
| 2408 | XLOC_007764 | chr11 | - | 129744395 | 129744811 | 129744395 | 129744475 | 129744687 | 212 | 129744811 | 2408 | 19,10 | 83,89 | 45,51 | 79,84 | 361 | 149 | 2.37E-07 | 1.46E-05 | 0.086,0.044 | 0.19,0.2 | -0.13 |
| 3788 | XLOC_046425 | chr9 | - | 130690343 | 130690762 | 130690343 | 130690498 | 130690711 | 213 | 130690762 | 3788 | 0,0 | 2,1 | 4,1 | 6,5 | 362 | 149 | 0.006244736 | 0.048709679 | 0.0,0.0 | 0.215,0.076 | -0.145 |
| 3376 | XLOC_009413 | chr12 | - | 56814746 | 56815277 | 56814746 | 56814918 | 56815134 | 216 | 56815277 | 3376 | 108,103 | 399,382 | 328,324 | 394,337 | 365 | 149 | 0 | 0 | 0.1,0.099 | 0.254,0.282 | -0.169 |
| 8 | XLOC_048030 | chrX | - | 100653357 | 100653934 | 100653357 | 100653555 | 100653772 | 217 | 100653934 | 8 | 10,8 | 87,73 | 33,54 | 64,68 | 366 | 149 | 1.02E-10 | 1.74E-08 | 0.045,0.043 | 0.173,0.244 | -0.164 |
| 6795 | XLOC_004772 | chr10 | + | 13167417 | 13167836 | 13167417 | 13167567 | 13167784 | 217 | 13167836 | 6795 | 7,6 | 1,3 | 6,10 | 0,1 | 366 | 149 | 0.003395139 | 0.030926268 | 0.74,0.449 | 1.0,0.803 | -0.307 |
| 14712 | XLOC_021319 | chr19 | + | 50169956 | 50170418 | 50169956 | 50170056 | 50170275 | 219 | 50170418 | 14712 | 37,31 | 127,135 | 57,68 | 83,94 | 368 | 149 | 6.72E-07 | 3.39E-05 | 0.106,0.085 | 0.218,0.227 | -0.127 |
| 13495 | XLOC_005284 | chr10 | - | 88827482 | 88827914 | 88827482 | 88827599 | 88827819 | 220 | 88827914 | 13495 | 1,0 | 0,3 | 2,1 | 0,1 | 369 | 149 | 0.000753265 | 0.00988709 | 1.0,0.0 | 1.0,0.288 | -0.144 |
| 16120 | XLOC_000275 | chr1 | + | 45478963 | 45479367 | 45478963 | 45479026 | 45479246 | 220 | 45479367 | 16120 | 5,3 | 17,30 | 25,21 | 36,54 | 369 | 149 | 0.002446297 | 0.024256554 | 0.106,0.039 | 0.219,0.136 | -0.105 |
| 1177 | XLOC_020248 | chr18 | - | 60985314 | 60987270 | 60985314 | 60986185 | 60986406 | 221 | 60987270 | 1177 | 18,11 | 31,23 | 40,55 | 10,25 | 370 | 149 | 9.38E-07 | 4.54E-05 | 0.19,0.161 | 0.617,0.47 | -0.368 |
| 720 | XLOC_027824 | chr21 | - | 47666220 | 47666880 | 47666220 | 47666318 | 47666541 | 223 | 47666880 | 720 | 82,93 | 61,77 | 135,156 | 53,62 | 372 | 149 | 0.00019926 | 0.003606821 | 0.35,0.326 | 0.505,0.502 | -0.166 |
| 5010 | XLOC_032596 | chr4 | - | 39507237 | 39507681 | 39507237 | 39507368 | 39507591 | 223 | 39507681 | 5010 | 0,0 | 4,1 | 5,2 | 7,10 | 372 | 149 | 0.001227133 | 0.014275746 | 0.0,0.0 | 0.222,0.074 | -0.148 |
| 13940 | XLOC_000343 | chr1 | + | 76216135 | 76216576 | 76216135 | 76216231 | 76216454 | 223 | 76216576 | 13940 | 1,2 | 5,1 | 6,6 | 1,1 | 372 | 149 | 0.001763716 | 0.018949866 | 0.074,0.445 | 0.706,0.706 | -0.446 |
| 15070 | XLOC_020938 | chr19 | + | 7599778 | 7600088 | 7599778 | 7599788 | 7600012 | 224 | 7600088 | 15070 | 0,2 | 3,0 | 6,4 | 1,0 | 373 | 149 | 0.003349595 | 0.030666227 | 0.0,1.0 | 0.706,1.0 | -0.353 |
| 9990 | XLOC_007571 | chr11 | - | 64670759 | 64671132 | 64670759 | 64670836 | 64671061 | 225 | 64671132 | 9990 | 7,9 | 38,46 | 36,14 | 41,28 | 374 | 149 | 5.77E-05 | 0.001340492 | 0.068,0.072 | 0.259,0.166 | -0.143 |
| 6540 | XLOC_000844 | chr1 | - | 2316068 | 2316504 | 2316068 | 2316190 | 2316416 | 226 | 2316504 | 6540 | 10,7 | 2,1 | 7,3 | 0,0 | 375 | 149 | 0.005613908 | 0.04530279 | 0.665,0.736 | 1.0,1.0 | -0.299 |
| 7003 | XLOC_021821 | chr19 | - | 49950280 | 49950674 | 49950280 | 49950356 | 49950583 | 227 | 49950674 | 7003 | 33,31 | 3,3 | 20,27 | 0,0 | 376 | 149 | 0.000701409 | 0.009361665 | 0.813,0.804 | 1.0,1.0 | -0.192 |
| 5322 | XLOC_029818 | chr3 | - | 48453265 | 48453713 | 48453265 | 48453418 | 48453646 | 228 | 48453713 | 5322 | 12,16 | 38,41 | 16,30 | 23,27 | 377 | 149 | 0.002713297 | 0.026145685 | 0.111,0.134 | 0.216,0.305 | -0.138 |
| 15337 | XLOC_034803 | chr5 | + | 176520138 | 176520502 | 176520138 | 176520178 | 176520406 | 228 | 176520502 | 15337 | 84,81 | 9,1 | 119,96 | 0,0 | 377 | 149 | 5.08E-05 | 0.00121946 | 0.787,0.97 | 1.0,1.0 | -0.121 |
| 23 | XLOC_047880 | chrX | + | 153641543 | 153641904 | 153641543 | 153641589 | 153641818 | 229 | 153641904 | 23 | 34,38 | 98,73 | 115,89 | 141,97 | 378 | 149 | 0.000202668 | 0.0036597 | 0.12,0.17 | 0.243,0.266 | -0.109 |
| 2899 | XLOC_014590 | chr15 | + | 42128672 | 42129138 | 42128672 | 42128749 | 42128978 | 229 | 42129138 | 2899 | 10,29 | 16,20 | 38,53 | 16,14 | 378 | 149 | 0.004075813 | 0.035561706 | 0.198,0.364 | 0.484,0.599 | -0.26 |
| 12147 | XLOC_000746 | chr1 | + | 224548196 | 224548667 | 224548196 | 224548265 | 224548494 | 229 | 224548667 | 12147 | 4,9 | 7,0 | 6,2 | 0,0 | 378 | 149 | 0.001665968 | 0.018178284 | 0.184,1.0 | 1.0,1.0 | -0.408 |
| 4506 | XLOC_047701 | chrX | + | 48434309 | 48434807 | 48434309 | 48434471 | 48434701 | 230 | 48434807 | 4506 | 60,60 | 43,34 | 137,157 | 40,59 | 379 | 149 | 0.002023397 | 0.021048361 | 0.354,0.41 | 0.574,0.511 | -0.16 |
| 7873 | XLOC_018791 | chr17 | - | 80047125 | 80047602 | 80047125 | 80047277 | 80047507 | 230 | 80047602 | 7873 | 14,8 | 0,7 | 14,15 | 0,0 | 379 | 149 | 0.001064078 | 0.012897549 | 1.0,0.31 | 1.0,1.0 | -0.345 |
| 9797 | XLOC_023552 | chr2 | + | 85598213 | 85598685 | 85598213 | 85598332 | 85598562 | 230 | 85598685 | 9797 | 2,0 | 4,12 | 2,11 | 8,9 | 379 | 149 | 0.00276052 | 0.02646877 | 0.164,0.0 | 0.089,0.325 | -0.125 |
| 3986 | XLOC_045902 | chr9 | + | 35800018 | 35800480 | 35800018 | 35800154 | 35800385 | 231 | 35800480 | 3986 | 0,1 | 6,4 | 5,3 | 5,15 | 380 | 149 | 0.005662861 | 0.045599975 | 0.0,0.089 | 0.282,0.073 | -0.133 |
| 262 | XLOC_038373 | chr6 | - | 99845590 | 99849506 | 99845590 | 99848187 | 99848419 | 232 | 99849506 | 262 | 364,303 | 141,143 | 337,330 | 52,92 | 381 | 149 | 0.000440597 | 0.006594491 | 0.502,0.453 | 0.717,0.584 | -0.173 |
| 1980 | XLOC_009388 | chr12 | - | 54107484 | 54108025 | 54107484 | 54107684 | 54107916 | 232 | 54108025 | 1980 | 139,108 | 175,163 | 210,144 | 92,87 | 381 | 149 | 2.88E-09 | 3.35E-07 | 0.237,0.206 | 0.472,0.393 | -0.211 |
| 2749 | XLOC_000004 | chr1 | + | 897008 | 898297 | 897008 | 897851 | 898083 | 232 | 898297 | 2749 | 18,11 | 20,27 | 50,69 | 29,27 | 381 | 149 | 0.00011585 | 0.002345372 | 0.26,0.137 | 0.403,0.5 | -0.253 |
| 5465 | XLOC_029402 | chr3 | + | 45043029 | 45043506 | 45043029 | 45043100 | 45043333 | 233 | 45043506 | 5465 | 8,6 | 4,0 | 4,9 | 0,0 | 382 | 149 | 0.004632813 | 0.039371061 | 0.438,1.0 | 1.0,1.0 | -0.281 |
| 10508 | XLOC_027816 | chr21 | - | 46274786 | 46275171 | 46274786 | 46274891 | 46275124 | 233 | 46275171 | 10508 | 13,16 | 5,1 | 7,8 | 0,0 | 382 | 149 | 6.96E-05 | 0.001561627 | 0.504,0.862 | 1.0,1.0 | -0.317 |
| 2890 | XLOC_016217 | chr16 | + | 2283011 | 2283991 | 2283011 | 2283648 | 2283882 | 234 | 2283991 | 2890 | 14,32 | 72,69 | 39,73 | 72,67 | 383 | 149 | 0.000667884 | 0.009060008 | 0.07,0.153 | 0.174,0.298 | -0.124 |
| 526 | XLOC_044238 | chr8 | - | 21963963 | 21965857 | 21963963 | 21965406 | 21965643 | 237 | 21965857 | 526 | 4,8 | 9,20 | 25,26 | 24,15 | 386 | 149 | 0.002694867 | 0.026020549 | 0.146,0.134 | 0.287,0.401 | -0.204 |
| 6489 | XLOC_014964 | chr15 | - | 68503600 | 68504079 | 68503600 | 68503656 | 68503893 | 237 | 68504079 | 6489 | 31,23 | 15,12 | 58,63 | 5,12 | 386 | 149 | 0.000637402 | 0.008708263 | 0.444,0.425 | 0.817,0.67 | -0.309 |
| 6690 | XLOC_023576 | chr2 | + | 96688421 | 96689121 | 96688421 | 96688501 | 96688739 | 238 | 96689121 | 6690 | 1,6 | 0,4 | 3,4 | 0,0 | 387 | 149 | 0.003045875 | 0.028492571 | 1.0,0.366 | 1.0,1.0 | -0.317 |
| 11984 | XLOC_009604 | chr12 | - | 124109333 | 124109767 | 124109333 | 124109409 | 124109647 | 238 | 124109767 | 11984 | 22,20 | 6,1 | 22,32 | 0,0 | 387 | 149 | 4.04E-05 | 0.001024736 | 0.585,0.885 | 1.0,1.0 | -0.265 |
| 8966 | XLOC_016541 | chr16 | + | 89976619 | 89977071 | 89976619 | 89976759 | 89976998 | 239 | 89977071 | 8966 | 30,18 | 8,11 | 23,35 | 0,4 | 388 | 149 | 0.001118796 | 0.013363373 | 0.59,0.386 | 1.0,0.771 | -0.397 |
| 3028 | XLOC_048047 | chrX | - | 106361995 | 106362795 | 106361995 | 106362132 | 106362372 | 240 | 106362795 | 3028 | 30,18 | 2,4 | 48,46 | 2,0 | 389 | 149 | 0.00576913 | 0.046134635 | 0.852,0.633 | 0.902,1.0 | -0.209 |
| 430 | XLOC_034597 | chr5 | + | 78908274 | 78908898 | 78908274 | 78908592 | 78908833 | 241 | 78908898 | 430 | 157,210 | 21,15 | 146,131 | 3,7 | 390 | 149 | 0.003155119 | 0.029295865 | 0.741,0.842 | 0.949,0.877 | -0.122 |
| 10847 | XLOC_011260 | chr13 | + | 20244979 | 20245429 | 20244979 | 20245104 | 20245345 | 241 | 20245429 | 10847 | 16,19 | 4,2 | 23,30 | 0,1 | 390 | 149 | 0.000468368 | 0.006902234 | 0.604,0.784 | 1.0,0.92 | -0.266 |
| 11603 | XLOC_009165 | chr12 | + | 123465366 | 123465816 | 123465366 | 123465434 | 123465675 | 241 | 123465816 | 11603 | 118,124 | 26,24 | 280,252 | 21,28 | 390 | 149 | 0.000698501 | 0.009347729 | 0.634,0.664 | 0.836,0.775 | -0.156 |
| 5100 | XLOC_032250 | chr4 | + | 3131689 | 3132150 | 3131689 | 3131789 | 3132031 | 242 | 3132150 | 5100 | 1,0 | 0,2 | 5,2 | 1,0 | 391 | 149 | 0.000717864 | 0.009538961 | 1.0,0.0 | 0.656,1.0 | -0.328 |
| 13139 | XLOC_014541 | chr15 | + | 30927282 | 30927836 | 30927282 | 30927446 | 30927689 | 243 | 30927836 | 13139 | 68,61 | 9,2 | 45,57 | 2,0 | 392 | 149 | 0.004292862 | 0.037094418 | 0.742,0.921 | 0.895,1.0 | -0.116 |
| 13562 | XLOC_029295 | chr3 | + | 5236470 | 5236966 | 5236470 | 5236650 | 5236893 | 243 | 5236966 | 13562 | 1,2 | 3,5 | 11,5 | 3,1 | 392 | 149 | 0.000495076 | 0.007188159 | 0.112,0.132 | 0.582,0.655 | -0.497 |

| | | | | | | | | | | | | | | | | | | | | | | |
|---|---|---|---|---|---|---|---|---|---|---|---|---|---|---|---|---|---|---|---|---|---|---|
| 8829 | XLOC_016188 | chr16 | + | 718357 | 718699 | 718357 | 718411 | 718655 | 244 | 718699 | 8829 | 84,105 | 5,11 | 93,122 | 0,0 | 393 | 149 | 1.98E-07 | 1.25E-05 | 0.864,0.784 | 1.0,1.0 | -0.176 |
| 22 | XLOC_047911 | chrX | - | 18942173 | 18942643 | 18942173 | 18942252 | 18942498 | 246 | 18942643 | 22 | 127,114 | 122,105 | 211,189 | 94,57 | 395 | 149 | 4.78E-07 | 2.63E-05 | 0.282,0.291 | 0.459,0.556 | -0.221 |
| 7897 | XLOC_013249 | chr14 | + | 96991646 | 96992096 | 96991646 | 96991728 | 96991974 | 246 | 96992096 | 7897 | 0,1 | 2,1 | 3,3 | 0,0 | 395 | 149 | 3.77E-07 | 2.16E-05 | 0.0,0.274 | 1.0,1.0 | -0.863 |
| 4940 | XLOC_038429 | chr6 | - | 131201283 | 131201629 | 131201283 | 131201346 | 131201593 | 247 | 131201629 | 4940 | 16,18 | 4,1 | 14,6 | 0,0 | 396 | 149 | 0.0003089 | 0.005045025 | 0.601,0.871 | 1.0,1.0 | -0.264 |
| 10749 | XLOC_023917 | chr2 | + | 242263630 | 242264077 | 242263630 | 242263656 | 242263903 | 247 | 242264077 | 10749 | 10,12 | 3,1 | 14,13 | 0,1 | 396 | 149 | 0.006404871 | 0.049551076 | 0.556,0.819 | 1.0,0.83 | -0.228 |
| 11223 | XLOC_018789 | chr17 | - | 80008537 | 80008892 | 80008537 | 80008640 | 80008887 | 247 | 80008892 | 11223 | 30,12 | 2,3 | 31,34 | 0,0 | 396 | 149 | 0.00047319 | 0.006963821 | 0.849,0.601 | 1.0,1.0 | -0.275 |
| 7649 | XLOC_026673 | chr20 | + | 329892 | 330476 | 329892 | 330007 | 330259 | 252 | 330476 | 7649 | 1,0 | 4,3 | 1,3 | 1,0 | 401 | 149 | 0.000261121 | 0.004458328 | 0.085,0.0 | 0.271,1.0 | -0.593 |
| 1288 | XLOC_018666 | chr17 | - | 61779973 | 61781407 | 61779973 | 61781111 | 61781364 | 253 | 61781407 | 1288 | 22,19 | 29,31 | 57,62 | 29,25 | 402 | 149 | 1.31E-05 | 0.000409879 | 0.219,0.185 | 0.421,0.479 | -0.248 |
| 1582 | XLOC_045894 | chr9 | + | 34660500 | 34660933 | 34660500 | 34660597 | 34660850 | 253 | 34660933 | 1582 | 13,11 | 7,1 | 14,12 | 1,0 | 402 | 149 | 0.000664811 | 0.009041666 | 0.408,0.803 | 0.838,1.0 | -0.314 |
| 2171 | XLOC_041357 | chr7 | + | 154795156 | 154798693 | 154795156 | 154795418 | 154795671 | 253 | 154798693 | 2171 | 376,288 | 27,29 | 270,244 | 2,2 | 402 | 149 | 1.00E-10 | 1.74E-08 | 0.838,0.786 | 0.98,0.978 | -0.167 |
| 9180 | XLOC_016214 | chr16 | + | 2200947 | 2201611 | 2200947 | 2201316 | 2201569 | 253 | 2201611 | 9180 | 7,8 | 5,1 | 8,8 | 0,0 | 402 | 149 | 5.78E-06 | 0.000209406 | 0.342,0.748 | 1.0,1.0 | -0.455 |
| 2888 | XLOC_016217 | chr16 | + | 2283011 | 2283648 | 2283011 | 2283184 | 2283438 | 254 | 2283648 | 2888 | 20,34 | 87,99 | 58,78 | 107,88 | 403 | 149 | 4.31E-05 | 0.001077557 | 0.078,0.113 | 0.167,0.247 | -0.112 |
| 3205 | XLOC_000825 | chr1 | - | 1233392 | 1233801 | 1233392 | 1233493 | 1233749 | 256 | 1233801 | 3205 | 30,51 | 53,47 | 106,118 | 78,54 | 405 | 149 | 0.002591709 | 0.025216352 | 0.172,0.285 | 0.333,0.446 | -0.161 |
| 837 | XLOC_004889 | chr10 | + | 75560397 | 75560953 | 75560397 | 75560517 | 75560774 | 257 | 75560953 | 837 | 17,36 | 81,44 | 55,68 | 52,53 | 406 | 149 | 0.000319489 | 0.005161928 | 0.072,0.231 | 0.28,0.32 | -0.149 |
| 1910 | XLOC_013040 | chr14 | + | 21464059 | 21464451 | 21464059 | 21464110 | 21464367 | 257 | 21464451 | 1910 | 73,56 | 220,327 | 231,203 | 265,283 | 406 | 149 | 1.67E-15 | 8.95E-13 | 0.109,0.059 | 0.242,0.208 | -0.141 |
| 9006 | XLOC_016452 | chr16 | + | 68389625 | 68390118 | 68389625 | 68389786 | 68390043 | 257 | 68390118 | 9006 | 3,1 | 2,2 | 6,2 | 0,0 | 406 | 149 | 2.39E-06 | 9.94E-05 | 0.355,0.155 | 1.0,1.0 | -0.745 |
| 5726 | XLOC_028548 | chr22 | + | 50354965 | 50355420 | 50354965 | 50355016 | 50355275 | 259 | 50355420 | 5726 | 45,20 | 7,0 | 33,45 | 0,0 | 408 | 149 | 0.000252934 | 0.004363107 | 0.701,1.0 | 1.0,1.0 | -0.149 |
| 14553 | XLOC_021229 | chr19 | + | 42364844 | 42365273 | 42364844 | 42364915 | 42365174 | 259 | 42365273 | 14553 | 3,6 | 5,4 | 2,1 | 0,0 | 408 | 149 | 4.88E-09 | 5.28E-07 | 0.18,0.354 | 1.0,1.0 | -0.733 |
| 491 | XLOC_032542 | chr4 | - | 2243246 | 2243898 | 2243246 | 2243517 | 2243777 | 260 | 2243898 | 491 | 74,72 | 16,20 | 141,110 | 6,6 | 409 | 149 | 6.09E-07 | 3.17E-05 | 0.628,0.567 | 0.895,0.87 | -0.285 |
| 12013 | XLOC_021828 | chr19 | - | 50364705 | 50365128 | 50364705 | 50364767 | 50365028 | 261 | 50365128 | 12013 | 82,79 | 9,3 | 55,44 | 1,0 | 410 | 149 | 0.000359688 | 0.005675864 | 0.768,0.905 | 0.952,1.0 | -0.139 |
| 185 | XLOC_041384 | chr7 | - | 2582837 | 2583596 | 2582837 | 2582957 | 2583223 | 266 | 2583596 | 185 | 65,55 | 126,138 | 119,115 | 130,127 | 415 | 149 | 5.08E-05 | 0.00121946 | 0.156,0.125 | 0.247,0.245 | -0.105 |
| 1427 | XLOC_018607 | chr17 | - | 45915017 | 45915816 | 45915017 | 45915368 | 45915635 | 267 | 45915816 | 1427 | 54,89 | 133,152 | 75,67 | 56,54 | 416 | 149 | 5.96E-07 | 3.11E-05 | 0.127,0.173 | 0.324,0.308 | -0.166 |
| 6327 | XLOC_016250 | chr16 | + | 3335719 | 3336149 | 3335719 | 3335754 | 3336022 | 268 | 3336149 | 6327 | 23,28 | 52,63 | 26,51 | 33,31 | 417 | 149 | 0.001083394 | 0.013026704 | 0.136,0.137 | 0.22,0.37 | -0.158 |
| 13374 | XLOC_030032 | chr3 | - | 129115882 | 129116378 | 129115882 | 129115983 | 129116253 | 270 | 129116378 | 13374 | 24,17 | 8,6 | 25,31 | 0,2 | 419 | 149 | 3.39E-05 | 0.000895878 | 0.516,0.502 | 1.0,0.846 | -0.414 |
| 2775 | XLOC_000718 | chr1 | + | 213414044 | 213416001 | 213414044 | 213415640 | 213415911 | 271 | 213416001 | 2775 | 23,29 | 52,48 | 71,55 | 58,61 | 420 | 149 | 0.00180834 | 0.019306727 | 0.136,0.177 | 0.303,0.242 | -0.116 |
| 14127 | XLOC_000344 | chr1 | + | 76259768 | 76262471 | 76259768 | 76259922 | 76260194 | 272 | 76262471 | 14127 | 22,21 | 6,1 | 44,51 | 1,1 | 421 | 149 | 0.003994123 | 0.03495032 | 0.565,0.881 | 0.94,0.948 | -0.221 |
| 2527 | XLOC_007071 | chr11 | + | 47182677 | 47184358 | 47182677 | 47182741 | 47183014 | 273 | 47184358 | 2527 | 36,64 | 72,86 | 67,77 | 40,59 | 422 | 149 | 4.27E-05 | 0.001071261 | 0.15,0.208 | 0.372,0.315 | -0.165 |
| 9606 | XLOC_014896 | chr15 | - | 50881393 | 50881853 | 50881393 | 50881513 | 50881788 | 275 | 50881853 | 9606 | 0,2 | 2,0 | 3,5 | 0,0 | 424 | 149 | 0.003579337 | 0.032317159 | 0.0,1.0 | 1.0,1.0 | -0.5 |
| 9977 | XLOC_021623 | chr19 | - | 19414156 | 19414644 | 19414156 | 19414257 | 19414532 | 275 | 19414644 | 9977 | 27,31 | 89,86 | 56,75 | 61,53 | 424 | 149 | 4.31E-08 | 3.41E-06 | 0.096,0.112 | 0.244,0.332 | -0.184 |
| 2471 | XLOC_038275 | chr6 | - | 43014248 | 43014845 | 43014248 | 43014339 | 43014617 | 278 | 43014845 | 2471 | 64,45 | 37,38 | 88,70 | 22,12 | 427 | 149 | 1.19E-05 | 0.000377779 | 0.376,0.292 | 0.583,0.671 | -0.293 |
| 3086 | XLOC_018154 | chr17 | + | 48944121 | 48945644 | 48944121 | 48944578 | 48944856 | 278 | 48945644 | 3086 | 4,2 | 10,19 | 9,8 | 4,14 | 427 | 149 | 0.005836501 | 0.04645114 | 0.122,0.035 | 0.44,0.166 | -0.224 |
| 1320 | XLOC_018401 | chr17 | - | 7982758 | 7983259 | 7982758 | 7982842 | 7983122 | 280 | 7983259 | 1320 | 5,13 | 0,9 | 7,10 | 1,0 | 429 | 149 | 0.006319276 | 0.049118899 | 1.0,0.334 | 0.709,1.0 | -0.188 |
| 644 | XLOC_029298 | chr3 | + | 9471510 | 9475634 | 9471510 | 9475246 | 9475528 | 282 | 9475634 | 644 | 21,32 | 65,65 | 54,56 | 35,28 | 431 | 149 | 1.24E-09 | 1.62E-07 | 0.1,0.145 | 0.348,0.409 | -0.256 |
| 14376 | XLOC_038236 | chr6 | - | 34263014 | 34263462 | 34263014 | 34263132 | 34263417 | 285 | 34263462 | 14376 | 2,3 | 2,1 | 8,9 | 1,0 | 434 | 149 | 0.000872398 | 0.011130287 | 0.256,0.507 | 0.733,1.0 | -0.485 |
| 15746 | XLOC_047699 | chrX | + | 48380267 | 48380640 | 48380267 | 48380295 | 48380580 | 285 | 48380640 | 15746 | 10,10 | 7,4 | 25,27 | 2,4 | 434 | 149 | 0.003996114 | 0.03495032 | 0.329,0.462 | 0.811,0.699 | -0.359 |
| 3159 | XLOC_018763 | chr17 | - | 78187602 | 78188127 | 78187602 | 78187684 | 78187970 | 286 | 78188127 | 3159 | 22,50 | 125,118 | 87,112 | 71,84 | 435 | 149 | 4.52E-14 | 1.79E-11 | 0.057,0.127 | 0.296,0.314 | -0.213 |
| 15882 | XLOC_000712 | chr1 | + | 212529967 | 212530386 | 212529967 | 212530018 | 212530304 | 286 | 212530386 | 15882 | 8,8 | 2,5 | 14,13 | 0,2 | 435 | 149 | 0.005199306 | 0.042759957 | 0.578,0.354 | 1.0,0.69 | -0.379 |
| 14375 | XLOC_038236 | chr6 | - | 34263014 | 34263462 | 34263014 | 34263130 | 34263417 | 287 | 34263462 | 14375 | 2,3 | 4,0 | 8,9 | 1,0 | 436 | 149 | 0.003402387 | 0.030961105 | 0.146,1.0 | 1.0,1.0 | -0.427 |
| 9509 | XLOC_015039 | chr15 | - | 85070011 | 85070657 | 85070011 | 85070107 | 85070396 | 289 | 85070657 | 9509 | 40,25 | 5,0 | 54,48 | 0,0 | 438 | 149 | 0.002796739 | 0.026643043 | 0.731,1.0 | 1.0,1.0 | -0.135 |
| 13263 | XLOC_000995 | chr1 | - | 33098862 | 33099328 | 33098862 | 33098955 | 33099245 | 290 | 33099328 | 13263 | 3,3 | 1,1 | 4,6 | 1,0 | 439 | 149 | 0.003076095 | 0.02873952 | 0.505,0.505 | 0.576,1.0 | -0.283 |
| 3416 | XLOC_018359 | chr17 | - | 4872444 | 4872996 | 4872444 | 4872632 | 4872924 | 292 | 4872996 | 3416 | 18,15 | 41,60 | 48,34 | 45,51 | 441 | 149 | 0.000294767 | 0.004856164 | 0.129,0.078 | 0.265,0.184 | -0.121 |
| 11117 | XLOC_009338 | chr12 | - | 49233216 | 49233692 | 49233216 | 49233334 | 49233626 | 292 | 49233692 | 11117 | 0,0 | 3,1 | 1,6 | 10,3 | 441 | 149 | 0.001654911 | 0.018077827 | 0.0,0.0 | 0.033,0.403 | -0.218 |
| 3176 | XLOC_023503 | chr2 | + | 66795798 | 66796277 | 66795798 | 66795888 | 66796181 | 293 | 66796277 | 3176 | 0,1 | 2,1 | 12,2 | 5,3 | 442 | 149 | 0.001809922 | 0.0193084 | 0.0,0.252 | 0.447,0.183 | -0.189 |
| 11788 | XLOC_008873 | chr12 | + | 49908337 | 49908778 | 49908337 | 49908396 | 49908691 | 295 | 49908778 | 11788 | 1,0 | 0,3 | 3,4 | 0,1 | 444 | 149 | 0.000320848 | 0.005178319 | 1.0,0.0 | 1.0,0.573 | -0.286 |
| 13915 | XLOC_046078 | chr9 | + | 131389278 | 131389777 | 131389278 | 131389369 | 131389664 | 295 | 131389777 | 13915 | 0,1 | 2,0 | 3,2 | 0,0 | 444 | 149 | 0.000363764 | 0.005717595 | 0.0,1.0 | 1.0,1.0 | -0.5 |
| 9685 | XLOC_014777 | chr15 | + | 92705985 | 92706683 | 92705985 | 92706228 | 92706525 | 297 | 92706683 | 9685 | 50,37 | 3,9 | 42,31 | 1,0 | 446 | 149 | 0.000108986 | 0.002227409 | 0.848,0.579 | 0.933,1.0 | -0.253 |
| 11346 | XLOC_009470 | chr12 | - | 80180153 | 80180638 | 80180153 | 80180258 | 80180556 | 298 | 80180638 | 11346 | 0,3 | 1,1 | 2,3 | 0,1 | 447 | 149 | 0.00216054 | 0.022033115 | 0.0,0.5 | 1.0,0.5 | -0.5 |
| 3262 | XLOC_029673 | chr3 | + | 183860839 | 183861353 | 183860839 | 183860930 | 183861229 | 299 | 183861353 | 3262 | 64,85 | 157,153 | 209,217 | 189,145 | 448 | 149 | 3.64E-10 | 5.56E-08 | 0.119,0.156 | 0.269,0.332 | -0.163 |
| 4499 | XLOC_044293 | chr8 | - | 48846193 | 48846650 | 48846193 | 48846224 | 48846524 | 300 | 48846650 | 4499 | 4,11 | 2,8 | 11,9 | 0,0 | 449 | 149 | 2.78E-08 | 2.38E-06 | 0.399,0.313 | 1.0,1.0 | -0.644 |
| 3030 | XLOC_016775 | chr16 | - | 67863667 | 67864451 | 67863667 | 67863991 | 67864292 | 301 | 67864451 | 3030 | 251,197 | 42,44 | 267,247 | 24,28 | 450 | 149 | 0.002055827 | 0.021224263 | 0.664,0.597 | 0.786,0.745 | -0.135 |
| 6028 | XLOC_046152 | chr9 | + | 140108418 | 140108896 | 140108418 | 140108520 | 140108821 | 301 | 140108896 | 6028 | 19,38 | 5,1 | 38,34 | 0,0 | 450 | 149 | 0.000106267 | 0.002180722 | 0.557,0.926 | 1.0,1.0 | -0.258 |
| 11898 | XLOC_018295 | chr17 | + | 79986113 | 79986559 | 79986113 | 79986187 | 79986488 | 301 | 79986559 | 11898 | 36,41 | 4,1 | 27,35 | 0,0 | 450 | 149 | 0.002451821 | 0.024280349 | 0.749,0.931 | 1.0,1.0 | -0.16 |
| 10175 | XLOC_013336 | chr14 | - | 24910879 | 24911378 | 24910879 | 24911001 | 24911303 | 302 | 24911378 | 10175 | 30,47 | 14,19 | 41,36 | 3,1 | 451 | 149 | 1.79E-06 | 7.89E-05 | 0.415,0.45 | 0.819,0.922 | -0.438 |
| 2198 | XLOC_009410 | chr12 | - | 56716398 | 56716993 | 56716398 | 56716542 | 56716845 | 303 | 56716993 | 2198 | 238,186 | 78,59 | 313,281 | 68,49 | 452 | 149 | 0.00383442 | 0.033956493 | 0.501,0.51 | 0.603,0.654 | -0.123 |
| 865 | XLOC_018679 | chr17 | - | 62968871 | 62969635 | 62968871 | 62969045 | 62969349 | 304 | 62969635 | 865 | 12,23 | 9,2 | 9,10 | 1,0 | 453 | 149 | 0.001280803 | 0.014799361 | 0.305,0.791 | 0.747,1.0 | -0.325 |
| 1458 | XLOC_016862 | chr16 | - | 89836573 | 89837042 | 89836573 | 89836667 | 89836971 | 304 | 89837042 | 1458 | 110,103 | 101,96 | 271,282 | 90,87 | 453 | 149 | 8.16E-11 | 1.51E-08 | 0.264,0.261 | 0.498,0.516 | -0.244 |
| 12615 | XLOC_007213 | chr11 | + | 72533532 | 72534006 | 72533532 | 72533588 | 72533892 | 304 | 72534006 | 12615 | 33,39 | 19,14 | 78,77 | 9,11 | 453 | 149 | 0.000152613 | 0.002916902 | 0.364,0.478 | 0.74,0.697 | -0.297 |
| 11687 | XLOC_005390 | chr10 | - | 115612481 | 115614118 | 115612481 | 115613606 | 115613912 | 306 | 115614118 | 11687 | 9,14 | 2,3 | 12,13 | 0,0 | 455 | 149 | 8.77E-05 | 0.001874037 | 0.596,0.604 | 1.0,1.0 | -0.4 |
| 6246 | XLOC_020995 | chr19 | + | 13002306 | 13002788 | 13002306 | 13002336 | 13002644 | 308 | 13002788 | 6246 | 29,27 | 58,71 | 84,87 | 47,48 | 457 | 149 | 9.98E-11 | 1.74E-08 | 0.14,0.11 | 0.368,0.371 | -0.244 |
| 9058 | XLOC_016403 | chr16 | + | 53355074 | 53355542 | 53355074 | 53355129 | 53355437 | 308 | 53355542 | 9058 | 2,8 | 4,1 | 6,10 | 0,0 | 457 | 149 | 4.14E-06 | 0.000157355 | 0.14,0.723 | 1.0,1.0 | -0.569 |
| 132 | XLOC_044027 | chr8 | + | 48641494 | 48642027 | 48641494 | 48641664 | 48641973 | 309 | 48642027 | 132 | 25,28 | 5,8 | 13,21 | 0,0 | 458 | 149 | 1.10E-06 | 5.22E-05 | 0.619,0.532 | 1.0,1.0 | -0.424 |
| 5481 | XLOC_029378 | chr3 | + | 38538011 | 38538420 | 38538011 | 38538021 | 38538333 | 312 | 38538420 | 5481 | 3,8 | 1,3 | 1,2 | 0,0 | 461 | 149 | 9.76E-05 | 0.002029683 | 0.492,0.463 | 1.0,1.0 | -0.522 |
| 11996 | XLOC_032428 | chr4 | + | 119199924 | 119200586 | 119199924 | 119199947 | 119200261 | 314 | 119200586 | 11996 | 154,188 | 16,10 | 365,407 | 5,13 | 463 | 149 | 0.000419466 | 0.006360491 | 0.756,0.858 | 0.959,0.91 | -0.128 |
| 13566 | XLOC_021518 | chr19 | - | 8654335 | 8654875 | 8654335 | 8654458 | 8654772 | 314 | 8654875 | 13566 | 8,12 | 6,3 | 10,10 | 1,0 | 463 | 149 | 0.001306755 | 0.014993295 | 0.563,0.563 | 0.763,1.0 | -0.319 |
| 701 | XLOC_028537 | chr22 | + | 46495382 | 46509758 | 46495382 | 46505233 | 46505550 | 317 | 46509758 | 701 | 24,33 | 14,15 | 42,51 | 3,10 | 466 | 149 | 0.001303875 | 0.014971668 | 0.354,0.413 | 0.817,0.62 | -0.335 |
| 5697 | XLOC_028555 | chr22 | + | 51143391 | 51143881 | 51143391 | 51143524 | 51143842 | 318 | 51143881 | 5697 | 8,17 | 0,4 | 11,8 | 0,0 | 467 | 149 | 0.00521893 | 0.042851063 | 1.0,0.576 | 1.0,1.0 | -0.212 |
| 14923 | XLOC_018785 | chr17 | - | 79894623 | 79895123 | 79894623 | 79894737 | 79895055 | 318 | 79895123 | 14923 | 100,100 | 0,13 | 135,156 | 1,0 | 467 | 149 | 0.000119626 | 0.002405628 | 1.0,0.711 | 0.977,1.0 | -0.133 |
| 3249 | XLOC_040979 | chr7 | + | 889156 | 889670 | 889156 | 889240 | 889559 | 319 | 889670 | 3249 | 136,163 | 228,230 | 246,169 | 160,149 | 468 | 149 | 8.37E-07 | 4.11E-05 | 0.16,0.184 | 0.329,0.265 | -0.125 |
| 11684 | XLOC_028526 | chr22 | + | 44330485 | 44330964 | 44330485 | 44330546 | 44330865 | 319 | 44330964 | 11684 | 15,26 | 3,1 | 19,34 | 0,0 | 468 | 149 | 0.005053433 | 0.041834749 | 0.614,0.892 | 1.0,1.0 | -0.247 |
| 1344 | XLOC_018152 | chr17 | + | 48628358 | 48629040 | 48628358 | 48628551 | 48628871 | 320 | 48629040 | 1344 | 49,48 | 66,62 | 95,89 | 65,63 | 469 | 149 | 0.001355737 | 0.015449244 | 0.191,0.197 | 0.317,0.31 | -0.119 |
| 1952 | XLOC_044467 | chr8 | - | 145750199 | 145753171 | 145750199 | 145752023 | 145752343 | 320 | 145753171 | 1952 | 206,234 | 43,51 | 429,466 | 35,38 | 469 | 149 | 5.80E-07 | 3.05E-05 | 0.603,0.593 | 0.796,0.796 | -0.198 |

| | | | | | | | | | | | | | | | | | | | | | | |
|---|---|---|---|---|---|---|---|---|---|---|---|---|---|---|---|---|---|---|---|---|---|---|
| 2591 | XLOC_024206 | chr2 | - | 128628387 | 128628933 | 128628387 | 128628502 | 128628822 | 320 | 128628933 | 2591 | 10,9 | 75,88 | 33,53 | 96,67 | 469 | 149 | 7.24E-08 | 5.37E-06 | 0.041,0.031 | 0.098,0.201 | -0.114 |
| 5239 | XLOC_029869 | chr3 | - | 52029057 | 52029479 | 52029057 | 52029122 | 52029442 | 320 | 52029479 | 5239 | 9,4 | 5,1 | 6,8 | 0,1 | 469 | 149 | 0.00069128 | 0.009267594 | 0.364,0.56 | 1.0,0.718 | -0.397 |
| 2231 | XLOC_009247 | chr12 | - | 9573326 | 9574486 | 9573326 | 9574087 | 9574413 | 326 | 9574486 | 2231 | 190,118 | 61,38 | 303,318 | 67,54 | 475 | 149 | 0.006040391 | 0.047645284 | 0.494,0.493 | 0.587,0.649 | -0.124 |
| 3969 | XLOC_045897 | chr9 | + | 35606833 | 35607406 | 35606833 | 35606980 | 35607308 | 328 | 35607406 | 3969 | 10,18 | 4,0 | 3,16 | 0,0 | 477 | 149 | 0.005836483 | 0.04645114 | 0.438,1.0 | 1.0,1.0 | -0.281 |
| 2838 | XLOC_009549 | chr12 | - | 112277460 | 112278281 | 112277460 | 112277807 | 112278136 | 329 | 112278281 | 2838 | 246,208 | 43,35 | 433,401 | 22,18 | 478 | 149 | 5.40E-09 | 5.72E-07 | 0.641,0.649 | 0.86,0.874 | -0.222 |
| 1208 | XLOC_023637 | chr2 | + | 114398929 | 114399441 | 114398929 | 114399027 | 114399357 | 330 | 114399441 | 1208 | 2,0 | 3,7 | 2,7 | 2,4 | 479 | 149 | 0.002467022 | 0.024333733 | 0.172,0.0 | 0.237,0.352 | -0.208 |
| 3405 | XLOC_021451 | chr19 | - | 1609289 | 1611848 | 1609289 | 1611368 | 1611702 | 334 | 1611848 | 3405 | 486,609 | 101,98 | 900,838 | 96,47 | 483 | 149 | 3.45E-06 | 0.000133758 | 0.597,0.657 | 0.743,0.846 | -0.167 |
| 6812 | XLOC_007584 | chr11 | - | 65427209 | 65427595 | 65427209 | 65427259 | 65427594 | 335 | 65427595 | 6812 | 48,38 | 6,0 | 35,68 | 0,0 | 484 | 149 | 0.000675314 | 0.009126844 | 0.711,1.0 | 1.0,1.0 | -0.145 |
| 3374 | XLOC_017907 | chr17 | + | 7358602 | 7359260 | 7358602 | 7358775 | 7359112 | 337 | 7359260 | 3374 | 17,24 | 6,6 | 16,37 | 0,1 | 486 | 149 | 3.10E-06 | 0.000123622 | 0.465,0.551 | 1.0,0.919 | -0.452 |
| 11526 | XLOC_023782 | chr2 | + | 197584897 | 197585392 | 197584897 | 197584958 | 197585298 | 340 | 197585392 | 11526 | 0,1 | 1,2 | 5,10 | 5,7 | 489 | 149 | 0.002007425 | 0.020930873 | 0.0,0.132 | 0.234,0.303 | -0.203 |
| 2894 | XLOC_016217 | chr16 | + | 2287767 | 2289507 | 2287767 | 2288019 | 2288362 | 343 | 2289507 | 2894 | 3,4 | 16,11 | 9,17 | 5,8 | 492 | 149 | 2.87E-05 | 0.000782193 | 0.054,0.099 | 0.353,0.392 | -0.296 |
| 3333 | XLOC_000851 | chr1 | - | 3547163 | 3548221 | 3547163 | 3547681 | 3548029 | 348 | 3548221 | 3333 | 37,21 | 29,24 | 76,48 | 24,23 | 497 | 149 | 0.001973841 | 0.020729023 | 0.277,0.208 | 0.487,0.385 | -0.194 |
| 8515 | XLOC_017884 | chr17 | + | 5284683 | 5285164 | 5284683 | 5284800 | 5285148 | 348 | 5285164 | 8515 | 24,21 | 3,2 | 29,29 | 1,0 | 497 | 149 | 0.002011847 | 0.020946998 | 0.706,0.759 | 0.897,1.0 | -0.216 |
| 2179 | XLOC_007058 | chr11 | + | 43904132 | 43904738 | 43904132 | 43904281 | 43904634 | 353 | 43904738 | 2179 | 5,2 | 29,27 | 34,23 | 48,63 | 502 | 149 | 8.57E-05 | 0.001836694 | 0.049,0.022 | 0.174,0.098 | -0.101 |
| 694 | XLOC_028592 | chr22 | - | 22313503 | 22314108 | 22313503 | 22313604 | 22313958 | 354 | 22314108 | 694 | 5,14 | 36,30 | 28,59 | 29,24 | 503 | 149 | 4.68E-06 | 0.000175446 | 0.04,0.121 | 0.222,0.421 | -0.241 |
| 2803 | XLOC_005184 | chr10 | - | 31605341 | 31607978 | 31605341 | 31607469 | 31607824 | 355 | 31607978 | 2803 | 93,100 | 21,18 | 121,151 | 8,14 | 504 | 149 | 0.001422951 | 0.016020976 | 0.567,0.622 | 0.817,0.761 | -0.194 |
| 4092 | XLOC_046494 | chr9 | - | 140006141 | 140006621 | 140006141 | 140006206 | 140006563 | 357 | 140006621 | 4092 | 70,71 | 17,35 | 69,83 | 6,1 | 506 | 149 | 3.13E-06 | 0.000123873 | 0.548,0.374 | 0.772,0.961 | -0.406 |
| 2551 | XLOC_007119 | chr11 | + | 62549584 | 62550297 | 62549584 | 62549805 | 62550164 | 359 | 62550297 | 2551 | 35,37 | 68,74 | 106,75 | 48,39 | 508 | 149 | 3.42E-11 | 7.25E-09 | 0.131,0.128 | 0.393,0.361 | -0.247 |
| 1993 | XLOC_000836 | chr1 | - | 1601102 | 1601590 | 1601102 | 1601175 | 1601536 | 361 | 1601590 | 1993 | 92,67 | 35,35 | 268,281 | 18,22 | 510 | 149 | 2.56E-14 | 1.10E-11 | 0.434,0.359 | 0.813,0.789 | -0.404 |
| 603 | XLOC_029564 | chr3 | + | 128207960 | 128208912 | 128207960 | 128208258 | 128208621 | 363 | 128208912 | 603 | 61,54 | 38,53 | 101,102 | 41,29 | 512 | 149 | 0.000494998 | 0.007188159 | 0.318,0.229 | 0.418,0.506 | -0.188 |
| 1106 | XLOC_021261 | chr19 | + | 45571272 | 45571738 | 45571272 | 45571312 | 45571677 | 365 | 45571738 | 1106 | 121,121 | 99,91 | 206,217 | 85,75 | 514 | 149 | 1.08E-05 | 0.000349737 | 0.262,0.278 | 0.413,0.456 | -0.164 |
| 10180 | XLOC_013336 | chr14 | - | 24911551 | 24912031 | 24911551 | 24911658 | 24912023 | 365 | 24912031 | 10180 | 103,90 | 4,4 | 97,107 | 0,1 | 514 | 149 | 0.001174677 | 0.013836716 | 0.882,0.867 | 1.0,0.969 | -0.11 |
| 102 | XLOC_044433 | chr8 | - | 144362237 | 144363167 | 144362237 | 144362685 | 144363051 | 366 | 144363167 | 102 | 107,113 | 31,20 | 109,110 | 7,5 | 515 | 149 | 1.25E-05 | 0.000396829 | 0.5,0.62 | 0.818,0.864 | -0.281 |
| 2582 | XLOC_006946 | chr11 | + | 581491 | 582081 | 581491 | 581597 | 581964 | 367 | 582081 | 2582 | 18,5 | 6,0 | 27,18 | 0,1 | 516 | 149 | 0.003800596 | 0.03378757 | 0.464,1.0 | 1.0,0.839 | -0.188 |
| 3381 | XLOC_001222 | chr1 | - | 109823729 | 109824682 | 109823729 | 109823873 | 109824240 | 367 | 109824682 | 3381 | 87,67 | 209,216 | 143,125 | 127,124 | 516 | 149 | 1.98E-11 | 4.44E-09 | 0.107,0.082 | 0.245,0.225 | -0.14 |
| 501 | XLOC_046480 | chr9 | - | 139326275 | 139327527 | 139326275 | 139327038 | 139327407 | 369 | 139327527 | 501 | 24,13 | 26,13 | 38,17 | 16,4 | 518 | 149 | 0.004153821 | 0.036033318 | 0.21,0.223 | 0.406,0.55 | -0.262 |
| 2589 | XLOC_000006 | chr1 | + | 982199 | 982834 | 982199 | 982337 | 982706 | 369 | 982834 | 2589 | 85,88 | 302,291 | 254,297 | 254,282 | 518 | 149 | 0 | 0 | 0.075,0.08 | 0.223,0.233 | -0.151 |
| 1690 | XLOC_014869 | chr15 | - | 43698141 | 43700286 | 43698141 | 43699768 | 43700140 | 372 | 43700286 | 1690 | 273,260 | 58,51 | 382,408 | 43,52 | 521 | 149 | 0.003177959 | 0.029435258 | 0.574,0.593 | 0.718,0.692 | -0.122 |
| 1898 | XLOC_013183 | chr14 | + | 73577538 | 73578510 | 73577538 | 73577863 | 73578235 | 372 | 73578510 | 1898 | 280,378 | 343,320 | 419,326 | 208,213 | 521 | 149 | 7.35E-06 | 0.000253332 | 0.189,0.253 | 0.366,0.304 | -0.114 |
| 11793 | XLOC_000548 | chr1 | + | 155295205 | 155295701 | 155295205 | 155295281 | 155295654 | 373 | 155295701 | 11793 | 23,17 | 5,1 | 26,7 | 0,1 | 522 | 149 | 0.003473505 | 0.031456029 | 0.568,0.829 | 1.0,0.666 | -0.135 |
| 996 | XLOC_026793 | chr20 | + | 42158923 | 42159674 | 42158923 | 42159059 | 42159437 | 378 | 42159674 | 996 | 20,13 | 2,2 | 15,14 | 0,0 | 527 | 149 | 0.001559373 | 0.01720916 | 0.739,0.648 | 1.0,1.0 | -0.306 |
| 2671 | XLOC_001160 | chr1 | - | 78412214 | 78414985 | 78412214 | 78414459 | 78414839 | 380 | 78414985 | 2671 | 672,575 | 833,725 | 928,849 | 529,549 | 529 | 149 | 2.11E-10 | 3.41E-08 | 0.185,0.183 | 0.331,0.303 | -0.133 |
| 1706 | XLOC_014811 | chr15 | - | 28459221 | 28459948 | 28459221 | 28459418 | 28459800 | 382 | 28459948 | 1706 | 9,2 | 64,25 | 18,34 | 38,50 | 531 | 149 | 2.32E-06 | 9.81E-05 | 0.038,0.022 | 0.117,0.16 | -0.109 |
| 2693 | XLOC_007775 | chr11 | - | 134122733 | 134123314 | 134122733 | 134122810 | 134123192 | 382 | 134123314 | 2693 | 47,58 | 30,21 | 67,39 | 9,8 | 531 | 149 | 0.001380774 | 0.015641338 | 0.305,0.437 | 0.676,0.578 | -0.256 |
| 1482 | XLOC_016684 | chr16 | - | 28890054 | 28891076 | 28890054 | 28890425 | 28890809 | 384 | 28891076 | 1482 | 26,13 | 3,5 | 10,20 | 0,1 | 533 | 149 | 0.000550044 | 0.007786903 | 0.708,0.421 | 1.0,0.848 | -0.359 |
| 128 | XLOC_044112 | chr8 | + | 101225317 | 101226146 | 101225317 | 101225656 | 101226046 | 390 | 101226146 | 128 | 0,0 | 13,13 | 6,2 | 19,4 | 539 | 149 | 0.00149377 | 0.016607017 | 0.0,0.0 | 0.08,0.121 | -0.101 |
| 15728 | XLOC_009117 | chr12 | + | 113725967 | 113726652 | 113725967 | 113726039 | 113726430 | 391 | 113726652 | 15728 | 56,50 | 5,0 | 66,53 | 0,0 | 540 | 149 | 0.005024864 | 0.041713027 | 0.756,1.0 | 1.0,1.0 | -0.122 |
| 1449 | XLOC_008841 | chr12 | + | 31242336 | 31243028 | 31242336 | 31242424 | 31242819 | 395 | 31243028 | 1449 | 198,165 | 117,155 | 259,215 | 113,90 | 544 | 149 | 0.000978165 | 0.012070188 | 0.317,0.226 | 0.386,0.396 | -0.119 |
| 2740 | XLOC_005357 | chr10 | - | 103344358 | 103345235 | 103344358 | 103344676 | 103345072 | 396 | 103345235 | 2740 | 39,33 | 60,54 | 56,62 | 45,34 | 545 | 149 | 0.000223377 | 0.003939084 | 0.151,0.143 | 0.254,0.333 | -0.146 |
| 1785 | XLOC_000030 | chr1 | + | 3394438 | 3395176 | 3394438 | 3394590 | 3394987 | 397 | 3395176 | 1785 | 59,64 | 150,121 | 127,125 | 116,104 | 546 | 149 | 7.31E-08 | 5.39E-06 | 0.097,0.126 | 0.23,0.247 | -0.127 |
| 6963 | XLOC_016235 | chr16 | + | 2818505 | 2819285 | 2818505 | 2818600 | 2818997 | 397 | 2819285 | 6963 | 355,308 | 59,82 | 650,487 | 38,32 | 546 | 149 | 3.44E-09 | 3.91E-07 | 0.621,0.506 | 0.824,0.806 | -0.251 |
| 2950 | XLOC_028389 | chr22 | + | 20127242 | 20128231 | 20127242 | 20127740 | 20128139 | 399 | 20128231 | 2950 | 41,53 | 30,33 | 120,98 | 36,41 | 548 | 149 | 0.0064655 | 0.04982277 | 0.271,0.304 | 0.475,0.394 | -0.147 |
| 228 | XLOC_041046 | chr7 | + | 27138692 | 27140058 | 27138692 | 27138985 | 27139397 | 412 | 27140058 | 228 | 22,21 | 33,26 | 42,50 | 17,18 | 561 | 149 | 8.89E-06 | 0.000295782 | 0.15,0.177 | 0.396,0.425 | -0.247 |
| 2195 | XLOC_009411 | chr12 | - | 56748561 | 56749135 | 56748561 | 56748647 | 56749059 | 412 | 56749135 | 2195 | 58,62 | 46,37 | 130,107 | 45,30 | 561 | 149 | 0.000276418 | 0.004614737 | 0.251,0.308 | 0.434,0.486 | -0.18 |
| 125 | XLOC_044167 | chr8 | + | 144462234 | 144462953 | 144462234 | 144462345 | 144462758 | 413 | 144462953 | 125 | 234,215 | 38,25 | 215,252 | 12,8 | 562 | 149 | 8.58E-06 | 0.000288207 | 0.62,0.695 | 0.826,0.893 | -0.202 |
| 7557 | XLOC_024366 | chr2 | - | 201778017 | 201778715 | 201778017 | 201778136 | 201778550 | 414 | 201778715 | 7557 | 14,38 | 6,0 | 78,87 | 0,0 | 563 | 149 | 0.001080935 | 0.013007544 | 0.382,1.0 | 1.0,1.0 | -0.309 |
| 1119 | XLOC_044280 | chr8 | - | 38125888 | 38126508 | 38125888 | 38125979 | 38126399 | 420 | 38126508 | 1119 | 37,61 | 64,54 | 64,57 | 46,26 | 569 | 149 | 0.00404822 | 0.035341452 | 0.131,0.228 | 0.267,0.365 | -0.137 |
| 2602 | XLOC_007095 | chr11 | + | 60702723 | 60704627 | 60702723 | 60702915 | 60703335 | 420 | 60704627 | 2602 | 74,89 | 150,133 | 98,76 | 83,61 | 569 | 149 | 3.69E-05 | 0.000954575 | 0.114,0.149 | 0.236,0.246 | -0.109 |
| 15850 | XLOC_000295 | chr1 | + | 52221991 | 52222487 | 52221991 | 52222030 | 52222452 | 422 | 52222487 | 15850 | 8,12 | 7,2 | 10,21 | 1,0 | 571 | 149 | 9.83E-05 | 0.002042848 | 0.23,0.61 | 0.723,1.0 | -0.441 |
| 12307 | XLOC_000147 | chr1 | + | 27087346 | 27087964 | 27087346 | 27087446 | 27087874 | 428 | 27087964 | 12307 | 15,12 | 4,1 | 7,18 | 0,0 | 577 | 149 | 2.93E-05 | 0.000794017 | 0.492,0.756 | 1.0,1.0 | -0.376 |
| 534 | XLOC_037824 | chr6 | + | 41753865 | 41755563 | 41753865 | 41754070 | 41754499 | 429 | 41755563 | 534 | 62,43 | 33,23 | 90,108 | 10,17 | 578 | 149 | 6.17E-07 | 3.18E-05 | 0.326,0.325 | 0.699,0.621 | -0.334 |
| 2829 | XLOC_005019 | chr10 | + | 103368591 | 103369918 | 103368591 | 103368694 | 103369123 | 429 | 103369918 | 2829 | 12,25 | 85,111 | 50,80 | 63,69 | 578 | 149 | 2.89E-13 | 9.88E-11 | 0.035,0.055 | 0.17,0.23 | -0.155 |
| 466 | XLOC_035162 | chr5 | - | 176958282 | 176959201 | 176958282 | 176958522 | 176958953 | 431 | 176959201 | 466 | 25,46 | 24,37 | 60,45 | 27,15 | 580 | 149 | 0.004170182 | 0.036133566 | 0.211,0.242 | 0.363,0.435 | -0.173 |
| 2181 | XLOC_009489 | chr12 | - | 95451506 | 95452265 | 95451506 | 95451654 | 95452085 | 431 | 95452265 | 2181 | 163,164 | 53,53 | 292,276 | 49,59 | 580 | 149 | 0.003362811 | 0.03073252 | 0.441,0.443 | 0.605,0.546 | -0.134 |
| 6291 | XLOC_016187 | chr16 | + | 708645 | 709126 | 708645 | 708653 | 709085 | 432 | 709126 | 6291 | 82,53 | 7,0 | 81,89 | 0,0 | 581 | 149 | 0.000331616 | 0.005317871 | 0.75,1.0 | 1.0,1.0 | -0.125 |
| 2512 | XLOC_007359 | chr11 | + | 134126441 | 134127151 | 134126441 | 134126542 | 134126981 | 439 | 134127151 | 2512 | 8,8 | 47,45 | 24,23 | 15,22 | 588 | 149 | 5.35E-09 | 5.71E-07 | 0.041,0.043 | 0.288,0.209 | -0.207 |
| 2261 | XLOC_014987 | chr15 | - | 75197004 | 75197572 | 75197004 | 75197053 | 75197494 | 441 | 75197572 | 2261 | 48,56 | 82,80 | 88,67 | 57,50 | 590 | 149 | 5.62E-05 | 0.001317054 | 0.129,0.15 | 0.281,0.253 | -0.128 |
| 8423 | XLOC_017964 | chr17 | + | 19283094 | 19283814 | 19283094 | 19283260 | 19283707 | 447 | 19283814 | 8423 | 14,13 | 0,4 | 8,11 | 0,0 | 596 | 149 | 0.004361866 | 0.037556488 | 1.0,0.448 | 1.0,1.0 | -0.276 |
| 2371 | XLOC_026876 | chr20 | + | 62319880 | 62320485 | 62319880 | 62319958 | 62320407 | 449 | 62320485 | 2371 | 33,22 | 58,90 | 79,110 | 61,101 | 598 | 149 | 4.50E-08 | 3.51E-06 | 0.124,0.057 | 0.244,0.213 | -0.138 |
| 5621 | XLOC_028680 | chr22 | - | 39136271 | 39137055 | 39136271 | 39136437 | 39136893 | 456 | 39137055 | 5621 | 17,11 | 0,6 | 35,23 | 0,0 | 605 | 149 | 0.00102029 | 0.012497718 | 1.0,0.311 | 1.0,1.0 | -0.345 |
| 5409 | XLOC_029564 | chr3 | + | 128208621 | 128209567 | 128208621 | 128208912 | 128209369 | 457 | 128209567 | 5409 | 20,16 | 3,9 | 20,19 | 0,0 | 606 | 149 | 2.17E-07 | 1.35E-05 | 0.621,0.304 | 1.0,1.0 | -0.537 |
| 10120 | XLOC_013470 | chr14 | - | 77270128 | 77270758 | 77270128 | 77270255 | 77270717 | 462 | 77270758 | 10120 | 5,6 | 2,2 | 8,19 | 1,0 | 611 | 149 | 0.000317932 | 0.005155618 | 0.379,0.422 | 0.661,1.0 | -0.43 |
| 940 | XLOC_023708 | chr2 | + | 170550968 | 170551757 | 170550968 | 170551251 | 170551724 | 473 | 170551757 | 940 | 3,5 | 4,7 | 19,7 | 4,2 | 622 | 149 | 0.001873827 | 0.019849366 | 0.152,0.146 | 0.532,0.456 | -0.345 |
| 492 | XLOC_032538 | chr4 | - | 1719242 | 1720353 | 1719242 | 1719402 | 1719878 | 476 | 1720353 | 492 | 82,64 | 64,46 | 89,67 | 26,15 | 625 | 149 | 6.06E-06 | 0.00021695 | 0.234,0.249 | 0.449,0.516 | -0.241 |
| 103 | XLOC_044433 | chr8 | - | 144363051 | 144363914 | 144363051 | 144363167 | 144363651 | 484 | 144363914 | 103 | 92,90 | 59,70 | 151,157 | 28,28 | 633 | 149 | 2.56E-11 | 5.58E-09 | 0.268,0.232 | 0.559,0.569 | -0.314 |
| 13736 | XLOC_014788 | chr15 | + | 99500289 | 99507746 | 99500289 | 99500705 | 99501191 | 486 | 99507746 | 13736 | 81,89 | 10,15 | 119,91 | 5,4 | 635 | 149 | 0.000975772 | 0.012050538 | 0.655,0.582 | 0.848,0.842 | -0.226 |
| 2024 | XLOC_047884 | chrX | + | 153689438 | 153690688 | 153689438 | 153689978 | 153690467 | 489 | 153690688 | 2024 | 47,69 | 31,32 | 79,80 | 25,15 | 638 | 149 | 0.005811901 | 0.046328891 | 0.261,0.335 | 0.425,0.555 | -0.192 |
| 6591 | XLOC_023924 | chr2 | - | 38828 | 42305 | 38828 | 41627 | 42117 | 490 | 42305 | 6591 | 5,4 | 3,2 | 1,9 | 0,0 | 639 | 149 | 9.32E-07 | 4.52E-05 | 0.28,0.318 | 1.0,1.0 | -0.701 |
| 7636 | XLOC_020063 | chr18 | + | 21406665 | 21407431 | 21406665 | 21406781 | 21407277 | 496 | 21407431 | 7636 | 17,18 | 3,6 | 25,34 | 0,0 | 645 | 149 | 5.33E-08 | 4.09E-06 | 0.567,0.409 | 1.0,1.0 | -0.512 |
| 3079 | XLOC_001411 | chr1 | - | 169820232 | 169827318 | 169820232 | 169822913 | 169823410 | 497 | 169827318 | 3079 | 51,33 | 41,24 | 103,74 | 31,36 | 646 | 149 | 0.005963101 | 0.047184097 | 0.223,0.241 | 0.434,0.322 | -0.146 |
| 3186 | XLOC_007240 | chr11 | + | 82783664 | 82808540 | 82783664 | 82783846 | 82784344 | 498 | 82808540 | 3186 | 70,72 | 95,101 | 101,68 | 53,65 | 647 | 149 | 0.001863462 | 0.019767415 | 0.145,0.141 | 0.305,0.194 | -0.107 |

| | | | | | | | | | | | | | | | | | | | | | | |
|---|---|---|---|---|---|---|---|---|---|---|---|---|---|---|---|---|---|---|---|---|---|---|
| 1968 | XLOC_029502 | chr3 | + | 100064428 | 100065099 | 100064428 | 100064522 | 100065024 | 502 | 100065099 | 1968 | 123,93 | 147,160 | 302,252 | 171,185 | 651 | 149 | 8.91E-08 | 6.38E-06 | 0.161,0.117 | 0.288,0.238 | -0.124 |
| 7 | XLOC_048071 | chrX | - | 122756955 | 122757822 | 122756955 | 122757134 | 122757637 | 503 | 122757822 | 7 | 227,233 | 164,183 | 302,292 | 117,140 | 652 | 149 | 0.000101474 | 0.002099556 | 0.24,0.225 | 0.371,0.323 | -0.114 |
| 13669 | XLOC_041095 | chr7 | + | 44796023 | 44796748 | 44796023 | 44796138 | 44796641 | 503 | 44796748 | 13669 | 43,51 | 9,20 | 53,48 | 1,4 | 652 | 149 | 0.000179373 | 0.003322824 | 0.522,0.368 | 0.924,0.733 | -0.384 |
| 3312 | XLOC_027761 | chr21 | - | 34127137 | 34132305 | 34127137 | 34131580 | 34132087 | 507 | 34132305 | 3312 | 67,59 | 68,45 | 206,181 | 32,40 | 656 | 149 | 2.08E-13 | 7.39E-11 | 0.183,0.229 | 0.594,0.507 | -0.344 |
| 1199 | XLOC_020063 | chr18 | + | 21406665 | 21407431 | 21406665 | 21406763 | 21407277 | 514 | 21407431 | 1199 | 57,70 | 42,36 | 104,104 | 29,29 | 663 | 149 | 0.000529242 | 0.007560163 | 0.234,0.304 | 0.446,0.446 | -0.177 |
| 297 | XLOC_038044 | chr6 | + | 157800198 | 157809091 | 157800198 | 157803298 | 157803814 | 516 | 157809091 | 297 | 44,70 | 5,0 | 74,41 | 0,0 | 665 | 149 | 0.003382954 | 0.030877668 | 0.663,1.0 | 1.0,1.0 | -0.168 |
| 1175 | XLOC_024366 | chr2 | - | 201778017 | 201778715 | 201778017 | 201778136 | 201778653 | 517 | 201778715 | 1175 | 58,92 | 86,63 | 257,248 | 50,58 | 666 | 149 | 3.16E-13 | 1.04E-10 | 0.131,0.246 | 0.535,0.489 | -0.324 |
| 6067 | XLOC_026782 | chr20 | + | 37075330 | 37076266 | 37075330 | 37075591 | 37076108 | 517 | 37076266 | 6067 | 30,21 | 3,2 | 44,34 | 0,0 | 666 | 149 | 0.000194958 | 0.003547609 | 0.691,0.701 | 1.0,1.0 | -0.304 |
| 3202 | XLOC_032226 | chr4 | + | 1243043 | 1246399 | 1243043 | 1244161 | 1244683 | 522 | 1246399 | 3202 | 102,124 | 26,21 | 256,249 | 12,8 | 671 | 149 | 1.14E-08 | 1.06E-06 | 0.466,0.567 | 0.826,0.874 | -0.334 |
| 3329 | XLOC_000753 | chr1 | + | 226032198 | 226033232 | 226032198 | 226032324 | 226032846 | 522 | 226033232 | 3329 | 153,171 | 200,188 | 429,382 | 181,190 | 671 | 149 | 4.60E-12 | 1.17E-09 | 0.145,0.168 | 0.345,0.309 | -0.17 |
| 3316 | XLOC_009117 | chr12 | + | 113725967 | 113726652 | 113725967 | 113726039 | 113726562 | 523 | 113726652 | 3316 | 197,141 | 108,100 | 277,227 | 62,94 | 672 | 149 | 0.001450047 | 0.016265177 | 0.288,0.238 | 0.498,0.349 | -0.16 |
| 5827 | XLOC_027790 | chr21 | - | 38444733 | 38445459 | 38444733 | 38444837 | 38445360 | 523 | 38445459 | 5827 | 42,33 | 6,11 | 48,25 | 0,1 | 672 | 149 | 2.85E-06 | 0.000116072 | 0.608,0.399 | 1.0,0.847 | -0.42 |
| 4454 | XLOC_041046 | chr7 | + | 27138692 | 27140058 | 27138692 | 27138858 | 27139397 | 539 | 27140058 | 4454 | 14,13 | 4,4 | 34,33 | 0,0 | 688 | 149 | 4.07E-08 | 3.29E-06 | 0.431,0.413 | 1.0,1.0 | -0.578 |
| 2260 | XLOC_014987 | chr15 | - | 75190467 | 75195124 | 75190467 | 75194404 | 75194944 | 540 | 75195124 | 2260 | 319,293 | 18,39 | 369,404 | 15,15 | 689 | 149 | 0.005072071 | 0.041919834 | 0.793,0.619 | 0.842,0.853 | -0.141 |
| 176 | XLOC_041435 | chr7 | - | 27168126 | 27181704 | 27168126 | 27169190 | 27169736 | 546 | 27181704 | 176 | 0,0 | 6,3 | 2,7 | 2,3 | 695 | 149 | 4.23E-05 | 0.001062518 | 0.0,0.0 | 0.177,0.333 | -0.255 |
| 63 | XLOC_046315 | chr9 | - | 95060538 | 95061236 | 95060538 | 95060618 | 95061165 | 547 | 95061236 | 63 | 38,50 | 135,147 | 175,200 | 184,228 | 696 | 149 | 1.52E-10 | 2.52E-08 | 0.057,0.068 | 0.169,0.158 | -0.101 |
| 4 | XLOC_048096 | chrX | - | 134184969 | 134186258 | 134184969 | 134185398 | 134185946 | 548 | 134186258 | 4 | 596,645 | 422,374 | 10,911,051 | 201,184 | 697 | 149 | 0 | 0 | 0.232,0.269 | 0.537,0.55 | -0.293 |
| 8330 | XLOC_018106 | chr17 | + | 43239251 | 43240002 | 43239251 | 43239306 | 43239854 | 548 | 43240002 | 8330 | 28,14 | 2,3 | 21,16 | 0,0 | 697 | 149 | 5.36E-05 | 0.001270009 | 0.75,0.499 | 1.0,1.0 | -0.375 |
| 385 | XLOC_034900 | chr5 | - | 54518003 | 54518851 | 54518003 | 54518227 | 54518778 | 551 | 54518851 | 385 | 15,20 | 15,11 | 47,48 | 7,14 | 700 | 149 | 0.001978217 | 0.020750582 | 0.176,0.279 | 0.588,0.422 | -0.277 |
| 9264 | XLOC_016339 | chr16 | + | 28847253 | 28848525 | 28847253 | 28847497 | 28848048 | 551 | 28848525 | 9264 | 369,327 | 27,15 | 333,380 | 5,5 | 700 | 149 | 6.71E-06 | 0.000235865 | 0.744,0.823 | 0.934,0.942 | -0.154 |
| 2420 | XLOC_001011 | chr1 | - | 35496147 | 35497208 | 35496147 | 35496314 | 35496867 | 553 | 35497208 | 2420 | 13,34 | 2,2 | 20,21 | 0,0 | 702 | 149 | 0.002142595 | 0.021894641 | 0.58,0.783 | 1.0,1.0 | -0.319 |
| 9061 | XLOC_021326 | chr19 | + | 50381754 | 50382717 | 50381754 | 50381829 | 50382382 | 553 | 50382717 | 9061 | 47,50 | 3,2 | 27,14 | 0,0 | 702 | 149 | 0.001960661 | 0.020638389 | 0.769,0.841 | 1.0,1.0 | -0.195 |
| 1570 | XLOC_016713 | chr16 | - | 31504298 | 31505108 | 31504298 | 31504369 | 31504923 | 554 | 31505108 | 1570 | 177,163 | 7,2 | 215,159 | 0,0 | 703 | 149 | 0.000129488 | 0.002549425 | 0.843,0.945 | 1.0,1.0 | -0.106 |
| 3167 | XLOC_017914 | chr17 | + | 7761923 | 7768020 | 7761923 | 7762142 | 7762699 | 557 | 7768020 | 3167 | 61,78 | 84,91 | 102,124 | 65,39 | 706 | 149 | 3.16E-05 | 0.000838972 | 0.133,0.153 | 0.249,0.402 | -0.182 |
| 1771 | XLOC_000505 | chr1 | + | 150965676 | 150969350 | 150965676 | 150967175 | 150967735 | 560 | 150969350 | 1771 | 50,31 | 16,20 | 68,45 | 5,0 | 709 | 149 | 3.26E-06 | 0.000127805 | 0.396,0.246 | 0.741,1.0 | -0.55 |
| 11858 | XLOC_015047 | chr15 | - | 89876326 | 89878071 | 89876326 | 89877144 | 89877705 | 561 | 89878071 | 11858 | 46,29 | 3,5 | 52,48 | 0,0 | 710 | 149 | 3.22E-06 | 0.000126606 | 0.763,0.549 | 1.0,1.0 | -0.344 |
| 2757 | XLOC_014776 | chr15 | + | 91504254 | 91506740 | 91504254 | 91504284 | 91504847 | 563 | 91506740 | 2757 | 686,724 | 351,322 | 1,044,938 | 327,242 | 712 | 149 | 1.87E-05 | 0.000541046 | 0.29,0.32 | 0.401,0.448 | -0.119 |
| 6048 | XLOC_026806 | chr20 | + | 43629032 | 43629994 | 43629032 | 43629227 | 43629807 | 580 | 43629994 | 6048 | 24,17 | 8,2 | 30,29 | 2,0 | 729 | 149 | 0.001462601 | 0.016357206 | 0.38,0.635 | 0.754,1.0 | -0.369 |
| 2125 | XLOC_028670 | chr22 | - | 38694099 | 38694939 | 38694099 | 38694209 | 38694790 | 581 | 38694939 | 2125 | 13,171,464 | 133,159 | 13,081,277 | 30,44 | 730 | 149 | 2.22E-16 | 1.34E-13 | 0.669,0.653 | 0.899,0.856 | -0.216 |
| 2139 | XLOC_018266 | chr17 | + | 77079842 | 77080768 | 77079842 | 77080037 | 77080619 | 582 | 77080768 | 2139 | 20,28 | 10,20 | 80,55 | 18,15 | 731 | 149 | 0.002598362 | 0.025264741 | 0.29,0.222 | 0.475,0.428 | -0.196 |
| 2940 | XLOC_000699 | chr1 | + | 207940357 | 207941168 | 207940357 | 207940540 | 207941123 | 583 | 207941168 | 2940 | 12,181,293 | 279,363 | 28,362,421 | 313,282 | 732 | 149 | 6.08E-12 | 1.52E-09 | 0.471,0.42 | 0.648,0.636 | -0.197 |
| 2138 | XLOC_018266 | chr17 | + | 77079842 | 77080768 | 77079842 | 77080033 | 77080619 | 586 | 77080768 | 2138 | 20,28 | 61,53 | 81,55 | 61,54 | 735 | 149 | 5.92E-06 | 0.000213692 | 0.062,0.097 | 0.212,0.171 | -0.112 |
| 15292 | XLOC_001334 | chr1 | - | 153905958 | 153906778 | 153905958 | 153906023 | 153906610 | 587 | 153906778 | 15292 | 109,88 | 8,0 | 154,136 | 1,0 | 736 | 149 | 0.000792704 | 0.010305837 | 0.734,1.0 | 0.969,1.0 | -0.117 |
| 498 | XLOC_046480 | chr9 | - | 139324728 | 139325569 | 139324728 | 139324865 | 139325453 | 588 | 139325569 | 498 | 15,15 | 67,55 | 54,40 | 46,52 | 737 | 149 | 9.15E-08 | 6.48E-06 | 0.043,0.052 | 0.192,0.135 | -0.116 |
| 9230 | XLOC_016177 | chr16 | + | 456714 | 457560 | 456714 | 456833 | 457424 | 591 | 457560 | 9230 | 10,25 | 1,6 | 24,26 | 0,0 | 740 | 149 | 5.45E-07 | 2.91E-05 | 0.668,0.456 | 1.0,1.0 | -0.438 |
| 133 | XLOC_044017 | chr8 | + | 42176069 | 42176939 | 42176069 | 42176193 | 42176787 | 594 | 42176939 | 133 | 42,34 | 93,49 | 115,134 | 71,51 | 743 | 149 | 1.06E-09 | 1.48E-07 | 0.083,0.122 | 0.245,0.345 | -0.192 |
| 2853 | XLOC_020911 | chr19 | + | 4501202 | 4502209 | 4501202 | 4501329 | 4501925 | 596 | 4502209 | 2853 | 50,62 | 42,48 | 93,100 | 36,49 | 745 | 149 | 0.004646214 | 0.039440383 | 0.192,0.205 | 0.341,0.29 | -0.117 |
| 1509 | XLOC_016525 | chr16 | + | 88740011 | 88741311 | 88740011 | 88740167 | 88740771 | 604 | 88741311 | 1509 | 6,9 | 4,0 | 11,10 | 0,0 | 753 | 149 | 0.002743528 | 0.026365349 | 0.229,1.0 | 1.0,1.0 | -0.385 |
| 1574 | XLOC_016339 | chr16 | + | 28847253 | 28848525 | 28847253 | 28847443 | 28848048 | 605 | 28848525 | 1574 | 14,401,395 | 46,37 | 17,921,872 | 5,7 | 754 | 149 | 1.11E-16 | 7.26E-14 | 0.861,0.882 | 0.986,0.981 | -0.112 |
| 1566 | XLOC_000120 | chr1 | + | 22827984 | 22828934 | 22827984 | 22828177 | 22828791 | 614 | 22828934 | 1566 | 43,36 | 28,23 | 106,106 | 38,29 | 763 | 149 | 0.003216611 | 0.029701823 | 0.231,0.234 | 0.353,0.416 | -0.152 |
| 355 | XLOC_035165 | chr5 | - | 177651642 | 177652430 | 177651642 | 177651730 | 177652355 | 625 | 177652430 | 355 | 62,60 | 102,74 | 109,52 | 53,52 | 774 | 149 | 0.001127896 | 0.013422314 | 0.105,0.135 | 0.284,0.161 | -0.102 |
| 3017 | XLOC_029452 | chr3 | + | 52324975 | 52325938 | 52324975 | 52325127 | 52325762 | 635 | 52325938 | 3017 | 18,24 | 12,34 | 29,24 | 9,7 | 784 | 149 | 0.000633256 | 0.008661183 | 0.222,0.118 | 0.38,0.395 | -0.218 |
| 16230 | XLOC_009618 | chr12 | - | 133403153 | 133405283 | 133403153 | 133403553 | 133404192 | 639 | 133405283 | 16230 | 33,37 | 16,8 | 25,16 | 2,0 | 788 | 149 | 0.000974543 | 0.012045252 | 0.281,0.467 | 0.703,1.0 | -0.477 |
| 12428 | XLOC_007509 | chr11 | - | 47598962 | 47600541 | 47598962 | 47599580 | 47600236 | 656 | 47600541 | 12428 | 5,4 | 1,1 | 8,6 | 1,0 | 805 | 149 | 0.003227742 | 0.029786313 | 0.481,0.425 | 0.597,1.0 | -0.346 |
| 10956 | XLOC_038262 | chr6 | - | 41039334 | 41040352 | 41039334 | 41039414 | 41040073 | 659 | 41040352 | 10956 | 36,36 | 21,20 | 33,49 | 8,9 | 808 | 149 | 0.002476096 | 0.024407227 | 0.24,0.249 | 0.432,0.501 | -0.222 |
| 3031 | XLOC_016775 | chr16 | - | 67863667 | 67865261 | 67863667 | 67864451 | 67865118 | 667 | 67865261 | 3031 | 77,72 | 88,104 | 147,192 | 56,71 | 816 | 149 | 4.30E-12 | 1.12E-09 | 0.138,0.112 | 0.324,0.331 | -0.203 |
| 11893 | XLOC_011461 | chr13 | - | 41834628 | 41835961 | 41834628 | 41835051 | 41835718 | 667 | 41835961 | 11893 | 17,16 | 3,5 | 40,34 | 2,1 | 816 | 149 | 0.002916735 | 0.027569316 | 0.509,0.369 | 0.785,0.861 | -0.384 |
| 1354 | XLOC_018071 | chr17 | + | 40838920 | 40839999 | 40838920 | 40839064 | 40839737 | 673 | 40839999 | 1354 | 34,68 | 30,35 | 21,27 | 2,2 | 822 | 149 | 6.74E-06 | 0.000236209 | 0.17,0.26 | 0.656,0.71 | -0.468 |
| 8006 | XLOC_018641 | chr17 | - | 55943834 | 55944776 | 55943834 | 55943905 | 55944589 | 684 | 55944776 | 8006 | 22,29 | 6,0 | 13,21 | 0,0 | 833 | 149 | 0.00087872 | 0.011173042 | 0.396,1.0 | 1.0,1.0 | -0.302 |
| 651 | XLOC_028756 | chr22 | - | 51220615 | 51222070 | 51220615 | 51220779 | 51221472 | 693 | 51222070 | 651 | 49,43 | 8,4 | 51,55 | 0,2 | 842 | 149 | 0.000366617 | 0.005729728 | 0.52,0.655 | 1.0,0.83 | -0.328 |
| 3454 | XLOC_005363 | chr10 | - | 103866339 | 103868934 | 103866339 | 103868080 | 103868785 | 705 | 103868934 | 3454 | 619,607 | 81,57 | 766,775 | 53,51 | 854 | 149 | 0.003736763 | 0.033338308 | 0.571,0.65 | 0.716,0.726 | -0.11 |
| 609 | XLOC_029512 | chr3 | + | 101571934 | 101573527 | 101571934 | 101572707 | 101573421 | 714 | 101573527 | 609 | 14,16 | 25,37 | 44,19 | 17,23 | 863 | 149 | 0.001716746 | 0.018644978 | 0.088,0.069 | 0.309,0.125 | -0.139 |
| 9983 | XLOC_041043 | chr7 | + | 26241280 | 26242229 | 26241280 | 26241446 | 26242160 | 714 | 26242229 | 9983 | 89,65 | 1,4 | 149,144 | 0,1 | 863 | 149 | 0.005068043 | 0.041919834 | 0.939,0.737 | 1.0,0.961 | -0.142 |
| 3105 | XLOC_000628 | chr1 | + | 180165396 | 180169004 | 180165396 | 180165734 | 180166467 | 733 | 180169004 | 3105 | 27,113,172 | 196,219 | 23,012,105 | 69,85 | 882 | 149 | 2.70E-07 | 1.59E-05 | 0.7,0.71 | 0.849,0.807 | -0.123 |
| 3004 | XLOC_000907 | chr1 | - | 16944679 | 16946519 | 16944679 | 16945598 | 16946339 | 741 | 16946519 | 3004 | 16,35 | 50,38 | 55,47 | 18,26 | 890 | 149 | 1.07E-06 | 5.13E-05 | 0.051,0.134 | 0.338,0.232 | -0.193 |
| 12996 | XLOC_000991 | chr1 | - | 32681706 | 32682952 | 32681706 | 32682118 | 32682859 | 741 | 32682952 | 12996 | 21,24 | 4,0 | 15,28 | 0,0 | 890 | 149 | 0.004176005 | 0.036163195 | 0.468,1.0 | 1.0,1.0 | -0.266 |
| 2784 | XLOC_016600 | chr16 | - | 3431568 | 3434941 | 3431568 | 3433711 | 3434458 | 747 | 3434941 | 2784 | 34,36 | 39,41 | 102,87 | 33,49 | 896 | 149 | 9.01E-05 | 0.001903505 | 0.127,0.127 | 0.339,0.228 | -0.157 |
| 2291 | XLOC_046145 | chr9 | + | 139745206 | 139746391 | 139745206 | 139745557 | 139746308 | 751 | 139746391 | 2291 | 13,5 | 11,6 | 55,41 | 16,10 | 900 | 149 | 0.001075093 | 0.012957966 | 0.164,0.121 | 0.363,0.404 | -0.241 |
| 2318 | XLOC_021921 | chr19 | - | 59080882 | 59081894 | 59080882 | 59080953 | 59081710 | 757 | 59081894 | 2318 | 109,106 | 96,113 | 226,175 | 102,84 | 906 | 149 | 6.86E-06 | 0.000239691 | 0.157,0.134 | 0.267,0.255 | -0.115 |
| 2403 | XLOC_000749 | chr1 | + | 225156460 | 225158905 | 225156460 | 225156576 | 225157335 | 759 | 225158905 | 2403 | 14,17 | 6,4 | 95,78 | 8,4 | 908 | 149 | 0.001796167 | 0.019216177 | 0.277,0.411 | 0.661,0.762 | -0.368 |
| 7328 | XLOC_021165 | chr19 | + | 37341336 | 37342806 | 37341336 | 37341870 | 37342629 | 759 | 37342806 | 7328 | 2,3 | 2,2 | 1,5 | 0,0 | 908 | 149 | 2.68E-07 | 1.59E-05 | 0.141,0.198 | 1.0,1.0 | -0.831 |
| 3240 | XLOC_001334 | chr1 | - | 153914344 | 153915606 | 153914344 | 153914589 | 153915353 | 764 | 153915606 | 3240 | 172,136 | 12,7 | 223,123 | 4,0 | 913 | 149 | 0.000176489 | 0.003289643 | 0.701,0.76 | 0.901,1.0 | -0.22 |
| 1097 | XLOC_029980 | chr3 | - | 119155637 | 119156950 | 119155637 | 119155825 | 119156601 | 776 | 119156950 | 1097 | 104,106 | 94,54 | 209,184 | 64,50 | 925 | 149 | 5.73E-06 | 0.000208231 | 0.151,0.24 | 0.345,0.372 | -0.163 |
| 2729 | XLOC_018559 | chr17 | - | 40854894 | 40855838 | 40854894 | 40854979 | 40855757 | 778 | 40855838 | 2729 | 196,211 | 21,18 | 322,315 | 17,12 | 927 | 149 | 0.005269922 | 0.04315197 | 0.6,0.653 | 0.753,0.808 | -0.154 |
| 902 | XLOC_023985 | chr2 | - | 27883850 | 27885148 | 27883850 | 27884255 | 27885045 | 790 | 27885148 | 902 | 446,459 | 117,150 | 531,479 | 83,80 | 939 | 149 | 4.68E-05 | 0.001147905 | 0.377,0.327 | 0.504,0.487 | -0.144 |
| 8009 | XLOC_018641 | chr17 | - | 55943834 | 55944776 | 55943834 | 55943905 | 55944698 | 793 | 55944776 | 8009 | 22,29 | 0,6 | 14,22 | 0,0 | 942 | 149 | 0.000632536 | 0.008661183 | 1.0,0.433 | 1.0,1.0 | -0.283 |
| 5810 | XLOC_027814 | chr21 | - | 45752915 | 45754354 | 45752915 | 45753145 | 45753943 | 798 | 45754354 | 5810 | 12,8 | 0,4 | 7,15 | 0,0 | 947 | 149 | 0.002479101 | 0.024411711 | 1.0,0.239 | 1.0,1.0 | -0.381 |
| 1473 | XLOC_004859 | chr10 | + | 70516029 | 70517134 | 70516029 | 70516240 | 70517050 | 810 | 70517134 | 1473 | 380,339 | 208,221 | 360,370 | 117,98 | 959 | 149 | 4.25E-07 | 2.39E-05 | 0.221,0.192 | 0.323,0.37 | -0.14 |
| 780 | XLOC_026863 | chr20 | + | 60754237 | 60758004 | 60754237 | 60754412 | 60755224 | 812 | 60758004 | 780 | 408,451 | 7,18 | 461,432 | 2,4 | 961 | 149 | 0.000255953 | 0.004400051 | 0.9,0.795 | 0.973,0.944 | -0.111 |
| 2645 | XLOC_016664 | chr16 | - | 21829976 | 21831764 | 21829976 | 21830802 | 21831615 | 813 | 21831764 | 2645 | 27,16 | 6,4 | 38,27 | 2,1 | 962 | 149 | 0.004463428 | 0.038255777 | 0.411,0.383 | 0.746,0.807 | -0.379 |
| 590 | XLOC_029664 | chr3 | + | 180326488 | 180327791 | 180326488 | 180326598 | 180327417 | 819 | 180327791 | 590 | 445,508 | 46,37 | 462,477 | 24,21 | 968 | 149 | 0.005550649 | 0.044912785 | 0.598,0.679 | 0.748,0.778 | -0.124 |

| | | | | | | | | | | | | | | | | | | | | | |
|---|---|---|---|---|---|---|---|---|---|---|---|---|---|---|---|---|---|---|---|---|---|
| 2215 | XLOC_009376 | chr12 | - | 53564206 | 53565225 | 53564206 | 53564286 | 53565109 | 823 | 53565225 | 2215 | 135,107 | 12,10 | 199,134 | 7,3 | 972 | 149 | 0.002760903 | 0.02646877 | 0.633,0.621 | 0.813,0.873 | -0.216 |
| 3424 | XLOC_007076 | chr11 | + | 47436838 | 47438150 | 47436838 | 47437047 | 47437875 | 828 | 47438150 | 3424 | 321,408 | 8,12 | 430,314 | 2,2 | 977 | 149 | 0.000186998 | 0.003426658 | 0.86,0.838 | 0.97,0.96 | -0.116 |
| 9064 | XLOC_021326 | chr19 | + | 50381754 | 50382717 | 50381754 | 50381829 | 50382662 | 833 | 50382717 | 9064 | 60,60 | 1,5 | 32,15 | 0,0 | 982 | 149 | 0.000173008 | 0.003252984 | 0.901,0.645 | 1.0,1.0 | -0.227 |
| 5215 | XLOC_029956 | chr3 | - | 108300954 | 108301930 | 108300954 | 108300985 | 108301823 | 838 | 108301930 | 5215 | 87,61 | 2,3 | 56,47 | 0,0 | 987 | 149 | 0.001699631 | 0.018472433 | 0.868,0.754 | 1.0,1.0 | -0.189 |
| 16086 | XLOC_034614 | chr5 | + | 87565061 | 87578683 | 87565061 | 87565504 | 87566342 | 838 | 87578683 | 16086 | 4,11 | 2,1 | 11,11 | 0,0 | 987 | 149 | 0.001376608 | 0.015616088 | 0.232,0.624 | 1.0,1.0 | -0.572 |
| 1466 | XLOC_016820 | chr16 | - | 75268786 | 75270896 | 75268786 | 75269884 | 75270725 | 841 | 75270896 | 1466 | 60,86 | 4,4 | 122,131 | 1,1 | 990 | 149 | 0.001498089 | 0.016630449 | 0.693,0.764 | 0.948,0.952 | -0.222 |
| 364 | XLOC_035102 | chr5 | - | 148871471 | 148876423 | 148871471 | 148873184 | 148874036 | 852 | 148876423 | 364 | 343,334 | 16,8 | 310,362 | 3,4 | 1001 | 149 | 0.001648036 | 0.018015807 | 0.761,0.861 | 0.939,0.931 | -0.124 |
| 503 | XLOC_032343 | chr4 | + | 68586139 | 68588555 | 68586139 | 68586339 | 68587194 | 855 | 68588555 | 503 | 37,34 | 1,9 | 71,35 | 1,0 | 1004 | 149 | 0.000226751 | 0.003984564 | 0.846,0.359 | 0.913,1.0 | -0.354 |
| 40 | XLOC_046502 | chr9 | - | 140469203 | 140470619 | 140469203 | 140469298 | 140470160 | 862 | 140470619 | 40 | 280,259 | 33,45 | 360,352 | 21,16 | 1011 | 149 | 1.44E-05 | 0.000435211 | 0.556,0.459 | 0.716,0.764 | -0.232 |
| 1748 | XLOC_006939 | chr11 | + | 290389 | 292095 | 290389 | 291113 | 291975 | 862 | 292095 | 1748 | 213,202 | 20,19 | 348,329 | 20,7 | 1011 | 149 | 0.004435137 | 0.038100131 | 0.611,0.61 | 0.719,0.874 | -0.186 |
| 9438 | XLOC_014944 | chr15 | - | 65113565 | 65114590 | 65113565 | 65113596 | 65114464 | 868 | 65114590 | 9438 | 50,61 | 4,3 | 74,65 | 0,1 | 1017 | 149 | 0.00031807 | 0.005155618 | 0.647,0.749 | 1.0,0.905 | -0.255 |
| 522 | XLOC_037917 | chr6 | + | 88049833 | 88054575 | 88049833 | 88050055 | 88050943 | 888 | 88054575 | 522 | 63,68 | 7,2 | 89,65 | 0,0 | 1037 | 149 | 5.23E-06 | 0.000191994 | 0.564,0.83 | 1.0,1.0 | -0.303 |
| 1583 | XLOC_016758 | chr16 | - | 67212101 | 67213465 | 67212101 | 67212407 | 67213304 | 897 | 67213465 | 1583 | 195,240 | 7,4 | 216,218 | 0,1 | 1046 | 149 | 0.000103605 | 0.002140687 | 0.799,0.895 | 1.0,0.969 | -0.137 |
| 2131 | XLOC_007597 | chr11 | - | 66133901 | 66135019 | 66133901 | 66134035 | 66134934 | 899 | 66135019 | 2131 | 268,266 | 143,125 | 384,306 | 108,85 | 1048 | 149 | 8.54E-05 | 0.001833049 | 0.21,0.232 | 0.336,0.339 | -0.117 |
| 10127 | XLOC_013436 | chr14 | - | 69584855 | 69585947 | 69584855 | 69584992 | 69585910 | 918 | 69585947 | 10127 | 81,84 | 12,14 | 76,98 | 4,5 | 1067 | 149 | 0.002766603 | 0.026489651 | 0.485,0.456 | 0.726,0.732 | -0.258 |
| 539 | XLOC_030174 | chr3 | - | 197338397 | 197342556 | 197338397 | 197341493 | 197342434 | 941 | 197342556 | 539 | 517,505 | 28,21 | 796,645 | 11,10 | 1090 | 149 | 7.36E-06 | 0.000253332 | 0.716,0.767 | 0.908,0.898 | -0.161 |
| 833 | XLOC_029685 | chr3 | + | 184556501 | 184557540 | 184556501 | 184556534 | 184557485 | 951 | 184557540 | 833 | 20,14 | 16,9 | 66,47 | 16,13 | 1100 | 149 | 0.001607869 | 0.017666597 | 0.145,0.174 | 0.358,0.329 | -0.184 |
| 822 | XLOC_018271 | chr17 | + | 78084739 | 78085899 | 78084739 | 78084824 | 78085781 | 957 | 78085899 | 822 | 233,248 | 155,163 | 479,479 | 175,156 | 1106 | 149 | 2.63E-06 | 0.00010833 | 0.168,0.17 | 0.269,0.293 | -0.112 |
| 3116 | XLOC_005198 | chr10 | - | 43046004 | 43047262 | 43046004 | 43046063 | 43047020 | 957 | 43047262 | 3116 | 20,23 | 7,9 | 24,18 | 0,1 | 1106 | 149 | 7.59E-06 | 0.000259386 | 0.278,0.256 | 1.0,0.708 | -0.587 |
| 2659 | XLOC_011393 | chr13 | + | 115007595 | 115008823 | 115007595 | 115007755 | 115008731 | 976 | 115008823 | 2659 | 514,552 | 258,297 | 842,700 | 169,161 | 1125 | 149 | 4.12E-12 | 1.09E-09 | 0.209,0.198 | 0.398,0.365 | -0.178 |
| 1978 | XLOC_009388 | chr12 | - | 54104919 | 54106990 | 54104919 | 54105905 | 54106883 | 978 | 54106990 | 1978 | 432,431 | 116,127 | 716,697 | 41,59 | 1127 | 149 | 6.66E-16 | 3.85E-13 | 0.33,0.31 | 0.698,0.61 | -0.334 |
| 2160 | XLOC_021499 | chr19 | - | 6413123 | 6415312 | 6413123 | 6414131 | 6415151 | 1020 | 6415312 | 2160 | 19,221,731 | 61,95 | 39,984,001 | 80,90 | 1169 | 149 | 8.29E-05 | 0.001795144 | 0.801,0.699 | 0.864,0.85 | -0.107 |
| 14390 | XLOC_027820 | chr21 | - | 46686350 | 46687628 | 46686350 | 46686475 | 46687504 | 1029 | 46687628 | 14390 | 32,31 | 1,4 | 35,34 | 0,0 | 1178 | 149 | 4.36E-05 | 0.001084669 | 0.802,0.495 | 1.0,1.0 | -0.351 |
| 2603 | XLOC_041436 | chr7 | - | 27201389 | 27204586 | 27201389 | 27203460 | 27204496 | 1036 | 27204586 | 2603 | 90,120 | 43,39 | 183,144 | 35,26 | 1185 | 149 | 0.000556342 | 0.007858199 | 0.208,0.279 | 0.397,0.411 | -0.161 |
| 2571 | XLOC_014712 | chr15 | + | 75644942 | 75646207 | 75644942 | 75645042 | 75646079 | 1037 | 75646207 | 2571 | 106,106 | 8,5 | 177,142 | 2,3 | 1186 | 149 | 0.004394625 | 0.037795279 | 0.625,0.727 | 0.917,0.856 | -0.211 |
| 2307 | XLOC_008778 | chr12 | + | 9464326 | 9466893 | 9464326 | 9464427 | 9465470 | 1043 | 9466893 | 2307 | 116,91 | 6,3 | 157,146 | 1,0 | 1192 | 149 | 3.51E-05 | 0.000922934 | 0.707,0.791 | 0.952,1.0 | -0.227 |
| 76 | XLOC_046113 | chr9 | + | 135930330 | 135931481 | 135930330 | 135930351 | 135931398 | 1047 | 135931481 | 76 | 175,166 | 24,16 | 463,431 | 12,21 | 1196 | 149 | 5.29E-05 | 0.001255961 | 0.476,0.564 | 0.828,0.719 | -0.253 |
| 3157 | XLOC_018763 | chr17 | - | 78183459 | 78186073 | 78183459 | 78184810 | 78185869 | 1059 | 78186073 | 3157 | 382,428 | 59,60 | 426,378 | 36,21 | 1208 | 149 | 0.000380922 | 0.005888823 | 0.444,0.468 | 0.593,0.689 | -0.185 |
| 2743 | XLOC_017938 | chr17 | + | 16219994 | 16221222 | 16219994 | 16220026 | 16221088 | 1062 | 16221222 | 2743 | 60,56 | 12,14 | 160,131 | 12,13 | 1211 | 149 | 0.002093098 | 0.021519576 | 0.381,0.33 | 0.621,0.554 | -0.232 |
| 1490 | XLOC_035181 | chr5 | - | 180272366 | 180278493 | 180272366 | 180275082 | 180276153 | 1071 | 180278493 | 1490 | 561,535 | 21,21 | 712,678 | 4,6 | 1220 | 149 | 1.25E-07 | 8.30E-06 | 0.765,0.757 | 0.956,0.932 | -0.183 |
| 2015 | XLOC_011561 | chr13 | - | 113978518 | 113980161 | 113978518 | 113978892 | 113979975 | 1083 | 113980161 | 2015 | 103,114 | 80,101 | 225,249 | 75,67 | 1232 | 149 | 1.62E-09 | 2.01E-07 | 0.135,0.12 | 0.266,0.31 | -0.161 |
| 134 | XLOC_044017 | chr8 | + | 42177102 | 42178362 | 42177102 | 42177164 | 42178252 | 1088 | 42178362 | 134 | 106,117 | 72,71 | 261,230 | 77,89 | 1237 | 149 | 0.00016919 | 0.003197179 | 0.151,0.166 | 0.29,0.237 | -0.105 |
| 1434 | XLOC_034813 | chr5 | + | 178309709 | 178316339 | 178309709 | 178311519 | 178312614 | 1095 | 178316339 | 1434 | 73,45 | 28,19 | 98,111 | 16,7 | 1244 | 149 | 0.000492815 | 0.007173519 | 0.238,0.221 | 0.423,0.655 | -0.31 |
| 2760 | XLOC_005338 | chr10 | - | 101370036 | 101373681 | 101370036 | 101372356 | 101373452 | 1096 | 101373681 | 2760 | 319,312 | 67,35 | 614,618 | 43,55 | 1245 | 149 | 0.004137151 | 0.035950912 | 0.363,0.516 | 0.631,0.574 | -0.163 |
| 14173 | XLOC_007142 | chr11 | + | 64852586 | 64854080 | 64852586 | 64852786 | 64853886 | 1100 | 64854080 | 14173 | 25,16 | 2,3 | 18,39 | 1,0 | 1249 | 149 | 0.000854974 | 0.010949508 | 0.599,0.389 | 0.682,1.0 | -0.347 |
| 2771 | XLOC_001135 | chr1 | - | 59154709 | 59155921 | 59154709 | 59154788 | 59155897 | 1109 | 59155921 | 2771 | 152,203 | 35,55 | 406,327 | 40,35 | 1258 | 149 | 3.56E-06 | 0.000137281 | 0.34,0.304 | 0.546,0.525 | -0.214 |
| 1370 | XLOC_017998 | chr17 | + | 28958772 | 28960408 | 28958772 | 28958997 | 28960115 | 1118 | 28960408 | 1370 | 192,171 | 43,54 | 206,154 | 18,19 | 1267 | 149 | 5.17E-05 | 0.001233985 | 0.344,0.271 | 0.574,0.488 | -0.223 |
| 870 | XLOC_024190 | chr2 | - | 118693070 | 118694315 | 118693070 | 118693155 | 118694277 | 1122 | 118694315 | 870 | 53,44 | 58,64 | 189,187 | 77,58 | 1271 | 149 | 6.32E-11 | 1.22E-08 | 0.097,0.075 | 0.223,0.274 | -0.163 |
| 206 | XLOC_041217 | chr7 | + | 99950995 | 99952863 | 99950995 | 99951635 | 99952765 | 1130 | 99952863 | 206 | 107,130 | 126,102 | 343,268 | 76,77 | 1279 | 149 | 1.11E-16 | 7.26E-14 | 0.09,0.129 | 0.345,0.288 | -0.207 |
| 528 | XLOC_032657 | chr4 | - | 83343838 | 83347282 | 83343838 | 83346036 | 83347189 | 1153 | 83347282 | 528 | 26,172,600 | 623,596 | 34,953,592 | 500,457 | 1302 | 149 | 6.72E-07 | 3.39E-05 | 0.325,0.333 | 0.444,0.474 | -0.13 |
| 12561 | XLOC_007305 | chr11 | + | 117109316 | 117110576 | 117109316 | 117109362 | 117110516 | 1154 | 117110576 | 12561 | 49,38 | 9,7 | 50,63 | 3,2 | 1303 | 149 | 0.002930355 | 0.027599961 | 0.384,0.383 | 0.656,0.783 | -0.336 |
| 2402 | XLOC_000749 | chr1 | + | 225155132 | 225156576 | 225155132 | 225155285 | 225156460 | 1175 | 225156576 | 2402 | 39,37 | 6,4 | 211,157 | 5,4 | 1324 | 149 | 0.000684556 | 0.009210285 | 0.422,0.51 | 0.826,0.815 | -0.355 |
| 583 | XLOC_029727 | chr3 | - | 9428391 | 9433510 | 9428391 | 9432181 | 9433362 | 1181 | 9433510 | 583 | 490,436 | 48,34 | 478,385 | 13,9 | 1330 | 149 | 2.68E-07 | 1.59E-05 | 0.534,0.59 | 0.805,0.827 | -0.254 |
| 175 | XLOC_041438 | chr7 | - | 27228813 | 27238061 | 27228813 | 27233177 | 27234404 | 1227 | 27238061 | 175 | 21,20 | 4,0 | 14,16 | 0,0 | 1376 | 149 | 0.002497984 | 0.024542568 | 0.362,1.0 | 1.0,1.0 | -0.319 |
| 917 | XLOC_009559 | chr12 | - | 115117309 | 115118951 | 115117309 | 115117456 | 115118683 | 1227 | 115118951 | 917 | 231,207 | 116,117 | 487,535 | 158,138 | 1376 | 149 | 2.35E-05 | 0.000658335 | 0.177,0.161 | 0.25,0.296 | -0.104 |
| 6995 | XLOC_020849 | chr19 | + | 798014 | 799443 | 798014 | 798171 | 799412 | 1241 | 799443 | 6995 | 10,13 | 2,1 | 24,13 | 0,0 | 1390 | 149 | 0.002074555 | 0.021358975 | 0.349,0.582 | 1.0,1.0 | -0.534 |
| 7984 | XLOC_001539 | chr1 | - | 226109903 | 226111471 | 226109903 | 226110083 | 226111336 | 1253 | 226111471 | 7984 | 31,47 | 6,0 | 57,70 | 0,0 | 1402 | 149 | 0.000779767 | 0.010181646 | 0.354,1.0 | 1.0,1.0 | -0.323 |
| 3191 | XLOC_000476 | chr1 | + | 146739084 | 146740537 | 146739084 | 146739177 | 146740440 | 1263 | 146740537 | 3191 | 242,253 | 101,119 | 381,283 | 68,88 | 1412 | 149 | 0.00102557 | 0.012532659 | 0.202,0.183 | 0.372,0.253 | -0.12 |
| 9696 | XLOC_014760 | chr15 | + | 89824995 | 89826481 | 89824995 | 89825066 | 89826362 | 1296 | 89826481 | 9696 | 67,73 | 2,6 | 98,82 | 0,0 | 1445 | 149 | 2.70E-06 | 0.000110681 | 0.775,0.556 | 1.0,1.0 | -0.335 |
| 2388 | XLOC_005079 | chr10 | + | 124187791 | 124193261 | 124187791 | 124187832 | 124189139 | 1307 | 124193261 | 2388 | 38,64 | 14,16 | 138,122 | 8,13 | 1456 | 149 | 0.000174454 | 0.003267916 | 0.217,0.29 | 0.638,0.49 | -0.311 |
| 147 | XLOC_041749 | chr7 | - | 141356343 | 141364040 | 141356343 | 141362651 | 141363971 | 1320 | 141364040 | 147 | 236,176 | 67,83 | 204,230 | 45,43 | 1469 | 149 | 0.002047788 | 0.021199463 | 0.263,0.177 | 0.315,0.352 | -0.114 |
| 8555 | XLOC_021853 | chr19 | - | 52566421 | 52570863 | 52566421 | 52566425 | 52567746 | 1321 | 52570863 | 8555 | 8,17 | 2,1 | 15,20 | 0,0 | 1470 | 149 | 0.00200704 | 0.020930873 | 0.288,0.633 | 1.0,1.0 | -0.539 |
| 5470 | XLOC_029391 | chr3 | + | 42702976 | 42704619 | 42702976 | 42703124 | 42704465 | 1341 | 42704619 | 5470 | 1,0 | 0,2 | 7,5 | 0,0 | 1490 | 149 | 1.13E-05 | 0.000363808 | 1.0,0.0 | 1.0,1.0 | -0.5 |
| 5124 | XLOC_041515 | chr7 | - | 64465336 | 64467093 | 64465336 | 64465572 | 64466918 | 1346 | 64467093 | 5124 | 2,9 | 0,3 | 10,9 | 0,0 | 1495 | 149 | 0.002457601 | 0.024295557 | 1.0,0.23 | 1.0,1.0 | -0.385 |
| 11413 | XLOC_030086 | chr3 | - | 156868070 | 156869719 | 156868070 | 156868170 | 156869517 | 1347 | 156869719 | 11413 | 160,173 | 8,1 | 307,261 | 0,0 | 1496 | 149 | 2.15E-05 | 0.000610852 | 0.666,0.945 | 1.0,1.0 | -0.195 |
| 8873 | XLOC_016718 | chr16 | - | 46989257 | 46991132 | 46989257 | 46989534 | 46990918 | 1384 | 46991132 | 8873 | 311,267 | 10,15 | 379,387 | 0,2 | 1533 | 149 | 8.25E-09 | 8.11E-07 | 0.751,0.634 | 1.0,0.95 | -0.282 |
| 14935 | XLOC_004882 | chr10 | + | 74883971 | 74886585 | 74883971 | 74885001 | 74886392 | 1391 | 74886585 | 14935 | 23,16 | 5,0 | 15,34 | 0,0 | 1540 | 149 | 0.000886605 | 0.011244785 | 0.308,1.0 | 1.0,1.0 | -0.346 |
| 2178 | XLOC_009511 | chr12 | - | 104166091 | 104171859 | 104166091 | 104169408 | 104170821 | 1413 | 104171859 | 2178 | 139,162 | 3,2 | 129,111 | 0,0 | 1562 | 149 | 0.003165715 | 0.029339913 | 0.815,0.885 | 1.0,1.0 | -0.15 |
| 2522 | XLOC_013112 | chr14 | + | 50790662 | 50798961 | 50790662 | 50790878 | 50792326 | 1448 | 50798961 | 2522 | 84,68 | 3,2 | 86,111 | 0,0 | 1597 | 149 | 0.000366821 | 0.005729728 | 0.723,0.76 | 1.0,1.0 | -0.258 |
| 15758 | XLOC_000882 | chr1 | - | 10516563 | 10523266 | 10516563 | 10521759 | 10523221 | 1462 | 10523266 | 15758 | 158,171 | 6,0 | 215,231 | 0,0 | 1611 | 149 | 0.000787926 | 0.010261457 | 0.709,1.0 | 1.0,1.0 | -0.145 |
| 3419 | XLOC_004807 | chr10 | + | 30726104 | 30728203 | 30726104 | 30726335 | 30727844 | 1509 | 30728203 | 3419 | 52,52 | 64,77 | 46,68 | 29,13 | 1658 | 149 | 4.07E-05 | 0.001029275 | 0.068,0.057 | 0.125,0.32 | -0.16 |
| 1836 | XLOC_001538 | chr1 | - | 226046895 | 226048697 | 226046895 | 226047049 | 226048559 | 1510 | 226048697 | 1836 | 151,126 | 81,103 | 424,428 | 72,122 | 1659 | 149 | 3.74E-09 | 4.16E-07 | 0.143,0.099 | 0.346,0.24 | -0.172 |
| 5867 | XLOC_034684 | chr5 | + | 134124202 | 134125912 | 134124202 | 134124281 | 134125795 | 1514 | 134125912 | 5867 | 31,24 | 17,16 | 45,40 | 12,4 | 1663 | 149 | 0.004816185 | 0.040359362 | 0.14,0.118 | 0.251,0.473 | -0.233 |
| 52 | XLOC_046427 | chr9 | - | 130823531 | 130826179 | 130823531 | 130824105 | 130825652 | 1547 | 130826179 | 52 | 72,127 | 34,51 | 255,206 | 35,46 | 1696 | 149 | 6.74E-05 | 0.001521194 | 0.157,0.18 | 0.39,0.282 | -0.167 |
| 2511 | XLOC_007242 | chr11 | + | 82922336 | 82925725 | 82922336 | 82922472 | 82924020 | 1548 | 82925725 | 2511 | 235,244 | 6,7 | 116,82 | 0,1 | 1697 | 149 | 0.000548653 | 0.007778364 | 0.775,0.754 | 1.0,0.878 | -0.175 |
| 9031 | XLOC_016435 | chr16 | + | 67195637 | 67197755 | 67195637 | 67195840 | 67197414 | 1574 | 67197755 | 9031 | 176,186 | 9,1 | 161,129 | 0,0 | 1723 | 149 | 5.61E-05 | 0.001316921 | 0.628,0.941 | 1.0,1.0 | -0.216 |
| 3288 | XLOC_028580 | chr22 | - | 20386516 | 20388425 | 20386516 | 20386675 | 20388299 | 1624 | 20388425 | 3288 | 21,13 | 2,1 | 60,42 | 0,0 | 1773 | 149 | 0.000260125 | 0.004446362 | 0.469,0.522 | 1.0,1.0 | -0.504 |
| 7037 | XLOC_021763 | chr19 | - | 44788580 | 44795350 | 44788580 | 44793349 | 44794992 | 1643 | 44795350 | 7037 | 9,2 | 2,2 | 2,1 | 0,0 | 1792 | 149 | 7.57E-08 | 5.45E-06 | 0.272,0.077 | 1.0,1.0 | -0.826 |
| 432 | XLOC_034597 | chr5 | + | 78936927 | 78938733 | 78936927 | 78937019 | 78938666 | 1647 | 78938733 | 432 | 68,41 | 52,37 | 61,67 | 17,20 | 1796 | 149 | 3.63E-05 | 0.000944131 | 0.098,0.084 | 0.229,0.217 | -0.132 |
| 1034 | XLOC_023585 | chr2 | + | 97858717 | 97860491 | 97858717 | 97858790 | 97860462 | 1672 | 97860491 | 1034 | 8,7 | 11,18 | 11,12 | 7,4 | 1821 | 149 | 0.004769617 | 0.04008077 | 0.056,0.031 | 0.114,0.197 | -0.112 |
| 1417 | XLOC_046504 | chr9 | - | 140509716 | 140512643 | 140509716 | 140510651 | 140512385 | 1734 | 140512643 | 1417 | 116,123 | 85,67 | 163,166 | 28,30 | 1883 | 149 | 8.50E-11 | 1.54E-08 | 0.097,0.127 | 0.315,0.305 | -0.198 |

| | | | | | | | | | | | | | | | | | | | | | |
|---|---|---|---|---|---|---|---|---|---|---|---|---|---|---|---|---|---|---|---|---|---|
| 3548 | XLOC_048083 | chrX | - | 130919580 | 130928143 | 130919580 | 130926262 | 130928021 | 1759 | 130928143 | 3548 | 0,4 | 3,2 | 12,16 | 1,0 | 1908 | 149 | 7.76E-06 | 0.000264201 | 0.0,0.135 | 0.484,1.0 | -0.674 |
| 2203 | XLOC_030086 | chr3 | - | 156868070 | 156870030 | 156868070 | 156868170 | 156869965 | 1795 | 156870030 | 2203 | 12,761,303 | 112,81 | 24,012,171 | 81,60 | 1944 | 149 | 6.53E-08 | 4.91E-06 | 0.466,0.552 | 0.694,0.735 | -0.205 |
| 2099 | XLOC_030015 | chr3 | - | 126178495 | 126181063 | 126178495 | 126178580 | 126180377 | 1797 | 126181063 | 2099 | 699,639 | 57,36 | 831,684 | 23,11 | 1946 | 149 | 2.35E-06 | 9.85E-05 | 0.484,0.576 | 0.734,0.826 | -0.25 |
| 768 | XLOC_026983 | chr20 | - | 34326889 | 34328809 | 34326889 | 34326939 | 34328745 | 1806 | 34328809 | 768 | 26,082,872 | 307,319 | 28,512,877 | 162,160 | 1955 | 149 | 6.20E-09 | 6.43E-07 | 0.393,0.407 | 0.573,0.578 | -0.175 |
| 1744 | XLOC_041183 | chr7 | + | 92078065 | 92080074 | 92078065 | 92078191 | 92080014 | 1823 | 92080074 | 1744 | 379,387 | 74,75 | 688,574 | 76,78 | 1972 | 149 | 0.003396503 | 0.030926268 | 0.279,0.281 | 0.406,0.357 | -0.101 |
| 2098 | XLOC_000380 | chr1 | + | 95709766 | 95712213 | 95709766 | 95710271 | 95712097 | 1826 | 95712213 | 2098 | 314,266 | 3,8 | 292,257 | 2,0 | 1975 | 149 | 0.001779653 | 0.019076435 | 0.888,0.715 | 0.917,1.0 | -0.157 |
| 21 | XLOC_047911 | chrX | - | 18915280 | 18917344 | 18915280 | 18915451 | 18917290 | 1839 | 18917344 | 21 | 829,707 | 101,86 | 11,071,119 | 83,76 | 1988 | 149 | 0.000380908 | 0.005888823 | 0.381,0.381 | 0.5,0.525 | -0.131 |
| 747 | XLOC_034684 | chr5 | + | 134124202 | 134126242 | 134124202 | 134124281 | 134126159 | 1878 | 134126242 | 747 | 305,268 | 273,215 | 394,439 | 175,112 | 2027 | 149 | 6.44E-10 | 9.49E-08 | 0.076,0.084 | 0.142,0.224 | -0.103 |
| 2096 | XLOC_000380 | chr1 | + | 95709766 | 95712213 | 95709766 | 95710207 | 95712097 | 1890 | 95712213 | 2096 | 359,300 | 9,9 | 328,283 | 0,3 | 2039 | 149 | 0.000271474 | 0.00458821 | 0.745,0.709 | 1.0,0.873 | -0.21 |
| 263 | XLOC_038373 | chr6 | - | 99850415 | 99852578 | 99850415 | 99850586 | 99852478 | 1892 | 99852578 | 263 | 24,452,523 | 71,43 | 21,982,111 | 25,20 | 2041 | 149 | 0.000208763 | 0.003742804 | 0.715,0.811 | 0.865,0.885 | -0.112 |
| 15005 | XLOC_021620 | chr19 | - | 19102148 | 19104549 | 19102148 | 19102551 | 19104446 | 1895 | 19104549 | 15005 | 172,157 | 5,1 | 192,153 | 0,0 | 2044 | 149 | 0.000159147 | 0.003030244 | 0.715,0.92 | 1.0,1.0 | -0.182 |
| 2236 | XLOC_009218 | chr12 | - | 6571410 | 6574107 | 6571410 | 6572135 | 6574055 | 1920 | 6574107 | 2236 | 399,434 | 4,5 | 289,253 | 0,0 | 2069 | 149 | 0.00021668 | 0.003855951 | 0.878,0.862 | 1.0,1.0 | -0.13 |
| 454 | XLOC_041145 | chr7 | + | 75042066 | 75044301 | 75042066 | 75042210 | 75044162 | 1952 | 75044301 | 454 | 475,513 | 52,53 | 658,649 | 27,17 | 2101 | 149 | 1.13E-07 | 7.68E-06 | 0.393,0.407 | 0.633,0.73 | -0.281 |
| 2092 | XLOC_037614 | chr6 | + | 348768 | 351482 | 348768 | 348841 | 350821 | 1980 | 351482 | 2092 | 357,363 | 42,39 | 391,408 | 16,16 | 2129 | 149 | 4.93E-06 | 0.000183004 | 0.373,0.394 | 0.631,0.641 | -0.253 |
| 4409 | XLOC_041489 | chr7 | - | 44116107 | 44118410 | 44116107 | 44116354 | 44118338 | 1984 | 44118410 | 4409 | 6,9 | 4,1 | 19,14 | 0,0 | 2133 | 149 | 7.01E-10 | 1.01E-07 | 0.095,0.386 | 1.0,1.0 | -0.76 |
| 2943 | XLOC_000133 | chr1 | + | 24973157 | 24975520 | 24973157 | 24973280 | 24975349 | 2069 | 24975520 | 2943 | 16,971,709 | 216,183 | 19,621,625 | 77,75 | 2218 | 149 | 2.11E-13 | 7.39E-11 | 0.345,0.386 | 0.631,0.593 | -0.246 |
| 15004 | XLOC_021620 | chr19 | - | 19102148 | 19104549 | 19102148 | 19102349 | 19104446 | 2097 | 19104549 | 15004 | 172,157 | 4,2 | 192,153 | 0,0 | 2246 | 149 | 0.000181006 | 0.003343346 | 0.74,0.839 | 1.0,1.0 | -0.211 |
| 15009 | XLOC_021620 | chr19 | - | 19102148 | 19104549 | 19102148 | 19102349 | 19104456 | 2107 | 19104549 | 15009 | 149,141 | 2,2 | 175,154 | 0,0 | 2256 | 149 | 0.005030498 | 0.041736763 | 0.831,0.823 | 1.0,1.0 | -0.173 |
| 15570 | XLOC_001218 | chr1 | - | 109468392 | 109477564 | 109468392 | 109475123 | 109477241 | 2118 | 109477564 | 15570 | 75,53 | 2,9 | 73,71 | 0,0 | 2267 | 149 | 6.17E-07 | 3.18E-05 | 0.711,0.279 | 1.0,1.0 | -0.505 |
| 3282 | XLOC_001218 | chr1 | - | 109468392 | 109477564 | 109468392 | 109475123 | 109477246 | 2123 | 109477564 | 3282 | 536,468 | 95,109 | 760,660 | 86,72 | 2272 | 149 | 6.12E-05 | 0.001400363 | 0.27,0.22 | 0.367,0.375 | -0.126 |
| 2621 | XLOC_000342 | chr1 | + | 75229577 | 75232352 | 75229577 | 75229834 | 75231967 | 2133 | 75232352 | 2621 | 11,691,003 | 37,52 | 17,161,771 | 25,22 | 2282 | 149 | 1.16E-06 | 5.48E-05 | 0.674,0.557 | 0.818,0.84 | -0.213 |
| 913 | XLOC_023912 | chr2 | + | 241500746 | 241503936 | 241500746 | 241500940 | 241503090 | 2150 | 241503936 | 913 | 66,71 | 3,1 | 86,84 | 0,0 | 2299 | 149 | 0.001482182 | 0.016518693 | 0.588,0.821 | 1.0,1.0 | -0.296 |
| 220 | XLOC_041198 | chr7 | + | 98530863 | 98533298 | 98530863 | 98531025 | 98533201 | 2176 | 98533298 | 220 | 63,44 | 64,70 | 194,135 | 40,43 | 2325 | 149 | 2.89E-15 | 1.40E-12 | 0.059,0.039 | 0.237,0.167 | -0.153 |
| 3226 | XLOC_000629 | chr1 | + | 180240969 | 180252105 | 180240969 | 180241141 | 180243319 | 2178 | 180252105 | 3226 | 276,239 | 4,8 | 224,160 | 2,0 | 2327 | 149 | 0.002493284 | 0.024512406 | 0.815,0.657 | 0.878,1.0 | -0.203 |
| 1476 | XLOC_034983 | chr5 | - | 94031559 | 94046632 | 94031559 | 94044314 | 94046522 | 2208 | 94046632 | 1476 | 264,276 | 26,19 | 1,018,881 | 26,25 | 2357 | 149 | 3.56E-06 | 0.000137281 | 0.391,0.479 | 0.712,0.69 | -0.266 |
| 11008 | XLOC_000380 | chr1 | + | 95709766 | 95712213 | 95709766 | 95709880 | 95712097 | 2217 | 95712213 | 11008 | 311,274 | 0,6 | 287,251 | 0,0 | 2366 | 149 | 0.001207853 | 0.014138932 | 1.0,0.742 | 1.0,1.0 | -0.129 |
| 14038 | XLOC_021000 | chr19 | + | 13261200 | 13265742 | 13261200 | 13261507 | 13263757 | 2250 | 13265742 | 14038 | 124,109 | 0,5 | 174,133 | 0,0 | 2399 | 149 | 0.001178263 | 0.01385726 | 1.0,0.575 | 1.0,1.0 | -0.213 |
| 14902 | XLOC_001012 | chr1 | - | 35643589 | 35652723 | 35643589 | 35650194 | 35652547 | 2353 | 35652723 | 14902 | 116,112 | 21,22 | 151,169 | 2,6 | 2502 | 149 | 4.21E-07 | 2.37E-05 | 0.248,0.233 | 0.818,0.627 | -0.482 |
| 2147 | XLOC_011380 | chr13 | + | 111932884 | 111935710 | 111932884 | 111933024 | 111935485 | 2461 | 111935710 | 2147 | 381,402 | 49,56 | 801,739 | 54,59 | 2610 | 149 | 0.000508596 | 0.007334906 | 0.307,0.291 | 0.459,0.417 | -0.139 |
| 524 | XLOC_037689 | chr6 | + | 26428870 | 26432480 | 26428870 | 26429013 | 26431499 | 2486 | 26432480 | 524 | 128,140 | 2,5 | 155,157 | 0,1 | 2635 | 149 | 0.0003595 | 0.005675864 | 0.784,0.613 | 1.0,0.899 | -0.251 |
| 3980 | XLOC_046188 | chr9 | - | 14677060 | 14680160 | 14677060 | 14677498 | 14680030 | 2532 | 14680160 | 3980 | 3,5 | 0,4 | 2,2 | 0,0 | 2681 | 149 | 0.00012026 | 0.002411943 | 1.0,0.065 | 1.0,1.0 | -0.468 |
| 195 | XLOC_041301 | chr7 | + | 129100141 | 129102881 | 129100141 | 129100213 | 129102806 | 2593 | 129102881 | 195 | 164,215 | 65,54 | 350,324 | 50,34 | 2742 | 149 | 4.54E-07 | 2.51E-05 | 0.121,0.178 | 0.276,0.341 | -0.159 |
| 1232 | XLOC_000461 | chr1 | + | 145281954 | 145293580 | 145281954 | 145290511 | 145293113 | 2602 | 145293580 | 1232 | 381,433 | 24,17 | 945,734 | 9,19 | 2751 | 149 | 0.000896484 | 0.011331863 | 0.462,0.58 | 0.85,0.677 | -0.243 |
| 13255 | XLOC_001307 | chr1 | - | 150944087 | 150947366 | 150944087 | 150944437 | 150947094 | 2657 | 150947366 | 13255 | 1,1 | 0,2 | 10,2 | 0,0 | 2806 | 149 | 1.17E-05 | 0.000374604 | 1.0,0.026 | 1.0,1.0 | -0.487 |
| 64 | XLOC_046278 | chr9 | - | 74975543 | 74978522 | 74975543 | 74975703 | 74978385 | 2682 | 74978522 | 64 | 642,573 | 203,227 | 714,678 | 102,116 | 2831 | 149 | 1.63E-09 | 2.01E-07 | 0.143,0.117 | 0.269,0.235 | -0.122 |
| 12413 | XLOC_020127 | chr18 | + | 59942606 | 59945686 | 59942606 | 59942706 | 59945448 | 2742 | 59945686 | 12413 | 2,4 | 2,0 | 2,6 | 0,0 | 2891 | 149 | 0.00057956 | 0.008079468 | 0.049,1.0 | 1.0,1.0 | -0.476 |
| 1247 | XLOC_013404 | chr14 | - | 59937884 | 59942724 | 59937884 | 59939808 | 59942586 | 2778 | 59942724 | 1247 | 240,255 | 43,53 | 438,469 | 42,44 | 2927 | 149 | 0.000122114 | 0.002429626 | 0.221,0.197 | 0.347,0.352 | -0.14 |
| 1190 | XLOC_020232 | chr18 | - | 47794348 | 47799108 | 47794348 | 47796188 | 47799046 | 2858 | 47799108 | 1190 | 766,806 | 47,29 | 10,481,086 | 19,25 | 3007 | 149 | 0.000654448 | 0.008916852 | 0.447,0.579 | 0.732,0.683 | -0.195 |
| 521 | XLOC_045969 | chr9 | + | 96859602 | 96864033 | 96859602 | 96860861 | 96863847 | 2986 | 96864033 | 521 | 381,384 | 54,57 | 814,704 | 24,26 | 3135 | 149 | 2.75E-14 | 1.15E-11 | 0.251,0.243 | 0.617,0.563 | -0.343 |
| 14772 | XLOC_045867 | chr9 | + | 6493825 | 6497360 | 6493825 | 6493932 | 6496959 | 3027 | 6497360 | 14772 | 150,131 | 5,7 | 236,195 | 4,0 | 3176 | 149 | 0.004560712 | 0.038934297 | 0.585,0.468 | 0.735,1.0 | -0.341 |
| 13616 | XLOC_018612 | chr17 | - | 46629388 | 46632980 | 46629388 | 46629849 | 46632892 | 3043 | 46632980 | 13616 | 2,1 | 0,2 | 6,2 | 0,0 | 3192 | 149 | 3.11E-05 | 0.00082753 | 1.0,0.023 | 1.0,1.0 | -0.489 |
| 2657 | XLOC_041202 | chr7 | + | 99074025 | 99085371 | 99074025 | 99074103 | 99077283 | 3180 | 99085371 | 2657 | 300,273 | 37,31 | 338,265 | 5,5 | 3329 | 149 | 4.69E-11 | 9.30E-09 | 0.266,0.283 | 0.752,0.703 | -0.453 |
| 2652 | XLOC_016868 | chr16 | - | 90104128 | 90110397 | 90104128 | 90106825 | 90110055 | 3230 | 90110397 | 2652 | 349,331 | 33,26 | 400,453 | 12,10 | 3379 | 149 | 3.11E-06 | 0.000123685 | 0.318,0.36 | 0.595,0.666 | -0.292 |
| 2467 | XLOC_007508 | chr11 | - | 47487344 | 47493909 | 47487344 | 47490443 | 47493742 | 3299 | 47493909 | 2467 | 40,374,414 | 99,115 | 50,584,502 | 74,77 | 3448 | 149 | 0.001252525 | 0.014518066 | 0.638,0.624 | 0.747,0.716 | -0.101 |
| 642 | XLOC_029298 | chr3 | + | 9471510 | 9475634 | 9471510 | 9471735 | 9475053 | 3318 | 9475634 | 642 | 172,179 | 30,35 | 324,316 | 9,12 | 3467 | 149 | 3.73E-11 | 7.79E-09 | 0.198,0.18 | 0.607,0.531 | -0.38 |
| 5838 | XLOC_027757 | chr21 | - | 33735464 | 33739064 | 33735464 | 33735638 | 33738962 | 3324 | 33739064 | 5838 | 88,91 | 4,0 | 150,134 | 0,0 | 3473 | 149 | 0.003509129 | 0.031759517 | 0.486,1.0 | 1.0,1.0 | -0.257 |
| 5837 | XLOC_027757 | chr21 | - | 33735464 | 33739064 | 33735464 | 33735581 | 33738926 | 3345 | 33739064 | 5837 | 82,94 | 6,0 | 129,130 | 0,0 | 3494 | 149 | 0.000645484 | 0.008810676 | 0.368,1.0 | 1.0,1.0 | -0.316 |
| 14024 | XLOC_021412 | chr19 | + | 58264658 | 58272912 | 58264658 | 58264747 | 58268239 | 3492 | 58272912 | 14024 | 149,121 | 4,2 | 108,127 | 1,0 | 3641 | 149 | 0.001287316 | 0.014837546 | 0.604,0.712 | 0.815,1.0 | -0.25 |
| 14326 | XLOC_023692 | chr2 | + | 160615736 | 160625224 | 160615736 | 160615869 | 160619390 | 3521 | 160625224 | 14326 | 6,6 | 10,4 | 5,11 | 0,0 | 3670 | 149 | 9.03E-09 | 8.66E-07 | 0.024,0.057 | 1.0,1.0 | -0.96 |
| 14486 | XLOC_009610 | chr12 | - | 125267228 | 125271049 | 125267228 | 125267357 | 125270986 | 3629 | 125271049 | 14486 | 14,581,497 | 36,53 | 14,061,302 | 25,20 | 3778 | 149 | 0.005946249 | 0.047125124 | 0.615,0.527 | 0.689,0.72 | -0.133 |
| 2302 | XLOC_000336 | chr1 | + | 70694104 | 70698060 | 70694104 | 70694238 | 70697950 | 3712 | 70698060 | 2302 | 67,736,522 | 367,315 | 84,668,025 | 240,252 | 3861 | 149 | 3.65E-06 | 0.000140563 | 0.416,0.444 | 0.577,0.551 | -0.134 |
| 3311 | XLOC_027761 | chr21 | - | 34127523 | 34131580 | 34127523 | 34127664 | 34131390 | 3726 | 34131580 | 3311 | 362,381 | 75,70 | 915,858 | 49,46 | 3875 | 149 | 1.97E-14 | 8.69E-12 | 0.157,0.173 | 0.418,0.418 | -0.253 |
| 2911 | XLOC_000386 | chr1 | + | 100609624 | 100613882 | 100609624 | 100609699 | 100613449 | 3750 | 100613882 | 2911 | 819,706 | 36,40 | 16,831,432 | 17,16 | 3899 | 149 | 2.45E-12 | 6.83E-10 | 0.465,0.403 | 0.791,0.774 | -0.348 |
| 680 | XLOC_028673 | chr22 | - | 38883883 | 38888120 | 38883883 | 38884120 | 38888060 | 3940 | 38888120 | 680 | 1,512,215,556 | 917,786 | 2,036,518,421 | 722,686 | 4089 | 149 | 7.63E-05 | 0.001681276 | 0.375,0.419 | 0.507,0.495 | -0.104 |
| 2912 | XLOC_000386 | chr1 | + | 100609624 | 100613882 | 100609624 | 100609699 | 100613648 | 3949 | 100613882 | 2912 | 948,805 | 17,24 | 18,361,582 | 6,5 | 4098 | 149 | 1.72E-09 | 2.10E-07 | 0.67,0.549 | 0.918,0.92 | -0.309 |
| 3346 | XLOC_032661 | chr4 | - | 83816844 | 83821376 | 83816844 | 83816927 | 83821229 | 4302 | 83821376 | 3346 | 902,805 | 26,35 | 903,927 | 17,11 | 4451 | 149 | 0.001588753 | 0.017482093 | 0.537,0.435 | 0.64,0.738 | -0.203 |
| 641 | XLOC_029298 | chr3 | + | 9470506 | 9475246 | 9470506 | 9470693 | 9475053 | 4360 | 9475246 | 641 | 229,242 | 19,24 | 397,383 | 14,8 | 4509 | 149 | 9.00E-05 | 0.001903505 | 0.285,0.25 | 0.484,0.613 | -0.281 |
| 2121 | XLOC_011529 | chr13 | - | 79884552 | 79894845 | 79884552 | 79889128 | 79893669 | 4541 | 79894845 | 2121 | 10,991,077 | 12,22 | 13,811,396 | 1,4 | 4690 | 149 | 7.56E-08 | 5.45E-06 | 0.744,0.609 | 0.978,0.917 | -0.271 |
| 7812 | XLOC_041258 | chr7 | + | 107383228 | 107389492 | 107383228 | 107383595 | 107389324 | 5729 | 107389492 | 7812 | 61,28 | 5,2 | 59,56 | 1,0 | 5878 | 149 | 0.00024745 | 0.004278324 | 0.236,0.262 | 0.599,1.0 | -0.55 |
| 3006 | XLOC_000907 | chr1 | - | 16944679 | 16952500 | 16944679 | 16946519 | 16952269 | 5750 | 16952500 | 3006 | 266,244 | 41,30 | 495,519 | 39,29 | 5899 | 149 | 0.000362258 | 0.005701256 | 0.141,0.17 | 0.243,0.311 | -0.122 |
| 2505 | XLOC_007264 | chr11 | + | 95555034 | 95561191 | 95555034 | 95555142 | 95560949 | 5807 | 95561191 | 2505 | 10,501,126 | 5,6 | 907,861 | 2,0 | 5956 | 149 | 0.003033533 | 0.028412453 | 0.84,0.824 | 0.919,1.0 | -0.128 |
| 2919 | XLOC_034863 | chr5 | - | 34159948 | 34176062 | 34159948 | 34169258 | 34175549 | 6291 | 34176062 | 2919 | 281,218 | 17,18 | 391,344 | 8,6 | 6440 | 149 | 3.18E-05 | 0.000843346 | 0.277,0.219 | 0.531,0.57 | -0.302 |
| 2404 | XLOC_000749 | chr1 | + | 225231587 | 225239497 | 225231587 | 225231685 | 225239152 | 7467 | 225239497 | 2404 | 9,11 | 1,2 | 4,1 | 0,0 | 7616 | 149 | 1.40E-07 | 9.09E-06 | 0.15,0.097 | 1.0,1.0 | -0.877 |
| 595 | XLOC_029597 | chr3 | + | 140681968 | 140689835 | 140681968 | 140682069 | 140689768 | 7699 | 140689835 | 595 | 26,602,571 | 163,161 | 41,243,705 | 66,97 | 7848 | 149 | 1.44E-09 | 1.84E-07 | 0.237,0.233 | 0.543,0.42 | -0.247 |
| 10933 | XLOC_014966 | chr15 | - | 70358352 | 70366946 | 70358352 | 70358554 | 70366841 | 8287 | 70366946 | 10933 | 280,255 | 8,6 | 277,274 | 0,1 | 8436 | 149 | 2.35E-06 | 9.85E-05 | 0.382,0.429 | 1.0,0.829 | -0.509 |
| 3353 | XLOC_005082 | chr10 | + | 124647784 | 124658477 | 124647784 | 124648239 | 124657308 | 9069 | 124658477 | 3353 | 458,407 | 5,4 | 656,698 | 0,1 | 9218 | 149 | 1.29E-05 | 0.000405287 | 0.597,0.622 | 1.0,0.919 | -0.35 |
| 14699 | XLOC_038025 | chr6 | + | 147660472 | 147674509 | 147660472 | 147660520 | 147674458 | 13938 | 147674509 | 14699 | 17,13 | 1,1 | 46,42 | 0,0 | 14087 | 149 | 2.32E-06 | 9.81E-05 | 0.152,0.121 | 1.0,1.0 | -0.863 |
| 1596 | XLOC_037960 | chr6 | + | 111805000 | 111824886 | 111805000 | 111806064 | 111824498 | 18434 | 111824886 | 1596 | 135,173 | 4,4 | 147,82 | 1,0 | 18583 | 149 | 0.001541002 | 0.017018909 | 0.213,0.257 | 0.541,1.0 | -0.535 |

**Table S1 (2)  The list of retained introns in SAP30BP-knockdown HeLa cells**

See rMATS (http://rnaseq-mats.sourceforge.net/) for the format of this table.

| ID | Gene ID | Chromosome | Strand | RI-Exon Start | RI-Exon End | Upstream ES | Upstream EE | Downstream ES | Intron (nt) | Downstream EE | ID.1 | IJC SAMPLE 1 | SJC SAMPLE 1 | IJC SAMPLE 2 | SJC SAMPLE 2 | Inc. Form (nt) | Skip Form (nt) | P Value | FDR | Inc. Level 1 | Inc. Level 2 | Inc. Level Differ. |
|---|---|---|---|---|---|---|---|---|---|---|---|---|---|---|---|---|---|---|---|---|---|---|
| 350 | XLOC_035171 | chr5 | - | 179044799 | 179045061 | 179044799 | 179044933 | 179044989 | 56 | 179045061 | 350 | 257,227 | 17,401,655 | 15,111,069 | 1,294,930 | 205 | 149 | 0 | 0 | 0.097,0.091 | 0.459,0.455 | -0.363 |
| 9393 | XLOC_009615 | chr12 | - | 132624967 | 132625280 | 132624967 | 132625100 | 132625164 | 64 | 132625280 | 9393 | 1,1 | 1,3 | 5,10 | 2,0 | 213 | 149 | 0.000691717 | 0.004508131 | 0.412,0.189 | 0.636,1.0 | -0.518 |
| 12853 | XLOC_027092 | chr20 | - | 62572169 | 62572561 | 62572169 | 62572396 | 62572462 | 66 | 62572561 | 12853 | 27,22 | 112,118 | 34,48 | 59,64 | 215 | 149 | 7.23E-07 | 1.10E-05 | 0.143,0.114 | 0.285,0.342 | -0.185 |
| 94 | XLOC_044466 | chr8 | - | 145737526 | 145737944 | 145737526 | 145737707 | 145737774 | 67 | 145737944 | 94 | 79,62 | 232,218 | 169,89 | 61,33 | 216 | 149 | 0 | 0 | 0.19,0.164 | 0.656,0.65 | -0.476 |
| 3701 | XLOC_046496 | chr9 | - | 140074632 | 140074995 | 140074632 | 140074836 | 140074903 | 67 | 140074995 | 3701 | 22,12 | 111,74 | 111,63 | 51,37 | 216 | 149 | 0 | 0 | 0.12,0.101 | 0.6,0.54 | -0.46 |
| 4057 | XLOC_044185 | chr8 | + | 145725306 | 145725774 | 145725306 | 145725582 | 145725649 | 67 | 145725774 | 4057 | 133,129 | 73,97 | 230,128 | 16,12 | 216 | 149 | 0 | 0 | 0.557,0.478 | 0.908,0.88 | -0.376 |
| 4411 | XLOC_044293 | chr8 | - | 48690985 | 48691360 | 48690985 | 48691221 | 48691288 | 67 | 48691360 | 4411 | 25,23 | 677,646 | 394,281 | 546,448 | 216 | 149 | 0 | 0 | 0.025,0.024 | 0.332,0.302 | -0.292 |
| 8348 | XLOC_016862 | chr16 | - | 89865358 | 89865640 | 89865358 | 89865506 | 89865573 | 67 | 89865640 | 8348 | 26,32 | 2,6 | 34,18 | 0,0 | 216 | 149 | 0.000122204 | 0.001018827 | 0.9,0.786 | 1.0,1.0 | -0.157 |
| 877 | XLOC_044459 | chr8 | - | 145579959 | 145580373 | 145579959 | 145580191 | 145580259 | 68 | 145580373 | 877 | 36,31 | 49,37 | 130,53 | 24,18 | 217 | 149 | 5.88E-10 | 1.59E-08 | 0.335,0.365 | 0.788,0.669 | -0.379 |
| 8396 | XLOC_016188 | chr16 | + | 722159 | 722384 | 722159 | 722185 | 722253 | 68 | 722384 | 8396 | 2,10 | 3,1 | 34,20 | 0,1 | 217 | 149 | 0.003034246 | 0.015403544 | 0.314,0.873 | 1.0,0.932 | -0.372 |
| 1539 | XLOC_016368 | chr16 | + | 30767687 | 30768036 | 30767687 | 30767841 | 30767910 | 69 | 30768036 | 1539 | 77,71 | 65,75 | 70,54 | 17,6 | 218 | 149 | 3.24E-09 | 7.60E-08 | 0.447,0.393 | 0.738,0.86 | -0.379 |
| 4065 | XLOC_044183 | chr8 | + | 145617018 | 145617370 | 145617018 | 145617222 | 145617291 | 69 | 145617370 | 4065 | 8,6 | 26,31 | 16,5 | 13,9 | 218 | 149 | 0.004679893 | 0.021972781 | 0.174,0.117 | 0.457,0.275 | -0.221 |
| 6070 | XLOC_016187 | chr16 | + | 705584 | 705889 | 705584 | 705703 | 705772 | 69 | 705889 | 6070 | 18,18 | 111,99 | 113,45 | 19,17 | 218 | 149 | 0 | 0 | 0.1,0.111 | 0.803,0.644 | -0.618 |
| 11048 | XLOC_017988 | chr17 | + | 27074839 | 27075196 | 27074839 | 27074965 | 27075034 | 69 | 27075196 | 11048 | 42,47 | 241,209 | 247,148 | 308,264 | 218 | 149 | 2.63E-13 | 1.20E-11 | 0.106,0.133 | 0.354,0.277 | -0.196 |
| 12066 | XLOC_016556 | chr16 | - | 424004 | 424408 | 424004 | 424151 | 424220 | 69 | 424408 | 12066 | 4,2 | 63,96 | 22,4 | 51,27 | 218 | 149 | 1.13E-06 | 1.63E-05 | 0.042,0.014 | 0.228,0.092 | -0.132 |
| 3272 | XLOC_018288 | chr17 | + | 79661844 | 79662097 | 79661844 | 79661883 | 79661953 | 70 | 79662097 | 3272 | 254,254 | 394,338 | 762,519 | 253,206 | 219 | 149 | 0 | 0 | 0.305,0.338 | 0.672,0.632 | -0.331 |
| 4186 | XLOC_041656 | chr7 | - | 102962958 | 102963241 | 102962958 | 102963079 | 102963149 | 70 | 102963241 | 4186 | 95,152 | 258,207 | 1,075,884 | 211,151 | 219 | 149 | 0 | 0 | 0.2,0.333 | 0.776,0.799 | -0.521 |
| 9333 | XLOC_006939 | chr11 | + | 294080 | 294478 | 294080 | 294196 | 294266 | 70 | 294478 | 9333 | 6,2 | 39,45 | 6,10 | 16,18 | 219 | 149 | 0.000628267 | 0.004163779 | 0.095,0.029 | 0.203,0.274 | -0.177 |
| 9844 | XLOC_013061 | chr14 | + | 24607410 | 24607777 | 24607410 | 24607612 | 24607682 | 70 | 24607777 | 9844 | 8,4 | 2,3 | 13,10 | 0,1 | 219 | 149 | 0.000930537 | 0.005738852 | 0.731,0.476 | 1.0,0.872 | -0.333 |
| 13695 | XLOC_007193 | chr11 | + | 67817340 | 67817721 | 67817340 | 67817533 | 67817603 | 70 | 67817721 | 13695 | 58,75 | 741,748 | 110,108 | 478,318 | 219 | 149 | 1.55E-13 | 7.33E-12 | 0.051,0.064 | 0.135,0.188 | -0.104 |
| 666 | XLOC_028745 | chr22 | - | 50656346 | 50656831 | 50656346 | 50656546 | 50656617 | 71 | 50656831 | 666 | 26,29 | 31,18 | 54,37 | 9,4 | 220 | 149 | 9.04E-07 | 1.34E-05 | 0.362,0.522 | 0.803,0.862 | -0.391 |
| 1850 | XLOC_027086 | chr20 | - | 62329998 | 62332054 | 62329998 | 62331882 | 62331953 | 71 | 62332054 | 1850 | 42,39 | 166,196 | 155,105 | 51,41 | 220 | 149 | 0 | 0 | 0.146,0.119 | 0.673,0.634 | -0.521 |
| 1874 | XLOC_013277 | chr14 | + | 105177273 | 105177523 | 105177273 | 105177344 | 105177415 | 71 | 105177523 | 1874 | 11,19 | 223,181 | 59,43 | 132,94 | 220 | 149 | 2.22E-15 | 1.32E-13 | 0.032,0.066 | 0.232,0.237 | -0.185 |
| 2901 | XLOC_016538 | chr16 | + | 89759681 | 89761196 | 89759681 | 89759734 | 89759805 | 71 | 89761196 | 2901 | 198,215 | 66,66 | 255,156 | 25,1 | 220 | 149 | 1.85E-11 | 6.39E-10 | 0.67,0.688 | 0.874,0.991 | -0.253 |
| 3206 | XLOC_029428 | chr3 | + | 49737904 | 49738142 | 49737904 | 49737997 | 49738068 | 71 | 49738142 | 3206 | 26,17 | 92,90 | 88,81 | 111,83 | 220 | 149 | 6.12E-10 | 1.65E-08 | 0.161,0.113 | 0.349,0.398 | -0.236 |
| 3988 | XLOC_044443 | chr8 | - | 144921188 | 144921513 | 144921188 | 144921272 | 144921343 | 71 | 144921513 | 3988 | 14,8 | 145,102 | 28,13 | 22,27 | 220 | 149 | 1.38E-08 | 2.89E-07 | 0.061,0.05 | 0.463,0.246 | -0.299 |
| 4785 | XLOC_034648 | chr5 | + | 112889463 | 112889729 | 112889463 | 112889551 | 112889622 | 71 | 112889729 | 4785 | 0,0 | 3,1 | 4,2 | 1,1 | 220 | 149 | 9.18E-07 | 1.36E-05 | 0.0,0.0 | 0.73,0.575 | -0.652 |
| 9153 | XLOC_016375 | chr16 | + | 31050861 | 31051135 | 31050861 | 31050972 | 31051043 | 71 | 31051135 | 9153 | 20,21 | 175,130 | 86,68 | 165,71 | 220 | 149 | 3.30E-11 | 1.11E-09 | 0.072,0.099 | 0.261,0.393 | -0.242 |
| 9858 | XLOC_018294 | chr17 | + | 79974675 | 79975009 | 79974675 | 79974745 | 79974816 | 71 | 79975009 | 9858 | 135,125 | 204,220 | 104,100 | 102,64 | 220 | 149 | 0.000184041 | 0.001444806 | 0.309,0.278 | 0.408,0.514 | -0.167 |
| 9975 | XLOC_016854 | chr16 | - | 88792954 | 88793206 | 88792954 | 88793051 | 88793122 | 71 | 88793206 | 9975 | 16,6 | 470,542 | 76,51 | 367,241 | 220 | 149 | 0 | 0 | 0.023,0.007 | 0.123,0.125 | -0.109 |
| 11746 | XLOC_007541 | chr11 | - | 62371401 | 62371673 | 62371401 | 62371513 | 62371584 | 71 | 62371673 | 11746 | 27,37 | 163,133 | 54,37 | 50,43 | 220 | 149 | 1.09E-11 | 3.89E-10 | 0.101,0.159 | 0.422,0.368 | -0.265 |
| 12193 | XLOC_000006 | chr1 | + | 981451 | 981645 | 981451 | 981468 | 981539 | 71 | 981645 | 12193 | 41,48 | 294,353 | 113,63 | 134,142 | 220 | 149 | 5.95E-12 | 2.20E-10 | 0.086,0.084 | 0.364,0.231 | -0.212 |
| 13174 | XLOC_044455 | chr8 | - | 145153983 | 145154305 | 145153983 | 145154108 | 145154179 | 71 | 145154305 | 13174 | 93,77 | 85,98 | 115,75 | 39,39 | 220 | 149 | 1.42E-05 | 0.000154626 | 0.426,0.347 | 0.666,0.566 | -0.23 |
| 13306 | XLOC_032424 | chr4 | + | 113568208 | 113568705 | 113568208 | 113568283 | 113568354 | 71 | 113568705 | 13306 | 9,7 | 114,80 | 55,40 | 81,100 | 220 | 149 | 5.95E-11 | 1.89E-09 | 0.051,0.056 | 0.315,0.213 | -0.211 |
| 13668 | XLOC_018727 | chr17 | - | 73909993 | 73910373 | 73909993 | 73910131 | 73910202 | 71 | 73910373 | 13668 | 14,19 | 91,63 | 21,9 | 21,15 | 220 | 149 | 0.000156767 | 0.001256896 | 0.094,0.17 | 0.404,0.289 | -0.215 |
| 13673 | XLOC_026823 | chr20 | + | 47682710 | 47683047 | 47682710 | 47682830 | 47682901 | 71 | 47683047 | 13673 | 2,11 | 449,391 | 90,57 | 432,330 | 220 | 149 | 0 | 0 | 0.003,0.019 | 0.124,0.105 | -0.103 |
| 14264 | XLOC_000825 | chr1 | - | 1233392 | 1233801 | 1233392 | 1233493 | 1233564 | 71 | 1233801 | 14264 | 7,6 | 3,3 | 9,9 | 0,1 | 220 | 149 | 0.000462795 | 0.003226085 | 0.612,0.575 | 1.0,0.859 | -0.336 |
| 14695 | XLOC_016224 | chr16 | + | 2578069 | 2578560 | 2578069 | 2578381 | 2578452 | 71 | 2578560 | 14695 | 2,5 | 97,77 | 28,22 | 26,15 | 220 | 149 | 0 | 0 | 0.014,0.042 | 0.422,0.498 | -0.432 |
| 460 | XLOC_046094 | chr9 | + | 132636854 | 132637234 | 132636854 | 132637035 | 132637107 | 72 | 132637234 | 460 | 6,1 | 27,22 | 10,4 | 8,10 | 221 | 149 | 0.003416075 | 0.017067566 | 0.13,0.03 | 0.457,0.212 | -0.255 |
| 897 | XLOC_024051 | chr2 | - | 61725807 | 61726048 | 61725807 | 61725927 | 61725999 | 72 | 61726048 | 897 | 19,46 | 572,563 | 274,201 | 448,339 | 221 | 149 | 0 | 0 | 0.022,0.052 | 0.292,0.286 | -0.252 |
| 1647 | XLOC_016761 | chr16 | - | 67272104 | 67272350 | 67272104 | 67272210 | 67272282 | 72 | 67272350 | 1647 | 27,15 | 60,51 | 50,24 | 51,37 | 221 | 149 | 0.003080756 | 0.0156004 | 0.233,0.165 | 0.398,0.304 | -0.152 |
| 2481 | XLOC_007374 | chr11 | - | 800761 | 801120 | 800761 | 800912 | 800984 | 72 | 801120 | 2481 | 16,9 | 19,13 | 15,18 | 7,0 | 221 | 149 | 0.001076563 | 0.006494787 | 0.362,0.318 | 0.591,1.0 | -0.456 |
| 2534 | XLOC_007173 | chr11 | + | 66372809 | 66373063 | 66372809 | 66372887 | 66372959 | 72 | 66373063 | 2534 | 8,6 | 165,135 | 97,48 | 70,52 | 221 | 149 | 0 | 0 | 0.032,0.029 | 0.483,0.384 | -0.403 |
| 3271 | XLOC_018288 | chr17 | + | 79660532 | 79660782 | 79660532 | 79660611 | 79660683 | 72 | 79660782 | 3271 | 103,112 | 405,398 | 480,299 | 264,211 | 221 | 149 | 0 | 0 | 0.146,0.159 | 0.551,0.489 | -0.368 |
| 8093 | XLOC_017896 | chr17 | + | 7128136 | 7128617 | 7128136 | 7128203 | 7128275 | 72 | 7128617 | 8093 | 121,186 | 19,131,957 | 542,326 | 14,251,229 | 221 | 149 | 0 | 0 | 0.041,0.06 | 0.204,0.152 | -0.128 |
| 8395 | XLOC_016188 | chr16 | + | 722159 | 722384 | 722159 | 722181 | 722253 | 72 | 722384 | 8395 | 4,12 | 328,286 | 54,33 | 191,174 | 221 | 149 | 0 | 0 | 0.008,0.028 | 0.16,0.113 | -0.119 |
| 8569 | XLOC_016435 | chr16 | + | 67201205 | 67201502 | 67201205 | 67201305 | 67201377 | 72 | 67201502 | 8569 | 22,28 | 4,4 | 42,32 | 0,0 | 221 | 149 | 4.91E-05 | 0.000463485 | 0.788,0.825 | 1.0,1.0 | -0.194 |
| 9808 | XLOC_013110 | chr14 | + | 50246312 | 50246528 | 50246312 | 50246393 | 50246465 | 72 | 50246528 | 9808 | 64,66 | 179,203 | 108,84 | 50,29 | 221 | 149 | 0 | 0 | 0.194,0.18 | 0.593,0.661 | -0.44 |
| 12678 | XLOC_018146 | chr17 | + | 48152723 | 48153099 | 48152723 | 48152890 | 48152962 | 72 | 48153099 | 12678 | 37,36 | 233,253 | 57,35 | 73,56 | 221 | 149 | 1.27E-13 | 6.13E-12 | 0.097,0.088 | 0.345,0.296 | -0.228 |
| 233 | XLOC_041771 | chr7 | - | 150773999 | 150774323 | 150773999 | 150774114 | 150774187 | 73 | 150774323 | 233 | 127,112 | 177,191 | 151,92 | 83,58 | 222 | 149 | 5.11E-09 | 1.16E-07 | 0.325,0.282 | 0.55,0.516 | -0.23 |
| 235 | XLOC_041771 | chr7 | - | 150773999 | 150774488 | 150773999 | 150774323 | 150774396 | 73 | 150774488 | 235 | 189,230 | 152,141 | 288,159 | 69,41 | 222 | 149 | 2.13E-10 | 6.18E-09 | 0.455,0.523 | 0.737,0.722 | -0.241 |
| 550 | XLOC_030002 | chr3 | - | 123649948 | 123650373 | 123649948 | 123650081 | 123650154 | 73 | 123650373 | 550 | 8,14 | 201,204 | 48,24 | 56,65 | 222 | 149 | 4.52E-14 | 2.25E-12 | 0.026,0.044 | 0.365,0.199 | -0.247 |
| 567 | XLOC_029841 | chr3 | - | 49160564 | 49161064 | 49160564 | 49160806 | 49160879 | 73 | 49161064 | 567 | 16,20 | 233,257 | 50,23 | 84,59 | 222 | 149 | 3.00E-15 | 1.76E-13 | 0.044,0.05 | 0.285,0.207 | -0.199 |
| 1292 | XLOC_028571 | chr22 | - | 19183776 | 19184167 | 19183776 | 19183926 | 19183999 | 73 | 19184167 | 1292 | 42,38 | 74,104 | 53,31 | 57,44 | 222 | 149 | 0.008253582 | 0.035280066 | 0.276,0.197 | 0.384,0.321 | -0.116 |
| 1674 | XLOC_021565 | chr19 | - | 12825860 | 12826138 | 12825860 | 12825983 | 12826056 | 73 | 12826138 | 1674 | 28,38 | 80,103 | 50,50 | 85,66 | 222 | 149 | 0.004327121 | 0.020618061 | 0.19,0.198 | 0.283,0.337 | -0.116 |
| 4041 | XLOC_046494 | chr9 | - | 140007657 | 140007930 | 140007657 | 140007739 | 140007812 | 73 | 140007930 | 4041 | 137,117 | 261,309 | 201,162 | 144,95 | 222 | 149 | 1.11E-16 | 7.47E-15 | 0.261,0.203 | 0.484,0.534 | -0.277 |
| 4307 | XLOC_041292 | chr7 | + | 128394319 | 128394776 | 128394319 | 128394509 | 128394582 | 73 | 128394776 | 4307 | 13,10 | 7,12 | 56,38 | 14,2 | 222 | 149 | 0.000906959 | 0.005617975 | 0.555,0.359 | 0.729,0.927 | -0.371 |
| 5367 | XLOC_028755 | chr22 | - | 51065018 | 51065468 | 51065018 | 51065188 | 51065261 | 73 | 51065468 | 5367 | 13,12 | 67,71 | 30,19 | 26,34 | 222 | 149 | 9.64E-06 | 0.000110484 | 0.115,0.102 | 0.436,0.273 | -0.246 |
| 5653 | XLOC_032798 | chr4 | - | 174253971 | 174254373 | 174253971 | 174254146 | 174254219 | 73 | 174254373 | 5653 | 204,205 | 18,431,905 | 330,226 | 852,624 | 222 | 149 | 0 | 0 | 0.069,0.067 | 0.206,0.196 | -0.133 |
| 5946 | XLOC_041630 | chr7 | - | 100280634 | 100281061 | 100280634 | 100280853 | 100280926 | 73 | 100281061 | 5946 | 29,27 | 240,284 | 142,92 | 217,156 | 222 | 149 | 0 | 0 | 0.075,0.06 | 0.305,0.284 | -0.227 |
| 6067 | XLOC_016187 | chr16 | + | 703340 | 703575 | 703340 | 703451 | 703524 | 73 | 703575 | 6067 | 23,28 | 96,135 | 56,32 | 75,55 | 222 | 149 | 8.12E-07 | 1.22E-05 | 0.139,0.122 | 0.334,0.281 | -0.177 |

| | | | | | | | | | | | | | | | | | | | | | | |
|---|---|---|---|---|---|---|---|---|---|---|---|---|---|---|---|---|---|---|---|---|---|---|
| 7334 | XLOC_016780 | chr16 | - | 68266273 | 68266587 | 68266273 | 68266406 | 68266479 | 73 | 68266587 | 7334 | 25,19 | 175,181 | 32,51 | 87,92 | 222 | 149 | 7.94E-09 | 1.74E-07 | 0.087,0.066 | 0.198,0.271 | -0.158 |
| 9057 | XLOC_014986 | chr15 | - | 75130492 | 75130707 | 75130492 | 75130533 | 75130606 | 73 | 75130707 | 9057 | 12,17 | 54,67 | 30,31 | 56,23 | 222 | 149 | 0.000467882 | 0.003258332 | 0.13,0.146 | 0.264,0.475 | -0.231 |
| 10404 | XLOC_044097 | chr8 | + | 95888301 | 95888721 | 95888301 | 95888375 | 95888448 | 73 | 95888721 | 10404 | 9,13 | 72,93 | 30,32 | 71,59 | 222 | 149 | 3.01E-06 | 3.91E-05 | 0.077,0.086 | 0.221,0.267 | -0.163 |
| 12790 | XLOC_016570 | chr16 | - | 1827747 | 1828031 | 1827747 | 1827873 | 1827946 | 73 | 1828031 | 12790 | 50,50 | 286,225 | 50,55 | 160,100 | 222 | 149 | 0.000327262 | 0.002393537 | 0.105,0.13 | 0.173,0.27 | -0.104 |
| 14368 | XLOC_000820 | chr1 | - | 892273 | 892539 | 892273 | 892405 | 892478 | 73 | 892539 | 14368 | 23,11 | 300,287 | 125,94 | 305,280 | 222 | 149 | 0 | 0 | 0.049,0.025 | 0.216,0.184 | -0.163 |
| 14584 | XLOC_044437 | chr8 | - | 144658581 | 144659012 | 144658581 | 144658742 | 144658815 | 73 | 144659012 | 14584 | 11,25 | 384,368 | 118,69 | 310,237 | 222 | 149 | 0 | 0 | 0.019,0.044 | 0.203,0.163 | -0.151 |
| 14831 | XLOC_001123 | chr1 | - | 53736594 | 53737018 | 53736594 | 53736825 | 53736898 | 73 | 53737018 | 14831 | 4,6 | 102,109 | 23,34 | 106,58 | 222 | 149 | 9.88E-09 | 2.13E-07 | 0.026,0.036 | 0.127,0.282 | -0.173 |
| 447 | XLOC_047682 | chrX | + | 44941820 | 44942034 | 44941820 | 44941885 | 44941959 | 74 | 44942034 | 447 | 4,0 | 3,12 | 17,16 | 6,4 | 223 | 149 | 0.00056079 | 0.003789493 | 0.471,0.0 | 0.654,0.728 | -0.456 |
| 878 | XLOC_044459 | chr8 | - | 145580467 | 145580781 | 145580467 | 145580575 | 145580649 | 74 | 145580781 | 878 | 66,57 | 29,39 | 117,82 | 5,9 | 223 | 149 | 4.44E-11 | 1.46E-09 | 0.603,0.494 | 0.94,0.859 | -0.351 |
| 1308 | XLOC_018550 | chr17 | - | 40342641 | 40342921 | 40342641 | 40342775 | 40342849 | 74 | 40342921 | 1308 | 10,9 | 37,30 | 29,23 | 17,11 | 223 | 149 | 2.67E-08 | 5.40E-07 | 0.153,0.167 | 0.533,0.583 | -0.398 |
| 1568 | XLOC_016179 | chr16 | + | 602300 | 602771 | 602300 | 602530 | 602604 | 74 | 602771 | 1568 | 36,38 | 138,134 | 152,78 | 95,110 | 223 | 149 | 7.13E-07 | 1.09E-05 | 0.148,0.159 | 0.517,0.321 | -0.266 |
| 1646 | XLOC_016761 | chr16 | - | 67263083 | 67263893 | 67263083 | 67263621 | 67263695 | 74 | 67263893 | 1646 | 30,46 | 100,121 | 100,76 | 35,32 | 223 | 149 | 0 | 0 | 0.167,0.203 | 0.656,0.613 | -0.45 |
| 2338 | XLOC_021828 | chr19 | - | 50365627 | 50365865 | 50365627 | 50365720 | 50365794 | 74 | 50365865 | 2338 | 6,15 | 171,154 | 22,37 | 140,93 | 223 | 149 | 1.63E-06 | 2.25E-05 | 0.023,0.061 | 0.095,0.21 | -0.111 |
| 2721 | XLOC_016200 | chr16 | + | 1550347 | 1550700 | 1550347 | 1550489 | 1550563 | 74 | 1550700 | 2721 | 18,24 | 88,80 | 22,23 | 39,37 | 223 | 149 | 0.001702059 | 0.00955033 | 0.12,0.167 | 0.274,0.293 | -0.14 |
| 3006 | XLOC_009522 | chr12 | - | 107386564 | 107386814 | 107386564 | 107386668 | 107386742 | 74 | 107386814 | 3006 | 14,21 | 72,83 | 63,43 | 14,11 | 223 | 149 | 0 | 0 | 0.115,0.145 | 0.75,0.723 | -0.606 |
| 3795 | XLOC_046296 | chr9 | - | 86593118 | 86593367 | 86593118 | 86593213 | 86593287 | 74 | 86593367 | 3795 | 0,0 | 1,2 | 1,9 | 2,3 | 223 | 149 | 7.92E-06 | 9.23E-05 | 0.0,0.0 | 0.25,0.667 | -0.459 |
| 5916 | XLOC_044184 | chr8 | + | 145693930 | 145694217 | 145693930 | 145694033 | 145694107 | 74 | 145694217 | 5916 | 5,5 | 109,78 | 14,7 | 31,15 | 223 | 149 | 2.32E-07 | 3.93E-06 | 0.03,0.041 | 0.232,0.238 | -0.199 |
| 6076 | XLOC_016187 | chr16 | + | 708952 | 709126 | 708952 | 709011 | 709085 | 74 | 709126 | 6076 | 60,47 | 105,90 | 80,51 | 38,39 | 223 | 149 | 2.19E-06 | 2.95E-05 | 0.276,0.259 | 0.584,0.466 | -0.258 |
| 6659 | XLOC_041547 | chr7 | - | 75616254 | 75616737 | 75616254 | 75616603 | 75616677 | 74 | 75616737 | 6659 | 27,16 | 174,165 | 66,58 | 60,46 | 223 | 149 | 0 | 0 | 0.094,0.061 | 0.424,0.457 | -0.363 |
| 12033 | XLOC_007159 | chr11 | + | 65632471 | 65632794 | 65632471 | 65632616 | 65632690 | 74 | 65632794 | 12033 | 29,22 | 114,116 | 79,59 | 90,58 | 223 | 149 | 7.78E-12 | 2.83E-10 | 0.145,0.112 | 0.37,0.405 | -0.259 |
| 12395 | XLOC_016203 | chr16 | + | 1816529 | 1816802 | 1816529 | 1816649 | 1816723 | 74 | 1816802 | 12395 | 2,7 | 43,38 | 23,10 | 28,11 | 223 | 149 | 1.86E-07 | 3.23E-06 | 0.03,0.11 | 0.354,0.378 | -0.296 |
| 14180 | XLOC_046382 | chr9 | - | 116151707 | 116151966 | 116151707 | 116151804 | 116151878 | 74 | 116151966 | 14180 | 0,0 | 29,38 | 7,5 | 21,15 | 223 | 149 | 1.06E-06 | 1.55E-05 | 0.0,0.0 | 0.182,0.182 | -0.182 |
| 14446 | XLOC_034803 | chr5 | + | 176520138 | 176520502 | 176520138 | 176520332 | 176520406 | 74 | 176520502 | 14446 | 12,11 | 32,38 | 24,22 | 24,15 | 223 | 149 | 0.000115505 | 0.000969845 | 0.2,0.162 | 0.401,0.495 | -0.267 |
| 14693 | XLOC_016224 | chr16 | + | 2577865 | 2578149 | 2577865 | 2577986 | 2578060 | 74 | 2578149 | 14693 | 14,14 | 89,57 | 41,29 | 14,14 | 223 | 149 | 0 | 0 | 0.095,0.141 | 0.662,0.581 | -0.503 |
| 95 | XLOC_044466 | chr8 | - | 145738024 | 145738521 | 145738024 | 145738154 | 145738229 | 75 | 145738521 | 95 | 97,99 | 272,272 | 287,153 | 149,101 | 224 | 149 | 0 | 0 | 0.192,0.195 | 0.562,0.502 | -0.339 |
| 566 | XLOC_029841 | chr3 | - | 49154491 | 49155003 | 49154491 | 49154794 | 49154869 | 75 | 49155003 | 566 | 5,7 | 63,41 | 17,13 | 9,16 | 224 | 149 | 1.15E-07 | 2.09E-06 | 0.05,0.102 | 0.557,0.351 | -0.378 |
| 1441 | XLOC_021139 | chr19 | + | 36104631 | 36104802 | 36104631 | 36104694 | 36104769 | 75 | 36104802 | 1441 | 14,15 | 100,114 | 39,12 | 44,39 | 224 | 149 | 5.29E-05 | 0.000494168 | 0.085,0.08 | 0.371,0.17 | -0.188 |
| 1852 | XLOC_020020 | chr18 | + | 2700536 | 2700916 | 2700536 | 2700657 | 2700732 | 75 | 2700916 | 1852 | 4,0 | 119,108 | 81,62 | 128,95 | 224 | 149 | 0 | 0 | 0.022,0.0 | 0.296,0.303 | -0.288 |
| 2151 | XLOC_018789 | chr17 | - | 80007552 | 80007882 | 80007552 | 80007718 | 80007793 | 75 | 80007882 | 2151 | 74,79 | 111,126 | 109,75 | 88,55 | 224 | 149 | 0.000131746 | 0.001083659 | 0.307,0.294 | 0.452,0.476 | -0.163 |
| 2742 | XLOC_000004 | chr1 | + | 897008 | 898297 | 897008 | 897130 | 897205 | 75 | 898297 | 2742 | 6,11 | 40,38 | 50,42 | 15,5 | 224 | 149 | 1.11E-16 | 7.47E-15 | 0.091,0.161 | 0.689,0.848 | -0.642 |
| 5410 | XLOC_028745 | chr22 | - | 50662569 | 50662841 | 50662569 | 50662685 | 50662760 | 75 | 50662841 | 5410 | 4,1 | 78,83 | 19,10 | 71,51 | 224 | 149 | 5.85E-07 | 9.15E-06 | 0.033,0.008 | 0.151,0.115 | -0.113 |
| 5773 | XLOC_028750 | chr22 | - | 50943485 | 50943746 | 50943485 | 50943582 | 50943657 | 75 | 50943746 | 5773 | 19,20 | 180,196 | 26,23 | 66,57 | 224 | 149 | 2.14E-07 | 3.65E-06 | 0.066,0.064 | 0.208,0.212 | -0.145 |
| 6068 | XLOC_016187 | chr16 | + | 703575 | 703803 | 703575 | 703670 | 703745 | 75 | 703803 | 6068 | 2,9 | 113,122 | 46,31 | 38,36 | 224 | 149 | 0 | 0 | 0.012,0.047 | 0.446,0.364 | -0.376 |
| 7655 | XLOC_018655 | chr17 | - | 58151106 | 58151520 | 58151106 | 58151247 | 58151322 | 75 | 58151520 | 7655 | 7,15 | 5,0 | 2,4 | 0,0 | 224 | 149 | 0.003187817 | 0.016061913 | 0.482,1.0 | 1.0,1.0 | -0.259 |
| 8547 | XLOC_016440 | chr16 | + | 67314620 | 67314908 | 67314620 | 67314709 | 67314784 | 75 | 67314908 | 8547 | 34,39 | 67,94 | 35,41 | 37,28 | 224 | 149 | 0.00024054 | 0.001821526 | 0.252,0.216 | 0.386,0.493 | -0.206 |
| 8640 | XLOC_000840 | chr1 | - | 1685462 | 1685822 | 1685462 | 1685647 | 1685722 | 75 | 1685822 | 8640 | 16,12 | 187,172 | 61,51 | 150,106 | 224 | 149 | 2.78E-14 | 1.40E-12 | 0.054,0.044 | 0.213,0.242 | -0.178 |
| 8830 | XLOC_001347 | chr1 | - | 155144826 | 155145104 | 155144826 | 155144965 | 155145040 | 75 | 155145104 | 8830 | 15,12 | 7,9 | 38,19 | 2,5 | 224 | 149 | 0.003049151 | 0.01546809 | 0.588,0.47 | 0.927,0.717 | -0.293 |
| 9352 | XLOC_018788 | chr17 | - | 79994066 | 79994313 | 79994066 | 79994184 | 79994259 | 75 | 79994313 | 9352 | 46,55 | 303,272 | 93,70 | 175,168 | 224 | 149 | 1.01E-08 | 2.16E-07 | 0.092,0.119 | 0.261,0.217 | -0.134 |
| 11236 | XLOC_008778 | chr12 | + | 9465212 | 9466893 | 9465212 | 9465625 | 9465700 | 75 | 9466893 | 11236 | 15,16 | 39,48 | 37,22 | 16,15 | 224 | 149 | 2.28E-07 | 3.88E-06 | 0.204,0.181 | 0.606,0.494 | -0.358 |
| 13587 | XLOC_041243 | chr7 | + | 102108495 | 102108824 | 102108495 | 102108634 | 102108709 | 75 | 102108824 | 13587 | 1,1 | 81,92 | 51,17 | 66,42 | 224 | 149 | 0 | 0 | 0.008,0.007 | 0.339,0.212 | -0.268 |
| 15103 | XLOC_007143 | chr11 | + | 64875013 | 64875494 | 64875013 | 64875199 | 64875274 | 75 | 64875494 | 15103 | 13,6 | 321,323 | 117,89 | 363,232 | 224 | 149 | 0 | 0 | 0.026,0.012 | 0.177,0.203 | -0.171 |
| 0 | XLOC_048133 | chrX | - | 153735533 | 153735821 | 153735533 | 153735660 | 153735736 | 76 | 153735821 | 0 | 18,13 | 182,178 | 49,40 | 99,72 | 225 | 149 | 1.49E-14 | 7.96E-13 | 0.061,0.046 | 0.247,0.269 | -0.205 |
| 123 | XLOC_044167 | chr8 | + | 144461484 | 144462953 | 144461484 | 144462155 | 144462231 | 76 | 144462953 | 123 | 51,60 | 42,52 | 36,24 | 4,4 | 225 | 149 | 9.70E-08 | 1.78E-06 | 0.446,0.433 | 0.856,0.799 | -0.388 |
| 804 | XLOC_041630 | chr7 | - | 100286073 | 100287178 | 100286073 | 100286138 | 100286214 | 76 | 100287178 | 804 | 39,40 | 123,111 | 91,54 | 68,45 | 225 | 149 | 4.50E-11 | 1.47E-09 | 0.174,0.193 | 0.47,0.443 | -0.273 |
| 818 | XLOC_013286 | chr14 | + | 105916394 | 105916661 | 105916394 | 105916521 | 105916597 | 76 | 105916661 | 818 | 56,84 | 235,229 | 164,130 | 275,204 | 225 | 149 | 4.05E-06 | 5.11E-05 | 0.136,0.195 | 0.283,0.297 | -0.124 |
| 2028 | XLOC_011375 | chr13 | + | 111286891 | 111287123 | 111286891 | 111286942 | 111287018 | 76 | 111287123 | 2028 | 86,113 | 94,107 | 177,133 | 70,59 | 225 | 149 | 3.46E-07 | 5.68E-06 | 0.377,0.412 | 0.626,0.599 | -0.218 |
| 2528 | XLOC_024028 | chr2 | - | 48061717 | 48062010 | 48061717 | 48061850 | 48061926 | 76 | 48062010 | 2528 | 0,0 | 92,69 | 42,23 | 88,75 | 225 | 149 | 0 | 0 | 0.0,0.0 | 0.24,0.169 | -0.205 |
| 2978 | XLOC_024343 | chr2 | - | 190618641 | 190619071 | 190618641 | 190618790 | 190618866 | 76 | 190619071 | 2978 | 3,0 | 32,30 | 5,4 | 30,5 | 225 | 149 | 0.005862312 | 0.026519358 | 0.058,0.0 | 0.099,0.346 | -0.193 |
| 3195 | XLOC_028665 | chr22 | - | 38228964 | 38229213 | 38228964 | 38229089 | 38229165 | 76 | 38229213 | 3195 | 3,1 | 67,80 | 9,10 | 42,50 | 225 | 149 | 1.28E-05 | 0.000140757 | 0.029,0.008 | 0.124,0.117 | -0.102 |
| 3273 | XLOC_018288 | chr17 | + | 79663636 | 79664028 | 79663636 | 79663777 | 79663853 | 76 | 79664028 | 3273 | 79,51 | 376,389 | 217,167 | 519,364 | 225 | 149 | 1.46E-10 | 4.35E-09 | 0.122,0.08 | 0.217,0.233 | -0.124 |
| 3423 | XLOC_007143 | chr11 | + | 64880691 | 64881066 | 64880691 | 64880886 | 64880962 | 76 | 64881066 | 3423 | 0,1 | 9,18 | 9,0 | 3,3 | 225 | 149 | 0.004147396 | 0.019933536 | 0.0,0.035 | 0.665,0.0 | -0.315 |
| 3994 | XLOC_044443 | chr8 | - | 144922100 | 144922300 | 144922100 | 144922202 | 144922278 | 76 | 144922300 | 3994 | 4,10 | 139,143 | 7,6 | 48,18 | 225 | 149 | 0.003292803 | 0.016531907 | 0.019,0.044 | 0.088,0.181 | -0.103 |
| 5744 | XLOC_026977 | chr20 | - | 33865523 | 33865912 | 33865523 | 33865735 | 33865811 | 76 | 33865912 | 5744 | 4,1 | 13,20 | 6,13 | 12,17 | 225 | 149 | 0.003061388 | 0.015519018 | 0.169,0.032 | 0.249,0.336 | -0.192 |
| 5828 | XLOC_046152 | chr9 | + | 140109268 | 140109654 | 140109268 | 140109470 | 140109546 | 76 | 140109654 | 5828 | 22,20 | 25,27 | 72,48 | 19,16 | 225 | 149 | 5.60E-07 | 8.80E-06 | 0.368,0.329 | 0.715,0.665 | -0.341 |
| 6884 | XLOC_021470 | chr19 | - | 3544559 | 3544937 | 3544559 | 3544730 | 3544806 | 76 | 3544937 | 6884 | 116,96 | 11,731,120 | 27,782,026 | 1,176,831 | 225 | 149 | 0 | 0 | 0.061,0.054 | 0.61,0.618 | -0.556 |
| 7694 | XLOC_018586 | chr17 | - | 43011600 | 43011981 | 43011600 | 43011798 | 43011874 | 76 | 43011981 | 7694 | 12,10 | 98,138 | 50,30 | 80,60 | 225 | 149 | 2.79E-12 | 1.11E-10 | 0.075,0.046 | 0.293,0.249 | -0.211 |
| 8468 | XLOC_016560 | chr16 | - | 772945 | 773161 | 772945 | 773018 | 773094 | 76 | 773161 | 8468 | 18,24 | 1,4 | 42,8 | 0,0 | 225 | 149 | 0.003476244 | 0.017305248 | 0.923,0.799 | 1.0,1.0 | -0.139 |
| 9888 | XLOC_021573 | chr19 | - | 13220582 | 13220811 | 13220582 | 13220652 | 13220728 | 76 | 13220811 | 9888 | 21,40 | 119,117 | 113,99 | 149,72 | 225 | 149 | 2.85E-08 | 5.73E-07 | 0.105,0.185 | 0.334,0.477 | -0.26 |
| 10048 | XLOC_016199 | chr16 | + | 1412470 | 1412743 | 1412470 | 1412535 | 1412611 | 76 | 1412743 | 10048 | 58,81 | 150,168 | 173,97 | 145,106 | 225 | 149 | 7.78E-08 | 1.46E-06 | 0.204,0.242 | 0.441,0.377 | -0.186 |
| 11888 | XLOC_009066 | chr12 | + | 104376918 | 104377167 | 104376918 | 104376996 | 104377072 | 76 | 104377167 | 11888 | 32,24 | 222,208 | 149,59 | 124,102 | 225 | 149 | 2.59E-13 | 1.19E-11 | 0.087,0.071 | 0.443,0.277 | -0.281 |
| 12619 | XLOC_027004 | chr20 | - | 43251469 | 43251788 | 43251469 | 43251571 | 43251647 | 76 | 43251788 | 12619 | 21,25 | 169,141 | 44,25 | 86,50 | 225 | 149 | 1.95E-07 | 3.35E-06 | 0.076,0.105 | 0.253,0.249 | -0.161 |
| 12787 | XLOC_016570 | chr16 | - | 1820885 | 1821172 | 1820885 | 1820994 | 1821070 | 76 | 1821172 | 12787 | 57,48 | 283,283 | 89,72 | 191,168 | 225 | 149 | 3.05E-07 | 5.07E-06 | 0.118,0.101 | 0.236,0.221 | -0.119 |
| 352 | XLOC_035165 | chr5 | - | 177651448 | 177651730 | 177651448 | 177651565 | 177651642 | 77 | 177651730 | 352 | 46,49 | 55,43 | 72,25 | 13,12 | 226 | 149 | 0.000198727 | 0.001543426 | 0.355,0.429 | 0.785,0.579 | -0.29 |
| 1060 | XLOC_021627 | chr19 | - | 19767468 | 19767958 | 19767468 | 19767725 | 19767802 | 77 | 19767958 | 1060 | 46,34 | 86,70 | 256,182 | 59,48 | 226 | 149 | 0 | 0 | 0.261,0.243 | 0.741,0.714 | -0.476 |
| 1707 | XLOC_014811 | chr15 | - | 28474605 | 28474993 | 28474605 | 28474742 | 28474819 | 77 | 28474993 | 1707 | 2,3 | 49,59 | 25,19 | 54,31 | 226 | 149 | 4.35E-10 | 1.20E-08 | 0.026,0.032 | 0.234,0.288 | -0.232 |

| | | | | | | | | | | | | | | | | | | | | | | |
|---|---|---|---|---|---|---|---|---|---|---|---|---|---|---|---|---|---|---|---|---|---|---|
| 1737 | XLOC_014574 | chr15 | + | 41106140 | 41106921 | 41106140 | 41106268 | 41106345 | 77 | 41106921 | 1737 | 116,149 | 115,111 | 146,136 | 10,12 | 226 | 149 | 0 | 0 | 0.399,0.469 | 0.906,0.882 | -0.46 |
| 2499 | XLOC_014807 | chr15 | - | 23326275 | 23326666 | 23326275 | 23326418 | 23326495 | 77 | 23326666 | 2499 | 2,0 | 23,23 | 9,12 | 13,19 | 226 | 149 | 3.16E-07 | 5.24E-06 | 0.054,0.0 | 0.313,0.294 | -0.276 |
| 3188 | XLOC_021440 | chr19 | - | 1011578 | 1011797 | 1011578 | 1011662 | 1011739 | 77 | 1011797 | 3188 | 66,39 | 362,311 | 114,87 | 261,227 | 226 | 149 | 3.38E-09 | 7.91E-08 | 0.107,0.076 | 0.224,0.202 | -0.122 |
| 3264 | XLOC_044437 | chr8 | - | 144657360 | 144657695 | 144657360 | 144657515 | 144657592 | 77 | 144657695 | 3264 | 90,83 | 435,389 | 159,123 | 301,208 | 226 | 149 | 6.13E-11 | 1.94E-09 | 0.12,0.123 | 0.258,0.281 | -0.148 |
| 3707 | XLOC_046491 | chr9 | - | 139934808 | 139935022 | 139934808 | 139934888 | 139934965 | 77 | 139935022 | 3707 | 28,27 | 255,286 | 50,39 | 135,97 | 226 | 149 | 2.69E-10 | 7.71E-09 | 0.068,0.059 | 0.196,0.21 | -0.14 |
| 3754 | XLOC_046457 | chr9 | - | 135772570 | 135772993 | 135772570 | 135772732 | 135772809 | 77 | 135772993 | 3754 | 16,3 | 61,57 | 19,29 | 37,32 | 226 | 149 | 1.59E-05 | 0.000169842 | 0.147,0.034 | 0.253,0.374 | -0.223 |
| 4618 | XLOC_037956 | chr6 | + | 111318186 | 111318467 | 111318186 | 111318308 | 111318385 | 77 | 111318467 | 4618 | 0,1 | 3,0 | 5,4 | 0,0 | 226 | 149 | 0.000552707 | 0.003751026 | 0.0,1.0 | 1.0,1.0 | -0.5 |
| 4931 | XLOC_046480 | chr9 | - | 139357763 | 139358192 | 139357763 | 139358053 | 139358130 | 77 | 139358192 | 4931 | 6,4 | 138,128 | 27,19 | 81,96 | 226 | 149 | 2.19E-09 | 5.34E-08 | 0.028,0.02 | 0.18,0.115 | -0.123 |
| 5002 | XLOC_044458 | chr8 | - | 145540684 | 145540914 | 145540684 | 145540772 | 145540849 | 77 | 145540914 | 5002 | 63,78 | 88,97 | 169,120 | 48,25 | 226 | 149 | 0 | 0 | 0.321,0.346 | 0.699,0.76 | -0.396 |
| 5385 | XLOC_028749 | chr22 | - | 50897683 | 50898087 | 50897683 | 50897821 | 50897898 | 77 | 50898087 | 5385 | 13,22 | 127,117 | 54,30 | 87,64 | 226 | 149 | 1.21E-08 | 2.55E-07 | 0.063,0.11 | 0.29,0.236 | -0.177 |
| 5388 | XLOC_028749 | chr22 | - | 50900224 | 50900708 | 50900224 | 50900298 | 50900375 | 77 | 50900708 | 5388 | 4,7 | 160,137 | 40,33 | 139,100 | 226 | 149 | 3.79E-13 | 1.70E-11 | 0.016,0.033 | 0.159,0.179 | -0.144 |
| 5437 | XLOC_032512 | chr4 | + | 185606565 | 185606838 | 185606565 | 185606652 | 185606729 | 77 | 185606838 | 5437 | 0,3 | 16,26 | 3,5 | 6,3 | 226 | 149 | 0.000519209 | 0.003550984 | 0.0,0.071 | 0.248,0.524 | -0.351 |
| 7191 | XLOC_016577 | chr16 | - | 2147148 | 2147429 | 2147148 | 2147242 | 2147319 | 77 | 2147429 | 7191 | 26,15 | 104,96 | 19,12 | 26,22 | 226 | 149 | 0.000256357 | 0.001934043 | 0.142,0.093 | 0.325,0.264 | -0.177 |
| 7212 | XLOC_006937 | chr11 | + | 212596 | 212981 | 212596 | 212759 | 212836 | 77 | 212981 | 7212 | 19,18 | 275,287 | 61,51 | 163,152 | 226 | 149 | 1.89E-15 | 1.13E-13 | 0.044,0.04 | 0.198,0.181 | -0.148 |
| 7262 | XLOC_020233 | chr18 | - | 47811293 | 47811721 | 47811293 | 47811617 | 47811694 | 77 | 47811721 | 7262 | 148,134 | 157,127 | 351,223 | 76,54 | 226 | 149 | 0 | 0 | 0.383,0.41 | 0.753,0.731 | -0.346 |
| 9361 | XLOC_018788 | chr17 | - | 79994419 | 79994605 | 79994419 | 79994519 | 79994596 | 77 | 79994605 | 9361 | 56,82 | 217,199 | 80,66 | 127,104 | 226 | 149 | 0.000269056 | 0.002014779 | 0.145,0.214 | 0.293,0.295 | -0.114 |
| 10171 | XLOC_021256 | chr19 | + | 45404464 | 45404709 | 45404464 | 45404567 | 45404644 | 77 | 45404709 | 10171 | 33,5 | 7,6 | 52,33 | 3,0 | 226 | 149 | 0.000141093 | 0.001148504 | 0.757,0.355 | 0.92,1.0 | -0.404 |
| 10273 | XLOC_011414 | chr13 | - | 25458035 | 25458383 | 25458035 | 25458221 | 25458298 | 77 | 25458383 | 10273 | 14,14 | 59,51 | 59,59 | 33,42 | 226 | 149 | 1.47E-12 | 6.21E-11 | 0.135,0.153 | 0.541,0.481 | -0.367 |
| 10499 | XLOC_000546 | chr1 | + | 155279579 | 155279996 | 155279579 | 155279756 | 155279833 | 77 | 155279996 | 10499 | 81,56 | 51,51 | 66,59 | 13,9 | 226 | 149 | 9.51E-08 | 1.75E-06 | 0.512,0.42 | 0.77,0.812 | -0.325 |
| 11447 | XLOC_030033 | chr3 | - | 129152692 | 129152976 | 129152692 | 129152827 | 129152904 | 77 | 129152976 | 11447 | 16,9 | 134,141 | 83,52 | 166,159 | 226 | 149 | 6.26E-11 | 1.98E-09 | 0.073,0.04 | 0.248,0.177 | -0.156 |
| 12320 | XLOC_007559 | chr11 | - | 63992014 | 63992410 | 63992014 | 63992189 | 63992266 | 77 | 63992410 | 12320 | 28,13 | 6,2 | 11,6 | 1,0 | 226 | 149 | 0.003368474 | 0.016863863 | 0.755,0.811 | 0.879,1.0 | -0.156 |
| 12444 | XLOC_018790 | chr17 | - | 80018576 | 80018830 | 80018576 | 80018659 | 80018736 | 77 | 80018830 | 12444 | 4,7 | 6,4 | 58,30 | 0,0 | 226 | 149 | 3.48E-10 | 9.76E-09 | 0.305,0.536 | 1.0,1.0 | -0.58 |
| 12478 | XLOC_034753 | chr5 | + | 149747251 | 149747480 | 149747251 | 149747329 | 149747406 | 77 | 149747480 | 12478 | 7,2 | 8,0 | 2,1 | 0,0 | 226 | 149 | 0.000655819 | 0.004319588 | 0.366,1.0 | 1.0,1.0 | -0.317 |
| 12703 | XLOC_016481 | chr16 | + | 74336109 | 74336301 | 74336109 | 74336190 | 74336267 | 77 | 74336301 | 12703 | 5,5 | 0,4 | 13,14 | 0,0 | 226 | 149 | 0.007490073 | 0.032509365 | 1.0,0.452 | 1.0,1.0 | -0.274 |
| 13346 | XLOC_004968 | chr10 | + | 94396914 | 94397302 | 94396914 | 94397066 | 94397143 | 77 | 94397302 | 13346 | 9,7 | 137,148 | 49,29 | 68,62 | 226 | 149 | 0 | 0 | 0.042,0.03 | 0.322,0.236 | -0.243 |
| 13361 | XLOC_021567 | chr19 | - | 12883426 | 12883853 | 12883426 | 12883516 | 12883593 | 77 | 12883853 | 13361 | 3,4 | 2,4 | 2,1 | 0,0 | 226 | 149 | 3.69E-06 | 4.67E-05 | 0.497,0.397 | 1.0,1.0 | -0.553 |
| 13648 | XLOC_018386 | chr17 | - | 7224878 | 7225301 | 7224878 | 7225106 | 7225183 | 77 | 7225301 | 13648 | 20,19 | 104,113 | 76,66 | 101,77 | 226 | 149 | 5.93E-12 | 2.20E-10 | 0.113,0.1 | 0.332,0.361 | -0.24 |
| 14299 | XLOC_016770 | chr16 | - | 67691408 | 67691717 | 67691408 | 67691587 | 67691664 | 77 | 67691717 | 14299 | 13,17 | 114,126 | 42,41 | 67,37 | 226 | 149 | 1.69E-11 | 5.86E-10 | 0.07,0.082 | 0.292,0.422 | -0.281 |
| 391 | XLOC_034801 | chr5 | + | 176083700 | 176086189 | 176083700 | 176083817 | 176083895 | 78 | 176086189 | 391 | 36,30 | 41,75 | 57,12 | 20,9 | 227 | 149 | 0.000562057 | 0.003794419 | 0.366,0.208 | 0.652,0.467 | -0.273 |
| 1180 | XLOC_020233 | chr18 | - | 47810074 | 47810471 | 47810074 | 47810185 | 47810263 | 78 | 47810471 | 1180 | 16,18 | 203,190 | 86,64 | 247,212 | 227 | 149 | 1.96E-10 | 5.74E-09 | 0.049,0.059 | 0.186,0.165 | -0.121 |
| 1467 | XLOC_016837 | chr16 | - | 84215351 | 84215664 | 84215351 | 84215469 | 84215547 | 78 | 84215664 | 1467 | 44,46 | 34,39 | 93,89 | 23,29 | 227 | 149 | 1.24E-05 | 0.000137119 | 0.459,0.436 | 0.726,0.668 | -0.25 |
| 1501 | XLOC_016559 | chr16 | - | 730588 | 733087 | 730588 | 732874 | 732952 | 78 | 733087 | 1501 | 18,17 | 128,154 | 114,89 | 88,72 | 227 | 149 | 0 | 0 | 0.085,0.068 | 0.46,0.448 | -0.378 |
| 2286 | XLOC_029430 | chr3 | + | 50154506 | 50154812 | 50154506 | 50154604 | 50154682 | 78 | 50154812 | 2286 | 26,43 | 186,232 | 153,103 | 229,194 | 227 | 149 | 1.21E-13 | 5.87E-12 | 0.084,0.108 | 0.305,0.258 | -0.185 |
| 3987 | XLOC_044443 | chr8 | - | 144921054 | 144921290 | 144921054 | 144921110 | 144921188 | 78 | 144921290 | 3987 | 35,30 | 93,87 | 21,41 | 18,27 | 227 | 149 | 3.02E-07 | 5.02E-06 | 0.198,0.185 | 0.434,0.499 | -0.275 |
| 5135 | XLOC_029841 | chr3 | - | 49154491 | 49155003 | 49154491 | 49154791 | 49154869 | 78 | 49155003 | 5135 | 3,2 | 8,8 | 12,8 | 0,0 | 227 | 149 | 7.74E-12 | 2.82E-10 | 0.198,0.141 | 1.0,1.0 | -0.831 |
| 5295 | XLOC_029456 | chr3 | + | 52727206 | 52727560 | 52727206 | 52727345 | 52727423 | 78 | 52727560 | 5295 | 95,94 | 493,561 | 572,408 | 518,416 | 227 | 149 | 0 | 0 | 0.112,0.099 | 0.42,0.392 | -0.3 |
| 5938 | XLOC_041630 | chr7 | - | 100279486 | 100279843 | 100279486 | 100279616 | 100279694 | 78 | 100279843 | 5938 | 41,42 | 234,207 | 86,68 | 147,125 | 227 | 149 | 1.63E-09 | 4.08E-08 | 0.103,0.118 | 0.277,0.263 | -0.16 |
| 8958 | XLOC_014944 | chr15 | - | 65113180 | 65113464 | 65113180 | 65113287 | 65113365 | 78 | 65113464 | 8958 | 19,24 | 32,48 | 34,29 | 17,28 | 227 | 149 | 0.003079622 | 0.015600251 | 0.28,0.247 | 0.568,0.405 | -0.223 |
| 9600 | XLOC_007211 | chr11 | + | 71940967 | 71941312 | 71940967 | 71941063 | 71941141 | 78 | 71941312 | 9600 | 4,6 | 0,5 | 7,6 | 0,0 | 227 | 149 | 0.001885622 | 0.010354058 | 1.0,0.441 | 1.0,1.0 | -0.279 |
| 10684 | XLOC_023711 | chr2 | + | 170663270 | 170663581 | 170663270 | 170663402 | 170663480 | 78 | 170663581 | 10684 | 49,55 | 474,396 | 228,142 | 453,311 | 227 | 149 | 0 | 0 | 0.064,0.084 | 0.248,0.231 | -0.165 |
| 10703 | XLOC_021207 | chr19 | + | 39905628 | 39906079 | 39905628 | 39905900 | 39905978 | 78 | 39906079 | 10703 | 0,0 | 26,19 | 3,5 | 24,15 | 227 | 149 | 0.000319256 | 0.002341041 | 0.0,0.0 | 0.076,0.18 | -0.128 |
| 10982 | XLOC_029823 | chr3 | - | 48669195 | 48669477 | 48669195 | 48669234 | 48669312 | 78 | 48669477 | 10982 | 7,5 | 46,62 | 20,14 | 33,23 | 227 | 149 | 1.91E-06 | 2.61E-05 | 0.091,0.05 | 0.285,0.285 | -0.214 |
| 11030 | XLOC_004857 | chr10 | + | 70101383 | 70101608 | 70101383 | 70101437 | 70101515 | 78 | 70101608 | 11030 | 59,59 | 914,963 | 218,229 | 793,561 | 227 | 149 | 0 | 0 | 0.041,0.039 | 0.153,0.211 | -0.142 |
| 14799 | XLOC_000821 | chr1 | - | 934912 | 935167 | 934912 | 934993 | 935071 | 78 | 935167 | 14799 | 207,233 | 89,100 | 659,504 | 113,85 | 227 | 149 | 1.58E-09 | 3.98E-08 | 0.604,0.605 | 0.793,0.796 | -0.19 |
| 124 | XLOC_044167 | chr8 | + | 144461484 | 144462953 | 144461484 | 144462155 | 144462234 | 79 | 144462953 | 124 | 52,60 | 71,38 | 36,24 | 6,1 | 228 | 149 | 7.85E-07 | 1.19E-05 | 0.324,0.508 | 0.797,0.94 | -0.453 |
| 928 | XLOC_023754 | chr2 | + | 182783324 | 182783609 | 182783324 | 182783402 | 182783481 | 79 | 182783609 | 928 | 11,37 | 179,176 | 147,108 | 241,183 | 228 | 149 | 1.74E-14 | 9.25E-13 | 0.039,0.121 | 0.285,0.278 | -0.201 |
| 1673 | XLOC_021565 | chr19 | - | 12812411 | 12812665 | 12812411 | 12812486 | 12812565 | 79 | 12812665 | 1673 | 19,25 | 85,133 | 59,33 | 85,59 | 228 | 149 | 5.17E-07 | 8.17E-06 | 0.127,0.109 | 0.312,0.268 | -0.172 |
| 1757 | XLOC_000897 | chr1 | - | 16269893 | 16270210 | 16269893 | 16270014 | 16270093 | 79 | 16270210 | 1757 | 2,0 | 43,27 | 4,8 | 21,34 | 228 | 149 | 0.000296176 | 0.002191143 | 0.029,0.0 | 0.111,0.133 | -0.107 |
| 2501 | XLOC_007282 | chr11 | + | 108098321 | 108098615 | 108098321 | 108098423 | 108098502 | 79 | 108098615 | 2501 | 8,8 | 24,39 | 30,14 | 44,15 | 228 | 149 | 0.00121806 | 0.007203797 | 0.179,0.118 | 0.308,0.379 | -0.195 |
| 2605 | XLOC_029893 | chr3 | - | 56694758 | 56695045 | 56694758 | 56694862 | 56694941 | 79 | 56695045 | 2605 | 0,0 | 117,98 | 18,16 | 83,70 | 228 | 149 | 1.90E-12 | 7.74E-11 | 0.0,0.0 | 0.124,0.13 | -0.127 |
| 2889 | XLOC_016217 | chr16 | + | 2284859 | 2285144 | 2284859 | 2285002 | 2285081 | 79 | 2285144 | 2889 | 32,31 | 169,182 | 162,83 | 122,104 | 228 | 149 | 7.77E-16 | 4.73E-14 | 0.11,0.1 | 0.465,0.343 | -0.299 |
| 3799 | XLOC_046241 | chr9 | - | 35738033 | 35738266 | 35738033 | 35738149 | 35738228 | 79 | 35738266 | 3799 | 61,50 | 73,56 | 157,100 | 51,36 | 228 | 149 | 1.03E-09 | 2.67E-08 | 0.353,0.368 | 0.668,0.645 | -0.296 |
| 5467 | XLOC_028629 | chr22 | - | 30690784 | 30691040 | 30690784 | 30690850 | 30690929 | 79 | 30691040 | 5467 | 0,0 | 30,27 | 1,4 | 18,7 | 228 | 149 | 0.009436308 | 0.039335589 | 0.0,0.0 | 0.035,0.272 | -0.154 |
| 5914 | XLOC_044184 | chr8 | + | 145693063 | 145693370 | 145693063 | 145693162 | 145693241 | 79 | 145693370 | 5914 | 11,22 | 126,91 | 27,8 | 38,40 | 228 | 149 | 0.007352658 | 0.032020223 | 0.054,0.136 | 0.317,0.116 | -0.121 |
| 10044 | XLOC_016199 | chr16 | + | 1402102 | 1402414 | 1402102 | 1402160 | 1402239 | 79 | 1402414 | 10044 | 40,63 | 201,209 | 158,119 | 165,136 | 228 | 149 | 5.66E-15 | 3.22E-13 | 0.115,0.165 | 0.385,0.364 | -0.234 |
| 10374 | XLOC_009601 | chr12 | - | 123812253 | 123812570 | 123812253 | 123812359 | 123812438 | 79 | 123812570 | 10374 | 0,2 | 138,115 | 30,26 | 111,101 | 228 | 149 | 1.11E-15 | 6.67E-14 | 0.0,0.011 | 0.15,0.144 | -0.141 |
| 10477 | XLOC_016348 | chr16 | + | 29813828 | 29814258 | 29813828 | 29814010 | 29814089 | 79 | 29814258 | 10477 | 14,20 | 5,7 | 39,20 | 0,1 | 228 | 149 | 1.50E-06 | 2.11E-05 | 0.647,0.651 | 1.0,0.929 | -0.316 |
| 10825 | XLOC_009411 | chr12 | - | 56742707 | 56743045 | 56742707 | 56742867 | 56742946 | 79 | 56743045 | 10825 | 7,8 | 4,0 | 10,3 | 0,0 | 228 | 149 | 0.009180486 | 0.038473592 | 0.534,1.0 | 1.0,1.0 | -0.233 |
| 10952 | XLOC_009247 | chr12 | - | 9571370 | 9571781 | 9571370 | 9571525 | 9571604 | 79 | 9571781 | 10952 | 9,18 | 13,17 | 26,31 | 12,10 | 228 | 149 | 0.004549428 | 0.021474457 | 0.311,0.409 | 0.586,0.67 | -0.268 |
| 11396 | XLOC_021828 | chr19 | - | 50365300 | 50365494 | 50365300 | 50365362 | 50365441 | 79 | 50365494 | 11396 | 15,22 | 160,112 | 48,21 | 77,78 | 228 | 149 | 4.15E-05 | 0.000400934 | 0.058,0.114 | 0.289,0.15 | -0.133 |
| 11458 | XLOC_000241 | chr1 | + | 41474331 | 41474562 | 41474331 | 41474386 | 41474465 | 79 | 41474562 | 11458 | 216,188 | 376,378 | 658,402 | 231,220 | 228 | 149 | 0 | 0 | 0.273,0.245 | 0.651,0.544 | -0.339 |
| 12604 | XLOC_021144 | chr19 | + | 36222808 | 36223230 | 36222808 | 36223036 | 36223115 | 79 | 36223230 | 12604 | 2,0 | 77,92 | 12,24 | 94,88 | 228 | 149 | 5.07E-09 | 1.15E-07 | 0.017,0.0 | 0.077,0.151 | -0.105 |
| 12901 | XLOC_005025 | chr10 | + | 104159026 | 104159475 | 104159026 | 104159254 | 104159333 | 79 | 104159475 | 12901 | 29,17 | 327,296 | 122,109 | 342,364 | 228 | 149 | 4.44E-16 | 2.74E-14 | 0.055,0.036 | 0.189,0.164 | -0.131 |
| 13179 | XLOC_044455 | chr8 | - | 145154179 | 145154514 | 145154179 | 145154333 | 145154412 | 79 | 145154514 | 13179 | 19,22 | 137,107 | 78,51 | 43,52 | 228 | 149 | 3.11E-14 | 1.56E-12 | 0.083,0.118 | 0.542,0.391 | -0.366 |
| 13614 | XLOC_001082 | chr1 | - | 45797615 | 45797982 | 45797615 | 45797758 | 45797837 | 79 | 45797982 | 13614 | 4,5 | 41,40 | 47,12 | 22,19 | 228 | 149 | 5.02E-07 | 7.96E-06 | 0.06,0.076 | 0.583,0.292 | -0.369 |

| | | | | | | | | | | | | | | | | | | | | | | |
|---|---|---|---|---|---|---|---|---|---|---|---|---|---|---|---|---|---|---|---|---|---|---|
| 13835 | XLOC_006953 | chr11 | + | 828574 | 828963 | 828574 | 828772 | 828851 | 79 | 828963 | 13835 | 0,1 | 20,25 | 11,1 | 9,5 | 228 | 149 | 1.04E-05 | 0.000117584 | 0.0,0.025 | 0.444,0.116 | -0.268 |
| 15002 | XLOC_007183 | chr11 | + | 67074312 | 67074584 | 67074312 | 67074433 | 67074512 | 79 | 67074584 | 15002 | 86,87 | 72,59 | 110,66 | 45,14 | 228 | 149 | 0.000626759 | 0.004155739 | 0.438,0.491 | 0.615,0.755 | -0.221 |
| 232 | XLOC_041771 | chr7 | - | 150773608 | 150774114 | 150773608 | 150773919 | 150773999 | 80 | 150774114 | 232 | 31,45 | 260,211 | 108,81 | 137,111 | 229 | 149 | 0 | 0 | 0.072,0.122 | 0.339,0.322 | -0.234 |
| 418 | XLOC_034648 | chr5 | + | 112889286 | 112889551 | 112889286 | 112889383 | 112889463 | 80 | 112889551 | 418 | 0,7 | 51,43 | 44,8 | 33,28 | 229 | 149 | 1.18E-05 | 0.000132002 | 0.0,0.096 | 0.465,0.157 | -0.263 |
| 553 | XLOC_030002 | chr3 | - | 123674689 | 123674962 | 123674689 | 123674812 | 123674892 | 80 | 123674962 | 553 | 79,86 | 46,69 | 72,42 | 16,19 | 229 | 149 | 0.006569211 | 0.029193506 | 0.528,0.448 | 0.745,0.59 | -0.179 |
| 1058 | XLOC_021627 | chr19 | - | 19767468 | 19767725 | 19767468 | 19767575 | 19767655 | 80 | 19767725 | 1058 | 109,78 | 66,55 | 251,213 | 41,18 | 229 | 149 | 5.88E-15 | 3.32E-13 | 0.518,0.48 | 0.799,0.885 | -0.343 |
| 1524 | XLOC_016440 | chr16 | + | 67320651 | 67321080 | 67320651 | 67320822 | 67320902 | 80 | 67321080 | 1524 | 67,44 | 73,74 | 60,45 | 38,36 | 229 | 149 | 0.006152273 | 0.027636309 | 0.374,0.279 | 0.507,0.449 | -0.151 |
| 3724 | XLOC_046490 | chr9 | - | 139905424 | 139905752 | 139905424 | 139905557 | 139905637 | 80 | 139905752 | 3724 | 43,65 | 211,175 | 62,50 | 99,55 | 229 | 149 | 3.24E-06 | 4.17E-05 | 0.117,0.195 | 0.29,0.372 | -0.175 |
| 4238 | XLOC_041495 | chr7 | - | 44609407 | 44609832 | 44609407 | 44609502 | 44609582 | 80 | 44609832 | 4238 | 11,13 | 11,8 | 31,25 | 6,8 | 229 | 149 | 0.008921454 | 0.03760003 | 0.394,0.514 | 0.771,0.67 | -0.267 |
| 4821 | XLOC_034488 | chr5 | + | 5447541 | 5448010 | 5447541 | 5447873 | 5447953 | 80 | 5448010 | 4821 | 4,4 | 164,183 | 117,90 | 273,170 | 229 | 149 | 0 | 0 | 0.016,0.014 | 0.218,0.256 | -0.222 |
| 6212 | XLOC_024159 | chr2 | - | 99787801 | 99788109 | 99787801 | 99787892 | 99787972 | 80 | 99788109 | 6212 | 23,13 | 209,175 | 109,62 | 169,106 | 229 | 149 | 0 | 0 | 0.067,0.046 | 0.296,0.276 | -0.23 |
| 8182 | XLOC_018607 | chr17 | - | 45915701 | 45916062 | 45915701 | 45915816 | 45915896 | 80 | 45916062 | 8182 | 44,31 | 151,171 | 35,17 | 51,39 | 229 | 149 | 0.000426097 | 0.002995402 | 0.159,0.106 | 0.309,0.221 | -0.133 |
| 8568 | XLOC_016435 | chr16 | + | 67201022 | 67201305 | 67201022 | 67201125 | 67201205 | 80 | 67201305 | 8568 | 24,14 | 13,9 | 31,30 | 1,5 | 229 | 149 | 0.000123081 | 0.001024321 | 0.546,0.503 | 0.953,0.796 | -0.35 |
| 9100 | XLOC_021565 | chr19 | - | 12825858 | 12826352 | 12825858 | 12826138 | 12826218 | 80 | 12826352 | 9100 | 2,7 | 108,124 | 32,30 | 93,85 | 229 | 149 | 2.47E-12 | 9.88E-11 | 0.012,0.035 | 0.183,0.187 | -0.162 |
| 9302 | XLOC_009456 | chr12 | - | 72013705 | 72014095 | 72013705 | 72013909 | 72013989 | 80 | 72014095 | 9302 | 2,3 | 0,4 | 31,12 | 0,0 | 229 | 149 | 0.006528443 | 0.029067213 | 1.0,0.328 | 1.0,1.0 | -0.336 |
| 10215 | XLOC_007565 | chr11 | - | 64564453 | 64564640 | 64564453 | 64564495 | 64564575 | 80 | 64564640 | 10215 | 23,23 | 82,98 | 23,14 | 39,24 | 229 | 149 | 0.004262669 | 0.020339124 | 0.154,0.132 | 0.277,0.275 | -0.133 |
| 11523 | XLOC_000780 | chr1 | + | 231409667 | 231410126 | 231409667 | 231409808 | 231409888 | 80 | 231410126 | 11523 | 29,18 | 5,2 | 31,36 | 0,0 | 229 | 149 | 0.000148157 | 0.00119743 | 0.791,0.854 | 1.0,1.0 | -0.177 |
| 12956 | XLOC_004959 | chr10 | + | 91477131 | 91477479 | 91477131 | 91477252 | 91477332 | 80 | 91477479 | 12956 | 2,0 | 72,91 | 26,11 | 58,56 | 229 | 149 | 2.03E-11 | 6.98E-10 | 0.018,0.0 | 0.226,0.113 | -0.161 |
| 13615 | XLOC_001082 | chr1 | - | 45797849 | 45798160 | 45797849 | 45797982 | 45798062 | 80 | 45798160 | 13615 | 5,8 | 53,49 | 23,12 | 14,19 | 229 | 149 | 1.45E-06 | 2.04E-05 | 0.058,0.096 | 0.517,0.291 | -0.327 |
| 15225 | XLOC_048116 | chrX | - | 153052256 | 153052608 | 153052256 | 153052425 | 153052505 | 80 | 153052608 | 15225 | 0,2 | 4,5 | 5,1 | 1,0 | 229 | 149 | 4.76E-06 | 5.90E-05 | 0.0,0.207 | 0.765,1.0 | -0.779 |
| 492 | XLOC_032442 | chr4 | + | 128804416 | 128804708 | 128804416 | 128804512 | 128804593 | 81 | 128804708 | 492 | 3,5 | 111,111 | 13,10 | 71,28 | 230 | 149 | 3.32E-06 | 4.26E-05 | 0.017,0.028 | 0.106,0.188 | -0.124 |
| 565 | XLOC_029841 | chr3 | - | 49147616 | 49148037 | 49147616 | 49147795 | 49147876 | 81 | 49148037 | 565 | 15,9 | 128,135 | 20,17 | 63,51 | 230 | 149 | 2.09E-05 | 0.00021658 | 0.071,0.041 | 0.171,0.178 | -0.118 |
| 1329 | XLOC_000536 | chr1 | + | 154962634 | 154963004 | 154962634 | 154962733 | 154962814 | 81 | 154963004 | 1329 | 16,23 | 45,57 | 44,37 | 34,35 | 230 | 149 | 1.54E-05 | 0.00016517 | 0.187,0.207 | 0.456,0.406 | -0.234 |
| 1562 | XLOC_016204 | chr16 | + | 1825797 | 1826385 | 1825797 | 1825987 | 1826068 | 81 | 1826385 | 1562 | 0,1 | 18,22 | 7,6 | 17,10 | 230 | 149 | 4.46E-06 | 5.58E-05 | 0.0,0.029 | 0.211,0.28 | -0.231 |
| 1780 | XLOC_000300 | chr1 | + | 53343353 | 53344440 | 53343353 | 53343515 | 53343596 | 81 | 53344440 | 1780 | 14,16 | 36,37 | 47,63 | 36,19 | 230 | 149 | 9.90E-06 | 0.000112877 | 0.201,0.219 | 0.458,0.682 | -0.36 |
| 1854 | XLOC_013342 | chr14 | - | 31361332 | 31364954 | 31361332 | 31364786 | 31364867 | 81 | 31364954 | 1854 | 0,0 | 55,56 | 11,9 | 65,39 | 230 | 149 | 2.21E-07 | 3.77E-06 | 0.0,0.0 | 0.099,0.13 | -0.115 |
| 1861 | XLOC_013325 | chr14 | - | 24032588 | 24032847 | 24032588 | 24032711 | 24032792 | 81 | 24032847 | 1861 | 31,45 | 49,42 | 67,32 | 29,20 | 230 | 149 | 0.001916613 | 0.010483472 | 0.291,0.41 | 0.599,0.509 | -0.204 |
| 1989 | XLOC_007440 | chr11 | - | 17156365 | 17156706 | 17156365 | 17156576 | 17156657 | 81 | 17156706 | 1989 | 12,16 | 105,120 | 106,81 | 99,97 | 230 | 149 | 0 | 0 | 0.069,0.08 | 0.41,0.351 | -0.306 |
| 2243 | XLOC_009165 | chr12 | + | 123467015 | 123467744 | 123467015 | 123467085 | 123467166 | 81 | 123467744 | 2243 | 121,123 | 803,782 | 209,163 | 490,437 | 230 | 149 | 1.76E-12 | 7.34E-11 | 0.089,0.092 | 0.216,0.195 | -0.115 |
| 3277 | XLOC_000803 | chr1 | + | 242012032 | 242012518 | 242012032 | 242012329 | 242012410 | 81 | 242012518 | 3277 | 21,42 | 92,120 | 82,58 | 72,83 | 230 | 149 | 1.27E-06 | 1.82E-05 | 0.129,0.185 | 0.425,0.312 | -0.211 |
| 3489 | XLOC_048139 | chrX | - | 154293760 | 154294064 | 154293760 | 154293812 | 154293893 | 81 | 154294064 | 3489 | 0,0 | 24,12 | 5,2 | 3,1 | 230 | 149 | 2.20E-07 | 3.75E-06 | 0.0,0.0 | 0.519,0.564 | -0.541 |
| 3784 | XLOC_046345 | chr9 | - | 100368424 | 100368758 | 100368424 | 100368543 | 100368624 | 81 | 100368758 | 3784 | 14,14 | 1,5 | 10,13 | 1,0 | 230 | 149 | 0.003577287 | 0.017720806 | 0.901,0.645 | 0.866,1.0 | -0.16 |
| 3952 | XLOC_044466 | chr8 | - | 145739311 | 145739614 | 145739311 | 145739491 | 145739572 | 81 | 145739614 | 3952 | 5,10 | 173,161 | 31,63 | 124,88 | 230 | 149 | 3.34E-12 | 1.29E-10 | 0.018,0.039 | 0.139,0.317 | -0.2 |
| 4144 | XLOC_045898 | chr9 | + | 35660561 | 35661014 | 35660561 | 35660855 | 35660936 | 81 | 35661014 | 4144 | 250,186 | 69,53 | 402,257 | 52,31 | 230 | 149 | 1.20E-05 | 0.000133018 | 0.701,0.695 | 0.834,0.843 | -0.141 |
| 5383 | XLOC_028749 | chr22 | - | 50893444 | 50893673 | 50893444 | 50893571 | 50893652 | 81 | 50893673 | 5383 | 17,15 | 171,144 | 76,31 | 142,90 | 230 | 149 | 1.07E-10 | 3.24E-09 | 0.061,0.063 | 0.257,0.182 | -0.158 |
| 7589 | XLOC_018725 | chr17 | - | 73888095 | 73888581 | 73888095 | 73888266 | 73888347 | 81 | 73888581 | 7589 | 84,70 | 121,98 | 99,78 | 37,19 | 230 | 149 | 3.20E-12 | 1.25E-10 | 0.31,0.316 | 0.634,0.727 | -0.367 |
| 8393 | XLOC_016188 | chr16 | + | 721720 | 722181 | 721720 | 722002 | 722083 | 81 | 722181 | 8393 | 15,25 | 412,353 | 103,73 | 198,177 | 230 | 149 | 0 | 0 | 0.023,0.044 | 0.252,0.211 | -0.198 |
| 8420 | XLOC_020860 | chr19 | + | 1063542 | 1063862 | 1063542 | 1063677 | 1063758 | 81 | 1063862 | 8420 | 3,0 | 12,8 | 9,3 | 8,3 | 230 | 149 | 0.000830112 | 0.005216938 | 0.139,0.0 | 0.422,0.393 | -0.338 |
| 9740 | XLOC_013303 | chr14 | - | 21868333 | 21868771 | 21868333 | 21868490 | 21868571 | 81 | 21868771 | 9740 | 0,0 | 2,4 | 2,2 | 6,4 | 230 | 149 | 0.006473117 | 0.028839065 | 0.0,0.0 | 0.178,0.245 | -0.211 |
| 10706 | XLOC_032606 | chr4 | - | 48140705 | 48141039 | 48140705 | 48140839 | 48140920 | 81 | 48141039 | 10706 | 0,7 | 37,22 | 26,15 | 22,16 | 230 | 149 | 3.45E-07 | 5.67E-06 | 0.0,0.171 | 0.434,0.378 | -0.321 |
| 12823 | XLOC_040996 | chr7 | + | 4830019 | 4830518 | 4830019 | 4830222 | 4830303 | 81 | 4830518 | 12823 | 12,29 | 55,73 | 48,29 | 47,37 | 230 | 149 | 4.04E-05 | 0.000391858 | 0.124,0.205 | 0.398,0.337 | -0.203 |
| 13782 | XLOC_029849 | chr3 | - | 49721982 | 49722388 | 49721982 | 49722089 | 49722170 | 81 | 49722388 | 13782 | 2,6 | 19,8 | 4,7 | 2,3 | 230 | 149 | 0.002981855 | 0.015175761 | 0.064,0.327 | 0.564,0.602 | -0.387 |
| 13786 | XLOC_029849 | chr3 | - | 49724383 | 49724718 | 49724383 | 49724500 | 49724581 | 81 | 49724718 | 13786 | 43,53 | 11,3 | 130,66 | 2,1 | 230 | 149 | 5.51E-05 | 0.000511938 | 0.717,0.92 | 0.977,0.977 | -0.158 |
| 1453 | XLOC_023617 | chr2 | + | 109367719 | 109368159 | 109367719 | 109367901 | 109367983 | 82 | 109368159 | 1453 | 6,14 | 181,158 | 57,56 | 127,110 | 231 | 149 | 0 | 0 | 0.021,0.054 | 0.225,0.247 | -0.198 |
| 1774 | XLOC_018149 | chr17 | + | 48452545 | 48453554 | 48452545 | 48453344 | 48453426 | 82 | 48453554 | 1774 | 15,17 | 55,96 | 63,21 | 35,37 | 231 | 149 | 4.80E-05 | 0.000454765 | 0.15,0.103 | 0.537,0.268 | -0.276 |
| 2656 | XLOC_011393 | chr13 | + | 115037658 | 115038417 | 115037658 | 115037956 | 115038038 | 82 | 115038417 | 2656 | 240,189 | 144,141 | 289,224 | 41,29 | 231 | 149 | 0 | 0 | 0.518,0.464 | 0.82,0.833 | -0.336 |
| 3156 | XLOC_023563 | chr2 | + | 86716440 | 86716782 | 86716440 | 86716560 | 86716642 | 82 | 86716782 | 3156 | 11,8 | 220,220 | 45,38 | 130,141 | 231 | 149 | 8.66E-15 | 4.82E-13 | 0.031,0.023 | 0.183,0.148 | -0.138 |
| 3961 | XLOC_044462 | chr8 | - | 145649434 | 145649651 | 145649434 | 145649515 | 145649597 | 82 | 145649651 | 3961 | 52,66 | 315,283 | 134,113 | 225,179 | 231 | 149 | 1.79E-12 | 7.42E-11 | 0.096,0.131 | 0.278,0.289 | -0.17 |
| 5511 | XLOC_028552 | chr22 | + | 50960771 | 50961146 | 50960771 | 50960844 | 50960926 | 82 | 50961146 | 5511 | 3,4 | 9,29 | 9,13 | 12,13 | 231 | 149 | 0.000733478 | 0.004700051 | 0.177,0.082 | 0.326,0.392 | -0.229 |
| 6617 | XLOC_001199 | chr1 | - | 93804633 | 93805093 | 93804633 | 93804831 | 93804913 | 82 | 93805093 | 6617 | 8,11 | 35,29 | 13,9 | 3,8 | 231 | 149 | 0.00045802 | 0.003195954 | 0.128,0.197 | 0.737,0.421 | -0.416 |
| 8356 | XLOC_016837 | chr16 | - | 84215197 | 84215676 | 84215197 | 84215269 | 84215351 | 82 | 84215676 | 8356 | 17,15 | 95,68 | 54,40 | 66,45 | 231 | 149 | 2.55E-09 | 6.15E-08 | 0.103,0.125 | 0.345,0.364 | -0.24 |
| 9096 | XLOC_021565 | chr19 | - | 12816305 | 12816644 | 12816305 | 12816413 | 12816495 | 82 | 12816644 | 9096 | 11,14 | 115,153 | 100,59 | 110,72 | 231 | 149 | 0 | 0 | 0.058,0.056 | 0.37,0.346 | -0.301 |
| 10047 | XLOC_016199 | chr16 | + | 1411686 | 1412132 | 1411686 | 1411956 | 1412038 | 82 | 1412132 | 10047 | 20,10 | 241,226 | 66,50 | 248,163 | 231 | 149 | 1.23E-11 | 4.34E-10 | 0.051,0.028 | 0.147,0.165 | -0.116 |
| 11212 | XLOC_046145 | chr9 | + | 139750708 | 139751027 | 139750708 | 139750833 | 139750915 | 82 | 139751027 | 11212 | 10,21 | 22,3 | 16,16 | 3,0 | 231 | 149 | 0.011456182 | 0.046466302 | 0.227,0.819 | 0.775,1.0 | -0.364 |
| 11691 | XLOC_007557 | chr11 | - | 63766309 | 63766472 | 63766309 | 63766344 | 63766426 | 82 | 63766472 | 11691 | 89,82 | 543,484 | 316,232 | 619,468 | 231 | 149 | 8.88E-16 | 5.36E-14 | 0.096,0.099 | 0.248,0.242 | -0.147 |
| 13028 | XLOC_045938 | chr9 | + | 72965033 | 72965387 | 72965033 | 72965209 | 72965291 | 82 | 72965387 | 13028 | 9,6 | 111,112 | 39,22 | 53,48 | 231 | 149 | 5.25E-13 | 2.30E-11 | 0.05,0.033 | 0.322,0.228 | -0.234 |
| 14294 | XLOC_016688 | chr16 | - | 30004779 | 30005119 | 30004779 | 30004878 | 30004960 | 82 | 30005119 | 14294 | 19,20 | 146,144 | 24,13 | 30,42 | 231 | 149 | 9.13E-05 | 0.000793224 | 0.077,0.082 | 0.34,0.166 | -0.173 |
| 14489 | XLOC_005187 | chr10 | - | 32575853 | 32576198 | 32575853 | 32575943 | 32576025 | 82 | 32576198 | 14489 | 1,1 | 29,42 | 21,22 | 42,28 | 231 | 149 | 7.50E-11 | 2.33E-09 | 0.022,0.015 | 0.244,0.336 | -0.272 |
| 15077 | XLOC_000399 | chr1 | + | 109238664 | 109238973 | 109238664 | 109238816 | 109238898 | 82 | 109238973 | 15077 | 6,5 | 280,230 | 96,87 | 247,229 | 231 | 149 | 0 | 0 | 0.014,0.014 | 0.2,0.197 | -0.184 |
| 425 | XLOC_034609 | chr5 | + | 81600433 | 81601300 | 81600433 | 81600669 | 81600752 | 83 | 81601300 | 425 | 7,3 | 3,10 | 12,4 | 2,1 | 232 | 149 | 0.010232552 | 0.042120177 | 0.6,0.162 | 0.794,0.72 | -0.376 |
| 1425 | XLOC_029858 | chr3 | - | 50326877 | 50327193 | 50326877 | 50327059 | 50327142 | 83 | 50327193 | 1425 | 20,28 | 101,137 | 116,92 | 198,147 | 232 | 149 | 2.26E-08 | 4.61E-07 | 0.113,0.116 | 0.273,0.287 | -0.166 |
| 2370 | XLOC_026876 | chr20 | + | 62321439 | 62321794 | 62321439 | 62321563 | 62321646 | 83 | 62321794 | 2370 | 3,8 | 62,47 | 12,9 | 51,22 | 232 | 149 | 0.007850549 | 0.033844109 | 0.03,0.099 | 0.131,0.208 | -0.105 |
| 2448 | XLOC_007549 | chr11 | - | 62592754 | 62593011 | 62592754 | 62592811 | 62592894 | 83 | 62593011 | 2448 | 2,2 | 117,112 | 58,39 | 101,70 | 232 | 149 | 0 | 0 | 0.011,0.011 | 0.269,0.264 | -0.256 |
| 3953 | XLOC_044466 | chr8 | - | 145740533 | 145740959 | 145740533 | 145740626 | 145740709 | 83 | 145740959 | 3953 | 4,6 | 187,207 | 115,53 | 159,74 | 232 | 149 | 0 | 0 | 0.014,0.018 | 0.317,0.315 | -0.3 |
| 4135 | XLOC_047697 | chrX | + | 48339674 | 48339916 | 48339674 | 48339730 | 48339813 | 83 | 48339916 | 4135 | 11,12 | 245,251 | 85,46 | 189,153 | 232 | 149 | 0 | 0 | 0.028,0.03 | 0.224,0.162 | -0.164 |

| | | | | | | | | | | | | | | | | | | | | | | |
|---|---|---|---|---|---|---|---|---|---|---|---|---|---|---|---|---|---|---|---|---|---|---|
| 4435 | XLOC_038498 | chr6 | - | 165831840 | 165832217 | 165831840 | 165832039 | 165832122 | 83 | 165832217 | 4435 | 0,0 | 4,12 | 7,5 | 4,3 | 232 | 149 | 4.57E-08 | 8.87E-07 | 0.0,0.0 | 0.529,0.517 | -0.523 |
| 5857 | XLOC_026790 | chr20 | + | 39795115 | 39795591 | 39795115 | 39795235 | 39795318 | 83 | 39795591 | 5857 | 2,1 | 85,78 | 3,8 | 37,24 | 232 | 149 | 0.00010719 | 0.000909766 | 0.015,0.008 | 0.049,0.176 | -0.101 |
| 6203 | XLOC_024212 | chr2 | - | 131829427 | 131829827 | 131829427 | 131829511 | 131829594 | 83 | 131829827 | 6203 | 1,1 | 5,8 | 16,3 | 6,9 | 232 | 149 | 0.00463307 | 0.021796416 | 0.114,0.074 | 0.631,0.176 | -0.309 |
| 6326 | XLOC_000123 | chr1 | + | 23408719 | 23409037 | 23408719 | 23408859 | 23408942 | 83 | 23409037 | 6326 | 28,49 | 19,14 | 100,63 | 7,7 | 232 | 149 | 4.43E-05 | 0.000424494 | 0.486,0.692 | 0.902,0.853 | -0.288 |
| 8036 | XLOC_017921 | chr17 | + | 8347244 | 8347675 | 8347244 | 8347494 | 8347577 | 83 | 8347675 | 8036 | 33,52 | 13,9 | 19,12 | 0,0 | 232 | 149 | 3.06E-06 | 3.96E-05 | 0.62,0.788 | 1.0,1.0 | -0.296 |
| 9749 | XLOC_013277 | chr14 | + | 105174868 | 105175069 | 105174868 | 105174924 | 105175007 | 83 | 105175069 | 9749 | 27,35 | 285,244 | 91,56 | 197,136 | 232 | 149 | 2.49E-12 | 9.93E-11 | 0.057,0.084 | 0.229,0.209 | -0.148 |
| 11732 | XLOC_007156 | chr11 | + | 65403644 | 65404143 | 65403644 | 65403778 | 65403861 | 83 | 65404143 | 11732 | 42,49 | 137,111 | 189,157 | 107,103 | 232 | 149 | 0 | 0 | 0.165,0.221 | 0.531,0.495 | -0.32 |
| 11796 | XLOC_007519 | chr11 | - | 57472009 | 57472482 | 57472009 | 57472107 | 57472190 | 83 | 57472482 | 11796 | 2,0 | 7,1 | 1,3 | 0,0 | 232 | 149 | 2.34E-10 | 6.77E-09 | 0.155,0.0 | 1.0,1.0 | -0.922 |
| 11860 | XLOC_007421 | chr11 | - | 9860836 | 9861243 | 9860836 | 9860961 | 9861044 | 83 | 9861243 | 11860 | 4,1 | 2,5 | 13,7 | 0,1 | 232 | 149 | 1.57E-05 | 0.000167638 | 0.562,0.114 | 1.0,0.818 | -0.571 |
| 12506 | XLOC_016216 | chr16 | + | 2256497 | 2256930 | 2256497 | 2256660 | 2256743 | 83 | 2256930 | 12506 | 0,0 | 4,1 | 4,2 | 3,1 | 232 | 149 | 1.25E-05 | 0.000138628 | 0.0,0.0 | 0.461,0.562 | -0.512 |
| 12902 | XLOC_005025 | chr10 | + | 104159836 | 104160094 | 104159836 | 104159951 | 104160034 | 83 | 104160094 | 12902 | 19,25 | 223,243 | 145,101 | 325,242 | 232 | 149 | 3.33E-16 | 2.11E-14 | 0.052,0.062 | 0.223,0.211 | -0.16 |
| 3932 | XLOC_046164 | chr9 | + | 140508432 | 140508927 | 140508432 | 140508666 | 140508750 | 84 | 140508927 | 3932 | 5,2 | 169,167 | 16,9 | 46,52 | 233 | 149 | 6.19E-10 | 1.67E-08 | 0.019,0.008 | 0.182,0.1 | -0.128 |
| 4675 | XLOC_035171 | chr5 | - | 179044984 | 179045320 | 179044984 | 179045061 | 179045145 | 84 | 179045320 | 4675 | 36,27 | 17,841,700 | 747,517 | 18,441,253 | 233 | 149 | 0 | 0 | 0.013,0.01 | 0.206,0.209 | -0.196 |
| 9440 | XLOC_016447 | chr16 | + | 67911406 | 67911748 | 67911406 | 67911559 | 67911643 | 84 | 67911748 | 9440 | 5,5 | 128,153 | 25,18 | 102,52 | 233 | 149 | 3.19E-10 | 8.99E-09 | 0.024,0.02 | 0.135,0.181 | -0.136 |
| 11122 | XLOC_030154 | chr3 | - | 194159405 | 194159701 | 194159405 | 194159500 | 194159584 | 84 | 194159701 | 11122 | 7,3 | 1,3 | 44,34 | 0,0 | 233 | 149 | 0.00124044 | 0.007314714 | 0.817,0.39 | 1.0,1.0 | -0.397 |
| 11700 | XLOC_007557 | chr11 | - | 63767001 | 63767220 | 63767001 | 63767032 | 63767116 | 84 | 63767220 | 11700 | 17,23 | 5,3 | 38,46 | 3,0 | 233 | 149 | 0.004834398 | 0.022608015 | 0.685,0.831 | 0.89,1.0 | -0.187 |
| 12338 | XLOC_009461 | chr12 | - | 75900095 | 75900385 | 75900095 | 75900179 | 75900263 | 84 | 75900385 | 12338 | 9,10 | 165,122 | 92,63 | 192,148 | 233 | 149 | 4.44E-16 | 2.74E-14 | 0.034,0.05 | 0.235,0.214 | -0.182 |
| 12955 | XLOC_004959 | chr10 | + | 91477131 | 91477479 | 91477131 | 91477248 | 91477332 | 84 | 91477479 | 12955 | 2,0 | 2,4 | 26,11 | 8,0 | 233 | 149 | 0.000210522 | 0.001622926 | 0.39,0.0 | 0.675,1.0 | -0.643 |
| 13110 | XLOC_021324 | chr19 | + | 50363312 | 50363593 | 50363312 | 50363440 | 50363524 | 84 | 50363593 | 13110 | 46,63 | 386,419 | 203,132 | 372,330 | 233 | 149 | 3.33E-16 | 2.11E-14 | 0.071,0.088 | 0.259,0.204 | -0.152 |
| 13188 | XLOC_021330 | chr19 | + | 50880809 | 50881127 | 50880809 | 50880905 | 50880989 | 84 | 50881127 | 13188 | 4,4 | 347,359 | 84,85 | 290,207 | 233 | 149 | 0 | 0 | 0.007,0.007 | 0.156,0.208 | -0.175 |
| 13194 | XLOC_044330 | chr8 | - | 74204923 | 74205226 | 74204923 | 74205032 | 74205116 | 84 | 74205226 | 13194 | 40,45 | 6,12 | 39,24 | 3,1 | 233 | 149 | 0.009103643 | 0.038162883 | 0.81,0.706 | 0.893,0.939 | -0.158 |
| 13370 | XLOC_028389 | chr22 | + | 20127000 | 20127415 | 20127000 | 20127158 | 20127242 | 84 | 20127415 | 13370 | 1,4 | 81,52 | 18,10 | 54,41 | 233 | 149 | 6.88E-07 | 1.05E-05 | 0.008,0.047 | 0.176,0.135 | -0.128 |
| 14643 | XLOC_000803 | chr1 | + | 242012032 | 242012507 | 242012032 | 242012329 | 242012413 | 84 | 242012507 | 14643 | 21,47 | 14,14 | 82,60 | 3,2 | 233 | 149 | 5.96E-08 | 1.14E-06 | 0.49,0.682 | 0.946,0.95 | -0.362 |
| 1009 | XLOC_021882 | chr19 | - | 55752606 | 55753007 | 55752606 | 55752759 | 55752844 | 85 | 55753007 | 1009 | 37,34 | 230,207 | 195,125 | 256,243 | 234 | 149 | 2.59E-14 | 1.31E-12 | 0.093,0.095 | 0.327,0.247 | -0.193 |
| 1476 | XLOC_016755 | chr16 | - | 66839679 | 66839929 | 66839679 | 66839729 | 66839814 | 85 | 66839929 | 1476 | 62,32 | 371,408 | 124,87 | 229,158 | 234 | 149 | 0 | 0 | 0.096,0.048 | 0.256,0.26 | -0.186 |
| 1740 | XLOC_009456 | chr12 | - | 72013705 | 72014095 | 72013705 | 72013904 | 72013989 | 85 | 72014095 | 1740 | 3,3 | 183,232 | 38,19 | 157,131 | 234 | 149 | 1.40E-14 | 7.54E-13 | 0.01,0.008 | 0.134,0.085 | -0.101 |
| 5030 | XLOC_041533 | chr7 | - | 72717596 | 72717911 | 72717596 | 72717737 | 72717822 | 85 | 72717911 | 5030 | 2,6 | 251,214 | 64,38 | 283,206 | 234 | 149 | 0 | 0 | 0.005,0.018 | 0.126,0.105 | -0.104 |
| 5391 | XLOC_028749 | chr22 | - | 50900633 | 50901111 | 50900633 | 50900902 | 50900987 | 85 | 50901111 | 5391 | 16,1 | 151,156 | 47,22 | 130,93 | 234 | 149 | 2.41E-10 | 6.96E-09 | 0.063,0.004 | 0.187,0.131 | -0.126 |
| 5407 | XLOC_028745 | chr22 | - | 50656617 | 50656971 | 50656617 | 50656831 | 50656916 | 85 | 50656971 | 5407 | 3,1 | 31,37 | 33,24 | 13,12 | 234 | 149 | 0 | 0 | 0.058,0.017 | 0.618,0.56 | -0.551 |
| 6694 | XLOC_016235 | chr16 | + | 2809569 | 2809997 | 2809569 | 2809662 | 2809747 | 85 | 2809997 | 6694 | 306,406 | 137,147 | 627,414 | 9,10 | 234 | 149 | 0 | 0 | 0.587,0.638 | 0.978,0.963 | -0.358 |
| 7135 | XLOC_000569 | chr1 | + | 160322949 | 160323291 | 160322949 | 160323027 | 160323112 | 85 | 160323291 | 7135 | 2,8 | 4,5 | 37,19 | 7,2 | 234 | 149 | 0.004350148 | 0.020699845 | 0.241,0.505 | 0.771,0.858 | -0.442 |
| 7287 | XLOC_020232 | chr18 | - | 47799046 | 47799321 | 47799046 | 47799108 | 47799193 | 85 | 47799321 | 7287 | 7,7 | 182,175 | 63,82 | 224,126 | 234 | 149 | 2.22E-16 | 1.46E-14 | 0.024,0.025 | 0.152,0.293 | -0.198 |
| 9922 | XLOC_000550 | chr1 | + | 155581343 | 155581618 | 155581343 | 155581394 | 155581479 | 85 | 155581618 | 9922 | 1,0 | 12,2 | 4,2 | 4,15 | 234 | 149 | 0.004003209 | 0.019388655 | 0.05,0.0 | 0.389,0.078 | -0.209 |
| 10276 | XLOC_000041 | chr1 | + | 6648119 | 6648502 | 6648119 | 6648256 | 6648341 | 85 | 6648502 | 10276 | 0,0 | 3,2 | 1,3 | 1,0 | 234 | 149 | 9.34E-06 | 0.000107388 | 0.0,0.0 | 0.389,1.0 | -0.695 |
| 12305 | XLOC_000991 | chr1 | - | 32682859 | 32683178 | 32682859 | 32682952 | 32683037 | 85 | 32683178 | 12305 | 0,2 | 16,28 | 2,8 | 7,2 | 234 | 149 | 0.000226437 | 0.001727635 | 0.0,0.044 | 0.154,0.718 | -0.414 |
| 12531 | XLOC_007434 | chr11 | - | 14541738 | 14542029 | 14541738 | 14541757 | 14541842 | 85 | 14542029 | 12531 | 45,24 | 0,7 | 36,32 | 0,0 | 234 | 149 | 0.000835991 | 0.005242765 | 1.0,0.686 | 1.0,1.0 | -0.157 |
| 14273 | XLOC_029428 | chr3 | + | 49736908 | 49737205 | 49736908 | 49737015 | 49737100 | 85 | 49737205 | 14273 | 2,0 | 77,117 | 34,25 | 180,127 | 234 | 149 | 7.85E-11 | 2.43E-09 | 0.016,0.0 | 0.107,0.111 | -0.101 |
| 14679 | XLOC_016576 | chr16 | - | 2089820 | 2090242 | 2089820 | 2090048 | 2090133 | 85 | 2090242 | 14679 | 6,5 | 171,156 | 65,22 | 226,138 | 234 | 149 | 7.29E-12 | 2.66E-10 | 0.022,0.02 | 0.155,0.092 | -0.103 |
| 351 | XLOC_035168 | chr5 | - | 178029485 | 178030935 | 178029485 | 178030758 | 178030844 | 86 | 178030935 | 351 | 4,5 | 84,53 | 12,24 | 55,40 | 235 | 149 | 7.11E-06 | 8.39E-05 | 0.029,0.056 | 0.122,0.276 | -0.157 |
| 2291 | XLOC_046145 | chr9 | + | 139749213 | 139749562 | 139749213 | 139749302 | 139749388 | 86 | 139749562 | 2291 | 8,9 | 7,9 | 8,20 | 3,3 | 235 | 149 | 0.0060583 | 0.027274902 | 0.42,0.388 | 0.628,0.809 | -0.315 |
| 4326 | XLOC_041217 | chr7 | + | 99950619 | 99950893 | 99950619 | 99950746 | 99950832 | 86 | 99950893 | 4326 | 21,17 | 112,116 | 79,55 | 66,67 | 235 | 149 | 1.35E-14 | 7.33E-13 | 0.106,0.085 | 0.431,0.342 | -0.291 |
| 8347 | XLOC_016862 | chr16 | - | 89865358 | 89865640 | 89865358 | 89865487 | 89865573 | 86 | 89865640 | 8347 | 37,51 | 182,193 | 59,23 | 43,35 | 235 | 149 | 4.53E-07 | 7.27E-06 | 0.114,0.144 | 0.465,0.294 | -0.251 |
| 8352 | XLOC_016837 | chr16 | - | 84213551 | 84213934 | 84213551 | 84213689 | 84213775 | 86 | 84213934 | 8352 | 2,3 | 59,61 | 12,17 | 47,45 | 235 | 149 | 8.83E-07 | 1.31E-05 | 0.021,0.03 | 0.139,0.193 | -0.141 |
| 9097 | XLOC_021565 | chr19 | - | 12816823 | 12817181 | 12816823 | 12816923 | 12817009 | 86 | 12817181 | 9097 | 23,22 | 5,5 | 28,16 | 0,0 | 235 | 149 | 6.72E-06 | 7.99E-05 | 0.745,0.736 | 1.0,1.0 | -0.26 |
| 9397 | XLOC_009615 | chr12 | - | 132626663 | 132626862 | 132626663 | 132626735 | 132626821 | 86 | 132626862 | 9397 | 0,0 | 41,46 | 26,14 | 61,34 | 235 | 149 | 4.98E-12 | 1.86E-10 | 0.0,0.0 | 0.213,0.207 | -0.21 |
| 10065 | XLOC_028486 | chr22 | + | 39066888 | 39067202 | 39066888 | 39066994 | 39067080 | 86 | 39067202 | 10065 | 3,9 | 3,1 | 20,8 | 1,0 | 235 | 149 | 0.003803841 | 0.018646876 | 0.388,0.851 | 0.927,1.0 | -0.344 |
| 10602 | XLOC_007390 | chr11 | - | 3382972 | 3383282 | 3382972 | 3383119 | 3383205 | 86 | 3383282 | 10602 | 14,16 | 21,27 | 46,13 | 7,2 | 235 | 149 | 7.55E-10 | 2.01E-08 | 0.297,0.273 | 0.806,0.805 | -0.521 |
| 12509 | XLOC_016216 | chr16 | + | 2257035 | 2257408 | 2257035 | 2257111 | 2257197 | 86 | 2257408 | 12509 | 1,0 | 13,20 | 4,2 | 19,1 | 235 | 149 | 0.004916234 | 0.022952719 | 0.047,0.0 | 0.118,0.559 | -0.315 |
| 13057 | XLOC_004755 | chr10 | + | 5762528 | 5762737 | 5762528 | 5762631 | 5762717 | 86 | 5762737 | 13057 | 1,4 | 123,141 | 59,55 | 140,124 | 235 | 149 | 0 | 0 | 0.005,0.018 | 0.211,0.219 | -0.203 |
| 13083 | XLOC_021582 | chr19 | - | 14521027 | 14521417 | 14521027 | 14521146 | 14521232 | 86 | 14521417 | 13083 | 7,12 | 12,26 | 16,8 | 6,6 | 235 | 149 | 0.003517735 | 0.017462554 | 0.27,0.226 | 0.628,0.458 | -0.295 |
| 14153 | XLOC_000269 | chr1 | + | 44680210 | 44680570 | 44680210 | 44680288 | 44680374 | 86 | 44680570 | 14153 | 17,13 | 16,11 | 78,39 | 7,4 | 235 | 149 | 6.72E-09 | 1.49E-07 | 0.403,0.428 | 0.876,0.861 | -0.453 |
| 1193 | XLOC_024434 | chr2 | - | 230656581 | 230656942 | 230656581 | 230656781 | 230656868 | 87 | 230656942 | 1193 | 1,13 | 295,232 | 134,58 | 288,196 | 236 | 149 | 0 | 0 | 0.002,0.034 | 0.227,0.157 | -0.174 |
| 1232 | XLOC_005276 | chr10 | - | 79745660 | 79745917 | 79745660 | 79745743 | 79745830 | 87 | 79745917 | 1232 | 12,0 | 59,34 | 34,20 | 61,65 | 236 | 149 | 0.000328476 | 0.002399934 | 0.114,0.0 | 0.26,0.163 | -0.155 |
| 3051 | XLOC_048132 | chrX | - | 153715589 | 153717421 | 153715589 | 153716835 | 153716922 | 87 | 153717421 | 3051 | 244,288 | 31,21 | 159,124 | 4,1 | 236 | 149 | 2.08E-06 | 2.82E-05 | 0.832,0.896 | 0.962,0.987 | -0.11 |
| 7091 | XLOC_021040 | chr19 | + | 17626981 | 17627283 | 17626981 | 17627089 | 17627176 | 87 | 17627283 | 7091 | 10,6 | 4,1 | 3,3 | 1,0 | 236 | 149 | 0.009982167 | 0.041330444 | 0.612,0.791 | 0.654,1.0 | -0.125 |
| 7538 | XLOC_018791 | chr17 | - | 80042560 | 80043032 | 80042560 | 80042587 | 80042674 | 87 | 80043032 | 7538 | 29,15 | 17,23 | 49,25 | 4,9 | 236 | 149 | 0.000980972 | 0.006001372 | 0.519,0.292 | 0.886,0.637 | -0.356 |
| 8476 | XLOC_016559 | chr16 | - | 733166 | 733342 | 733166 | 733234 | 733321 | 87 | 733342 | 8476 | 7,9 | 155,141 | 44,29 | 175,114 | 236 | 149 | 1.17E-08 | 2.48E-07 | 0.028,0.039 | 0.137,0.138 | -0.104 |
| 9560 | XLOC_013545 | chr14 | - | 105616788 | 105617248 | 105616788 | 105617114 | 105617201 | 87 | 105617248 | 9560 | 55,44 | 135,101 | 97,110 | 141,90 | 236 | 149 | 0.000479228 | 0.003316092 | 0.205,0.216 | 0.303,0.436 | -0.159 |
| 14694 | XLOC_016224 | chr16 | + | 2577986 | 2578381 | 2577986 | 2578149 | 2578236 | 87 | 2578381 | 14694 | 2,3 | 129,84 | 11,7 | 39,26 | 236 | 149 | 1.36E-07 | 2.43E-06 | 0.01,0.022 | 0.151,0.145 | -0.132 |
| 546 | XLOC_030112 | chr3 | - | 176765087 | 176765337 | 176765087 | 176765185 | 176765273 | 88 | 176765337 | 546 | 1,0 | 231,239 | 24,25 | 131,114 | 237 | 149 | 0 | 0 | 0.003,0.0 | 0.103,0.121 | -0.11 |
| 2466 | XLOC_007508 | chr11 | - | 47487344 | 47493909 | 47487344 | 47493654 | 47493742 | 88 | 47493909 | 2466 | 135,102 | 124,125 | 308,230 | 50,60 | 237 | 149 | 0 | 0 | 0.406,0.339 | 0.795,0.707 | -0.378 |
| 2507 | XLOC_007251 | chr11 | + | 93462779 | 93463135 | 93462779 | 93462947 | 93463035 | 88 | 93463135 | 2507 | 11,17 | 132,156 | 45,30 | 99,80 | 237 | 149 | 3.19E-09 | 7.50E-08 | 0.05,0.064 | 0.222,0.191 | -0.15 |
| 4520 | XLOC_038081 | chr6 | - | 3739029 | 3739470 | 3739029 | 3739203 | 3739291 | 88 | 3739470 | 4520 | 7,4 | 1,6 | 2,6 | 0,0 | 237 | 149 | 2.37E-05 | 0.000241513 | 0.815,0.295 | 1.0,1.0 | -0.445 |
| 5144 | XLOC_029841 | chr3 | - | 49161827 | 49162199 | 49161827 | 49162045 | 49162133 | 88 | 49162199 | 5144 | 7,1 | 177,285 | 28,10 | 101,63 | 237 | 149 | 1.55E-13 | 7.33E-12 | 0.024,0.002 | 0.148,0.091 | -0.106 |
| 6709 | XLOC_021882 | chr19 | - | 55751193 | 55751504 | 55751193 | 55751279 | 55751367 | 88 | 55751504 | 6709 | 13,13 | 0,5 | 18,8 | 0,0 | 237 | 149 | 0.002327981 | 0.012323116 | 1.0,0.62 | 1.0,1.0 | -0.19 |

| | | | | | | | | | | | | | | | | | | | | | | |
|---|---|---|---|---|---|---|---|---|---|---|---|---|---|---|---|---|---|---|---|---|---|---|
| 6769 | XLOC_029383 | chr3 | + | 40502898 | 40503152 | 40502898 | 40503010 | 40503098 | 88 | 40503152 | 6769 | 3,8 | 4,1 | 7,222 | 0,0 | 237 | 149 | 0.000118905 | 0.000996026 | 0.32,0.834 | 1.0,1.0 | -0.423 |
| 7798 | XLOC_021306 | chr19 | + | 49652404 | 49652582 | 49652404 | 49652425 | 49652513 | 88 | 49652582 | 7798 | 0,0 | 6,3 | 2,2 | 5,5 | 237 | 149 | 0.005472561 | 0.025094884 | 0.0,0.0 | 0.201,0.201 | -0.201 |
| 7998 | XLOC_023740 | chr2 | + | 178081403 | 178081599 | 178081403 | 178081499 | 178081587 | 88 | 178081599 | 7998 | 25,19 | 356,372 | 141,116 | 432,310 | 237 | 149 | 0 | 0 | 0.042,0.031 | 0.17,0.19 | -0.143 |
| 8037 | XLOC_017920 | chr17 | + | 8192575 | 8192928 | 8192575 | 8192732 | 8192820 | 88 | 8192928 | 8037 | 49,52 | 254,229 | 82,71 | 112,87 | 237 | 149 | 5.82E-13 | 2.54E-11 | 0.108,0.125 | 0.315,0.339 | -0.211 |
| 8124 | XLOC_017856 | chr17 | + | 2908444 | 2908734 | 2908444 | 2908574 | 2908662 | 88 | 2908734 | 8124 | 5,6 | 19,18 | 29,14 | 13,14 | 237 | 149 | 0.000182638 | 0.001437792 | 0.142,0.173 | 0.584,0.386 | -0.328 |
| 8402 | XLOC_016779 | chr16 | - | 67979853 | 67980262 | 67979853 | 67979985 | 67980073 | 88 | 67980262 | 8402 | 0,3 | 111,108 | 16,5 | 38,25 | 237 | 149 | 2.75E-10 | 7.86E-09 | 0.0,0.017 | 0.209,0.112 | -0.152 |
| 8435 | XLOC_016647 | chr16 | - | 15718708 | 15719022 | 15718708 | 15718784 | 15718872 | 88 | 15719022 | 8435 | 4,9 | 63,69 | 25,32 | 64,59 | 237 | 149 | 4.30E-07 | 6.93E-06 | 0.038,0.076 | 0.197,0.254 | -0.169 |
| 9084 | XLOC_014958 | chr15 | - | 66793286 | 66793682 | 66793286 | 66793443 | 66793531 | 88 | 66793682 | 9084 | 38,58 | 16,18 | 61,60 | 6,12 | 237 | 149 | 0.00887486 | 0.037448676 | 0.599,0.67 | 0.865,0.759 | -0.177 |
| 10210 | XLOC_007565 | chr11 | - | 64559696 | 64559894 | 64559696 | 64559747 | 64559835 | 88 | 64559894 | 10210 | 26,27 | 58,47 | 19,14 | 16,6 | 237 | 149 | 0.003192772 | 0.01608114 | 0.22,0.265 | 0.427,0.595 | -0.269 |
| 10239 | XLOC_011518 | chr13 | - | 76111634 | 76112100 | 76111634 | 76111811 | 76111899 | 88 | 76112100 | 10239 | 3,3 | 532,557 | 148,139 | 520,402 | 237 | 149 | 0 | 0 | 0.004,0.003 | 0.152,0.179 | -0.162 |
| 10549 | XLOC_013060 | chr14 | + | 24584414 | 24584601 | 24584414 | 24584502 | 24584590 | 88 | 24584601 | 10549 | 5,6 | 34,47 | 11,9 | 32,18 | 237 | 149 | 0.007929795 | 0.034112852 | 0.085,0.074 | 0.178,0.239 | -0.129 |
| 12222 | XLOC_041138 | chr7 | + | 73111618 | 73112024 | 73111618 | 73111846 | 73111934 | 88 | 73112024 | 12222 | 36,42 | 32,28 | 61,86 | 21,16 | 237 | 149 | 0.000134444 | 0.001103288 | 0.414,0.485 | 0.646,0.772 | -0.26 |
| 12837 | XLOC_001074 | chr1 | - | 44466438 | 44466723 | 44466438 | 44466539 | 44466627 | 88 | 44466723 | 12837 | 1,0 | 117,118 | 44,41 | 130,93 | 237 | 149 | 0 | 0 | 0.005,0.0 | 0.175,0.217 | -0.194 |
| 13617 | XLOC_001082 | chr1 | - | 45798956 | 45799169 | 45798956 | 45798996 | 45799084 | 88 | 45799169 | 13617 | 5,2 | 97,76 | 9,15 | 50,24 | 237 | 149 | 2.21E-06 | 2.98E-05 | 0.031,0.016 | 0.102,0.282 | -0.168 |
| 13636 | XLOC_018386 | chr17 | - | 7216530 | 7217003 | 7216530 | 7216610 | 7216698 | 88 | 7217003 | 13636 | 10,24 | 216,236 | 76,50 | 223,177 | 237 | 149 | 1.54E-11 | 5.40E-10 | 0.028,0.06 | 0.176,0.151 | -0.119 |
| 3587 | XLOC_047954 | chrX | - | 48935301 | 48935593 | 48935301 | 48935406 | 48935495 | 89 | 48935593 | 3587 | 117,114 | 182,193 | 185,128 | 45,18 | 238 | 149 | 0 | 0 | 0.287,0.27 | 0.72,0.817 | -0.49 |
| 3785 | XLOC_046315 | chr9 | - | 95085517 | 95085785 | 95085517 | 95085633 | 95085722 | 89 | 95085785 | 3785 | 4,7 | 9,17 | 14,22 | 7,6 | 238 | 149 | 5.49E-05 | 0.000510219 | 0.218,0.205 | 0.556,0.697 | -0.415 |
| 8089 | XLOC_017896 | chr17 | + | 7127647 | 7128033 | 7127647 | 7127871 | 7127960 | 89 | 7128033 | 8089 | 263,194 | 13,251,419 | 643,423 | 1,071,821 | 238 | 149 | 0 | 0 | 0.111,0.079 | 0.273,0.244 | -0.164 |
| 12336 | XLOC_020062 | chr18 | + | 21110170 | 21110576 | 21110170 | 21110261 | 21110350 | 89 | 21110576 | 12336 | 37,24 | 77,59 | 82,35 | 50,44 | 238 | 149 | 0.000828214 | 0.005207867 | 0.231,0.203 | 0.507,0.332 | -0.202 |
| 13095 | XLOC_000917 | chr1 | - | 19472286 | 19472600 | 19472286 | 19472427 | 19472516 | 89 | 19472600 | 13095 | 0,0 | 7,11 | 6,6 | 10,5 | 238 | 149 | 1.83E-06 | 2.51E-05 | 0.0,0.0 | 0.273,0.429 | -0.351 |
| 13505 | XLOC_000967 | chr1 | - | 27682878 | 27683175 | 27682878 | 27682991 | 27683080 | 89 | 27683175 | 13505 | 0,1 | 37,40 | 9,0 | 5,1 | 238 | 149 | 0.000341524 | 0.002476046 | 0.0,0.015 | 0.53,0.0 | -0.258 |
| 2770 | XLOC_001135 | chr1 | - | 59154709 | 59156089 | 59154709 | 59155921 | 59156011 | 90 | 59156089 | 2770 | 9,6 | 57,82 | 85,88 | 85,52 | 239 | 149 | 0 | 0 | 0.09,0.044 | 0.384,0.513 | -0.382 |
| 5298 | XLOC_029454 | chr3 | + | 52512170 | 52512593 | 52512170 | 52512298 | 52512388 | 90 | 52512593 | 5298 | 1,0 | 23,16 | 6,1 | 5,2 | 239 | 149 | 1.73E-05 | 0.000183063 | 0.026,0.0 | 0.428,0.238 | -0.32 |
| 6424 | XLOC_023633 | chr2 | + | 113410278 | 113410518 | 113410278 | 113410375 | 113410465 | 90 | 113410518 | 6424 | 22,35 | 14,6 | 59,38 | 5,3 | 239 | 149 | 0.004943802 | 0.023058557 | 0.495,0.784 | 0.88,0.888 | -0.245 |
| 6932 | XLOC_021369 | chr19 | + | 54710143 | 54710592 | 54710143 | 54710330 | 54710420 | 90 | 54710592 | 6932 | 68,53 | 362,323 | 201,158 | 364,315 | 239 | 149 | 4.65E-13 | 2.07E-11 | 0.105,0.093 | 0.256,0.238 | -0.148 |
| 8153 | XLOC_009276 | chr12 | - | 21654107 | 21654576 | 21654107 | 21654306 | 21654396 | 90 | 21654576 | 8153 | 130,145 | 38,24 | 65,70 | 1,6 | 239 | 149 | 4.55E-05 | 0.000433358 | 0.681,0.79 | 0.976,0.879 | -0.192 |
| 9586 | XLOC_013517 | chr14 | - | 96798641 | 96798989 | 96798641 | 96798747 | 96798837 | 90 | 96798989 | 9586 | 0,0 | 16,13 | 17,10 | 14,15 | 239 | 149 | 2.71E-09 | 6.50E-08 | 0.0,0.0 | 0.431,0.294 | -0.362 |
| 9606 | XLOC_007211 | chr11 | + | 71941141 | 71941512 | 71941141 | 71941315 | 71941405 | 90 | 71941512 | 9606 | 53,24 | 891,770 | 117,80 | 429,324 | 239 | 149 | 0 | 0 | 0.036,0.019 | 0.145,0.133 | -0.112 |
| 13891 | XLOC_007127 | chr11 | + | 63970483 | 63970700 | 63970483 | 63970506 | 63970596 | 90 | 63970700 | 13891 | 5,16 | 5,0 | 5,6 | 0,0 | 239 | 149 | 0.003933758 | 0.019137605 | 0.384,1.0 | 1.0,1.0 | -0.308 |
| 14919 | XLOC_017909 | chr17 | + | 7454115 | 7454291 | 7454115 | 7454169 | 7454259 | 90 | 7454291 | 14919 | 28,19 | 59,65 | 30,23 | 21,20 | 239 | 149 | 1.72E-05 | 0.000182295 | 0.228,0.154 | 0.471,0.418 | -0.254 |
| 14980 | XLOC_001222 | chr1 | - | 109823391 | 109823873 | 109823391 | 109823668 | 109823758 | 90 | 109823873 | 14980 | 136,111 | 105,123 | 60,59 | 32,25 | 239 | 149 | 0.00245026 | 0.012859539 | 0.447,0.36 | 0.539,0.595 | -0.163 |
| 14993 | XLOC_001522 | chr1 | - | 220213526 | 220213878 | 220213526 | 220213611 | 220213701 | 90 | 220213878 | 14993 | 0,0 | 4,1 | 1,2 | 0,0 | 239 | 149 | 7.14E-10 | 1.91E-08 | 0.0,0.0 | 1.0,1.0 | -1 |
| 121 | XLOC_044167 | chr8 | + | 144460438 | 144461248 | 144460438 | 144460990 | 144461081 | 91 | 144461248 | 121 | 34,23 | 42,72 | 28,23 | 24,6 | 240 | 149 | 0.003963788 | 0.01926379 | 0.334,0.165 | 0.42,0.704 | -0.312 |
| 419 | XLOC_034648 | chr5 | + | 112928999 | 112929247 | 112928999 | 112929087 | 112929178 | 91 | 112929247 | 419 | 30,23 | 51,47 | 62,46 | 39,45 | 240 | 149 | 0.000813043 | 0.005128451 | 0.268,0.233 | 0.497,0.388 | -0.192 |
| 6043 | XLOC_020995 | chr19 | + | 13002084 | 13002474 | 13002084 | 13002209 | 13002300 | 91 | 13002474 | 6043 | 0,1 | 97,98 | 49,11 | 103,81 | 240 | 149 | 5.06E-13 | 2.22E-11 | 0.0,0.006 | 0.228,0.078 | -0.15 |
| 7387 | XLOC_037628 | chr6 | + | 4056554 | 4056922 | 4056554 | 4056669 | 4056760 | 91 | 4056922 | 7387 | 22,22 | 21,20 | 52,28 | 9,11 | 240 | 149 | 0.000479794 | 0.00331838 | 0.394,0.406 | 0.782,0.612 | -0.297 |
| 8231 | XLOC_013326 | chr14 | - | 24423075 | 24423522 | 24423075 | 24423242 | 24423333 | 91 | 24423522 | 8231 | 9,4 | 10,3 | 1,3 | 1,0 | 240 | 149 | 0.00453848 | 0.021437106 | 0.358,0.453 | 0.383,1.0 | -0.286 |
| 9890 | XLOC_021573 | chr19 | - | 13223475 | 13223825 | 13223475 | 13223631 | 13223722 | 91 | 13223825 | 9890 | 23,9 | 24,9 | 30,37 | 16,6 | 240 | 149 | 0.006719032 | 0.02972835 | 0.373,0.383 | 0.538,0.793 | -0.287 |
| 11121 | XLOC_030154 | chr3 | - | 194158039 | 194158299 | 194158039 | 194158193 | 194158284 | 91 | 194158299 | 11121 | 9,9 | 316,382 | 48,64 | 282,240 | 240 | 149 | 0 | 0 | 0.017,0.014 | 0.096,0.142 | -0.103 |
| 11476 | XLOC_026876 | chr20 | + | 62326190 | 62326574 | 62326190 | 62326327 | 62326418 | 91 | 62326574 | 11476 | 8,18 | 61,50 | 20,18 | 27,31 | 240 | 149 | 0.001525084 | 0.008688486 | 0.075,0.183 | 0.315,0.265 | -0.161 |
| 13066 | XLOC_017919 | chr17 | + | 8159122 | 8159477 | 8159122 | 8159228 | 8159319 | 91 | 8159477 | 13066 | 4,1 | 7,9 | 13,4 | 2,0 | 240 | 149 | 1.29E-06 | 1.84E-05 | 0.262,0.065 | 0.801,1.0 | -0.737 |
| 14949 | XLOC_000082 | chr1 | + | 12052335 | 12052747 | 12052335 | 12052520 | 12052611 | 91 | 12052747 | 14949 | 1,3 | 13,7 | 12,5 | 13,1 | 240 | 149 | 0.003277676 | 0.016461815 | 0.046,0.21 | 0.364,0.756 | -0.432 |
| 349 | XLOC_035171 | chr5 | - | 179043869 | 179044111 | 179043869 | 179043959 | 179044051 | 92 | 179044111 | 349 | 311,323 | 19,182,000 | 928,682 | 16,631,271 | 241 | 149 | 0 | 0 | 0.091,0.091 | 0.257,0.249 | -0.162 |
| 3282 | XLOC_001211 | chr1 | - | 100573183 | 100573560 | 100573183 | 100573273 | 100573365 | 92 | 100573560 | 3282 | 16,33 | 201,192 | 98,90 | 159,91 | 241 | 149 | 0 | 0 | 0.047,0.096 | 0.276,0.379 | -0.256 |
| 4167 | XLOC_041744 | chr7 | - | 140051676 | 140051981 | 140051676 | 140051769 | 140051861 | 92 | 140051981 | 4167 | 0,5 | 2,8 | 8,2 | 2,1 | 241 | 149 | 0.008235986 | 0.035246566 | 0.0,0.279 | 0.712,0.553 | -0.493 |
| 5806 | XLOC_026867 | chr20 | + | 60963341 | 60963745 | 60963341 | 60963420 | 60963512 | 92 | 60963745 | 5806 | 180,125 | 67,596,952 | 31,162,360 | 75,836,093 | 241 | 149 | 0 | 0 | 0.016,0.011 | 0.203,0.193 | -0.184 |
| 6116 | XLOC_028393 | chr22 | + | 20936945 | 20937254 | 20936945 | 20937034 | 20937126 | 92 | 20937254 | 6116 | 5,4 | 115,110 | 18,4 | 41,33 | 241 | 149 | 5.02E-06 | 6.16E-05 | 0.026,0.022 | 0.213,0.07 | -0.118 |
| 6479 | XLOC_023462 | chr2 | + | 46803699 | 46804018 | 46803699 | 46803795 | 46803887 | 92 | 46804018 | 6479 | 3,7 | 7,1 | 20,31 | 0,0 | 241 | 149 | 7.05E-06 | 8.33E-05 | 0.209,0.812 | 1.0,1.0 | -0.489 |
| 6999 | XLOC_021235 | chr19 | + | 42818650 | 42819003 | 42818650 | 42818851 | 42818943 | 92 | 42819003 | 6999 | 0,0 | 3,2 | 1,5 | 17,3 | 241 | 149 | 0.002083568 | 0.011252862 | 0.0,0.0 | 0.035,0.507 | -0.271 |
| 7892 | XLOC_018249 | chr17 | + | 73998299 | 73998693 | 73998299 | 73998499 | 73998591 | 92 | 73998693 | 7892 | 6,5 | 43,32 | 25,5 | 9,12 | 241 | 149 | 0.000893967 | 0.005554554 | 0.079,0.088 | 0.632,0.205 | -0.335 |
| 8038 | XLOC_017920 | chr17 | + | 8192575 | 8192928 | 8192575 | 8192732 | 8192824 | 92 | 8192928 | 8038 | 43,51 | 5,8 | 69,64 | 2,0 | 241 | 149 | 4.68E-05 | 0.000444203 | 0.842,0.798 | 0.955,1.0 | -0.157 |
| 8880 | XLOC_016344 | chr16 | + | 28967255 | 28967658 | 28967255 | 28967379 | 28967471 | 92 | 28967658 | 8880 | 65,74 | 260,255 | 69,66 | 97,77 | 241 | 149 | 3.89E-09 | 9.03E-08 | 0.134,0.152 | 0.305,0.346 | -0.182 |
| 8946 | XLOC_014664 | chr15 | + | 63848865 | 63849121 | 63848865 | 63848947 | 63849039 | 92 | 63849121 | 8946 | 1,0 | 4,1 | 4,1 | 1,7 | 241 | 149 | 0.011537133 | 0.04672754 | 0.134,0.0 | 0.712,0.081 | -0.329 |
| 10510 | XLOC_009358 | chr12 | - | 50387718 | 50388112 | 50387718 | 50387821 | 50387913 | 92 | 50388112 | 10510 | 18,27 | 8,9 | 21,20 | 0,0 | 241 | 149 | 6.92E-08 | 1.31E-06 | 0.582,0.65 | 1.0,1.0 | -0.384 |
| 10794 | XLOC_018530 | chr17 | - | 38209550 | 38209932 | 38209550 | 38209633 | 38209725 | 92 | 38209932 | 10794 | 0,2 | 4,3 | 5,5 | 4,4 | 241 | 149 | 0.007742287 | 0.033459095 | 0.0,0.292 | 0.436,0.436 | -0.29 |
| 11342 | XLOC_000252 | chr1 | + | 43825863 | 43826262 | 43825863 | 43826060 | 43826152 | 92 | 43826262 | 11342 | 9,1 | 4,2 | 5,3 | 0,0 | 241 | 149 | 4.66E-06 | 5.79E-05 | 0.582,0.236 | 1.0,1.0 | -0.591 |
| 14595 | XLOC_000697 | chr1 | + | 207495580 | 207495912 | 207495580 | 207495634 | 207495726 | 92 | 207495912 | 14595 | 9,10 | 7,6 | 69,70 | 9,9 | 241 | 149 | 0.000339585 | 0.00246452 | 0.443,0.507 | 0.826,0.828 | -0.352 |
| 14771 | XLOC_001039 | chr1 | - | 38463359 | 38463804 | 38463359 | 38463542 | 38463634 | 92 | 38463804 | 14771 | 11,10 | 135,158 | 46,27 | 114,82 | 241 | 149 | 2.31E-10 | 6.70E-09 | 0.048,0.038 | 0.2,0.169 | -0.142 |
| 2567 | XLOC_014712 | chr15 | + | 75644451 | 75644728 | 75644451 | 75644571 | 75644664 | 93 | 75644728 | 2567 | 5,10 | 23,34 | 19,20 | 33,10 | 242 | 149 | 0.003738464 | 0.018403259 | 0.118,0.153 | 0.262,0.552 | -0.272 |
| 4750 | XLOC_034842 | chr5 | - | 1632647 | 1633021 | 1632647 | 1632866 | 1632959 | 93 | 1633021 | 4750 | 66,52 | 20,6 | 37,35 | 0,4 | 242 | 149 | 0.004261142 | 0.020339124 | 0.67,0.842 | 1.0,0.843 | -0.165 |
| 6873 | XLOC_021479 | chr19 | - | 4110356 | 4110653 | 4110356 | 4110413 | 4110506 | 93 | 4110653 | 6873 | 9,4 | 0,11 | 11,6 | 0,0 | 242 | 149 | 0.001430161 | 0.00823071 | 1.0,0.183 | 1.0,1.0 | -0.408 |
| 7995 | XLOC_023740 | chr2 | + | 178080477 | 178080912 | 178080477 | 178080624 | 178080717 | 93 | 178080912 | 7995 | 5,13 | 38,0 | 27,16 | 0,0 | 242 | 149 | 0.000656059 | 0.004319588 | 0.075,1.0 | 1.0,1.0 | -0.463 |
| 8682 | XLOC_016239 | chr16 | + | 3016140 | 3016414 | 3016140 | 3016232 | 3016325 | 93 | 3016414 | 8682 | 8,9 | 16,12 | 12,5 | 2,3 | 242 | 149 | 0.005201495 | 0.023998926 | 0.235,0.316 | 0.787,0.506 | -0.371 |
| 8855 | XLOC_030079 | chr3 | - | 153998366 | 153998651 | 153998366 | 153998458 | 153998551 | 93 | 153998651 | 8855 | 3,9 | 123,137 | 79,62 | 148,115 | 242 | 149 | 0 | 0 | 0.015,0.039 | 0.247,0.249 | -0.221 |
| 9696 | XLOC_013334 | chr14 | - | 24737571 | 24737825 | 24737571 | 24737626 | 24737719 | 93 | 24737825 | 9696 | 10,2 | 78,90 | 15,27 | 61,51 | 242 | 149 | 1.40E-06 | 1.98E-05 | 0.073,0.013 | 0.131,0.246 | -0.146 |

| | | | | | | | | | | | | | | | | | | | | | | |
|---|---|---|---|---|---|---|---|---|---|---|---|---|---|---|---|---|---|---|---|---|---|---|
| 11109 | XLOC_020148 | chr18 | - | 265302 | 265527 | 265302 | 265363 | 265456 | 93 | 265527 | 11109 | 2,0 | 118,119 | 52,22 | 91,91 | 242 | 149 | 0 | 0 | 0.01,0.0 | 0.26,0.13 | -0.19 |
| 14035 | XLOC_041406 | chr7 | - | 12376613 | 12376921 | 12376613 | 12376709 | 12376802 | 93 | 12376921 | 14035 | 4,11 | 17,10 | 18,11 | 2,3 | 242 | 149 | 5.98E-05 | 0.000547361 | 0.127,0.404 | 0.847,0.693 | -0.504 |
| 1033 | XLOC_023585 | chr2 | + | 97873547 | 97873743 | 97873547 | 97873576 | 97873670 | 94 | 97873743 | 1033 | 1,2 | 4,14 | 3,3 | 2,2 | 243 | 149 | 0.006404804 | 0.028570767 | 0.133,0.081 | 0.479,0.479 | -0.372 |
| 2060 | XLOC_023860 | chr2 | + | 220097800 | 220098155 | 220097800 | 220097913 | 220098007 | 94 | 220098155 | 2060 | 32,21 | 203,239 | 87,61 | 69,60 | 243 | 149 | 0 | 0 | 0.088,0.051 | 0.436,0.384 | -0.341 |
| 4768 | XLOC_034736 | chr5 | + | 145887195 | 145887519 | 145887195 | 145887341 | 145887435 | 94 | 145887519 | 4768 | 14,10 | 14,17 | 18,9 | 2,5 | 243 | 149 | 0.005623709 | 0.025645348 | 0.38,0.265 | 0.847,0.525 | -0.363 |
| 8282 | XLOC_021139 | chr19 | + | 36104769 | 36105036 | 36104769 | 36104802 | 36104896 | 94 | 36105036 | 8282 | 2,5 | 4,1 | 9,5 | 0,1 | 243 | 149 | 0.00100512 | 0.006123946 | 0.235,0.754 | 1.0,0.754 | -0.383 |
| 10244 | XLOC_018507 | chr17 | - | 34868500 | 34868726 | 34868500 | 34868622 | 34868716 | 94 | 34868726 | 10244 | 19,21 | 7,7 | 20,14 | 0,0 | 243 | 149 | 5.15E-07 | 8.14E-06 | 0.625,0.648 | 1.0,1.0 | -0.363 |
| 10246 | XLOC_018507 | chr17 | - | 34883194 | 34883530 | 34883194 | 34883287 | 34883381 | 94 | 34883530 | 10246 | 5,2 | 4,8 | 37,20 | 5,8 | 243 | 149 | 0.0015484 | 0.00880002 | 0.434,0.133 | 0.819,0.605 | -0.428 |
| 10493 | XLOC_046108 | chr9 | + | 134385129 | 134385528 | 134385129 | 134385195 | 134385289 | 94 | 134385528 | 10493 | 1,3 | 19,34 | 25,7 | 20,20 | 243 | 149 | 1.58E-05 | 0.000168757 | 0.031,0.051 | 0.434,0.177 | -0.265 |
| 13089 | XLOC_011478 | chr13 | - | 50123593 | 50124036 | 50123593 | 50123784 | 50123878 | 94 | 50124036 | 13089 | 5,4 | 4,9 | 5,3 | 0,1 | 243 | 149 | 0.000156322 | 0.001254395 | 0.434,0.214 | 1.0,0.648 | -0.5 |
| 14987 | XLOC_000367 | chr1 | + | 90470561 | 90470766 | 90470561 | 90470597 | 90470691 | 94 | 90470766 | 14987 | 11,21 | 116,87 | 65,37 | 124,84 | 243 | 149 | 1.88E-06 | 2.57E-05 | 0.055,0.129 | 0.243,0.213 | -0.136 |
| 117 | XLOC_044185 | chr8 | + | 145725649 | 145725995 | 145725649 | 145725774 | 145725869 | 95 | 145725995 | 117 | 69,97 | 102,71 | 95,63 | 19,29 | 244 | 149 | 0.000227426 | 0.0017328 | 0.292,0.455 | 0.753,0.57 | -0.288 |
| 5353 | XLOC_029298 | chr3 | + | 9475811 | 9476169 | 9475811 | 9475865 | 9475960 | 95 | 9476169 | 5353 | 13,10 | 1,3 | 14,9 | 0,0 | 244 | 149 | 0.006793313 | 0.029998933 | 0.888,0.671 | 1.0,1.0 | -0.22 |
| 6400 | XLOC_023741 | chr2 | + | 178301471 | 178301782 | 178301471 | 178301612 | 178301707 | 95 | 178301782 | 6400 | 0,0 | 90,93 | 21,17 | 113,84 | 244 | 149 | 7.49E-11 | 2.33E-09 | 0.0,0.0 | 0.102,0.11 | -0.106 |
| 7434 | XLOC_037805 | chr6 | + | 35436915 | 35437306 | 35436915 | 35437062 | 35437157 | 95 | 35437306 | 7434 | 455,451 | 298,334 | 1,123,877 | 250,221 | 244 | 149 | 0 | 0 | 0.483,0.452 | 0.733,0.708 | -0.253 |
| 8067 | XLOC_017896 | chr17 | + | 7125255 | 7125621 | 7125255 | 7125400 | 7125495 | 95 | 7125621 | 8067 | 839,722 | 12,491,246 | 1,216,799 | 799,606 | 244 | 149 | 1.16E-13 | 5.66E-12 | 0.291,0.261 | 0.482,0.446 | -0.188 |
| 8914 | XLOC_026746 | chr20 | + | 31015933 | 31016225 | 31015933 | 31016032 | 31016127 | 95 | 31016225 | 8914 | 31,42 | 4,5 | 27,21 | 1,0 | 244 | 149 | 0.001180854 | 0.007013122 | 0.826,0.837 | 0.943,1.0 | -0.14 |
| 9152 | XLOC_014850 | chr15 | - | 41072125 | 41072550 | 41072125 | 41072195 | 41072290 | 95 | 41072550 | 9152 | 0,0 | 1,5 | 1,1 | 0,0 | 244 | 149 | 3.79E-09 | 8.82E-08 | 0.0,0.0 | 1.0,1.0 | -1 |
| 9478 | XLOC_007774 | chr11 | - | 134026646 | 134026990 | 134026646 | 134026817 | 134026912 | 95 | 134026990 | 9478 | 35,23 | 13,9 | 18,15 | 0,0 | 244 | 149 | 1.22E-07 | 2.22E-06 | 0.622,0.609 | 1.0,1.0 | -0.385 |
| 10397 | XLOC_046127 | chr9 | + | 138393689 | 138394107 | 138393689 | 138393819 | 138393914 | 95 | 138394107 | 10397 | 13,17 | 2,5 | 45,38 | 0,1 | 244 | 149 | 4.96E-05 | 0.000467156 | 0.799,0.675 | 1.0,0.959 | -0.242 |
| 7814 | XLOC_018393 | chr17 | - | 7495388 | 7495672 | 7495388 | 7495475 | 7495571 | 96 | 7495672 | 7814 | 12,20 | 21,28 | 62,33 | 10,3 | 245 | 149 | 3.69E-12 | 1.41E-10 | 0.258,0.303 | 0.79,0.87 | -0.55 |
| 9272 | XLOC_014584 | chr15 | + | 41793383 | 41793757 | 41793383 | 41793564 | 41793660 | 96 | 41793757 | 9272 | 1,1 | 0,4 | 1,1 | 0,0 | 245 | 149 | 0.00188691 | 0.010357104 | 1.0,0.132 | 1.0,1.0 | -0.434 |
| 10545 | XLOC_021824 | chr19 | - | 50165401 | 50165874 | 50165401 | 50165585 | 50165681 | 96 | 50165874 | 10545 | 27,47 | 244,203 | 58,59 | 94,86 | 245 | 149 | 3.57E-12 | 1.37E-10 | 0.063,0.123 | 0.273,0.294 | -0.19 |
| 11568 | XLOC_018736 | chr17 | - | 74387836 | 74388063 | 74387836 | 74387894 | 74387990 | 96 | 74388063 | 11568 | 14,7 | 149,139 | 55,37 | 165,144 | 245 | 149 | 1.38E-08 | 2.89E-07 | 0.054,0.03 | 0.169,0.135 | -0.11 |
| 14626 | XLOC_001370 | chr1 | - | 156518118 | 156518540 | 156518118 | 156518276 | 156518372 | 96 | 156518540 | 14626 | 4,2 | 28,37 | 13,6 | 14,17 | 245 | 149 | 0.000170332 | 0.00134994 | 0.08,0.032 | 0.361,0.177 | -0.213 |
| 15086 | XLOC_030106 | chr3 | - | 170585801 | 170586172 | 170585801 | 170585990 | 170586086 | 96 | 170586172 | 15086 | 14,8 | 191,229 | 38,47 | 161,154 | 245 | 149 | 2.33E-11 | 7.93E-10 | 0.043,0.021 | 0.126,0.157 | -0.11 |
| 1860 | XLOC_013325 | chr14 | - | 24032408 | 24032711 | 24032408 | 24032491 | 24032588 | 97 | 24032711 | 1860 | 17,15 | 1,6 | 27,18 | 0,0 | 246 | 149 | 7.64E-05 | 0.000675054 | 0.911,0.602 | 1.0,1.0 | -0.244 |
| 3957 | XLOC_044466 | chr8 | - | 145741147 | 145741534 | 145741147 | 145741274 | 145741371 | 97 | 145741534 | 3957 | 26,20 | 225,266 | 97,67 | 181,102 | 246 | 149 | 0 | 0 | 0.065,0.044 | 0.245,0.285 | -0.211 |
| 5174 | XLOC_029837 | chr3 | - | 49053251 | 49053631 | 49053251 | 49053493 | 49053590 | 97 | 49053631 | 5174 | 46,20 | 22,36 | 21,19 | 6,0 | 246 | 149 | 0.001442733 | 0.008299682 | 0.559,0.252 | 0.679,1.0 | -0.434 |
| 6256 | XLOC_024101 | chr2 | - | 74657594 | 74657804 | 74657594 | 74657645 | 74657742 | 97 | 74657804 | 6256 | 5,13 | 121,120 | 20,33 | 52,45 | 246 | 149 | 1.96E-10 | 5.74E-09 | 0.024,0.062 | 0.189,0.308 | -0.206 |
| 7269 | XLOC_021195 | chr19 | + | 38895956 | 38896275 | 38895956 | 38896061 | 38896158 | 97 | 38896275 | 7269 | 3,2 | 28,27 | 7,10 | 9,9 | 246 | 149 | 5.05E-06 | 6.21E-05 | 0.061,0.043 | 0.32,0.402 | -0.309 |
| 9379 | XLOC_018149 | chr17 | + | 48456812 | 48457137 | 48456812 | 48456928 | 48457025 | 97 | 48457137 | 9379 | 28,21 | 2,8 | 12,19 | 0,0 | 246 | 149 | 4.83E-05 | 0.000457123 | 0.895,0.614 | 1.0,1.0 | -0.246 |
| 11823 | XLOC_001313 | chr1 | - | 151317581 | 151317916 | 151317581 | 151317664 | 151317761 | 97 | 151317916 | 11823 | 5,1 | 4,1 | 8,5 | 0,0 | 246 | 149 | 9.92E-06 | 0.000112929 | 0.431,0.377 | 1.0,1.0 | -0.596 |
| 12490 | XLOC_041304 | chr7 | + | 131122375 | 131122756 | 131122375 | 131122446 | 131122543 | 97 | 131122756 | 12490 | 5,7 | 1,4 | 7,1 | 0,0 | 246 | 149 | 5.50E-05 | 0.000510614 | 0.752,0.515 | 1.0,1.0 | -0.367 |
| 14152 | XLOC_000269 | chr1 | + | 44680021 | 44680288 | 44680021 | 44680113 | 44680210 | 97 | 44680288 | 14152 | 17,12 | 7,14 | 69,47 | 3,1 | 246 | 149 | 2.35E-08 | 4.79E-07 | 0.595,0.342 | 0.933,0.966 | -0.481 |
| 14816 | XLOC_041705 | chr7 | - | 128034942 | 128035424 | 128034942 | 128035087 | 128035184 | 97 | 128035424 | 14816 | 8,8 | 317,357 | 122,63 | 272,209 | 246 | 149 | 0 | 0 | 0.015,0.013 | 0.214,0.154 | -0.17 |
| 4141 | XLOC_045898 | chr9 | + | 35660561 | 35660855 | 35660561 | 35660644 | 35660742 | 98 | 35660855 | 4141 | 78,83 | 118,125 | 173,158 | 151,83 | 247 | 149 | 0.000115412 | 0.00096964 | 0.285,0.286 | 0.409,0.535 | -0.186 |
| 6882 | XLOC_021471 | chr19 | - | 3613055 | 3613487 | 3613055 | 3613363 | 3613461 | 98 | 3613487 | 6882 | 8,0 | 35,20 | 41,43 | 65,44 | 247 | 149 | 4.43E-06 | 5.55E-05 | 0.121,0.0 | 0.276,0.371 | -0.263 |
| 9033 | XLOC_015040 | chr15 | - | 85189289 | 85189579 | 85189289 | 85189316 | 85189414 | 98 | 85189579 | 9033 | 42,43 | 30,31 | 92,63 | 32,21 | 247 | 149 | 0.00275577 | 0.014178409 | 0.458,0.456 | 0.634,0.644 | -0.182 |
| 13638 | XLOC_018386 | chr17 | - | 7216698 | 7217003 | 7216698 | 7216788 | 7216886 | 98 | 7217003 | 13638 | 7,5 | 191,216 | 49,40 | 190,181 | 247 | 149 | 1.77E-14 | 9.30E-13 | 0.022,0.014 | 0.135,0.118 | -0.108 |
| 14958 | XLOC_001361 | chr1 | - | 155989803 | 155990137 | 155989803 | 155989958 | 155990056 | 98 | 155990137 | 14958 | 1,1 | 1,1 | 2,2 | 1,0 | 247 | 149 | 0.002758118 | 0.014182917 | 0.376,0.376 | 0.547,1.0 | -0.398 |
| 15180 | XLOC_009345 | chr12 | - | 49435884 | 49435884 | 49435884 | 49435773 | 49435871 | 98 | 49435884 | 15180 | 7,17 | 86,107 | 36,25 | 95,83 | 247 | 149 | 0.000119221 | 0.00099808 | 0.047,0.087 | 0.186,0.154 | -0.103 |
| 4042 | XLOC_044306 | chr8 | - | 59512309 | 59512552 | 59512309 | 59512429 | 59512528 | 99 | 59512552 | 4042 | 6,5 | 88,77 | 35,17 | 86,69 | 248 | 149 | 3.98E-07 | 6.48E-06 | 0.039,0.038 | 0.196,0.129 | -0.124 |
| 4444 | XLOC_041635 | chr7 | - | 100850858 | 100851114 | 100850858 | 100851005 | 100851104 | 99 | 100851114 | 4444 | 9,6 | 0,4 | 16,14 | 0,0 | 248 | 149 | 0.008475248 | 0.0360856 | 1.0,0.474 | 1.0,1.0 | -0.263 |
| 4823 | XLOC_034487 | chr5 | + | 1802262 | 1802488 | 1802262 | 1802335 | 1802434 | 99 | 1802488 | 4823 | 6,9 | 3,2 | 7,8 | 0,0 | 248 | 149 | 9.34E-05 | 0.000809142 | 0.546,0.73 | 1.0,1.0 | -0.362 |
| 5258 | XLOC_029576 | chr3 | + | 132203206 | 132203516 | 132203206 | 132203332 | 132203431 | 99 | 132203516 | 5258 | 0,1 | 3,4 | 4,3 | 3,8 | 248 | 149 | 0.002867815 | 0.014653455 | 0.0,0.131 | 0.445,0.184 | -0.249 |
| 7799 | XLOC_018773 | chr17 | - | 79478788 | 79479168 | 79478788 | 79478829 | 79478928 | 99 | 79479168 | 7799 | 126,117 | 10,24 | 211,126 | 1,13 | 248 | 149 | 0.001471921 | 0.008433243 | 0.883,0.745 | 0.992,0.853 | -0.108 |
| 3858 | XLOC_046087 | chr9 | + | 131831006 | 131831408 | 131831006 | 131831254 | 131831354 | 100 | 131831408 | 3858 | 3,7 | 11,27 | 14,16 | 9,16 | 249 | 149 | 0.000345427 | 0.002500495 | 0.14,0.134 | 0.482,0.374 | -0.291 |
| 3991 | XLOC_044443 | chr8 | - | 144921464 | 144921721 | 144921464 | 144921580 | 144921680 | 100 | 144921721 | 3991 | 0,0 | 139,109 | 18,4 | 49,24 | 249 | 149 | 4.50E-12 | 1.70E-10 | 0.0,0.0 | 0.18,0.091 | -0.136 |
| 4984 | XLOC_034868 | chr5 | - | 36207271 | 36207519 | 36207271 | 36207367 | 36207467 | 100 | 36207519 | 4984 | 20,25 | 4,1 | 29,17 | 0,1 | 249 | 149 | 0.010592781 | 0.043400595 | 0.749,0.937 | 1.0,0.91 | -0.112 |
| 5173 | XLOC_029837 | chr3 | - | 49053251 | 49053493 | 49053251 | 49053305 | 49053405 | 100 | 49053493 | 5173 | 151,153 | 76,92 | 151,93 | 36,33 | 249 | 149 | 0.001137152 | 0.006804368 | 0.543,0.499 | 0.715,0.628 | -0.15 |
| 5891 | XLOC_026684 | chr20 | + | 2842260 | 2842579 | 2842260 | 2842350 | 2842450 | 100 | 2842579 | 5891 | 3,3 | 70,60 | 6,15 | 40,39 | 249 | 149 | 6.92E-05 | 0.000620847 | 0.025,0.029 | 0.082,0.187 | -0.108 |
| 10910 | XLOC_009356 | chr12 | - | 50196575 | 50196948 | 50196575 | 50196779 | 50196879 | 100 | 50196948 | 10910 | 6,10 | 36,26 | 46,57 | 55,42 | 249 | 149 | 1.32E-05 | 0.000144694 | 0.091,0.187 | 0.334,0.448 | -0.252 |
| 14812 | XLOC_013106 | chr14 | + | 45579096 | 45579418 | 45579096 | 45579196 | 45579296 | 100 | 45579418 | 14812 | 72,87 | 19,32 | 106,97 | 19,14 | 249 | 149 | 0.010211235 | 0.042044687 | 0.694,0.619 | 0.77,0.806 | -0.132 |
| 1931 | XLOC_028602 | chr22 | - | 24176704 | 24179341 | 24176704 | 24179149 | 24179250 | 101 | 24179341 | 1931 | 36,44 | 21,16 | 73,45 | 9,7 | 250 | 149 | 0.00033586 | 0.002445026 | 0.505,0.621 | 0.829,0.793 | -0.248 |
| 3914 | XLOC_045946 | chr9 | + | 80866823 | 80867214 | 80866823 | 80866959 | 80867060 | 101 | 80867214 | 3914 | 11,9 | 2,2 | 18,27 | 0,0 | 250 | 149 | 0.001859362 | 0.010249717 | 0.766,0.728 | 1.0,1.0 | -0.253 |
| 5702 | XLOC_027051 | chr20 | - | 57572657 | 57573009 | 57572657 | 57572808 | 57572909 | 101 | 57573009 | 5702 | 10,21 | 4,14 | 4,3 | 0,1 | 250 | 149 | 0.00164164 | 0.009259136 | 0.598,0.472 | 1.0,0.641 | -0.286 |
| 5802 | XLOC_026868 | chr20 | + | 61299150 | 61299536 | 61299150 | 61299262 | 61299363 | 101 | 61299536 | 5802 | 127,95 | 532,607 | 299,210 | 266,304 | 250 | 149 | 0 | 0 | 0.125,0.085 | 0.401,0.292 | -0.242 |
| 5809 | XLOC_026862 | chr20 | + | 60708110 | 60708531 | 60708110 | 60708244 | 60708345 | 101 | 60708531 | 5809 | 11,8 | 14,28 | 13,4 | 0,0 | 250 | 149 | 8.81E-11 | 2.72E-09 | 0.319,0.146 | 1.0,1.0 | -0.768 |
| 10123 | XLOC_005260 | chr10 | - | 75006196 | 75006557 | 75006196 | 75006344 | 75006446 | 101 | 75006557 | 10123 | 1,3 | 27,22 | 24,16 | 60,57 | 250 | 149 | 0.000476238 | 0.003303491 | 0.022,0.075 | 0.193,0.143 | -0.119 |
| 11471 | XLOC_026876 | chr20 | + | 62320859 | 62321218 | 62320859 | 62321001 | 62321102 | 101 | 62321218 | 11471 | 2,2 | 34,50 | 11,4 | 14,33 | 250 | 149 | 0.000530297 | 0.003618471 | 0.034,0.023 | 0.319,0.067 | -0.165 |
| 12391 | XLOC_026690 | chr20 | + | 3783376 | 3783619 | 3783376 | 3783455 | 3783556 | 101 | 3783619 | 12391 | 5,7 | 6,8 | 11,9 | 0,3 | 250 | 149 | 0.001353785 | 0.007877835 | 0.332,0.343 | 1.0,0.641 | -0.483 |
| 12706 | XLOC_005370 | chr10 | - | 104860419 | 104860859 | 104860419 | 104860700 | 104860801 | 101 | 104860859 | 12706 | 2,3 | 17,18 | 5,4 | 0,1 | 250 | 149 | 4.95E-08 | 9.59E-07 | 0.066,0.09 | 1.0,0.704 | -0.774 |
| 4327 | XLOC_041217 | chr7 | + | 99950619 | 99951106 | 99950619 | 99950893 | 99950995 | 102 | 99951106 | 4327 | 16,11 | 81,89 | 41,37 | 18,26 | 251 | 149 | 4.44E-16 | 2.74E-14 | 0.105,0.068 | 0.575,0.458 | -0.43 |
| 4779 | XLOC_041459 | chr7 | - | 30618621 | 30618930 | 30618621 | 30618744 | 30618846 | 102 | 30618930 | 4779 | 1,7 | 5,9 | 7,7 | 2,3 | 251 | 149 | 0.009311164 | 0.038919268 | 0.106,0.316 | 0.675,0.581 | -0.417 |

| | | | | | | | | | | | | | | | | | | | | | | |
|---|---|---|---|---|---|---|---|---|---|---|---|---|---|---|---|---|---|---|---|---|---|---|
| 7943 | XLOC_018130 | chr17 | + | 46055869 | 46056283 | 46055869 | 46056092 | 46056194 | 102 | 46056283 | 7943 | 4,1 | 0,4 | 10,2 | 0,0 | 251 | 149 | 0.007194287 | 0.03140801 | 1.0,0.129 | 1.0,1.0 | -0.435 |
| 12423 | XLOC_011393 | chr13 | + | 115002119 | 115002381 | 115002119 | 115002174 | 115002276 | 102 | 115002381 | 12423 | 17,10 | 22,22 | 36,20 | 18,9 | 251 | 149 | 0.000418064 | 0.002944794 | 0.314,0.212 | 0.543,0.569 | -0.293 |
| 14775 | XLOC_000426 | chr1 | + | 113217176 | 113217541 | 113217176 | 113217351 | 113217453 | 102 | 113217541 | 14775 | 23,52 | 77,77 | 40,35 | 37,32 | 251 | 149 | 0.002227995 | 0.011891455 | 0.151,0.286 | 0.391,0.394 | -0.174 |
| 2306 | XLOC_015047 | chr15 | - | 89870142 | 89870580 | 89870142 | 89870294 | 89870397 | 103 | 89870580 | 2306 | 27,33 | 178,194 | 63,52 | 180,118 | 252 | 149 | 9.94E-06 | 0.000113066 | 0.082,0.091 | 0.171,0.207 | -0.103 |
| 3993 | XLOC_044443 | chr8 | - | 144921907 | 144922300 | 144921907 | 144921997 | 144922100 | 103 | 144922300 | 3993 | 27,36 | 110,114 | 20,18 | 38,18 | 252 | 149 | 0.003900436 | 0.019008235 | 0.127,0.157 | 0.237,0.372 | -0.162 |
| 4136 | XLOC_047697 | chrX | + | 48339674 | 48340103 | 48339674 | 48339916 | 48340019 | 103 | 48340103 | 4136 | 33,25 | 247,279 | 93,49 | 230,168 | 252 | 149 | 1.05E-09 | 2.71E-08 | 0.073,0.05 | 0.193,0.147 | -0.108 |
| 4375 | XLOC_041073 | chr7 | + | 35919205 | 35919550 | 35919205 | 35919329 | 35919432 | 103 | 35919550 | 4375 | 3,6 | 2,2 | 4,9 | 0,1 | 252 | 149 | 0.002909565 | 0.014856034 | 0.47,0.639 | 1.0,0.842 | -0.367 |
| 5523 | XLOC_028551 | chr22 | + | 50878126 | 50878478 | 50878126 | 50878286 | 50878389 | 103 | 50878478 | 5523 | 6,1 | 82,85 | 28,9 | 52,31 | 252 | 149 | 7.42E-10 | 1.98E-08 | 0.041,0.007 | 0.241,0.147 | -0.17 |
| 11086 | XLOC_018726 | chr17 | - | 73897807 | 73898235 | 73897807 | 73897997 | 73898100 | 103 | 73898235 | 11086 | 1,1 | 9,17 | 11,6 | 5,28 | 252 | 149 | 0.001330968 | 0.007757845 | 0.062,0.034 | 0.565,0.112 | -0.29 |
| 11559 | XLOC_007764 | chr11 | - | 129754639 | 129754990 | 129754639 | 129754777 | 129754880 | 103 | 129754990 | 11559 | 2,1 | 1,7 | 14,8 | 0,4 | 252 | 149 | 0.002500437 | 0.013083982 | 0.542,0.078 | 1.0,0.542 | -0.461 |
| 12092 | XLOC_023535 | chr2 | + | 74438867 | 74439236 | 74438867 | 74438993 | 74439096 | 103 | 74439236 | 12092 | 1,3 | 5,4 | 12,9 | 2,0 | 252 | 149 | 6.71E-06 | 7.99E-05 | 0.106,0.307 | 0.78,1.0 | -0.683 |
| 3930 | XLOC_046164 | chr9 | + | 140507864 | 140508221 | 140507864 | 140507962 | 140508066 | 104 | 140508221 | 3930 | 3,2 | 193,247 | 12,29 | 80,53 | 253 | 149 | 6.44E-15 | 3.59E-13 | 0.009,0.005 | 0.081,0.244 | -0.155 |
| 6402 | XLOC_023728 | chr2 | + | 174228524 | 174228889 | 174228524 | 174228623 | 174228727 | 104 | 174228889 | 6402 | 129,156 | 41,49 | 304,228 | 6,17 | 253 | 149 | 6.66E-16 | 4.07E-14 | 0.649,0.652 | 0.968,0.888 | -0.277 |
| 7729 | XLOC_034818 | chr5 | + | 179142899 | 179143295 | 179142899 | 179143001 | 179143105 | 104 | 179143295 | 7729 | 5,5 | 2,3 | 12,5 | 0,1 | 253 | 149 | 0.000803954 | 0.005080199 | 0.596,0.495 | 1.0,0.746 | -0.328 |
| 8373 | XLOC_016786 | chr16 | - | 69402278 | 69402498 | 69402278 | 69402385 | 69402489 | 104 | 69402498 | 8373 | 2,0 | 2,6 | 11,9 | 8,5 | 253 | 149 | 0.002699773 | 0.013938283 | 0.371,0.0 | 0.447,0.515 | -0.295 |
| 9596 | XLOC_007211 | chr11 | + | 71940469 | 71940796 | 71940469 | 71940602 | 71940706 | 104 | 71940796 | 9596 | 24,28 | 770,809 | 222,62 | 368,258 | 253 | 149 | 0 | 0 | 0.018,0.02 | 0.262,0.124 | -0.174 |
| 11432 | XLOC_035150 | chr5 | - | 175956274 | 175956645 | 175956274 | 175956419 | 175956523 | 104 | 175956645 | 11432 | 4,5 | 6,3 | 12,6 | 2,0 | 253 | 149 | 0.000696788 | 0.004532815 | 0.282,0.495 | 0.779,1.0 | -0.501 |
| 12260 | XLOC_028747 | chr22 | - | 50719192 | 50719618 | 50719192 | 50719413 | 50719517 | 104 | 50719618 | 12260 | 3,4 | 5,3 | 20,10 | 2,0 | 253 | 149 | 2.83E-05 | 0.000283509 | 0.261,0.44 | 0.855,1.0 | -0.577 |
| 14770 | XLOC_000731 | chr1 | + | 220307743 | 220308019 | 220307743 | 220307852 | 220307956 | 104 | 220308019 | 14770 | 4,1 | 1,2 | 3,8 | 0,0 | 253 | 149 | 0.006267781 | 0.02809598 | 0.702,0.227 | 1.0,1.0 | -0.536 |
| 1387 | XLOC_017956 | chr17 | + | 18166399 | 18169893 | 18166399 | 18166918 | 18167023 | 105 | 18169893 | 1387 | 7,9 | 7,4 | 42,32 | 1,0 | 254 | 149 | 1.53E-08 | 3.17E-07 | 0.37,0.569 | 0.961,1.0 | -0.511 |
| 4055 | XLOC_044243 | chr8 | - | 23298905 | 23299236 | 23298905 | 23299000 | 23299105 | 105 | 23299236 | 4055 | 24,29 | 4,4 | 7,11 | 0,0 | 254 | 149 | 0.000181837 | 0.001433885 | 0.779,0.81 | 1.0,1.0 | -0.206 |
| 5336 | XLOC_029380 | chr3 | + | 39430852 | 39431107 | 39430852 | 39430880 | 39430985 | 105 | 39431107 | 5336 | 0,1 | 2,1 | 6,2 | 2,1 | 254 | 149 | 0.000613908 | 0.004087798 | 0.0,0.37 | 0.638,0.54 | -0.404 |
| 6938 | XLOC_029980 | chr3 | - | 119156434 | 119156950 | 119156434 | 119156496 | 119156601 | 105 | 119156950 | 6938 | 14,12 | 2,3 | 37,42 | 0,0 | 254 | 149 | 0.000417579 | 0.002942846 | 0.804,0.701 | 1.0,1.0 | -0.248 |
| 7734 | XLOC_018570 | chr17 | - | 41891378 | 41891695 | 41891378 | 41891459 | 41891564 | 105 | 41891695 | 7734 | 18,16 | 79,87 | 28,16 | 33,34 | 254 | 149 | 0.000160398 | 0.001282654 | 0.118,0.097 | 0.332,0.216 | -0.167 |
| 8524 | XLOC_018589 | chr17 | - | 43197539 | 43197821 | 43197539 | 43197586 | 43197691 | 105 | 43197821 | 8524 | 2,0 | 0,3 | 20,12 | 0,0 | 254 | 149 | 0.003824511 | 0.018741695 | 1.0,0.0 | 1.0,1.0 | -0.5 |
| 11524 | XLOC_000780 | chr1 | + | 231409667 | 231410126 | 231409667 | 231409808 | 231409913 | 105 | 231410126 | 11524 | 29,18 | 2,4 | 31,36 | 0,0 | 254 | 149 | 0.000318953 | 0.002341028 | 0.895,0.725 | 1.0,1.0 | -0.19 |
| 12564 | XLOC_013366 | chr14 | - | 45701711 | 45702023 | 45701711 | 45701830 | 45701935 | 105 | 45702023 | 12564 | 0,0 | 1,4 | 1,1 | 0,0 | 254 | 149 | 4.75E-09 | 1.09E-07 | 0.0,0.0 | 1.0,1.0 | -1 |
| 13133 | XLOC_007605 | chr11 | - | 67120458 | 67120760 | 67120458 | 67120548 | 67120653 | 105 | 67120760 | 13133 | 30,25 | 14,9 | 8,5 | 0,1 | 254 | 149 | 0.000478133 | 0.003311755 | 0.557,0.62 | 1.0,0.746 | -0.284 |
| 13451 | XLOC_018153 | chr17 | + | 48827861 | 48828213 | 48827861 | 48828002 | 48828107 | 105 | 48828213 | 13451 | 915,899 | 219,188 | 13,811,161 | 124,104 | 254 | 149 | 4.65E-12 | 1.74E-10 | 0.71,0.737 | 0.867,0.868 | -0.144 |
| 8658 | XLOC_001193 | chr1 | - | 92732096 | 92732298 | 92732096 | 92732150 | 92732256 | 106 | 92732298 | 8658 | 13,7 | 5,1 | 35,12 | 1,0 | 255 | 149 | 0.000184307 | 0.00144609 | 0.603,0.804 | 0.953,1.0 | -0.273 |
| 9140 | XLOC_014859 | chr15 | - | 42456912 | 42457311 | 42456912 | 42457087 | 42457193 | 106 | 42457311 | 9140 | 6,13 | 0,4 | 6,3 | 0,0 | 255 | 149 | 0.009278814 | 0.038828055 | 1.0,0.655 | 1.0,1.0 | -0.172 |
| 13714 | XLOC_016476 | chr16 | + | 71881702 | 71881974 | 71881702 | 71881806 | 71881912 | 106 | 71881974 | 13714 | 2,5 | 40,31 | 10,6 | 29,13 | 255 | 149 | 0.003374613 | 0.016888612 | 0.028,0.086 | 0.168,0.212 | -0.133 |
| 3986 | XLOC_044443 | chr8 | - | 144920638 | 144921110 | 144920638 | 144920947 | 144921054 | 107 | 144921110 | 3986 | 28,41 | 86,87 | 23,33 | 20,21 | 256 | 149 | 5.84E-06 | 7.07E-05 | 0.159,0.215 | 0.401,0.478 | -0.253 |
| 4649 | XLOC_041476 | chr7 | - | 38767940 | 38768186 | 38767940 | 38768034 | 38768141 | 107 | 38768186 | 4649 | 0,3 | 1,6 | 4,7 | 0,4 | 256 | 149 | 0.007075054 | 0.030954458 | 0.0,0.225 | 1.0,0.505 | -0.64 |
| 5014 | XLOC_041226 | chr7 | + | 100456445 | 100456783 | 100456445 | 100456564 | 100456671 | 107 | 100456783 | 5014 | 1,8 | 70,66 | 9,9 | 23,12 | 256 | 149 | 6.74E-06 | 8.00E-05 | 0.008,0.066 | 0.186,0.304 | -0.208 |
| 7627 | XLOC_034626 | chr5 | + | 96092881 | 96093198 | 96092881 | 96093040 | 96093147 | 107 | 96093198 | 7627 | 10,5 | 6,2 | 20,6 | 0,0 | 256 | 149 | 6.56E-07 | 1.01E-05 | 0.492,0.593 | 1.0,1.0 | -0.458 |
| 13067 | XLOC_017919 | chr17 | + | 8159319 | 8159725 | 8159319 | 8159477 | 8159584 | 107 | 8159725 | 13067 | 13,13 | 5,0 | 27,20 | 0,0 | 256 | 149 | 0.002073458 | 0.011211119 | 0.602,1.0 | 1.0,1.0 | -0.199 |
| 5130 | XLOC_029841 | chr3 | - | 49153246 | 49153595 | 49153246 | 49153371 | 49153479 | 108 | 49153595 | 5130 | 8,3 | 40,40 | 15,3 | 20,15 | 257 | 149 | 0.00986862 | 0.040944335 | 0.104,0.042 | 0.303,0.104 | -0.131 |
| 5447 | XLOC_028668 | chr22 | - | 38512081 | 38512417 | 38512081 | 38512218 | 38512326 | 108 | 38512417 | 5447 | 7,1 | 4,6 | 2,3 | 1,0 | 257 | 149 | 0.001160524 | 0.006914582 | 0.504,0.088 | 0.537,1.0 | -0.472 |
| 7922 | XLOC_018203 | chr17 | + | 62529051 | 62529309 | 62529051 | 62529126 | 62529234 | 108 | 62529309 | 7922 | 2,3 | 123,88 | 18,29 | 110,71 | 257 | 149 | 3.04E-10 | 8.61E-09 | 0.009,0.019 | 0.087,0.191 | -0.125 |
| 8275 | XLOC_018254 | chr17 | + | 74740403 | 74740719 | 74740403 | 74740548 | 74740656 | 108 | 74740719 | 8275 | 0,1 | 8,9 | 2,2 | 4,3 | 257 | 149 | 0.006003384 | 0.027053539 | 0.0,0.061 | 0.225,0.279 | -0.222 |
| 10844 | XLOC_009410 | chr12 | - | 56717878 | 56718277 | 56717878 | 56717969 | 56718077 | 108 | 56718277 | 10844 | 6,8 | 51,54 | 32,14 | 19,25 | 257 | 149 | 2.80E-06 | 3.67E-05 | 0.064,0.079 | 0.494,0.245 | -0.298 |
| 11991 | XLOC_007182 | chr11 | + | 67057563 | 67057917 | 67057563 | 67057684 | 67057792 | 108 | 67057917 | 11991 | 13,21 | 82,73 | 88,63 | 46,37 | 257 | 149 | 0 | 0 | 0.084,0.143 | 0.526,0.497 | -0.398 |
| 14100 | XLOC_000759 | chr1 | + | 227077458 | 227077834 | 227077458 | 227077627 | 227077735 | 108 | 227077834 | 14100 | 5,1 | 1,2 | 8,2 | 0,0 | 257 | 149 | 0.008583557 | 0.036425956 | 0.744,0.225 | 1.0,1.0 | -0.516 |
| 15241 | XLOC_000263 | chr1 | + | 44422302 | 44422648 | 44422302 | 44422373 | 44422481 | 108 | 44422648 | 15241 | 0,0 | 43,42 | 20,28 | 43,15 | 257 | 149 | 5.52E-12 | 2.05E-10 | 0.0,0.0 | 0.212,0.52 | -0.366 |
| 3220 | XLOC_017994 | chr17 | + | 27938893 | 27939291 | 27938893 | 27939024 | 27939133 | 109 | 27939291 | 3220 | 6,6 | 21,16 | 51,34 | 26,36 | 258 | 149 | 0.000206124 | 0.001591629 | 0.142,0.178 | 0.531,0.353 | -0.282 |
| 5199 | XLOC_029810 | chr3 | - | 47323629 | 47324064 | 47323629 | 47323875 | 47323984 | 109 | 47324064 | 5199 | 0,1 | 3,0 | 3,3 | 0,0 | 258 | 149 | 0.000374608 | 0.002681497 | 0.0,1.0 | 1.0,1.0 | -0.5 |
| 5783 | XLOC_026904 | chr20 | - | 2819489 | 2819912 | 2819489 | 2819728 | 2819837 | 109 | 2819912 | 5783 | 44,42 | 58,54 | 34,27 | 15,13 | 258 | 149 | 0.00018713 | 0.001464135 | 0.305,0.31 | 0.567,0.545 | -0.249 |
| 6540 | XLOC_004772 | chr10 | + | 13167784 | 13168039 | 13167784 | 13167836 | 13167945 | 109 | 13168039 | 6540 | 11,4 | 1,4 | 2,2 | 0,0 | 258 | 149 | 0.000203504 | 0.001573976 | 0.864,0.366 | 1.0,1.0 | -0.385 |
| 6981 | XLOC_021277 | chr19 | + | 47251771 | 47252141 | 47251771 | 47251922 | 47252031 | 109 | 47252141 | 6981 | 18,8 | 6,4 | 5,8 | 0,0 | 258 | 149 | 6.20E-06 | 7.44E-05 | 0.634,0.536 | 1.0,1.0 | -0.415 |
| 7897 | XLOC_018245 | chr17 | + | 73616303 | 73616580 | 73616303 | 73616437 | 73616546 | 109 | 73616580 | 7897 | 9,20 | 84,122 | 7,9 | 15,15 | 258 | 149 | 0.001748444 | 0.009760911 | 0.058,0.086 | 0.212,0.257 | -0.162 |
| 10561 | XLOC_046244 | chr9 | - | 35812893 | 35813335 | 35812893 | 35813153 | 35813262 | 109 | 35813335 | 10561 | 9,2 | 3,7 | 39,31 | 2,3 | 258 | 149 | 0.000493033 | 0.003394978 | 0.634,0.142 | 0.918,0.856 | -0.499 |
| 14332 | XLOC_023671 | chr2 | + | 135920148 | 135920421 | 135920148 | 135920208 | 135920317 | 109 | 135920421 | 14332 | 9,11 | 3,4 | 18,14 | 1,0 | 258 | 149 | 0.000137687 | 0.001124023 | 0.634,0.614 | 0.912,1.0 | -0.332 |
| 5595 | XLOC_028382 | chr22 | + | 19481849 | 19482125 | 19481849 | 19481905 | 19482015 | 110 | 19482125 | 5595 | 11,5 | 1,4 | 5,3 | 0,0 | 259 | 149 | 0.000205548 | 0.001588046 | 0.864,0.418 | 1.0,1.0 | -0.359 |
| 6255 | XLOC_024101 | chr2 | - | 74655757 | 74656066 | 74655757 | 74655864 | 74655974 | 110 | 74656066 | 6255 | 10,16 | 86,86 | 19,19 | 43,30 | 259 | 149 | 4.49E-05 | 0.000429128 | 0.063,0.097 | 0.203,0.267 | -0.155 |
| 6902 | XLOC_021444 | chr19 | - | 1235310 | 1235564 | 1235310 | 1235389 | 1235499 | 110 | 1235564 | 6902 | 0,1 | 7,9 | 3,8 | 4,14 | 259 | 149 | 0.000130378 | 0.001074913 | 0.0,0.06 | 0.301,0.247 | -0.244 |
| 8099 | XLOC_017875 | chr17 | + | 4855122 | 4855442 | 4855122 | 4855209 | 4855319 | 110 | 4855442 | 8099 | 32,29 | 16,31 | 50,26 | 11,4 | 259 | 149 | 0.000437988 | 0.003065263 | 0.535,0.35 | 0.723,0.789 | -0.314 |
| 9996 | XLOC_016854 | chr16 | - | 88801384 | 88801654 | 88801384 | 88801432 | 88801542 | 110 | 88801654 | 9996 | 10,3 | 0,8 | 8,12 | 0,0 | 259 | 149 | 0.002106853 | 0.011369919 | 1.0,0.177 | 1.0,1.0 | -0.411 |
| 11750 | XLOC_007541 | chr11 | - | 62378107 | 62378464 | 62378107 | 62378240 | 62378350 | 110 | 62378464 | 11750 | 5,7 | 13,6 | 11,10 | 1,3 | 259 | 149 | 0.000701565 | 0.00455759 | 0.181,0.402 | 0.864,0.657 | -0.469 |
| 13774 | XLOC_000827 | chr1 | - | 1277046 | 1277340 | 1277046 | 1277185 | 1277295 | 110 | 1277340 | 13774 | 25,13 | 2,2 | 20,11 | 0,0 | 259 | 149 | 0.010838292 | 0.044214095 | 0.878,0.789 | 1.0,1.0 | -0.166 |
| 4480 | XLOC_038322 | chr6 | - | 74227866 | 74228333 | 74227866 | 74227965 | 74228076 | 111 | 74228333 | 4480 | 175,177 | 948,936 | 220,193 | 0,0 | 260 | 149 | 0 | 0 | 0.096,0.098 | 1.0,1.0 | -0.903 |
| 5107 | XLOC_029867 | chr3 | - | 51994881 | 51995286 | 51994881 | 51994914 | 51995025 | 111 | 51995286 | 5107 | 0,1 | 75,51 | 5,1 | 13,11 | 260 | 149 | 0.001505542 | 0.008591019 | 0.0,0.011 | 0.181,0.05 | -0.11 |
| 6985 | XLOC_021261 | chr19 | + | 45567269 | 45567888 | 45567269 | 45567489 | 45567600 | 111 | 45567888 | 6985 | 21,27 | 3,2 | 30,32 | 0,0 | 260 | 149 | 0.002735722 | 0.01410095 | 0.8,0.886 | 1.0,1.0 | -0.157 |
| 10843 | XLOC_009410 | chr12 | - | 56717878 | 56718277 | 56717878 | 56717966 | 56718077 | 111 | 56718277 | 10843 | 6,9 | 28,27 | 33,14 | 15,5 | 260 | 149 | 6.61E-09 | 1.47E-07 | 0.109,0.16 | 0.558,0.616 | -0.452 |
| 3944 | XLOC_044466 | chr8 | - | 145737063 | 145737445 | 145737063 | 145737172 | 145737284 | 112 | 145737445 | 3944 | 6,6 | 6,4 | 23,12 | 1,2 | 261 | 149 | 0.000233082 | 0.001770745 | 0.363,0.461 | 0.929,0.774 | -0.44 |

| | | | | | | | | | | | | | | | | | | | | | | |
|---|---|---|---|---|---|---|---|---|---|---|---|---|---|---|---|---|---|---|---|---|---|---|
| 4946 | XLOC_032302 | chr4 | + | 39274599 | 39274808 | 39274599 | 39274681 | 39274793 | 112 | 39274808 | 4946 | 27,31 | 5,8 | 33,16 | 0,0 | 261 | 149 | 1.58E-06 | 2.19E-05 | 0.755,0.689 | 1.0,1.0 | -0.278 |
| 5955 | XLOC_041630 | chr7 | - | 100281863 | 100282239 | 100281863 | 100281960 | 100282072 | 112 | 100282239 | 5955 | 29,23 | 174,177 | 95,74 | 156,99 | 261 | 149 | 1.25E-14 | 6.84E-13 | 0.087,0.069 | 0.258,0.299 | -0.2 |
| 6083 | XLOC_016187 | chr16 | + | 716654 | 716989 | 716654 | 716792 | 716904 | 112 | 716989 | 6083 | 51,38 | 182,153 | 108,45 | 74,51 | 261 | 149 | 4.21E-11 | 1.39E-09 | 0.138,0.124 | 0.454,0.335 | -0.264 |
| 6618 | XLOC_001199 | chr1 | - | 93804745 | 93805236 | 93804745 | 93805093 | 93805205 | 112 | 93805236 | 6618 | 3,1 | 37,18 | 5,2 | 4,3 | 261 | 149 | 0.000802337 | 0.005072247 | 0.044,0.031 | 0.416,0.276 | -0.308 |
| 10028 | XLOC_007152 | chr11 | + | 65293391 | 65293818 | 65293391 | 65293514 | 65293626 | 112 | 65293818 | 10028 | 5,3 | 8,0 | 11,20 | 0,0 | 261 | 149 | 0.000967457 | 0.005932869 | 0.263,1.0 | 1.0,1.0 | -0.369 |
| 10813 | XLOC_001315 | chr1 | - | 151380808 | 151381304 | 151380808 | 151381057 | 151381169 | 112 | 151381304 | 10813 | 4,6 | 103,144 | 57,28 | 83,73 | 261 | 149 | 1.11E-16 | 7.47E-15 | 0.022,0.023 | 0.282,0.18 | -0.208 |
| 11954 | XLOC_043958 | chr8 | + | 21959486 | 21959826 | 21959486 | 21959573 | 21959685 | 112 | 21959826 | 11954 | 22,8 | 3,16 | 13,11 | 0,1 | 261 | 149 | 0.000182047 | 0.001433946 | 0.807,0.222 | 1.0,0.863 | -0.417 |
| 15003 | XLOC_007183 | chr11 | + | 67074776 | 67075208 | 67074776 | 67074907 | 67075019 | 112 | 67075208 | 15003 | 106,95 | 56,68 | 94,68 | 33,14 | 261 | 149 | 0.001759139 | 0.00980826 | 0.519,0.444 | 0.619,0.735 | -0.196 |
| 4203 | XLOC_041585 | chr7 | - | 91874909 | 91875183 | 91874909 | 91874990 | 91875103 | 113 | 91875183 | 4203 | 36,36 | 10,0 | 21,21 | 0,0 | 262 | 149 | 0.000392353 | 0.002796922 | 0.672,1.0 | 1.0,1.0 | -0.164 |
| 5132 | XLOC_029841 | chr3 | - | 49153883 | 49154288 | 49153883 | 49154040 | 49154153 | 113 | 49154288 | 5132 | 11,15 | 10,18 | 29,9 | 9,2 | 262 | 149 | 0.001620761 | 0.009167003 | 0.385,0.322 | 0.647,0.719 | -0.33 |
| 7594 | XLOC_018718 | chr17 | - | 73500275 | 73500771 | 73500275 | 73500387 | 73500500 | 113 | 73500771 | 7594 | 0,0 | 36,13 | 8,4 | 27,26 | 262 | 149 | 4.32E-05 | 0.000414753 | 0.0,0.0 | 0.144,0.08 | -0.112 |
| 8741 | XLOC_016191 | chr16 | + | 778796 | 779088 | 778796 | 778895 | 779008 | 113 | 779088 | 8741 | 79,84 | 317,314 | 135,79 | 222,145 | 262 | 149 | 3.19E-07 | 5.28E-06 | 0.124,0.132 | 0.257,0.237 | -0.119 |
| 9651 | XLOC_016566 | chr16 | - | 1417656 | 1418115 | 1417656 | 1417859 | 1417972 | 113 | 1418115 | 9651 | 25,31 | 9,5 | 67,35 | 0,6 | 262 | 149 | 0.008960885 | 0.037721185 | 0.612,0.779 | 1.0,0.768 | -0.189 |
| 12245 | XLOC_018124 | chr17 | + | 45696374 | 45696688 | 45696374 | 45696530 | 45696643 | 113 | 45696688 | 12245 | 64,61 | 21,23 | 47,25 | 0,0 | 262 | 149 | 3.10E-11 | 1.04E-09 | 0.634,0.601 | 1.0,1.0 | -0.383 |
| 6312 | XLOC_023989 | chr2 | - | 32168073 | 32168452 | 32168073 | 32168256 | 32168370 | 114 | 32168452 | 6312 | 0,1 | 3,0 | 2,1 | 0,0 | 263 | 149 | 0.000354196 | 0.0025535 | 0.0,1.0 | 1.0,1.0 | -0.5 |
| 7402 | XLOC_024042 | chr2 | - | 55792037 | 55792347 | 55792037 | 55792177 | 55792291 | 114 | 55792347 | 7402 | 3,4 | 1,2 | 9,6 | 0,0 | 263 | 149 | 0.011635444 | 0.047071719 | 0.63,0.531 | 1.0,1.0 | -0.419 |
| 8315 | XLOC_038369 | chr6 | - | 97717777 | 97718070 | 97717777 | 97717868 | 97717982 | 114 | 97718070 | 8315 | 0,0 | 2,6 | 1,4 | 7,1 | 263 | 149 | 0.003696674 | 0.018229095 | 0.0,0.0 | 0.075,0.694 | -0.384 |
| 9064 | XLOC_014986 | chr15 | - | 75134620 | 75135015 | 75134620 | 75134761 | 75134875 | 114 | 75135015 | 9064 | 8,3 | 4,8 | 17,6 | 1,1 | 263 | 149 | 0.000174108 | 0.00137832 | 0.531,0.175 | 0.906,0.773 | -0.487 |
| 9513 | XLOC_000212 | chr1 | + | 36759451 | 36759842 | 36759451 | 36759536 | 36759650 | 114 | 36759842 | 9513 | 35,57 | 73,64 | 72,68 | 38,35 | 263 | 149 | 1.58E-05 | 0.000169182 | 0.214,0.335 | 0.518,0.524 | -0.246 |
| 11853 | XLOC_007438 | chr11 | - | 17035531 | 17035955 | 17035531 | 17035748 | 17035862 | 114 | 17035955 | 11853 | 0,4 | 28,14 | 5,9 | 23,15 | 263 | 149 | 0.009441115 | 0.039344015 | 0.0,0.139 | 0.11,0.254 | -0.112 |
| 12730 | XLOC_017993 | chr17 | + | 27896335 | 27896643 | 27896335 | 27896377 | 27896491 | 114 | 27896643 | 12730 | 8,15 | 3,1 | 10,2 | 0,0 | 263 | 149 | 0.005940701 | 0.026813845 | 0.602,0.895 | 1.0,1.0 | -0.252 |
| 13016 | XLOC_018381 | chr17 | - | 7133126 | 7133456 | 7133126 | 7133262 | 7133376 | 114 | 7133456 | 13016 | 80,34 | 85,94 | 53,40 | 49,24 | 263 | 149 | 0.01155704 | 0.046781333 | 0.348,0.17 | 0.38,0.486 | -0.174 |
| 13616 | XLOC_001082 | chr1 | - | 45798768 | 45799169 | 45798768 | 45798842 | 45798956 | 114 | 45799169 | 13616 | 26,27 | 78,48 | 37,20 | 31,15 | 263 | 149 | 0.000126129 | 0.001044761 | 0.159,0.242 | 0.403,0.43 | -0.216 |
| 14386 | XLOC_007445 | chr11 | - | 18320167 | 18320517 | 18320167 | 18320224 | 18320338 | 114 | 18320517 | 14386 | 4,2 | 4,0 | 2,1 | 0,0 | 263 | 149 | 0.005303998 | 0.024400121 | 0.362,1.0 | 1.0,1.0 | -0.319 |
| 1104 | XLOC_021261 | chr19 | + | 45567269 | 45567888 | 45567269 | 45567485 | 45567600 | 115 | 45567888 | 1104 | 46,56 | 117,123 | 76,72 | 74,58 | 264 | 149 | 4.77E-07 | 7.63E-06 | 0.182,0.204 | 0.367,0.412 | -0.196 |
| 4955 | XLOC_032289 | chr4 | + | 25396292 | 25396535 | 25396292 | 25396335 | 25396450 | 115 | 25396535 | 4955 | 8,11 | 50,56 | 35,27 | 53,26 | 264 | 149 | 1.08E-06 | 1.56E-05 | 0.083,0.1 | 0.272,0.37 | -0.23 |
| 5424 | XLOC_028745 | chr22 | - | 50694036 | 50694310 | 50694036 | 50694123 | 50694238 | 115 | 50694310 | 5424 | 21,18 | 69,81 | 37,17 | 45,38 | 264 | 149 | 0.001891574 | 0.010374641 | 0.147,0.111 | 0.317,0.202 | -0.131 |
| 6211 | XLOC_024188 | chr2 | - | 114384349 | 114384713 | 114384349 | 114384499 | 114384614 | 115 | 114384713 | 6211 | 38,40 | 2,4 | 42,43 | 0,0 | 264 | 149 | 0.001848604 | 0.010210463 | 0.915,0.849 | 1.0,1.0 | -0.118 |
| 7233 | XLOC_038282 | chr6 | - | 43536217 | 43536474 | 43536217 | 43536332 | 43536447 | 115 | 43536474 | 7233 | 0,2 | 1,2 | 8,1 | 0,0 | 264 | 149 | 0.00010575 | 0.00090079 | 0.0,0.361 | 1.0,1.0 | -0.82 |
| 10514 | XLOC_023660 | chr2 | + | 131116840 | 131117219 | 131116840 | 131117065 | 131117180 | 115 | 131117219 | 10514 | 8,7 | 13,13 | 10,9 | 4,5 | 264 | 149 | 0.009199864 | 0.038531934 | 0.258,0.233 | 0.585,0.504 | -0.299 |
| 11173 | XLOC_009592 | chr12 | - | 122967825 | 122968137 | 122967825 | 122967891 | 122968006 | 115 | 122968137 | 11173 | 13,31 | 65,67 | 29,30 | 31,26 | 264 | 149 | 5.35E-05 | 0.000498194 | 0.101,0.207 | 0.346,0.394 | -0.216 |
| 11199 | XLOC_008857 | chr12 | + | 44164840 | 44165168 | 44164840 | 44164907 | 44165022 | 115 | 44165168 | 11199 | 4,1 | 2,1 | 4,2 | 0,0 | 264 | 149 | 0.006019465 | 0.027117353 | 0.53,0.361 | 1.0,1.0 | -0.554 |
| 12177 | XLOC_021540 | chr19 | - | 10676390 | 10676705 | 10676390 | 10676466 | 10676581 | 115 | 10676705 | 12177 | 3,6 | 89,82 | 16,8 | 41,38 | 264 | 149 | 8.90E-06 | 0.000102548 | 0.019,0.04 | 0.18,0.106 | -0.113 |
| 13437 | XLOC_009179 | chr12 | + | 132391436 | 132391707 | 132391436 | 132391469 | 132391584 | 115 | 132391707 | 13437 | 12,13 | 2,3 | 11,2 | 1,0 | 264 | 149 | 0.006404303 | 0.028570767 | 0.772,0.71 | 0.861,1.0 | -0.19 |
| 13779 | XLOC_004752 | chr10 | + | 1169922 | 1170292 | 1169922 | 1170042 | 1170157 | 115 | 1170292 | 13779 | 3,3 | 3,5 | 9,2 | 0,1 | 264 | 149 | 0.000167432 | 0.001330689 | 0.361,0.253 | 1.0,0.53 | -0.458 |
| 2480 | XLOC_007374 | chr11 | - | 799814 | 800244 | 799814 | 800014 | 800130 | 116 | 800244 | 2480 | 39,48 | 31,21 | 49,16 | 9,3 | 265 | 149 | 0.001498231 | 0.00856314 | 0.414,0.562 | 0.754,0.75 | -0.264 |
| 2928 | XLOC_023538 | chr2 | + | 74717151 | 74717600 | 74717151 | 74717254 | 74717370 | 116 | 74717600 | 2928 | 25,20 | 66,51 | 31,24 | 37,15 | 265 | 149 | 0.000969619 | 0.005943546 | 0.176,0.181 | 0.32,0.474 | -0.219 |
| 4583 | XLOC_037801 | chr6 | + | 35253904 | 35254176 | 35253904 | 35253985 | 35254101 | 116 | 35254176 | 4583 | 8,8 | 4,1 | 7,7 | 0,0 | 265 | 149 | 0.000122014 | 0.00101784 | 0.529,0.818 | 1.0,1.0 | -0.327 |
| 5432 | XLOC_028716 | chr22 | - | 43032484 | 43032852 | 43032484 | 43032604 | 43032720 | 116 | 43032852 | 5432 | 5,6 | 2,2 | 12,3 | 0,0 | 265 | 149 | 0.000385778 | 0.002754469 | 0.584,0.628 | 1.0,1.0 | -0.394 |
| 8782 | XLOC_016652 | chr16 | - | 18863374 | 18863784 | 18863374 | 18863523 | 18863639 | 116 | 18863784 | 8782 | 2,0 | 2,1 | 4,3 | 0,0 | 265 | 149 | 0.000103623 | 0.000883734 | 0.36,0.0 | 1.0,1.0 | -0.82 |
| 14948 | XLOC_000082 | chr1 | + | 12041866 | 12042171 | 12041866 | 12041910 | 12042026 | 116 | 12042171 | 14948 | 12,9 | 1,3 | 13,12 | 1,0 | 265 | 149 | 0.011128341 | 0.045214724 | 0.871,0.628 | 0.88,1.0 | -0.19 |
| 231 | XLOC_040986 | chr7 | + | 2394158 | 2395055 | 2394158 | 2394751 | 2394868 | 117 | 2395055 | 231 | 812,690 | 138,123 | 16,311,318 | 103,70 | 266 | 149 | 2.02E-14 | 1.05E-12 | 0.767,0.759 | 0.899,0.913 | -0.143 |
| 5050 | XLOC_030173 | chr3 | - | 197024056 | 197024297 | 197024056 | 197024106 | 197024223 | 117 | 197024297 | 5050 | 1,1 | 18,5 | 6,2 | 6,10 | 266 | 149 | 0.007830064 | 0.03377642 | 0.03,0.101 | 0.359,0.101 | -0.164 |
| 5498 | XLOC_028577 | chr22 | - | 20102997 | 20103303 | 20102997 | 20103077 | 20103194 | 117 | 20103303 | 5498 | 9,18 | 0,5 | 40,25 | 0,0 | 266 | 149 | 0.001682037 | 0.009456771 | 1.0,0.668 | 1.0,1.0 | -0.166 |
| 8387 | XLOC_016188 | chr16 | + | 720280 | 720773 | 720280 | 720557 | 720674 | 117 | 720773 | 8387 | 17,7 | 239,226 | 40,35 | 165,122 | 266 | 149 | 3.90E-11 | 1.30E-09 | 0.038,0.017 | 0.12,0.138 | -0.102 |
| 9483 | XLOC_044454 | chr8 | - | 145008169 | 145008634 | 145008169 | 145008368 | 145008485 | 117 | 145008634 | 9483 | 26,17 | 7,0 | 43,43 | 0,1 | 266 | 149 | 0.000960158 | 0.005898355 | 0.675,1.0 | 1.0,0.96 | -0.142 |
| 11711 | XLOC_007548 | chr11 | - | 62563752 | 62564039 | 62563752 | 62563814 | 62563931 | 117 | 62564039 | 11711 | 6,2 | 1,4 | 2,4 | 0,0 | 266 | 149 | 3.16E-05 | 0.000314069 | 0.771,0.219 | 1.0,1.0 | -0.505 |
| 13217 | XLOC_021801 | chr19 | - | 48752779 | 48753135 | 48752779 | 48752890 | 48753007 | 117 | 48753135 | 13217 | 1,0 | 42,38 | 8,16 | 47,26 | 266 | 149 | 1.90E-07 | 3.28E-06 | 0.013,0.0 | 0.087,0.256 | -0.165 |
| 1932 | XLOC_028602 | chr22 | - | 24180897 | 24181318 | 24180897 | 24180963 | 24181081 | 118 | 24181318 | 1932 | 20,11 | 26,27 | 63,56 | 38,20 | 267 | 149 | 7.92E-05 | 0.000699459 | 0.3,0.185 | 0.481,0.61 | -0.303 |
| 7640 | XLOC_018673 | chr17 | - | 62499920 | 62500205 | 62499920 | 62499986 | 62500104 | 118 | 62500205 | 7640 | 19,14 | 1,5 | 48,20 | 0,0 | 267 | 149 | 0.000101547 | 0.000869705 | 0.914,0.61 | 1.0,1.0 | -0.238 |
| 9912 | XLOC_028602 | chr22 | - | 24180538 | 24180960 | 24180538 | 24180632 | 24180750 | 118 | 24180960 | 9912 | 12,18 | 28,30 | 40,30 | 29,22 | 267 | 149 | 0.000882379 | 0.005489799 | 0.193,0.251 | 0.435,0.432 | -0.211 |
| 9945 | XLOC_011394 | chr13 | + | 115051733 | 115052104 | 115051733 | 115051875 | 115051993 | 118 | 115052104 | 9945 | 88,99 | 204,203 | 103,89 | 104,125 | 267 | 149 | 0.000373612 | 0.00267784 | 0.194,0.214 | 0.356,0.284 | -0.116 |
| 11319 | XLOC_021921 | chr19 | - | 59080643 | 59080966 | 59080643 | 59080764 | 59080882 | 118 | 59080966 | 11319 | 15,11 | 99,101 | 19,21 | 53,39 | 267 | 149 | 4.06E-05 | 0.000393095 | 0.078,0.057 | 0.167,0.231 | -0.132 |
| 13969 | XLOC_000695 | chr1 | + | 206905183 | 206905429 | 206905183 | 206905264 | 206905382 | 118 | 206905429 | 13969 | 11,19 | 26,34 | 45,30 | 0,3 | 267 | 149 | 0 | 0 | 0.191,0.238 | 1.0,0.848 | -0.709 |
| 5133 | XLOC_029841 | chr3 | - | 49153883 | 49154288 | 49153883 | 49154040 | 49154159 | 119 | 49154288 | 5133 | 11,19 | 87,93 | 29,10 | 44,46 | 268 | 149 | 0.006537417 | 0.029097996 | 0.066,0.102 | 0.268,0.108 | -0.104 |
| 6826 | XLOC_021606 | chr19 | - | 17438246 | 17438677 | 17438246 | 17438378 | 17438497 | 119 | 17438677 | 6826 | 18,15 | 34,37 | 23,14 | 14,13 | 268 | 149 | 0.00205134 | 0.011112803 | 0.227,0.184 | 0.477,0.375 | -0.22 |
| 14274 | XLOC_029428 | chr3 | + | 49737659 | 49737913 | 49737659 | 49737785 | 49737904 | 119 | 49737913 | 14274 | 10,8 | 90,95 | 90,35 | 113,96 | 268 | 149 | 1.68E-09 | 4.19E-08 | 0.058,0.045 | 0.307,0.169 | -0.186 |
| 14371 | XLOC_047875 | chrX | + | 152959359 | 152959713 | 152959359 | 152959472 | 152959591 | 119 | 152959713 | 14371 | 21,19 | 10,7 | 21,34 | 3,1 | 268 | 149 | 0.000486918 | 0.003362709 | 0.539,0.601 | 0.796,0.95 | -0.303 |
| 2107 | XLOC_046244 | chr9 | - | 35812893 | 35813335 | 35812893 | 35813142 | 35813262 | 120 | 35813335 | 2107 | 19,8 | 195,179 | 63,51 | 81,37 | 269 | 149 | 0 | 0 | 0.051,0.024 | 0.301,0.433 | -0.33 |
| 3966 | XLOC_044462 | chr8 | - | 145651257 | 145651591 | 145651257 | 145651442 | 145651562 | 120 | 145651591 | 3966 | 46,20 | 0,6 | 29,27 | 0,0 | 269 | 149 | 0.001990356 | 0.010832292 | 1.0,0.649 | 1.0,1.0 | -0.175 |
| 5909 | XLOC_044184 | chr8 | + | 145692119 | 145692493 | 145692119 | 145692221 | 145692341 | 120 | 145692493 | 5909 | 1,0 | 80,77 | 0,10 | 25,17 | 269 | 149 | 8.92E-05 | 0.000777467 | 0.007,0.0 | 0.0,0.246 | -0.119 |
| 7239 | XLOC_028478 | chr22 | + | 38150922 | 38151160 | 38150922 | 38150991 | 38151111 | 120 | 38151160 | 7239 | 3,5 | 3,3 | 2,4 | 1,0 | 269 | 149 | 0.00221871 | 0.011851303 | 0.356,0.48 | 0.526,1.0 | -0.345 |
| 11721 | XLOC_007546 | chr11 | - | 62482780 | 62483079 | 62482780 | 62482851 | 62482971 | 120 | 62483079 | 11721 | 19,13 | 732,697 | 142,156 | 627,500 | 269 | 149 | 0 | 0 | 0.014,0.01 | 0.111,0.147 | -0.117 |
| 13696 | XLOC_009383 | chr12 | - | 53877132 | 53877549 | 53877132 | 53877270 | 53877390 | 120 | 53877549 | 13696 | 1,13 | 22,27 | 7,5 | 0,2 | 269 | 149 | 3.00E-05 | 0.000298742 | 0.025,0.211 | 1.0,0.581 | -0.672 |
| 7254 | XLOC_014977 | chr15 | - | 73855576 | 73855924 | 73855576 | 73855598 | 73855719 | 121 | 73855924 | 7254 | 16,11 | 6,12 | 30,33 | 0,2 | 270 | 149 | 3.57E-07 | 5.86E-06 | 0.595,0.336 | 1.0,0.901 | -0.485 |

| | | | | | | | | | | | | | | | | | | | | | | |
|---|---|---|---|---|---|---|---|---|---|---|---|---|---|---|---|---|---|---|---|---|---|---|
| 8772 | XLOC_018408 | chr17 | - | 8113271 | 8113567 | 8113271 | 8113373 | 8113494 | 121 | 8113567 | 8772 | 39,39 | 3,11 | 45,53 | 0,0 | 270 | 149 | 1.01E-06 | 1.48E-05 | 0.878,0.662 | 1.0,1.0 | -0.23 |
| 2125 | XLOC_021249 | chr19 | + | 44660404 | 44666933 | 44660404 | 44660551 | 44660673 | 122 | 44666933 | 2125 | 25,18 | 4,1 | 50,42 | 0,0 | 271 | 149 | 0.001832486 | 0.010149096 | 0.775,0.908 | 1.0,1.0 | -0.158 |
| 2631 | XLOC_007559 | chr11 | - | 63991571 | 63991813 | 63991571 | 63991634 | 63991756 | 122 | 63991813 | 2631 | 113,75 | 16,11 | 57,43 | 3,2 | 271 | 149 | 0.007649336 | 0.033093355 | 0.795,0.789 | 0.913,0.922 | -0.125 |
| 9724 | XLOC_024080 | chr2 | - | 70520509 | 70520972 | 70520509 | 70520627 | 70520749 | 122 | 70520972 | 9724 | 5,3 | 1,9 | 11,14 | 0,0 | 271 | 149 | 1.21E-06 | 1.73E-05 | 0.733,0.155 | 1.0,1.0 | -0.556 |
| 12267 | XLOC_028747 | chr22 | - | 50721139 | 50721612 | 50721139 | 50721310 | 50721432 | 122 | 50721612 | 12267 | 19,13 | 24,14 | 32,20 | 9,11 | 271 | 149 | 0.003967757 | 0.019267803 | 0.303,0.338 | 0.662,0.5 | -0.26 |
| 12547 | XLOC_013224 | chr14 | + | 89034201 | 89034497 | 89034201 | 89034260 | 89034382 | 122 | 89034497 | 12547 | 11,9 | 3,3 | 2,4 | 0,1 | 271 | 149 | 0.005958872 | 0.026883872 | 0.668,0.623 | 1.0,0.687 | -0.198 |
| 13275 | XLOC_000373 | chr1 | + | 93586105 | 93586451 | 93586105 | 93586229 | 93586351 | 122 | 93586451 | 13275 | 23,22 | 5,3 | 53,41 | 3,0 | 271 | 149 | 0.004266732 | 0.020350913 | 0.717,0.801 | 0.907,1.0 | -0.195 |
| 4102 | XLOC_046107 | chr9 | + | 134311845 | 134312282 | 134311845 | 134312109 | 134312232 | 123 | 134312282 | 4102 | 3,1 | 1,4 | 8,1 | 1,0 | 272 | 149 | 0.000309716 | 0.002275815 | 0.622,0.12 | 0.814,1.0 | -0.536 |
| 4384 | XLOC_041008 | chr7 | + | 6681455 | 6681739 | 6681455 | 6681593 | 6681716 | 123 | 6681739 | 4384 | 15,11 | 0,4 | 85,52 | 0,0 | 272 | 149 | 0.011942475 | 0.048134583 | 1.0,0.601 | 1.0,1.0 | -0.2 |
| 7307 | XLOC_014730 | chr15 | + | 79178482 | 79178737 | 79178482 | 79178569 | 79178692 | 123 | 79178737 | 7307 | 3,3 | 7,3 | 9,8 | 0,3 | 272 | 149 | 0.001707574 | 0.009577469 | 0.19,0.354 | 1.0,0.594 | -0.525 |
| 8254 | XLOC_020871 | chr19 | + | 1358361 | 1358808 | 1358361 | 1358463 | 1358586 | 123 | 1358808 | 8254 | 9,12 | 57,46 | 21,12 | 14,12 | 272 | 149 | 6.39E-07 | 9.90E-06 | 0.08,0.125 | 0.451,0.354 | -0.3 |
| 13745 | XLOC_004988 | chr10 | + | 97816649 | 97817004 | 97816649 | 97816757 | 97816880 | 123 | 97817004 | 13745 | 2,0 | 7,8 | 9,10 | 6,4 | 272 | 149 | 9.82E-06 | 0.000112123 | 0.135,0.0 | 0.451,0.578 | -0.447 |
| 14927 | XLOC_017909 | chr17 | + | 7479628 | 7480010 | 7479628 | 7479718 | 7479841 | 123 | 7480010 | 14927 | 38,25 | 14,15 | 71,62 | 10,10 | 272 | 149 | 0.001319777 | 0.007702151 | 0.598,0.477 | 0.795,0.773 | -0.247 |
| 7352 | XLOC_000997 | chr1 | - | 33263192 | 33263444 | 33263192 | 33263239 | 33263363 | 124 | 33263444 | 7352 | 3,2 | 6,5 | 9,1 | 0,0 | 273 | 149 | 1.03E-08 | 2.19E-07 | 0.214,0.179 | 1.0,1.0 | -0.803 |
| 7521 | XLOC_018791 | chr17 | - | 80037685 | 80038114 | 80037685 | 80037891 | 80038015 | 124 | 80038114 | 7521 | 34,28 | 10,7 | 39,30 | 0,0 | 273 | 149 | 2.20E-08 | 4.51E-07 | 0.65,0.686 | 1.0,1.0 | -0.332 |
| 10332 | XLOC_011260 | chr13 | + | 20235837 | 20236217 | 20235837 | 20235978 | 20236102 | 124 | 20236217 | 10332 | 3,3 | 11,5 | 1,5 | 1,0 | 273 | 149 | 0.000664302 | 0.004359638 | 0.13,0.247 | 0.353,1.0 | -0.488 |
| 11940 | XLOC_014807 | chr15 | - | 23311641 | 23312308 | 23311641 | 23311912 | 23312036 | 124 | 23312308 | 11940 | 52,64 | 10,0 | 48,29 | 1,0 | 273 | 149 | 0.002132119 | 0.011493099 | 0.739,1.0 | 0.963,1.0 | -0.112 |
| 11953 | XLOC_043958 | chr8 | + | 21959224 | 21959514 | 21959224 | 21959362 | 21959486 | 124 | 21959514 | 11953 | 15,30 | 12,7 | 14,24 | 0,3 | 273 | 149 | 0.002677192 | 0.01384474 | 0.406,0.701 | 1.0,0.814 | -0.354 |
| 12891 | XLOC_018545 | chr17 | - | 40139608 | 40139928 | 40139608 | 40139712 | 40139836 | 124 | 40139928 | 12891 | 6,8 | 11,6 | 26,14 | 5,6 | 273 | 149 | 0.005785484 | 0.026222205 | 0.229,0.421 | 0.739,0.56 | -0.324 |
| 14776 | XLOC_000426 | chr1 | + | 113217318 | 113217541 | 113217318 | 113217351 | 113217475 | 124 | 113217541 | 14776 | 11,7 | 8,12 | 10,12 | 0,4 | 273 | 149 | 0.003524377 | 0.017483238 | 0.429,0.241 | 1.0,0.621 | -0.476 |
| 10160 | XLOC_007440 | chr11 | - | 17112880 | 17113197 | 17112880 | 17113006 | 17113131 | 125 | 17113197 | 10160 | 30,20 | 6,2 | 41,23 | 2,0 | 274 | 149 | 0.004485476 | 0.021208026 | 0.731,0.845 | 0.918,1.0 | -0.171 |
| 12969 | XLOC_045959 | chr9 | + | 90342275 | 90342644 | 90342275 | 90342383 | 90342508 | 125 | 90342644 | 12969 | 10,13 | 3,4 | 10,6 | 0,0 | 274 | 149 | 2.17E-05 | 0.000223622 | 0.644,0.639 | 1.0,1.0 | -0.359 |
| 13849 | XLOC_007153 | chr11 | + | 65338052 | 65338441 | 65338052 | 65338231 | 65338356 | 125 | 65338441 | 13849 | 1,0 | 4,8 | 6,4 | 9,13 | 274 | 149 | 0.001486136 | 0.00850778 | 0.12,0.0 | 0.266,0.143 | -0.145 |
| 1962 | XLOC_023611 | chr2 | + | 105959304 | 105959680 | 105959304 | 105959431 | 105959557 | 126 | 105959680 | 1962 | 292,320 | 59,78 | 341,264 | 15,20 | 275 | 149 | 5.89E-11 | 1.88E-09 | 0.728,0.69 | 0.925,0.877 | -0.192 |
| 7624 | XLOC_018692 | chr17 | - | 67249716 | 67250010 | 67249716 | 67249808 | 67249934 | 126 | 67250010 | 7624 | 2,0 | 18,17 | 7,1 | 8,5 | 275 | 149 | 0.001047772 | 0.006348212 | 0.057,0.0 | 0.322,0.098 | -0.182 |
| 8350 | XLOC_016862 | chr16 | - | 89877114 | 89877465 | 89877114 | 89877210 | 89877336 | 126 | 89877465 | 8350 | 20,18 | 225,264 | 50,48 | 140,90 | 275 | 149 | 6.22E-15 | 3.48E-13 | 0.046,0.036 | 0.162,0.224 | -0.152 |
| 10500 | XLOC_000546 | chr1 | + | 155279854 | 155280244 | 155279854 | 155279996 | 155280122 | 126 | 155280244 | 10500 | 45,48 | 3,3 | 53,36 | 0,0 | 275 | 149 | 0.002670828 | 0.013821107 | 0.89,0.897 | 1.0,1.0 | -0.107 |
| 4434 | XLOC_038498 | chr6 | - | 165831840 | 165832217 | 165831840 | 165831995 | 165832122 | 127 | 165832217 | 4434 | 0,3 | 16,14 | 10,6 | 3,1 | 276 | 149 | 1.08E-08 | 2.29E-07 | 0.0,0.104 | 0.643,0.764 | -0.651 |
| 4708 | XLOC_035086 | chr5 | - | 141009426 | 141009687 | 141009426 | 141009483 | 141009610 | 127 | 141009687 | 4708 | 14,36 | 135,112 | 61,51 | 98,55 | 276 | 149 | 3.52E-08 | 6.93E-07 | 0.053,0.148 | 0.252,0.334 | -0.193 |
| 6727 | XLOC_020849 | chr19 | + | 804531 | 804915 | 804531 | 804702 | 804829 | 127 | 804915 | 6727 | 31,22 | 1,4 | 36,30 | 0,0 | 276 | 149 | 0.003537528 | 0.017536158 | 0.944,0.748 | 1.0,1.0 | -0.154 |
| 7467 | XLOC_017854 | chr17 | + | 2267352 | 2267670 | 2267352 | 2267477 | 2267604 | 127 | 2267670 | 7467 | 14,5 | 2,4 | 13,11 | 0,0 | 276 | 149 | 2.30E-05 | 0.000235905 | 0.791,0.403 | 1.0,1.0 | -0.403 |
| 8026 | XLOC_014991 | chr15 | - | 75651960 | 75652310 | 75651960 | 75652116 | 75652243 | 127 | 75652310 | 8026 | 7,9 | 150,139 | 47,30 | 71,84 | 276 | 149 | 2.00E-14 | 1.04E-12 | 0.025,0.034 | 0.263,0.162 | -0.183 |
| 8997 | XLOC_009231 | chr12 | - | 7079098 | 7079443 | 7079098 | 7079231 | 7079358 | 127 | 7079443 | 8997 | 18,9 | 2,6 | 37,36 | 1,1 | 276 | 149 | 0.000814729 | 0.005136795 | 0.829,0.447 | 0.952,0.951 | -0.314 |
| 14716 | XLOC_000760 | chr1 | + | 227171252 | 227171697 | 227171252 | 227171334 | 227171461 | 127 | 227171697 | 14716 | 34,10 | 71,80 | 70,41 | 105,56 | 276 | 149 | 0.000653555 | 0.004307117 | 0.205,0.063 | 0.265,0.283 | -0.14 |
| 3187 | XLOC_021440 | chr19 | - | 1011578 | 1011797 | 1011578 | 1011611 | 1011739 | 128 | 1011797 | 3187 | 199,152 | 86,76 | 231,185 | 54,52 | 277 | 149 | 0.000623987 | 0.0041426 | 0.555,0.518 | 0.697,0.657 | -0.141 |
| 4333 | XLOC_041200 | chr7 | + | 98988526 | 98988876 | 98988526 | 98988672 | 98988800 | 128 | 98988876 | 4333 | 8,6 | 0,5 | 16,19 | 0,0 | 277 | 149 | 0.002293221 | 0.012184783 | 1.0,0.392 | 1.0,1.0 | -0.304 |
| 4995 | XLOC_037824 | chr6 | + | 41751199 | 41751532 | 41751199 | 41751343 | 41751471 | 128 | 41751532 | 4995 | 38,43 | 1,5 | 18,15 | 0,0 | 277 | 149 | 0.002224544 | 0.011877971 | 0.953,0.822 | 1.0,1.0 | -0.113 |
| 6403 | XLOC_023728 | chr2 | + | 174231010 | 174231409 | 174231010 | 174231160 | 174231288 | 128 | 174231409 | 6403 | 2,4 | 2,1 | 6,3 | 0,0 | 277 | 149 | 0.008350531 | 0.035619012 | 0.35,0.683 | 1.0,1.0 | -0.484 |
| 9216 | XLOC_014753 | chr15 | + | 85633962 | 85634412 | 85633962 | 85634146 | 85634274 | 128 | 85634412 | 9216 | 11,15 | 2,4 | 9,6 | 0,1 | 277 | 149 | 0.003756384 | 0.018449815 | 0.747,0.669 | 1.0,0.763 | -0.173 |
| 9316 | XLOC_030041 | chr3 | - | 131220374 | 131220768 | 131220374 | 131220559 | 131220687 | 128 | 131220768 | 9316 | 0,0 | 5,6 | 1,3 | 5,1 | 277 | 149 | 0.011472502 | 0.046519136 | 0.0,0.0 | 0.097,0.617 | -0.357 |
| 13600 | XLOC_027820 | chr21 | - | 46698017 | 46698296 | 46698017 | 46698084 | 46698212 | 128 | 46698296 | 13600 | 0,7 | 30,42 | 4,14 | 5,9 | 277 | 149 | 1.15E-06 | 1.65E-05 | 0.0,0.082 | 0.301,0.456 | -0.338 |
| 14810 | XLOC_013106 | chr14 | + | 45565751 | 45566208 | 45565751 | 45565961 | 45566089 | 128 | 45566208 | 14810 | 61,58 | 32,24 | 123,139 | 36,31 | 277 | 149 | 0.009409163 | 0.039245604 | 0.506,0.565 | 0.648,0.707 | -0.142 |
| 5216 | XLOC_029718 | chr3 | - | 3214189 | 3214609 | 3214189 | 3214330 | 3214459 | 129 | 3214609 | 5216 | 2,0 | 32,39 | 6,11 | 20,20 | 278 | 149 | 9.45E-07 | 1.39E-05 | 0.032,0.0 | 0.139,0.228 | -0.167 |
| 11608 | XLOC_005129 | chr10 | - | 12212715 | 12212943 | 12212715 | 12212769 | 12212898 | 129 | 12212943 | 11608 | 64,37 | 352,340 | 146,143 | 397,304 | 278 | 149 | 2.96E-11 | 9.99E-10 | 0.089,0.055 | 0.165,0.201 | -0.111 |
| 14440 | XLOC_001451 | chr1 | - | 192997004 | 192997278 | 192997004 | 192997085 | 192997214 | 129 | 192997278 | 14440 | 2,2 | 47,49 | 19,6 | 41,30 | 278 | 149 | 3.18E-06 | 4.11E-05 | 0.022,0.021 | 0.199,0.097 | -0.127 |
| 10092 | XLOC_005308 | chr10 | - | 95461930 | 95462304 | 95461930 | 95462001 | 95462131 | 130 | 95462304 | 10092 | 9,8 | 3,4 | 12,4 | 1,0 | 279 | 149 | 0.000543988 | 0.003697179 | 0.616,0.516 | 0.865,1.0 | -0.366 |
| 10501 | XLOC_000546 | chr1 | + | 155279854 | 155280244 | 155279854 | 155279996 | 155280126 | 130 | 155280244 | 10501 | 45,48 | 20,22 | 53,36 | 4,7 | 279 | 149 | 0.000480627 | 0.00332251 | 0.546,0.538 | 0.876,0.733 | -0.262 |
| 8289 | XLOC_034679 | chr5 | + | 133724015 | 133724279 | 133724015 | 133724104 | 133724235 | 131 | 133724279 | 8289 | 3,4 | 3,4 | 10,6 | 0,0 | 280 | 149 | 1.52E-07 | 2.68E-06 | 0.347,0.347 | 1.0,1.0 | -0.653 |
| 8800 | XLOC_037650 | chr6 | + | 13601065 | 13601353 | 13601065 | 13601181 | 13601312 | 131 | 13601353 | 8800 | 7,2 | 3,2 | 1,3 | 0,0 | 280 | 149 | 7.96E-06 | 9.27E-05 | 0.554,0.347 | 1.0,1.0 | -0.549 |
| 13947 | XLOC_001351 | chr1 | - | 155221292 | 155221699 | 155221292 | 155221392 | 155221523 | 131 | 155221699 | 13947 | 20,19 | 94,104 | 46,39 | 57,34 | 280 | 149 | 2.97E-10 | 8.44E-09 | 0.102,0.089 | 0.3,0.379 | -0.244 |
| 4405 | XLOC_046186 | chr9 | - | 13121744 | 13122169 | 13121744 | 13121953 | 13122085 | 132 | 13122169 | 4405 | 1,0 | 0,2 | 3,2 | 0,0 | 281 | 149 | 0.00142641 | 0.008225884 | 1.0,0.0 | 1.0,1.0 | -0.5 |
| 8996 | XLOC_009231 | chr12 | - | 7079098 | 7079443 | 7079098 | 7079226 | 7079358 | 132 | 7079443 | 8996 | 18,9 | 13,4 | 37,36 | 7,4 | 281 | 149 | 0.001792032 | 0.009952369 | 0.423,0.544 | 0.737,0.827 | -0.298 |
| 9147 | XLOC_047718 | chrX | + | 53113616 | 53114048 | 53113616 | 53113804 | 53113936 | 132 | 53114048 | 9147 | 26,16 | 43,37 | 28,10 | 13,10 | 281 | 149 | 0.004831013 | 0.022599666 | 0.243,0.187 | 0.533,0.347 | -0.225 |
| 10986 | XLOC_009211 | chr12 | - | 2973848 | 2974111 | 2973848 | 2973918 | 2974050 | 132 | 2974111 | 10986 | 11,2 | 13,13 | 13,8 | 3,0 | 281 | 149 | 3.72E-05 | 0.000364393 | 0.31,0.075 | 0.697,1.0 | -0.656 |
| 11786 | XLOC_007532 | chr11 | - | 61188525 | 61189080 | 61188525 | 61188729 | 61188861 | 132 | 61189080 | 11786 | 33,36 | 12,8 | 38,32 | 3,0 | 281 | 149 | 0.00011089 | 0.000936661 | 0.593,0.705 | 0.87,1.0 | -0.286 |
| 2058 | XLOC_017999 | chr17 | + | 29095851 | 29096803 | 29095851 | 29095885 | 29096018 | 133 | 29096803 | 2058 | 124,175 | 19,34 | 65,66 | 1,1 | 282 | 149 | 4.14E-08 | 8.08E-07 | 0.775,0.731 | 0.972,0.972 | -0.219 |
| 10202 | XLOC_000083 | chr1 | + | 12089897 | 12090181 | 12089897 | 12089951 | 12090084 | 133 | 12090181 | 10202 | 28,46 | 83,74 | 69,46 | 37,21 | 282 | 149 | 2.88E-09 | 6.85E-08 | 0.151,0.247 | 0.496,0.536 | -0.317 |
| 13995 | XLOC_013315 | chr14 | - | 23533313 | 23533692 | 23533313 | 23533430 | 23533563 | 133 | 23533692 | 13995 | 79,63 | 101,101 | 138,90 | 102,93 | 282 | 149 | 0.006295179 | 0.028188589 | 0.292,0.248 | 0.417,0.338 | -0.107 |
| 3479 | XLOC_021524 | chr19 | - | 9529180 | 9529571 | 9529180 | 9529307 | 9529441 | 134 | 9529571 | 3479 | 47,47 | 15,15 | 107,54 | 8,8 | 283 | 149 | 0.000746442 | 0.004770134 | 0.623,0.623 | 0.876,0.78 | -0.205 |
| 4295 | XLOC_041366 | chr7 | + | 157046360 | 157046836 | 157046360 | 157046513 | 157046647 | 134 | 157046836 | 4295 | 0,2 | 1,4 | 10,8 | 0,0 | 283 | 149 | 4.91E-09 | 1.13E-07 | 0.0,0.208 | 1.0,1.0 | -0.896 |
| 5887 | XLOC_026714 | chr20 | + | 13782131 | 13782519 | 13782131 | 13782329 | 13782463 | 134 | 13782519 | 5887 | 1,0 | 2,2 | 12,1 | 5,5 | 283 | 149 | 0.003580243 | 0.017727016 | 0.208,0.0 | 0.558,0.095 | -0.223 |
| 10078 | XLOC_035087 | chr5 | - | 141023784 | 141024184 | 141023784 | 141024005 | 141024139 | 134 | 141024184 | 10078 | 10,6 | 54,47 | 26,20 | 17,17 | 283 | 149 | 2.03E-10 | 5.93E-09 | 0.089,0.063 | 0.446,0.382 | -0.338 |
| 13402 | XLOC_000529 | chr1 | + | 154213988 | 154214273 | 154213988 | 154214041 | 154214175 | 134 | 154214273 | 13402 | 47,28 | 44,38 | 61,54 | 33,29 | 283 | 149 | 0.003854005 | 0.018833948 | 0.36,0.28 | 0.493,0.495 | -0.174 |
| 5484 | XLOC_028587 | chr22 | - | 21983604 | 21984050 | 21983604 | 21983696 | 21983831 | 135 | 21984050 | 5484 | 80,43 | 48,39 | 125,86 | 37,35 | 284 | 149 | 0.001900593 | 0.010411978 | 0.466,0.366 | 0.639,0.563 | -0.185 |
| 10222 | XLOC_020976 | chr19 | + | 11560080 | 11560500 | 11560080 | 11560243 | 11560378 | 135 | 11560500 | 10222 | 48,88 | 4,8 | 459,337 | 4,6 | 284 | 149 | 7.62E-05 | 0.000674252 | 0.863,0.852 | 0.984,0.967 | -0.118 |

| | | | | | | | | | | | | | | | | | | | | | | |
|---|---|---|---|---|---|---|---|---|---|---|---|---|---|---|---|---|---|---|---|---|---|---|
| 11359 | XLOC_000372 | chr1 | + | 93299101 | 93299506 | 93299101 | 93299217 | 93299352 | 135 | 93299506 | 11359 | 23,31 | 13,13 | 86,77 | 3,11 | 284 | 149 | 9.75E-06 | 0.000111552 | 0.481,0.556 | 0.938,0.786 | -0.344 |
| 11408 | XLOC_000229 | chr1 | + | 40214378 | 40214760 | 40214378 | 40214439 | 40214574 | 135 | 40214760 | 11408 | 1,2 | 6,4 | 2,2 | 0,0 | 284 | 149 | 4.17E-10 | 1.15E-08 | 0.08,0.208 | 1.0,1.0 | -0.856 |
| 8634 | XLOC_016352 | chr16 | + | 30001863 | 30002287 | 30001863 | 30001968 | 30002104 | 136 | 30002287 | 8634 | 11,9 | 5,4 | 8,10 | 0,0 | 285 | 149 | 2.49E-06 | 3.30E-05 | 0.535,0.541 | 1.0,1.0 | -0.462 |
| 10336 | XLOC_000396 | chr1 | + | 104068309 | 104068884 | 104068309 | 104068552 | 104068688 | 136 | 104068884 | 10336 | 42,40 | 5,11 | 39,28 | 2,1 | 285 | 149 | 0.003724561 | 0.018347394 | 0.815,0.655 | 0.911,0.936 | -0.189 |
| 10893 | XLOC_007073 | chr11 | + | 47259391 | 47259734 | 47259391 | 47259552 | 47259688 | 136 | 47259734 | 10893 | 21,20 | 171,228 | 45,32 | 96,54 | 285 | 149 | 3.95E-12 | 1.50E-10 | 0.06,0.044 | 0.197,0.237 | -0.165 |
| 11115 | XLOC_041587 | chr7 | - | 92207468 | 92207707 | 92207468 | 92207496 | 92207632 | 136 | 92207707 | 11115 | 2,0 | 105,114 | 22,26 | 119,91 | 285 | 149 | 3.00E-12 | 1.17E-10 | 0.01,0.0 | 0.088,0.13 | -0.104 |
| 13303 | XLOC_032424 | chr4 | + | 113567506 | 113567820 | 113567506 | 113567607 | 113567743 | 136 | 113567820 | 13303 | 3,5 | 127,123 | 29,16 | 99,74 | 285 | 149 | 6.95E-10 | 1.87E-08 | 0.012,0.021 | 0.133,0.102 | -0.101 |
| 14866 | XLOC_021276 | chr19 | + | 47109083 | 47109442 | 47109083 | 47109114 | 47109250 | 136 | 47109442 | 14866 | 17,13 | 2,4 | 6,7 | 1,0 | 285 | 149 | 0.006669232 | 0.029535767 | 0.816,0.63 | 0.758,1.0 | -0.156 |
| 11409 | XLOC_021106 | chr19 | + | 33073100 | 33073364 | 33073100 | 33073138 | 33073275 | 137 | 33073364 | 11409 | 16,23 | 4,2 | 40,27 | 0,0 | 286 | 149 | 9.27E-05 | 0.000804015 | 0.676,0.857 | 1.0,1.0 | -0.234 |
| 14592 | XLOC_016793 | chr16 | - | 70288293 | 70288637 | 70288293 | 70288386 | 70288523 | 137 | 70288637 | 14592 | 5,1 | 2,4 | 9,4 | 1,1 | 286 | 149 | 0.008332585 | 0.035563946 | 0.566,0.115 | 0.824,0.676 | -0.41 |
| 14766 | XLOC_000574 | chr1 | + | 161140469 | 161140739 | 161140469 | 161140559 | 161140696 | 137 | 161140739 | 14766 | 4,9 | 89,122 | 23,16 | 67,23 | 286 | 149 | 1.02E-08 | 2.17E-07 | 0.023,0.037 | 0.152,0.266 | -0.179 |
| 5580 | XLOC_041629 | chr7 | - | 100176021 | 100176346 | 100176021 | 100176134 | 100176272 | 138 | 100176346 | 5580 | 3,7 | 85,84 | 20,13 | 59,33 | 287 | 149 | 3.94E-07 | 6.43E-06 | 0.018,0.041 | 0.15,0.17 | -0.131 |
| 10234 | XLOC_018796 | chr17 | - | 80407045 | 80407331 | 80407045 | 80407168 | 80407306 | 138 | 80407331 | 10234 | 58,40 | 39,34 | 45,33 | 5,0 | 287 | 149 | 3.82E-09 | 8.87E-08 | 0.436,0.379 | 0.824,1.0 | -0.504 |
| 10380 | XLOC_029789 | chr3 | - | 39142278 | 39142593 | 39142278 | 39142368 | 39142506 | 138 | 39142593 | 10380 | 5,15 | 22,35 | 24,14 | 16,18 | 287 | 149 | 0.002424998 | 0.012755401 | 0.106,0.182 | 0.438,0.288 | -0.219 |
| 11360 | XLOC_000372 | chr1 | + | 93299101 | 93299506 | 93299101 | 93299217 | 93299355 | 138 | 93299506 | 11360 | 23,31 | 17,8 | 86,77 | 14,6 | 287 | 149 | 0.00331163 | 0.016614619 | 0.413,0.668 | 0.761,0.869 | -0.274 |
| 13645 | XLOC_018386 | chr17 | - | 7221594 | 7221993 | 7221594 | 7221675 | 7221813 | 138 | 7221993 | 13645 | 13,9 | 68,63 | 98,43 | 120,67 | 287 | 149 | 1.99E-09 | 4.92E-08 | 0.09,0.069 | 0.298,0.25 | -0.195 |
| 6528 | XLOC_023399 | chr2 | + | 27260657 | 27260969 | 27260657 | 27260760 | 27260899 | 139 | 27260969 | 6528 | 7,11 | 1,3 | 21,13 | 0,0 | 288 | 149 | 0.001909458 | 0.01044838 | 0.784,0.655 | 1.0,1.0 | -0.28 |
| 7420 | XLOC_034882 | chr5 | - | 40834023 | 40834367 | 40834023 | 40834143 | 40834282 | 139 | 40834367 | 7420 | 8,9 | 9,0 | 102,65 | 0,0 | 288 | 149 | 0.000647217 | 0.004271331 | 0.315,1.0 | 1.0,1.0 | -0.343 |
| 8221 | XLOC_016438 | chr16 | + | 67262397 | 67262739 | 67262397 | 67262560 | 67262699 | 139 | 67262739 | 8221 | 9,12 | 25,10 | 64,45 | 27,20 | 288 | 149 | 0.001087848 | 0.006551674 | 0.157,0.383 | 0.551,0.538 | -0.274 |
| 8452 | XLOC_026945 | chr20 | - | 23375558 | 23375899 | 23375558 | 23375636 | 23375775 | 139 | 23375899 | 8452 | 4,13 | 13,15 | 18,17 | 2,4 | 288 | 149 | 8.35E-06 | 9.68E-05 | 0.137,0.31 | 0.823,0.687 | -0.531 |
| 10399 | XLOC_046127 | chr9 | + | 138393689 | 138394107 | 138393689 | 138393819 | 138393958 | 139 | 138394107 | 10399 | 6,9 | 4,0 | 29,22 | 0,0 | 288 | 149 | 0.007124155 | 0.031149983 | 0.437,1.0 | 1.0,1.0 | -0.281 |
| 13626 | XLOC_016775 | chr16 | - | 67866130 | 67866446 | 67866130 | 67866218 | 67866357 | 139 | 67866446 | 13626 | 32,20 | 124,124 | 42,47 | 118,71 | 288 | 149 | 0.000414151 | 0.002923069 | 0.118,0.077 | 0.156,0.255 | -0.108 |
| 5579 | XLOC_041629 | chr7 | - | 100175781 | 100176088 | 100175781 | 100175881 | 100176021 | 140 | 100176088 | 5579 | 13,15 | 97,82 | 26,28 | 44,33 | 289 | 149 | 8.67E-08 | 1.60E-06 | 0.065,0.086 | 0.234,0.304 | -0.194 |
| 7747 | XLOC_018533 | chr17 | - | 38552234 | 38552717 | 38552234 | 38552394 | 38552534 | 140 | 38552717 | 7747 | 4,9 | 1,2 | 5,1 | 0,0 | 289 | 149 | 0.004389507 | 0.020844532 | 0.673,0.699 | 1.0,1.0 | -0.314 |
| 7867 | XLOC_018304 | chr17 | + | 80680679 | 80680987 | 80680679 | 80680762 | 80680902 | 140 | 80680987 | 7867 | 14,15 | 7,2 | 20,20 | 0,0 | 289 | 149 | 3.49E-06 | 4.44E-05 | 0.508,0.795 | 1.0,1.0 | -0.349 |
| 10789 | XLOC_009489 | chr12 | - | 95450972 | 95451419 | 95450972 | 95451188 | 95451328 | 140 | 95451419 | 10789 | 0,5 | 9,15 | 14,8 | 5,8 | 289 | 149 | 0.000299606 | 0.00221304 | 0.0,0.147 | 0.591,0.34 | -0.392 |
| 12985 | XLOC_000277 | chr1 | + | 45807144 | 45807415 | 45807144 | 45807241 | 45807381 | 140 | 45807415 | 12985 | 6,5 | 2,2 | 14,7 | 0,1 | 289 | 149 | 0.00172107 | 0.009634037 | 0.607,0.563 | 1.0,0.783 | -0.306 |
| 8177 | XLOC_029429 | chr3 | + | 50098233 | 50098654 | 50098233 | 50098431 | 50098572 | 141 | 50098654 | 8177 | 2,8 | 113,124 | 39,45 | 130,67 | 290 | 149 | 1.55E-13 | 7.33E-12 | 0.009,0.032 | 0.134,0.257 | -0.175 |
| 8343 | XLOC_016862 | chr16 | - | 89836374 | 89836667 | 89836374 | 89836432 | 89836573 | 141 | 89836667 | 8343 | 31,26 | 160,146 | 48,43 | 131,78 | 290 | 149 | 5.44E-05 | 0.000506002 | 0.091,0.084 | 0.158,0.221 | -0.102 |
| 12204 | XLOC_007382 | chr11 | - | 1774762 | 1775131 | 1774762 | 1774891 | 1775032 | 141 | 1775131 | 12204 | 6,1 | 6,0 | 2,1 | 0,0 | 290 | 149 | 0.00077751 | 0.004937395 | 0.339,1.0 | 1.0,1.0 | -0.331 |
| 12265 | XLOC_028747 | chr22 | - | 50720358 | 50720742 | 50720358 | 50720471 | 50720612 | 141 | 50720742 | 12265 | 6,3 | 2,2 | 15,9 | 0,0 | 290 | 149 | 0.000268966 | 0.002014779 | 0.607,0.435 | 1.0,1.0 | -0.479 |
| 13229 | XLOC_016538 | chr16 | + | 89760976 | 89761460 | 89760976 | 89761196 | 89761338 | 142 | 89761460 | 13229 | 76,93 | 149,129 | 94,62 | 64,41 | 291 | 149 | 1.79E-06 | 2.45E-05 | 0.207,0.27 | 0.429,0.436 | -0.194 |
| 7419 | XLOC_034882 | chr5 | - | 40834023 | 40834367 | 40834023 | 40834139 | 40834282 | 143 | 40834367 | 7419 | 8,9 | 30,24 | 102,65 | 59,53 | 292 | 149 | 1.69E-07 | 2.96E-06 | 0.12,0.161 | 0.469,0.385 | -0.286 |
| 10284 | XLOC_035144 | chr5 | - | 173036240 | 173036673 | 173036240 | 173036437 | 173036580 | 143 | 173036673 | 10284 | 55,51 | 51,52 | 101,71 | 34,19 | 292 | 149 | 3.36E-07 | 5.53E-06 | 0.355,0.334 | 0.603,0.656 | -0.285 |
| 12689 | XLOC_016244 | chr16 | + | 3115462 | 3115656 | 3115462 | 3115495 | 3115638 | 143 | 3115656 | 12689 | 20,29 | 3,2 | 66,32 | 0,0 | 292 | 149 | 0.001875024 | 0.01031994 | 0.773,0.881 | 1.0,1.0 | -0.173 |
| 4275 | XLOC_041420 | chr7 | - | 21945705 | 21946074 | 21945705 | 21945762 | 21945906 | 144 | 21946074 | 4275 | 0,4 | 4,3 | 6,1 | 0,0 | 293 | 149 | 2.18E-09 | 5.34E-08 | 0.0,0.404 | 1.0,1.0 | -0.798 |
| 9822 | XLOC_021503 | chr19 | - | 6730997 | 6731245 | 6730997 | 6731065 | 6731209 | 144 | 6731245 | 9822 | 24,43 | 5,1 | 46,38 | 0,1 | 293 | 149 | 0.003161418 | 0.015957367 | 0.709,0.956 | 1.0,0.951 | -0.143 |
| 2572 | XLOC_014712 | chr15 | + | 75646993 | 75648036 | 75646993 | 75647159 | 75647304 | 145 | 75648036 | 2572 | 7,5 | 38,40 | 21,13 | 21,27 | 294 | 149 | 8.53E-05 | 0.000745849 | 0.085,0.06 | 0.336,0.196 | -0.194 |
| 8199 | XLOC_021878 | chr19 | - | 55604150 | 55604430 | 55604150 | 55604233 | 55604378 | 145 | 55604430 | 8199 | 4,0 | 3,2 | 3,6 | 0,2 | 294 | 149 | 0.009038109 | 0.037989645 | 0.403,0.0 | 1.0,0.603 | -0.6 |
| 11992 | XLOC_007182 | chr11 | + | 67057792 | 67058187 | 67057792 | 67057917 | 67058062 | 145 | 67058187 | 11992 | 6,10 | 21,16 | 15,13 | 10,8 | 294 | 149 | 0.002537824 | 0.013232141 | 0.126,0.241 | 0.432,0.452 | -0.259 |
| 5842 | XLOC_026812 | chr20 | + | 44441758 | 44442103 | 44441758 | 44441929 | 44442075 | 146 | 44442103 | 5842 | 25,22 | 20,20 | 24,17 | 5,6 | 295 | 149 | 0.003779391 | 0.018546329 | 0.387,0.357 | 0.708,0.589 | -0.276 |
| 3158 | XLOC_018763 | chr17 | - | 78183459 | 78186073 | 78183459 | 78185722 | 78185869 | 147 | 78186073 | 3158 | 68,87 | 5,5 | 131,132 | 0,0 | 296 | 149 | 2.60E-05 | 0.000263149 | 0.873,0.898 | 1.0,1.0 | -0.115 |
| 4420 | XLOC_047701 | chrX | + | 48433948 | 48434354 | 48433948 | 48434055 | 48434202 | 147 | 48434354 | 4420 | 92,71 | 28,36 | 326,145 | 44,34 | 296 | 149 | 0.001858119 | 0.010246861 | 0.623,0.498 | 0.789,0.682 | -0.175 |
| 8202 | XLOC_021878 | chr19 | - | 55604150 | 55604430 | 55604150 | 55604236 | 55604383 | 147 | 55604430 | 8202 | 4,0 | 36,35 | 8,10 | 42,22 | 296 | 149 | 0.000638236 | 0.004219945 | 0.053,0.0 | 0.087,0.186 | -0.11 |
| 11602 | XLOC_007722 | chr11 | - | 116732917 | 116733293 | 116732917 | 116733043 | 116733190 | 147 | 116733293 | 11602 | 4,2 | 4,0 | 9,1 | 0,0 | 296 | 149 | 0.004737415 | 0.022220697 | 0.335,1.0 | 1.0,1.0 | -0.333 |
| 11868 | XLOC_007403 | chr11 | - | 6638325 | 6638659 | 6638325 | 6638384 | 6638531 | 147 | 6638659 | 11868 | 15,9 | 205,233 | 39,17 | 91,78 | 296 | 149 | 4.55E-11 | 1.49E-09 | 0.036,0.019 | 0.177,0.099 | -0.111 |
| 15233 | XLOC_018472 | chr17 | - | 26960300 | 26960680 | 26960300 | 26960419 | 26960566 | 147 | 26960680 | 15233 | 46,53 | 298,304 | 125,78 | 154,115 | 296 | 149 | 0 | 0 | 0.072,0.081 | 0.29,0.255 | -0.196 |
| 6032 | XLOC_001238 | chr1 | - | 111662236 | 111662581 | 111662236 | 111662350 | 111662498 | 148 | 111662581 | 6032 | 9,13 | 2,4 | 6,15 | 1,0 | 297 | 149 | 0.00146084 | 0.008390175 | 0.693,0.62 | 0.751,1.0 | -0.219 |
| 9804 | XLOC_029885 | chr3 | - | 53852964 | 53853358 | 53852964 | 53853067 | 53853215 | 148 | 53853358 | 9804 | 2,4 | 9,2 | 2,3 | 1,0 | 297 | 149 | 0.001107327 | 0.00665197 | 0.1,0.501 | 0.501,1.0 | -0.45 |
| 11828 | XLOC_001313 | chr1 | - | 151318897 | 151319342 | 151318897 | 151319082 | 151319230 | 148 | 151319342 | 11828 | 3,0 | 14,9 | 4,7 | 8,3 | 297 | 149 | 0.002207435 | 0.011804468 | 0.097,0.0 | 0.201,0.539 | -0.322 |
| 12199 | XLOC_024206 | chr2 | - | 128624473 | 128624825 | 128624473 | 128624570 | 128624718 | 148 | 128624825 | 12199 | 10,8 | 25,32 | 31,36 | 23,21 | 297 | 149 | 9.88E-07 | 1.45E-05 | 0.167,0.111 | 0.403,0.462 | -0.293 |
| 14027 | XLOC_029433 | chr3 | + | 50307557 | 50307924 | 50307557 | 50307715 | 50307863 | 148 | 50307924 | 14027 | 20,30 | 147,154 | 261,156 | 502,370 | 297 | 149 | 5.59E-09 | 1.26E-07 | 0.064,0.089 | 0.207,0.175 | -0.115 |
| 3894 | XLOC_046005 | chr9 | + | 111697917 | 111698291 | 111697917 | 111697969 | 111698118 | 149 | 111698291 | 3894 | 16,26 | 7,4 | 52,47 | 3,3 | 298 | 149 | 0.008248152 | 0.035278211 | 0.533,0.765 | 0.897,0.887 | -0.243 |
| 12345 | XLOC_041378 | chr7 | - | 1523336 | 1523806 | 1523336 | 1523489 | 1523638 | 149 | 1523806 | 12345 | 11,17 | 116,139 | 49,55 | 151,124 | 298 | 149 | 9.52E-08 | 1.75E-06 | 0.045,0.058 | 0.14,0.182 | -0.11 |
| 12667 | XLOC_044333 | chr8 | - | 74867881 | 74868289 | 74867881 | 74867996 | 74868145 | 149 | 74868289 | 12667 | 19,21 | 3,3 | 11,13 | 1,0 | 298 | 149 | 0.004233303 | 0.020218782 | 0.76,0.778 | 0.846,1.0 | -0.154 |
| 7067 | XLOC_021069 | chr19 | + | 19309768 | 19310043 | 19309768 | 19309812 | 19309962 | 150 | 19310043 | 7067 | 0,15 | 5,0 | 2,12 | 0,0 | 299 | 149 | 0.002574413 | 0.013376909 | 0.0,1.0 | 1.0,1.0 | -0.5 |
| 11527 | XLOC_023422 | chr2 | + | 29149333 | 29149689 | 29149333 | 29149420 | 29149570 | 150 | 29149689 | 11527 | 3,5 | 6,4 | 11,9 | 0,0 | 299 | 149 | 1.61E-09 | 4.05E-08 | 0.199,0.384 | 1.0,1.0 | -0.709 |
| 66 | XLOC_046241 | chr9 | - | 35738228 | 35738629 | 35738228 | 35738371 | 35738522 | 151 | 35738629 | 66 | 121,111 | 45,64 | 224,182 | 51,22 | 300 | 149 | 6.39E-05 | 0.000580174 | 0.572,0.463 | 0.686,0.804 | -0.228 |
| 1260 | XLOC_018792 | chr17 | - | 80151631 | 80152015 | 80151631 | 80151706 | 80151857 | 151 | 80152015 | 1260 | 73,76 | 39,52 | 104,65 | 32,13 | 300 | 149 | 0.00054733 | 0.003716321 | 0.482,0.421 | 0.617,0.713 | -0.214 |
| 15127 | XLOC_000531 | chr1 | + | 154245955 | 154246279 | 154245955 | 154246074 | 154246225 | 151 | 154246279 | 15127 | 4,10 | 7,5 | 17,14 | 2,1 | 300 | 149 | 0.000508397 | 0.003483788 | 0.221,0.498 | 0.808,0.874 | -0.481 |
| 4668 | XLOC_035177 | chr5 | - | 179674438 | 179674728 | 179674438 | 179674510 | 179674662 | 152 | 179674728 | 4668 | 13,2 | 10,5 | 30,26 | 1,6 | 301 | 149 | 0.000269316 | 0.002015656 | 0.392,0.165 | 0.937,0.682 | -0.531 |
| 6624 | XLOC_016192 | chr16 | + | 838655 | 839129 | 838655 | 838775 | 838927 | 152 | 839129 | 6624 | 0,1 | 12,12 | 1,4 | 5,11 | 301 | 149 | 0.008528346 | 0.036235207 | 0.0,0.04 | 0.09,0.153 | -0.101 |
| 6800 | XLOC_021710 | chr19 | - | 39321717 | 39322147 | 39321717 | 39321806 | 39321959 | 153 | 39322147 | 6800 | 1,0 | 6,2 | 6,4 | 2,1 | 302 | 149 | 7.32E-06 | 8.62E-05 | 0.076,0.0 | 0.597,0.664 | -0.593 |
| 12269 | XLOC_028747 | chr22 | - | 50721432 | 50721874 | 50721432 | 50721612 | 50721765 | 153 | 50721874 | 12269 | 20,24 | 1,5 | 41,40 | 0,2 | 302 | 149 | 0.008872668 | 0.037448676 | 0.908,0.703 | 1.0,0.908 | -0.148 |
| 12454 | XLOC_001053 | chr1 | - | 40544231 | 40544510 | 40544231 | 40544330 | 40544483 | 153 | 40544510 | 12454 | 1,4 | 6,1 | 11,5 | 0,0 | 302 | 149 | 4.98E-07 | 7.92E-06 | 0.076,0.664 | 1.0,1.0 | -0.63 |

| | | | | | | | | | | | | | | | | | | | | | | |
|---|---|---|---|---|---|---|---|---|---|---|---|---|---|---|---|---|---|---|---|---|---|---|
| 14991 | XLOC_000016 | chr1 | + | 1451013 | 1451468 | 1451013 | 1451238 | 1451391 | 153 | 1451468 | 14991 | 10,3 | 5,6 | 28,11 | 5,0 | 302 | 149 | 0.002258006 | 0.012029356 | 0.497,0.198 | 0.734,1.0 | -0.519 |
| 5892 | XLOC_026684 | chr20 | + | 2842421 | 2842776 | 2842421 | 2842545 | 2842699 | 154 | 2842776 | 5892 | 9,4 | 81,62 | 16,19 | 33,33 | 303 | 149 | 2.99E-07 | 4.99E-06 | 0.052,0.031 | 0.193,0.221 | -0.166 |
| 7395 | XLOC_028532 | chr22 | + | 45717927 | 45718307 | 45717927 | 45718096 | 45718251 | 155 | 45718307 | 7395 | 0,2 | 5,6 | 11,4 | 0,5 | 304 | 149 | 0.000408525 | 0.002889204 | 0.0,0.14 | 1.0,0.282 | -0.571 |
| 8851 | XLOC_046057 | chr9 | + | 130165933 | 130166346 | 130165933 | 130166082 | 130166237 | 155 | 130166346 | 8851 | 9,8 | 55,35 | 21,17 | 41,16 | 304 | 149 | 0.000346697 | 0.002507115 | 0.074,0.101 | 0.201,0.342 | -0.184 |
| 9506 | XLOC_021058 | chr19 | + | 18684102 | 18684557 | 18684102 | 18684213 | 18684368 | 155 | 18684557 | 9506 | 6,8 | 0,5 | 17,4 | 0,0 | 304 | 149 | 0.001544093 | 0.008779078 | 1.0,0.44 | 1.0,1.0 | -0.28 |
| 13393 | XLOC_029987 | chr3 | - | 121386907 | 121387355 | 121386907 | 121387005 | 121387160 | 155 | 121387355 | 13393 | 0,0 | 10,4 | 1,4 | 6,2 | 304 | 149 | 0.002768097 | 0.014226286 | 0.0,0.0 | 0.076,0.495 | -0.285 |
| 8340 | XLOC_016862 | chr16 | - | 89808927 | 89809346 | 89808927 | 89809051 | 89809207 | 156 | 89809346 | 8340 | 10,9 | 1,4 | 4,10 | 0,0 | 305 | 149 | 0.000129514 | 0.001069036 | 0.83,0.524 | 1.0,1.0 | -0.323 |
| 7844 | XLOC_018346 | chr17 | - | 3785550 | 3785889 | 3785550 | 3785664 | 3785821 | 157 | 3785889 | 7844 | 6,6 | 18,13 | 14,22 | 8,6 | 306 | 149 | 5.33E-05 | 0.000497016 | 0.14,0.183 | 0.46,0.641 | -0.389 |
| 10142 | XLOC_000887 | chr1 | - | 11137386 | 11137708 | 11137386 | 11137500 | 11137657 | 157 | 11137708 | 10142 | 51,40 | 183,188 | 110,92 | 171,111 | 306 | 149 | 8.02E-10 | 2.12E-08 | 0.119,0.094 | 0.239,0.288 | -0.157 |
| 11727 | XLOC_043957 | chr8 | + | 21937922 | 21938136 | 21937922 | 21937971 | 21938128 | 157 | 21938136 | 11727 | 32,23 | 12,5 | 12,3 | 0,0 | 306 | 149 | 1.32E-05 | 0.000145156 | 0.565,0.691 | 1.0,1.0 | -0.372 |
| 6464 | XLOC_028458 | chr22 | + | 32007751 | 32008035 | 32007751 | 32007826 | 32007984 | 158 | 32008035 | 6464 | 58,61 | 12,20 | 95,65 | 5,2 | 307 | 149 | 2.82E-06 | 3.69E-05 | 0.701,0.597 | 0.902,0.94 | -0.272 |
| 6559 | XLOC_023559 | chr2 | + | 85848329 | 85848702 | 85848329 | 85848449 | 85848607 | 158 | 85848702 | 6559 | 1,0 | 4,11 | 10,0 | 5,1 | 307 | 149 | 0.00166867 | 0.009392837 | 0.108,0.0 | 0.493,0.0 | -0.193 |
| 6901 | XLOC_021444 | chr19 | - | 1235310 | 1235564 | 1235310 | 1235341 | 1235499 | 158 | 1235564 | 6901 | 1,2 | 4,9 | 3,11 | 0,0 | 307 | 149 | 3.34E-11 | 1.12E-09 | 0.108,0.097 | 1.0,1.0 | -0.897 |
| 7227 | XLOC_047975 | chrX | - | 57004724 | 57005093 | 57004724 | 57004854 | 57005012 | 158 | 57005093 | 7227 | 0,0 | 3,1 | 9,6 | 13,0 | 307 | 149 | 1.57E-05 | 0.000167638 | 0.0,0.0 | 0.252,1.0 | -0.626 |
| 8335 | XLOC_016862 | chr16 | - | 89805289 | 89805674 | 89805289 | 89805382 | 89805540 | 158 | 89805674 | 8335 | 169,164 | 128,146 | 228,193 | 110,83 | 307 | 149 | 0.000150296 | 0.001211547 | 0.391,0.353 | 0.501,0.53 | -0.144 |
| 13919 | XLOC_001072 | chr1 | - | 43917498 | 43917990 | 43917498 | 43917717 | 43917875 | 158 | 43917990 | 13919 | 28,32 | 1,6 | 36,27 | 0,0 | 307 | 149 | 0.000160479 | 0.001282654 | 0.931,0.721 | 1.0,1.0 | -0.174 |
| 14206 | XLOC_018794 | chr17 | - | 80203851 | 80204341 | 80203851 | 80203915 | 80204073 | 158 | 80204341 | 14206 | 105,139 | 83,87 | 187,164 | 15,15 | 307 | 149 | 0 | 0 | 0.38,0.437 | 0.858,0.841 | -0.441 |
| 6069 | XLOC_016187 | chr16 | + | 705115 | 705468 | 705115 | 705147 | 705306 | 159 | 705468 | 6069 | 11,12 | 89,103 | 39,38 | 64,44 | 308 | 149 | 1.42E-11 | 5.00E-10 | 0.056,0.053 | 0.228,0.295 | -0.207 |
| 6978 | XLOC_021281 | chr19 | + | 47761586 | 47761967 | 47761586 | 47761660 | 47761819 | 159 | 47761967 | 6978 | 1,7 | 13,4 | 1,2 | 1,0 | 308 | 149 | 0.006542786 | 0.029112717 | 0.036,0.458 | 0.326,1.0 | -0.416 |
| 8261 | XLOC_047679 | chrX | + | 41064555 | 41064936 | 41064555 | 41064746 | 41064905 | 159 | 41064936 | 8261 | 0,1 | 2,1 | 8,3 | 3,0 | 308 | 149 | 0.000227775 | 0.001734158 | 0.0,0.326 | 0.563,1.0 | -0.618 |
| 8345 | XLOC_016862 | chr16 | - | 89851261 | 89851567 | 89851261 | 89851372 | 89851531 | 159 | 89851567 | 8345 | 0,7 | 4,1 | 7,17 | 1,1 | 308 | 149 | 0.005144612 | 0.023790883 | 0.0,0.772 | 0.772,0.892 | -0.446 |
| 9236 | XLOC_014711 | chr15 | + | 75631377 | 75631685 | 75631377 | 75631445 | 75631604 | 159 | 75631685 | 9236 | 71,54 | 395,426 | 97,66 | 234,144 | 308 | 149 | 4.09E-10 | 1.13E-08 | 0.08,0.058 | 0.167,0.181 | -0.105 |
| 4567 | XLOC_037837 | chr6 | + | 43143443 | 43143904 | 43143443 | 43143705 | 43143865 | 160 | 43143904 | 4567 | 2,5 | 2,3 | 14,1 | 0,0 | 309 | 149 | 2.50E-06 | 3.32E-05 | 0.325,0.446 | 1.0,1.0 | -0.615 |
| 8724 | XLOC_016204 | chr16 | + | 1825239 | 1825685 | 1825239 | 1825409 | 1825569 | 160 | 1825685 | 8724 | 0,5 | 29,42 | 8,2 | 8,7 | 309 | 149 | 0.000557172 | 0.00377407 | 0.0,0.054 | 0.325,0.121 | -0.196 |
| 9072 | XLOC_016207 | chr16 | + | 2011502 | 2011986 | 2011502 | 2011637 | 2011797 | 160 | 2011986 | 9072 | 186,218 | 507,490 | 581,391 | 169,96 | 309 | 149 | 0 | 0 | 0.15,0.177 | 0.624,0.663 | -0.48 |
| 9593 | XLOC_007211 | chr11 | + | 71939770 | 71940080 | 71939770 | 71939891 | 71940051 | 160 | 71940080 | 9593 | 87,107 | 74,67 | 193,129 | 25,10 | 309 | 149 | 0 | 0 | 0.362,0.435 | 0.788,0.862 | -0.426 |
| 13647 | XLOC_018386 | chr17 | - | 7224679 | 7224979 | 7224679 | 7224778 | 7224938 | 160 | 7224979 | 13647 | 1,4 | 4,0 | 1,5 | 0,0 | 309 | 149 | 0.005179971 | 0.023938719 | 0.108,1.0 | 1.0,1.0 | -0.446 |
| 7590 | XLOC_005345 | chr10 | - | 101945270 | 101945645 | 101945270 | 101945394 | 101945555 | 161 | 101945645 | 7590 | 2,4 | 2,4 | 11,8 | 1,0 | 310 | 149 | 1.41E-05 | 0.000153473 | 0.325,0.325 | 0.841,1.0 | -0.595 |
| 9519 | XLOC_007716 | chr11 | - | 113678672 | 113679159 | 113678672 | 113678858 | 113679019 | 161 | 113679159 | 9519 | 0,1 | 1,3 | 4,1 | 0,0 | 310 | 149 | 4.62E-09 | 1.06E-07 | 0.0,0.138 | 1.0,1.0 | -0.931 |
| 12618 | XLOC_032703 | chr4 | - | 104115526 | 104115841 | 104115526 | 104115598 | 104115759 | 161 | 104115841 | 12618 | 0,0 | 38,32 | 4,10 | 18,30 | 310 | 149 | 2.43E-06 | 3.22E-05 | 0.0,0.0 | 0.097,0.138 | -0.118 |
| 12859 | XLOC_027092 | chr20 | - | 62577202 | 62577620 | 62577202 | 62577352 | 62577513 | 161 | 62577620 | 12859 | 47,22 | 6,1 | 38,32 | 1,0 | 310 | 149 | 0.001211956 | 0.007170698 | 0.79,0.914 | 0.948,1.0 | -0.122 |
| 4507 | XLOC_038232 | chr6 | - | 33545311 | 33545685 | 33545311 | 33545412 | 33545574 | 162 | 33545685 | 4507 | 0,0 | 1,6 | 2,8 | 5,0 | 311 | 149 | 5.15E-05 | 0.000483038 | 0.0,0.0 | 0.161,1.0 | -0.581 |
| 7566 | XLOC_018771 | chr17 | - | 79226608 | 79227040 | 79226608 | 79226710 | 79226872 | 162 | 79227040 | 7566 | 2,1 | 5,2 | 12,6 | 0,0 | 311 | 149 | 2.10E-09 | 5.16E-08 | 0.161,0.193 | 1.0,1.0 | -0.823 |
| 9396 | XLOC_009615 | chr12 | - | 132626394 | 132626862 | 132626394 | 132626501 | 132626663 | 162 | 132626862 | 9396 | 14,5 | 30,38 | 58,40 | 42,19 | 311 | 149 | 8.49E-08 | 1.58E-06 | 0.183,0.059 | 0.398,0.502 | -0.329 |
| 11394 | XLOC_021828 | chr19 | - | 50365028 | 50365494 | 50365028 | 50365138 | 50365300 | 162 | 50365494 | 11394 | 51,44 | 155,112 | 76,63 | 80,64 | 311 | 149 | 2.82E-07 | 4.72E-06 | 0.136,0.158 | 0.313,0.32 | -0.169 |
| 8336 | XLOC_016862 | chr16 | - | 89805289 | 89805674 | 89805289 | 89805382 | 89805545 | 163 | 89805674 | 8336 | 78,91 | 7,15 | 98,95 | 6,1 | 312 | 149 | 0.001772765 | 0.009868648 | 0.842,0.743 | 0.886,0.978 | -0.139 |
| 9520 | XLOC_023855 | chr2 | + | 219544416 | 219544803 | 219544416 | 219544452 | 219544615 | 163 | 219544803 | 9520 | 0,2 | 21,14 | 6,1 | 10,4 | 312 | 149 | 0.002881411 | 0.014717602 | 0.0,0.064 | 0.223,0.107 | -0.133 |
| 4912 | XLOC_032534 | chr4 | - | 959851 | 960315 | 959851 | 960089 | 960253 | 164 | 960315 | 4912 | 3,1 | 2,2 | 4,2 | 0,0 | 313 | 149 | 1.01E-05 | 0.00011492 | 0.417,0.192 | 1.0,1.0 | -0.696 |
| 7077 | XLOC_044384 | chr8 | - | 103306182 | 103306585 | 103306182 | 103306343 | 103306507 | 164 | 103306585 | 7077 | 0,2 | 1,1 | 2,2 | 0,0 | 313 | 149 | 0.008998078 | 0.037855186 | 0.0,0.488 | 1.0,1.0 | -0.756 |
| 5409 | XLOC_028745 | chr22 | - | 50658150 | 50658444 | 50658150 | 50658220 | 50658385 | 165 | 50658444 | 5409 | 6,4 | 56,79 | 22,10 | 45,36 | 314 | 149 | 4.64E-06 | 5.77E-05 | 0.048,0.023 | 0.188,0.116 | -0.116 |
| 6073 | XLOC_016187 | chr16 | + | 708182 | 708614 | 708182 | 708344 | 708509 | 165 | 708614 | 6073 | 5,2 | 4,2 | 5,2 | 0,0 | 314 | 149 | 1.18E-06 | 1.70E-05 | 0.372,0.322 | 1.0,1.0 | -0.653 |
| 402 | XLOC_034712 | chr5 | + | 140021899 | 140022256 | 140021899 | 140022004 | 140022170 | 166 | 140022256 | 402 | 34,24 | 40,46 | 42,28 | 28,21 | 315 | 149 | 0.006434439 | 0.0286939 | 0.287,0.198 | 0.415,0.387 | -0.159 |
| 3912 | XLOC_045958 | chr9 | + | 90254268 | 90254602 | 90254268 | 90254398 | 90254564 | 166 | 90254602 | 3912 | 11,6 | 142,105 | 15,7 | 25,23 | 315 | 149 | 1.45E-06 | 2.04E-05 | 0.035,0.026 | 0.221,0.126 | -0.143 |
| 5203 | XLOC_029782 | chr3 | - | 33438236 | 33438627 | 33438236 | 33438370 | 33438536 | 166 | 33438627 | 5203 | 7,7 | 8,13 | 44,14 | 13,7 | 315 | 149 | 0.000881756 | 0.00548834 | 0.293,0.203 | 0.616,0.486 | -0.303 |
| 13571 | XLOC_037622 | chr6 | + | 2779496 | 2780107 | 2779496 | 2779726 | 2779892 | 166 | 2780107 | 13571 | 68,61 | 2,6 | 56,48 | 0,0 | 315 | 149 | 0.000373718 | 0.00267784 | 0.941,0.828 | 1.0,1.0 | -0.116 |
| 5641 | XLOC_027720 | chr21 | + | 46591524 | 46591847 | 46591524 | 46591599 | 46591766 | 167 | 46591847 | 5641 | 48,40 | 7,12 | 90,54 | 4,6 | 316 | 149 | 0.009206936 | 0.038550122 | 0.764,0.611 | 0.914,0.809 | -0.174 |
| 13659 | XLOC_018386 | chr17 | - | 7228550 | 7229032 | 7228550 | 7228792 | 7228959 | 167 | 7229032 | 13659 | 14,28 | 57,72 | 66,42 | 73,58 | 316 | 149 | 6.32E-05 | 0.000574837 | 0.104,0.155 | 0.299,0.255 | -0.148 |
| 5262 | XLOC_029569 | chr3 | + | 129225251 | 129225679 | 129225251 | 129225392 | 129225560 | 168 | 129225679 | 5262 | 0,0 | 3,1 | 2,2 | 0,1 | 317 | 149 | 1.34E-05 | 0.000147353 | 0.0,0.0 | 1.0,0.485 | -0.742 |
| 10079 | XLOC_035087 | chr5 | - | 141024139 | 141024505 | 141024139 | 141024257 | 141024425 | 168 | 141024505 | 10079 | 11,13 | 41,49 | 29,25 | 18,13 | 317 | 149 | 3.47E-09 | 8.10E-08 | 0.112,0.111 | 0.431,0.475 | -0.341 |
| 2131 | XLOC_007597 | chr11 | - | 66136530 | 66137003 | 66136530 | 66136670 | 66136839 | 169 | 66137003 | 2131 | 97,101 | 128,111 | 121,93 | 54,54 | 318 | 149 | 1.64E-06 | 2.26E-05 | 0.262,0.299 | 0.512,0.447 | -0.199 |
| 2293 | XLOC_046145 | chr9 | + | 139749882 | 139750601 | 139749882 | 139750300 | 139750469 | 169 | 139750601 | 2293 | 5,4 | 15,6 | 12,12 | 2,1 | 318 | 149 | 7.04E-07 | 1.08E-05 | 0.135,0.238 | 0.738,0.849 | -0.607 |
| 11971 | XLOC_007214 | chr11 | + | 73074791 | 73075284 | 73074791 | 73075047 | 73075216 | 169 | 73075284 | 11971 | 13,13 | 3,1 | 14,8 | 0,0 | 318 | 149 | 0.004425922 | 0.020961085 | 0.67,0.859 | 1.0,1.0 | -0.236 |
| 12846 | XLOC_021518 | chr19 | - | 8660656 | 8661103 | 8660656 | 8660787 | 8660956 | 169 | 8661103 | 12846 | 3,3 | 28,23 | 8,4 | 5,3 | 318 | 149 | 2.34E-05 | 0.000239292 | 0.048,0.058 | 0.428,0.385 | -0.353 |
| 1046 | XLOC_021694 | chr19 | - | 37672053 | 37678146 | 37672053 | 37677830 | 37678000 | 170 | 37678146 | 1046 | 13,12 | 4,1 | 10,2 | 0,0 | 319 | 149 | 0.000242394 | 0.00183164 | 0.603,0.849 | 1.0,1.0 | -0.274 |
| 5446 | XLOC_028673 | chr22 | - | 38895405 | 38895718 | 38895405 | 38895504 | 38895674 | 170 | 38895718 | 5446 | 1,0 | 1,3 | 1,7 | 0,2 | 319 | 149 | 0.000138651 | 0.001130582 | 0.318,0.0 | 1.0,0.62 | -0.651 |
| 8643 | XLOC_000840 | chr1 | - | 1687941 | 1688318 | 1687941 | 1688047 | 1688217 | 170 | 1688318 | 8643 | 38,26 | 8,3 | 33,18 | 0,2 | 319 | 149 | 0.004044006 | 0.019552721 | 0.689,0.802 | 1.0,0.808 | -0.158 |
| 9987 | XLOC_016854 | chr16 | - | 88799995 | 88800458 | 88799995 | 88800153 | 88800323 | 170 | 88800458 | 9987 | 47,28 | 8,4 | 88,56 | 1,4 | 319 | 149 | 0.002695355 | 0.013933565 | 0.733,0.766 | 0.976,0.867 | -0.172 |
| 1018 | XLOC_021778 | chr19 | - | 46274607 | 46274923 | 46274607 | 46274654 | 46274825 | 171 | 46274923 | 1018 | 18,31 | 97,94 | 40,32 | 41,42 | 320 | 149 | 5.96E-07 | 9.30E-06 | 0.08,0.133 | 0.312,0.262 | -0.181 |
| 1868 | XLOC_013325 | chr14 | - | 24034327 | 24035369 | 24034327 | 24035101 | 24035272 | 171 | 24035369 | 1868 | 54,63 | 25,27 | 62,37 | 10,4 | 320 | 149 | 0.000277114 | 0.002067447 | 0.501,0.521 | 0.743,0.812 | -0.267 |
| 5705 | XLOC_027036 | chr20 | - | 49557401 | 49557746 | 49557401 | 49557470 | 49557641 | 171 | 49557746 | 5705 | 6,0 | 7,5 | 9,5 | 0,1 | 320 | 149 | 2.28E-06 | 3.06E-05 | 0.285,0.0 | 1.0,0.7 | -0.708 |
| 5820 | XLOC_026848 | chr20 | + | 57242256 | 57242653 | 57242256 | 57242374 | 57242545 | 171 | 57242653 | 5820 | 14,22 | 30,16 | 41,31 | 11,7 | 320 | 149 | 5.82E-05 | 0.000536241 | 0.179,0.39 | 0.634,0.673 | -0.369 |
| 6401 | XLOC_023741 | chr2 | + | 178306847 | 178307214 | 178306847 | 178306963 | 178307134 | 171 | 178307214 | 6401 | 1,0 | 16,10 | 15,9 | 34,30 | 320 | 149 | 5.25E-05 | 0.000490709 | 0.028,0.0 | 0.17,0.123 | -0.133 |
| 9598 | XLOC_007211 | chr11 | + | 71940706 | 71941028 | 71940706 | 71940796 | 71940967 | 171 | 71941028 | 9598 | 100,92 | 828,875 | 278,173 | 406,309 | 320 | 149 | 0 | 0 | 0.053,0.047 | 0.242,0.207 | -0.174 |
| 1526 | XLOC_016432 | chr16 | + | 66997393 | 66997838 | 66997393 | 66997532 | 66997704 | 172 | 66997838 | 1526 | 59,58 | 9,10 | 162,93 | 4,2 | 321 | 149 | 8.09E-06 | 9.40E-05 | 0.753,0.729 | 0.949,0.956 | -0.211 |
| 2991 | XLOC_013457 | chr14 | - | 74756729 | 74757202 | 74756729 | 74756821 | 74756993 | 172 | 74757202 | 2991 | 20,31 | 57,58 | 27,18 | 31,17 | 321 | 149 | 0.009093321 | 0.038130929 | 0.14,0.199 | 0.288,0.33 | -0.139 |

| | | | | | | | | | | | | | | | | | | | | | | |
|---|---|---|---|---|---|---|---|---|---|---|---|---|---|---|---|---|---|---|---|---|---|---|
| 3176 | XLOC_026791 | chr20 | + | 39985679 | 39986087 | 39985679 | 39985828 | 39986000 | 172 | 39986087 | 3176 | 28,38 | 29,19 | 28,25 | 5,3 | 321 | 149 | 0.00018371 | 0.00144312 | 0.309,0.481 | 0.722,0.795 | -0.363 |
| 6262 | XLOC_024079 | chr2 | - | 70456190 | 70456446 | 70456190 | 70456223 | 70456395 | 172 | 70456446 | 6262 | 90,84 | 81,92 | 186,152 | 113,86 | 321 | 149 | 0.002036179 | 0.011043379 | 0.34,0.298 | 0.433,0.451 | -0.123 |
| 12858 | XLOC_027092 | chr20 | - | 62577202 | 62577620 | 62577202 | 62577341 | 62577513 | 172 | 62577620 | 12858 | 47,22 | 4,7 | 38,32 | 0,2 | 321 | 149 | 0.001476228 | 0.008454487 | 0.845,0.593 | 1.0,0.881 | -0.222 |
| 5794 | XLOC_026877 | chr20 | + | 62368885 | 62369255 | 62368885 | 62369000 | 62369173 | 173 | 62369255 | 5794 | 3,3 | 25,34 | 4,9 | 11,21 | 322 | 149 | 0.006575191 | 0.02920984 | 0.053,0.039 | 0.144,0.165 | -0.108 |
| 11374 | XLOC_032428 | chr4 | + | 119199924 | 119200586 | 119199924 | 119199947 | 119200120 | 173 | 119200586 | 11374 | 46,38 | 1,5 | 130,62 | 1,0 | 322 | 149 | 0.00171788 | 0.009623808 | 0.955,0.779 | 0.984,1.0 | -0.125 |
| 3412 | XLOC_044165 | chr8 | + | 144356873 | 144359124 | 144356873 | 144356972 | 144357147 | 175 | 144359124 | 3412 | 48,23 | 5,1 | 38,41 | 1,0 | 324 | 149 | 0.003905624 | 0.019026949 | 0.815,0.914 | 0.946,1.0 | -0.108 |
| 4783 | XLOC_034699 | chr5 | + | 138268266 | 138268647 | 138268266 | 138268401 | 138268576 | 175 | 138268647 | 4783 | 5,10 | 1,3 | 29,20 | 0,0 | 324 | 149 | 0.000498721 | 0.003430802 | 0.697,0.605 | 1.0,1.0 | -0.349 |
| 11610 | XLOC_021877 | chr19 | - | 55558472 | 55558856 | 55558472 | 55558579 | 55558754 | 175 | 55558856 | 11610 | 25,31 | 4,4 | 19,28 | 0,0 | 324 | 149 | 4.24E-05 | 0.000409029 | 0.742,0.781 | 1.0,1.0 | -0.238 |
| 14542 | XLOC_007550 | chr11 | - | 62602634 | 62603004 | 62602634 | 62602727 | 62602902 | 175 | 62603004 | 14542 | 18,23 | 7,0 | 44,24 | 0,1 | 324 | 149 | 0.002602266 | 0.013496805 | 0.542,1.0 | 1.0,0.917 | -0.188 |
| 3857 | XLOC_046087 | chr9 | + | 131831006 | 131831408 | 131831006 | 131831178 | 131831354 | 176 | 131831408 | 3857 | 4,8 | 4,0 | 7,7 | 0,0 | 325 | 149 | 0.006125792 | 0.027534868 | 0.314,1.0 | 1.0,1.0 | -0.343 |
| 4453 | XLOC_041635 | chr7 | - | 100859188 | 100859583 | 100859188 | 100859301 | 100859477 | 176 | 100859583 | 4453 | 46,43 | 9,9 | 67,71 | 5,4 | 325 | 149 | 0.005898185 | 0.026636114 | 0.701,0.687 | 0.86,0.891 | -0.181 |
| 9594 | XLOC_007211 | chr11 | + | 71939770 | 71940080 | 71939770 | 71939891 | 71940067 | 176 | 71940080 | 9594 | 61,76 | 46,72 | 152,102 | 11,12 | 325 | 149 | 0 | 0 | 0.378,0.326 | 0.864,0.796 | -0.478 |
| 15240 | XLOC_018472 | chr17 | - | 26969886 | 26970244 | 26969886 | 26970005 | 26970181 | 176 | 26970244 | 15240 | 21,13 | 152,218 | 69,45 | 129,98 | 325 | 149 | 1.71E-14 | 9.11E-13 | 0.06,0.027 | 0.197,0.174 | -0.142 |
| 6915 | XLOC_038364 | chr6 | - | 91260186 | 91260504 | 91260186 | 91260268 | 91260445 | 177 | 91260504 | 6915 | 6,3 | 0,4 | 16,12 | 0,0 | 326 | 149 | 0.005749054 | 0.026082201 | 1.0,0.255 | 1.0,1.0 | -0.373 |
| 11500 | XLOC_000830 | chr1 | - | 1333611 | 1334051 | 1333611 | 1333798 | 1333976 | 178 | 1334051 | 11500 | 2,4 | 5,4 | 12,12 | 0,0 | 327 | 149 | 2.31E-10 | 6.69E-09 | 0.154,0.313 | 1.0,1.0 | -0.766 |
| 13509 | XLOC_046097 | chr9 | + | 133346220 | 133346664 | 133346220 | 133346291 | 133346469 | 178 | 133346664 | 13509 | 24,37 | 7,2 | 59,32 | 1,0 | 327 | 149 | 0.000121644 | 0.001015356 | 0.61,0.894 | 0.964,1.0 | -0.23 |
| 10957 | XLOC_009247 | chr12 | - | 9576819 | 9577236 | 9576819 | 9576944 | 9577123 | 179 | 9577236 | 10957 | 11,10 | 17,18 | 17,16 | 6,3 | 328 | 149 | 6.77E-05 | 0.00060793 | 0.227,0.202 | 0.563,0.708 | -0.421 |
| 12279 | XLOC_028747 | chr22 | - | 50723001 | 50723342 | 50723001 | 50723094 | 50723273 | 179 | 50723342 | 12279 | 7,15 | 2,2 | 98,53 | 1,0 | 328 | 149 | 0.002347584 | 0.01240364 | 0.614,0.773 | 0.978,1.0 | -0.295 |
| 4926 | XLOC_032385 | chr4 | + | 88372479 | 88372887 | 88372479 | 88372560 | 88372740 | 180 | 88372887 | 4926 | 3,0 | 0,5 | 1,2 | 0,0 | 329 | 149 | 0.000874721 | 0.00544695 | 1.0,0.0 | 1.0,1.0 | -0.5 |
| 7832 | XLOC_018703 | chr17 | - | 72769714 | 72770078 | 72769714 | 72769824 | 72770004 | 180 | 72770078 | 7832 | 0,3 | 13,7 | 17,11 | 21,9 | 329 | 149 | 0.000339004 | 0.002461974 | 0.0,0.163 | 0.268,0.356 | -0.23 |
| 3548 | XLOC_048083 | chrX | - | 130928021 | 130928494 | 130928021 | 130928143 | 130928324 | 181 | 130928494 | 3548 | 1,0 | 8,3 | 7,6 | 12,15 | 330 | 149 | 0.00090846 | 0.005622343 | 0.053,0.0 | 0.208,0.153 | -0.154 |
| 6349 | XLOC_023898 | chr2 | + | 234356503 | 234356827 | 234356503 | 234356551 | 234356732 | 181 | 234356827 | 6349 | 5,14 | 67,68 | 90,41 | 62,28 | 330 | 149 | 0 | 0 | 0.033,0.085 | 0.396,0.398 | -0.338 |
| 6979 | XLOC_021281 | chr19 | + | 47761586 | 47761967 | 47761586 | 47761660 | 47761842 | 182 | 47761967 | 6979 | 23,21 | 2,5 | 21,12 | 0,0 | 331 | 149 | 5.62E-05 | 0.000520658 | 0.838,0.654 | 1.0,1.0 | -0.254 |
| 8723 | XLOC_016204 | chr16 | + | 1824947 | 1825409 | 1824947 | 1825133 | 1825315 | 182 | 1825409 | 8723 | 38,29 | 11,9 | 18,8 | 2,0 | 331 | 149 | 0.004273884 | 0.020378143 | 0.609,0.592 | 0.802,1.0 | -0.3 |
| 10953 | XLOC_009247 | chr12 | - | 9571370 | 9571781 | 9571370 | 9571525 | 9571707 | 182 | 9571781 | 10953 | 16,29 | 36,20 | 52,42 | 15,13 | 331 | 149 | 0.000188974 | 0.001474516 | 0.167,0.395 | 0.609,0.593 | -0.32 |
| 13411 | XLOC_024005 | chr2 | - | 38976381 | 38976847 | 38976381 | 38976488 | 38976670 | 182 | 38976847 | 13411 | 267,305 | 118,139 | 365,298 | 75,102 | 331 | 149 | 0.00536553 | 0.024659091 | 0.505,0.497 | 0.687,0.568 | -0.126 |
| 14408 | XLOC_001334 | chr1 | - | 153913379 | 153913870 | 153913379 | 153913524 | 153913706 | 182 | 153913870 | 14408 | 32,33 | 66,60 | 193,107 | 40,30 | 331 | 149 | 0 | 0 | 0.179,0.198 | 0.685,0.616 | -0.462 |
| 4050 | XLOC_044256 | chr8 | - | 28213925 | 28214296 | 28213925 | 28214003 | 28214186 | 183 | 28214296 | 4050 | 10,4 | 3,4 | 4,11 | 0,0 | 332 | 149 | 4.20E-07 | 6.79E-06 | 0.599,0.31 | 1.0,1.0 | -0.545 |
| 4825 | XLOC_034481 | chr5 | + | 647919 | 648338 | 647919 | 648031 | 648214 | 183 | 648338 | 4825 | 12,10 | 18,17 | 77,52 | 45,39 | 332 | 149 | 0.002960211 | 0.015085357 | 0.23,0.209 | 0.434,0.374 | -0.185 |
| 4985 | XLOC_034868 | chr5 | - | 36225358 | 36225725 | 36225358 | 36225460 | 36225643 | 183 | 36225725 | 4985 | 16,21 | 4,1 | 11,14 | 0,0 | 332 | 149 | 0.000732291 | 0.004695267 | 0.642,0.904 | 1.0,1.0 | -0.227 |
| 11424 | XLOC_038395 | chr6 | - | 109766973 | 109767615 | 109766973 | 109767155 | 109767338 | 183 | 109767615 | 11424 | 0,1 | 3,1 | 1,1 | 0,0 | 332 | 149 | 4.59E-09 | 1.06E-07 | 0.0,0.31 | 1.0,1.0 | -0.845 |
| 1841 | XLOC_007551 | chr11 | - | 62621272 | 62621536 | 62621272 | 62621305 | 62621489 | 184 | 62621536 | 1841 | 334,312 | 565,530 | 594,470 | 563,404 | 333 | 149 | 1.66E-07 | 2.90E-06 | 0.209,0.208 | 0.321,0.342 | -0.123 |
| 10250 | XLOC_026869 | chr20 | + | 61429938 | 61430400 | 61429938 | 61430020 | 61430204 | 184 | 61430400 | 10250 | 18,17 | 15,18 | 7,14 | 0,4 | 333 | 149 | 0.006336936 | 0.028330657 | 0.349,0.297 | 1.0,0.61 | -0.482 |
| 2168 | XLOC_009600 | chr12 | - | 123745344 | 123746350 | 123745344 | 123745490 | 123745675 | 185 | 123746350 | 2168 | 230,215 | 26,8 | 184,144 | 0,3 | 334 | 149 | 2.07E-06 | 2.80E-05 | 0.798,0.923 | 1.0,0.955 | -0.117 |
| 11231 | XLOC_008778 | chr12 | + | 9446711 | 9447114 | 9446711 | 9446842 | 9447027 | 185 | 9447114 | 11231 | 13,12 | 23,33 | 39,20 | 14,14 | 334 | 149 | 3.65E-05 | 0.000358002 | 0.201,0.14 | 0.554,0.389 | -0.301 |
| 12857 | XLOC_027092 | chr20 | - | 62577202 | 62577620 | 62577202 | 62577328 | 62577513 | 185 | 62577620 | 12857 | 105,45 | 141,127 | 91,71 | 72,41 | 334 | 149 | 6.04E-06 | 7.27E-05 | 0.249,0.136 | 0.361,0.436 | -0.206 |
| 15276 | XLOC_009486 | chr12 | - | 94706347 | 94706793 | 94706347 | 94706504 | 94706689 | 185 | 94706793 | 15276 | 0,1 | 3,2 | 5,1 | 2,5 | 334 | 149 | 0.008949963 | 0.037686441 | 0.0,0.182 | 0.527,0.082 | -0.213 |
| 5903 | XLOC_026682 | chr20 | + | 2636783 | 2637195 | 2636783 | 2636860 | 2637046 | 186 | 2637195 | 5903 | 159,136 | 157,144 | 244,216 | 128,130 | 335 | 149 | 6.66E-05 | 0.000600291 | 0.311,0.296 | 0.459,0.425 | -0.139 |
| 6319 | XLOC_023980 | chr2 | - | 27590288 | 27590689 | 27590288 | 27590426 | 27590612 | 186 | 27590689 | 6319 | 10,7 | 126,135 | 29,39 | 104,90 | 335 | 149 | 7.30E-10 | 1.95E-08 | 0.034,0.023 | 0.11,0.162 | -0.108 |
| 10868 | XLOC_001244 | chr1 | - | 113246005 | 113246428 | 113246005 | 113246079 | 113246265 | 186 | 113246428 | 10868 | 11,4 | 2,3 | 5,5 | 0,1 | 335 | 149 | 0.003517223 | 0.017462554 | 0.71,0.372 | 1.0,0.69 | -0.304 |
| 13383 | XLOC_037875 | chr6 | + | 52957228 | 52957622 | 52957228 | 52957345 | 52957531 | 186 | 52957622 | 13383 | 12,12 | 189,132 | 246,194 | 157,97 | 335 | 149 | 0 | 0 | 0.027,0.039 | 0.411,0.471 | -0.408 |
| 7763 | XLOC_018482 | chr17 | - | 27445092 | 27445453 | 27445092 | 27445207 | 27445394 | 187 | 27445453 | 7763 | 10,6 | 106,101 | 31,37 | 33,30 | 336 | 149 | 0 | 0 | 0.04,0.026 | 0.294,0.354 | -0.291 |
| 8338 | XLOC_016862 | chr16 | - | 89805536 | 89805961 | 89805536 | 89805697 | 89805885 | 188 | 89805961 | 8338 | 199,186 | 213,188 | 350,265 | 175,122 | 337 | 149 | 4.70E-08 | 9.12E-07 | 0.292,0.304 | 0.469,0.49 | -0.181 |
| 9101 | XLOC_014945 | chr15 | - | 65261277 | 65261699 | 65261277 | 65261403 | 65261591 | 188 | 65261699 | 9101 | 0,0 | 4,5 | 6,1 | 0,5 | 337 | 149 | 0.002539878 | 0.013232141 | 0.0,0.0 | 1.0,0.081 | -0.54 |
| 8112 | XLOC_017871 | chr17 | + | 4719986 | 4720350 | 4719986 | 4720041 | 4720231 | 190 | 4720350 | 8112 | 13,2 | 88,104 | 15,8 | 12,30 | 339 | 149 | 2.34E-05 | 0.000239648 | 0.061,0.008 | 0.355,0.105 | -0.195 |
| 1324 | XLOC_005229 | chr10 | - | 58112902 | 58118220 | 58112902 | 58117946 | 58118137 | 191 | 58118220 | 1324 | 96,69 | 360,362 | 110,107 | 133,81 | 340 | 149 | 2.22E-16 | 1.46E-14 | 0.105,0.077 | 0.266,0.367 | -0.226 |
| 3969 | XLOC_044462 | chr8 | - | 145652291 | 145652654 | 145652291 | 145652362 | 145652554 | 192 | 145652654 | 3969 | 37,26 | 2,3 | 14,11 | 0,0 | 341 | 149 | 0.003946658 | 0.01919375 | 0.89,0.791 | 1.0,1.0 | -0.159 |
| 7886 | XLOC_018261 | chr17 | + | 76200908 | 76201271 | 76200908 | 76200981 | 76201173 | 192 | 76201271 | 7886 | 21,16 | 9,10 | 40,14 | 0,0 | 341 | 149 | 3.13E-10 | 8.85E-09 | 0.505,0.411 | 1.0,1.0 | -0.542 |
| 8739 | XLOC_016191 | chr16 | + | 778503 | 778872 | 778503 | 778604 | 778796 | 192 | 778872 | 8739 | 36,19 | 38,23 | 64,28 | 20,7 | 341 | 149 | 4.84E-06 | 5.99E-05 | 0.293,0.265 | 0.583,0.636 | -0.33 |
| 10385 | XLOC_023860 | chr2 | + | 220098436 | 220098910 | 220098436 | 220098665 | 220098857 | 192 | 220098910 | 10385 | 3,5 | 15,16 | 14,9 | 6,1 | 341 | 149 | 1.50E-05 | 0.000161842 | 0.08,0.12 | 0.505,0.797 | -0.551 |
| 12584 | XLOC_000173 | chr1 | + | 28887379 | 28887772 | 28887379 | 28887432 | 28887624 | 192 | 28887772 | 12584 | 7,12 | 2,3 | 11,2 | 0,0 | 341 | 149 | 0.000166313 | 0.001323661 | 0.605,0.636 | 1.0,1.0 | -0.379 |
| 6333 | XLOC_017917 | chr17 | + | 7850931 | 7851297 | 7850931 | 7851040 | 7851233 | 193 | 7851297 | 6333 | 17,17 | 130,170 | 57,34 | 96,35 | 342 | 149 | 1.32E-13 | 6.34E-12 | 0.054,0.042 | 0.206,0.297 | -0.204 |
| 7195 | XLOC_016577 | chr16 | - | 2162788 | 2163285 | 2162788 | 2162964 | 2163157 | 193 | 2163285 | 7195 | 8,2 | 21,13 | 9,8 | 6,7 | 342 | 149 | 0.002703512 | 0.013950199 | 0.142,0.063 | 0.395,0.332 | -0.261 |
| 10363 | XLOC_021489 | chr19 | - | 5610005 | 5610432 | 5610005 | 5610106 | 5610299 | 193 | 5610432 | 10363 | 10,9 | 10,25 | 29,14 | 7,2 | 342 | 149 | 4.47E-06 | 5.59E-05 | 0.303,0.136 | 0.643,0.753 | -0.478 |
| 1377 | XLOC_014991 | chr15 | - | 75651668 | 75652116 | 75651668 | 75651766 | 75651960 | 194 | 75652116 | 1377 | 316,279 | 32,53 | 314,206 | 13,13 | 343 | 149 | 5.62E-05 | 0.00052064 | 0.811,0.696 | 0.913,0.873 | -0.139 |
| 12195 | XLOC_000006 | chr1 | + | 986105 | 986423 | 986105 | 986217 | 986411 | 194 | 986423 | 12195 | 2,0 | 0,4 | 15,4 | 1,0 | 343 | 149 | 0.002002039 | 0.010887485 | 1.0,0.0 | 0.867,1.0 | -0.433 |
| 13499 | XLOC_007598 | chr11 | - | 66204272 | 66204734 | 66204272 | 66204380 | 66204574 | 194 | 66204734 | 13499 | 4,15 | 17,29 | 27,18 | 16,17 | 343 | 149 | 0.001080111 | 0.00651063 | 0.093,0.183 | 0.423,0.315 | -0.231 |
| 5706 | XLOC_027036 | chr20 | - | 49557401 | 49557746 | 49557401 | 49557470 | 49557665 | 195 | 49557746 | 5706 | 6,1 | 17,28 | 9,5 | 9,2 | 344 | 149 | 0.000414698 | 0.002925466 | 0.133,0.015 | 0.302,0.52 | -0.337 |
| 7515 | XLOC_018797 | chr17 | - | 80584871 | 80585167 | 80584871 | 80584884 | 80585079 | 195 | 80585167 | 7515 | 39,29 | 15,14 | 20,23 | 3,2 | 344 | 149 | 0.003352461 | 0.016806076 | 0.53,0.473 | 0.743,0.833 | -0.286 |
| 10008 | XLOC_044467 | chr8 | - | 145753407 | 145753819 | 145753407 | 145753456 | 145753651 | 195 | 145753819 | 10008 | 38,38 | 4,3 | 83,46 | 0,0 | 344 | 149 | 0.000120496 | 0.001006963 | 0.804,0.846 | 1.0,1.0 | -0.175 |
| 11054 | XLOC_038073 | chr6 | - | 485008 | 486764 | 485008 | 486403 | 486598 | 195 | 486764 | 11054 | 90,97 | 6,4 | 112,88 | 0,0 | 344 | 149 | 9.88E-05 | 0.000849213 | 0.867,0.913 | 1.0,1.0 | -0.11 |
| 3811 | XLOC_046241 | chr9 | - | 35740568 | 35741061 | 35740568 | 35740625 | 35740821 | 196 | 35741061 | 3811 | 4,7 | 102,98 | 120,79 | 91,72 | 345 | 149 | 0 | 0 | 0.017,0.03 | 0.363,0.322 | -0.319 |
| 9313 | XLOC_018520 | chr17 | - | 37009274 | 37009967 | 37009274 | 37009358 | 37009554 | 196 | 37009967 | 9313 | 23,24 | 4,2 | 21,31 | 0,1 | 345 | 149 | 0.001414367 | 0.008169777 | 0.713,0.838 | 1.0,0.93 | -0.19 |
| 7196 | XLOC_016577 | chr16 | - | 2162788 | 2163285 | 2162788 | 2162964 | 2163161 | 197 | 2163285 | 7196 | 8,3 | 53,45 | 10,8 | 12,14 | 346 | 149 | 5.03E-05 | 0.000472687 | 0.061,0.028 | 0.264,0.197 | -0.186 |
| 12523 | XLOC_011543 | chr13 | - | 100622022 | 100626405 | 100622022 | 100622183 | 100622380 | 197 | 100626405 | 12523 | 0,0 | 24,11 | 6,4 | 2,5 | 346 | 149 | 3.95E-07 | 6.44E-06 | 0.0,0.0 | 0.564,0.256 | -0.41 |

| | | | | | | | | | | | | | | | | | | | | | | |
|---|---|---|---|---|---|---|---|---|---|---|---|---|---|---|---|---|---|---|---|---|---|---|
| 13527 | XLOC_013457 | chr14 | - | 74755845 | 74756821 | 74755845 | 74755988 | 74756185 | 197 | 74756821 | 13527 | 16,14 | 5,2 | 14,7 | 0,1 | 346 | 149 | 0.001793318 | 0.009955592 | 0.579,0.751 | 1.0,0.751 | -0.21 |
| 5278 | XLOC_029529 | chr3 | + | 113666866 | 113667309 | 113666866 | 113667037 | 113667235 | 198 | 113667309 | 5278 | 9,13 | 4,0 | 7,1 | 0,0 | 347 | 149 | 0.007216942 | 0.031477725 | 0.491,1.0 | 1.0,1.0 | -0.254 |
| 10574 | XLOC_013429 | chr14 | - | 68059031 | 68059435 | 68059031 | 68059153 | 68059351 | 198 | 68059435 | 10574 | 0,0 | 2,4 | 9,7 | 11,8 | 347 | 149 | 3.96E-05 | 0.000384567 | 0.0,0.0 | 0.26,0.273 | -0.267 |
| 1565 | XLOC_016191 | chr16 | + | 779008 | 779836 | 779008 | 779088 | 779287 | 199 | 779836 | 1565 | 364,295 | 313,323 | 639,366 | 179,114 | 348 | 149 | 0 | 0 | 0.332,0.281 | 0.605,0.579 | -0.285 |
| 2128 | XLOC_007597 | chr11 | - | 66133596 | 66134035 | 66133596 | 66133702 | 66133901 | 199 | 66134035 | 2128 | 78,77 | 106,130 | 128,106 | 30,30 | 348 | 149 | 0 | 0 | 0.24,0.202 | 0.646,0.602 | -0.403 |
| 13246 | XLOC_009336 | chr12 | - | 49166290 | 49166597 | 49166290 | 49166326 | 49166525 | 199 | 49166597 | 13246 | 20,16 | 3,2 | 23,15 | 0,0 | 348 | 149 | 0.000659957 | 0.004339187 | 0.741,0.774 | 1.0,1.0 | -0.242 |
| 4424 | XLOC_038512 | chr6 | - | 170887645 | 170888050 | 170887645 | 170887744 | 170887944 | 200 | 170888050 | 4424 | 6,19 | 53,52 | 90,61 | 40,44 | 349 | 149 | 2.29E-12 | 9.28E-11 | 0.046,0.135 | 0.49,0.372 | -0.341 |
| 7383 | XLOC_041349 | chr7 | + | 150769028 | 150769521 | 150769028 | 150769223 | 150769423 | 200 | 150769521 | 7383 | 9,8 | 3,5 | 7,18 | 0,0 | 349 | 149 | 1.88E-07 | 3.25E-06 | 0.562,0.406 | 1.0,1.0 | -0.516 |
| 10983 | XLOC_029823 | chr3 | - | 48669312 | 48669770 | 48669312 | 48669477 | 48669677 | 200 | 48669770 | 10983 | 16,32 | 52,58 | 140,82 | 28,20 | 349 | 149 | 0 | 0 | 0.116,0.191 | 0.681,0.636 | -0.505 |
| 11741 | XLOC_007543 | chr11 | - | 62407098 | 62407538 | 62407098 | 62407203 | 62407403 | 200 | 62407538 | 11741 | 13,17 | 2,9 | 15,6 | 0,1 | 349 | 149 | 0.000558676 | 0.003780633 | 0.735,0.446 | 1.0,0.719 | -0.269 |
| 12286 | XLOC_007578 | chr11 | - | 64900630 | 64901089 | 64900630 | 64900740 | 64900940 | 200 | 64901089 | 12286 | 0,6 | 2,1 | 23,8 | 0,2 | 349 | 149 | 0.007215792 | 0.031477725 | 0.0,0.719 | 1.0,0.631 | -0.456 |
| 13290 | XLOC_023538 | chr2 | + | 74719430 | 74719815 | 74719430 | 74719572 | 74719772 | 200 | 74719815 | 13290 | 21,14 | 55,75 | 35,35 | 26,41 | 349 | 149 | 1.09E-06 | 1.57E-05 | 0.14,0.074 | 0.365,0.267 | -0.209 |
| 15027 | XLOC_021451 | chr19 | - | 1615422 | 1615820 | 1615422 | 1615484 | 1615684 | 200 | 1615820 | 15027 | 7,12 | 17,14 | 17,16 | 5,1 | 349 | 149 | 6.52E-05 | 0.000589946 | 0.15,0.268 | 0.592,0.872 | -0.523 |
| 3705 | XLOC_046491 | chr9 | - | 139934447 | 139934888 | 139934447 | 139934607 | 139934808 | 201 | 139934888 | 3705 | 69,82 | 62,64 | 41,33 | 3,13 | 350 | 149 | 0.002065833 | 0.01117845 | 0.321,0.353 | 0.853,0.519 | -0.349 |
| 11887 | XLOC_009066 | chr12 | + | 104376597 | 104376996 | 104376597 | 104376712 | 104376913 | 201 | 104376996 | 11887 | 63,35 | 231,222 | 151,121 | 136,137 | 350 | 149 | 0 | 0 | 0.104,0.063 | 0.321,0.273 | -0.214 |
| 1342 | XLOC_018241 | chr17 | + | 73570690 | 73571275 | 73570690 | 73570749 | 73570951 | 202 | 73571275 | 1342 | 129,88 | 142,185 | 100,72 | 105,49 | 351 | 149 | 0.011531906 | 0.046719768 | 0.278,0.168 | 0.288,0.384 | -0.113 |
| 2290 | XLOC_046145 | chr9 | + | 139748662 | 139749128 | 139748662 | 139748788 | 139748990 | 202 | 139749128 | 2290 | 8,7 | 12,16 | 14,16 | 8,6 | 351 | 149 | 0.00177075 | 0.009861318 | 0.221,0.157 | 0.426,0.531 | -0.29 |
| 5788 | XLOC_026895 | chr20 | - | 480168 | 480578 | 480168 | 480274 | 480476 | 202 | 480578 | 5788 | 1,0 | 2,1 | 2,6 | 0,4 | 351 | 149 | 0.001405358 | 0.008127709 | 0.175,0.0 | 1.0,0.389 | -0.607 |
| 6320 | XLOC_023980 | chr2 | - | 27590612 | 27591069 | 27590612 | 27590689 | 27590891 | 202 | 27591069 | 6320 | 17,29 | 110,133 | 61,41 | 99,86 | 351 | 149 | 1.02E-06 | 1.50E-05 | 0.062,0.085 | 0.207,0.168 | -0.114 |
| 6595 | XLOC_046049 | chr9 | + | 127962698 | 127963181 | 127962698 | 127962856 | 127963058 | 202 | 127963181 | 6595 | 24,31 | 19,20 | 69,33 | 16,10 | 351 | 149 | 0.000962407 | 0.005909599 | 0.349,0.397 | 0.647,0.583 | -0.242 |
| 9918 | XLOC_034700 | chr5 | + | 138609800 | 138610118 | 138609800 | 138609850 | 138610052 | 202 | 138610118 | 9918 | 2,4 | 6,1 | 8,4 | 0,0 | 351 | 149 | 2.09E-07 | 3.58E-06 | 0.124,0.629 | 1.0,1.0 | -0.623 |
| 13785 | XLOC_029849 | chr3 | - | 49723749 | 49724235 | 49723749 | 49723914 | 49724116 | 202 | 49724235 | 13785 | 73,47 | 6,9 | 214,154 | 6,1 | 351 | 149 | 5.91E-05 | 0.000542955 | 0.838,0.689 | 0.938,0.985 | -0.198 |
| 15221 | XLOC_048116 | chrX | - | 153051093 | 153051727 | 153051093 | 153051464 | 153051666 | 202 | 153051727 | 15221 | 21,14 | 0,4 | 39,19 | 0,0 | 351 | 149 | 0.011622694 | 0.047033611 | 1.0,0.598 | 1.0,1.0 | -0.201 |
| 442 | XLOC_034514 | chr5 | + | 35709049 | 35710594 | 35709049 | 35709223 | 35709426 | 203 | 35710594 | 442 | 17,23 | 2,4 | 2,2 | 0,0 | 352 | 149 | 0.000266031 | 0.00199556 | 0.783,0.709 | 1.0,1.0 | -0.254 |
| 6792 | XLOC_009222 | chr12 | - | 6673056 | 6673414 | 6673056 | 6673112 | 6673315 | 203 | 6673414 | 6792 | 1,1 | 8,4 | 8,6 | 0,1 | 352 | 149 | 6.89E-09 | 1.53E-07 | 0.05,0.096 | 1.0,0.717 | -0.786 |
| 6508 | XLOC_023405 | chr2 | + | 27448572 | 27449041 | 27448572 | 27448794 | 27448998 | 204 | 27449041 | 6508 | 5,15 | 225,216 | 73,64 | 189,137 | 353 | 149 | 0 | 0 | 0.009,0.028 | 0.14,0.165 | -0.134 |
| 12897 | XLOC_046038 | chr9 | + | 123933436 | 123933826 | 123933436 | 123933481 | 123933685 | 204 | 123933826 | 12897 | 0,1 | 2,0 | 2,2 | 1,0 | 353 | 149 | 0.001188072 | 0.007044722 | 0.0,1.0 | 0.458,1.0 | -0.229 |
| 13059 | XLOC_004753 | chr10 | + | 3141466 | 3141891 | 3141466 | 3141544 | 3141748 | 204 | 3141891 | 13059 | 1,5 | 17,21 | 27,27 | 65,37 | 353 | 149 | 0.001170567 | 0.006961179 | 0.024,0.091 | 0.149,0.235 | -0.135 |
| 1103 | XLOC_021261 | chr19 | + | 45556048 | 45556435 | 45556048 | 45556150 | 45556355 | 205 | 45556435 | 1103 | 22,11 | 272,219 | 118,41 | 131,93 | 354 | 149 | 0 | 0 | 0.033,0.021 | 0.275,0.157 | -0.189 |
| 2532 | XLOC_007173 | chr11 | + | 66366586 | 66367107 | 66366586 | 66366724 | 66366929 | 205 | 66367107 | 2532 | 89,136 | 79,92 | 147,99 | 54,42 | 354 | 149 | 0.000287809 | 0.002138204 | 0.322,0.384 | 0.534,0.498 | -0.163 |
| 4680 | XLOC_035171 | chr5 | - | 179047888 | 179048349 | 179047888 | 179048036 | 179048242 | 206 | 179048349 | 4680 | 832,718 | 17,331,697 | 16,971,143 | 16,421,297 | 355 | 149 | 3.32E-12 | 1.29E-10 | 0.168,0.151 | 0.303,0.27 | -0.127 |
| 5986 | XLOC_047690 | chrX | + | 47085981 | 47086462 | 47085981 | 47086102 | 47086308 | 206 | 47086462 | 5986 | 16,21 | 4,2 | 108,89 | 3,0 | 355 | 149 | 0.001389606 | 0.008049794 | 0.627,0.815 | 0.938,1.0 | -0.248 |
| 11099 | XLOC_008988 | chr12 | + | 64889483 | 64889838 | 64889483 | 64889555 | 64889761 | 206 | 64889838 | 11099 | 0,6 | 5,9 | 16,8 | 3,2 | 355 | 149 | 5.68E-05 | 0.000525273 | 0.0,0.219 | 0.691,0.627 | -0.549 |
| 11176 | XLOC_008874 | chr12 | + | 50036712 | 50037180 | 50036712 | 50036799 | 50037005 | 206 | 50037180 | 11176 | 3,5 | 3,4 | 9,1 | 0,0 | 355 | 149 | 5.14E-07 | 8.13E-06 | 0.296,0.344 | 1.0,1.0 | -0.68 |
| 13265 | XLOC_017927 | chr17 | + | 12032455 | 12032901 | 12032455 | 12032604 | 12032810 | 206 | 12032901 | 13265 | 0,0 | 3,2 | 6,2 | 0,0 | 355 | 149 | 4.95E-11 | 1.60E-09 | 0.0,0.0 | 1.0,1.0 | -1 |
| 1843 | XLOC_007551 | chr11 | - | 62622604 | 62623342 | 62622604 | 62622688 | 62622895 | 207 | 62623342 | 1843 | 183,209 | 366,349 | 411,333 | 390,308 | 356 | 149 | 2.30E-07 | 3.90E-06 | 0.173,0.2 | 0.306,0.312 | -0.122 |
| 2197 | XLOC_009410 | chr12 | - | 56720096 | 56720743 | 56720096 | 56720193 | 56720400 | 207 | 56720743 | 2197 | 27,38 | 92,80 | 41,29 | 35,24 | 356 | 149 | 4.08E-06 | 5.14E-05 | 0.109,0.166 | 0.329,0.336 | -0.195 |
| 2470 | XLOC_021203 | chr19 | + | 39873847 | 39876316 | 39873847 | 39873931 | 39874138 | 207 | 39876316 | 2470 | 136,145 | 105,91 | 299,259 | 57,39 | 356 | 149 | 3.33E-16 | 2.11E-14 | 0.352,0.4 | 0.687,0.735 | -0.335 |
| 6599 | XLOC_021230 | chr19 | + | 42396095 | 42396443 | 42396095 | 42396194 | 42396401 | 207 | 42396443 | 6599 | 48,27 | 153,131 | 83,67 | 74,88 | 356 | 149 | 3.93E-10 | 1.09E-08 | 0.116,0.079 | 0.319,0.242 | -0.183 |
| 9617 | XLOC_007211 | chr11 | + | 71944083 | 71944566 | 71944083 | 71944277 | 71944484 | 207 | 71944566 | 9617 | 3,7 | 2,1 | 26,20 | 1,0 | 356 | 149 | 0.00876462 | 0.037087325 | 0.386,0.746 | 0.916,1.0 | -0.392 |
| 13510 | XLOC_046097 | chr9 | + | 133346469 | 133346902 | 133346469 | 133346664 | 133346871 | 207 | 133346902 | 13510 | 24,39 | 5,7 | 67,33 | 3,1 | 356 | 149 | 0.002518049 | 0.013151775 | 0.668,0.7 | 0.903,0.932 | -0.234 |
| 14630 | XLOC_016334 | chr16 | + | 27355914 | 27356341 | 27355914 | 27355982 | 27356189 | 207 | 27356341 | 14630 | 2,0 | 2,7 | 2,2 | 0,0 | 356 | 149 | 2.88E-12 | 1.14E-10 | 0.295,0.0 | 1.0,1.0 | -0.853 |
| 14826 | XLOC_007017 | chr11 | + | 18418026 | 18418515 | 18418026 | 18418157 | 18418365 | 208 | 18418515 | 14826 | 14,17 | 13,11 | 64,13 | 3,0 | 357 | 149 | 1.55E-09 | 3.92E-08 | 0.31,0.392 | 0.899,1.0 | -0.599 |
| 454 | XLOC_041145 | chr7 | + | 75044869 | 75045469 | 75044869 | 75045048 | 75045258 | 210 | 75045469 | 454 | 211,241 | 53,34 | 422,277 | 23,20 | 359 | 149 | 3.36E-06 | 4.31E-05 | 0.623,0.746 | 0.884,0.852 | -0.183 |
| 4219 | XLOC_041532 | chr7 | - | 72469989 | 72470382 | 72469989 | 72470073 | 72470283 | 210 | 72470382 | 4219 | 0,4 | 12,17 | 8,13 | 1,3 | 359 | 149 | 2.36E-09 | 5.71E-08 | 0.0,0.089 | 0.769,0.643 | -0.661 |
| 4590 | XLOC_037801 | chr6 | + | 35262301 | 35262684 | 35262301 | 35262346 | 35262556 | 210 | 35262684 | 4590 | 55,48 | 1,13 | 63,39 | 1,1 | 359 | 149 | 0.003847358 | 0.018823129 | 0.958,0.605 | 0.963,0.942 | -0.171 |
| 11911 | XLOC_000835 | chr1 | - | 1647784 | 1648250 | 1647784 | 1647917 | 1648127 | 210 | 1648250 | 11911 | 8,4 | 3,1 | 11,9 | 0,0 | 359 | 149 | 0.000263498 | 0.001980515 | 0.525,0.624 | 1.0,1.0 | -0.425 |
| 1267 | XLOC_018771 | chr17 | - | 79218117 | 79220861 | 79218117 | 79220473 | 79220684 | 211 | 79220861 | 1267 | 45,32 | 149,126 | 73,42 | 47,31 | 360 | 149 | 1.14E-14 | 6.26E-13 | 0.111,0.095 | 0.391,0.359 | -0.272 |
| 13514 | XLOC_020165 | chr18 | - | 9593765 | 9594232 | 9593765 | 9593872 | 9594084 | 212 | 9594232 | 13514 | 23,12 | 1,4 | 22,9 | 0,0 | 361 | 149 | 0.000503019 | 0.003453641 | 0.905,0.553 | 1.0,1.0 | -0.271 |
| 1499 | XLOC_016560 | chr16 | - | 775412 | 775880 | 775412 | 775580 | 775793 | 213 | 775880 | 1499 | 8,10 | 18,30 | 16,21 | 19,17 | 362 | 149 | 0.005089477 | 0.023574009 | 0.155,0.121 | 0.257,0.337 | -0.159 |
| 3765 | XLOC_046425 | chr9 | - | 130690343 | 130690762 | 130690343 | 130690498 | 130690711 | 213 | 130690762 | 3765 | 0,0 | 2,1 | 4,1 | 1,1 | 362 | 149 | 0.000215521 | 0.001656038 | 0.0,0.0 | 0.622,0.292 | -0.457 |
| 4335 | XLOC_041489 | chr7 | - | 44113729 | 44114129 | 44113729 | 44113783 | 44113996 | 213 | 44114129 | 4335 | 24,45 | 5,1 | 48,30 | 0,0 | 362 | 149 | 0.000371492 | 0.002664592 | 0.664,0.949 | 1.0,1.0 | -0.194 |
| 6383 | XLOC_023816 | chr2 | + | 207025822 | 207026196 | 207025822 | 207025856 | 207026069 | 213 | 207026196 | 6383 | 30,36 | 22,16 | 126,67 | 29,19 | 362 | 149 | 0.005081113 | 0.02354928 | 0.359,0.481 | 0.641,0.592 | -0.197 |
| 9218 | XLOC_014753 | chr15 | + | 85663761 | 85664245 | 85663761 | 85663814 | 85664027 | 213 | 85664245 | 9218 | 11,18 | 14,5 | 17,22 | 4,0 | 362 | 149 | 0.006823451 | 0.030090657 | 0.244,0.597 | 0.636,1.0 | -0.398 |
| 165 | XLOC_041517 | chr7 | - | 65439512 | 65440058 | 65439512 | 65439691 | 65439905 | 214 | 65440058 | 165 | 83,76 | 232,188 | 93,78 | 75,75 | 363 | 149 | 3.96E-10 | 1.10E-08 | 0.128,0.142 | 0.337,0.299 | -0.183 |
| 9292 | XLOC_014566 | chr15 | + | 40681366 | 40681812 | 40681366 | 40681492 | 40681706 | 214 | 40681812 | 9292 | 19,8 | 5,4 | 1,8 | 0,1 | 363 | 149 | 0.002611645 | 0.013540479 | 0.609,0.451 | 1.0,0.767 | -0.353 |
| 13574 | XLOC_000240 | chr1 | + | 41232267 | 41232646 | 41232267 | 41232375 | 41232589 | 214 | 41232646 | 13574 | 0,7 | 34,22 | 8,7 | 19,14 | 363 | 149 | 0.008504458 | 0.036188149 | 0.0,0.116 | 0.147,0.17 | -0.101 |
| 156 | XLOC_041620 | chr7 | - | 99752884 | 99754172 | 99752884 | 99753078 | 99753293 | 215 | 99754172 | 156 | 22,19 | 21,43 | 47,37 | 29,19 | 364 | 149 | 0.001835804 | 0.010158358 | 0.3,0.153 | 0.399,0.444 | -0.195 |
| 1348 | XLOC_018130 | chr17 | + | 46052879 | 46053379 | 46052879 | 46053019 | 46053234 | 215 | 46053379 | 1348 | 644,646 | 160,160 | 448,396 | 53,35 | 364 | 149 | 3.34E-08 | 6.62E-07 | 0.622,0.623 | 0.776,0.822 | -0.176 |
| 3833 | XLOC_046149 | chr9 | + | 139972501 | 139973129 | 139972501 | 139972738 | 139972953 | 215 | 139973129 | 3833 | 17,5 | 5,6 | 7,3 | 0,0 | 364 | 149 | 6.87E-07 | 1.05E-05 | 0.582,0.254 | 1.0,1.0 | -0.582 |
| 9314 | XLOC_023615 | chr2 | + | 109085435 | 109085788 | 109085435 | 109085540 | 109085755 | 215 | 109085788 | 9314 | 1,2 | 0,6 | 3,5 | 0,1 | 364 | 149 | 0.00387833 | 0.018913552 | 1.0,0.12 | 1.0,0.672 | -0.276 |
| 3372 | XLOC_009413 | chr12 | - | 56814746 | 56815277 | 56814746 | 56814918 | 56815134 | 216 | 56815277 | 3372 | 108,103 | 399,382 | 87,89 | 149,133 | 365 | 149 | 1.60E-07 | 2.81E-06 | 0.1,0.099 | 0.192,0.215 | -0.104 |
| 6511 | XLOC_023405 | chr2 | + | 27459616 | 27460034 | 27459616 | 27459698 | 27459914 | 216 | 27460034 | 6511 | 0,0 | 3,6 | 7,3 | 12,5 | 365 | 149 | 0.000126705 | 0.001048306 | 0.0,0.0 | 0.192,0.197 | -0.195 |
| 3653 | XLOC_047705 | chrX | + | 48758466 | 48758882 | 48758466 | 48758578 | 48758795 | 217 | 48758882 | 3653 | 4,6 | 6,3 | 22,19 | 4,0 | 366 | 149 | 0.000900895 | 0.005587765 | 0.213,0.449 | 0.691,1.0 | -0.514 |
| 5810 | XLOC_026855 | chr20 | + | 57564544 | 57565043 | 57564544 | 57564695 | 57564912 | 217 | 57565043 | 5810 | 34,32 | 376,321 | 133,111 | 355,267 | 366 | 149 | 1.11E-16 | 7.47E-15 | 0.036,0.039 | 0.132,0.145 | -0.101 |

| | | | | | | | | | | | | | | | | | | | | | | |
|---|---|---|---|---|---|---|---|---|---|---|---|---|---|---|---|---|---|---|---|---|---|---|
| 6539 | XLOC_004772 | chr10 | + | 13167417 | 13167836 | 13167417 | 13167567 | 13167784 | 217 | 13167836 | 6539 | 7,6 | 1,3 | 5,5 | 0,0 | 366 | 149 | 0.000492122 | 0.003390364 | 0.74,0.449 | 1.0,1.0 | -0.405 |
| 8483 | XLOC_016558 | chr16 | - | 684892 | 685340 | 684892 | 685063 | 685280 | 217 | 685340 | 8483 | 224,232 | 55,54 | 235,110 | 20,19 | 366 | 149 | 0.003752756 | 0.018447676 | 0.624,0.636 | 0.827,0.702 | -0.134 |
| 14379 | XLOC_034781 | chr5 | + | 169017759 | 169018228 | 169017759 | 169017833 | 169018051 | 218 | 169018228 | 14379 | 2,8 | 3,3 | 8,13 | 0,1 | 367 | 149 | 8.38E-05 | 0.000734613 | 0.213,0.52 | 1.0,0.841 | -0.554 |
| 2701 | XLOC_001249 | chr1 | - | 114438378 | 114439087 | 114438378 | 114438660 | 114438879 | 219 | 114439087 | 2701 | 6,9 | 71,68 | 31,13 | 60,37 | 368 | 149 | 5.97E-06 | 7.19E-05 | 0.033,0.051 | 0.173,0.125 | -0.107 |
| 7468 | XLOC_017854 | chr17 | + | 2267599 | 2267957 | 2267599 | 2267670 | 2267889 | 219 | 2267957 | 7468 | 25,9 | 1,7 | 23,15 | 0,0 | 368 | 149 | 7.52E-05 | 0.000667347 | 0.91,0.342 | 1.0,1.0 | -0.374 |
| 2253 | XLOC_014864 | chr15 | - | 43028072 | 43028978 | 43028072 | 43028279 | 43028499 | 220 | 43028978 | 2253 | 4,9 | 18,25 | 20,6 | 11,11 | 369 | 149 | 0.008559342 | 0.036345035 | 0.082,0.127 | 0.423,0.18 | -0.197 |
| 1144 | XLOC_020936 | chr19 | + | 7535011 | 7537466 | 7535011 | 7535201 | 7535422 | 221 | 7537466 | 1144 | 118,110 | 24,22 | 134,101 | 1,6 | 370 | 149 | 1.21E-07 | 2.20E-06 | 0.664,0.668 | 0.982,0.871 | -0.26 |
| 1176 | XLOC_020248 | chr18 | - | 60985314 | 60987270 | 60985314 | 60986185 | 60986406 | 221 | 60987270 | 1176 | 18,11 | 31,23 | 20,13 | 6,10 | 370 | 149 | 0.003008645 | 0.015290067 | 0.19,0.161 | 0.573,0.344 | -0.283 |
| 5260 | XLOC_029569 | chr3 | + | 129179696 | 129180147 | 129179696 | 129179849 | 129180070 | 221 | 129180147 | 5260 | 2,3 | 6,6 | 12,5 | 0,0 | 370 | 149 | 1.78E-10 | 5.26E-09 | 0.118,0.168 | 1.0,1.0 | -0.857 |
| 6175 | XLOC_024344 | chr2 | - | 190647405 | 190647849 | 190647405 | 190647518 | 190647739 | 221 | 190647849 | 6175 | 59,41 | 4,5 | 23,16 | 0,0 | 370 | 149 | 0.000183617 | 0.00144312 | 0.856,0.768 | 1.0,1.0 | -0.188 |
| 14426 | XLOC_007630 | chr11 | - | 72404375 | 72404850 | 72404375 | 72404515 | 72404736 | 221 | 72404850 | 14426 | 29,28 | 66,91 | 81,52 | 83,56 | 370 | 149 | 6.88E-06 | 8.15E-05 | 0.15,0.11 | 0.282,0.272 | -0.147 |
| 13231 | XLOC_016538 | chr16 | + | 89761338 | 89761787 | 89761338 | 89761478 | 89761700 | 222 | 89761787 | 13231 | 27,26 | 220,201 | 92,60 | 115,77 | 371 | 149 | 0 | 0 | 0.047,0.049 | 0.243,0.238 | -0.193 |
| 13244 | XLOC_046445 | chr9 | - | 132580456 | 132580926 | 132580456 | 132580576 | 132580798 | 222 | 132580926 | 13244 | 18,16 | 3,3 | 21,11 | 0,0 | 371 | 149 | 0.000103085 | 0.000879679 | 0.707,0.682 | 1.0,1.0 | -0.305 |
| 967 | XLOC_023405 | chr2 | + | 27454847 | 27455562 | 27454847 | 27455092 | 27455315 | 223 | 27455562 | 967 | 36,36 | 116,127 | 113,82 | 95,103 | 372 | 149 | 1.54E-09 | 3.91E-08 | 0.111,0.102 | 0.323,0.242 | -0.176 |
| 2252 | XLOC_014864 | chr15 | - | 43028072 | 43028978 | 43028072 | 43028276 | 43028499 | 223 | 43028978 | 2252 | 4,9 | 17,29 | 20,8 | 10,1 | 372 | 149 | 5.21E-05 | 0.00048835 | 0.086,0.111 | 0.445,0.762 | -0.505 |
| 808 | XLOC_046147 | chr9 | + | 139887091 | 139887426 | 139887091 | 139887152 | 139887376 | 224 | 139887426 | 808 | 337,310 | 35,44 | 293,245 | 21,8 | 373 | 149 | 0.000468138 | 0.003258506 | 0.794,0.738 | 0.848,0.924 | -0.12 |
| 1394 | XLOC_017920 | chr17 | + | 8192820 | 8193973 | 8192820 | 8192906 | 8193130 | 224 | 8193973 | 1394 | 273,218 | 128,127 | 225,235 | 42,26 | 373 | 149 | 1.05E-10 | 3.21E-09 | 0.46,0.407 | 0.682,0.783 | -0.299 |
| 5801 | XLOC_026873 | chr20 | + | 61910250 | 61910658 | 61910250 | 61910347 | 61910571 | 224 | 61910658 | 5801 | 0,4 | 2,2 | 1,3 | 0,1 | 373 | 149 | 0.00311182 | 0.015735136 | 0.0,0.444 | 1.0,0.545 | -0.55 |
| 11650 | XLOC_005023 | chr10 | + | 103921013 | 103921419 | 103921013 | 103921088 | 103921312 | 224 | 103921419 | 11650 | 19,8 | 36,46 | 19,12 | 8,13 | 373 | 149 | 0.000748632 | 0.004779805 | 0.174,0.065 | 0.487,0.269 | -0.259 |
| 12511 | XLOC_000638 | chr1 | + | 183496760 | 183497162 | 183496760 | 183496866 | 183497090 | 224 | 183497162 | 12511 | 1,0 | 10,3 | 3,7 | 11,3 | 373 | 149 | 0.000688206 | 0.004491463 | 0.038,0.0 | 0.098,0.482 | -0.271 |
| 14214 | XLOC_020938 | chr19 | + | 7599778 | 7600088 | 7599778 | 7599788 | 7600012 | 224 | 7600088 | 14214 | 0,2 | 3,0 | 5,1 | 0,0 | 373 | 149 | 0.001126468 | 0.00674973 | 0.0,1.0 | 1.0,1.0 | -0.5 |
| 3995 | XLOC_044442 | chr8 | - | 144873558 | 144873854 | 144873558 | 144873610 | 144873835 | 225 | 144873854 | 3995 | 80,88 | 8,8 | 229,149 | 3,3 | 374 | 149 | 6.95E-05 | 0.000622857 | 0.799,0.814 | 0.968,0.952 | -0.153 |
| 9019 | XLOC_007137 | chr11 | + | 64135603 | 64136069 | 64135603 | 64135732 | 64135957 | 225 | 64136069 | 9019 | 18,8 | 5,5 | 38,28 | 0,1 | 374 | 149 | 9.09E-07 | 1.35E-05 | 0.589,0.389 | 1.0,0.918 | -0.47 |
| 768 | XLOC_026983 | chr20 | - | 34326889 | 34328809 | 34326889 | 34328519 | 34328745 | 226 | 34328809 | 768 | 204,242 | 367,316 | 557,375 | 395,316 | 375 | 149 | 5.94E-08 | 1.14E-06 | 0.181,0.233 | 0.359,0.32 | -0.133 |
| 1299 | XLOC_018586 | chr17 | - | 43002084 | 43003867 | 43002084 | 43003554 | 43003781 | 227 | 43003867 | 1299 | 322,372 | 19,20 | 474,379 | 0,2 | 376 | 149 | 4.54E-12 | 1.71E-10 | 0.87,0.881 | 1.0,0.987 | -0.118 |
| 10181 | XLOC_029301 | chr3 | + | 9798450 | 9799122 | 9798450 | 9798500 | 9798727 | 227 | 9799122 | 10181 | 5,15 | 1,9 | 8,4 | 1,0 | 376 | 149 | 0.000831289 | 0.005220227 | 0.665,0.398 | 0.76,1.0 | -0.348 |
| 10961 | XLOC_041487 | chr7 | - | 44001193 | 44001689 | 44001193 | 44001344 | 44001571 | 227 | 44001689 | 10961 | 1,0 | 0,3 | 1,2 | 0,0 | 376 | 149 | 0.000126404 | 0.001046424 | 1.0,0.0 | 1.0,1.0 | -0.5 |
| 839 | XLOC_007373 | chr11 | - | 770992 | 771426 | 770992 | 771104 | 771332 | 228 | 771426 | 839 | 64,60 | 56,58 | 151,75 | 46,43 | 377 | 149 | 0.001840751 | 0.010174922 | 0.311,0.29 | 0.565,0.408 | -0.186 |
| 9389 | XLOC_021381 | chr19 | + | 56170227 | 56170711 | 56170227 | 56170347 | 56170575 | 228 | 56170711 | 9389 | 13,5 | 3,4 | 40,35 | 0,4 | 377 | 149 | 0.003959114 | 0.019247699 | 0.631,0.331 | 1.0,0.776 | -0.407 |
| 9741 | XLOC_013303 | chr14 | - | 21905059 | 21905494 | 21905059 | 21905124 | 21905352 | 228 | 21905494 | 9741 | 4,5 | 6,10 | 21,14 | 4,4 | 377 | 149 | 6.69E-05 | 0.00060237 | 0.209,0.165 | 0.675,0.58 | -0.44 |
| 14785 | XLOC_001533 | chr1 | - | 224420596 | 224420991 | 224420596 | 224420674 | 224420902 | 228 | 224420991 | 14785 | 2,1 | 0,8 | 2,3 | 0,1 | 377 | 149 | 0.005635573 | 0.025682865 | 1.0,0.047 | 1.0,0.542 | -0.248 |
| 23 | XLOC_047880 | chrX | + | 153641543 | 153641904 | 153641543 | 153641589 | 153641818 | 229 | 153641904 | 23 | 34,38 | 98,73 | 59,38 | 63,47 | 378 | 149 | 0.000852245 | 0.005330495 | 0.12,0.17 | 0.27,0.242 | -0.111 |
| 2896 | XLOC_014590 | chr15 | + | 42128672 | 42129138 | 42128672 | 42128749 | 42128978 | 229 | 42129138 | 2896 | 10,29 | 16,20 | 38,14 | 11,6 | 378 | 149 | 0.008296752 | 0.035443141 | 0.198,0.364 | 0.577,0.479 | -0.247 |
| 5550 | XLOC_028506 | chr22 | + | 41623522 | 41623873 | 41623522 | 41623578 | 41623807 | 229 | 41623873 | 5550 | 1,1 | 3,0 | 3,3 | 0,0 | 378 | 149 | 0.001371777 | 0.007963422 | 0.116,1.0 | 1.0,1.0 | -0.442 |
| 2254 | XLOC_038073 | chr6 | - | 485008 | 486764 | 485008 | 486403 | 486633 | 230 | 486764 | 2254 | 219,199 | 16,11 | 267,251 | 6,1 | 379 | 149 | 2.76E-05 | 0.000278352 | 0.843,0.877 | 0.946,0.99 | -0.108 |
| 7546 | XLOC_018791 | chr17 | - | 80047125 | 80047602 | 80047125 | 80047277 | 80047507 | 230 | 80047602 | 7546 | 14,8 | 0,7 | 14,4 | 1,0 | 379 | 149 | 0.009916658 | 0.041095353 | 1.0,0.31 | 0.846,1.0 | -0.268 |
| 3939 | XLOC_045902 | chr9 | + | 35800018 | 35800480 | 35800018 | 35800154 | 35800385 | 231 | 35800480 | 3939 | 0,1 | 6,4 | 7,8 | 17,5 | 380 | 149 | 0.000533002 | 0.003633004 | 0.0,0.089 | 0.139,0.386 | -0.218 |
| 10692 | XLOC_009568 | chr12 | - | 118589731 | 118590214 | 118589731 | 118589800 | 118590031 | 231 | 118590214 | 10692 | 0,0 | 1,2 | 2,1 | 0,0 | 380 | 149 | 3.00E-09 | 7.12E-08 | 0.0,0.0 | 1.0,1.0 | -1 |
| 261 | XLOC_038373 | chr6 | - | 99845590 | 99849506 | 99845590 | 99848187 | 99848419 | 232 | 99849506 | 261 | 364,303 | 141,143 | 328,249 | 86,47 | 381 | 149 | 9.62E-05 | 0.000829289 | 0.502,0.453 | 0.599,0.674 | -0.159 |
| 2745 | XLOC_000004 | chr1 | + | 897008 | 898297 | 897008 | 897851 | 898083 | 232 | 898297 | 2745 | 18,11 | 20,27 | 42,27 | 18,12 | 381 | 149 | 0.000110784 | 0.000936325 | 0.26,0.137 | 0.477,0.468 | -0.274 |
| 6160 | XLOC_013166 | chr14 | + | 68038578 | 68038994 | 68038578 | 68038619 | 68038851 | 232 | 68038994 | 6160 | 20,4 | 71,61 | 87,63 | 184,92 | 381 | 149 | 3.93E-06 | 4.96E-05 | 0.099,0.025 | 0.156,0.211 | -0.121 |
| 5324 | XLOC_029402 | chr3 | + | 45043029 | 45043506 | 45043029 | 45043100 | 45043333 | 233 | 45043506 | 5324 | 8,6 | 4,0 | 24,4 | 0,0 | 382 | 149 | 0.004625028 | 0.021765833 | 0.438,1.0 | 1.0,1.0 | -0.281 |
| 7469 | XLOC_017854 | chr17 | + | 2267873 | 2268351 | 2267873 | 2267957 | 2268190 | 233 | 2268351 | 7469 | 20,8 | 186,156 | 44,39 | 71,80 | 382 | 149 | 3.77E-15 | 2.17E-13 | 0.04,0.02 | 0.195,0.16 | -0.147 |
| 10019 | XLOC_027816 | chr21 | - | 46274791 | 46275171 | 46274791 | 46274891 | 46275124 | 233 | 46275171 | 10019 | 13,16 | 5,1 | 8,6 | 0,1 | 382 | 149 | 0.00560222 | 0.02556401 | 0.504,0.862 | 1.0,0.701 | -0.167 |
| 802 | XLOC_041630 | chr7 | - | 100280212 | 100280853 | 100280212 | 100280400 | 100280634 | 234 | 100280853 | 802 | 210,160 | 71,78 | 364,202 | 49,51 | 383 | 149 | 0.000624186 | 0.0041426 | 0.535,0.444 | 0.743,0.606 | -0.185 |
| 2887 | XLOC_016217 | chr16 | + | 2283011 | 2283991 | 2283011 | 2283648 | 2283882 | 234 | 2283991 | 2887 | 14,32 | 72,69 | 46,43 | 86,44 | 383 | 149 | 0.002591873 | 0.013452784 | 0.07,0.153 | 0.172,0.275 | -0.112 |
| 3186 | XLOC_000249 | chr1 | + | 43292581 | 43296786 | 43292581 | 43296204 | 43296438 | 234 | 43296786 | 3186 | 9,6 | 19,11 | 17,15 | 9,2 | 383 | 149 | 0.000931382 | 0.005741557 | 0.156,0.175 | 0.424,0.745 | -0.419 |
| 9595 | XLOC_007211 | chr11 | + | 71940177 | 71940602 | 71940177 | 71940274 | 71940508 | 234 | 71940602 | 9595 | 87,76 | 747,783 | 266,141 | 366,244 | 383 | 149 | 0 | 0 | 0.043,0.036 | 0.22,0.184 | -0.163 |
| 9755 | XLOC_001328 | chr1 | - | 153517787 | 153518264 | 153517787 | 153517994 | 153518228 | 234 | 153518264 | 9755 | 39,13 | 103,77 | 64,54 | 51,38 | 383 | 149 | 1.20E-11 | 4.27E-10 | 0.128,0.062 | 0.328,0.356 | -0.247 |
| 11612 | XLOC_007252 | chr11 | + | 93474871 | 93475183 | 93474871 | 93474894 | 93475128 | 234 | 93475183 | 11612 | 6,9 | 16,4 | 28,14 | 3,0 | 383 | 149 | 1.95E-05 | 0.000203529 | 0.127,0.467 | 0.784,1.0 | -0.595 |
| 12468 | XLOC_000519 | chr1 | + | 151534193 | 151534641 | 151534193 | 151534332 | 151534566 | 234 | 151534641 | 12468 | 4,2 | 9,3 | 1,1 | 0,0 | 383 | 149 | 5.34E-07 | 8.43E-06 | 0.147,0.206 | 1.0,1.0 | -0.824 |
| 1306 | XLOC_024153 | chr2 | - | 98274889 | 98275466 | 98274889 | 98275106 | 98275341 | 235 | 98275466 | 1306 | 70,93 | 112,118 | 101,54 | 57,43 | 384 | 149 | 0.000108106 | 0.000916438 | 0.195,0.234 | 0.407,0.328 | -0.153 |
| 3129 | XLOC_024455 | chr2 | - | 239146842 | 239148209 | 239146842 | 239147892 | 239148127 | 235 | 239148209 | 3129 | 0,6 | 4,12 | 5,2 | 0,0 | 384 | 149 | 1.27E-08 | 2.67E-07 | 0.0,0.162 | 1.0,1.0 | -0.919 |
| 8714 | XLOC_016213 | chr16 | + | 2134566 | 2135027 | 2134566 | 2134716 | 2134951 | 235 | 2135027 | 8714 | 2,3 | 117,107 | 26,17 | 68,34 | 384 | 149 | 1.12E-14 | 6.16E-13 | 0.007,0.011 | 0.129,0.162 | -0.137 |
| 9355 | XLOC_018788 | chr17 | - | 79994057 | 79994519 | 79994057 | 79994184 | 79994419 | 235 | 79994519 | 9355 | 67,87 | 4,4 | 102,82 | 1,0 | 384 | 149 | 0.001001449 | 0.00610685 | 0.867,0.894 | 0.975,1.0 | -0.107 |
| 9925 | XLOC_000550 | chr1 | + | 155582208 | 155582766 | 155582208 | 155582361 | 155582596 | 235 | 155582766 | 9925 | 5,10 | 4,2 | 6,6 | 0,0 | 384 | 149 | 1.98E-06 | 2.69E-05 | 0.327,0.66 | 1.0,1.0 | -0.506 |
| 13263 | XLOC_018585 | chr17 | - | 42931273 | 42931724 | 42931273 | 42931401 | 42931636 | 235 | 42931724 | 13263 | 25,24 | 5,3 | 98,81 | 1,1 | 384 | 149 | 4.83E-05 | 0.000457123 | 0.66,0.756 | 0.974,0.969 | -0.264 |
| 2353 | XLOC_035150 | chr5 | - | 175958159 | 175959517 | 175958159 | 175959130 | 175959366 | 236 | 175959517 | 2353 | 42,48 | 38,27 | 264,217 | 70,45 | 385 | 149 | 3.04E-06 | 3.95E-05 | 0.3,0.408 | 0.593,0.651 | -0.268 |
| 523 | XLOC_044238 | chr8 | - | 21963963 | 21965857 | 21963963 | 21965406 | 21965643 | 237 | 21965857 | 523 | 4,8 | 9,20 | 4,5 | 0,0 | 386 | 149 | 2.24E-08 | 4.58E-07 | 0.146,0.134 | 1.0,1.0 | -0.86 |
| 5944 | XLOC_041630 | chr7 | - | 100280212 | 100280853 | 100280212 | 100280397 | 100280634 | 237 | 100280853 | 5944 | 130,107 | 16,8 | 247,129 | 4,2 | 386 | 149 | 5.14E-06 | 6.30E-05 | 0.758,0.838 | 0.96,0.961 | -0.162 |
| 6244 | XLOC_014964 | chr15 | - | 68503600 | 68504079 | 68503600 | 68503656 | 68503893 | 237 | 68504079 | 6244 | 31,23 | 15,12 | 69,56 | 9,3 | 386 | 149 | 1.70E-05 | 0.000179585 | 0.444,0.425 | 0.747,0.878 | -0.378 |
| 7720 | XLOC_018577 | chr17 | - | 42297833 | 42298302 | 42297833 | 42297870 | 42298107 | 237 | 42298302 | 7720 | 15,20 | 0,10 | 15,6 | 0,0 | 386 | 149 | 0.000434897 | 0.003046655 | 1.0,0.436 | 1.0,1.0 | -0.282 |
| 11149 | XLOC_017867 | chr17 | + | 4621210 | 4621640 | 4621210 | 4621318 | 4621555 | 237 | 4621640 | 11149 | 62,58 | 144,122 | 129,70 | 43,25 | 386 | 149 | 0 | 0 | 0.143,0.155 | 0.537,0.519 | -0.379 |
| 13160 | XLOC_016217 | chr16 | + | 2283438 | 2283991 | 2283438 | 2283648 | 2283885 | 237 | 2283991 | 13160 | 10,28 | 14,29 | 31,40 | 8,16 | 386 | 149 | 0.000319088 | 0.002341028 | 0.216,0.272 | 0.599,0.491 | -0.301 |
| 13576 | XLOC_047966 | chrX | - | 53578957 | 53579402 | 53578957 | 53579034 | 53579271 | 237 | 53579402 | 13576 | 5,10 | 6,5 | 13,10 | 0,0 | 386 | 149 | 9.89E-09 | 2.13E-07 | 0.243,0.436 | 1.0,1.0 | -0.66 |

| | | | | | | | | | | | | | | | | | | | | | | |
|---|---|---|---|---|---|---|---|---|---|---|---|---|---|---|---|---|---|---|---|---|---|---|
| 5529 | XLOC_028549 | chr22 | + | 50635877 | 50636316 | 50635877 | 50636017 | 50636255 | 238 | 50636316 | 5529 | 24,21 | 23,20 | 55,37 | 26,8 | 387 | 149 | 0.004947014 | 0.023065924 | 0.287,0.288 | 0.449,0.64 | -0.257 |
| 6444 | XLOC_023576 | chr2 | + | 96688421 | 96689121 | 96688421 | 96688501 | 96688739 | 238 | 96689121 | 6444 | 1,6 | 0,4 | 1,1 | 0,0 | 387 | 149 | 0.003047363 | 0.015464572 | 1.0,0.366 | 1.0,1.0 | -0.317 |
| 2127 | XLOC_007597 | chr11 | - | 66133596 | 66134035 | 66133596 | 66133662 | 66133901 | 239 | 66134035 | 2127 | 127,118 | 29,31 | 160,125 | 19,17 | 388 | 149 | 0.006301171 | 0.02820648 | 0.627,0.594 | 0.764,0.738 | -0.14 |
| 4916 | XLOC_032498 | chr4 | + | 175237320 | 175237718 | 175237320 | 175237441 | 175237680 | 239 | 175237718 | 4916 | 41,35 | 32,40 | 62,42 | 35,14 | 388 | 149 | 0.012012687 | 0.048348581 | 0.33,0.252 | 0.405,0.535 | -0.179 |
| 5075 | XLOC_030002 | chr3 | - | 123674892 | 123675209 | 123674892 | 123674962 | 123675201 | 239 | 123675209 | 5075 | 135,163 | 58,71 | 89,92 | 12,23 | 388 | 149 | 0.001696922 | 0.00952529 | 0.472,0.469 | 0.74,0.606 | -0.203 |
| 12798 | XLOC_005246 | chr10 | - | 70276507 | 70276939 | 70276507 | 70276600 | 70276840 | 240 | 70276939 | 12798 | 5,6 | 136,104 | 20,24 | 53,37 | 389 | 149 | 8.41E-13 | 3.63E-11 | 0.014,0.022 | 0.126,0.199 | -0.145 |
| 427 | XLOC_034597 | chr5 | + | 78908274 | 78908898 | 78908274 | 78908592 | 78908833 | 241 | 78908898 | 427 | 157,210 | 21,15 | 321,194 | 11,4 | 390 | 149 | 0.000120143 | 0.001005204 | 0.741,0.842 | 0.918,0.949 | -0.142 |
| 4125 | XLOC_043986 | chr8 | + | 28906470 | 28906818 | 28906470 | 28906565 | 28906806 | 241 | 28906818 | 4125 | 1,7 | 24,34 | 44,30 | 17,9 | 390 | 149 | 3.33E-16 | 2.11E-14 | 0.016,0.073 | 0.497,0.56 | -0.484 |
| 10335 | XLOC_011260 | chr13 | + | 20244979 | 20245429 | 20244979 | 20245104 | 20245345 | 241 | 20245429 | 10335 | 16,19 | 4,2 | 14,11 | 0,0 | 390 | 149 | 4.40E-05 | 0.000422254 | 0.604,0.784 | 1.0,1.0 | -0.306 |
| 11015 | XLOC_009165 | chr12 | + | 123465366 | 123465816 | 123465366 | 123465434 | 123465675 | 241 | 123465816 | 11015 | 118,124 | 26,24 | 126,116 | 12,4 | 390 | 149 | 0.000199405 | 0.00154651 | 0.634,0.664 | 0.8,0.917 | -0.21 |
| 1784 | XLOC_009615 | chr12 | - | 132626042 | 132626501 | 132626042 | 132626151 | 132626394 | 243 | 132626501 | 1784 | 37,24 | 33,35 | 116,67 | 40,23 | 392 | 149 | 3.56E-06 | 4.52E-05 | 0.299,0.207 | 0.524,0.525 | -0.271 |
| 5106 | XLOC_029867 | chr3 | - | 51993618 | 51994039 | 51993618 | 51993667 | 51993910 | 243 | 51994039 | 5106 | 5,2 | 39,56 | 11,7 | 19,12 | 392 | 149 | 9.73E-06 | 0.000111386 | 0.046,0.013 | 0.18,0.181 | -0.151 |
| 12841 | XLOC_029295 | chr3 | + | 5236470 | 5236966 | 5236470 | 5236650 | 5236893 | 243 | 5236966 | 12841 | 1,2 | 3,5 | 10,3 | 1,1 | 392 | 149 | 0.000137677 | 0.001124023 | 0.112,0.132 | 0.792,0.533 | -0.541 |
| 13518 | XLOC_000575 | chr1 | + | 161173226 | 161173627 | 161173226 | 161173333 | 161173576 | 243 | 161173627 | 13518 | 0,3 | 1,1 | 3,2 | 1,0 | 392 | 149 | 0.001713968 | 0.009605825 | 0.0,0.533 | 0.533,1.0 | -0.5 |
| 4014 | XLOC_044427 | chr8 | - | 141762001 | 141762415 | 141762001 | 141762086 | 141762330 | 244 | 141762415 | 4014 | 0,2 | 2,1 | 3,4 | 1,0 | 393 | 149 | 0.000131178 | 0.00108025 | 0.0,0.431 | 0.532,1.0 | -0.55 |
| 6201 | XLOC_024255 | chr2 | - | 160253578 | 160253900 | 160253578 | 160253611 | 160253855 | 244 | 160253900 | 6201 | 1,5 | 7,11 | 7,5 | 1,0 | 393 | 149 | 4.86E-07 | 7.75E-06 | 0.051,0.147 | 0.726,1.0 | -0.764 |
| 7255 | XLOC_007078 | chr11 | + | 47600510 | 47600886 | 47600510 | 47600576 | 47600820 | 244 | 47600886 | 7255 | 3,7 | 0,4 | 6,9 | 0,0 | 393 | 149 | 0.003715289 | 0.018308103 | 1.0,0.399 | 1.0,1.0 | -0.3 |
| 8301 | XLOC_035183 | chr5 | - | 180664224 | 180664719 | 180664224 | 180664364 | 180664608 | 244 | 180664719 | 8301 | 70,60 | 2,4 | 336,287 | 0,0 | 393 | 149 | 0.001741297 | 0.009727983 | 0.93,0.85 | 1.0,1.0 | -0.11 |
| 880 | XLOC_044459 | chr8 | - | 145581287 | 145582195 | 145581287 | 145581446 | 145581691 | 245 | 145582195 | 880 | 175,165 | 12,14 | 246,128 | 5,0 | 394 | 149 | 1.26E-05 | 0.000139694 | 0.847,0.817 | 0.949,1.0 | -0.142 |
| 2248 | XLOC_018052 | chr17 | + | 38349614 | 38351065 | 38349614 | 38349669 | 38349915 | 246 | 38351065 | 2248 | 0,0 | 8,6 | 4,1 | 0,2 | 395 | 149 | 0.003857136 | 0.018842729 | 0.0,0.0 | 1.0,0.159 | -0.58 |
| 2972 | XLOC_005350 | chr10 | - | 102249459 | 102250640 | 102249459 | 102249518 | 102249764 | 246 | 102250640 | 2972 | 4,4 | 10,3 | 22,14 | 4,8 | 395 | 149 | 0.009060478 | 0.03802708 | 0.131,0.335 | 0.675,0.398 | -0.303 |
| 3591 | XLOC_047948 | chrX | - | 48752320 | 48752737 | 48752320 | 48752384 | 48752630 | 246 | 48752737 | 3591 | 0,3 | 2,3 | 9,7 | 0,1 | 395 | 149 | 5.24E-06 | 6.40E-05 | 0.0,0.274 | 1.0,0.725 | -0.726 |
| 5578 | XLOC_041629 | chr7 | - | 100175444 | 100175873 | 100175444 | 100175535 | 100175781 | 246 | 100175873 | 5578 | 29,29 | 99,100 | 32,34 | 40,37 | 395 | 149 | 1.12E-05 | 0.000125187 | 0.1,0.099 | 0.232,0.257 | -0.145 |
| 7564 | XLOC_013249 | chr14 | + | 96991646 | 96992096 | 96991646 | 96991728 | 96991974 | 246 | 96992096 | 7564 | 0,1 | 2,1 | 6,6 | 0,2 | 395 | 149 | 4.97E-05 | 0.000468403 | 0.0,0.274 | 1.0,0.531 | -0.629 |
| 69 | XLOC_046238 | chr9 | - | 35662942 | 35663389 | 35662942 | 35663071 | 35663318 | 247 | 35663389 | 69 | 92,126 | 103,112 | 61,45 | 26,28 | 396 | 149 | 0.004055935 | 0.019596977 | 0.252,0.297 | 0.469,0.377 | -0.149 |
| 4122 | XLOC_043988 | chr8 | + | 29961877 | 29962371 | 29961877 | 29962002 | 29962249 | 247 | 29962371 | 4122 | 2,0 | 1,3 | 6,2 | 0,0 | 396 | 149 | 8.66E-08 | 1.60E-06 | 0.429,0.0 | 1.0,1.0 | -0.785 |
| 4830 | XLOC_038429 | chr6 | - | 131201283 | 131201629 | 131201283 | 131201346 | 131201593 | 247 | 131201629 | 4830 | 16,18 | 4,1 | 17,10 | 0,0 | 396 | 149 | 0.000307263 | 0.002263679 | 0.601,0.871 | 1.0,1.0 | -0.264 |
| 10685 | XLOC_018789 | chr17 | - | 80008537 | 80009218 | 80008537 | 80008640 | 80008887 | 247 | 80009218 | 10685 | 30,12 | 2,3 | 21,21 | 0,0 | 396 | 149 | 0.000483188 | 0.003338586 | 0.849,0.601 | 1.0,1.0 | -0.275 |
| 10231 | XLOC_028694 | chr22 | - | 41243973 | 41244373 | 41243973 | 41244048 | 41244297 | 249 | 41244373 | 10231 | 0,0 | 10,12 | 3,1 | 1,0 | 398 | 149 | 5.85E-05 | 0.000538598 | 0.0,0.0 | 0.529,1.0 | -0.764 |
| 908 | XLOC_017917 | chr17 | + | 7847440 | 7847956 | 7847440 | 7847543 | 7847793 | 250 | 7847956 | 908 | 96,134 | 8,1 | 79,64 | 0,0 | 399 | 149 | 0.000704669 | 0.004567254 | 0.818,0.98 | 1.0,1.0 | -0.101 |
| 11174 | XLOC_008874 | chr12 | + | 50036362 | 50036799 | 50036362 | 50036458 | 50036709 | 251 | 50036799 | 11174 | 14,5 | 2,6 | 4,1 | 0,0 | 400 | 149 | 2.29E-06 | 3.06E-05 | 0.723,0.237 | 1.0,1.0 | -0.52 |
| 12367 | XLOC_018568 | chr17 | - | 41622342 | 41622738 | 41622342 | 41622390 | 41622641 | 251 | 41622738 | 12367 | 8,6 | 72,63 | 25,9 | 25,25 | 400 | 149 | 1.68E-06 | 2.31E-05 | 0.04,0.034 | 0.271,0.118 | -0.158 |
| 14147 | XLOC_000684 | chr1 | + | 204091028 | 204091527 | 204091028 | 204091113 | 204091364 | 251 | 204091527 | 14147 | 23,18 | 66,110 | 85,48 | 38,26 | 400 | 149 | 0 | 0 | 0.115,0.057 | 0.455,0.407 | -0.345 |
| 3240 | XLOC_001334 | chr1 | - | 153915031 | 153915606 | 153915031 | 153915101 | 153915353 | 252 | 153915606 | 3240 | 52,43 | 82,70 | 234,160 | 66,41 | 401 | 149 | 0 | 0 | 0.191,0.186 | 0.568,0.592 | -0.391 |
| 3609 | XLOC_047885 | chrX | + | 153692770 | 153693211 | 153692770 | 153692844 | 153693096 | 252 | 153693211 | 3609 | 12,9 | 31,26 | 31,21 | 20,14 | 401 | 149 | 1.91E-05 | 0.00019981 | 0.126,0.114 | 0.365,0.358 | -0.241 |
| 14708 | XLOC_013186 | chr14 | + | 73730964 | 73731434 | 73730964 | 73731043 | 73731295 | 252 | 73731434 | 14708 | 4,7 | 28,34 | 13,4 | 9,6 | 401 | 149 | 0.000349196 | 0.002523898 | 0.05,0.071 | 0.349,0.199 | -0.214 |
| 1583 | XLOC_045894 | chr9 | + | 34660500 | 34660933 | 34660500 | 34660597 | 34660850 | 253 | 34660933 | 1583 | 13,11 | 7,1 | 20,7 | 1,0 | 402 | 149 | 0.000559409 | 0.003781966 | 0.408,0.803 | 0.881,1.0 | -0.335 |
| 2722 | XLOC_016200 | chr16 | + | 1556484 | 1557371 | 1556484 | 1557052 | 1557306 | 254 | 1557371 | 2722 | 25,28 | 94,90 | 37,31 | 75,35 | 403 | 149 | 0.001825003 | 0.010119559 | 0.09,0.103 | 0.154,0.247 | -0.104 |
| 2885 | XLOC_016217 | chr16 | + | 2283011 | 2283648 | 2283011 | 2283184 | 2283438 | 254 | 2283648 | 2885 | 20,34 | 87,99 | 68,63 | 120,79 | 403 | 149 | 7.10E-05 | 0.000634647 | 0.078,0.113 | 0.173,0.228 | -0.105 |
| 9186 | XLOC_014806 | chr15 | - | 23191453 | 23191942 | 23191453 | 23191579 | 23191833 | 254 | 23191942 | 9186 | 14,10 | 45,27 | 11,20 | 16,19 | 403 | 149 | 0.006647784 | 0.029477754 | 0.103,0.12 | 0.203,0.28 | -0.13 |
| 9547 | XLOC_001344 | chr1 | - | 154933949 | 154937052 | 154933949 | 154936407 | 154936661 | 254 | 154937052 | 9547 | 23,18 | 3,1 | 20,15 | 0,0 | 403 | 149 | 0.006378399 | 0.028479967 | 0.739,0.869 | 1.0,1.0 | -0.196 |
| 11175 | XLOC_008874 | chr12 | + | 50036362 | 50036799 | 50036362 | 50036458 | 50036712 | 254 | 50036799 | 11175 | 8,5 | 1,3 | 3,1 | 1,0 | 403 | 149 | 0.011200487 | 0.045493571 | 0.747,0.381 | 0.526,1.0 | -0.199 |
| 11499 | XLOC_000830 | chr1 | - | 1333611 | 1334051 | 1333611 | 1333722 | 1333976 | 254 | 1334051 | 11499 | 9,10 | 228,205 | 43,29 | 97,94 | 403 | 149 | 2.22E-16 | 1.46E-14 | 0.014,0.018 | 0.141,0.102 | -0.105 |
| 14902 | XLOC_021131 | chr19 | + | 35230052 | 35230514 | 35230052 | 35230173 | 35230427 | 254 | 35230514 | 14902 | 6,3 | 19,14 | 11,8 | 3,7 | 403 | 149 | 0.000827978 | 0.005207867 | 0.105,0.073 | 0.575,0.297 | -0.347 |
| 2524 | XLOC_007071 | chr11 | + | 47182677 | 47184358 | 47182677 | 47182741 | 47182996 | 255 | 47184358 | 2524 | 33,58 | 8,8 | 37,34 | 0,3 | 404 | 149 | 0.004366377 | 0.020742129 | 0.603,0.728 | 1.0,0.807 | -0.238 |
| 6317 | XLOC_023980 | chr2 | - | 27589951 | 27590426 | 27589951 | 27590068 | 27590323 | 255 | 27590426 | 6317 | 43,40 | 106,117 | 75,77 | 93,66 | 404 | 149 | 1.82E-06 | 2.50E-05 | 0.13,0.112 | 0.229,0.301 | -0.144 |
| 10474 | XLOC_020931 | chr19 | + | 6746188 | 6746572 | 6746188 | 6746207 | 6746462 | 255 | 6746572 | 10474 | 21,20 | 49,49 | 10,17 | 2,5 | 404 | 149 | 4.25E-07 | 6.85E-06 | 0.136,0.131 | 0.648,0.556 | -0.469 |
| 836 | XLOC_004889 | chr10 | + | 75560397 | 75560953 | 75560397 | 75560517 | 75560774 | 257 | 75560953 | 836 | 17,36 | 81,44 | 42,40 | 29,30 | 406 | 149 | 8.97E-05 | 0.000780238 | 0.072,0.231 | 0.347,0.329 | -0.186 |
| 1131 | XLOC_021130 | chr19 | + | 35173681 | 35174143 | 35173681 | 35173802 | 35174059 | 257 | 35174143 | 1131 | 54,75 | 62,53 | 74,52 | 23,19 | 406 | 149 | 8.08E-05 | 0.000711711 | 0.242,0.342 | 0.541,0.501 | -0.229 |
| 1197 | XLOC_000264 | chr1 | + | 44436267 | 44436861 | 44436267 | 44436380 | 44436637 | 257 | 44436861 | 1197 | 48,62 | 55,69 | 400,330 | 126,85 | 406 | 149 | 3.71E-14 | 1.86E-12 | 0.243,0.248 | 0.538,0.588 | -0.317 |
| 1427 | XLOC_037839 | chr6 | + | 43405671 | 43406331 | 43405671 | 43405751 | 43406008 | 257 | 43406331 | 1427 | 3,5 | 4,3 | 37,14 | 2,3 | 406 | 149 | 0.00146791 | 0.008413673 | 0.216,0.38 | 0.872,0.631 | -0.454 |
| 7913 | XLOC_018780 | chr17 | - | 79803140 | 79803618 | 79803140 | 79803179 | 79803436 | 257 | 79803618 | 7913 | 21,22 | 2,3 | 16,11 | 0,0 | 406 | 149 | 0.00092746 | 0.005722661 | 0.794,0.729 | 1.0,1.0 | -0.238 |
| 8423 | XLOC_016687 | chr16 | - | 29922798 | 29923188 | 29922798 | 29922878 | 29923135 | 257 | 29923188 | 8423 | 7,6 | 2,2 | 4,2 | 0,0 | 406 | 149 | 0.000233013 | 0.001770745 | 0.562,0.524 | 1.0,1.0 | -0.457 |
| 9898 | XLOC_007066 | chr11 | + | 46389201 | 46389633 | 46389201 | 46389297 | 46389554 | 257 | 46389633 | 9898 | 2,2 | 17,13 | 5,12 | 14,12 | 406 | 149 | 0.002546195 | 0.013254668 | 0.041,0.053 | 0.116,0.268 | -0.145 |
| 12088 | XLOC_035160 | chr5 | - | 176918010 | 176918421 | 176918010 | 176918147 | 176918404 | 257 | 176918421 | 12088 | 12,8 | 6,3 | 7,7 | 0,0 | 406 | 149 | 9.98E-07 | 1.47E-05 | 0.423,0.495 | 1.0,1.0 | -0.541 |
| 14903 | XLOC_021131 | chr19 | + | 35230052 | 35230514 | 35230052 | 35230173 | 35230430 | 257 | 35230514 | 14903 | 6,3 | 13,14 | 11,8 | 5,8 | 406 | 149 | 0.004949987 | 0.023072168 | 0.145,0.073 | 0.447,0.268 | -0.249 |
| 1206 | XLOC_021387 | chr19 | + | 56925327 | 56925809 | 56925327 | 56925454 | 56925713 | 259 | 56925809 | 1206 | 1,0 | 16,2 | 2,9 | 6,4 | 408 | 149 | 0.000132609 | 0.00109013 | 0.022,0.0 | 0.109,0.451 | -0.269 |
| 2459 | XLOC_007532 | chr11 | - | 61187941 | 61189080 | 61187941 | 61188602 | 61188861 | 259 | 61189080 | 2459 | 75,75 | 5,2 | 114,96 | 0,0 | 408 | 149 | 0.000707363 | 0.004572122 | 0.846,0.932 | 1.0,1.0 | -0.111 |
| 5534 | XLOC_028548 | chr22 | + | 50354965 | 50355420 | 50354965 | 50355016 | 50355275 | 259 | 50355420 | 5534 | 45,20 | 7,0 | 104,75 | 0,2 | 408 | 149 | 0.003149933 | 0.015910767 | 0.701,1.0 | 1.0,0.932 | -0.115 |
| 488 | XLOC_032542 | chr4 | - | 2243246 | 2243898 | 2243246 | 2243517 | 2243777 | 260 | 2243898 | 488 | 74,72 | 16,20 | 182,101 | 4,3 | 409 | 149 | 1.96E-10 | 5.74E-09 | 0.628,0.567 | 0.943,0.925 | -0.337 |
| 1236 | XLOC_016253 | chr16 | + | 3454425 | 3455927 | 3454425 | 3454648 | 3454908 | 260 | 3455927 | 1236 | 11,18 | 10,6 | 17,15 | 0,3 | 409 | 149 | 0.005025247 | 0.023361281 | 0.286,0.522 | 1.0,0.646 | -0.419 |
| 11921 | XLOC_007319 | chr11 | + | 119002539 | 119003007 | 119002539 | 119002692 | 119002952 | 260 | 119003007 | 11921 | 1,4 | 1,4 | 3,5 | 0,0 | 409 | 149 | 8.14E-07 | 1.22E-05 | 0.267,0.267 | 1.0,1.0 | -0.733 |
| 11392 | XLOC_021828 | chr19 | - | 50364705 | 50365128 | 50364705 | 50364767 | 50365028 | 261 | 50365128 | 11392 | 82,79 | 9,3 | 66,41 | 0,0 | 410 | 149 | 2.91E-05 | 0.00029165 | 0.768,0.905 | 1.0,1.0 | -0.163 |
| 1282 | XLOC_018712 | chr17 | - | 73234062 | 73234843 | 73234062 | 73234470 | 73234732 | 262 | 73234843 | 1282 | 44,37 | 64,71 | 160,142 | 79,64 | 411 | 149 | 1.06E-10 | 3.22E-09 | 0.2,0.159 | 0.423,0.446 | -0.255 |
| 4713 | XLOC_035081 | chr5 | - | 139940669 | 139941034 | 139940669 | 139940686 | 139940949 | 263 | 139941034 | 4713 | 4,4 | 39,42 | 14,8 | 19,11 | 412 | 149 | 4.68E-06 | 5.80E-05 | 0.036,0.033 | 0.21,0.208 | -0.174 |

| | | | | | | | | | | | | | | | | | | | | | | |
|---|---|---|---|---|---|---|---|---|---|---|---|---|---|---|---|---|---|---|---|---|---|---|
| 4373 | XLOC_041076 | chr7 | + | 36464924 | 36465349 | 36464924 | 36465046 | 36465310 | 264 | 36465349 | 4373 | 0,0 | 1,3 | 5,5 | 0,0 | 413 | 149 | 8.13E-11 | 2.51E-09 | 0.0,0.0 | 1.0,1.0 | -1 |
| 1385 | XLOC_017956 | chr17 | + | 18166399 | 18166918 | 18166399 | 18166562 | 18166827 | 265 | 18166918 | 1385 | 12,12 | 3,2 | 60,21 | 2,0 | 414 | 149 | 0.00375711 | 0.018449815 | 0.59,0.683 | 0.915,1.0 | -0.321 |
| 2647 | XLOC_016203 | chr16 | + | 1817149 | 1817736 | 1817149 | 1817298 | 1817564 | 266 | 1817736 | 2647 | 0,4 | 40,22 | 18,20 | 33,20 | 415 | 149 | 3.58E-08 | 7.04E-07 | 0.0,0.061 | 0.164,0.264 | -0.184 |
| 6406 | XLOC_023725 | chr2 | + | 173431583 | 173431987 | 173431583 | 173431661 | 173431927 | 266 | 173431987 | 6406 | 3,2 | 2,5 | 21,5 | 0,0 | 415 | 149 | 2.64E-10 | 7.59E-09 | 0.35,0.126 | 1.0,1.0 | -0.762 |
| 1401 | XLOC_017896 | chr17 | + | 7125985 | 7126556 | 7125985 | 7126184 | 7126451 | 267 | 7126556 | 1401 | 12,621,175 | 15,661,655 | 21,471,458 | 924,700 | 416 | 149 | 0 | 0 | 0.224,0.203 | 0.454,0.427 | -0.227 |
| 1429 | XLOC_018607 | chr17 | - | 45915017 | 45915816 | 45915017 | 45915368 | 45915635 | 267 | 45915816 | 1429 | 54,89 | 133,152 | 113,63 | 27,28 | 416 | 149 | 1.11E-12 | 4.77E-11 | 0.127,0.173 | 0.6,0.446 | -0.373 |
| 1538 | XLOC_016368 | chr16 | + | 30762426 | 30762924 | 30762426 | 30762602 | 30762869 | 267 | 30762924 | 1538 | 137,135 | 65,75 | 83,54 | 24,15 | 416 | 149 | 0.008909619 | 0.037561357 | 0.43,0.392 | 0.553,0.563 | -0.147 |
| 2479 | XLOC_020078 | chr18 | + | 32822354 | 32823257 | 32822354 | 32822848 | 32823115 | 267 | 32823257 | 2479 | 11,2 | 27,19 | 31,12 | 29,17 | 416 | 149 | 0.001023324 | 0.006216087 | 0.127,0.036 | 0.277,0.202 | -0.158 |
| 3160 | XLOC_018763 | chr17 | - | 78188413 | 78188937 | 78188413 | 78188564 | 78188831 | 267 | 78188937 | 3160 | 15,32 | 63,88 | 51,37 | 29,18 | 416 | 149 | 1.81E-12 | 7.50E-11 | 0.079,0.115 | 0.386,0.424 | -0.308 |
| 7474 | XLOC_017854 | chr17 | + | 2275636 | 2276080 | 2275636 | 2275782 | 2276049 | 267 | 2276080 | 7474 | 42,26 | 0,8 | 16,26 | 0,0 | 416 | 149 | 0.000543441 | 0.003695245 | 1.0,0.538 | 1.0,1.0 | -0.231 |
| 6114 | XLOC_016250 | chr16 | + | 3335719 | 3336149 | 3335719 | 3335754 | 3336022 | 268 | 3336149 | 6114 | 23,28 | 52,63 | 78,48 | 43,30 | 417 | 149 | 1.00E-08 | 2.15E-07 | 0.136,0.137 | 0.393,0.364 | -0.242 |
| 7437 | XLOC_000461 | chr1 | + | 145290428 | 145290850 | 145290428 | 145290511 | 145290780 | 269 | 145290850 | 7437 | 66,76 | 12,18 | 134,88 | 9,6 | 418 | 149 | 0.000566015 | 0.003813851 | 0.662,0.601 | 0.841,0.839 | -0.209 |
| 9650 | XLOC_013423 | chr14 | - | 65544061 | 65544754 | 65544061 | 65544361 | 65544630 | 269 | 65544754 | 9650 | 24,11 | 15,12 | 36,38 | 13,10 | 418 | 149 | 0.011998116 | 0.048317475 | 0.363,0.246 | 0.497,0.575 | -0.232 |
| 6516 | XLOC_023405 | chr2 | + | 27463434 | 27463823 | 27463434 | 27463508 | 27463778 | 270 | 27463823 | 6516 | 3,4 | 4,0 | 14,18 | 0,0 | 419 | 149 | 0.003966837 | 0.019267803 | 0.211,1.0 | 1.0,1.0 | -0.394 |
| 11556 | XLOC_007764 | chr11 | - | 129753964 | 129754419 | 129753964 | 129754038 | 129754308 | 270 | 129754419 | 11556 | 2,4 | 0,6 | 14,8 | 0,0 | 419 | 149 | 0.001317537 | 0.007692261 | 1.0,0.192 | 1.0,1.0 | -0.404 |
| 12658 | XLOC_030032 | chr3 | - | 129115882 | 129116378 | 129115882 | 129115983 | 129116253 | 270 | 129116378 | 12658 | 24,17 | 8,6 | 27,19 | 2,2 | 419 | 149 | 0.007986972 | 0.034348355 | 0.516,0.502 | 0.828,0.772 | -0.291 |
| 14227 | XLOC_044016 | chr8 | + | 42022588 | 42023064 | 42022588 | 42022674 | 42022944 | 270 | 42023064 | 14227 | 12,5 | 39,33 | 12,12 | 16,12 | 419 | 149 | 0.001120781 | 0.00672421 | 0.099,0.051 | 0.211,0.262 | -0.161 |
| 4655 | XLOC_047755 | chrX | + | 72899160 | 72899544 | 72899160 | 72899217 | 72899488 | 271 | 72899544 | 4655 | 1,7 | 32,35 | 5,17 | 9,5 | 420 | 149 | 8.48E-05 | 0.000742544 | 0.011,0.066 | 0.165,0.547 | -0.318 |
| 35 | XLOC_047846 | chrX | + | 134480943 | 134483239 | 134480943 | 134482762 | 134483034 | 272 | 134483239 | 35 | 61,65 | 10,11 | 75,40 | 2,0 | 421 | 149 | 5.34E-06 | 6.52E-05 | 0.683,0.677 | 0.93,1.0 | -0.285 |
| 5778 | XLOC_028750 | chr22 | - | 50945210 | 50946121 | 50945210 | 50945464 | 50945736 | 272 | 50946121 | 5778 | 60,52 | 5,8 | 58,22 | 2,0 | 421 | 149 | 0.002125075 | 0.011463877 | 0.809,0.697 | 0.911,1.0 | -0.203 |
| 11383 | XLOC_023965 | chr2 | - | 25043252 | 25043717 | 25043252 | 25043320 | 25043592 | 272 | 25043717 | 11383 | 104,104 | 9,13 | 104,87 | 5,0 | 421 | 149 | 0.001022152 | 0.006211643 | 0.804,0.739 | 0.88,1.0 | -0.168 |
| 2525 | XLOC_007071 | chr11 | + | 47182677 | 47184358 | 47182677 | 47182741 | 47183014 | 273 | 47184358 | 2525 | 36,64 | 72,86 | 43,35 | 22,16 | 422 | 149 | 2.59E-06 | 3.42E-05 | 0.15,0.208 | 0.408,0.436 | -0.243 |
| 3009 | XLOC_000106 | chr1 | + | 17274825 | 17275421 | 17274825 | 17275019 | 17275293 | 274 | 17275421 | 3009 | 2,3 | 10,5 | 9,4 | 0,0 | 423 | 149 | 1.25E-10 | 3.74E-09 | 0.066,0.174 | 1.0,1.0 | -0.88 |
| 986 | XLOC_016192 | chr16 | + | 841850 | 842347 | 841850 | 841948 | 842223 | 275 | 842347 | 986 | 207,206 | 103,71 | 310,163 | 52,27 | 424 | 149 | 1.95E-06 | 2.65E-05 | 0.414,0.505 | 0.677,0.68 | -0.219 |
| 14696 | XLOC_016224 | chr16 | + | 2578712 | 2579096 | 2578712 | 2578719 | 2578994 | 275 | 2579096 | 14696 | 83,81 | 85,67 | 109,68 | 20,20 | 424 | 149 | 1.16E-08 | 2.45E-07 | 0.255,0.298 | 0.657,0.544 | -0.324 |
| 6644 | XLOC_007505 | chr11 | - | 47194976 | 47195433 | 47194976 | 47195033 | 47195309 | 276 | 47195433 | 6644 | 66,84 | 173,174 | 121,93 | 66,44 | 425 | 149 | 0 | 0 | 0.118,0.145 | 0.391,0.426 | -0.277 |
| 13236 | XLOC_000581 | chr1 | + | 161721147 | 161721571 | 161721147 | 161721167 | 161721444 | 277 | 161721571 | 13236 | 7,0 | 4,3 | 10,3 | 2,0 | 426 | 149 | 0.005233433 | 0.024122642 | 0.38,0.0 | 0.636,1.0 | -0.628 |
| 1279 | XLOC_018746 | chr17 | - | 74729864 | 74732546 | 74729864 | 74731957 | 74732235 | 278 | 74732546 | 1279 | 12,531,309 | 221,207 | 18,661,593 | 121,117 | 427 | 149 | 1.06E-11 | 3.79E-10 | 0.664,0.688 | 0.843,0.826 | -0.159 |
| 8776 | XLOC_004892 | chr10 | + | 75863577 | 75864028 | 75863577 | 75863686 | 75863964 | 278 | 75864028 | 8776 | 17,9 | 0,4 | 8,4 | 0,0 | 427 | 149 | 0.00632467 | 0.028284772 | 1.0,0.44 | 1.0,1.0 | -0.28 |
| 8948 | XLOC_014664 | chr15 | + | 63880516 | 63881014 | 63880516 | 63880622 | 63880900 | 278 | 63881014 | 8948 | 43,43 | 129,95 | 168,99 | 59,40 | 427 | 149 | 0 | 0 | 0.104,0.136 | 0.498,0.463 | -0.361 |
| 3938 | XLOC_045966 | chr9 | + | 95874162 | 95874571 | 95874162 | 95874220 | 95874499 | 279 | 95874571 | 3938 | 53,39 | 101,88 | 65,41 | 36,21 | 428 | 149 | 3.09E-09 | 7.31E-08 | 0.154,0.134 | 0.386,0.405 | -0.252 |
| 1322 | XLOC_018401 | chr17 | - | 7982758 | 7983259 | 7982758 | 7982842 | 7983122 | 280 | 7983259 | 1322 | 5,13 | 0,9 | 8,4 | 0,0 | 429 | 149 | 0.000529921 | 0.003618471 | 1.0,0.334 | 1.0,1.0 | -0.333 |
| 13975 | XLOC_000910 | chr1 | - | 17330831 | 17331316 | 17330831 | 17330906 | 17331186 | 280 | 17331316 | 13975 | 0,0 | 10,8 | 2,4 | 13,6 | 429 | 149 | 0.006291274 | 0.028180037 | 0.0,0.0 | 0.051,0.188 | -0.119 |
| 9047 | XLOC_015009 | chr15 | - | 78585180 | 78585614 | 78585180 | 78585227 | 78585508 | 281 | 78585614 | 9047 | 10,7 | 4,4 | 20,11 | 2,0 | 430 | 149 | 0.000981177 | 0.006001372 | 0.464,0.377 | 0.776,1.0 | -0.468 |
| 12674 | XLOC_045913 | chr9 | + | 37492278 | 37492712 | 37492278 | 37492372 | 37492653 | 281 | 37492712 | 12674 | 2,1 | 0,3 | 2,1 | 0,0 | 430 | 149 | 0.004877916 | 0.022796429 | 1.0,0.104 | 1.0,1.0 | -0.448 |
| 2482 | XLOC_007374 | chr11 | - | 802681 | 803587 | 802681 | 802891 | 803173 | 282 | 803587 | 2482 | 38,30 | 10,28 | 33,18 | 4,1 | 431 | 149 | 0.001851449 | 0.01021806 | 0.568,0.27 | 0.74,0.862 | -0.382 |
| 5332 | XLOC_041127 | chr7 | + | 66773563 | 66774126 | 66773563 | 66773639 | 66773921 | 282 | 66774126 | 5332 | 19,22 | 5,2 | 8,7 | 1,0 | 431 | 149 | 0.00478896 | 0.022425224 | 0.568,0.792 | 0.734,1.0 | -0.187 |
| 7988 | XLOC_018028 | chr17 | + | 34958981 | 34959813 | 34958981 | 34959075 | 34959357 | 282 | 34959813 | 7988 | 10,8 | 2,8 | 7,5 | 1,0 | 431 | 149 | 0.00081984 | 0.005162106 | 0.634,0.257 | 0.708,1.0 | -0.408 |
| 1004 | XLOC_000573 | chr1 | + | 161133953 | 161134402 | 161133953 | 161134040 | 161134323 | 283 | 161134402 | 1004 | 135,149 | 110,115 | 205,142 | 85,71 | 432 | 149 | 0.00062011 | 0.004125204 | 0.297,0.309 | 0.454,0.408 | -0.128 |
| 4663 | XLOC_047953 | chrX | - | 48890650 | 48891055 | 48890650 | 48890696 | 48890979 | 283 | 48891055 | 4663 | 3,3 | 5,11 | 6,3 | 1,0 | 432 | 149 | 1.45E-05 | 0.00015729 | 0.171,0.086 | 0.674,1.0 | -0.708 |
| 13974 | XLOC_000910 | chr1 | - | 17330831 | 17331316 | 17330831 | 17330903 | 17331186 | 283 | 17331316 | 13974 | 0,0 | 1,7 | 2,4 | 7,0 | 432 | 149 | 0.004664762 | 0.021923607 | 0.0,0.0 | 0.09,1.0 | -0.545 |
| 2992 | XLOC_013457 | chr14 | - | 74759450 | 74759951 | 74759450 | 74759572 | 74759856 | 284 | 74759951 | 2992 | 37,39 | 36,30 | 60,52 | 11,4 | 433 | 149 | 6.87E-08 | 1.30E-06 | 0.261,0.309 | 0.652,0.817 | -0.449 |
| 97 | XLOC_044466 | chr8 | - | 145741371 | 145742574 | 145741371 | 145742148 | 145742433 | 285 | 145742574 | 97 | 224,202 | 189,158 | 318,274 | 77,58 | 434 | 149 | 1.11E-16 | 7.47E-15 | 0.289,0.305 | 0.586,0.619 | -0.306 |
| 8737 | XLOC_016191 | chr16 | + | 777411 | 778029 | 777411 | 777679 | 777964 | 285 | 778029 | 8737 | 44,41 | 13,14 | 77,48 | 10,1 | 434 | 149 | 0.00141148 | 0.008158443 | 0.537,0.501 | 0.726,0.943 | -0.316 |
| 14824 | XLOC_047699 | chrX | + | 48380267 | 48380640 | 48380267 | 48380295 | 48380580 | 285 | 48380640 | 14824 | 10,10 | 7,4 | 42,22 | 3,1 | 434 | 149 | 0.000112993 | 0.000950447 | 0.329,0.462 | 0.828,0.883 | -0.46 |
| 3159 | XLOC_018763 | chr17 | - | 78187602 | 78188127 | 78187602 | 78187684 | 78187970 | 286 | 78188127 | 3159 | 22,50 | 125,118 | 75,66 | 70,59 | 435 | 149 | 1.45E-10 | 4.32E-09 | 0.057,0.127 | 0.268,0.277 | -0.181 |
| 6055 | XLOC_018159 | chr17 | + | 53120618 | 53121031 | 53120618 | 53120686 | 53120972 | 286 | 53121031 | 6055 | 0,0 | 1,2 | 7,4 | 5,1 | 435 | 149 | 2.38E-05 | 0.000243123 | 0.0,0.0 | 0.324,0.578 | -0.451 |
| 14942 | XLOC_000712 | chr1 | + | 212529967 | 212530386 | 212529967 | 212530018 | 212530304 | 286 | 212530386 | 14942 | 8,8 | 2,5 | 5,7 | 0,0 | 435 | 149 | 2.17E-07 | 3.71E-06 | 0.578,0.354 | 1.0,1.0 | -0.534 |
| 4429 | XLOC_038506 | chr6 | - | 170063936 | 170064382 | 170063936 | 170063974 | 170064261 | 287 | 170064382 | 4429 | 67,60 | 27,28 | 135,86 | 5,12 | 436 | 149 | 1.09E-06 | 1.57E-05 | 0.459,0.423 | 0.902,0.71 | -0.365 |
| 11441 | XLOC_046430 | chr9 | - | 130884640 | 130885413 | 130884640 | 130884925 | 130885212 | 287 | 130885413 | 11441 | 41,29 | 3,2 | 45,30 | 1,0 | 436 | 149 | 0.005627168 | 0.025652836 | 0.824,0.832 | 0.939,1.0 | -0.142 |
| 1015 | XLOC_021778 | chr19 | - | 46274228 | 46274923 | 46274228 | 46274318 | 46274607 | 289 | 46274923 | 1015 | 49,49 | 67,39 | 77,46 | 20,13 | 438 | 149 | 2.03E-07 | 3.49E-06 | 0.199,0.299 | 0.567,0.546 | -0.307 |
| 2938 | XLOC_000133 | chr1 | + | 24973157 | 24975520 | 24973157 | 24973280 | 24973569 | 289 | 24975520 | 2938 | 123,155 | 138,116 | 150,149 | 34,25 | 438 | 149 | 6.11E-15 | 3.44E-13 | 0.233,0.313 | 0.6,0.67 | -0.362 |
| 8300 | XLOC_035183 | chr5 | - | 180664224 | 180664719 | 180664224 | 180664319 | 180664608 | 289 | 180664719 | 8300 | 70,60 | 31,30 | 336,287 | 64,47 | 438 | 149 | 5.36E-06 | 6.54E-05 | 0.434,0.405 | 0.641,0.675 | -0.239 |
| 10053 | XLOC_021199 | chr19 | + | 39218585 | 39219021 | 39218585 | 39218666 | 39218955 | 289 | 39219021 | 10053 | 9,5 | 2,2 | 8,3 | 0,0 | 438 | 149 | 0.000241719 | 0.001828495 | 0.605,0.46 | 1.0,1.0 | -0.468 |
| 12981 | XLOC_021262 | chr19 | + | 45582528 | 45583287 | 45582528 | 45582875 | 45583164 | 289 | 45583287 | 12981 | 15,23 | 4,0 | 30,27 | 0,0 | 438 | 149 | 0.009003762 | 0.037859898 | 0.561,1.0 | 1.0,1.0 | -0.22 |
| 9170 | XLOC_014840 | chr15 | - | 40329186 | 40329556 | 40329186 | 40329219 | 40329509 | 290 | 40329556 | 9170 | 2,2 | 27,11 | 14,8 | 14,22 | 439 | 149 | 0.000345706 | 0.002501231 | 0.025,0.058 | 0.253,0.11 | -0.14 |
| 10558 | XLOC_007087 | chr11 | + | 58377310 | 58377807 | 58377310 | 58377480 | 58377770 | 290 | 58377807 | 10558 | 0,0 | 2,5 | 1,5 | 1,3 | 439 | 149 | 0.000174591 | 0.001379961 | 0.0,0.0 | 0.253,0.361 | -0.307 |
| 14606 | XLOC_018288 | chr17 | + | 79660899 | 79661356 | 79660899 | 79660995 | 79661285 | 290 | 79661356 | 14606 | 39,38 | 6,4 | 91,66 | 0,0 | 439 | 149 | 3.74E-07 | 6.12E-06 | 0.688,0.763 | 1.0,1.0 | -0.274 |
| 7411 | XLOC_021318 | chr19 | + | 50149364 | 50149821 | 50149364 | 50149459 | 50149750 | 291 | 50149821 | 7411 | 3,4 | 8,4 | 19,5 | 5,0 | 440 | 149 | 0.002184273 | 0.0117161 | 0.113,0.253 | 0.563,1.0 | -0.599 |
| 778 | XLOC_026868 | chr20 | + | 61299363 | 61299893 | 61299363 | 61299536 | 61299828 | 292 | 61299893 | 778 | 214,210 | 486,487 | 407,317 | 263,235 | 441 | 149 | 0 | 0 | 0.13,0.127 | 0.343,0.313 | -0.2 |
| 1559 | XLOC_016220 | chr16 | + | 2514039 | 2515054 | 2514039 | 2514230 | 2514522 | 292 | 2515054 | 1559 | 110,103 | 10,26 | 136,89 | 3,3 | 441 | 149 | 3.78E-05 | 0.000368734 | 0.788,0.572 | 0.939,0.909 | -0.244 |
| 1829 | XLOC_014707 | chr15 | + | 75189351 | 75192421 | 75189351 | 75189560 | 75189852 | 292 | 75192421 | 1829 | 17,14 | 121,128 | 41,41 | 34,26 | 441 | 149 | 0 | 0 | 0.045,0.036 | 0.289,0.348 | -0.278 |
| 6696 | XLOC_016235 | chr16 | + | 2809747 | 2810500 | 2809747 | 2810009 | 2810301 | 292 | 2810500 | 6696 | 312,331 | 27,35 | 473,369 | 5,4 | 441 | 149 | 2.89E-13 | 1.31E-11 | 0.796,0.762 | 0.97,0.969 | -0.191 |
| 10597 | XLOC_009338 | chr12 | - | 49233216 | 49233692 | 49233216 | 49233334 | 49233626 | 292 | 49233692 | 10597 | 0,0 | 3,1 | 1,5 | 3,1 | 441 | 149 | 0.001251958 | 0.007367254 | 0.0,0.0 | 0.101,0.628 | -0.364 |
| 9385 | XLOC_044303 | chr8 | - | 56986228 | 56986718 | 56986228 | 56986325 | 56986618 | 293 | 56986718 | 9385 | 57,80 | 6,3 | 126,113 | 0,0 | 442 | 149 | 1.44E-05 | 0.000157123 | 0.762,0.9 | 1.0,1.0 | -0.169 |

| | | | | | | | | | | | | | | | | | | | | | | |
|---|---|---|---|---|---|---|---|---|---|---|---|---|---|---|---|---|---|---|---|---|---|---|
| 3430 | XLOC_001116 | chr1 | - | 52812413 | 52819319 | 52812413 | 52818895 | 52819189 | 294 | 52819319 | 3430 | 25,23 | 88,92 | 55,35 | 37,37 | 443 | 149 | 2.00E-09 | 4.92E-08 | 0.087,0.078 | 0.333,0.241 | -0.205 |
| 5489 | XLOC_028585 | chr22 | - | 21355544 | 21356302 | 21355544 | 21355700 | 21355995 | 295 | 21356302 | 5489 | 78,76 | 6,7 | 86,54 | 0,0 | 444 | 149 | 2.42E-06 | 3.22E-05 | 0.814,0.785 | 1.0,1.0 | -0.201 |
| 13184 | XLOC_046078 | chr9 | + | 131389282 | 131389777 | 131389282 | 131389369 | 131389664 | 295 | 131389777 | 13184 | 0,1 | 2,0 | 2,6 | 0,0 | 444 | 149 | 0.000363764 | 0.002613142 | 0.0,1.0 | 1.0,1.0 | -0.5 |
| 14217 | XLOC_020938 | chr19 | + | 7599635 | 7600463 | 7599635 | 7600099 | 7600394 | 295 | 7600463 | 14217 | 5,3 | 1,2 | 1,2 | 0,0 | 444 | 149 | 0.00392175 | 0.019085759 | 0.627,0.335 | 1.0,1.0 | -0.519 |
| 2694 | XLOC_020866 | chr19 | + | 1244313 | 1244976 | 1244313 | 1244439 | 1244735 | 296 | 1244976 | 2694 | 854,826 | 317,244 | 22,031,740 | 323,272 | 445 | 149 | 4.83E-11 | 1.57E-09 | 0.474,0.531 | 0.695,0.682 | -0.186 |
| 4954 | XLOC_032289 | chr4 | + | 25395931 | 25396346 | 25395931 | 25395996 | 25396292 | 296 | 25396346 | 4954 | 11,28 | 51,46 | 61,41 | 55,27 | 445 | 149 | 1.62E-05 | 0.000171831 | 0.067,0.169 | 0.271,0.337 | -0.186 |
| 5936 | XLOC_030105 | chr3 | - | 169846127 | 169846525 | 169846127 | 169846175 | 169846471 | 296 | 169846525 | 5936 | 1,0 | 8,1 | 7,3 | 8,3 | 445 | 149 | 0.000507137 | 0.003478533 | 0.04,0.0 | 0.227,0.251 | -0.219 |
| 8312 | XLOC_008841 | chr12 | + | 31255908 | 31256329 | 31255908 | 31255954 | 31256250 | 296 | 31256329 | 8312 | 55,63 | 187,182 | 115,61 | 131,91 | 445 | 149 | 3.48E-07 | 5.72E-06 | 0.09,0.104 | 0.227,0.183 | -0.108 |
| 3237 | XLOC_001334 | chr1 | - | 153912906 | 153913524 | 153912906 | 153913082 | 153913379 | 297 | 153913524 | 3237 | 56,50 | 54,82 | 228,174 | 38,29 | 446 | 149 | 0 | 0 | 0.257,0.169 | 0.667,0.667 | -0.454 |
| 4972 | XLOC_032231 | chr4 | + | 1806056 | 1806696 | 1806056 | 1806253 | 1806550 | 297 | 1806696 | 4972 | 31,44 | 12,12 | 13,9 | 2,0 | 446 | 149 | 0.01174363 | 0.047416787 | 0.463,0.551 | 0.685,1.0 | -0.336 |
| 14845 | XLOC_007607 | chr11 | - | 67207256 | 67207723 | 67207256 | 67207314 | 67207611 | 297 | 67207723 | 14845 | 33,26 | 1,5 | 34,49 | 0,1 | 446 | 149 | 0.000985123 | 0.006020081 | 0.917,0.635 | 1.0,0.942 | -0.195 |
| 14838 | XLOC_044013 | chr8 | + | 41466934 | 41467328 | 41466934 | 41467004 | 41467302 | 298 | 41467328 | 14838 | 0,1 | 4,0 | 2,2 | 0,0 | 447 | 149 | 3.77E-05 | 0.000367525 | 0.0,1.0 | 1.0,1.0 | -0.5 |
| 12032 | XLOC_007159 | chr11 | + | 65632190 | 65632616 | 65632190 | 65632286 | 65632585 | 299 | 65632616 | 12032 | 106,88 | 37,19 | 115,77 | 6,7 | 448 | 149 | 1.20E-05 | 0.000133617 | 0.488,0.606 | 0.864,0.785 | -0.278 |
| 754 | XLOC_037851 | chr6 | + | 44197643 | 44198218 | 44197643 | 44197783 | 44198083 | 300 | 44198218 | 754 | 70,80 | 161,142 | 76,73 | 84,66 | 449 | 149 | 0.000135353 | 0.001108839 | 0.126,0.158 | 0.231,0.268 | -0.107 |
| 4417 | XLOC_044293 | chr8 | - | 48846193 | 48846650 | 48846193 | 48846224 | 48846524 | 300 | 48846650 | 4417 | 4,11 | 2,8 | 23,15 | 0,0 | 449 | 149 | 3.12E-09 | 7.35E-08 | 0.399,0.313 | 1.0,1.0 | -0.644 |
| 13002 | XLOC_046501 | chr9 | - | 140346816 | 140347256 | 140346816 | 140346887 | 140347187 | 300 | 140347256 | 13002 | 11,14 | 1,4 | 12,5 | 1,0 | 449 | 149 | 0.003851444 | 0.018828569 | 0.785,0.537 | 0.799,1.0 | -0.238 |
| 1163 | XLOC_047987 | chrX | - | 64743935 | 64744494 | 64743935 | 64744142 | 64744443 | 301 | 64744494 | 1163 | 39,31 | 9,13 | 242,148 | 8,12 | 450 | 149 | 4.42E-06 | 5.54E-05 | 0.589,0.441 | 0.909,0.803 | -0.341 |
| 1265 | XLOC_018778 | chr17 | - | 79779984 | 79781187 | 79779984 | 79780783 | 79781084 | 301 | 79781187 | 1265 | 99,133 | 8,8 | 117,93 | 0,0 | 450 | 149 | 6.68E-07 | 1.03E-05 | 0.804,0.846 | 1.0,1.0 | -0.175 |
| 3028 | XLOC_016775 | chr16 | - | 67863667 | 67864451 | 67863667 | 67863991 | 67864292 | 301 | 67864451 | 3028 | 251,197 | 42,44 | 327,240 | 24,18 | 450 | 149 | 9.90E-06 | 0.000112877 | 0.664,0.597 | 0.819,0.815 | -0.186 |
| 831 | XLOC_013335 | chr14 | - | 24773749 | 24777126 | 24773749 | 24776721 | 24777023 | 302 | 24777126 | 831 | 209,227 | 4,13 | 334,195 | 0,0 | 451 | 149 | 6.26E-07 | 9.72E-06 | 0.945,0.852 | 1.0,1.0 | -0.102 |
| 1575 | XLOC_045894 | chr9 | + | 34646639 | 34647255 | 34646639 | 34646783 | 34647085 | 302 | 34647255 | 1575 | 31,40 | 52,58 | 69,25 | 21,14 | 451 | 149 | 5.84E-06 | 7.07E-05 | 0.165,0.186 | 0.521,0.371 | -0.271 |
| 9688 | XLOC_013336 | chr14 | - | 24910879 | 24911378 | 24910879 | 24911001 | 24911303 | 302 | 24911378 | 9688 | 30,47 | 14,19 | 47,23 | 3,4 | 451 | 149 | 0.001059961 | 0.006415438 | 0.415,0.45 | 0.838,0.655 | -0.314 |
| 14270 | XLOC_029428 | chr3 | + | 49735474 | 49735959 | 49735474 | 49735570 | 49735872 | 302 | 49735959 | 14270 | 15,18 | 94,92 | 99,58 | 141,133 | 451 | 149 | 1.13E-07 | 2.06E-06 | 0.05,0.061 | 0.188,0.126 | -0.102 |
| 2196 | XLOC_009410 | chr12 | - | 56716398 | 56716993 | 56716398 | 56716542 | 56716845 | 303 | 56716993 | 2196 | 238,186 | 78,59 | 175,146 | 24,19 | 452 | 149 | 1.85E-05 | 0.000194527 | 0.501,0.51 | 0.706,0.717 | -0.206 |
| 6522 | XLOC_023400 | chr2 | + | 27291499 | 27291962 | 27291499 | 27291612 | 27291915 | 303 | 27291962 | 6522 | 4,13 | 11,10 | 17,12 | 2,3 | 452 | 149 | 0.000472378 | 0.00328155 | 0.107,0.3 | 0.737,0.569 | -0.45 |
| 13728 | XLOC_047722 | chrX | + | 54838006 | 54838473 | 54838006 | 54838086 | 54838389 | 303 | 54838473 | 13728 | 0,2 | 4,0 | 4,4 | 1,0 | 452 | 149 | 0.001834256 | 0.010154919 | 0.0,1.0 | 0.569,1.0 | -0.284 |
| 1738 | XLOC_028752 | chr22 | - | 50965504 | 50968148 | 50965504 | 50965712 | 50966016 | 304 | 50968148 | 1738 | 59,33 | 92,83 | 25,17 | 25,11 | 453 | 149 | 0.00714482 | 0.031230671 | 0.174,0.116 | 0.248,0.337 | -0.147 |
| 5341 | XLOC_029378 | chr3 | + | 38540612 | 38542002 | 38540612 | 38540867 | 38541171 | 304 | 38542002 | 5341 | 0,1 | 1,1 | 8,1 | 0,0 | 453 | 149 | 7.42E-05 | 0.000659605 | 0.0,0.248 | 1.0,1.0 | -0.876 |
| 632 | XLOC_029385 | chr3 | + | 41280624 | 41282321 | 41280624 | 41280845 | 41281150 | 305 | 41282321 | 632 | 614,577 | 161,178 | 13,251,100 | 111,63 | 454 | 149 | 0 | 0 | 0.556,0.515 | 0.797,0.851 | -0.288 |
| 896 | XLOC_024067 | chr2 | - | 68546353 | 68547110 | 68546353 | 68546598 | 68546903 | 305 | 68547110 | 896 | 5,7 | 1,4 | 7,2 | 0,0 | 454 | 149 | 2.95E-06 | 3.84E-05 | 0.621,0.365 | 1.0,1.0 | -0.507 |
| 1214 | XLOC_014981 | chr15 | - | 74280939 | 74281598 | 74280939 | 74281143 | 74281448 | 305 | 74281598 | 1214 | 5,4 | 2,6 | 5,7 | 1,0 | 454 | 149 | 0.000247538 | 0.001868506 | 0.451,0.18 | 0.621,1.0 | -0.495 |
| 7568 | XLOC_018770 | chr17 | - | 79210779 | 79212947 | 79210779 | 79210881 | 79211186 | 305 | 79212947 | 7568 | 22,24 | 0,4 | 17,12 | 0,0 | 454 | 149 | 0.012068696 | 0.048560171 | 1.0,0.663 | 1.0,1.0 | -0.168 |
| 3955 | XLOC_044466 | chr8 | - | 145740709 | 145741181 | 145740709 | 145740841 | 145741147 | 306 | 145741181 | 3955 | 36,55 | 206,220 | 215,177 | 105,95 | 455 | 149 | 0 | 0 | 0.054,0.076 | 0.401,0.379 | -0.325 |
| 343 | XLOC_041541 | chr7 | - | 74299454 | 74300329 | 74299454 | 74299941 | 74300249 | 308 | 74300329 | 343 | 97,51 | 34,19 | 72,40 | 9,6 | 457 | 149 | 0.001446841 | 0.008319927 | 0.482,0.467 | 0.723,0.685 | -0.229 |
| 78 | XLOC_046093 | chr9 | + | 132571183 | 132575317 | 132571183 | 132571311 | 132571620 | 309 | 132575317 | 78 | 16,31 | 104,99 | 56,42 | 102,38 | 458 | 149 | 3.33E-06 | 4.27E-05 | 0.048,0.092 | 0.152,0.264 | -0.138 |
| 132 | XLOC_044027 | chr8 | + | 48641494 | 48642027 | 48641494 | 48641664 | 48641973 | 309 | 48642027 | 132 | 25,28 | 5,8 | 44,25 | 0,3 | 458 | 149 | 0.002576457 | 0.013382605 | 0.619,0.532 | 1.0,0.731 | -0.29 |
| 3041 | XLOC_001428 | chr1 | - | 179068450 | 179078576 | 179068450 | 179078033 | 179078342 | 309 | 179078576 | 3041 | 161,127 | 39,51 | 326,211 | 33,26 | 458 | 149 | 1.09E-05 | 0.000122456 | 0.573,0.448 | 0.763,0.725 | -0.234 |
| 3954 | XLOC_044466 | chr8 | - | 145740709 | 145741181 | 145740709 | 145740837 | 145741147 | 310 | 145741181 | 3954 | 41,56 | 36,57 | 216,179 | 55,25 | 459 | 149 | 2.91E-10 | 8.29E-09 | 0.27,0.242 | 0.56,0.699 | -0.373 |
| 883 | XLOC_024114 | chr2 | - | 85785562 | 85786198 | 85785562 | 85785728 | 85786039 | 311 | 85786198 | 883 | 95,137 | 39,39 | 166,114 | 30,16 | 460 | 149 | 0.001973253 | 0.010747169 | 0.441,0.532 | 0.642,0.698 | -0.183 |
| 556 | XLOC_030002 | chr3 | - | 123675201 | 123675642 | 123675201 | 123675274 | 123675586 | 312 | 123675642 | 556 | 223,261 | 49,55 | 118,160 | 12,6 | 461 | 149 | 3.23E-05 | 0.000320217 | 0.595,0.605 | 0.761,0.896 | -0.229 |
| 1448 | XLOC_016367 | chr16 | + | 30708740 | 30711302 | 30708740 | 30710916 | 30711228 | 312 | 30711302 | 1448 | 11,9 | 1,3 | 6,5 | 0,0 | 461 | 149 | 0.000804773 | 0.0050831 | 0.78,0.492 | 1.0,1.0 | -0.364 |
| 865 | XLOC_018679 | chr17 | - | 62968871 | 62969635 | 62968871 | 62969045 | 62969359 | 314 | 62969635 | 865 | 13,25 | 21,22 | 15,12 | 3,1 | 463 | 149 | 0.00013313 | 0.001093141 | 0.166,0.268 | 0.617,0.794 | -0.488 |
| 12845 | XLOC_021518 | chr19 | - | 8654335 | 8654875 | 8654335 | 8654458 | 8654772 | 314 | 8654875 | 12845 | 8,12 | 2,3 | 8,10 | 0,0 | 463 | 149 | 5.95E-05 | 0.000545802 | 0.563,0.563 | 1.0,1.0 | -0.437 |
| 13409 | XLOC_024005 | chr2 | - | 38973835 | 38974272 | 38973835 | 38973889 | 38974203 | 314 | 38974272 | 13409 | 34,43 | 6,0 | 43,58 | 1,0 | 463 | 149 | 0.003847792 | 0.018823129 | 0.646,1.0 | 0.933,1.0 | -0.144 |
| 12334 | XLOC_020062 | chr18 | + | 21109601 | 21110096 | 21109601 | 21109676 | 21109991 | 315 | 21110096 | 12334 | 44,15 | 78,68 | 62,61 | 75,44 | 464 | 149 | 5.99E-05 | 0.00054826 | 0.153,0.066 | 0.21,0.308 | -0.15 |
| 3418 | XLOC_000391 | chr1 | + | 101440365 | 101447410 | 101440365 | 101440458 | 101440774 | 316 | 101447410 | 3418 | 129,131 | 42,30 | 254,215 | 28,24 | 465 | 149 | 6.26E-05 | 0.000570932 | 0.496,0.583 | 0.744,0.742 | -0.204 |
| 879 | XLOC_044459 | chr8 | - | 145580649 | 145581162 | 145580649 | 145580781 | 145581098 | 317 | 145581162 | 879 | 166,160 | 33,23 | 240,154 | 5,10 | 466 | 149 | 8.21E-07 | 1.23E-05 | 0.617,0.69 | 0.939,0.831 | -0.232 |
| 2464 | XLOC_007511 | chr11 | - | 47752925 | 47754222 | 47752925 | 47753771 | 47754088 | 317 | 47754222 | 2464 | 382,360 | 108,76 | 446,304 | 62,38 | 466 | 149 | 0.000271763 | 0.002030747 | 0.531,0.602 | 0.697,0.719 | -0.141 |
| 11336 | XLOC_021026 | chr19 | + | 16973930 | 16974607 | 16973930 | 16974173 | 16974490 | 317 | 16974607 | 11336 | 33,28 | 6,5 | 23,34 | 0,2 | 466 | 149 | 0.001555677 | 0.008837821 | 0.637,0.642 | 1.0,0.845 | -0.283 |
| 2302 | XLOC_008778 | chr12 | + | 9462468 | 9463037 | 9462468 | 9462616 | 9462934 | 318 | 9463037 | 2302 | 34,22 | 23,24 | 53,31 | 19,5 | 467 | 149 | 0.002583866 | 0.013416155 | 0.32,0.226 | 0.471,0.664 | -0.294 |
| 7505 | XLOC_021416 | chr19 | + | 58757708 | 58758160 | 58757708 | 58757793 | 58758111 | 318 | 58758160 | 7505 | 30,28 | 33,29 | 50,18 | 9,3 | 467 | 149 | 1.38E-07 | 2.45E-06 | 0.225,0.236 | 0.639,0.657 | -0.418 |
| 14074 | XLOC_018785 | chr17 | - | 79894623 | 79895123 | 79894623 | 79894737 | 79895055 | 318 | 79895123 | 14074 | 100,100 | 0,13 | 130,80 | 0,0 | 467 | 149 | 6.16E-05 | 0.000562411 | 1.0,0.711 | 1.0,1.0 | -0.145 |
| 2410 | XLOC_007749 | chr11 | - | 124972027 | 124972705 | 124972027 | 124972213 | 124972532 | 319 | 124972705 | 2410 | 1,4 | 2,9 | 17,20 | 0,1 | 468 | 149 | 1.55E-10 | 4.60E-09 | 0.137,0.124 | 1.0,0.864 | -0.801 |
| 11093 | XLOC_028526 | chr22 | + | 44330485 | 44330964 | 44330485 | 44330546 | 44330865 | 319 | 44330964 | 11093 | 15,26 | 3,1 | 13,10 | 0,0 | 468 | 149 | 0.005127444 | 0.023719257 | 0.614,0.892 | 1.0,1.0 | -0.247 |
| 1950 | XLOC_044467 | chr8 | - | 145750199 | 145753171 | 145750199 | 145752023 | 145752343 | 320 | 145753171 | 1950 | 206,234 | 43,51 | 284,177 | 31,16 | 469 | 149 | 0.000382234 | 0.002730546 | 0.603,0.593 | 0.744,0.778 | -0.163 |
| 3490 | XLOC_048133 | chrX | - | 153735736 | 153736216 | 153735736 | 153735821 | 153736141 | 320 | 153736216 | 3490 | 32,25 | 149,125 | 77,51 | 96,85 | 469 | 149 | 2.75E-09 | 6.56E-08 | 0.064,0.06 | 0.203,0.16 | -0.119 |
| 5104 | XLOC_029869 | chr3 | - | 52029057 | 52029479 | 52029057 | 52029122 | 52029442 | 320 | 52029479 | 5104 | 9,4 | 5,1 | 18,16 | 0,1 | 469 | 149 | 4.60E-05 | 0.000437569 | 0.364,0.56 | 1.0,0.836 | -0.456 |
| 6951 | XLOC_021331 | chr19 | + | 50919867 | 50920354 | 50919867 | 50919980 | 50920301 | 321 | 50920354 | 6951 | 60,79 | 171,182 | 156,131 | 223,129 | 470 | 149 | 3.45E-06 | 4.40E-05 | 0.1,0.121 | 0.182,0.244 | -0.102 |
| 7666 | XLOC_018637 | chr17 | - | 54972882 | 54973313 | 54972882 | 54972965 | 54973286 | 321 | 54973313 | 7666 | 45,53 | 148,127 | 139,113 | 121,107 | 470 | 149 | 1.23E-10 | 3.68E-09 | 0.088,0.117 | 0.267,0.251 | -0.157 |
| 5340 | XLOC_029378 | chr3 | + | 38540416 | 38540867 | 38540416 | 38540480 | 38540802 | 322 | 38540867 | 5340 | 8,2 | 1,2 | 3,5 | 0,0 | 471 | 149 | 0.004619964 | 0.021749252 | 0.717,0.24 | 1.0,1.0 | -0.522 |
| 751 | XLOC_046136 | chr9 | + | 139443195 | 139446260 | 139443195 | 139443260 | 139443583 | 323 | 139446260 | 751 | 43,49 | 9,11 | 33,25 | 0,3 | 472 | 149 | 0.005715014 | 0.02596638 | 0.601,0.584 | 1.0,0.725 | -0.27 |
| 3191 | XLOC_000975 | chr1 | - | 28906044 | 28906493 | 28906044 | 28906099 | 28906423 | 324 | 28906493 | 3191 | 276,234 | 52,33 | 563,458 | 51,26 | 473 | 149 | 5.54E-05 | 0.000513909 | 0.626,0.691 | 0.777,0.847 | -0.154 |
| 12709 | XLOC_005370 | chr10 | - | 104865114 | 104865558 | 104865114 | 104865138 | 104865462 | 324 | 104865558 | 12709 | 0,1 | 30,22 | 1,4 | 7,3 | 473 | 149 | 0.009274353 | 0.038820892 | 0.0,0.014 | 0.043,0.296 | -0.162 |
| 6778 | XLOC_021732 | chr19 | - | 41246695 | 41248675 | 41246695 | 41248094 | 41248419 | 325 | 41248675 | 6778 | 68,67 | 2,7 | 27,28 | 0,0 | 474 | 149 | 0.000212047 | 0.001632011 | 0.914,0.751 | 1.0,1.0 | -0.167 |
| 2229 | XLOC_009247 | chr12 | - | 9573326 | 9574486 | 9573326 | 9574087 | 9574413 | 326 | 9574486 | 2229 | 190,118 | 61,38 | 195,147 | 18,11 | 475 | 149 | 7.08E-09 | 1.57E-07 | 0.494,0.493 | 0.773,0.807 | -0.297 |

| | | | | | | | | | | | | | | | | | | | | | | |
|---|---|---|---|---|---|---|---|---|---|---|---|---|---|---|---|---|---|---|---|---|---|---|
| 10786 | XLOC_009493 | chr12 | - | 96414467 | 96414915 | 96414467 | 96414536 | 96414862 | 326 | 96414915 | 10786 | 0,0 | 3,2 | 12,3 | 0,0 | 475 | 149 | 4.81E-12 | 1.80E-10 | 0.0,0.0 | 1.0,1.0 | -1 |
| 971 | XLOC_000458 | chr1 | + | 144606368 | 144609918 | 144606368 | 144606510 | 144606837 | 327 | 144609918 | 971 | 71,75 | 45,35 | 121,124 | 30,30 | 476 | 149 | 0.000307761 | 0.002264986 | 0.331,0.401 | 0.558,0.564 | -0.195 |
| 3428 | XLOC_001045 | chr1 | - | 40041430 | 40042559 | 40041430 | 40041895 | 40042223 | 328 | 40042559 | 3428 | 12,501,196 | 113,55 | 21,591,547 | 24,28 | 477 | 149 | 3.44E-15 | 1.98E-13 | 0.776,0.872 | 0.966,0.945 | -0.131 |
| 3919 | XLOC_045897 | chr9 | + | 35606833 | 35607406 | 35606833 | 35606980 | 35607308 | 328 | 35607406 | 3919 | 10,18 | 4,0 | 17,9 | 0,0 | 477 | 149 | 0.005832599 | 0.026401859 | 0.438,1.0 | 1.0,1.0 | -0.281 |
| 2835 | XLOC_009549 | chr12 | - | 112277460 | 112278281 | 112277460 | 112277807 | 112278136 | 329 | 112278281 | 2835 | 246,208 | 43,35 | 409,312 | 2,3 | 478 | 149 | 0 | 0 | 0.641,0.649 | 0.985,0.97 | -0.333 |
| 3440 | XLOC_005412 | chr10 | - | 121336122 | 121336715 | 121336122 | 121336262 | 121336591 | 329 | 121336715 | 3440 | 287,283 | 410,429 | 566,510 | 371,267 | 478 | 149 | 1.39E-13 | 6.65E-12 | 0.179,0.171 | 0.322,0.373 | -0.173 |
| 2868 | XLOC_047841 | chrX | + | 133559230 | 133562819 | 133559230 | 133559360 | 133559690 | 330 | 133562819 | 2868 | 131,121 | 90,107 | 243,198 | 32,13 | 479 | 149 | 0 | 0 | 0.312,0.26 | 0.703,0.826 | -0.478 |
| 6245 | XLOC_014964 | chr15 | - | 68503600 | 68504079 | 68503600 | 68503656 | 68503986 | 330 | 68504079 | 6245 | 25,17 | 3,1 | 50,43 | 0,0 | 479 | 149 | 0.003498501 | 0.017393519 | 0.722,0.841 | 1.0,1.0 | -0.219 |
| 11693 | XLOC_007557 | chr11 | - | 63766430 | 63766876 | 63766430 | 63766472 | 63766802 | 330 | 63766876 | 11693 | 279,252 | 37,48 | 452,332 | 26,30 | 479 | 149 | 0.000169406 | 0.00134411 | 0.701,0.62 | 0.844,0.775 | -0.149 |
| 619 | XLOC_029419 | chr3 | + | 48500757 | 48501315 | 48500757 | 48500853 | 48501185 | 332 | 48501315 | 619 | 2,2 | 34,36 | 2,6 | 12,7 | 481 | 149 | 0.003979772 | 0.019309085 | 0.018,0.017 | 0.049,0.21 | -0.112 |
| 3881 | XLOC_046066 | chr9 | + | 130569270 | 130569761 | 130569270 | 130569366 | 130569699 | 333 | 130569761 | 3881 | 8,18 | 4,1 | 11,10 | 1,0 | 482 | 149 | 0.003682731 | 0.018179381 | 0.382,0.848 | 0.773,1.0 | -0.272 |
| 3402 | XLOC_021451 | chr19 | - | 1609289 | 1611848 | 1609289 | 1611368 | 1611702 | 334 | 1611848 | 3402 | 486,609 | 101,98 | 740,564 | 28,15 | 483 | 149 | 0 | 0 | 0.597,0.657 | 0.891,0.921 | -0.279 |
| 3133 | XLOC_021059 | chr19 | + | 18699804 | 18700493 | 18699804 | 18699887 | 18700222 | 335 | 18700493 | 3133 | 147,115 | 130,99 | 125,74 | 36,39 | 484 | 149 | 0.000737046 | 0.004718634 | 0.258,0.263 | 0.517,0.369 | -0.182 |
| 4505 | XLOC_038237 | chr6 | - | 34392848 | 34393804 | 34392848 | 34392998 | 34393334 | 336 | 34393804 | 4505 | 4,3 | 1,3 | 1,8 | 0,0 | 485 | 149 | 1.53E-05 | 0.000164879 | 0.551,0.235 | 1.0,1.0 | -0.607 |
| 1227 | XLOC_021210 | chr19 | + | 39998469 | 40000464 | 39998469 | 39998554 | 39998891 | 337 | 40000464 | 1227 | 2,9 | 22,7 | 59,54 | 31,24 | 486 | 149 | 0.000199369 | 0.00154651 | 0.027,0.283 | 0.368,0.408 | -0.233 |
| 2668 | XLOC_001160 | chr1 | - | 78444568 | 78453654 | 78444568 | 78444743 | 78445080 | 337 | 78453654 | 2668 | 72,78 | 6,0 | 53,31 | 0,0 | 486 | 149 | 0.002548979 | 0.013259387 | 0.786,1.0 | 1.0,1.0 | -0.107 |
| 2906 | XLOC_000386 | chr1 | + | 100605963 | 100606575 | 100605963 | 100606070 | 100606407 | 337 | 100606575 | 2906 | 75,65 | 36,33 | 189,133 | 6,14 | 486 | 149 | 4.41E-11 | 1.46E-09 | 0.39,0.377 | 0.906,0.744 | -0.441 |
| 3370 | XLOC_017907 | chr17 | + | 7358602 | 7359260 | 7358602 | 7358775 | 7359112 | 337 | 7359260 | 3370 | 17,24 | 6,6 | 16,13 | 1,1 | 486 | 149 | 0.01218451 | 0.048942503 | 0.465,0.551 | 0.831,0.799 | -0.307 |
| 5374 | XLOC_028754 | chr22 | - | 51016977 | 51017684 | 51016977 | 51017088 | 51017425 | 337 | 51017684 | 5374 | 2,6 | 3,0 | 1,2 | 0,0 | 486 | 149 | 0.009529924 | 0.039678986 | 0.17,1.0 | 1.0,1.0 | -0.415 |
| 3380 | XLOC_001300 | chr1 | - | 150237440 | 150239056 | 150237440 | 150238594 | 150238932 | 338 | 150239056 | 3380 | 10,871,142 | 259,237 | 19,091,379 | 80,65 | 487 | 149 | 0 | 0 | 0.562,0.596 | 0.88,0.867 | -0.294 |
| 11113 | XLOC_041587 | chr7 | - | 92206968 | 92207496 | 92206968 | 92207125 | 92207463 | 338 | 92207496 | 11113 | 9,8 | 1,4 | 35,22 | 1,1 | 487 | 149 | 0.008835969 | 0.037351208 | 0.734,0.38 | 0.915,0.871 | -0.336 |
| 4634 | XLOC_037635 | chr6 | + | 7404650 | 7405254 | 7404650 | 7404788 | 7405128 | 340 | 7405254 | 4634 | 2,0 | 0,4 | 1,4 | 0,1 | 489 | 149 | 0.00164567 | 0.009278161 | 1.0,0.0 | 1.0,0.549 | -0.274 |
| 2891 | XLOC_016217 | chr16 | + | 2287767 | 2289507 | 2287767 | 2288019 | 2288362 | 343 | 2289507 | 2891 | 3,4 | 16,11 | 8,9 | 4,9 | 492 | 149 | 0.002516296 | 0.013147482 | 0.054,0.099 | 0.377,0.232 | -0.228 |
| 3033 | XLOC_041344 | chr7 | + | 150066759 | 150071492 | 150066759 | 150067973 | 150068316 | 343 | 150071492 | 3033 | 393,349 | 49,57 | 942,753 | 46,34 | 492 | 149 | 1.28E-09 | 3.29E-08 | 0.708,0.65 | 0.861,0.87 | -0.186 |
| 5850 | XLOC_026805 | chr20 | + | 43560982 | 43561826 | 43560982 | 43561354 | 43561697 | 343 | 43561826 | 5850 | 70,63 | 6,3 | 39,19 | 0,0 | 492 | 149 | 0.00015074 | 0.001214435 | 0.779,0.864 | 1.0,1.0 | -0.178 |
| 9351 | XLOC_000897 | chr1 | - | 16270302 | 16271334 | 16270302 | 16270450 | 16270794 | 344 | 16271334 | 9351 | 5,1 | 1,2 | 13,1 | 0,1 | 493 | 149 | 0.007006423 | 0.030739892 | 0.602,0.131 | 1.0,0.232 | -0.249 |
| 302 | XLOC_037951 | chr6 | + | 109762754 | 109763270 | 109762754 | 109762831 | 109763176 | 345 | 109763270 | 302 | 40,32 | 42,55 | 43,41 | 6,14 | 494 | 149 | 2.65E-06 | 3.49E-05 | 0.223,0.149 | 0.684,0.469 | -0.391 |
| 2959 | XLOC_004790 | chr10 | + | 22607200 | 22607642 | 22607200 | 22607235 | 22607581 | 346 | 22607642 | 2959 | 74,82 | 180,188 | 136,106 | 144,102 | 495 | 149 | 2.67E-07 | 4.48E-06 | 0.11,0.116 | 0.221,0.238 | -0.116 |
| 7413 | XLOC_021318 | chr19 | + | 50149364 | 50149821 | 50149364 | 50149459 | 50149805 | 346 | 50149821 | 7413 | 24,30 | 5,4 | 64,42 | 0,2 | 495 | 149 | 0.000202794 | 0.00156935 | 0.591,0.693 | 1.0,0.863 | -0.29 |
| 9317 | XLOC_008878 | chr12 | + | 50472283 | 50472763 | 50472283 | 50472360 | 50472706 | 346 | 50472763 | 9317 | 11,11 | 75,58 | 30,17 | 9,18 | 495 | 149 | 1.77E-07 | 3.09E-06 | 0.042,0.054 | 0.501,0.221 | -0.313 |
| 3150 | XLOC_001352 | chr1 | - | 155238083 | 155239507 | 155238083 | 155238150 | 155238498 | 348 | 155239507 | 3150 | 255,291 | 90,75 | 284,175 | 43,29 | 497 | 149 | 0.000502855 | 0.003453641 | 0.459,0.538 | 0.664,0.644 | -0.155 |
| 3331 | XLOC_000851 | chr1 | - | 3547163 | 3548221 | 3547163 | 3547681 | 3548029 | 348 | 3548221 | 3331 | 37,21 | 29,24 | 98,84 | 18,16 | 497 | 149 | 7.55E-09 | 1.66E-07 | 0.277,0.208 | 0.62,0.611 | -0.373 |
| 8097 | XLOC_017884 | chr17 | + | 5284683 | 5285164 | 5284683 | 5284800 | 5285148 | 348 | 5285164 | 8097 | 24,21 | 3,2 | 25,15 | 0,0 | 497 | 149 | 0.00049044 | 0.003383725 | 0.706,0.759 | 1.0,1.0 | -0.268 |
| 9766 | XLOC_021379 | chr19 | + | 56153422 | 56155140 | 56153422 | 56153514 | 56153862 | 348 | 56155140 | 9766 | 26,45 | 6,11 | 50,29 | 4,2 | 497 | 149 | 0.010455586 | 0.04294834 | 0.565,0.551 | 0.789,0.813 | -0.243 |
| 119 | XLOC_044167 | chr8 | + | 144460438 | 144460990 | 144460438 | 144460516 | 144460865 | 349 | 144460990 | 119 | 214,191 | 15,23 | 83,87 | 4,2 | 498 | 149 | 0.008112754 | 0.034804502 | 0.81,0.713 | 0.861,0.929 | -0.133 |
| 2677 | XLOC_026732 | chr20 | + | 23350220 | 23354006 | 23350220 | 23350378 | 23350727 | 349 | 23354006 | 2677 | 9,2 | 19,18 | 19,7 | 8,6 | 498 | 149 | 0.000187874 | 0.001468367 | 0.124,0.032 | 0.415,0.259 | -0.259 |
| 10932 | XLOC_009324 | chr12 | - | 48143712 | 48144203 | 48143712 | 48143773 | 48144123 | 350 | 48144203 | 10932 | 0,0 | 18,14 | 6,4 | 17,8 | 499 | 149 | 0.000167891 | 0.00133284 | 0.0,0.0 | 0.095,0.13 | -0.113 |
| 3898 | XLOC_045993 | chr9 | + | 104161797 | 104162292 | 104161797 | 104161812 | 104162165 | 353 | 104162292 | 3898 | 1,2 | 1,5 | 27,13 | 0,3 | 502 | 149 | 3.67E-05 | 0.000359898 | 0.229,0.106 | 1.0,0.563 | -0.614 |
| 2577 | XLOC_006946 | chr11 | + | 581491 | 582081 | 581491 | 581606 | 581961 | 355 | 582081 | 2577 | 18,5 | 53,54 | 61,39 | 74,30 | 504 | 149 | 2.22E-08 | 4.55E-07 | 0.091,0.027 | 0.196,0.278 | -0.178 |
| 1378 | XLOC_014991 | chr15 | - | 75652905 | 75653532 | 75652905 | 75653029 | 75653386 | 357 | 75653532 | 1378 | 218,184 | 47,35 | 220,180 | 26,14 | 506 | 149 | 0.001531785 | 0.008719627 | 0.577,0.608 | 0.714,0.791 | -0.16 |
| 4035 | XLOC_046494 | chr9 | - | 140006141 | 140006621 | 140006141 | 140006206 | 140006563 | 357 | 140006621 | 4035 | 70,71 | 17,35 | 59,44 | 9,0 | 506 | 149 | 0.002327948 | 0.012323116 | 0.548,0.374 | 0.659,1.0 | -0.368 |
| 5376 | XLOC_028754 | chr22 | - | 51018219 | 51018700 | 51018219 | 51018259 | 51018618 | 359 | 51018700 | 5376 | 40,47 | 3,2 | 28,20 | 0,0 | 508 | 149 | 0.003092652 | 0.015655029 | 0.796,0.873 | 1.0,1.0 | -0.165 |
| 1670 | XLOC_018164 | chr17 | + | 54939500 | 54946418 | 54939500 | 54939612 | 54939972 | 360 | 54946418 | 1670 | 26,49 | 66,58 | 48,39 | 23,11 | 509 | 149 | 1.12E-06 | 1.61E-05 | 0.103,0.198 | 0.379,0.509 | -0.293 |
| 987 | XLOC_017951 | chr17 | + | 17918470 | 17920080 | 17918470 | 17919509 | 17919871 | 362 | 17920080 | 987 | 16,18 | 0,11 | 90,45 | 0,0 | 511 | 149 | 0.000398314 | 0.002830232 | 1.0,0.323 | 1.0,1.0 | -0.339 |
| 1029 | XLOC_021732 | chr19 | - | 41249971 | 41250588 | 41249971 | 41250101 | 41250463 | 362 | 41250588 | 1029 | 13,17 | 56,54 | 9,21 | 14,18 | 511 | 149 | 0.000684693 | 0.004470608 | 0.063,0.084 | 0.158,0.254 | -0.133 |
| 15189 | XLOC_007075 | chr11 | + | 47310518 | 47311044 | 47310518 | 47310578 | 47310941 | 363 | 47311044 | 15189 | 9,5 | 34,29 | 16,10 | 10,5 | 512 | 149 | 1.69E-06 | 2.32E-05 | 0.072,0.048 | 0.318,0.368 | -0.283 |
| 2576 | XLOC_006946 | chr11 | + | 581491 | 582081 | 581491 | 581597 | 581961 | 364 | 582081 | 2576 | 18,5 | 5,18 | 67,40 | 13,9 | 513 | 149 | 0.007773119 | 0.033561527 | 0.511,0.075 | 0.6,0.563 | -0.288 |
| 3397 | XLOC_014828 | chr15 | - | 34651789 | 34652410 | 34651789 | 34651946 | 34652311 | 365 | 34652410 | 3397 | 203,219 | 82,144 | 461,294 | 108,64 | 514 | 149 | 7.11E-06 | 8.39E-05 | 0.418,0.306 | 0.553,0.571 | -0.2 |
| 2578 | XLOC_006946 | chr11 | + | 581491 | 582081 | 581491 | 581597 | 581964 | 367 | 582081 | 2578 | 18,5 | 6,0 | 71,40 | 0,0 | 516 | 149 | 0.000338532 | 0.002461655 | 0.464,1.0 | 1.0,1.0 | -0.268 |
| 961 | XLOC_023476 | chr2 | + | 55196996 | 55199148 | 55196996 | 55197431 | 55197799 | 368 | 55199148 | 961 | 25,26 | 0,6 | 8,5 | 0,0 | 517 | 149 | 0.00071433 | 0.004606612 | 1.0,0.555 | 1.0,1.0 | -0.222 |
| 498 | XLOC_046480 | chr9 | - | 139326275 | 139327527 | 139326275 | 139327038 | 139327407 | 369 | 139327527 | 498 | 24,13 | 26,13 | 37,28 | 13,4 | 518 | 149 | 0.000601698 | 0.004015963 | 0.21,0.223 | 0.45,0.668 | -0.343 |
| 526 | XLOC_032657 | chr4 | - | 83343838 | 83347282 | 83343838 | 83346820 | 83347189 | 369 | 83347282 | 526 | 11,251,059 | 115,123 | 47,863,500 | 183,109 | 518 | 149 | 0 | 0 | 0.738,0.712 | 0.883,0.902 | -0.168 |
| 2585 | XLOC_000006 | chr1 | + | 982199 | 982834 | 982199 | 982337 | 982706 | 369 | 982834 | 2585 | 85,88 | 302,291 | 243,184 | 193,101 | 518 | 149 | 0 | 0 | 0.075,0.08 | 0.266,0.344 | -0.227 |
| 3421 | XLOC_004836 | chr10 | + | 51746924 | 51749136 | 51746924 | 51748698 | 51749067 | 369 | 51749136 | 3421 | 53,49 | 27,21 | 125,91 | 27,15 | 518 | 149 | 0.000603032 | 0.00402107 | 0.361,0.402 | 0.571,0.636 | -0.222 |
| 10189 | XLOC_018411 | chr17 | - | 8133208 | 8133786 | 8133208 | 8133240 | 8133611 | 371 | 8133786 | 10189 | 47,26 | 9,2 | 45,32 | 0,0 | 520 | 149 | 1.93E-06 | 2.63E-05 | 0.599,0.788 | 1.0,1.0 | -0.306 |
| 1493 | XLOC_016619 | chr16 | - | 4853208 | 4855311 | 4853208 | 4854149 | 4854522 | 373 | 4855311 | 1493 | 583,575 | 35,55 | 879,725 | 26,4 | 522 | 149 | 4.69E-10 | 1.29E-08 | 0.826,0.749 | 0.906,0.981 | -0.156 |
| 11186 | XLOC_000548 | chr1 | + | 155295205 | 155295701 | 155295205 | 155295281 | 155295654 | 373 | 155295701 | 11186 | 23,17 | 5,1 | 17,11 | 0,1 | 522 | 149 | 0.002962488 | 0.015085357 | 0.568,0.829 | 1.0,0.758 | -0.181 |
| 2181 | XLOC_009465 | chr12 | - | 76738310 | 76742847 | 76738310 | 76741567 | 76741941 | 374 | 76742847 | 2181 | 38,25 | 33,15 | 35,48 | 9,6 | 523 | 149 | 0.000153676 | 0.001235267 | 0.247,0.322 | 0.526,0.695 | -0.326 |
| 2727 | XLOC_020868 | chr19 | + | 1271979 | 1273323 | 1271979 | 1272050 | 1272425 | 375 | 1273323 | 2727 | 320,372 | 393,363 | 725,485 | 359,243 | 524 | 149 | 1.83E-10 | 5.40E-09 | 0.188,0.226 | 0.365,0.362 | -0.156 |
| 654 | XLOC_028754 | chr22 | - | 51016977 | 51017684 | 51016977 | 51017088 | 51017464 | 376 | 51017684 | 654 | 57,47 | 28,19 | 38,31 | 5,6 | 525 | 149 | 0.003707074 | 0.018273998 | 0.366,0.412 | 0.683,0.595 | -0.25 |
| 1135 | XLOC_021070 | chr19 | + | 19455657 | 19456178 | 19455657 | 19455735 | 19456112 | 377 | 19456178 | 1135 | 20,35 | 61,75 | 137,80 | 70,66 | 526 | 149 | 2.41E-09 | 5.82E-08 | 0.085,0.117 | 0.357,0.256 | -0.205 |
| 993 | XLOC_026793 | chr20 | + | 42158923 | 42159674 | 42158923 | 42159059 | 42159437 | 378 | 42159674 | 993 | 20,13 | 2,2 | 18,7 | 0,0 | 527 | 149 | 0.001604232 | 0.009098903 | 0.739,0.648 | 1.0,1.0 | -0.306 |
| 15166 | XLOC_000275 | chr1 | + | 45478807 | 45479280 | 45478807 | 45478887 | 45479265 | 378 | 45479280 | 15166 | 73,78 | 24,16 | 157,124 | 4,15 | 527 | 149 | 0.000330945 | 0.002414224 | 0.462,0.58 | 0.917,0.7 | -0.287 |
| 1529 | XLOC_016432 | chr16 | + | 67000146 | 67000768 | 67000146 | 67000248 | 67000627 | 379 | 67000768 | 1529 | 60,53 | 42,28 | 207,176 | 25,27 | 528 | 149 | 4.49E-11 | 1.47E-09 | 0.287,0.348 | 0.7,0.648 | -0.356 |
| 3284 | XLOC_016224 | chr16 | + | 2578994 | 2580106 | 2578994 | 2579096 | 2579475 | 379 | 2580106 | 3284 | 66,86 | 75,112 | 129,83 | 13,20 | 528 | 149 | 2.53E-11 | 8.60E-10 | 0.199,0.178 | 0.737,0.539 | -0.45 |

| | | | | | | | | | | | | | | | | | | | | | | |
|---|---|---|---|---|---|---|---|---|---|---|---|---|---|---|---|---|---|---|---|---|---|---|
| 943 | XLOC_023683 | chr2 | + | 152266431 | 152267041 | 152266431 | 152266547 | 152266927 | 380 | 152267041 | 943 | 123,100 | 15,11 | 164,132 | 5,9 | 529 | 149 | 0.00936635 | 0.039101674 | 0.698,0.719 | 0.902,0.805 | -0.145 |
| 1649 | XLOC_016761 | chr16 | - | 67272805 | 67273357 | 67272805 | 67272870 | 67273250 | 380 | 67273357 | 1649 | 1,11 | 85,88 | 47,17 | 77,41 | 529 | 149 | 1.64E-11 | 5.71E-10 | 0.003,0.034 | 0.147,0.105 | -0.107 |
| 2667 | XLOC_001160 | chr1 | - | 78412214 | 78414985 | 78412214 | 78414459 | 78414839 | 380 | 78414985 | 2667 | 672,575 | 833,725 | 13,111,020 | 501,329 | 529 | 149 | 0 | 0 | 0.185,0.183 | 0.424,0.466 | -0.261 |
| 2570 | XLOC_014712 | chr15 | + | 75644942 | 75646618 | 75644942 | 75646207 | 75646590 | 383 | 75646618 | 2570 | 68,92 | 6,6 | 125,93 | 0,4 | 532 | 149 | 0.005290623 | 0.02434652 | 0.76,0.811 | 1.0,0.867 | -0.148 |
| 1484 | XLOC_016684 | chr16 | - | 28890054 | 28891076 | 28890054 | 28890425 | 28890809 | 384 | 28891076 | 1484 | 26,13 | 3,5 | 4,7 | 1,0 | 533 | 149 | 0.006268529 | 0.02809598 | 0.708,0.421 | 0.528,1.0 | -0.2 |
| 1742 | XLOC_023552 | chr2 | + | 85582198 | 85583019 | 85582198 | 85582293 | 85582677 | 384 | 85583019 | 1742 | 39,28 | 10,6 | 45,45 | 3,1 | 533 | 149 | 0.000558435 | 0.003780633 | 0.522,0.566 | 0.807,0.926 | -0.323 |
| 12477 | XLOC_020941 | chr19 | + | 7695708 | 7696207 | 7695708 | 7695732 | 7696117 | 385 | 7696207 | 12477 | 57,41 | 62,31 | 161,93 | 18,5 | 534 | 149 | 1.11E-16 | 7.47E-15 | 0.204,0.27 | 0.714,0.838 | -0.539 |
| 6959 | XLOC_021314 | chr19 | + | 49999705 | 50000582 | 49999705 | 50000064 | 50000450 | 386 | 50000582 | 6959 | 42,44 | 51,57 | 92,76 | 15,12 | 535 | 149 | 3.33E-16 | 2.11E-14 | 0.187,0.177 | 0.631,0.638 | -0.453 |
| 1073 | XLOC_021549 | chr19 | - | 11456195 | 11456979 | 11456195 | 11456413 | 11456802 | 389 | 11456979 | 1073 | 76,80 | 66,72 | 114,88 | 41,32 | 538 | 149 | 1.00E-05 | 0.000114022 | 0.242,0.235 | 0.435,0.432 | -0.195 |
| 128 | XLOC_044112 | chr8 | + | 101225317 | 101226146 | 101225317 | 101225656 | 101226046 | 390 | 101226146 | 128 | 0,0 | 13,13 | 12,6 | 23,17 | 539 | 149 | 1.60E-05 | 0.00017054 | 0.0,0.0 | 0.126,0.089 | -0.107 |
| 14806 | XLOC_009117 | chr12 | + | 113725967 | 113726652 | 113725967 | 113726039 | 113726430 | 391 | 113726652 | 14806 | 56,50 | 5,0 | 51,46 | 0,0 | 540 | 149 | 0.005060068 | 0.023469075 | 0.756,1.0 | 1.0,1.0 | -0.122 |
| 5230 | XLOC_029675 | chr3 | + | 183897956 | 183898438 | 183897956 | 183898039 | 183898432 | 393 | 183898438 | 5230 | 195,182 | 0,13 | 362,292 | 0,0 | 542 | 149 | 1.20E-05 | 0.000133039 | 1.0,0.794 | 1.0,1.0 | -0.103 |
| 2215 | XLOC_009356 | chr12 | - | 50184821 | 50186371 | 50184821 | 50185834 | 50186228 | 394 | 50186371 | 2215 | 68,70 | 21,27 | 214,139 | 28,26 | 543 | 149 | 0.001132276 | 0.006781649 | 0.47,0.416 | 0.677,0.595 | -0.193 |
| 2152 | XLOC_018789 | chr17 | - | 80007793 | 80008431 | 80007793 | 80007882 | 80008277 | 395 | 80008431 | 2152 | 204,227 | 95,113 | 266,227 | 67,48 | 544 | 149 | 1.01E-05 | 0.000114811 | 0.37,0.355 | 0.521,0.564 | -0.18 |
| 3214 | XLOC_021536 | chr19 | - | 10464203 | 10464910 | 10464203 | 10464322 | 10464717 | 395 | 10464910 | 3214 | 142,128 | 38,36 | 161,115 | 12,13 | 544 | 149 | 1.54E-05 | 0.000165306 | 0.506,0.493 | 0.786,0.708 | -0.247 |
| 3900 | XLOC_045993 | chr9 | + | 104161797 | 104162292 | 104161797 | 104161812 | 104162207 | 395 | 104162292 | 3900 | 2,7 | 9,28 | 28,13 | 6,7 | 544 | 149 | 4.22E-07 | 6.80E-06 | 0.057,0.064 | 0.561,0.337 | -0.389 |
| 5478 | XLOC_028610 | chr22 | - | 26866684 | 26867475 | 26866684 | 26866779 | 26867174 | 395 | 26867475 | 5478 | 2,3 | 0,5 | 2,3 | 0,0 | 544 | 149 | 0.001245048 | 0.007329643 | 1.0,0.141 | 1.0,1.0 | -0.429 |
| 2736 | XLOC_005357 | chr10 | - | 103344358 | 103345235 | 103344358 | 103344676 | 103345072 | 396 | 103345235 | 2736 | 39,33 | 60,54 | 77,39 | 36,46 | 545 | 149 | 0.009092941 | 0.038130929 | 0.151,0.143 | 0.369,0.188 | -0.131 |
| 4114 | XLOC_044029 | chr8 | + | 48881829 | 48882617 | 48881829 | 48881961 | 48882357 | 396 | 48882617 | 4114 | 9,17 | 32,30 | 9,5 | 4,0 | 545 | 149 | 0.008439337 | 0.035965226 | 0.071,0.134 | 0.381,1.0 | -0.588 |
| 6697 | XLOC_016235 | chr16 | + | 2818505 | 2819285 | 2818505 | 2818600 | 2818997 | 397 | 2819285 | 6697 | 355,308 | 59,82 | 420,306 | 7,0 | 546 | 149 | 0 | 0 | 0.621,0.506 | 0.942,1.0 | -0.407 |
| 1283 | XLOC_018712 | chr17 | - | 73236422 | 73237138 | 73236422 | 73236493 | 73236892 | 399 | 73237138 | 1283 | 67,70 | 83,69 | 137,101 | 82,62 | 548 | 149 | 0.001082402 | 0.006521655 | 0.18,0.216 | 0.312,0.307 | -0.111 |
| 2412 | XLOC_007746 | chr11 | - | 123599833 | 123600533 | 123599833 | 123599922 | 123600322 | 400 | 123600533 | 2412 | 4,11 | 18,20 | 32,14 | 17,9 | 549 | 149 | 6.57E-05 | 0.000594139 | 0.057,0.13 | 0.338,0.297 | -0.224 |
| 699 | XLOC_028542 | chr22 | + | 46751340 | 46751969 | 46751340 | 46751485 | 46751886 | 401 | 46751969 | 699 | 316,282 | 125,125 | 693,530 | 118,95 | 550 | 149 | 1.57E-09 | 3.96E-08 | 0.406,0.379 | 0.614,0.602 | -0.215 |
| 5590 | XLOC_028387 | chr22 | + | 20073209 | 20074844 | 20073209 | 20074206 | 20074610 | 404 | 20074844 | 5590 | 27,21 | 0,4 | 12,8 | 0,0 | 553 | 149 | 0.008732311 | 0.036990529 | 1.0,0.586 | 1.0,1.0 | -0.207 |
| 1865 | XLOC_013325 | chr14 | - | 24034327 | 24034910 | 24034327 | 24034408 | 24034814 | 406 | 24034910 | 1865 | 31,40 | 27,45 | 67,48 | 23,12 | 555 | 149 | 8.89E-06 | 0.000102548 | 0.236,0.193 | 0.439,0.518 | -0.264 |
| 2540 | XLOC_007155 | chr11 | + | 65387224 | 65387900 | 65387224 | 65387403 | 65387809 | 406 | 65387900 | 2540 | 52,52 | 83,75 | 136,166 | 72,66 | 555 | 149 | 2.04E-10 | 5.93E-09 | 0.144,0.157 | 0.336,0.403 | -0.219 |
| 2834 | XLOC_041095 | chr7 | + | 44796023 | 44796748 | 44796023 | 44796138 | 44796545 | 407 | 44796748 | 2834 | 45,42 | 56,53 | 75,60 | 17,29 | 556 | 149 | 1.96E-05 | 0.000204788 | 0.177,0.175 | 0.542,0.357 | -0.274 |
| 8378 | XLOC_008936 | chr12 | + | 56226742 | 56227325 | 56226742 | 56226822 | 56227230 | 408 | 56227325 | 8378 | 17,24 | 3,3 | 28,7 | 0,0 | 557 | 149 | 2.95E-05 | 0.000294522 | 0.603,0.682 | 1.0,1.0 | -0.357 |
| 541 | XLOC_030127 | chr3 | - | 183701523 | 183705705 | 183701523 | 183702682 | 183703091 | 409 | 183705705 | 541 | 49,57 | 6,5 | 94,79 | 4,0 | 558 | 149 | 0.003873122 | 0.018894681 | 0.686,0.753 | 0.863,1.0 | -0.212 |
| 36 | XLOC_047663 | chrX | + | 23801916 | 23802520 | 23801916 | 23802000 | 23802410 | 410 | 23802520 | 36 | 13,451,265 | 73,97 | 35,722,886 | 83,80 | 559 | 149 | 3.81E-10 | 1.06E-08 | 0.831,0.777 | 0.92,0.906 | -0.109 |
| 227 | XLOC_041046 | chr7 | + | 27138692 | 27140058 | 27138692 | 27138985 | 27139397 | 412 | 27140058 | 227 | 22,21 | 33,26 | 75,38 | 20,15 | 561 | 149 | 6.09E-07 | 9.49E-06 | 0.15,0.177 | 0.499,0.402 | -0.287 |
| 834 | XLOC_016187 | chr16 | + | 705772 | 706537 | 705772 | 705889 | 706301 | 412 | 706537 | 834 | 221,213 | 52,66 | 219,191 | 12,3 | 561 | 149 | 4.53E-13 | 2.02E-11 | 0.53,0.462 | 0.829,0.944 | -0.39 |
| 125 | XLOC_044167 | chr8 | + | 144462234 | 144462953 | 144462234 | 144462345 | 144462758 | 413 | 144462953 | 125 | 234,215 | 38,25 | 123,75 | 3,1 | 562 | 149 | 4.81E-07 | 7.69E-06 | 0.62,0.695 | 0.916,0.952 | -0.276 |
| 1518 | XLOC_016471 | chr16 | + | 70501276 | 70501869 | 70501276 | 70501374 | 70501788 | 414 | 70501869 | 1518 | 12,2 | 15,19 | 26,20 | 24,15 | 563 | 149 | 0.005034674 | 0.023373344 | 0.175,0.027 | 0.223,0.261 | -0.141 |
| 1739 | XLOC_014554 | chr15 | + | 37100524 | 37102368 | 37100524 | 37100690 | 37101104 | 414 | 37102368 | 1739 | 38,60 | 3,3 | 45,43 | 0,0 | 563 | 149 | 0.000563851 | 0.003802897 | 0.77,0.841 | 1.0,1.0 | -0.195 |
| 7245 | XLOC_024366 | chr2 | - | 201778017 | 201778715 | 201778017 | 201778136 | 201778550 | 414 | 201778715 | 7245 | 14,38 | 6,0 | 25,14 | 1,0 | 563 | 149 | 0.010131892 | 0.041775237 | 0.382,1.0 | 0.869,1.0 | -0.243 |
| 2747 | XLOC_000004 | chr1 | + | 898716 | 899388 | 898716 | 898884 | 899299 | 415 | 899388 | 2747 | 87,70 | 37,41 | 117,103 | 27,33 | 564 | 149 | 0.00947444 | 0.039459606 | 0.383,0.311 | 0.534,0.452 | -0.146 |
| 6766 | XLOC_029383 | chr3 | + | 40498836 | 40499482 | 40498836 | 40498965 | 40499380 | 415 | 40499482 | 6766 | 10,28 | 163,174 | 19,242 | 146,148 | 564 | 149 | 2.33E-05 | 0.000239125 | 0.016,0.041 | 0.033,0.302 | -0.139 |
| 1936 | XLOC_013242 | chr14 | + | 94568159 | 94569277 | 94568159 | 94568405 | 94568822 | 417 | 94569277 | 1936 | 12,18 | 17,20 | 23,29 | 2,14 | 566 | 149 | 0.005362567 | 0.024653494 | 0.157,0.192 | 0.752,0.353 | -0.378 |
| 1118 | XLOC_044280 | chr8 | - | 38125888 | 38126508 | 38125888 | 38125979 | 38126399 | 420 | 38126508 | 1118 | 37,61 | 64,54 | 93,66 | 35,21 | 569 | 149 | 2.09E-07 | 3.58E-06 | 0.131,0.228 | 0.41,0.451 | -0.251 |
| 2598 | XLOC_007095 | chr11 | + | 60702723 | 60704627 | 60702723 | 60702915 | 60703335 | 420 | 60704627 | 2598 | 74,89 | 150,133 | 96,67 | 31,33 | 569 | 149 | 1.11E-11 | 3.96E-10 | 0.114,0.149 | 0.448,0.347 | -0.266 |
| 1528 | XLOC_016432 | chr16 | + | 66997393 | 66998413 | 66997393 | 66997838 | 66998259 | 421 | 66998413 | 1528 | 126,105 | 19,11 | 314,205 | 15,7 | 570 | 149 | 0.000289148 | 0.002146091 | 0.634,0.714 | 0.845,0.884 | -0.191 |
| 14917 | XLOC_000295 | chr1 | + | 52221991 | 52222487 | 52221991 | 52222030 | 52222452 | 422 | 52222487 | 14917 | 8,12 | 7,2 | 33,16 | 0,0 | 571 | 149 | 4.20E-08 | 8.18E-07 | 0.23,0.61 | 1.0,1.0 | -0.58 |
| 11686 | XLOC_000147 | chr1 | + | 27087346 | 27087964 | 27087346 | 27087446 | 27087874 | 428 | 27087964 | 11686 | 15,12 | 4,1 | 12,6 | 0,0 | 577 | 149 | 2.94E-05 | 0.000294392 | 0.492,0.756 | 1.0,1.0 | -0.376 |
| 3217 | XLOC_005275 | chr10 | - | 79579078 | 79579776 | 79579078 | 79579223 | 79579652 | 429 | 79579776 | 3217 | 61,34 | 123,112 | 71,54 | 79,49 | 578 | 149 | 8.45E-06 | 9.80E-05 | 0.113,0.073 | 0.188,0.221 | -0.112 |
| 2485 | XLOC_000189 | chr1 | + | 32670794 | 32671898 | 32670794 | 32671324 | 32671754 | 430 | 32671898 | 2485 | 22,13 | 29,19 | 37,30 | 14,20 | 579 | 149 | 0.002751592 | 0.014162073 | 0.163,0.15 | 0.405,0.279 | -0.186 |
| 2731 | XLOC_005357 | chr10 | - | 103332467 | 103340173 | 103332467 | 103339574 | 103340004 | 430 | 103340173 | 2731 | 54,38 | 88,94 | 88,71 | 67,44 | 579 | 149 | 1.90E-07 | 3.29E-06 | 0.136,0.094 | 0.253,0.293 | -0.158 |
| 463 | XLOC_035162 | chr5 | - | 176958282 | 176959201 | 176958282 | 176958522 | 176958953 | 431 | 176959201 | 463 | 25,46 | 24,37 | 36,37 | 16,7 | 580 | 149 | 0.006123488 | 0.027533276 | 0.211,0.242 | 0.366,0.576 | -0.244 |
| 1339 | XLOC_018272 | chr17 | + | 78194714 | 78195593 | 78194714 | 78194914 | 78195346 | 432 | 78195593 | 1339 | 74,67 | 52,40 | 57,47 | 5,6 | 581 | 149 | 8.61E-09 | 1.87E-07 | 0.267,0.3 | 0.745,0.668 | -0.423 |
| 2273 | XLOC_041587 | chr7 | - | 92206968 | 92207496 | 92206968 | 92207031 | 92207463 | 432 | 92207496 | 2273 | 22,37 | 59,78 | 96,69 | 84,63 | 581 | 149 | 4.98E-06 | 6.14E-05 | 0.087,0.108 | 0.227,0.219 | -0.126 |
| 2591 | XLOC_020956 | chr19 | + | 9451777 | 9454807 | 9451777 | 9451860 | 9452292 | 432 | 9454807 | 2591 | 6,9 | 5,16 | 14,7 | 0,2 | 581 | 149 | 0.000753965 | 0.004809508 | 0.235,0.126 | 1.0,0.473 | -0.556 |
| 3432 | XLOC_001116 | chr1 | - | 52823452 | 52824133 | 52823452 | 52823570 | 52824002 | 432 | 52824133 | 3432 | 29,39 | 58,51 | 64,33 | 47,22 | 581 | 149 | 0.000681493 | 0.004453826 | 0.114,0.164 | 0.259,0.278 | -0.13 |
| 1844 | XLOC_023861 | chr2 | + | 220110617 | 220111598 | 220110617 | 220110807 | 220111243 | 436 | 220111598 | 1844 | 40,28 | 2,2 | 34,19 | 0,0 | 585 | 149 | 0.004458488 | 0.021094547 | 0.836,0.781 | 1.0,1.0 | -0.192 |
| 1021 | XLOC_021769 | chr19 | - | 45853009 | 45854979 | 45853009 | 45854288 | 45854725 | 437 | 45854979 | 1021 | 212,226 | 12,10 | 395,409 | 1,9 | 586 | 149 | 0.000186308 | 0.001459361 | 0.818,0.852 | 0.99,0.92 | -0.12 |
| 10651 | XLOC_007597 | chr11 | - | 66132886 | 66133492 | 66132886 | 66132968 | 66133406 | 438 | 66133492 | 10651 | 43,27 | 12,16 | 56,71 | 5,1 | 587 | 149 | 1.09E-05 | 0.000122176 | 0.476,0.3 | 0.74,0.947 | -0.455 |
| 6957 | XLOC_021314 | chr19 | + | 49999705 | 50000582 | 49999705 | 50000011 | 50000450 | 439 | 50000582 | 6957 | 66,65 | 99,95 | 95,81 | 27,23 | 588 | 149 | 5.88E-15 | 3.32E-13 | 0.145,0.148 | 0.471,0.472 | -0.325 |
| 2259 | XLOC_014987 | chr15 | - | 75197004 | 75197572 | 75197004 | 75197053 | 75197494 | 441 | 75197572 | 2259 | 48,56 | 82,80 | 88,82 | 64,52 | 590 | 149 | 2.99E-05 | 0.000298279 | 0.129,0.15 | 0.258,0.285 | -0.132 |
| 3554 | XLOC_048040 | chrX | - | 102939608 | 102940106 | 102939608 | 102939657 | 102940098 | 441 | 102940106 | 3554 | 193,224 | 449,478 | 433,305 | 257,160 | 590 | 149 | 0 | 0 | 0.098,0.106 | 0.298,0.325 | -0.209 |
| 3193 | XLOC_000975 | chr1 | - | 28907071 | 28907741 | 28907071 | 28907158 | 28907605 | 447 | 28907741 | 3193 | 71,65 | 23,7 | 103,65 | 3,5 | 596 | 149 | 0.002644972 | 0.013703206 | 0.436,0.699 | 0.896,0.765 | -0.263 |
| 8019 | XLOC_017964 | chr17 | + | 19283094 | 19283814 | 19283094 | 19283260 | 19283707 | 447 | 19283814 | 8019 | 14,13 | 0,4 | 13,9 | 0,0 | 596 | 149 | 0.004360809 | 0.020722646 | 1.0,0.448 | 1.0,1.0 | -0.276 |
| 2798 | XLOC_007401 | chr11 | - | 6496969 | 6499122 | 6496969 | 6498498 | 6498947 | 449 | 6499122 | 2798 | 165,134 | 268,264 | 468,339 | 290,223 | 598 | 149 | 2.30E-14 | 1.18E-12 | 0.133,0.112 | 0.287,0.275 | -0.159 |
| 208 | XLOC_041204 | chr7 | + | 99158155 | 99174973 | 99158155 | 99169415 | 99169867 | 452 | 99174973 | 208 | 133,138 | 46,37 | 169,114 | 27,16 | 601 | 149 | 0.001667875 | 0.009392104 | 0.418,0.48 | 0.608,0.639 | -0.174 |
| 9816 | XLOC_013063 | chr14 | + | 24633271 | 24634164 | 24633271 | 24633343 | 24633795 | 452 | 24634164 | 9816 | 9,16 | 1,5 | 14,4 | 0,0 | 601 | 149 | 2.64E-06 | 3.48E-05 | 0.691,0.442 | 1.0,1.0 | -0.433 |
| 5626 | XLOC_027790 | chr21 | - | 38444733 | 38445459 | 38444733 | 38444837 | 38445290 | 453 | 38445459 | 5626 | 21,25 | 1,5 | 9,9 | 1,0 | 602 | 149 | 0.00677157 | 0.02993267 | 0.839,0.553 | 0.69,1.0 | -0.149 |
| 11248 | XLOC_015047 | chr15 | - | 89876326 | 89878071 | 89876326 | 89877144 | 89877597 | 453 | 89878071 | 11248 | 46,29 | 5,0 | 55,36 | 0,0 | 602 | 149 | 0.001908276 | 0.01044596 | 0.695,1.0 | 1.0,1.0 | -0.153 |

| | | | | | | | | | | | | | | | | | | | | | | |
|---|---|---|---|---|---|---|---|---|---|---|---|---|---|---|---|---|---|---|---|---|---|---|
| 2327 | XLOC_029351 | chr3 | + | 25832480 | 25833488 | 25832480 | 25832806 | 25833260 | 454 | 25833488 | 2327 | 158,160 | 10,9 | 274,185 | 3,2 | 603 | 149 | 8.68E-05 | 0.00075812 | 0.796,0.815 | 0.958,0.958 | -0.152 |
| 4193 | XLOC_041620 | chr7 | - | 99753293 | 99754172 | 99753293 | 99753554 | 99754008 | 454 | 99754172 | 4193 | 20,16 | 12,1 | 35,20 | 0,2 | 603 | 149 | 0.005478942 | 0.02511493 | 0.292,0.798 | 1.0,0.712 | -0.311 |
| 6648 | XLOC_026793 | chr20 | + | 42157296 | 42158008 | 42157296 | 42157385 | 42157842 | 457 | 42158008 | 6648 | 6,1 | 5,3 | 5,2 | 1,0 | 606 | 149 | 0.001304023 | 0.007632293 | 0.228,0.076 | 0.551,1.0 | -0.624 |
| 1863 | XLOC_013325 | chr14 | - | 24033770 | 24034910 | 24033770 | 24033869 | 24034327 | 458 | 24034910 | 1863 | 23,34 | 41,56 | 34,30 | 31,14 | 607 | 149 | 0.002842361 | 0.014549717 | 0.121,0.13 | 0.212,0.345 | -0.153 |
| 1379 | XLOC_014991 | chr15 | - | 75654982 | 75656529 | 75654982 | 75655089 | 75655550 | 461 | 75656529 | 1379 | 151,112 | 2,9 | 133,101 | 0,1 | 610 | 149 | 0.000266065 | 0.00199556 | 0.949,0.752 | 1.0,0.961 | -0.13 |
| 1386 | XLOC_017956 | chr17 | + | 18166399 | 18169893 | 18166399 | 18166562 | 18167023 | 461 | 18169893 | 1386 | 18,19 | 68,60 | 94,45 | 23,14 | 610 | 149 | 0 | 0 | 0.061,0.072 | 0.5,0.44 | -0.403 |
| 1953 | XLOC_018345 | chr17 | - | 3710040 | 3716573 | 3710040 | 3715948 | 3716410 | 462 | 3716573 | 1953 | 513,497 | 35,25 | 700,499 | 14,10 | 611 | 149 | 1.25E-05 | 0.000138209 | 0.781,0.829 | 0.924,0.924 | -0.119 |
| 8932 | XLOC_008774 | chr12 | + | 8932630 | 8933335 | 8932630 | 8932708 | 8933170 | 462 | 8933335 | 8932 | 55,91 | 21,6 | 258,207 | 16,2 | 611 | 149 | 0.006441284 | 0.028715357 | 0.39,0.787 | 0.797,0.962 | -0.291 |
| 9625 | XLOC_013470 | chr14 | - | 77270128 | 77270758 | 77270128 | 77270255 | 77270717 | 462 | 77270758 | 9625 | 5,6 | 2,2 | 8,1 | 0,0 | 611 | 149 | 2.81E-05 | 0.000282511 | 0.379,0.422 | 1.0,1.0 | -0.6 |
| 633 | XLOC_029385 | chr3 | + | 41280624 | 41282321 | 41280624 | 41280845 | 41281309 | 464 | 41282321 | 633 | 1,001,938 | 122,129 | 20,051,673 | 57,37 | 613 | 149 | 0 | 0 | 0.666,0.639 | 0.895,0.917 | -0.253 |
| 2318 | XLOC_021921 | chr19 | - | 59080882 | 59082796 | 59080882 | 59081894 | 59082360 | 466 | 59082796 | 2318 | 204,270 | 63,56 | 258,244 | 19,12 | 615 | 149 | 1.60E-09 | 4.03E-08 | 0.44,0.539 | 0.767,0.831 | -0.309 |
| 693 | XLOC_028585 | chr22 | - | 21354935 | 21355700 | 21354935 | 21355076 | 21355544 | 468 | 21355700 | 693 | 26,28 | 87,96 | 83,50 | 91,64 | 617 | 149 | 5.57E-07 | 8.76E-06 | 0.067,0.066 | 0.181,0.159 | -0.103 |
| 2787 | XLOC_005240 | chr10 | - | 69832639 | 69835043 | 69832639 | 69832943 | 69833413 | 470 | 69835043 | 2787 | 98,98 | 275,253 | 162,132 | 150,107 | 619 | 149 | 3.18E-13 | 1.44E-11 | 0.079,0.085 | 0.206,0.229 | -0.136 |
| 2504 | XLOC_007262 | chr11 | + | 94799812 | 94804995 | 94799812 | 94801346 | 94801817 | 471 | 94804995 | 2504 | 243,283 | 31,45 | 266,230 | 8,5 | 620 | 149 | 5.97E-10 | 1.62E-08 | 0.653,0.602 | 0.889,0.917 | -0.276 |
| 8757 | XLOC_016177 | chr16 | + | 456714 | 457560 | 456714 | 456952 | 457424 | 472 | 457560 | 8757 | 10,25 | 2,2 | 11,8 | 0,0 | 621 | 149 | 0.001355731 | 0.007882943 | 0.545,0.75 | 1.0,1.0 | -0.353 |
| 938 | XLOC_023708 | chr2 | + | 170550968 | 170551757 | 170550968 | 170551251 | 170551724 | 473 | 170551757 | 938 | 3,5 | 4,7 | 23,13 | 3,0 | 622 | 149 | 3.25E-05 | 0.000321833 | 0.152,0.146 | 0.647,1.0 | -0.674 |
| 489 | XLOC_032538 | chr4 | - | 1719242 | 1720353 | 1719242 | 1719402 | 1719878 | 476 | 1720353 | 489 | 82,64 | 64,46 | 66,56 | 22,14 | 625 | 149 | 0.00013826 | 0.001128046 | 0.234,0.249 | 0.417,0.488 | -0.211 |
| 2697 | XLOC_008906 | chr12 | + | 53579170 | 53579861 | 53579170 | 53579260 | 53579737 | 477 | 53579861 | 2697 | 30,26 | 226,209 | 137,99 | 213,129 | 626 | 149 | 0 | 0 | 0.031,0.029 | 0.133,0.154 | -0.114 |
| 984 | XLOC_016192 | chr16 | + | 841160 | 842347 | 841160 | 841370 | 841850 | 480 | 842347 | 984 | 312,366 | 87,62 | 368,275 | 35,19 | 629 | 149 | 1.55E-05 | 0.000165988 | 0.459,0.583 | 0.714,0.774 | -0.223 |
| 2457 | XLOC_007532 | chr11 | - | 61187941 | 61189080 | 61187941 | 61188045 | 61188525 | 480 | 61189080 | 2457 | 120,136 | 11,6 | 179,159 | 2,0 | 629 | 149 | 3.53E-06 | 4.49E-05 | 0.721,0.843 | 0.955,1.0 | -0.196 |
| 2009 | XLOC_000083 | chr1 | + | 12089279 | 12089951 | 12089279 | 12089338 | 12089821 | 483 | 12089951 | 2009 | 96,89 | 63,47 | 185,118 | 26,31 | 632 | 149 | 1.40E-05 | 0.000153036 | 0.264,0.309 | 0.627,0.473 | -0.264 |
| 103 | XLOC_044433 | chr8 | - | 144363051 | 144363914 | 144363051 | 144363167 | 144363651 | 484 | 144363914 | 103 | 92,90 | 59,70 | 117,83 | 42,29 | 633 | 149 | 0.00069816 | 0.00453965 | 0.268,0.232 | 0.396,0.403 | -0.15 |
| 1899 | XLOC_013159 | chr14 | + | 65204073 | 65204942 | 65204073 | 65204093 | 65204579 | 486 | 65204942 | 1899 | 59,79 | 28,44 | 67,17 | 10,5 | 635 | 149 | 0.005490305 | 0.025158849 | 0.331,0.296 | 0.611,0.444 | -0.214 |
| 12999 | XLOC_014788 | chr15 | + | 99500289 | 99507746 | 99500289 | 99500705 | 99501191 | 486 | 99507746 | 12999 | 81,89 | 10,15 | 142,107 | 5,2 | 635 | 149 | 2.20E-05 | 0.000226607 | 0.655,0.582 | 0.87,0.926 | -0.279 |
| 2872 | XLOC_000563 | chr1 | + | 156703168 | 156704285 | 156703168 | 156703312 | 156703800 | 488 | 156704285 | 2872 | 44,39 | 50,57 | 37,30 | 21,18 | 637 | 149 | 0.003267416 | 0.016416121 | 0.171,0.138 | 0.292,0.28 | -0.132 |
| 6339 | XLOC_023924 | chr2 | - | 38828 | 42305 | 38828 | 41627 | 42117 | 490 | 42305 | 6339 | 5,4 | 3,2 | 2,7 | 0,0 | 639 | 149 | 9.64E-07 | 1.42E-05 | 0.28,0.318 | 1.0,1.0 | -0.701 |
| 1743 | XLOC_023552 | chr2 | + | 85617260 | 85618877 | 85617260 | 85617388 | 85617882 | 494 | 85618877 | 1743 | 62,57 | 21,23 | 63,48 | 9,6 | 643 | 149 | 0.001617108 | 0.009153675 | 0.406,0.365 | 0.619,0.65 | -0.249 |
| 6831 | XLOC_021591 | chr19 | - | 15483661 | 15484876 | 15483661 | 15484151 | 15484645 | 494 | 15484876 | 6831 | 26,31 | 1,3 | 30,32 | 0,0 | 643 | 149 | 0.004421694 | 0.020955567 | 0.858,0.705 | 1.0,1.0 | -0.219 |
| 722 | XLOC_027814 | chr21 | - | 45748064 | 45750802 | 45748064 | 45750209 | 45750705 | 496 | 45750802 | 722 | 30,34 | 39,45 | 55,82 | 10,27 | 645 | 149 | 6.54E-08 | 1.24E-06 | 0.151,0.149 | 0.56,0.412 | -0.336 |
| 7309 | XLOC_020063 | chr18 | + | 21406665 | 21407431 | 21406665 | 21406781 | 21407277 | 496 | 21407431 | 7309 | 17,18 | 3,6 | 16,8 | 0,1 | 645 | 149 | 0.00073148 | 0.004693637 | 0.567,0.409 | 1.0,0.649 | -0.337 |
| 2295 | XLOC_046145 | chr9 | + | 139753660 | 139755286 | 139753660 | 139754516 | 139755013 | 497 | 139755286 | 2295 | 5,9 | 18,22 | 12,11 | 4,9 | 646 | 149 | 0.001088371 | 0.006552031 | 0.06,0.086 | 0.409,0.22 | -0.241 |
| 1276 | XLOC_018746 | chr17 | - | 74729864 | 74731240 | 74729864 | 74730644 | 74731142 | 498 | 74731240 | 1276 | 40,703,869 | 191,196 | 53,004,298 | 72,41 | 647 | 149 | 0 | 0 | 0.831,0.82 | 0.944,0.96 | -0.127 |
| 3185 | XLOC_007240 | chr11 | + | 82783664 | 82808540 | 82783664 | 82783846 | 82784344 | 498 | 82808540 | 3185 | 70,72 | 95,101 | 99,66 | 36,22 | 647 | 149 | 4.86E-11 | 1.57E-09 | 0.145,0.141 | 0.388,0.409 | -0.255 |
| 1896 | XLOC_021723 | chr19 | - | 40596072 | 40596814 | 40596072 | 40596142 | 40596642 | 500 | 40596814 | 1896 | 77,83 | 1,7 | 45,57 | 0,0 | 649 | 149 | 0.000222171 | 0.001698822 | 0.946,0.731 | 1.0,1.0 | -0.161 |
| 7 | XLOC_048071 | chrX | - | 122756955 | 122757822 | 122756955 | 122757134 | 122757637 | 503 | 122757822 | 7 | 227,233 | 164,183 | 410,319 | 137,93 | 652 | 149 | 1.03E-09 | 2.67E-08 | 0.24,0.225 | 0.406,0.439 | -0.19 |
| 630 | XLOC_029390 | chr3 | + | 42684830 | 42685495 | 42684830 | 42684887 | 42685390 | 503 | 42685495 | 630 | 57,55 | 62,46 | 141,132 | 63,70 | 652 | 149 | 0.000479026 | 0.003316092 | 0.174,0.215 | 0.338,0.301 | -0.125 |
| 2368 | XLOC_016732 | chr16 | - | 54315815 | 54318224 | 54315815 | 54317652 | 54318157 | 505 | 54318224 | 2368 | 100,120 | 67,79 | 188,179 | 54,47 | 654 | 149 | 1.33E-06 | 1.90E-05 | 0.254,0.257 | 0.442,0.465 | -0.198 |
| 631 | XLOC_029390 | chr3 | + | 42684830 | 42685495 | 42684830 | 42684887 | 42685393 | 506 | 42685495 | 631 | 57,55 | 61,61 | 141,132 | 81,43 | 655 | 149 | 8.67E-05 | 0.000758023 | 0.175,0.17 | 0.284,0.411 | -0.175 |
| 1218 | XLOC_037628 | chr6 | + | 4060646 | 4061934 | 4060646 | 4061381 | 4061888 | 507 | 4061934 | 1218 | 520,574 | 71,82 | 934,721 | 76,49 | 656 | 149 | 0.000113563 | 0.00095467 | 0.625,0.614 | 0.736,0.77 | -0.134 |
| 240 | XLOC_041771 | chr7 | - | 150776586 | 150778987 | 150776586 | 150777009 | 150777523 | 514 | 150778987 | 240 | 315,252 | 90,95 | 276,220 | 61,44 | 663 | 149 | 0.009876842 | 0.040954385 | 0.44,0.373 | 0.504,0.529 | -0.11 |
| 296 | XLOC_038044 | chr6 | + | 157800198 | 157809091 | 157800198 | 157803298 | 157803814 | 516 | 157809091 | 296 | 44,70 | 5,0 | 30,18 | 0,0 | 665 | 149 | 0.003427854 | 0.017118666 | 0.663,1.0 | 1.0,1.0 | -0.168 |
| 2282 | XLOC_018441 | chr17 | - | 17715401 | 17716793 | 17715401 | 17716165 | 17716681 | 516 | 17716793 | 2282 | 213,219 | 457,474 | 466,339 | 192,136 | 665 | 149 | 0 | 0 | 0.095,0.094 | 0.352,0.358 | -0.26 |
| 5864 | XLOC_026782 | chr20 | + | 37075330 | 37076266 | 37075330 | 37075591 | 37076108 | 517 | 37076266 | 5864 | 30,21 | 3,2 | 32,19 | 0,0 | 666 | 149 | 0.000276013 | 0.002060321 | 0.691,0.701 | 1.0,1.0 | -0.304 |
| 3201 | XLOC_032226 | chr4 | + | 1243043 | 1246399 | 1243043 | 1244161 | 1244683 | 522 | 1246399 | 3201 | 102,124 | 26,21 | 112,102 | 8,1 | 671 | 149 | 4.67E-05 | 0.000443428 | 0.466,0.567 | 0.757,0.958 | -0.341 |
| 3327 | XLOC_000753 | chr1 | + | 226032198 | 226033232 | 226032198 | 226032324 | 226032846 | 522 | 226033232 | 3327 | 153,171 | 200,188 | 294,208 | 93,57 | 671 | 149 | 0 | 0 | 0.145,0.168 | 0.412,0.448 | -0.273 |
| 5627 | XLOC_027790 | chr21 | - | 38444733 | 38445459 | 38444733 | 38444837 | 38445360 | 523 | 38445459 | 5627 | 42,33 | 6,11 | 13,24 | 0,0 | 672 | 149 | 1.04E-06 | 1.52E-05 | 0.608,0.399 | 1.0,1.0 | -0.496 |
| 527 | XLOC_037824 | chr6 | + | 41751199 | 41751976 | 41751199 | 41751343 | 41751868 | 525 | 41751976 | 527 | 137,158 | 23,22 | 57,48 | 1,3 | 674 | 149 | 0.001177557 | 0.006996484 | 0.568,0.614 | 0.926,0.78 | -0.262 |
| 919 | XLOC_023823 | chr2 | + | 209168894 | 209169737 | 209168894 | 209169042 | 209169569 | 527 | 209169737 | 919 | 31,24 | 16,25 | 67,64 | 17,23 | 676 | 149 | 0.00426282 | 0.020339124 | 0.299,0.175 | 0.465,0.38 | -0.185 |
| 8471 | XLOC_016560 | chr16 | - | 775965 | 776763 | 775965 | 776085 | 776614 | 529 | 776763 | 8471 | 4,7 | 0,4 | 13,6 | 0,0 | 678 | 149 | 0.002515306 | 0.013147175 | 1.0,0.278 | 1.0,1.0 | -0.361 |
| 196 | XLOC_041279 | chr7 | + | 120595612 | 120597151 | 120595612 | 120595678 | 120596213 | 535 | 120597151 | 196 | 106,114 | 7,4 | 48,46 | 0,0 | 684 | 149 | 7.54E-05 | 0.000668199 | 0.767,0.861 | 1.0,1.0 | -0.186 |
| 9036 | XLOC_015039 | chr15 | - | 85070011 | 85070657 | 85070011 | 85070107 | 85070642 | 535 | 85070657 | 9036 | 37,25 | 7,1 | 17,27 | 0,0 | 684 | 149 | 1.40E-05 | 0.000153262 | 0.535,0.845 | 1.0,1.0 | -0.31 |
| 9380 | XLOC_018149 | chr17 | + | 48456812 | 48457862 | 48456812 | 48457137 | 48457672 | 535 | 48457862 | 9380 | 29,36 | 1,4 | 37,35 | 0,0 | 684 | 149 | 0.000350325 | 0.002530763 | 0.863,0.662 | 1.0,1.0 | -0.238 |
| 110 | XLOC_047939 | chrX | - | 46458596 | 46466638 | 46458596 | 46465805 | 46466342 | 537 | 46466638 | 110 | 33,57 | 12,19 | 130,79 | 5,1 | 686 | 149 | 3.02E-10 | 8.55E-09 | 0.374,0.395 | 0.85,0.945 | -0.513 |
| 2061 | XLOC_023860 | chr2 | + | 220098854 | 220100034 | 220098854 | 220099010 | 220099547 | 537 | 220100034 | 2061 | 457,468 | 202,230 | 287,290 | 105,60 | 686 | 149 | 0.010816169 | 0.044149349 | 0.329,0.306 | 0.373,0.512 | -0.125 |
| 509 | XLOC_032278 | chr4 | + | 15687858 | 15692034 | 15687858 | 15688085 | 15688623 | 538 | 15692034 | 509 | 265,246 | 25,23 | 204,149 | 8,4 | 687 | 149 | 0.00078556 | 0.004981795 | 0.697,0.699 | 0.847,0.89 | -0.171 |
| 4381 | XLOC_041046 | chr7 | + | 27138692 | 27140058 | 27138692 | 27138858 | 27139397 | 539 | 27140058 | 4381 | 14,13 | 4,4 | 42,24 | 3,1 | 688 | 149 | 0.005718009 | 0.02596638 | 0.431,0.413 | 0.752,0.839 | -0.373 |
| 90 | XLOC_046188 | chr9 | - | 14611045 | 14619680 | 14611045 | 14619096 | 14619636 | 540 | 14619680 | 90 | 1,0 | 18,14 | 10,6 | 17,8 | 689 | 149 | 1.19E-05 | 0.000133003 | 0.012,0.0 | 0.113,0.14 | -0.12 |
| 2258 | XLOC_014987 | chr15 | - | 75190467 | 75195124 | 75190467 | 75194404 | 75194944 | 540 | 75195124 | 2258 | 319,293 | 18,39 | 494,353 | 17,7 | 689 | 149 | 0.00017502 | 0.001381671 | 0.793,0.619 | 0.863,0.916 | -0.183 |
| 1949 | XLOC_016854 | chr16 | - | 88799014 | 88799862 | 88799014 | 88799140 | 88799685 | 545 | 88799862 | 1949 | 449,442 | 643,590 | 868,631 | 441,314 | 694 | 149 | 0 | 0 | 0.13,0.139 | 0.297,0.301 | -0.164 |
| 3200 | XLOC_032226 | chr4 | + | 1243043 | 1246399 | 1243043 | 1244138 | 1244683 | 545 | 1246399 | 3200 | 110,138 | 9,10 | 117,105 | 0,2 | 694 | 149 | 1.38E-05 | 0.000150703 | 0.724,0.748 | 1.0,0.919 | -0.224 |
| 10245 | XLOC_018507 | chr17 | - | 34868716 | 34869348 | 34868716 | 34868726 | 34869271 | 545 | 34869348 | 10245 | 57,63 | 3,3 | 98,52 | 0,0 | 694 | 149 | 0.000398393 | 0.002830232 | 0.803,0.818 | 1.0,1.0 | -0.19 |
| 175 | XLOC_041435 | chr7 | - | 27168126 | 27181704 | 27168126 | 27169190 | 27169736 | 546 | 27181704 | 175 | 0,0 | 6,3 | 5,2 | 2,2 | 695 | 149 | 0.000124879 | 0.001036844 | 0.0,0.0 | 0.349,0.177 | -0.263 |
| 1414 | XLOC_018103 | chr17 | + | 43212707 | 43213987 | 43212707 | 43213320 | 43213866 | 546 | 43213987 | 1414 | 70,62 | 6,4 | 102,84 | 0,1 | 695 | 149 | 1.99E-05 | 0.000207436 | 0.714,0.769 | 1.0,0.947 | -0.232 |
| 2652 | XLOC_014820 | chr15 | - | 32825412 | 32828263 | 32825412 | 32827577 | 32828124 | 547 | 32828263 | 2652 | 2,3 | 1,6 | 11,6 | 0,0 | 696 | 149 | 1.05E-11 | 3.77E-10 | 0.3,0.097 | 1.0,1.0 | -0.801 |
| 4 | XLOC_048096 | chrX | - | 134184969 | 134186258 | 134184969 | 134185398 | 134185946 | 548 | 134186258 | 4 | 596,645 | 422,374 | 14,131,064 | 251,181 | 697 | 149 | 0 | 0 | 0.232,0.269 | 0.546,0.557 | -0.301 |

| | | | | | | | | | | | | | | | | | | | | | | |
|---|---|---|---|---|---|---|---|---|---|---|---|---|---|---|---|---|---|---|---|---|---|---|
| 2932 | XLOC_046478 | chr9 | - | 139270066 | 139270933 | 139270066 | 139270252 | 139270800 | 548 | 139270933 | 2932 | 20,24 | 66,47 | 74,31 | 38,59 | 697 | 149 | 0.001954062 | 0.010659412 | 0.061,0.098 | 0.294,0.101 | -0.118 |
| 3446 | XLOC_030013 | chr3 | - | 125725011 | 125726068 | 125725011 | 125725407 | 125725956 | 549 | 125726068 | 3446 | 807,738 | 39,25 | 976,751 | 9,13 | 698 | 149 | 2.40E-06 | 3.20E-05 | 0.815,0.863 | 0.959,0.925 | -0.103 |
| 3112 | XLOC_001309 | chr1 | - | 151027491 | 151028469 | 151027491 | 151027602 | 151028152 | 550 | 151028469 | 3112 | 216,217 | 274,258 | 427,276 | 238,184 | 699 | 149 | 1.28E-07 | 2.30E-06 | 0.144,0.152 | 0.277,0.242 | -0.112 |
| 382 | XLOC_034900 | chr5 | - | 54518003 | 54518851 | 54518003 | 54518227 | 54518778 | 551 | 54518851 | 382 | 15,20 | 15,11 | 31,22 | 0,2 | 700 | 149 | 8.93E-07 | 1.33E-05 | 0.176,0.279 | 1.0,0.701 | -0.623 |
| 7490 | XLOC_020853 | chr19 | + | 990222 | 990995 | 990222 | 990364 | 990915 | 551 | 990995 | 7490 | 63,65 | 5,7 | 106,85 | 4,1 | 700 | 149 | 0.008509526 | 0.036194476 | 0.728,0.664 | 0.849,0.948 | -0.203 |
| 8795 | XLOC_016339 | chr16 | + | 28847253 | 28848525 | 28847253 | 28847497 | 28848048 | 551 | 28848525 | 8795 | 369,327 | 27,15 | 383,299 | 12,7 | 700 | 149 | 0.006658301 | 0.029505865 | 0.744,0.823 | 0.872,0.901 | -0.103 |
| 8599 | XLOC_021326 | chr19 | + | 50381754 | 50382717 | 50381754 | 50381829 | 50382382 | 553 | 50382717 | 8599 | 47,50 | 3,2 | 37,13 | 0,0 | 702 | 149 | 0.001848627 | 0.010210463 | 0.769,0.841 | 1.0,1.0 | -0.195 |
| 3166 | XLOC_017914 | chr17 | + | 7761923 | 7768020 | 7761923 | 7762142 | 7762699 | 557 | 7768020 | 3166 | 61,78 | 84,91 | 212,150 | 81,49 | 706 | 149 | 2.37E-12 | 9.55E-11 | 0.133,0.153 | 0.356,0.392 | -0.231 |
| 2335 | XLOC_005106 | chr10 | + | 135215032 | 135215749 | 135215032 | 135215094 | 135215652 | 558 | 135215749 | 2335 | 209,175 | 92,80 | 174,135 | 55,25 | 707 | 149 | 0.010018962 | 0.041458482 | 0.324,0.316 | 0.4,0.532 | -0.146 |
| 1771 | XLOC_000505 | chr1 | + | 150965676 | 150969350 | 150965676 | 150967175 | 150967735 | 560 | 150969350 | 1771 | 50,31 | 16,20 | 26,16 | 2,3 | 709 | 149 | 0.012390115 | 0.049683589 | 0.396,0.246 | 0.732,0.528 | -0.309 |
| 11249 | XLOC_015047 | chr15 | - | 89876326 | 89878071 | 89876326 | 89877144 | 89877705 | 561 | 89878071 | 11249 | 46,29 | 3,5 | 55,36 | 0,0 | 710 | 149 | 3.68E-06 | 4.66E-05 | 0.763,0.549 | 1.0,1.0 | -0.344 |
| 2753 | XLOC_014776 | chr15 | + | 91504254 | 91506740 | 91504254 | 91504284 | 91504847 | 563 | 91506740 | 2753 | 686,724 | 351,322 | 522,390 | 138,110 | 712 | 149 | 2.00E-05 | 0.000208401 | 0.29,0.32 | 0.442,0.426 | -0.129 |
| 5037 | XLOC_034483 | chr5 | + | 917122 | 918140 | 917122 | 917266 | 917833 | 567 | 918140 | 5037 | 9,431,033 | 72,71 | 17,031,408 | 20,23 | 716 | 149 | 0 | 0 | 0.732,0.752 | 0.947,0.927 | -0.195 |
| 314 | XLOC_037801 | chr6 | + | 35262232 | 35263816 | 35262232 | 35262346 | 35262920 | 574 | 35263816 | 314 | 85,88 | 161,130 | 116,67 | 49,28 | 723 | 149 | 4.83E-13 | 2.14E-11 | 0.098,0.122 | 0.328,0.33 | -0.219 |
| 902 | XLOC_023985 | chr2 | - | 27883850 | 27885148 | 27883850 | 27884468 | 27885045 | 577 | 27885148 | 902 | 387,387 | 9,17 | 366,352 | 5,1 | 726 | 149 | 0.000281676 | 0.002095951 | 0.898,0.824 | 0.938,0.986 | -0.101 |
| 2123 | XLOC_028670 | chr22 | - | 38694099 | 38694939 | 38694099 | 38694209 | 38694790 | 581 | 38694939 | 2123 | 13,171,464 | 133,159 | 19,361,405 | 55,36 | 730 | 149 | 0 | 0 | 0.669,0.653 | 0.878,0.888 | -0.222 |
| 431 | XLOC_034594 | chr5 | + | 76378272 | 76383444 | 76378272 | 76382729 | 76383311 | 582 | 76383444 | 431 | 31,30 | 4,3 | 26,13 | 0,0 | 731 | 149 | 1.41E-05 | 0.000153473 | 0.612,0.671 | 1.0,1.0 | -0.359 |
| 1269 | XLOC_018771 | chr17 | - | 79225292 | 79226483 | 79225292 | 79225412 | 79225994 | 582 | 79226483 | 1269 | 125,189 | 192,151 | 409,261 | 118,67 | 731 | 149 | 0 | 0 | 0.117,0.203 | 0.414,0.443 | -0.268 |
| 2937 | XLOC_000699 | chr1 | + | 207940357 | 207941168 | 207940357 | 207940540 | 207941123 | 583 | 207941168 | 2937 | 12,181,293 | 279,363 | 56,924,016 | 501,395 | 732 | 149 | 0 | 0 | 0.471,0.42 | 0.698,0.674 | -0.24 |
| 14401 | XLOC_001334 | chr1 | - | 153905958 | 153906778 | 153905958 | 153906023 | 153906610 | 587 | 153906778 | 14401 | 109,88 | 8,0 | 154,161 | 1,1 | 736 | 149 | 0.005644066 | 0.025713275 | 0.734,1.0 | 0.969,0.97 | -0.103 |
| 8756 | XLOC_016177 | chr16 | + | 456714 | 457560 | 456714 | 456833 | 457424 | 591 | 457560 | 8756 | 10,25 | 1,6 | 11,8 | 0,0 | 740 | 149 | 8.57E-07 | 1.28E-05 | 0.668,0.456 | 1.0,1.0 | -0.438 |
| 937 | XLOC_023708 | chr2 | + | 170550968 | 170551757 | 170550968 | 170551132 | 170551724 | 592 | 170551757 | 937 | 9,15 | 11,9 | 43,16 | 8,3 | 741 | 149 | 0.000554525 | 0.003761552 | 0.141,0.251 | 0.519,0.517 | -0.322 |
| 2715 | XLOC_021677 | chr19 | - | 36387814 | 36388758 | 36387814 | 36387960 | 36388552 | 592 | 36388758 | 2715 | 6,6 | 27,13 | 38,7 | 21,12 | 741 | 149 | 0.000625214 | 0.004147442 | 0.043,0.085 | 0.267,0.105 | -0.122 |
| 4545 | XLOC_037885 | chr6 | + | 64286340 | 64287727 | 64286340 | 64286890 | 64287482 | 592 | 64287727 | 4545 | 32,34 | 6,2 | 133,121 | 0,0 | 741 | 149 | 4.06E-07 | 6.60E-06 | 0.517,0.774 | 1.0,1.0 | -0.355 |
| 2849 | XLOC_020911 | chr19 | + | 4501202 | 4502209 | 4501202 | 4501329 | 4501922 | 593 | 4502209 | 2849 | 49,62 | 134,140 | 144,111 | 88,71 | 742 | 149 | 9.55E-15 | 5.29E-13 | 0.068,0.082 | 0.247,0.239 | -0.168 |
| 4758 | XLOC_034801 | chr5 | + | 176083700 | 176086189 | 176083700 | 176083925 | 176084518 | 593 | 176086189 | 4758 | 95,107 | 2,7 | 49,11 | 0,0 | 742 | 149 | 0.000241969 | 0.001829404 | 0.905,0.754 | 1.0,1.0 | -0.17 |
| 1567 | XLOC_016182 | chr16 | + | 630182 | 630972 | 630182 | 630263 | 630857 | 594 | 630972 | 1567 | 31,30 | 30,37 | 46,49 | 6,0 | 743 | 149 | 2.39E-07 | 4.04E-06 | 0.172,0.14 | 0.606,1.0 | -0.647 |
| 629 | XLOC_029390 | chr3 | + | 42661155 | 42672808 | 42661155 | 42672067 | 42672662 | 595 | 42672808 | 629 | 136,128 | 64,73 | 214,164 | 72,42 | 744 | 149 | 0.002284533 | 0.012143191 | 0.299,0.26 | 0.373,0.439 | -0.127 |
| 2850 | XLOC_020911 | chr19 | + | 4501202 | 4502209 | 4501202 | 4501329 | 4501925 | 596 | 4502209 | 2850 | 50,62 | 42,48 | 144,111 | 16,25 | 745 | 149 | 4.08E-08 | 7.96E-07 | 0.192,0.205 | 0.643,0.47 | -0.358 |
| 13761 | XLOC_000428 | chr1 | + | 113499050 | 113500344 | 113499050 | 113499578 | 113500180 | 602 | 113500344 | 13761 | 16,25 | 15,17 | 16,15 | 3,0 | 751 | 149 | 0.002385828 | 0.012565957 | 0.175,0.226 | 0.514,1.0 | -0.556 |
| 1510 | XLOC_016525 | chr16 | + | 88740011 | 88741311 | 88740011 | 88740167 | 88740771 | 604 | 88741311 | 1510 | 6,9 | 4,0 | 13,13 | 0,0 | 753 | 149 | 0.002742997 | 0.014133289 | 0.229,1.0 | 1.0,1.0 | -0.385 |
| 151 | XLOC_041655 | chr7 | - | 102918060 | 102918793 | 102918060 | 102918085 | 102918691 | 606 | 102918793 | 151 | 50,49 | 5,5 | 21,19 | 0,0 | 755 | 149 | 1.83E-05 | 0.000192183 | 0.664,0.659 | 1.0,1.0 | -0.339 |
| 3354 | XLOC_021131 | chr19 | + | 35223593 | 35225951 | 35223593 | 35225265 | 35225874 | 609 | 35225951 | 3354 | 47,69 | 12,11 | 45,36 | 3,2 | 758 | 149 | 0.009079799 | 0.038096853 | 0.435,0.552 | 0.747,0.78 | -0.27 |
| 3209 | XLOC_029851 | chr3 | - | 49881233 | 49882013 | 49881233 | 49881361 | 49881973 | 612 | 49882013 | 3209 | 74,49 | 38,47 | 119,78 | 49,21 | 761 | 149 | 0.006232125 | 0.027959435 | 0.276,0.17 | 0.322,0.421 | -0.148 |
| 1277 | XLOC_018746 | chr17 | - | 74731142 | 74732546 | 74731142 | 74731240 | 74731853 | 613 | 74732546 | 1277 | 20,802,141 | 279,254 | 32,662,627 | 160,172 | 762 | 149 | 7.60E-11 | 2.36E-09 | 0.593,0.622 | 0.8,0.749 | -0.167 |
| 1302 | XLOC_021006 | chr19 | + | 14029942 | 14030752 | 14029942 | 14030015 | 14030630 | 615 | 14030752 | 1302 | 70,47 | 63,70 | 183,102 | 48,37 | 764 | 149 | 9.45E-11 | 2.89E-09 | 0.178,0.116 | 0.426,0.35 | -0.241 |
| 334 | XLOC_037632 | chr6 | + | 7181356 | 7182835 | 7181356 | 7181479 | 7182102 | 623 | 7182835 | 334 | 75,90 | 56,72 | 249,212 | 79,76 | 772 | 149 | 2.92E-06 | 3.81E-05 | 0.205,0.194 | 0.378,0.35 | -0.164 |
| 353 | XLOC_035165 | chr5 | - | 177651642 | 177652430 | 177651642 | 177651730 | 177652355 | 625 | 177652430 | 353 | 62,60 | 102,74 | 49,35 | 34,23 | 774 | 149 | 0.001392564 | 0.008063627 | 0.105,0.135 | 0.217,0.227 | -0.102 |
| 3173 | XLOC_021620 | chr19 | - | 19101568 | 19105295 | 19101568 | 19104549 | 19105174 | 625 | 19105295 | 3173 | 505,497 | 134,157 | 899,747 | 88,89 | 774 | 149 | 3.80E-12 | 1.46E-10 | 0.42,0.379 | 0.663,0.618 | -0.241 |
| 1519 | XLOC_021375 | chr19 | + | 55997027 | 55998935 | 55997027 | 55997147 | 55997774 | 627 | 55998935 | 1519 | 55,48 | 121,141 | 173,114 | 26,35 | 776 | 149 | 0 | 0 | 0.08,0.061 | 0.561,0.385 | -0.403 |
| 12224 | XLOC_007095 | chr11 | + | 60701013 | 60702236 | 60701013 | 60701216 | 60701843 | 627 | 60702236 | 12224 | 4,7 | 0,3 | 21,11 | 0,0 | 776 | 149 | 0.011083494 | 0.045071172 | 1.0,0.309 | 1.0,1.0 | -0.346 |
| 12657 | XLOC_005357 | chr10 | - | 103347002 | 103347925 | 103347002 | 103347163 | 103347791 | 628 | 103347925 | 12657 | 57,36 | 0,6 | 55,26 | 0,0 | 777 | 149 | 0.000830396 | 0.005216938 | 1.0,0.535 | 1.0,1.0 | -0.232 |
| 1130 | XLOC_021130 | chr19 | + | 35168568 | 35169718 | 35168568 | 35169008 | 35169641 | 633 | 35169718 | 1130 | 103,73 | 5,7 | 86,60 | 1,1 | 782 | 149 | 0.002329508 | 0.01232658 | 0.797,0.665 | 0.942,0.92 | -0.2 |
| 956 | XLOC_023553 | chr2 | + | 85770023 | 85772077 | 85770023 | 85770157 | 85770792 | 635 | 85772077 | 956 | 24,732,599 | 464,470 | 51,474,147 | 233,181 | 784 | 149 | 0 | 0 | 0.503,0.512 | 0.808,0.813 | -0.303 |
| 2590 | XLOC_020956 | chr19 | + | 9448471 | 9449258 | 9448471 | 9448534 | 9449169 | 635 | 9449258 | 2590 | 17,7 | 10,9 | 11,6 | 2,0 | 784 | 149 | 0.01196953 | 0.048216108 | 0.244,0.129 | 0.511,1.0 | -0.569 |
| 1698 | XLOC_001548 | chr1 | - | 228295935 | 228296722 | 228295935 | 228296019 | 228296655 | 636 | 228296722 | 1698 | 233,211 | 63,44 | 368,242 | 34,17 | 785 | 149 | 1.38E-07 | 2.45E-06 | 0.412,0.477 | 0.673,0.73 | -0.257 |
| 3959 | XLOC_044466 | chr8 | - | 145741371 | 145742574 | 145741371 | 145741797 | 145742433 | 636 | 145742574 | 3959 | 150,148 | 18,43 | 231,173 | 13,7 | 785 | 149 | 0.000106039 | 0.00090216 | 0.613,0.395 | 0.771,0.824 | -0.293 |
| 15270 | XLOC_009618 | chr12 | - | 133403153 | 133405283 | 133403153 | 133403553 | 133404192 | 639 | 133405283 | 15270 | 33,37 | 16,8 | 11,11 | 1,0 | 788 | 149 | 0.00029139 | 0.002158971 | 0.281,0.467 | 0.675,1.0 | -0.464 |
| 2950 | XLOC_037875 | chr6 | + | 52957531 | 52958359 | 52957531 | 52957622 | 52958263 | 641 | 52958359 | 2950 | 457,439 | 80,65 | 791,610 | 78,51 | 790 | 149 | 0.000473079 | 0.003283195 | 0.519,0.56 | 0.657,0.693 | -0.135 |
| 750 | XLOC_046136 | chr9 | + | 139442293 | 139446260 | 139442293 | 139442553 | 139443195 | 642 | 139446260 | 750 | 87,64 | 18,9 | 55,37 | 0,2 | 791 | 149 | 5.72E-05 | 0.000528525 | 0.477,0.573 | 1.0,0.777 | -0.364 |
| 1456 | XLOC_009585 | chr12 | - | 121880794 | 121881596 | 121880794 | 121880833 | 121881482 | 649 | 121881596 | 1456 | 44,25 | 78,53 | 39,27 | 35,19 | 798 | 149 | 0.00109982 | 0.0066125 | 0.095,0.081 | 0.172,0.21 | -0.103 |
| 3408 | XLOC_000953 | chr1 | - | 25570040 | 25571792 | 25570040 | 25570989 | 25571640 | 651 | 25571792 | 3408 | 348,282 | 36,29 | 719,541 | 7,11 | 800 | 149 | 2.19E-14 | 1.12E-12 | 0.643,0.644 | 0.95,0.902 | -0.282 |
| 14437 | XLOC_028384 | chr22 | + | 19751677 | 19752636 | 19751677 | 19751849 | 19752503 | 654 | 19752636 | 14437 | 39,49 | 4,1 | 139,82 | 0,0 | 803 | 149 | 0.000535404 | 0.003645857 | 0.644,0.901 | 1.0,1.0 | -0.228 |
| 2120 | XLOC_011529 | chr13 | - | 79884552 | 79894845 | 79884552 | 79893014 | 79893669 | 655 | 79894845 | 2120 | 237,203 | 49,54 | 381,389 | 56,54 | 804 | 149 | 0.005659002 | 0.025764695 | 0.473,0.411 | 0.558,0.572 | -0.123 |
| 11806 | XLOC_007509 | chr11 | - | 47598962 | 47600541 | 47598962 | 47599580 | 47600236 | 656 | 47600541 | 11806 | 5,4 | 1,1 | 6,6 | 0,0 | 805 | 149 | 0.005757066 | 0.026107242 | 0.481,0.425 | 1.0,1.0 | -0.547 |
| 2074 | XLOC_000418 | chr1 | + | 110881963 | 110890509 | 110881963 | 110888271 | 110888929 | 658 | 110890509 | 2074 | 421,416 | 28,29 | 377,277 | 7,13 | 807 | 149 | 0.004570626 | 0.021552904 | 0.735,0.726 | 0.909,0.797 | -0.123 |
| 2424 | XLOC_021296 | chr19 | + | 48982379 | 48985843 | 48982379 | 48983345 | 48984006 | 661 | 48985843 | 2424 | 234,303 | 31,17 | 229,127 | 7,5 | 810 | 149 | 0.006661269 | 0.029509754 | 0.581,0.766 | 0.858,0.824 | -0.167 |
| 3030 | XLOC_016775 | chr16 | - | 67863667 | 67865261 | 67863667 | 67864451 | 67865118 | 667 | 67865261 | 3030 | 77,72 | 88,104 | 375,228 | 55,29 | 816 | 149 | 0 | 0 | 0.138,0.112 | 0.555,0.589 | -0.447 |
| 624 | XLOC_029391 | chr3 | + | 42704503 | 42705433 | 42704503 | 42704619 | 42705288 | 669 | 42705433 | 624 | 17,5 | 26,22 | 25,51 | 19,22 | 818 | 149 | 2.58E-05 | 0.000261008 | 0.106,0.04 | 0.193,0.297 | -0.172 |
| 12551 | XLOC_011299 | chr13 | + | 39612462 | 39613425 | 39612462 | 39612602 | 39613272 | 670 | 39613425 | 12551 | 33,26 | 4,2 | 23,19 | 1,0 | 819 | 149 | 0.001907559 | 0.01044596 | 0.6,0.703 | 0.807,1.0 | -0.252 |
| 524 | XLOC_032657 | chr4 | - | 83343838 | 83347282 | 83343838 | 83346036 | 83346715 | 679 | 83347282 | 524 | 15,641,609 | 74,69 | 59,164,565 | 71,56 | 828 | 149 | 2.22E-16 | 1.46E-14 | 0.792,0.808 | 0.937,0.936 | -0.137 |
| 7661 | XLOC_018641 | chr17 | - | 55943834 | 55944776 | 55943834 | 55943905 | 55944589 | 684 | 55944776 | 7661 | 22,29 | 6,0 | 10,9 | 0,0 | 833 | 149 | 0.000911193 | 0.005636783 | 0.396,1.0 | 1.0,1.0 | -0.302 |
| 1828 | XLOC_014707 | chr15 | + | 75188492 | 75189560 | 75188492 | 75188666 | 75189351 | 685 | 75189560 | 1828 | 63,64 | 135,104 | 104,97 | 26,22 | 834 | 149 | 0 | 0 | 0.077,0.099 | 0.417,0.441 | -0.341 |
| 1595 | XLOC_032748 | chr4 | - | 128838979 | 128842926 | 128838979 | 128841991 | 128842678 | 687 | 128842926 | 1595 | 97,98 | 23,25 | 190,137 | 8,11 | 836 | 149 | 2.31E-06 | 3.09E-05 | 0.429,0.411 | 0.809,0.689 | -0.329 |
| 7803 | XLOC_018402 | chr17 | - | 8026348 | 8027255 | 8026348 | 8026444 | 8027134 | 690 | 8027255 | 7803 | 44,23 | 2,2 | 30,3 | 0,0 | 839 | 149 | 0.003198206 | 0.016102765 | 0.796,0.671 | 1.0,1.0 | -0.266 |

| | | | | | | | | | | | | | | | | | | | | | | |
|---|---|---|---|---|---|---|---|---|---|---|---|---|---|---|---|---|---|---|---|---|---|---|
| 1870 | XLOC_013325 | chr14 | - | 24035486 | 24036524 | 24035486 | 24035628 | 24036319 | 691 | 24036524 | 1870 | 86,90 | 14,13 | 83,70 | 2,1 | 840 | 149 | 5.90E-06 | 7.14E-05 | 0.521,0.551 | 0.88,0.925 | -0.367 |
| 13342 | XLOC_000133 | chr1 | + | 24974514 | 24975520 | 24974514 | 24974657 | 24975349 | 692 | 24975520 | 13342 | 120,150 | 11,4 | 83,70 | 0,0 | 841 | 149 | 7.59E-06 | 8.90E-05 | 0.659,0.869 | 1.0,1.0 | -0.236 |
| 3152 | XLOC_001352 | chr1 | - | 155238498 | 155239507 | 155238498 | 155238583 | 155239278 | 695 | 155239507 | 3152 | 513,518 | 28,32 | 586,406 | 6,5 | 844 | 149 | 1.41E-08 | 2.94E-07 | 0.764,0.741 | 0.945,0.935 | -0.188 |
| 528 | XLOC_037824 | chr6 | + | 41751199 | 41752810 | 41751199 | 41751976 | 41752672 | 696 | 41752810 | 528 | 29,42 | 42,51 | 18,22 | 12,7 | 845 | 149 | 0.007044205 | 0.030857867 | 0.109,0.127 | 0.209,0.357 | -0.165 |
| 3058 | XLOC_029849 | chr3 | - | 49725182 | 49726860 | 49725182 | 49725330 | 49726030 | 700 | 49726860 | 3058 | 201,192 | 13,2 | 407,311 | 1,0 | 849 | 149 | 1.55E-06 | 2.15E-05 | 0.731,0.944 | 0.986,1.0 | -0.156 |
| 748 | XLOC_044144 | chr8 | + | 124789504 | 124790332 | 124789504 | 124789562 | 124790264 | 702 | 124790332 | 748 | 8,11 | 107,145 | 65,47 | 79,70 | 851 | 149 | 0 | 0 | 0.013,0.013 | 0.126,0.105 | -0.102 |
| 10801 | XLOC_009464 | chr12 | - | 76434231 | 76442244 | 76434231 | 76441498 | 76442200 | 702 | 76442244 | 10801 | 906,823 | 189,169 | 1,205,869 | 144,117 | 851 | 149 | 0.000200987 | 0.001557073 | 0.456,0.46 | 0.594,0.565 | -0.121 |
| 2903 | XLOC_016538 | chr16 | + | 89759805 | 89761196 | 89759805 | 89759875 | 89760580 | 705 | 89761196 | 2903 | 750,755 | 108,122 | 808,550 | 61,19 | 854 | 149 | 1.91E-06 | 2.61E-05 | 0.548,0.519 | 0.698,0.835 | -0.233 |
| 3451 | XLOC_005363 | chr10 | - | 103866339 | 103868934 | 103866339 | 103868080 | 103868785 | 705 | 103868934 | 3451 | 619,607 | 81,57 | 520,457 | 29,23 | 854 | 149 | 0.000139406 | 0.001136083 | 0.571,0.65 | 0.758,0.776 | -0.156 |
| 1592 | XLOC_047952 | chrX | - | 48831566 | 48832523 | 48831566 | 48831721 | 48832429 | 708 | 48832523 | 1592 | 216,219 | 158,127 | 324,232 | 90,56 | 857 | 149 | 3.58E-09 | 8.35E-08 | 0.192,0.231 | 0.385,0.419 | -0.191 |
| 1347 | XLOC_018130 | chr17 | + | 46051765 | 46052703 | 46051765 | 46051814 | 46052524 | 710 | 46052703 | 1347 | 13,591,473 | 188,182 | 1,052,831 | 47,47 | 859 | 149 | 1.91E-10 | 5.61E-09 | 0.556,0.584 | 0.795,0.754 | -0.204 |
| 5543 | XLOC_028538 | chr22 | + | 46546695 | 46547873 | 46546695 | 46547080 | 46547790 | 710 | 46547873 | 5543 | 4,0 | 0,2 | 1,2 | 0,0 | 859 | 149 | 0.001186966 | 0.007043497 | 1.0,0.0 | 1.0,1.0 | -0.5 |
| 6195 | XLOC_024263 | chr2 | - | 165697665 | 165698608 | 165697665 | 165697756 | 165698466 | 710 | 165698608 | 6195 | 22,21 | 2,2 | 9,11 | 1,0 | 859 | 149 | 0.008036652 | 0.03453454 | 0.656,0.646 | 0.61,1.0 | -0.154 |
| 2700 | XLOC_000685 | chr1 | + | 204506557 | 204507436 | 204506557 | 204506625 | 204507336 | 711 | 204507436 | 2700 | 115,82 | 58,68 | 206,166 | 58,37 | 860 | 149 | 2.54E-06 | 3.37E-05 | 0.256,0.173 | 0.381,0.437 | -0.195 |
| 605 | XLOC_029512 | chr3 | + | 101571934 | 101573527 | 101571934 | 101572707 | 101573421 | 714 | 101573527 | 605 | 14,16 | 25,37 | 28,30 | 17,27 | 863 | 149 | 0.001002429 | 0.006110189 | 0.088,0.069 | 0.221,0.161 | -0.113 |
| 9484 | XLOC_041043 | chr7 | + | 26241280 | 26242229 | 26241280 | 26241446 | 26242160 | 714 | 26242229 | 9484 | 89,65 | 1,4 | 201,175 | 0,1 | 863 | 149 | 0.003998945 | 0.019374648 | 0.939,0.737 | 1.0,0.968 | -0.146 |
| 7480 | XLOC_026718 | chr20 | + | 17949547 | 17951013 | 17949547 | 17949723 | 17950443 | 720 | 17951013 | 7480 | 29,43 | 6,0 | 33,55 | 0,0 | 869 | 149 | 0.000791182 | 0.005012951 | 0.453,1.0 | 1.0,1.0 | -0.273 |
| 2260 | XLOC_046262 | chr9 | - | 41951843 | 41954633 | 41951843 | 41953779 | 41954503 | 724 | 41954633 | 2260 | 33,24 | 3,3 | 38,24 | 0,0 | 873 | 149 | 1.66E-05 | 0.00017624 | 0.652,0.577 | 1.0,1.0 | -0.385 |
| 11010 | XLOC_009165 | chr12 | + | 123460357 | 123461314 | 123460357 | 123460414 | 123461147 | 733 | 123461314 | 11010 | 45,40 | 12,6 | 24,47 | 1,1 | 882 | 149 | 0.000220027 | 0.001686996 | 0.388,0.53 | 0.802,0.888 | -0.386 |
| 833 | XLOC_029685 | chr3 | + | 184566858 | 184567810 | 184566858 | 184566983 | 184567723 | 740 | 184567810 | 833 | 10,4 | 31,28 | 52,50 | 60,43 | 889 | 149 | 1.86E-06 | 2.55E-05 | 0.051,0.023 | 0.127,0.163 | -0.108 |
| 3002 | XLOC_000907 | chr1 | - | 16944679 | 16946519 | 16944679 | 16945598 | 16946339 | 741 | 16946519 | 3002 | 16,35 | 50,38 | 65,89 | 10,16 | 890 | 149 | 0 | 0 | 0.051,0.134 | 0.521,0.482 | -0.409 |
| 1855 | XLOC_021639 | chr19 | - | 23436859 | 23443736 | 23436859 | 23442918 | 23443665 | 747 | 23443736 | 1855 | 9,13 | 5,1 | 3,12 | 0,0 | 896 | 149 | 6.70E-07 | 1.03E-05 | 0.23,0.684 | 1.0,1.0 | -0.543 |
| 8745 | XLOC_000120 | chr1 | + | 22827984 | 22828934 | 22827984 | 22828042 | 22828791 | 749 | 22828934 | 8745 | 47,44 | 9,0 | 90,64 | 0,0 | 898 | 149 | 0.000277479 | 0.002069077 | 0.464,1.0 | 1.0,1.0 | -0.268 |
| 2289 | XLOC_046145 | chr9 | + | 139745206 | 139746391 | 139745206 | 139745557 | 139746308 | 751 | 139746391 | 2289 | 13,5 | 11,6 | 24,35 | 2,3 | 900 | 149 | 1.51E-06 | 2.11E-05 | 0.164,0.121 | 0.665,0.659 | -0.52 |
| 3122 | XLOC_041406 | chr7 | - | 12375766 | 12376709 | 12375766 | 12375862 | 12376613 | 751 | 12376709 | 3122 | 17,25 | 11,14 | 45,39 | 4,3 | 900 | 149 | 2.88E-06 | 3.76E-05 | 0.204,0.228 | 0.651,0.683 | -0.451 |
| 2816 | XLOC_005076 | chr10 | + | 122665387 | 122666367 | 122665387 | 122665533 | 122666287 | 754 | 122666367 | 2816 | 175,202 | 108,149 | 406,264 | 108,51 | 903 | 149 | 5.90E-11 | 1.88E-09 | 0.211,0.183 | 0.383,0.461 | -0.225 |
| 2316 | XLOC_021921 | chr19 | - | 59080882 | 59081894 | 59080882 | 59080953 | 59081710 | 757 | 59081894 | 2316 | 109,106 | 96,113 | 319,223 | 31,20 | 906 | 149 | 0 | 0 | 0.157,0.134 | 0.629,0.647 | -0.492 |
| 7035 | XLOC_021165 | chr19 | + | 37341336 | 37342806 | 37341336 | 37341870 | 37342629 | 759 | 37342806 | 7035 | 2,3 | 2,2 | 1,3 | 0,0 | 908 | 149 | 2.87E-07 | 4.81E-06 | 0.141,0.198 | 1.0,1.0 | -0.831 |
| 4519 | XLOC_038093 | chr6 | - | 10410133 | 10411782 | 10410133 | 10410568 | 10411334 | 766 | 10411782 | 4519 | 55,38 | 7,3 | 114,67 | 0,4 | 915 | 149 | 0.007849277 | 0.033844109 | 0.561,0.673 | 1.0,0.732 | -0.249 |
| 557 | XLOC_029956 | chr3 | - | 108300954 | 108301930 | 108300954 | 108301049 | 108301823 | 774 | 108301930 | 557 | 59,28 | 67,51 | 63,75 | 26,31 | 923 | 149 | 1.80E-07 | 3.14E-06 | 0.124,0.081 | 0.281,0.281 | -0.179 |
| 3114 | XLOC_005198 | chr10 | - | 43046004 | 43047262 | 43046004 | 43046063 | 43046847 | 784 | 43047262 | 3114 | 17,19 | 3,2 | 13,11 | 0,0 | 933 | 149 | 1.15E-05 | 0.000128405 | 0.475,0.603 | 1.0,1.0 | -0.461 |
| 10820 | XLOC_038312 | chr6 | - | 58275150 | 58276289 | 58275150 | 58275346 | 58276136 | 790 | 58276289 | 10820 | 14,10 | 5,8 | 28,2 | 0,0 | 939 | 149 | 2.84E-09 | 6.78E-08 | 0.308,0.166 | 1.0,1.0 | -0.763 |
| 7663 | XLOC_018641 | chr17 | - | 55943834 | 55944776 | 55943834 | 55943905 | 55944698 | 793 | 55944776 | 7663 | 22,29 | 0,6 | 10,10 | 0,0 | 942 | 149 | 0.000657679 | 0.004328238 | 1.0,0.433 | 1.0,1.0 | -0.283 |
| 1444 | XLOC_024384 | chr2 | - | 208444105 | 208446670 | 208444105 | 208445722 | 208446517 | 795 | 208446670 | 1444 | 64,85 | 5,4 | 27,13 | 0,0 | 944 | 149 | 5.21E-05 | 0.000488243 | 0.669,0.77 | 1.0,1.0 | -0.28 |
| 11659 | XLOC_007616 | chr11 | - | 68028804 | 68030386 | 68028804 | 68029193 | 68030005 | 812 | 68030386 | 11659 | 130,128 | 7,1 | 159,118 | 2,0 | 961 | 149 | 0.010449737 | 0.042939087 | 0.742,0.952 | 0.925,1.0 | -0.116 |
| 2641 | XLOC_016664 | chr16 | - | 21829976 | 21831764 | 21829976 | 21830802 | 21831615 | 813 | 21831764 | 2641 | 27,16 | 6,4 | 25,22 | 0,2 | 962 | 149 | 0.004479154 | 0.021185229 | 0.411,0.383 | 1.0,0.63 | -0.418 |
| 587 | XLOC_029664 | chr3 | + | 180326488 | 180327791 | 180326488 | 180326598 | 180327417 | 819 | 180327791 | 587 | 445,508 | 46,37 | 768,603 | 18,7 | 968 | 149 | 4.60E-11 | 1.50E-09 | 0.598,0.679 | 0.868,0.93 | -0.26 |
| 1972 | XLOC_009588 | chr12 | - | 122239612 | 122240690 | 122239612 | 122239714 | 122240536 | 822 | 122240690 | 1972 | 92,82 | 15,32 | 91,64 | 7,3 | 971 | 149 | 0.000572921 | 0.003854864 | 0.485,0.282 | 0.666,0.766 | -0.333 |
| 5480 | XLOC_028606 | chr22 | - | 24984181 | 24985233 | 24984181 | 24984297 | 24985123 | 826 | 24985233 | 5480 | 4,15 | 4,0 | 3,12 | 0,0 | 975 | 149 | 0.002153336 | 0.011585357 | 0.133,1.0 | 1.0,1.0 | -0.433 |
| 8601 | XLOC_021326 | chr19 | + | 50381754 | 50382717 | 50381754 | 50381829 | 50382657 | 828 | 50382717 | 8601 | 60,57 | 1,4 | 39,13 | 0,0 | 977 | 149 | 0.000746425 | 0.004770134 | 0.901,0.685 | 1.0,1.0 | -0.207 |
| 548 | XLOC_030068 | chr3 | - | 148747870 | 148750159 | 148747870 | 148749150 | 148749987 | 837 | 148750159 | 548 | 14,021,416 | 80,63 | 23,081,650 | 52,50 | 986 | 149 | 4.52E-05 | 0.000430866 | 0.726,0.773 | 0.87,0.833 | -0.102 |
| 5080 | XLOC_029956 | chr3 | - | 108300954 | 108301930 | 108300954 | 108300985 | 108301823 | 838 | 108301930 | 5080 | 87,61 | 2,3 | 82,83 | 0,0 | 987 | 149 | 0.00152916 | 0.008708195 | 0.868,0.754 | 1.0,1.0 | -0.189 |
| 15139 | XLOC_034614 | chr5 | + | 87565061 | 87578683 | 87565061 | 87565504 | 87566342 | 838 | 87578683 | 15139 | 4,11 | 2,1 | 7,7 | 0,0 | 987 | 149 | 0.001374421 | 0.007970319 | 0.232,0.624 | 1.0,1.0 | -0.572 |
| 3338 | XLOC_032661 | chr4 | - | 83813646 | 83819215 | 83813646 | 83816000 | 83816844 | 844 | 83819215 | 3338 | 142,135 | 104,90 | 196,192 | 94,57 | 993 | 149 | 0.001235283 | 0.007293435 | 0.17,0.184 | 0.238,0.336 | -0.11 |
| 362 | XLOC_035102 | chr5 | - | 148871471 | 148876423 | 148871471 | 148873184 | 148874036 | 852 | 148876423 | 362 | 343,334 | 16,8 | 357,270 | 1,2 | 1001 | 149 | 1.03E-05 | 0.000116851 | 0.761,0.861 | 0.982,0.953 | -0.157 |
| 846 | XLOC_016250 | chr16 | + | 3331074 | 3335239 | 3331074 | 3334205 | 3335058 | 853 | 3335239 | 846 | 112,88 | 58,38 | 215,182 | 39,33 | 1002 | 149 | 1.52E-06 | 2.13E-05 | 0.223,0.256 | 0.45,0.451 | -0.211 |
| 500 | XLOC_032343 | chr4 | + | 68586139 | 68588555 | 68586139 | 68586339 | 68587194 | 855 | 68588555 | 500 | 37,34 | 1,9 | 74,50 | 0,0 | 1004 | 149 | 6.20E-06 | 7.43E-05 | 0.846,0.359 | 1.0,1.0 | -0.397 |
| 1536 | XLOC_016387 | chr16 | + | 31717210 | 31728151 | 31717210 | 31717355 | 31718211 | 856 | 31728151 | 1536 | 7,8 | 22,7 | 18,29 | 16,8 | 1005 | 149 | 0.003227664 | 0.016229218 | 0.045,0.145 | 0.143,0.35 | -0.151 |
| 2211 | XLOC_009393 | chr12 | - | 54573273 | 54576407 | 54573273 | 54575431 | 54576287 | 856 | 54576407 | 2211 | 436,320 | 74,58 | 506,449 | 39,31 | 1005 | 149 | 1.55E-06 | 2.15E-05 | 0.466,0.45 | 0.658,0.682 | -0.212 |
| 14126 | XLOC_018763 | chr17 | - | 78183459 | 78185722 | 78183459 | 78184810 | 78185667 | 857 | 78185722 | 14126 | 83,98 | 4,6 | 76,77 | 1,0 | 1006 | 149 | 0.000145868 | 0.001181928 | 0.754,0.708 | 0.918,1.0 | -0.228 |
| 40 | XLOC_046502 | chr9 | - | 140469203 | 140470619 | 140469203 | 140469298 | 140470160 | 862 | 140470619 | 40 | 280,259 | 33,45 | 476,336 | 31,19 | 1011 | 149 | 0.000144065 | 0.001169328 | 0.556,0.459 | 0.694,0.723 | -0.201 |
| 8960 | XLOC_014944 | chr15 | - | 65113565 | 65114590 | 65113565 | 65113596 | 65114464 | 868 | 65114590 | 8960 | 50,61 | 4,3 | 42,29 | 0,0 | 1017 | 149 | 3.37E-05 | 0.000332657 | 0.647,0.749 | 1.0,1.0 | -0.302 |
| 1552 | XLOC_016298 | chr16 | + | 14354765 | 14360799 | 14354765 | 14355359 | 14356234 | 875 | 14360799 | 1552 | 326,304 | 4,12 | 276,218 | 1,0 | 1024 | 149 | 2.53E-05 | 0.00025601 | 0.922,0.787 | 0.976,1.0 | -0.133 |
| 522 | XLOC_045865 | chr9 | + | 5772891 | 5778429 | 5772891 | 5773080 | 5773957 | 877 | 5778429 | 522 | 121,97 | 70,74 | 146,98 | 42,33 | 1026 | 149 | 9.60E-05 | 0.000828157 | 0.201,0.16 | 0.335,0.301 | -0.138 |
| 1639 | XLOC_001283 | chr1 | - | 146626258 | 146633359 | 146626258 | 146632412 | 146633290 | 878 | 146633359 | 1639 | 101,94 | 64,59 | 179,174 | 45,47 | 1027 | 149 | 3.24E-06 | 4.17E-05 | 0.186,0.188 | 0.366,0.349 | -0.17 |
| 1129 | XLOC_021130 | chr19 | + | 35168568 | 35169718 | 35168568 | 35168761 | 35169641 | 880 | 35169718 | 1129 | 169,135 | 44,47 | 140,107 | 13,13 | 1029 | 149 | 2.11E-05 | 0.000218186 | 0.357,0.294 | 0.609,0.544 | -0.251 |
| 11808 | XLOC_007509 | chr11 | - | 47598962 | 47600541 | 47598962 | 47599580 | 47600463 | 883 | 47600541 | 11808 | 5,4 | 1,1 | 6,6 | 0,0 | 1032 | 149 | 0.002317632 | 0.012291368 | 0.419,0.366 | 1.0,1.0 | -0.608 |
| 519 | XLOC_037917 | chr6 | + | 88049833 | 88054575 | 88049833 | 88050055 | 88050943 | 888 | 88054575 | 519 | 63,68 | 7,2 | 79,82 | 0,0 | 1037 | 149 | 4.84E-06 | 5.99E-05 | 0.564,0.83 | 1.0,1.0 | -0.303 |
| 3586 | XLOC_047954 | chrX | - | 48934303 | 48935406 | 48934303 | 48934412 | 48935301 | 889 | 48935406 | 3586 | 125,140 | 5,20 | 157,131 | 4,0 | 1038 | 149 | 0.00059484 | 0.003976242 | 0.782,0.501 | 0.849,1.0 | -0.283 |
| 3031 | XLOC_041344 | chr7 | + | 150066759 | 150067973 | 150066759 | 150066957 | 150067848 | 891 | 150067973 | 3031 | 836,747 | 65,66 | 18,971,414 | 59,61 | 1040 | 149 | 5.61E-07 | 8.80E-06 | 0.648,0.619 | 0.822,0.769 | -0.162 |
| 17 | XLOC_047954 | chrX | - | 48934303 | 48935406 | 48934303 | 48934409 | 48935301 | 892 | 48935406 | 17 | 622,611 | 160,219 | 717,498 | 36,25 | 1041 | 149 | 0 | 0 | 0.358,0.285 | 0.74,0.74 | -0.418 |
| 1826 | XLOC_026692 | chr20 | + | 3844902 | 3856888 | 3844902 | 3845435 | 3846329 | 894 | 3856888 | 1826 | 34,26 | 109,101 | 81,58 | 70,52 | 1043 | 149 | 4.97E-11 | 1.60E-09 | 0.043,0.035 | 0.142,0.137 | -0.101 |
| 691 | XLOC_028592 | chr22 | - | 22311411 | 22313065 | 22311411 | 22311967 | 22312863 | 896 | 22313065 | 691 | 136,142 | 26,24 | 285,198 | 14,11 | 1045 | 149 | 2.19E-06 | 2.95E-05 | 0.427,0.458 | 0.744,0.72 | -0.289 |
| 1584 | XLOC_016758 | chr16 | - | 67212101 | 67213465 | 67212101 | 67212407 | 67213304 | 897 | 67213465 | 1584 | 195,240 | 7,4 | 176,111 | 1,0 | 1046 | 149 | 0.000417412 | 0.002942846 | 0.799,0.895 | 0.962,1.0 | -0.134 |
| 2129 | XLOC_007597 | chr11 | - | 66133901 | 66135019 | 66133901 | 66134035 | 66134934 | 899 | 66135019 | 2129 | 268,266 | 143,125 | 444,246 | 65,29 | 1048 | 149 | 8.88E-16 | 5.36E-14 | 0.21,0.232 | 0.493,0.547 | -0.299 |

| | | | | | | | | | | | | | | | | | | | | | | |
|---|---|---|---|---|---|---|---|---|---|---|---|---|---|---|---|---|---|---|---|---|---|---|
| 2059 | XLOC_021489 | chr19 | - | 5611130 | 5612578 | 5611130 | 5611641 | 5612550 | 909 | 5612578 | 2059 | 108,136 | 69,85 | 382,295 | 82,84 | 1058 | 149 | 1.88E-08 | 3.88E-07 | 0.181,0.184 | 0.396,0.331 | -0.181 |
| 1525 | XLOC_016435 | chr16 | + | 67195637 | 67197755 | 67195637 | 67195840 | 67196750 | 910 | 67197755 | 1525 | 217,214 | 21,24 | 232,149 | 0,7 | 1059 | 149 | 6.27E-05 | 0.00057125 | 0.592,0.556 | 1.0,0.75 | -0.301 |
| 2967 | XLOC_048031 | chrX | - | 100870065 | 100871756 | 100870065 | 100870750 | 100871661 | 911 | 100871756 | 2967 | 10,2 | 2,4 | 34,33 | 0,1 | 1060 | 149 | 7.66E-07 | 1.16E-05 | 0.413,0.066 | 1.0,0.823 | -0.672 |
| 3216 | XLOC_021355 | chr19 | + | 53883974 | 53890179 | 53883974 | 53887309 | 53888222 | 913 | 53890179 | 3216 | 68,73 | 3,3 | 88,71 | 0,1 | 1062 | 149 | 0.001303031 | 0.007629648 | 0.761,0.773 | 1.0,0.909 | -0.188 |
| 2264 | XLOC_018643 | chr17 | - | 56081011 | 56082402 | 56081011 | 56081362 | 56082283 | 921 | 56082402 | 2264 | 34,773,303 | 553,542 | 55,144,225 | 225,221 | 1070 | 149 | 0 | 0 | 0.467,0.459 | 0.773,0.727 | -0.287 |
| 723 | XLOC_027814 | chr21 | - | 45750705 | 45751897 | 45750705 | 45750802 | 45751725 | 923 | 45751897 | 723 | 81,65 | 28,23 | 125,96 | 18,10 | 1072 | 149 | 0.000197299 | 0.001533543 | 0.287,0.282 | 0.491,0.572 | -0.247 |
| 8816 | XLOC_016758 | chr16 | - | 67212101 | 67213465 | 67212101 | 67212409 | 67213332 | 923 | 67213465 | 8816 | 79,110 | 4,9 | 68,35 | 2,0 | 1072 | 149 | 0.006577023 | 0.02920984 | 0.733,0.629 | 0.825,1.0 | -0.231 |
| 38 | XLOC_047663 | chrX | + | 23801916 | 23803546 | 23801916 | 23802520 | 23803444 | 924 | 23803546 | 38 | 22,532,208 | 105,121 | 60,025,012 | 107,95 | 1073 | 149 | 4.88E-13 | 2.15E-11 | 0.749,0.717 | 0.886,0.88 | -0.15 |
| 2361 | XLOC_030033 | chr3 | - | 129149732 | 129151449 | 129149732 | 129150421 | 129151345 | 924 | 129151449 | 2361 | 179,185 | 72,63 | 534,312 | 78,65 | 1073 | 149 | 2.20E-05 | 0.000226607 | 0.257,0.29 | 0.487,0.4 | -0.17 |
| 14140 | XLOC_000682 | chr1 | + | 203770702 | 203772144 | 203770702 | 203770767 | 203771696 | 929 | 203772144 | 14140 | 230,268 | 0,10 | 341,282 | 0,0 | 1078 | 149 | 0.000102931 | 0.0008789 | 1.0,0.787 | 1.0,1.0 | -0.107 |
| 586 | XLOC_029664 | chr3 | + | 180325435 | 180326598 | 180325435 | 180325553 | 180326488 | 935 | 180326598 | 586 | 393,429 | 19,19 | 684,452 | 16,10 | 1084 | 149 | 0.009425114 | 0.039300528 | 0.74,0.756 | 0.855,0.861 | -0.11 |
| 1488 | XLOC_029641 | chr3 | + | 160118609 | 160119881 | 160118609 | 160118753 | 160119702 | 949 | 160119881 | 1488 | 219,238 | 284,310 | 225,194 | 111,86 | 1098 | 149 | 2.63E-12 | 1.05E-10 | 0.095,0.094 | 0.216,0.234 | -0.131 |
| 775 | XLOC_026894 | chr20 | - | 415979 | 419485 | 415979 | 418242 | 419193 | 951 | 419485 | 775 | 308,394 | 25,16 | 546,390 | 1,1 | 1100 | 149 | 8.12E-13 | 3.52E-11 | 0.625,0.769 | 0.987,0.981 | -0.287 |
| 832 | XLOC_029685 | chr3 | + | 184556501 | 184557540 | 184556501 | 184556534 | 184557485 | 951 | 184557540 | 832 | 20,14 | 16,9 | 152,132 | 21,28 | 1100 | 149 | 8.48E-06 | 9.81E-05 | 0.145,0.174 | 0.495,0.39 | -0.283 |
| 821 | XLOC_018271 | chr17 | + | 78084739 | 78085899 | 78084739 | 78084824 | 78085781 | 957 | 78085899 | 821 | 233,248 | 155,163 | 342,208 | 77,88 | 1106 | 149 | 0.000135581 | 0.001109395 | 0.168,0.17 | 0.374,0.242 | -0.139 |
| 1082 | XLOC_021444 | chr19 | - | 1229472 | 1231274 | 1229472 | 1230141 | 1231099 | 958 | 1231274 | 1082 | 65,54 | 25,24 | 143,85 | 12,9 | 1107 | 149 | 1.53E-07 | 2.69E-06 | 0.259,0.232 | 0.616,0.56 | -0.343 |
| 2893 | XLOC_020130 | chr18 | + | 60241289 | 60245879 | 60241289 | 60242783 | 60243744 | 961 | 60245879 | 2893 | 81,77 | 52,44 | 162,201 | 38,36 | 1110 | 149 | 1.01E-07 | 1.85E-06 | 0.173,0.19 | 0.364,0.428 | -0.215 |
| 2463 | XLOC_007511 | chr11 | - | 47752925 | 47754222 | 47752925 | 47753113 | 47754088 | 975 | 47754222 | 2463 | 11,091,100 | 99,115 | 1,053,783 | 59,43 | 1124 | 149 | 0.000402059 | 0.002852914 | 0.598,0.559 | 0.703,0.707 | -0.126 |
| 2655 | XLOC_011393 | chr13 | + | 115007595 | 115008823 | 115007595 | 115007755 | 115008731 | 976 | 115008823 | 2655 | 514,552 | 258,297 | 697,539 | 180,126 | 1125 | 149 | 8.33E-09 | 1.81E-07 | 0.209,0.198 | 0.339,0.362 | -0.147 |
| 1976 | XLOC_009388 | chr12 | - | 54104919 | 54106990 | 54104919 | 54105905 | 54106883 | 978 | 54106990 | 1976 | 432,431 | 116,127 | 544,378 | 61,66 | 1127 | 149 | 0.000144258 | 0.00117022 | 0.33,0.31 | 0.541,0.431 | -0.166 |
| 601 | XLOC_029529 | chr3 | + | 113677209 | 113681954 | 113677209 | 113677385 | 113678367 | 982 | 113681954 | 601 | 314,298 | 16,18 | 408,305 | 8,1 | 1131 | 149 | 2.15E-05 | 0.000222053 | 0.721,0.686 | 0.87,0.976 | -0.22 |
| 2023 | XLOC_011449 | chr13 | - | 37596540 | 37598330 | 37596540 | 37597181 | 37598171 | 990 | 37598330 | 2023 | 152,119 | 18,15 | 315,294 | 0,0 | 1139 | 149 | 0 | 0 | 0.525,0.509 | 1.0,1.0 | -0.483 |
| 760 | XLOC_026993 | chr20 | - | 37049652 | 37050888 | 37049652 | 37049745 | 37050738 | 993 | 37050888 | 760 | 203,223 | 27,25 | 495,350 | 13,22 | 1142 | 149 | 0.000213472 | 0.001642085 | 0.495,0.538 | 0.832,0.675 | -0.237 |
| 1659 | XLOC_047898 | chrX | - | 14024610 | 14027288 | 14024610 | 14025989 | 14026989 | 1000 | 14027288 | 1659 | 141,144 | 4,6 | 122,63 | 0,0 | 1149 | 149 | 2.51E-05 | 0.000255269 | 0.821,0.757 | 1.0,1.0 | -0.211 |
| 1133 | XLOC_021130 | chr19 | + | 35174059 | 35178584 | 35174059 | 35174143 | 35175156 | 1013 | 35178584 | 1133 | 171,160 | 79,65 | 155,149 | 38,24 | 1162 | 149 | 0.000374589 | 0.002681497 | 0.217,0.24 | 0.343,0.443 | -0.165 |
| 2158 | XLOC_021499 | chr19 | - | 6413123 | 6415312 | 6413123 | 6414131 | 6415151 | 1020 | 6415312 | 2158 | 19,221,731 | 61,95 | 44,243,234 | 38,29 | 1169 | 149 | 4.44E-16 | 2.74E-14 | 0.801,0.699 | 0.937,0.934 | -0.185 |
| 3059 | XLOC_001153 | chr1 | - | 71531360 | 71532617 | 71531360 | 71531435 | 71532458 | 1023 | 71532617 | 3059 | 13,991,340 | 148,125 | 20,301,646 | 105,79 | 1172 | 149 | 7.93E-07 | 1.20E-05 | 0.546,0.577 | 0.711,0.726 | -0.157 |
| 9044 | XLOC_047898 | chrX | - | 14024610 | 14027288 | 14024610 | 14025989 | 14027017 | 1028 | 14027288 | 9044 | 81,76 | 5,1 | 62,33 | 0,0 | 1177 | 149 | 0.000147224 | 0.001192224 | 0.672,0.906 | 1.0,1.0 | -0.211 |
| 737 | XLOC_027721 | chr21 | + | 46709265 | 46717412 | 46709265 | 46709353 | 46710385 | 1032 | 46717412 | 737 | 37,26 | 26,21 | 35,39 | 12,6 | 1181 | 149 | 0.003180872 | 0.016044094 | 0.152,0.135 | 0.269,0.451 | -0.216 |
| 2436 | XLOC_007615 | chr11 | - | 67933021 | 67934645 | 67933021 | 67933415 | 67934448 | 1033 | 67934645 | 2436 | 580,674 | 18,15 | 802,626 | 9,4 | 1182 | 149 | 0.000706914 | 0.004571315 | 0.802,0.85 | 0.918,0.952 | -0.109 |
| 2599 | XLOC_041436 | chr7 | - | 27201389 | 27204586 | 27201389 | 27203460 | 27204496 | 1036 | 27204586 | 2599 | 90,120 | 43,39 | 66,68 | 9,7 | 1185 | 149 | 5.80E-05 | 0.000534639 | 0.208,0.279 | 0.48,0.55 | -0.272 |
| 2568 | XLOC_014712 | chr15 | + | 75644942 | 75646207 | 75644942 | 75645042 | 75646079 | 1037 | 75646207 | 2568 | 106,106 | 8,5 | 109,103 | 0,0 | 1186 | 149 | 3.58E-07 | 5.87E-06 | 0.625,0.727 | 1.0,1.0 | -0.324 |
| 3469 | XLOC_016815 | chr16 | - | 74481268 | 74487232 | 74481268 | 74486029 | 74487068 | 1039 | 74487232 | 3469 | 10,931,175 | 109,112 | 23,321,796 | 24,55 | 1188 | 149 | 1.11E-16 | 7.47E-15 | 0.557,0.568 | 0.924,0.804 | -0.302 |
| 1209 | XLOC_023637 | chr2 | + | 114399607 | 114400918 | 114399607 | 114399722 | 114400766 | 1044 | 114400918 | 1209 | 52,41 | 2,2 | 12,6 | 0,0 | 1193 | 149 | 0.002539994 | 0.013232141 | 0.765,0.719 | 1.0,1.0 | -0.258 |
| 13997 | XLOC_013315 | chr14 | - | 23535174 | 23536321 | 23535174 | 23535217 | 23536264 | 1047 | 23536321 | 13997 | 88,89 | 4,1 | 152,126 | 0,0 | 1196 | 149 | 0.001349693 | 0.00785726 | 0.733,0.917 | 1.0,1.0 | -0.175 |
| 3014 | XLOC_018646 | chr17 | - | 56492686 | 56495003 | 56492686 | 56493323 | 56494377 | 1054 | 56495003 | 3014 | 245,282 | 10,11 | 231,166 | 5,0 | 1203 | 149 | 0.006135104 | 0.027567952 | 0.752,0.76 | 0.851,1.0 | -0.169 |
| 2005 | XLOC_007426 | chr11 | - | 10871986 | 10877804 | 10871986 | 10876633 | 10877690 | 1057 | 10877804 | 2005 | 324,341 | 81,74 | 435,287 | 57,30 | 1206 | 149 | 0.000126089 | 0.001044761 | 0.331,0.363 | 0.485,0.542 | -0.167 |
| 3157 | XLOC_018763 | chr17 | - | 78183459 | 78186073 | 78183459 | 78184810 | 78185869 | 1059 | 78186073 | 3157 | 382,428 | 59,60 | 497,419 | 26,17 | 1208 | 149 | 1.06E-08 | 2.26E-07 | 0.444,0.468 | 0.702,0.752 | -0.271 |
| 1554 | XLOC_027049 | chr20 | - | 56820838 | 56822562 | 56820838 | 56821308 | 56822377 | 1069 | 56822562 | 1554 | 39,45 | 11,14 | 104,71 | 6,5 | 1218 | 149 | 2.48E-05 | 0.000252241 | 0.303,0.282 | 0.68,0.635 | -0.365 |
| 1492 | XLOC_035181 | chr5 | - | 180272366 | 180278493 | 180272366 | 180275082 | 180276153 | 1071 | 180278493 | 1492 | 561,535 | 21,21 | 641,419 | 12,6 | 1220 | 149 | 0.003615195 | 0.017870986 | 0.765,0.757 | 0.867,0.895 | -0.12 |
| 4285 | XLOC_041384 | chr7 | - | 2579419 | 2580686 | 2579419 | 2579541 | 2580612 | 1071 | 2580686 | 4285 | 18,29 | 3,2 | 30,21 | 0,0 | 1220 | 149 | 7.04E-06 | 8.32E-05 | 0.423,0.639 | 1.0,1.0 | -0.469 |
| 8409 | XLOC_004859 | chr10 | + | 70481011 | 70482334 | 70481011 | 70481136 | 70482211 | 1075 | 70482334 | 8409 | 12,15 | 3,0 | 17,6 | 0,0 | 1224 | 149 | 0.012356766 | 0.049563931 | 0.327,1.0 | 1.0,1.0 | -0.337 |
| 2013 | XLOC_011561 | chr13 | - | 113978518 | 113980161 | 113978518 | 113978892 | 113979975 | 1083 | 113980161 | 2013 | 103,114 | 80,101 | 413,354 | 42,30 | 1232 | 149 | 0 | 0 | 0.135,0.12 | 0.543,0.588 | -0.438 |
| 2262 | XLOC_018643 | chr17 | - | 56067009 | 56068306 | 56067009 | 56067157 | 56068242 | 1085 | 56068306 | 2262 | 150,134 | 40,26 | 78,73 | 8,5 | 1234 | 149 | 0.002218671 | 0.011851303 | 0.312,0.384 | 0.541,0.638 | -0.242 |
| 9917 | XLOC_013242 | chr14 | + | 94567084 | 94568321 | 94567084 | 94567117 | 94568216 | 1099 | 94568321 | 9917 | 51,62 | 4,2 | 50,22 | 0,0 | 1248 | 149 | 4.30E-05 | 0.000418378 | 0.604,0.787 | 1.0,1.0 | -0.304 |
| 13413 | XLOC_007142 | chr11 | + | 64852586 | 64854080 | 64852586 | 64852786 | 64853886 | 1100 | 64854080 | 13413 | 25,16 | 2,3 | 29,8 | 0,0 | 1249 | 149 | 3.38E-06 | 4.32E-05 | 0.599,0.389 | 1.0,1.0 | -0.506 |
| 2767 | XLOC_001135 | chr1 | - | 59154709 | 59155921 | 59154709 | 59154788 | 59155897 | 1109 | 59155921 | 2767 | 152,203 | 35,55 | 436,362 | 56,39 | 1258 | 149 | 5.98E-05 | 0.000547296 | 0.34,0.304 | 0.48,0.524 | -0.18 |
| 3545 | XLOC_048083 | chrX | - | 130917914 | 130919292 | 130917914 | 130918067 | 130919180 | 1113 | 130919292 | 3545 | 0,3 | 1,2 | 8,4 | 0,1 | 1262 | 149 | 0.000725926 | 0.004662237 | 0.0,0.15 | 1.0,0.321 | -0.586 |
| 1955 | XLOC_000237 | chr1 | + | 40942969 | 40945160 | 40942969 | 40943566 | 40944687 | 1121 | 40945160 | 1955 | 7,5 | 3,1 | 24,7 | 1,0 | 1270 | 149 | 0.000166453 | 0.001323661 | 0.215,0.37 | 0.738,1.0 | -0.577 |
| 7637 | XLOC_018673 | chr17 | - | 62496666 | 62498187 | 62496666 | 62497000 | 62498127 | 1127 | 62498187 | 7637 | 515,496 | 14,7 | 723,599 | 1,2 | 1276 | 149 | 1.62E-06 | 2.24E-05 | 0.811,0.892 | 0.988,0.972 | -0.128 |
| 2030 | XLOC_011375 | chr13 | + | 111289475 | 111292474 | 111289475 | 111289596 | 111290724 | 1128 | 111292474 | 2030 | 282,241 | 130,113 | 695,467 | 91,56 | 1277 | 149 | 0 | 0 | 0.202,0.199 | 0.471,0.493 | -0.281 |
| 205 | XLOC_041217 | chr7 | + | 99950995 | 99952863 | 99950995 | 99951635 | 99952765 | 1130 | 99952863 | 205 | 107,130 | 126,102 | 193,144 | 92,26 | 1279 | 149 | 3.18E-05 | 0.000315502 | 0.09,0.129 | 0.196,0.392 | -0.185 |
| 1191 | XLOC_020232 | chr18 | - | 47794348 | 47799108 | 47794348 | 47797910 | 47799046 | 1136 | 47799108 | 1191 | 206,228 | 81,52 | 770,549 | 107,61 | 1285 | 149 | 4.11E-06 | 5.17E-05 | 0.228,0.337 | 0.455,0.511 | -0.2 |
| 2952 | XLOC_029987 | chr3 | - | 121437234 | 121438615 | 121437234 | 121437348 | 121438492 | 1144 | 121438615 | 2952 | 56,90 | 64,76 | 88,86 | 38,13 | 1293 | 149 | 0.000106479 | 0.000904818 | 0.092,0.12 | 0.211,0.433 | -0.216 |
| 8157 | XLOC_018376 | chr17 | - | 6529849 | 6532061 | 6529849 | 6530288 | 6531436 | 1148 | 6532061 | 8157 | 21,14 | 6,5 | 11,15 | 0,1 | 1297 | 149 | 0.000303737 | 0.002241212 | 0.287,0.243 | 1.0,0.633 | -0.551 |
| 477 | XLOC_032656 | chr4 | - | 83273437 | 83276554 | 83273437 | 83275307 | 83276456 | 1149 | 83276554 | 477 | 12,001,159 | 11,171,059 | 39,383,072 | 722,640 | 1298 | 149 | 0 | 0 | 0.11,0.112 | 0.385,0.355 | -0.259 |
| 649 | XLOC_028756 | chr22 | - | 51220615 | 51222070 | 51220615 | 51220779 | 51221928 | 1149 | 51222070 | 649 | 153,117 | 1,14 | 82,58 | 0,0 | 1298 | 149 | 7.50E-05 | 0.000665724 | 0.946,0.49 | 1.0,1.0 | -0.282 |
| 525 | XLOC_032657 | chr4 | - | 83343838 | 83347282 | 83343838 | 83346036 | 83347189 | 1153 | 83347282 | 525 | 26,172,600 | 623,596 | 104,807,890 | 823,693 | 1302 | 149 | 0 | 0 | 0.325,0.333 | 0.593,0.566 | -0.25 |
| 2163 | XLOC_024165 | chr2 | - | 101648729 | 101650220 | 101648729 | 101648847 | 101650005 | 1158 | 101650220 | 2163 | 278,291 | 87,112 | 413,341 | 77,63 | 1307 | 149 | 6.60E-05 | 0.000596427 | 0.267,0.229 | 0.379,0.382 | -0.133 |
| 1351 | XLOC_018098 | chr17 | + | 42808332 | 42809633 | 42808332 | 42808383 | 42809545 | 1162 | 42809633 | 1351 | 137,129 | 32,33 | 179,112 | 14,8 | 1311 | 149 | 8.25E-06 | 9.57E-05 | 0.327,0.308 | 0.592,0.614 | -0.285 |
| 2081 | XLOC_005256 | chr10 | - | 74092144 | 74095689 | 74092144 | 74094375 | 74095537 | 1162 | 74095689 | 2081 | 312,236 | 130,73 | 415,256 | 45,31 | 1311 | 149 | 1.48E-11 | 5.20E-10 | 0.214,0.269 | 0.512,0.484 | -0.257 |
| 2299 | XLOC_009325 | chr12 | - | 48177861 | 48179254 | 48177861 | 48178000 | 48179169 | 1169 | 48179254 | 2299 | 291,232 | 148,106 | 372,279 | 74,76 | 1318 | 149 | 2.95E-06 | 3.84E-05 | 0.182,0.198 | 0.362,0.293 | -0.138 |
| 2562 | XLOC_006990 | chr11 | + | 8987450 | 8990724 | 8987450 | 8988203 | 8989385 | 1182 | 8990724 | 2562 | 56,60 | 25,13 | 28,28 | 1,3 | 1331 | 149 | 0.007650627 | 0.033093355 | 0.2,0.341 | 0.758,0.511 | -0.364 |
| 2955 | XLOC_007142 | chr11 | + | 64852586 | 64854080 | 64852586 | 64852698 | 64853886 | 1188 | 64854080 | 2955 | 257,198 | 126,107 | 229,161 | 73,35 | 1337 | 149 | 0.000354858 | 0.00255697 | 0.185,0.171 | 0.259,0.339 | -0.121 |
| 148 | XLOC_041749 | chr7 | - | 141356343 | 141364040 | 141356343 | 141362779 | 141363971 | 1192 | 141364040 | 148 | 203,160 | 6,6 | 151,108 | 2,0 | 1341 | 149 | 0.002697444 | 0.013938283 | 0.79,0.748 | 0.893,1.0 | -0.177 |

| | | | | | | | | | | | | | | | | | | | | | | |
|---|---|---|---|---|---|---|---|---|---|---|---|---|---|---|---|---|---|---|---|---|---|---|
| 8012 | XLOC_017998 | chr17 | + | 28958772 | 28960408 | 28958772 | 28958913 | 28960115 | 1202 | 28960408 | 8012 | 181,152 | 7,3 | 88,104 | 1,0 | 1351 | 149 | 0.000674684 | 0.004411369 | 0.74,0.848 | 0.907,1.0 | -0.159 |
| 368 | XLOC_034970 | chr5 | - | 87493485 | 87494967 | 87493485 | 87493582 | 87494792 | 1210 | 87494967 | 368 | 135,119 | 42,62 | 309,236 | 85,39 | 1359 | 149 | 0.004968372 | 0.02315022 | 0.261,0.174 | 0.285,0.399 | -0.124 |
| 2082 | XLOC_005256 | chr10 | - | 74092144 | 74095689 | 74092144 | 74094375 | 74095591 | 1216 | 74095689 | 2082 | 361,282 | 61,62 | 466,276 | 34,17 | 1365 | 149 | 1.33E-07 | 2.38E-06 | 0.392,0.332 | 0.599,0.639 | -0.257 |
| 174 | XLOC_041438 | chr7 | - | 27228813 | 27238061 | 27228813 | 27233177 | 27234404 | 1227 | 27238061 | 174 | 21,20 | 4,0 | 6,9 | 0,0 | 1376 | 149 | 0.002499159 | 0.013082143 | 0.362,1.0 | 1.0,1.0 | -0.319 |
| 916 | XLOC_009559 | chr12 | - | 115117309 | 115118951 | 115117309 | 115117456 | 115118683 | 1227 | 115118951 | 916 | 231,207 | 116,117 | 829,620 | 219,173 | 1376 | 149 | 1.48E-06 | 2.08E-05 | 0.177,0.161 | 0.291,0.28 | -0.116 |
| 2430 | XLOC_007680 | chr11 | - | 93465409 | 93469470 | 93465409 | 93466563 | 93467790 | 1227 | 93469470 | 2430 | 16,061,545 | 266,253 | 54,384,342 | 538,441 | 1376 | 149 | 1.75E-05 | 0.000185213 | 0.395,0.398 | 0.523,0.516 | -0.123 |
| 784 | XLOC_026831 | chr20 | + | 49197801 | 49201292 | 49197801 | 49197997 | 49199228 | 1231 | 49201292 | 784 | 363,348 | 76,69 | 694,589 | 16,19 | 1380 | 149 | 0 | 0 | 0.34,0.353 | 0.824,0.77 | -0.45 |
| 894 | XLOC_024079 | chr2 | - | 70454866 | 70456223 | 70454866 | 70454954 | 70456190 | 1236 | 70456223 | 894 | 603,607 | 54,64 | 991,788 | 51,39 | 1385 | 149 | 0.00016768 | 0.001331912 | 0.546,0.505 | 0.676,0.685 | -0.155 |
| 3873 | XLOC_046076 | chr9 | + | 131256839 | 131258420 | 131256839 | 131256947 | 131258184 | 1237 | 131258420 | 3873 | 25,32 | 4,0 | 16,23 | 0,0 | 1386 | 149 | 0.003107063 | 0.01572234 | 0.402,1.0 | 1.0,1.0 | -0.299 |
| 6724 | XLOC_020849 | chr19 | + | 798014 | 799443 | 798014 | 798171 | 799412 | 1241 | 799443 | 6724 | 10,13 | 2,1 | 15,4 | 0,0 | 1390 | 149 | 0.002074555 | 0.011212762 | 0.349,0.582 | 1.0,1.0 | -0.534 |
| 583 | XLOC_029694 | chr3 | + | 186505591 | 186510608 | 186505591 | 186505671 | 186506913 | 1242 | 186510608 | 583 | 29,633,032 | 725,736 | 75,745,846 | 788,691 | 1391 | 149 | 2.55E-13 | 1.17E-11 | 0.304,0.306 | 0.507,0.475 | -0.186 |
| 7632 | XLOC_001539 | chr1 | - | 226109903 | 226111471 | 226109903 | 226110083 | 226111336 | 1253 | 226111471 | 7632 | 31,47 | 6,0 | 41,31 | 0,1 | 1402 | 149 | 0.007906212 | 0.034042511 | 0.354,1.0 | 1.0,0.767 | -0.206 |
| 168 | XLOC_041504 | chr7 | - | 50515878 | 50517421 | 50515878 | 50515987 | 50517242 | 1255 | 50517421 | 168 | 232,190 | 1,5 | 280,306 | 0,1 | 1404 | 149 | 0.004074446 | 0.0196528 | 0.961,0.801 | 1.0,0.97 | -0.104 |
| 3071 | XLOC_024009 | chr2 | - | 39995563 | 39997259 | 39995563 | 39995641 | 39996897 | 1256 | 39997259 | 3071 | 62,83 | 4,3 | 163,96 | 0,1 | 1405 | 149 | 9.84E-05 | 0.000847487 | 0.622,0.746 | 1.0,0.911 | -0.272 |
| 686 | XLOC_028620 | chr22 | - | 29659823 | 29661657 | 29659823 | 29660207 | 29661467 | 1260 | 29661657 | 686 | 471,521 | 5,9 | 521,351 | 0,1 | 1409 | 149 | 4.29E-05 | 0.000412965 | 0.909,0.86 | 1.0,0.974 | -0.102 |
| 1426 | XLOC_037839 | chr6 | + | 43399879 | 43402586 | 43399879 | 43401098 | 43402358 | 1260 | 43402586 | 1426 | 0,4 | 15,15 | 26,22 | 10,5 | 1409 | 149 | 6.94E-11 | 2.17E-09 | 0.0,0.027 | 0.216,0.318 | -0.254 |
| 3182 | XLOC_045880 | chr9 | + | 22112318 | 22114029 | 22112318 | 22112394 | 22113664 | 1270 | 22114029 | 3182 | 54,92 | 18,11 | 42,51 | 1,3 | 1419 | 149 | 0.003979944 | 0.019309085 | 0.24,0.468 | 0.815,0.641 | -0.374 |
| 2057 | XLOC_009302 | chr12 | - | 32898492 | 32900297 | 32898492 | 32898738 | 32900009 | 1271 | 32900297 | 2057 | 163,145 | 4,6 | 156,97 | 2,0 | 1420 | 149 | 0.004338614 | 0.02065888 | 0.81,0.717 | 0.891,1.0 | -0.182 |
| 4171 | XLOC_041714 | chr7 | - | 130628824 | 130630594 | 130628824 | 130629206 | 130630478 | 1272 | 130630594 | 4171 | 55,75 | 11,10 | 80,59 | 0,1 | 1421 | 149 | 3.81E-08 | 7.46E-07 | 0.344,0.44 | 1.0,0.861 | -0.538 |
| 170 | XLOC_041504 | chr7 | - | 50515878 | 50517421 | 50515878 | 50515987 | 50517260 | 1273 | 50517421 | 170 | 234,192 | 19,14 | 286,309 | 1,5 | 1422 | 149 | 8.00E-08 | 1.49E-06 | 0.563,0.59 | 0.968,0.866 | -0.341 |
| 33 | XLOC_047846 | chrX | + | 134480943 | 134483239 | 134480943 | 134481397 | 134482678 | 1281 | 134483239 | 33 | 144,211 | 10,14 | 175,146 | 6,0 | 1430 | 149 | 0.003590989 | 0.017769987 | 0.6,0.611 | 0.752,1.0 | -0.271 |
| 71 | XLOC_046210 | chr9 | - | 27559304 | 27561647 | 27559304 | 27560297 | 27561582 | 1285 | 27561647 | 71 | 84,65 | 18,14 | 100,52 | 4,1 | 1434 | 149 | 1.08E-05 | 0.0001218 | 0.327,0.325 | 0.722,0.844 | -0.457 |
| 9682 | XLOC_013354 | chr14 | - | 36157664 | 36159209 | 36157664 | 36157746 | 36159041 | 1295 | 36159209 | 9682 | 18,21 | 3,2 | 9,4 | 1,0 | 1444 | 149 | 0.010822983 | 0.044164399 | 0.382,0.52 | 0.482,1.0 | -0.29 |
| 9210 | XLOC_014760 | chr15 | + | 89824995 | 89826481 | 89824995 | 89825066 | 89826362 | 1296 | 89826481 | 9210 | 67,73 | 2,6 | 43,51 | 0,1 | 1445 | 149 | 0.000900367 | 0.005586944 | 0.775,0.556 | 1.0,0.84 | -0.254 |
| 1717 | XLOC_014760 | chr15 | + | 89824995 | 89826481 | 89824995 | 89825066 | 89826366 | 1300 | 89826481 | 1717 | 548,573 | 185,177 | 636,471 | 96,82 | 1449 | 149 | 9.28E-07 | 1.37E-05 | 0.233,0.25 | 0.405,0.371 | -0.147 |
| 3436 | XLOC_000375 | chr1 | + | 94016503 | 94020392 | 94016503 | 94016662 | 94017965 | 1303 | 94020392 | 3436 | 880,833 | 40,33 | 1,056,707 | 8,8 | 1452 | 149 | 2.72E-09 | 6.51E-08 | 0.693,0.721 | 0.931,0.901 | -0.209 |
| 354 | XLOC_035165 | chr5 | - | 177656940 | 177658524 | 177656940 | 177657100 | 177658405 | 1305 | 177658524 | 354 | 320,350 | 29,31 | 320,210 | 12,12 | 1454 | 149 | 0.010496889 | 0.043095224 | 0.531,0.536 | 0.732,0.642 | -0.153 |
| 2386 | XLOC_005079 | chr10 | + | 124187791 | 124193261 | 124187791 | 124187832 | 124189139 | 1307 | 124193261 | 2386 | 38,64 | 14,16 | 352,210 | 15,24 | 1456 | 149 | 0.000133058 | 0.001093141 | 0.217,0.29 | 0.706,0.472 | -0.335 |
| 116 | XLOC_044185 | chr8 | + | 145721843 | 145724223 | 145721843 | 145722836 | 145724152 | 1316 | 145724223 | 116 | 214,194 | 263,239 | 411,264 | 180,132 | 1465 | 149 | 1.87E-12 | 7.68E-11 | 0.076,0.076 | 0.188,0.169 | -0.102 |
| 1288 | XLOC_001539 | chr1 | - | 226109903 | 226111471 | 226109903 | 226110083 | 226111400 | 1317 | 226111471 | 1288 | 478,521 | 281,301 | 546,472 | 90,85 | 1466 | 149 | 0 | 0 | 0.147,0.15 | 0.381,0.361 | -0.223 |
| 147 | XLOC_041749 | chr7 | - | 141356343 | 141364040 | 141356343 | 141362651 | 141363971 | 1320 | 141364040 | 147 | 236,176 | 67,83 | 167,117 | 22,20 | 1469 | 149 | 4.18E-05 | 0.000403157 | 0.263,0.177 | 0.435,0.372 | -0.183 |
| 8132 | XLOC_021853 | chr19 | - | 52566421 | 52570863 | 52566421 | 52566425 | 52567746 | 1321 | 52570863 | 8132 | 8,17 | 2,1 | 28,6 | 0,0 | 1470 | 149 | 0.00200704 | 0.010910479 | 0.288,0.633 | 1.0,1.0 | -0.539 |
| 2749 | XLOC_001174 | chr1 | - | 86851140 | 86852769 | 86851140 | 86851273 | 86852597 | 1324 | 86852769 | 2749 | 241,199 | 41,48 | 337,263 | 18,7 | 1473 | 149 | 8.07E-09 | 1.76E-07 | 0.373,0.295 | 0.654,0.792 | -0.389 |
| 5327 | XLOC_029391 | chr3 | + | 42702976 | 42704619 | 42702976 | 42703124 | 42704465 | 1341 | 42704619 | 5327 | 1,0 | 0,2 | 2,11 | 0,0 | 1490 | 149 | 1.13E-05 | 0.000126229 | 1.0,0.0 | 1.0,1.0 | -0.5 |
| 4983 | XLOC_041515 | chr7 | - | 64465336 | 64467093 | 64465336 | 64465394 | 64466918 | 1346 | 64467093 | 4983 | 2,9 | 0,3 | 12,9 | 0,0 | 1495 | 149 | 0.002457581 | 0.012888383 | 1.0,0.23 | 1.0,1.0 | -0.385 |
| 10857 | XLOC_030086 | chr3 | - | 156868070 | 156869719 | 156868070 | 156868170 | 156869517 | 1347 | 156869719 | 10857 | 160,173 | 8,1 | 113,89 | 0,0 | 1496 | 149 | 0.000117269 | 0.000982904 | 0.666,0.945 | 1.0,1.0 | -0.195 |
| 13453 | XLOC_018153 | chr17 | + | 48828107 | 48834841 | 48828107 | 48828213 | 48829560 | 1347 | 48834841 | 13453 | 558,559 | 29,18 | 714,561 | 6,0 | 1496 | 149 | 9.34E-10 | 2.45E-08 | 0.657,0.756 | 0.922,1.0 | -0.255 |
| 3322 | XLOC_000882 | chr1 | - | 10516563 | 10523266 | 10516563 | 10521759 | 10523114 | 1355 | 10523266 | 3322 | 75,83 | 96,109 | 127,109 | 54,37 | 1504 | 149 | 1.08E-10 | 3.25E-09 | 0.072,0.07 | 0.189,0.226 | -0.137 |
| 3032 | XLOC_041344 | chr7 | + | 150066759 | 150071492 | 150066759 | 150066957 | 150068316 | 1359 | 150071492 | 3032 | 12,011,078 | 74,86 | 28,122,133 | 76,55 | 1508 | 149 | 3.79E-10 | 1.06E-08 | 0.616,0.553 | 0.785,0.793 | -0.205 |
| 3250 | XLOC_030082 | chr3 | - | 155538792 | 155546166 | 155538792 | 155544608 | 155545970 | 1362 | 155546166 | 3250 | 468,452 | 10,10 | 561,467 | 6,1 | 1511 | 149 | 0.003219657 | 0.016199224 | 0.822,0.817 | 0.902,0.979 | -0.121 |
| 80 | XLOC_046076 | chr9 | + | 131256788 | 131258420 | 131256788 | 131256947 | 131258319 | 1372 | 131258420 | 80 | 225,211 | 96,92 | 210,144 | 55,31 | 1521 | 149 | 0.001240189 | 0.007314714 | 0.187,0.183 | 0.272,0.313 | -0.107 |
| 622 | XLOC_029391 | chr3 | + | 42702976 | 42704619 | 42702976 | 42703124 | 42704503 | 1379 | 42704619 | 622 | 36,16 | 22,25 | 99,55 | 33,14 | 1528 | 149 | 6.48E-05 | 0.000588046 | 0.138,0.059 | 0.226,0.277 | -0.153 |
| 8421 | XLOC_016718 | chr16 | - | 46989257 | 46991132 | 46989257 | 46989534 | 46990918 | 1384 | 46991132 | 8421 | 311,267 | 10,15 | 308,211 | 2,2 | 1533 | 149 | 7.16E-05 | 0.000638235 | 0.751,0.634 | 0.937,0.911 | -0.232 |
| 510 | XLOC_032278 | chr4 | + | 15685395 | 15692034 | 15685395 | 15688771 | 15690173 | 1402 | 15692034 | 510 | 372,323 | 21,12 | 292,262 | 1,0 | 1551 | 149 | 7.65E-10 | 2.03E-08 | 0.63,0.721 | 0.966,1.0 | -0.307 |
| 2176 | XLOC_009511 | chr12 | - | 104166091 | 104171859 | 104166091 | 104169408 | 104170821 | 1413 | 104171859 | 2176 | 139,162 | 3,2 | 58,75 | 0,0 | 1562 | 149 | 0.004181535 | 0.020025712 | 0.815,0.885 | 1.0,1.0 | -0.15 |
| 37 | XLOC_047663 | chrX | + | 23801916 | 23803546 | 23801916 | 23802000 | 23803444 | 1444 | 23803546 | 37 | 34,923,365 | 422,439 | 93,147,635 | 702,600 | 1593 | 149 | 8.79E-06 | 0.000101559 | 0.436,0.418 | 0.554,0.543 | -0.121 |
| 2520 | XLOC_013112 | chr14 | + | 50790662 | 50798961 | 50790662 | 50790878 | 50792326 | 1448 | 50798961 | 2520 | 84,68 | 3,2 | 74,35 | 0,0 | 1597 | 149 | 0.000556525 | 0.003773308 | 0.723,0.76 | 1.0,1.0 | -0.258 |
| 14834 | XLOC_000882 | chr1 | - | 10516563 | 10523266 | 10516563 | 10521759 | 10523221 | 1462 | 10523266 | 14834 | 158,171 | 6,0 | 171,118 | 0,0 | 1611 | 149 | 0.000843771 | 0.005284514 | 0.709,1.0 | 1.0,1.0 | -0.145 |
| 153 | XLOC_041655 | chr7 | - | 102918691 | 102920833 | 102918691 | 102918793 | 102920279 | 1486 | 102920833 | 153 | 133,157 | 5,4 | 93,70 | 0,0 | 1635 | 149 | 3.36E-05 | 0.000331979 | 0.708,0.782 | 1.0,1.0 | -0.255 |
| 9311 | XLOC_041183 | chr7 | + | 92078065 | 92080074 | 92078065 | 92078527 | 92080014 | 1487 | 92080074 | 9311 | 45,35 | 4,8 | 35,18 | 0,0 | 1636 | 149 | 1.37E-07 | 2.43E-06 | 0.506,0.285 | 1.0,1.0 | -0.605 |
| 856 | XLOC_024364 | chr2 | - | 201724402 | 201726189 | 201724402 | 201724469 | 201725960 | 1491 | 201726189 | 856 | 15,211,374 | 203,203 | 27,772,354 | 157,140 | 1640 | 149 | 3.82E-12 | 1.46E-10 | 0.405,0.381 | 0.616,0.604 | -0.217 |
| 765 | XLOC_026983 | chr20 | - | 34326889 | 34328519 | 34326889 | 34326939 | 34328446 | 1507 | 34328519 | 765 | 25,752,838 | 97,76 | 42,363,267 | 61,49 | 1656 | 149 | 8.46E-07 | 1.27E-05 | 0.705,0.771 | 0.862,0.857 | -0.121 |
| 3417 | XLOC_004807 | chr10 | + | 30726104 | 30728203 | 30726104 | 30726335 | 30727844 | 1509 | 30728203 | 3417 | 52,52 | 64,77 | 101,55 | 24,29 | 1658 | 149 | 1.46E-07 | 2.58E-06 | 0.068,0.057 | 0.274,0.146 | -0.148 |
| 5661 | XLOC_034684 | chr5 | + | 134124202 | 134125912 | 134124202 | 134124281 | 134125795 | 1514 | 134125912 | 5661 | 31,24 | 17,16 | 78,48 | 22,14 | 1663 | 149 | 0.011908241 | 0.048010303 | 0.14,0.118 | 0.241,0.235 | -0.109 |
| 263 | XLOC_038373 | chr6 | - | 99912955 | 99914639 | 99912955 | 99913012 | 99914547 | 1535 | 99914639 | 263 | 49,33 | 19,11 | 62,34 | 7,5 | 1684 | 149 | 0.003864576 | 0.018866023 | 0.186,0.21 | 0.439,0.376 | -0.209 |
| 2793 | XLOC_005193 | chr10 | - | 38115792 | 38120240 | 38115792 | 38118616 | 38120159 | 1543 | 38120240 | 2793 | 156,186 | 4,3 | 306,244 | 0,1 | 1692 | 149 | 0.000430655 | 0.003022932 | 0.774,0.845 | 1.0,0.956 | -0.168 |
| 330 | XLOC_037686 | chr6 | + | 26391030 | 26397897 | 26391030 | 26391057 | 26392602 | 1545 | 26397897 | 330 | 37,31 | 29,34 | 22,34 | 6,4 | 1694 | 149 | 0.000153331 | 0.001233193 | 0.101,0.074 | 0.244,0.428 | -0.248 |
| 52 | XLOC_046427 | chr9 | - | 130823531 | 130826179 | 130823531 | 130824105 | 130825652 | 1547 | 130826179 | 52 | 72,127 | 34,51 | 227,171 | 20,30 | 1696 | 149 | 9.49E-06 | 0.000108953 | 0.157,0.18 | 0.499,0.334 | -0.248 |
| 1964 | XLOC_038295 | chr6 | - | 47251673 | 47254331 | 47251673 | 47252128 | 47253679 | 1551 | 47254331 | 1964 | 94,92 | 4,1 | 85,83 | 0,1 | 1700 | 149 | 0.005034794 | 0.023373344 | 0.673,0.89 | 1.0,0.879 | -0.158 |
| 1094 | XLOC_001359 | chr1 | - | 155881033 | 155885755 | 155881033 | 155884111 | 155885663 | 1552 | 155885755 | 1094 | 20,192,176 | 175,148 | 30,522,365 | 79,49 | 1701 | 149 | 1.11E-16 | 7.47E-15 | 0.503,0.563 | 0.772,0.809 | -0.258 |
| 62 | XLOC_046327 | chr9 | - | 98268259 | 98271538 | 98268259 | 98268881 | 98270442 | 1561 | 98271538 | 62 | 30,37 | 3,4 | 18,11 | 1,0 | 1710 | 149 | 0.003301675 | 0.016570559 | 0.466,0.446 | 0.611,1.0 | -0.349 |
| 2438 | XLOC_007664 | chr11 | - | 82901839 | 82903858 | 82901839 | 82902210 | 82903777 | 1567 | 82903858 | 2438 | 146,107 | 7,13 | 156,72 | 1,2 | 1716 | 149 | 0.0009123 | 0.005641159 | 0.644,0.417 | 0.931,0.758 | -0.314 |
| 2730 | XLOC_009361 | chr12 | - | 51373597 | 51375586 | 51373597 | 51373841 | 51375421 | 1580 | 51375586 | 2730 | 491,464 | 22,16 | 353,310 | 3,2 | 1729 | 149 | 1.04E-05 | 0.000117258 | 0.658,0.714 | 0.91,0.93 | -0.234 |
| 13903 | XLOC_000333 | chr1 | + | 67450254 | 67459428 | 67450254 | 67450484 | 67452085 | 1601 | 67459428 | 13903 | 139,166 | 9,0 | 143,123 | 0,0 | 1750 | 149 | 0.000289174 | 0.002146091 | 0.568,1.0 | 1.0,1.0 | -0.216 |
| 12435 | XLOC_001160 | chr1 | - | 78413137 | 78414985 | 78413137 | 78413237 | 78414839 | 1602 | 78414985 | 12435 | 524,427 | 0,13 | 1,033,798 | 0,3 | 1751 | 149 | 0.000567667 | 0.003823157 | 1.0,0.736 | 1.0,0.958 | -0.111 |

| | | | | | | | | | | | | | | | | | | | | | | |
|---|---|---|---|---|---|---|---|---|---|---|---|---|---|---|---|---|---|---|---|---|---|---|
| 2008 | XLOC_007426 | chr11 | - | 10871986 | 10879728 | 10871986 | 10877804 | 10879408 | 1604 | 10879728 | 2008 | 771,668 | 97,90 | 940,652 | 54,59 | 1753 | 149 | 0.001785775 | 0.009933242 | 0.403,0.387 | 0.597,0.484 | -0.145 |
| 13705 | XLOC_018094 | chr17 | + | 42264359 | 42266956 | 42264359 | 42264768 | 42266389 | 1621 | 42266956 | 13705 | 137,175 | 17,14 | 194,158 | 0,0 | 1770 | 149 | 3.75E-13 | 1.69E-11 | 0.404,0.513 | 1.0,1.0 | -0.541 |
| 3286 | XLOC_028580 | chr22 | - | 20386516 | 20388425 | 20386516 | 20386675 | 20388299 | 1624 | 20388425 | 3286 | 21,13 | 2,1 | 9,3 | 0,0 | 1773 | 149 | 0.000369543 | 0.002653305 | 0.469,0.522 | 1.0,1.0 | -0.504 |
| 2996 | XLOC_023692 | chr2 | + | 160619390 | 160625224 | 160619390 | 160619504 | 160621136 | 1632 | 160625224 | 2996 | 228,184 | 167,135 | 340,216 | 102,71 | 1781 | 149 | 2.20E-08 | 4.51E-07 | 0.103,0.102 | 0.218,0.203 | -0.108 |
| 3474 | XLOC_000525 | chr1 | + | 153609047 | 153610924 | 153609047 | 153609129 | 153610770 | 1641 | 153610924 | 3474 | 13,051,339 | 408,369 | 24,491,972 | 346,334 | 1790 | 149 | 2.75E-08 | 5.54E-07 | 0.21,0.232 | 0.371,0.33 | -0.13 |
| 1189 | XLOC_020232 | chr18 | - | 47794348 | 47797910 | 47794348 | 47796188 | 47797838 | 1650 | 47797910 | 1189 | 592,606 | 16,25 | 1,501,968 | 18,7 | 1799 | 149 | 9.32E-06 | 0.000107136 | 0.754,0.668 | 0.874,0.92 | -0.186 |
| 204 | XLOC_041217 | chr7 | + | 99950995 | 99952863 | 99950995 | 99951106 | 99952765 | 1659 | 99952863 | 204 | 400,450 | 12,20 | 447,341 | 11,0 | 1808 | 149 | 0.006870588 | 0.03023207 | 0.733,0.65 | 0.77,1.0 | -0.194 |
| 13207 | XLOC_020109 | chr18 | + | 51795969 | 51797855 | 51795969 | 51796069 | 51797729 | 1660 | 51797855 | 13207 | 150,146 | 5,0 | 88,68 | 0,0 | 1809 | 149 | 0.003014983 | 0.015316764 | 0.712,1.0 | 1.0,1.0 | -0.144 |
| 1031 | XLOC_023585 | chr2 | + | 97858717 | 97860491 | 97858717 | 97858790 | 97860462 | 1672 | 97860491 | 1031 | 8,7 | 11,18 | 11,7 | 4,1 | 1821 | 149 | 0.002304832 | 0.012228076 | 0.056,0.031 | 0.184,0.364 | -0.231 |
| 1034 | XLOC_023585 | chr2 | + | 97873670 | 97875446 | 97873670 | 97873743 | 97875417 | 1674 | 97875446 | 1034 | 14,14 | 4,1 | 29,27 | 1,0 | 1823 | 149 | 0.000233856 | 0.001773767 | 0.222,0.534 | 0.703,1.0 | -0.473 |
| 4338 | XLOC_041489 | chr7 | - | 44116107 | 44118410 | 44116107 | 44116228 | 44117920 | 1692 | 44118410 | 4338 | 3,4 | 1,2 | 11,19 | 1,0 | 1841 | 149 | 0.000271359 | 0.0020288 | 0.195,0.139 | 0.471,1.0 | -0.569 |
| 2876 | XLOC_001549 | chr1 | - | 228580948 | 228584748 | 228580948 | 228582953 | 228584647 | 1694 | 228584748 | 2876 | 115,119 | 152,129 | 618,533 | 149,91 | 1843 | 149 | 0 | 0 | 0.058,0.069 | 0.251,0.321 | -0.223 |
| 2863 | XLOC_007334 | chr11 | + | 124492611 | 124494897 | 124492611 | 124493129 | 124494826 | 1697 | 124494897 | 2863 | 112,112 | 25,25 | 264,257 | 29,28 | 1846 | 149 | 0.001855168 | 0.010234584 | 0.266,0.266 | 0.424,0.426 | -0.159 |
| 4518 | XLOC_038093 | chr6 | - | 10400680 | 10402843 | 10400680 | 10401024 | 10402724 | 1700 | 10402843 | 4518 | 81,63 | 1,4 | 85,50 | 0,0 | 1849 | 149 | 0.000166437 | 0.001323661 | 0.867,0.559 | 1.0,1.0 | -0.287 |
| 1934 | XLOC_013242 | chr14 | + | 94565637 | 94569277 | 94565637 | 94567117 | 94568822 | 1705 | 94569277 | 1934 | 52,84 | 3,9 | 67,56 | 0,0 | 1854 | 149 | 1.47E-07 | 2.59E-06 | 0.582,0.429 | 1.0,1.0 | -0.495 |
| 2226 | XLOC_000852 | chr1 | - | 3653484 | 3656951 | 3653484 | 3655077 | 3656796 | 1719 | 3656951 | 2226 | 290,293 | 29,23 | 534,372 | 16,12 | 1868 | 149 | 2.36E-05 | 0.000240895 | 0.444,0.504 | 0.727,0.712 | -0.246 |
| 2329 | XLOC_024177 | chr2 | - | 110300589 | 110303814 | 110300589 | 110301901 | 110303626 | 1725 | 110303814 | 2329 | 309,327 | 260,299 | 931,705 | 107,89 | 1874 | 149 | 0 | 0 | 0.086,0.08 | 0.409,0.386 | -0.315 |
| 6427 | XLOC_023633 | chr2 | + | 113414698 | 113416671 | 113414698 | 113414818 | 113416550 | 1732 | 113416671 | 6427 | 232,240 | 4,3 | 367,274 | 0,0 | 1881 | 149 | 0.000211681 | 0.001630085 | 0.821,0.864 | 1.0,1.0 | -0.157 |
| 1971 | XLOC_009588 | chr12 | - | 122237752 | 122239714 | 122237752 | 122237862 | 122239612 | 1750 | 122239714 | 1971 | 100,84 | 20,32 | 140,102 | 13,10 | 1899 | 149 | 0.000221391 | 0.001695608 | 0.282,0.171 | 0.458,0.445 | -0.225 |
| 2545 | XLOC_007687 | chr11 | - | 96120693 | 96123028 | 96120693 | 96120777 | 96122528 | 1751 | 96123028 | 2545 | 374,320 | 11,10 | 447,294 | 0,3 | 1900 | 149 | 1.53E-05 | 0.000164818 | 0.727,0.715 | 1.0,0.885 | -0.222 |
| 3547 | XLOC_048083 | chrX | - | 130919580 | 130928143 | 130919580 | 130926262 | 130928021 | 1759 | 130928143 | 3547 | 0,4 | 3,2 | 15,3 | 0,0 | 1908 | 149 | 1.14E-12 | 4.87E-11 | 0.0,0.135 | 1.0,1.0 | -0.932 |
| 1839 | XLOC_016566 | chr16 | - | 1413208 | 1416396 | 1413208 | 1414256 | 1416018 | 1762 | 1416396 | 1839 | 8,811,066 | 33,30 | 933,790 | 4,8 | 1911 | 149 | 5.84E-08 | 1.12E-06 | 0.675,0.735 | 0.948,0.885 | -0.211 |
| 2205 | XLOC_030086 | chr3 | - | 156870824 | 156876764 | 156870824 | 156874872 | 156876654 | 1782 | 156876764 | 2205 | 634,646 | 46,56 | 787,597 | 28,12 | 1931 | 149 | 6.60E-06 | 7.88E-05 | 0.515,0.471 | 0.684,0.793 | -0.246 |
| 2710 | XLOC_011299 | chr13 | + | 39616241 | 39618318 | 39616241 | 39616442 | 39618226 | 1784 | 39618318 | 2710 | 296,275 | 53,66 | 393,297 | 29,26 | 1933 | 149 | 8.34E-07 | 1.25E-05 | 0.301,0.243 | 0.511,0.468 | -0.218 |
| 2097 | XLOC_030015 | chr3 | - | 126178495 | 126181063 | 126178495 | 126178580 | 126180377 | 1797 | 126181063 | 2097 | 699,639 | 57,36 | 631,579 | 14,8 | 1946 | 149 | 1.04E-07 | 1.89E-06 | 0.484,0.576 | 0.775,0.847 | -0.281 |
| 8135 | XLOC_000282 | chr1 | + | 46466190 | 46468555 | 46466190 | 46466651 | 46468476 | 1825 | 46468555 | 8135 | 38,56 | 18,12 | 11,7 | 0,0 | 1974 | 149 | 5.21E-06 | 6.37E-05 | 0.137,0.26 | 1.0,1.0 | -0.801 |
| 2288 | XLOC_004977 | chr10 | + | 96361284 | 96373815 | 96361284 | 96367783 | 96369609 | 1826 | 96373815 | 2288 | 150,93 | 26,8 | 166,116 | 9,1 | 1975 | 149 | 0.004798335 | 0.022461678 | 0.303,0.467 | 0.582,0.897 | -0.355 |
| 2953 | XLOC_000866 | chr1 | - | 6649817 | 6655617 | 6649817 | 6653718 | 6655544 | 1826 | 6655617 | 2953 | 141,104 | 40,41 | 342,312 | 61,33 | 1975 | 149 | 0.000135059 | 0.001107689 | 0.21,0.161 | 0.297,0.416 | -0.171 |
| 2775 | XLOC_037728 | chr6 | + | 28248696 | 28251986 | 28248696 | 28249695 | 28251552 | 1857 | 28251986 | 2775 | 51,21 | 20,12 | 84,70 | 14,15 | 2006 | 149 | 0.003996602 | 0.019369944 | 0.159,0.115 | 0.308,0.257 | -0.145 |
| 830 | XLOC_014710 | chr15 | + | 75498339 | 75506632 | 75498339 | 75501217 | 75503089 | 1872 | 75506632 | 830 | 105,127 | 45,40 | 271,230 | 22,16 | 2021 | 149 | 1.72E-12 | 7.21E-11 | 0.147,0.19 | 0.476,0.515 | -0.327 |
| 11386 | XLOC_005106 | chr10 | + | 135213032 | 135215749 | 135213032 | 135213157 | 135215032 | 1875 | 135215749 | 11386 | 36,48 | 0,4 | 38,27 | 0,0 | 2024 | 149 | 0.00298677 | 0.015195301 | 1.0,0.469 | 1.0,1.0 | -0.266 |
| 262 | XLOC_038373 | chr6 | - | 99850415 | 99852578 | 99850415 | 99850586 | 99852478 | 1892 | 99852578 | 262 | 24,452,523 | 71,43 | 29,072,374 | 29,23 | 2041 | 149 | 4.48E-05 | 0.000428578 | 0.715,0.811 | 0.88,0.883 | -0.118 |
| 645 | XLOC_034841 | chr5 | - | 1571956 | 1574410 | 1571956 | 1572401 | 1574296 | 1895 | 1574410 | 645 | 321,314 | 36,23 | 768,573 | 9,10 | 2044 | 149 | 1.35E-11 | 4.76E-10 | 0.394,0.499 | 0.862,0.807 | -0.388 |
| 14160 | XLOC_021620 | chr19 | - | 19102148 | 19104549 | 19102148 | 19102551 | 19104446 | 1895 | 19104549 | 14160 | 172,157 | 5,1 | 278,203 | 0,0 | 2044 | 149 | 0.0001542 | 0.001238772 | 0.715,0.92 | 1.0,1.0 | -0.182 |
| 2994 | XLOC_013457 | chr14 | - | 74759450 | 74761901 | 74759450 | 74759951 | 74761850 | 1899 | 74761901 | 2994 | 159,161 | 30,31 | 247,186 | 16,12 | 2048 | 149 | 1.49E-05 | 0.000161104 | 0.278,0.274 | 0.529,0.53 | -0.254 |
| 2333 | XLOC_005106 | chr10 | + | 135213032 | 135215749 | 135213032 | 135213123 | 135215032 | 1909 | 135215749 | 2333 | 538,517 | 89,90 | 435,415 | 35,32 | 2058 | 149 | 1.45E-05 | 0.00015729 | 0.304,0.294 | 0.474,0.484 | -0.18 |
| 2234 | XLOC_009218 | chr12 | - | 6571410 | 6574107 | 6571410 | 6572135 | 6574055 | 1920 | 6574107 | 2234 | 399,434 | 4,5 | 218,119 | 0,0 | 2069 | 149 | 0.00033998 | 0.002466118 | 0.878,0.862 | 1.0,1.0 | -0.13 |
| 829 | XLOC_014710 | chr15 | + | 75498339 | 75506632 | 75498339 | 75501165 | 75503089 | 1924 | 75506632 | 829 | 121,140 | 17,25 | 285,251 | 3,6 | 2073 | 149 | 9.03E-11 | 2.78E-09 | 0.338,0.287 | 0.872,0.75 | -0.498 |
| 2472 | XLOC_014740 | chr15 | + | 83423380 | 83425537 | 83423380 | 83423536 | 83425481 | 1945 | 83425537 | 2472 | 183,158 | 23,55 | 353,225 | 28,11 | 2094 | 149 | 0.000227956 | 0.001734604 | 0.361,0.17 | 0.473,0.593 | -0.267 |
| 451 | XLOC_041145 | chr7 | + | 75042066 | 75044301 | 75042066 | 75042210 | 75044162 | 1952 | 75044301 | 451 | 475,513 | 52,53 | 660,535 | 49,23 | 2101 | 149 | 0.008801105 | 0.037214973 | 0.393,0.407 | 0.489,0.623 | -0.156 |
| 4339 | XLOC_041489 | chr7 | - | 44116107 | 44118410 | 44116107 | 44116354 | 44118338 | 1984 | 44118410 | 4339 | 6,9 | 4,1 | 14,13 | 0,0 | 2133 | 149 | 7.02E-10 | 1.88E-08 | 0.095,0.386 | 1.0,1.0 | -0.76 |
| 1650 | XLOC_015065 | chr15 | - | 100267570 | 100272212 | 100267570 | 100269936 | 100271922 | 1986 | 100272212 | 1650 | 200,196 | 12,22 | 213,173 | 5,2 | 2135 | 149 | 0.000152287 | 0.001226197 | 0.538,0.383 | 0.748,0.858 | -0.342 |
| 8730 | XLOC_028432 | chr22 | + | 28318005 | 28322853 | 28318005 | 28318329 | 28320334 | 2005 | 28322853 | 8730 | 16,10 | 2,1 | 32,21 | 1,0 | 2154 | 149 | 0.000683386 | 0.004464138 | 0.356,0.409 | 0.689,1.0 | -0.462 |
| 7151 | XLOC_020886 | chr19 | + | 2340895 | 2343290 | 2340895 | 2341012 | 2343055 | 2043 | 2343290 | 7151 | 35,28 | 3,1 | 16,15 | 0,0 | 2192 | 149 | 0.000124802 | 0.00103681 | 0.442,0.656 | 1.0,1.0 | -0.451 |
| 3450 | XLOC_013273 | chr14 | + | 104151322 | 104153548 | 104151322 | 104151373 | 104153417 | 2044 | 104153548 | 3450 | 826,814 | 12,8 | 667,543 | 0,0 | 2193 | 149 | 3.01E-07 | 5.01E-06 | 0.824,0.874 | 1.0,1.0 | -0.151 |
| 2473 | XLOC_048000 | chrX | - | 71822968 | 71825250 | 71822968 | 71823070 | 71825120 | 2050 | 71825250 | 2473 | 197,164 | 57,61 | 336,245 | 52,48 | 2199 | 149 | 0.000395536 | 0.002814186 | 0.19,0.154 | 0.305,0.257 | -0.109 |
| 2809 | XLOC_005113 | chr10 | - | 1094803 | 1102662 | 1094803 | 1100185 | 1102248 | 2063 | 1102662 | 2809 | 79,67 | 4,2 | 23,49 | 1,0 | 2212 | 149 | 0.004704326 | 0.022080157 | 0.571,0.693 | 0.608,1.0 | -0.172 |
| 696 | XLOC_028550 | chr22 | + | 50644745 | 50647145 | 50644745 | 50644891 | 50646964 | 2073 | 50647145 | 696 | 624,581 | 11,5 | 576,408 | 0,0 | 2222 | 149 | 1.31E-06 | 1.87E-05 | 0.792,0.886 | 1.0,1.0 | -0.161 |
| 2963 | XLOC_001594 | chr1 | - | 245009905 | 245012158 | 245009905 | 245010008 | 245012089 | 2081 | 245012158 | 2963 | 17,121,698 | 108,87 | 18,711,531 | 20,17 | 2230 | 149 | 0 | 0 | 0.514,0.566 | 0.862,0.857 | -0.319 |
| 1930 | XLOC_028602 | chr22 | - | 24176704 | 24179341 | 24176704 | 24177166 | 24179250 | 2084 | 24179341 | 1930 | 314,342 | 2,3 | 622,431 | 0,0 | 2233 | 149 | 0.00690241 | 0.030349546 | 0.913,0.884 | 1.0,1.0 | -0.101 |
| 3381 | XLOC_000367 | chr1 | + | 90470691 | 90473309 | 90470691 | 90470803 | 90472903 | 2100 | 90473309 | 3381 | 409,360 | 82,64 | 702,473 | 71,38 | 2249 | 149 | 1.54E-05 | 0.000165061 | 0.248,0.271 | 0.396,0.452 | -0.165 |
| 211 | XLOC_041489 | chr7 | - | 44116107 | 44118410 | 44116107 | 44116228 | 44118338 | 2110 | 44118410 | 211 | 91,89 | 36,28 | 203,167 | 19,19 | 2259 | 149 | 2.12E-07 | 3.63E-06 | 0.143,0.173 | 0.413,0.367 | -0.232 |
| 2912 | XLOC_041167 | chr7 | + | 86815144 | 86817617 | 86815144 | 86815296 | 86817407 | 2111 | 86817617 | 2912 | 242,181 | 89,88 | 650,496 | 98,76 | 2260 | 149 | 7.41E-11 | 2.31E-09 | 0.152,0.119 | 0.304,0.301 | -0.167 |
| 14663 | XLOC_001218 | chr1 | - | 109468392 | 109477564 | 109468392 | 109475123 | 109477241 | 2118 | 109477564 | 14663 | 75,53 | 2,9 | 26,12 | 0,0 | 2267 | 149 | 1.97E-06 | 2.68E-05 | 0.711,0.279 | 1.0,1.0 | -0.505 |
| 4697 | XLOC_035132 | chr5 | - | 159824448 | 159827393 | 159824448 | 159824522 | 159826645 | 2123 | 159827393 | 4697 | 4,8 | 2,0 | 6,3 | 0,0 | 2272 | 149 | 0.011724937 | 0.047365928 | 0.116,1.0 | 1.0,1.0 | -0.442 |
| 8050 | XLOC_007221 | chr11 | + | 74336452 | 74339062 | 74336452 | 74336618 | 74338742 | 2124 | 74339062 | 8050 | 31,31 | 1,9 | 30,29 | 1,0 | 2273 | 149 | 0.000705296 | 0.004568727 | 0.67,0.184 | 0.663,1.0 | -0.404 |
| 14157 | XLOC_021620 | chr19 | - | 19101568 | 19104549 | 19101568 | 19101958 | 19104085 | 2127 | 19104549 | 14157 | 146,178 | 4,5 | 321,225 | 0,0 | 2276 | 149 | 1.05E-06 | 1.53E-05 | 0.705,0.7 | 1.0,1.0 | -0.298 |
| 912 | XLOC_023912 | chr2 | + | 241500746 | 241503936 | 241500746 | 241500940 | 241503090 | 2150 | 241503936 | 912 | 66,71 | 3,1 | 23,44 | 0,0 | 2299 | 149 | 0.001508762 | 0.008604871 | 0.588,0.821 | 1.0,1.0 | -0.296 |
| 1765 | XLOC_000593 | chr1 | + | 166818174 | 166825519 | 166818174 | 166819660 | 166821821 | 2161 | 166825519 | 1765 | 865,783 | 177,162 | 1,055,822 | 128,106 | 2310 | 149 | 0.000225992 | 0.001726079 | 0.24,0.238 | 0.347,0.333 | -0.101 |
| 1563 | XLOC_028432 | chr22 | + | 28318005 | 28322853 | 28318005 | 28318166 | 28320334 | 2168 | 28322853 | 1563 | 206,238 | 139,142 | 517,384 | 91,76 | 2317 | 149 | 3.33E-16 | 2.11E-14 | 0.087,0.097 | 0.268,0.245 | -0.165 |
| 2926 | XLOC_000859 | chr1 | - | 6311397 | 6314207 | 6311397 | 6311590 | 6313777 | 2187 | 6314207 | 2926 | 119,134 | 9,16 | 303,253 | 5,9 | 2336 | 149 | 0.000358515 | 0.002581291 | 0.458,0.348 | 0.794,0.642 | -0.315 |
| 1440 | XLOC_021370 | chr19 | + | 54969492 | 54973077 | 54969492 | 54969700 | 54971945 | 2245 | 54973077 | 1440 | 49,565,077 | 108,92 | 69,024,875 | 50,25 | 2394 | 149 | 1.88E-12 | 7.68E-11 | 0.741,0.775 | 0.896,0.924 | -0.152 |
| 13292 | XLOC_021000 | chr19 | + | 13261200 | 13265742 | 13261200 | 13261507 | 13263757 | 2250 | 13265742 | 13292 | 124,109 | 0,5 | 117,102 | 0,0 | 2399 | 149 | 0.001221736 | 0.007222508 | 1.0,0.575 | 1.0,1.0 | -0.213 |
| 615 | XLOC_029460 | chr3 | + | 56600977 | 56604695 | 56600977 | 56601081 | 56603340 | 2259 | 56604695 | 615 | 24,19 | 7,8 | 75,30 | 4,2 | 2408 | 149 | 0.000137043 | 0.001120054 | 0.175,0.128 | 0.537,0.481 | -0.358 |
| 2654 | XLOC_041202 | chr7 | + | 99081652 | 99085371 | 99081652 | 99081766 | 99084098 | 2332 | 99085371 | 2654 | 61,59 | 8,5 | 80,67 | 1,0 | 2481 | 149 | 1.97E-06 | 2.68E-05 | 0.314,0.415 | 0.828,1.0 | -0.549 |

| | | | | | | | | | | | | | | | | | | | | | | |
|---|---|---|---|---|---|---|---|---|---|---|---|---|---|---|---|---|---|---|---|---|---|---|
| 3383 | XLOC_000367 | chr1 | + | 90470691 | 90475845 | 90470691 | 90473309 | 90475646 | 2337 | 90475845 | 3383 | 263,261 | 129,116 | 688,559 | 73,62 | 2486 | 149 | 0 | 0 | 0.109,0.119 | 0.361,0.351 | -0.242 |
| 14053 | XLOC_001012 | chr1 | - | 35643589 | 35652723 | 35643589 | 35650194 | 35652547 | 2353 | 35652723 | 14053 | 116,112 | 21,22 | 220,159 | 10,17 | 2502 | 149 | 0.005493512 | 0.025165381 | 0.248,0.233 | 0.567,0.358 | -0.222 |
| 14856 | XLOC_000928 | chr1 | - | 21099943 | 21102498 | 21099943 | 21100103 | 21102460 | 2357 | 21102498 | 14856 | 286,284 | 1,5 | 210,175 | 0,0 | 2506 | 149 | 0.000634199 | 0.00419915 | 0.944,0.772 | 1.0,1.0 | -0.142 |
| 14989 | XLOC_000367 | chr1 | + | 90470691 | 90473309 | 90470691 | 90470803 | 90473167 | 2364 | 90473309 | 14989 | 392,322 | 6,2 | 644,443 | 0,2 | 2513 | 149 | 0.002197751 | 0.011779446 | 0.795,0.905 | 1.0,0.929 | -0.114 |
| 3080 | XLOC_018253 | chr17 | + | 74722930 | 74725876 | 74722930 | 74723295 | 74725771 | 2476 | 74725876 | 3080 | 226,281 | 34,57 | 279,209 | 29,17 | 2625 | 149 | 0.002699848 | 0.013938283 | 0.274,0.219 | 0.353,0.411 | -0.136 |
| 521 | XLOC_037689 | chr6 | + | 26428870 | 26432480 | 26428870 | 26429013 | 26431499 | 2486 | 26432480 | 521 | 128,140 | 2,5 | 59,30 | 0,0 | 2635 | 149 | 2.15E-05 | 0.000222053 | 0.784,0.613 | 1.0,1.0 | -0.301 |
| 3170 | XLOC_021620 | chr19 | - | 19101568 | 19104549 | 19101568 | 19101958 | 19104446 | 2488 | 19104549 | 3170 | 26,722,451 | 31,21 | 44,163,386 | 6,3 | 2637 | 149 | 1.32E-12 | 5.61E-11 | 0.83,0.868 | 0.977,0.985 | -0.132 |
| 752 | XLOC_046346 | chr9 | - | 100444481 | 100447322 | 100444481 | 100444712 | 100447204 | 2492 | 100447322 | 752 | 120,76 | 10,2 | 139,133 | 0,3 | 2641 | 149 | 0.007989665 | 0.03434948 | 0.404,0.682 | 1.0,0.714 | -0.314 |
| 3171 | XLOC_021620 | chr19 | - | 19101568 | 19104549 | 19101568 | 19101958 | 19104456 | 2498 | 19104549 | 3171 | 26,862,465 | 13,34 | 44,493,416 | 4,0 | 2647 | 149 | 2.01E-13 | 9.32E-12 | 0.921,0.803 | 0.984,1.0 | -0.13 |
| 13861 | XLOC_018253 | chr17 | + | 74722930 | 74725876 | 74722930 | 74723260 | 74725771 | 2511 | 74725876 | 13861 | 200,247 | 65,68 | 242,188 | 25,34 | 2660 | 149 | 0.000576021 | 0.003872038 | 0.147,0.169 | 0.352,0.236 | -0.136 |
| 742 | XLOC_027692 | chr21 | + | 38117028 | 38123284 | 38117028 | 38117437 | 38119965 | 2528 | 38123284 | 742 | 50,18 | 56,65 | 393,257 | 49,24 | 2677 | 149 | 0 | 0 | 0.047,0.015 | 0.309,0.373 | -0.31 |
| 3934 | XLOC_046188 | chr9 | - | 14677060 | 14680160 | 14677060 | 14677498 | 14680030 | 2532 | 14680160 | 3934 | 3,5 | 0,4 | 2,3 | 0,0 | 2681 | 149 | 0.000120259 | 0.001005576 | 1.0,0.065 | 1.0,1.0 | -0.468 |
| 72 | XLOC_046202 | chr9 | - | 21995081 | 22014972 | 21995081 | 22006246 | 22008796 | 2550 | 22014972 | 72 | 231,215 | 205,219 | 724,627 | 159,121 | 2699 | 149 | 0 | 0 | 0.059,0.051 | 0.201,0.222 | -0.157 |
| 1234 | XLOC_000461 | chr1 | + | 145281954 | 145293580 | 145281954 | 145290511 | 145293113 | 2602 | 145293580 | 1234 | 381,433 | 24,17 | 810,658 | 9,17 | 2751 | 149 | 0.001243068 | 0.00732104 | 0.462,0.58 | 0.83,0.677 | -0.233 |
| 2998 | XLOC_023692 | chr2 | + | 160621136 | 160625224 | 160621136 | 160621185 | 160623794 | 2609 | 160625224 | 2998 | 310,324 | 153,123 | 631,483 | 100,65 | 2758 | 149 | 2.68E-13 | 1.22E-11 | 0.099,0.125 | 0.254,0.286 | -0.158 |
| 1713 | XLOC_014793 | chr15 | + | 100252594 | 100257268 | 100252594 | 100253040 | 100255672 | 2632 | 100257268 | 1713 | 16,191,594 | 15,15 | 813,733 | 0,1 | 2781 | 149 | 1.03E-06 | 1.50E-05 | 0.853,0.851 | 1.0,0.975 | -0.136 |
| 389 | XLOC_034819 | chr5 | + | 179198147 | 179204444 | 179198147 | 179198244 | 179200895 | 2651 | 179204444 | 389 | 151,140 | 59,49 | 215,176 | 26,27 | 2800 | 149 | 1.12E-06 | 1.62E-05 | 0.12,0.132 | 0.306,0.258 | -0.156 |
| 12545 | XLOC_001307 | chr1 | - | 150944087 | 150947366 | 150944087 | 150944437 | 150947094 | 2657 | 150947366 | 12545 | 1,1 | 0,2 | 13,7 | 0,0 | 2806 | 149 | 1.17E-05 | 0.000130781 | 1.0,0.026 | 1.0,1.0 | -0.487 |
| 64 | XLOC_046278 | chr9 | - | 74975543 | 74978522 | 74975543 | 74975703 | 74978385 | 2682 | 74978522 | 64 | 642,573 | 203,227 | 1,008,942 | 140,155 | 2831 | 149 | 6.55E-11 | 2.06E-09 | 0.143,0.117 | 0.275,0.242 | -0.129 |
| 2643 | XLOC_018004 | chr17 | + | 29546022 | 29548947 | 29546022 | 29546136 | 29548867 | 2731 | 29548947 | 2643 | 81,91 | 64,80 | 198,160 | 50,43 | 2880 | 149 | 5.28E-10 | 1.44E-08 | 0.061,0.056 | 0.17,0.161 | -0.107 |
| 11792 | XLOC_020127 | chr18 | + | 59942606 | 59945686 | 59942606 | 59942706 | 59945448 | 2742 | 59945686 | 11792 | 2,4 | 2,0 | 11,6 | 0,0 | 2891 | 149 | 0.00057956 | 0.00389027 | 0.049,1.0 | 1.0,1.0 | -0.476 |
| 1249 | XLOC_013404 | chr14 | - | 59937884 | 59942724 | 59937884 | 59939808 | 59942586 | 2778 | 59942724 | 1249 | 240,255 | 43,53 | 618,469 | 29,7 | 2927 | 149 | 8.65E-08 | 1.60E-06 | 0.221,0.197 | 0.52,0.773 | -0.438 |
| 2706 | XLOC_020183 | chr18 | - | 21226152 | 21229157 | 21226152 | 21226251 | 21229046 | 2795 | 21229157 | 2706 | 236,243 | 8,10 | 540,395 | 0,2 | 2944 | 149 | 1.45E-08 | 3.01E-07 | 0.599,0.552 | 1.0,0.909 | -0.379 |
| 1190 | XLOC_020232 | chr18 | - | 47794348 | 47799108 | 47794348 | 47796188 | 47799046 | 2858 | 47799108 | 1190 | 766,806 | 47,29 | 21,521,462 | 22,13 | 3007 | 149 | 3.32E-10 | 9.35E-09 | 0.447,0.579 | 0.829,0.848 | -0.326 |
| 1838 | XLOC_001538 | chr1 | - | 226066919 | 226071110 | 226066919 | 226067110 | 226070003 | 2893 | 226071110 | 1838 | 76,80 | 53,55 | 189,130 | 41,33 | 3042 | 149 | 7.88E-08 | 1.47E-06 | 0.066,0.067 | 0.184,0.162 | -0.106 |
| 1071 | XLOC_027825 | chr21 | - | 47737925 | 47744469 | 47737925 | 47738134 | 47741059 | 2925 | 47744469 | 1071 | 226,208 | 4,3 | 97,65 | 0,0 | 3074 | 149 | 0.000107486 | 0.000911724 | 0.733,0.771 | 1.0,1.0 | -0.248 |
| 540 | XLOC_030162 | chr3 | - | 195695089 | 195698315 | 195695089 | 195695256 | 195698193 | 2937 | 195698315 | 540 | 320,365 | 15,14 | 566,462 | 5,2 | 3086 | 149 | 7.26E-07 | 1.11E-05 | 0.507,0.557 | 0.845,0.918 | -0.349 |
| 13938 | XLOC_045867 | chr9 | + | 6493825 | 6497360 | 6493825 | 6493932 | 6496959 | 3027 | 6497360 | 13938 | 150,131 | 5,7 | 296,210 | 1,3 | 3176 | 149 | 0.00131108 | 0.007664064 | 0.585,0.468 | 0.933,0.767 | -0.324 |
| 87 | XLOC_045946 | chr9 | + | 80863677 | 80866959 | 80863677 | 80863789 | 80866823 | 3034 | 80866959 | 87 | 267,265 | 49,51 | 309,234 | 12,16 | 3183 | 149 | 2.15E-06 | 2.89E-05 | 0.203,0.196 | 0.547,0.406 | -0.277 |
| 3162 | XLOC_001270 | chr1 | - | 143426483 | 143465438 | 143426483 | 143462343 | 143465379 | 3036 | 143465438 | 3162 | 11,20 | 42,32 | 51,39 | 26,9 | 3185 | 149 | 1.29E-09 | 3.31E-08 | 0.012,0.028 | 0.084,0.169 | -0.106 |
| 12889 | XLOC_018612 | chr17 | - | 46629388 | 46632980 | 46629388 | 46629849 | 46632892 | 3043 | 46632980 | 12889 | 2,1 | 0,2 | 7,2 | 0,0 | 3192 | 149 | 3.11E-05 | 0.000309365 | 1.0,0.023 | 1.0,1.0 | -0.489 |
| 259 | XLOC_038404 | chr6 | - | 112397524 | 112400957 | 112397524 | 112397712 | 112400835 | 3123 | 112400957 | 259 | 195,188 | 26,41 | 594,454 | 53,45 | 3272 | 149 | 0.002232018 | 0.011904377 | 0.255,0.173 | 0.338,0.315 | -0.113 |
| 2653 | XLOC_041202 | chr7 | + | 99074025 | 99085371 | 99074025 | 99074103 | 99077283 | 3180 | 99085371 | 2653 | 300,273 | 37,31 | 250,206 | 4,5 | 3329 | 149 | 2.60E-08 | 5.26E-07 | 0.266,0.283 | 0.737,0.648 | -0.418 |
| 1369 | XLOC_018010 | chr17 | + | 30696617 | 30700472 | 30696617 | 30697101 | 30700417 | 3316 | 30700472 | 1369 | 13,621,337 | 12,13 | 19,551,517 | 2,1 | 3465 | 149 | 7.65E-08 | 1.44E-06 | 0.83,0.816 | 0.977,0.985 | -0.158 |
| 639 | XLOC_029298 | chr3 | + | 9471510 | 9475634 | 9471510 | 9471735 | 9475053 | 3318 | 9475634 | 639 | 172,179 | 30,35 | 367,301 | 37,18 | 3467 | 149 | 0.000668162 | 0.004376836 | 0.198,0.18 | 0.299,0.418 | -0.169 |
| 1815 | XLOC_024041 | chr2 | - | 55514857 | 55518911 | 55514857 | 55515491 | 55518811 | 3320 | 55518911 | 1815 | 10,831,111 | 4,14 | 729,571 | 2,0 | 3469 | 149 | 0.000740508 | 0.004736499 | 0.921,0.773 | 0.94,1.0 | -0.123 |
| 5637 | XLOC_027757 | chr21 | - | 33735464 | 33739064 | 33735464 | 33735638 | 33738962 | 3324 | 33739064 | 5637 | 88,91 | 4,0 | 319,208 | 0,0 | 3473 | 149 | 0.003497322 | 0.017393519 | 0.486,1.0 | 1.0,1.0 | -0.257 |
| 5524 | XLOC_028550 | chr22 | + | 50639436 | 50643453 | 50639436 | 50640019 | 50643347 | 3328 | 50643453 | 5524 | 11,11 | 5,0 | 25,13 | 0,0 | 3477 | 149 | 0.000390067 | 0.002783689 | 0.086,1.0 | 1.0,1.0 | -0.457 |
| 5636 | XLOC_027757 | chr21 | - | 33735464 | 33739064 | 33735464 | 33735581 | 33738926 | 3345 | 33739064 | 5636 | 82,94 | 6,0 | 290,220 | 1,0 | 3494 | 149 | 0.002954314 | 0.015062736 | 0.368,1.0 | 0.925,1.0 | -0.279 |
| 11894 | XLOC_001574 | chr1 | - | 234541648 | 234545319 | 234541648 | 234541845 | 234545261 | 3416 | 234545319 | 11894 | 56,50 | 1,3 | 54,48 | 0,0 | 3565 | 149 | 0.000144524 | 0.001171707 | 0.701,0.411 | 1.0,1.0 | -0.444 |
| 2819 | XLOC_018612 | chr17 | - | 46629388 | 46632980 | 46629388 | 46629452 | 46632892 | 3440 | 46632980 | 2819 | 175,151 | 4,2 | 346,241 | 0,0 | 3589 | 149 | 1.79E-05 | 0.00018802 | 0.645,0.758 | 1.0,1.0 | -0.298 |
| 298 | XLOC_037999 | chr6 | + | 133846291 | 133853416 | 133846291 | 133846392 | 133849862 | 3470 | 133853416 | 298 | 697,676 | 16,24 | 24,321,871 | 7,3 | 3619 | 149 | 1.75E-14 | 9.28E-13 | 0.642,0.537 | 0.935,0.963 | -0.36 |
| 14885 | XLOC_028488 | chr22 | + | 39126616 | 39130583 | 39126616 | 39126664 | 39130154 | 3490 | 39130583 | 14885 | 146,145 | 4,4 | 192,206 | 0,0 | 3639 | 149 | 1.08E-06 | 1.56E-05 | 0.599,0.597 | 1.0,1.0 | -0.402 |
| 1368 | XLOC_018010 | chr17 | + | 30696617 | 30700472 | 30696617 | 30696801 | 30700417 | 3616 | 30700472 | 1368 | 22,732,136 | 17,27 | 31,732,462 | 4,0 | 3765 | 149 | 2.97E-12 | 1.17E-10 | 0.841,0.758 | 0.969,1.0 | -0.185 |
| 13688 | XLOC_009610 | chr12 | - | 125267228 | 125271049 | 125267228 | 125267357 | 125270986 | 3629 | 125271049 | 13688 | 14,581,497 | 36,53 | 28,812,167 | 30,38 | 3778 | 149 | 0.000280786 | 0.00209043 | 0.615,0.527 | 0.791,0.692 | -0.171 |
| 12013 | XLOC_007176 | chr11 | + | 66606336 | 66610820 | 66606336 | 66606678 | 66610319 | 3641 | 66610820 | 12013 | 8,15 | 1,1 | 7,8 | 0,0 | 3790 | 149 | 0.010987712 | 0.044737042 | 0.239,0.371 | 1.0,1.0 | -0.695 |
| 8832 | XLOC_021777 | chr19 | - | 46213918 | 46220594 | 46213918 | 46216831 | 46220522 | 3691 | 46220594 | 8832 | 60,40 | 19,28 | 42,49 | 10,6 | 3840 | 149 | 0.005048859 | 0.023424781 | 0.109,0.053 | 0.14,0.241 | -0.11 |
| 8662 | XLOC_046352 | chr9 | - | 101494195 | 101498905 | 101494195 | 101494835 | 101498563 | 3728 | 101498905 | 8662 | 27,25 | 0,3 | 15,12 | 0,0 | 3877 | 149 | 0.004978531 | 0.023189906 | 1.0,0.243 | 1.0,1.0 | -0.379 |
| 2908 | XLOC_000386 | chr1 | + | 100609624 | 100613882 | 100609624 | 100609699 | 100613449 | 3750 | 100613882 | 2908 | 819,706 | 36,40 | 12,791,008 | 12,12 | 3899 | 149 | 4.04E-11 | 1.34E-09 | 0.465,0.403 | 0.803,0.762 | -0.348 |
| 421 | XLOC_034621 | chr5 | + | 94030535 | 94042283 | 94030535 | 94033622 | 94037399 | 3777 | 94042283 | 421 | 959,921 | 12,7 | 831,708 | 1,4 | 3926 | 149 | 0.006940346 | 0.030459451 | 0.752,0.833 | 0.969,0.87 | -0.127 |
| 445 | XLOC_043980 | chr8 | + | 27369352 | 27373330 | 27369352 | 27369427 | 27373234 | 3807 | 27373330 | 445 | 15,27 | 1,1 | 22,20 | 0,1 | 3956 | 149 | 0.010458071 | 0.04294834 | 0.361,0.504 | 1.0,0.43 | -0.282 |
| 2117 | XLOC_011529 | chr13 | - | 79884552 | 79893014 | 79884552 | 79889128 | 79892955 | 3827 | 79893014 | 2117 | 900,897 | 51,52 | 14,951,298 | 59,44 | 3976 | 149 | 0.010555983 | 0.043298808 | 0.398,0.393 | 0.487,0.525 | -0.11 |
| 2118 | XLOC_011529 | chr13 | - | 79884552 | 79894845 | 79884552 | 79889125 | 79892955 | 3830 | 79894845 | 2118 | 900,897 | 7,17 | 14,951,298 | 6,7 | 3979 | 149 | 0.006104504 | 0.027465407 | 0.828,0.664 | 0.903,0.874 | -0.143 |
| 9648 | XLOC_001538 | chr1 | - | 226066919 | 226071110 | 226066919 | 226067110 | 226071029 | 3919 | 226071110 | 9648 | 76,80 | 4,0 | 187,114 | 0,0 | 4068 | 149 | 0.002532262 | 0.013211356 | 0.41,1.0 | 1.0,1.0 | -0.295 |
| 1275 | XLOC_018752 | chr17 | - | 76670134 | 76676426 | 76670134 | 76672251 | 76676271 | 4020 | 76676426 | 1275 | 19,891,824 | 137,115 | 33,382,526 | 158,95 | 4169 | 149 | 0.001308428 | 0.00765173 | 0.342,0.362 | 0.43,0.487 | -0.107 |
| 1625 | XLOC_008774 | chr12 | + | 8925916 | 8933335 | 8925916 | 8926062 | 8930185 | 4123 | 8933335 | 1625 | 32,072,964 | 35,23 | 94,837,377 | 39,29 | 4272 | 149 | 5.84E-05 | 0.000538176 | 0.762,0.818 | 0.895,0.899 | -0.107 |
| 1752 | XLOC_032346 | chr4 | + | 71655211 | 71659621 | 71655211 | 71655309 | 71659481 | 4172 | 71659621 | 1752 | 368,339 | 3,4 | 421,275 | 0,0 | 4321 | 149 | 8.99E-05 | 0.000781375 | 0.809,0.745 | 1.0,1.0 | -0.223 |
| 1596 | XLOC_021902 | chr19 | - | 58317797 | 58324788 | 58317797 | 58320471 | 58324661 | 4190 | 58324788 | 1596 | 90,82 | 32,25 | 151,68 | 12,11 | 4339 | 149 | 0.000280475 | 0.002089215 | 0.088,0.101 | 0.302,0.175 | -0.144 |
| 1753 | XLOC_032346 | chr4 | + | 71655211 | 71659621 | 71655211 | 71655309 | 71659501 | 4192 | 71659621 | 1753 | 368,342 | 35,19 | 422,275 | 9,4 | 4341 | 149 | 1.60E-05 | 0.000170708 | 0.265,0.382 | 0.617,0.702 | -0.336 |
| 3343 | XLOC_032661 | chr4 | - | 83816844 | 83821376 | 83816844 | 83816927 | 83821229 | 4302 | 83821376 | 3343 | 902,805 | 26,35 | 1,065,695 | 15,8 | 4451 | 149 | 0.000147459 | 0.001193448 | 0.537,0.435 | 0.704,0.744 | -0.238 |
| 6471 | XLOC_016746 | chr16 | - | 58193491 | 58198051 | 58193491 | 58193655 | 58197957 | 4302 | 58198051 | 6471 | 251,270 | 4,3 | 258,161 | 0,0 | 4451 | 149 | 2.20E-05 | 0.000226607 | 0.677,0.751 | 1.0,1.0 | -0.286 |
| 859 | XLOC_024344 | chr2 | - | 190631241 | 190636628 | 190631241 | 190632114 | 190636433 | 4319 | 190636628 | 859 | 494,565 | 2,4 | 408,370 | 0,0 | 4468 | 149 | 0.001381911 | 0.008008508 | 0.892,0.825 | 1.0,1.0 | -0.141 |
| 638 | XLOC_029298 | chr3 | + | 9470506 | 9475246 | 9470506 | 9470693 | 9475053 | 4360 | 9475246 | 638 | 229,242 | 19,24 | 464,385 | 21,12 | 4509 | 149 | 0.001144603 | 0.006829416 | 0.285,0.25 | 0.422,0.515 | -0.201 |
| 2119 | XLOC_011529 | chr13 | - | 79884552 | 79894845 | 79884552 | 79889128 | 79893669 | 4541 | 79894845 | 2119 | 10,991,077 | 12,22 | 18,421,662 | 8,14 | 4690 | 149 | 0.004673539 | 0.021957548 | 0.744,0.609 | 0.88,0.79 | -0.158 |
| 3460 | XLOC_014666 | chr15 | + | 64429766 | 64436417 | 64429766 | 64430296 | 64434901 | 4605 | 64436417 | 3460 | 747,755 | 4,4 | 706,426 | 0,0 | 4754 | 149 | 0.000261252 | 0.001964677 | 0.854,0.855 | 1.0,1.0 | -0.145 |

| | | | | | | | | | | | | | | | | | | | | | | |
|---|---|---|---|---|---|---|---|---|---|---|---|---|---|---|---|---|---|---|---|---|---|---|
| 3342 | XLOC_032661 | chr4 | - | 83813646 | 83821376 | 83813646 | 83816000 | 83821229 | 5229 | 83821376 | 3342 | 1,027,927 | 19,11 | 1,234,872 | 0,0 | 5378 | 149 | 3.50E-12 | 1.35E-10 | 0.6,0.7 | 1.0,1.0 | -0.35 |
| 3004 | XLOC_000907 | chr1 | - | 16944679 | 16952500 | 16944679 | 16946519 | 16952269 | 5750 | 16952500 | 3004 | 266,244 | 41,30 | 1,137,990 | 39,29 | 5899 | 149 | 4.44E-13 | 1.98E-11 | 0.141,0.17 | 0.424,0.463 | -0.288 |
| 6531 | XLOC_023378 | chr2 | + | 17935203 | 17941371 | 17935203 | 17935234 | 17941195 | 5961 | 17941371 | 6531 | 502,467 | 4,9 | 466,331 | 2,0 | 6110 | 149 | 0.001222763 | 0.007225559 | 0.754,0.559 | 0.85,1.0 | -0.268 |
| 2446 | XLOC_007563 | chr11 | - | 64537727 | 64544098 | 64537727 | 64537880 | 64543969 | 6089 | 64544098 | 2446 | 33,183,404 | 50,60 | 56,584,276 | 42,37 | 6238 | 149 | 4.89E-05 | 0.000462391 | 0.613,0.575 | 0.763,0.734 | -0.154 |
| 2860 | XLOC_001166 | chr1 | - | 85397044 | 85403560 | 85397044 | 85397251 | 85403437 | 6186 | 85403560 | 2860 | 48,58 | 9,16 | 152,113 | 9,6 | 6335 | 149 | 3.77E-05 | 0.000367525 | 0.111,0.079 | 0.284,0.307 | -0.2 |
| 2916 | XLOC_034863 | chr5 | - | 34159948 | 34176062 | 34159948 | 34169258 | 34175549 | 6291 | 34176062 | 2916 | 281,218 | 17,18 | 569,404 | 7,12 | 6440 | 149 | 0.000594902 | 0.003976242 | 0.277,0.219 | 0.653,0.438 | -0.297 |
| 6227 | XLOC_005203 | chr10 | - | 44063346 | 44070045 | 44063346 | 44063469 | 44069918 | 6449 | 44070045 | 6227 | 322,301 | 1,6 | 672,546 | 0,0 | 6598 | 149 | 1.54E-05 | 0.000165061 | 0.879,0.531 | 1.0,1.0 | -0.295 |
| 13267 | XLOC_034863 | chr5 | - | 34169131 | 34176062 | 34169131 | 34169258 | 34175926 | 6668 | 34176062 | 13267 | 290,222 | 3,5 | 566,408 | 1,1 | 6817 | 149 | 0.000854254 | 0.005338333 | 0.679,0.493 | 0.925,0.899 | -0.326 |
| 3323 | XLOC_000420 | chr1 | + | 111717500 | 111724940 | 111717500 | 111717585 | 111724808 | 7223 | 111724940 | 3323 | 528,541 | 245,234 | 1,252,966 | 46,46 | 7372 | 149 | 0 | 0 | 0.042,0.045 | 0.355,0.298 | -0.283 |
| 2045 | XLOC_001156 | chr1 | - | 75667868 | 75677237 | 75667868 | 75669518 | 75677152 | 7634 | 75677237 | 2045 | 28,35 | 1,1 | 43,41 | 0,0 | 7783 | 149 | 0.001449205 | 0.008330128 | 0.349,0.401 | 1.0,1.0 | -0.625 |
| 592 | XLOC_029597 | chr3 | + | 140681968 | 140689835 | 140681968 | 140682069 | 140689768 | 7699 | 140689835 | 592 | 26,602,571 | 163,161 | 63,785,124 | 96,78 | 7848 | 149 | 0 | 0 | 0.237,0.233 | 0.558,0.555 | -0.322 |
| 2071 | XLOC_014966 | chr15 | - | 70358352 | 70366946 | 70358352 | 70358557 | 70366841 | 8284 | 70366946 | 2071 | 283,260 | 23,18 | 16,821,316 | 47,34 | 8433 | 149 | 4.57E-06 | 5.70E-05 | 0.179,0.203 | 0.387,0.406 | -0.206 |
| 10422 | XLOC_014966 | chr15 | - | 70358352 | 70366946 | 70358352 | 70358554 | 70366841 | 8287 | 70366946 | 10422 | 280,255 | 8,6 | 16,491,297 | 9,4 | 8436 | 149 | 1.49E-05 | 0.000161104 | 0.382,0.429 | 0.764,0.851 | -0.402 |
| 10423 | XLOC_014966 | chr15 | - | 70358352 | 70366946 | 70358352 | 70358554 | 70366871 | 8317 | 70366946 | 10423 | 284,261 | 17,15 | 16,641,306 | 19,30 | 8466 | 149 | 3.73E-05 | 0.000365049 | 0.227,0.234 | 0.607,0.434 | -0.29 |
| 1587 | XLOC_044021 | chr8 | + | 42873500 | 42885639 | 42873500 | 42873637 | 42882571 | 8934 | 42885639 | 1587 | 13,551,430 | 0,6 | 1,013,691 | 0,0 | 9083 | 149 | 0.002239271 | 0.011938554 | 1.0,0.796 | 1.0,1.0 | -0.102 |
| 2911 | XLOC_041167 | chr7 | + | 86783705 | 86792933 | 86783705 | 86783844 | 86792810 | 8966 | 86792933 | 2911 | 837,755 | 24,11 | 13,051,013 | 10,9 | 9115 | 149 | 0.00415315 | 0.019950658 | 0.363,0.529 | 0.681,0.648 | -0.219 |
| 207 | XLOC_041204 | chr7 | + | 99158155 | 99174973 | 99158155 | 99158318 | 99169867 | 11549 | 99174973 | 207 | 29,642,914 | 31,16 | 29,022,408 | 2,9 | 11698 | 149 | 0.000472033 | 0.003280767 | 0.549,0.699 | 0.949,0.773 | -0.237 |
| 3146 | XLOC_011557 | chr13 | - | 111545040 | 111558471 | 111545040 | 111546740 | 111558379 | 11639 | 111558471 | 3146 | 96,159,414 | 16,22 | 1,374,811,163 | 3,0 | 11788 | 149 | 8.53E-12 | 3.09E-10 | 0.884,0.844 | 0.983,1.0 | -0.128 |
| 3148 | XLOC_011557 | chr13 | - | 111545374 | 111558471 | 111545374 | 111545610 | 111558379 | 12769 | 111558471 | 3148 | 1,122,510,896 | 181,183 | 1,589,712,896 | 101,72 | 12918 | 149 | 6.22E-14 | 3.07E-12 | 0.417,0.407 | 0.645,0.674 | -0.247 |
| 13872 | XLOC_038025 | chr6 | + | 147660472 | 147674509 | 147660472 | 147660520 | 147674458 | 13938 | 147674509 | 13872 | 17,13 | 1,1 | 58,30 | 0,0 | 14087 | 149 | 2.32E-06 | 3.09E-05 | 0.152,0.121 | 1.0,1.0 | -0.863 |

**Table S1 (3) The list of overlapped retained introns in RBM17- and SAP30BP-knockdown HeLa cells**

See rMATS (http://rnaseq-mats.sourceforge.net/) for the format of this table.

| ID | Gene ID | Chromosome | Strand | RI-Exon Start | RI-Exon End | Upstream ES | Upstream EE | Downstream ES | Intron (nt) | Downstream EE | ID.1 | IJC SAMPLE 1 | SJC SAMPLE 1 | IJC SAMPLE 2 | SJC SAMPLE 2 | Inc. Form (nt) | Skip Form (nt) | P Value | FDR | Inc. Level 1 | Inc. Level 2 | Inc. Level.Differ. |
|---|---|---|---|---|---|---|---|---|---|---|---|---|---|---|---|---|---|---|---|---|---|---|
| 352 | XLOC_035171 | chr5 | - | 179044799 | 179045061 | 179044799 | 179044933 | 179044989 | 56 | 179045061 | 352 | 257,227 | 17,401,655 | 778,733 | 14,231,535 | 205 | 149 | 0 | 0 | 0.097,0.091 | 0.284,0.258 | -0.177 |
| 9885 | XLOC_009615 | chr12 | - | 132624967 | 132625280 | 132624967 | 132625100 | 132625164 | 64 | 132625280 | 9885 | 1,1 | 1,3 | 4,2 | 1,0 | 213 | 149 | 9.22E-05 | 0.001935548 | 0.412,0.189 | 0.737,1.0 | -0.568 |
| 3274 | XLOC_018288 | chr17 | + | 79661844 | 79662097 | 79661844 | 79661883 | 79661953 | 70 | 79662097 | 3274 | 254,254 | 394,338 | 501,467 | 501,335 | 219 | 149 | 0.000131947 | 0.002590726 | 0.305,0.338 | 0.405,0.487 | -0.124 |
| 15126 | XLOC_000825 | chr1 | - | 1233392 | 1233801 | 1233392 | 1233493 | 1233564 | 71 | 1233801 | 15126 | 7,6 | 3,3 | 31,51 | 0,2 | 220 | 149 | 0.000124699 | 0.00247131 | 0.612,0.575 | 1.0,0.945 | -0.379 |
| 2483 | XLOC_007374 | chr11 | - | 800761 | 801120 | 800761 | 800912 | 800984 | 72 | 801120 | 2483 | 16,9 | 19,13 | 38,41 | 8,10 | 221 | 149 | 1.93E-06 | 8.40E-05 | 0.362,0.318 | 0.762,0.734 | -0.408 |
| 9037 | XLOC_016435 | chr16 | + | 67201205 | 67201502 | 67201205 | 67201305 | 67201377 | 72 | 67201502 | 9037 | 22,28 | 4,4 | 66,53 | 0,2 | 221 | 149 | 0.000421669 | 0.006381025 | 0.788,0.825 | 1.0,0.947 | -0.167 |
| 10309 | XLOC_013110 | chr14 | + | 50246312 | 50246528 | 50246312 | 50246393 | 50246465 | 72 | 50246528 | 10309 | 64,66 | 179,203 | 123,134 | 138,138 | 221 | 149 | 9.46E-10 | 1.34E-07 | 0.194,0.18 | 0.375,0.396 | -0.199 |
| 234 | XLOC_041771 | chr7 | - | 150773999 | 150774323 | 150773999 | 150774114 | 150774187 | 73 | 150774323 | 234 | 127,112 | 177,191 | 163,213 | 192,165 | 222 | 149 | 0.005520697 | 0.04474263 | 0.325,0.282 | 0.363,0.464 | -0.11 |
| 450 | XLOC_047682 | chrX | + | 44941820 | 44942034 | 44941820 | 44941885 | 44941959 | 74 | 44942034 | 450 | 4,0 | 3,12 | 19,14 | 6,4 | 223 | 149 | 0.000543375 | 0.007718076 | 0.471,0.0 | 0.679,0.7 | -0.454 |
| 95 | XLOC_044466 | chr8 | - | 145738024 | 145738521 | 145738024 | 145738154 | 145738229 | 75 | 145738521 | 95 | 97,99 | 272,272 | 229,239 | 234,192 | 224 | 149 | 1.09E-14 | 4.96E-12 | 0.192,0.195 | 0.394,0.453 | -0.23 |
| 2980 | XLOC_024343 | chr2 | - | 190618641 | 190619071 | 190618641 | 190618790 | 190618866 | 76 | 190619071 | 2980 | 3,0 | 32,30 | 8,5 | 28,17 | 225 | 149 | 0.000766191 | 0.010013069 | 0.058,0.0 | 0.159,0.163 | -0.132 |
| 3275 | XLOC_018288 | chr17 | + | 79663636 | 79664028 | 79663636 | 79663777 | 79663853 | 76 | 79664028 | 3275 | 79,51 | 376,389 | 170,195 | 451,484 | 225 | 149 | 1.74E-08 | 1.53E-06 | 0.122,0.08 | 0.2,0.211 | -0.105 |
| 1737 | XLOC_014574 | chr15 | + | 41106140 | 41106921 | 41106140 | 41106268 | 41106345 | 77 | 41106921 | 1737 | 116,149 | 115,111 | 159,193 | 54,47 | 226 | 149 | 1.13E-09 | 1.54E-07 | 0.399,0.469 | 0.66,0.73 | -0.261 |
| 3764 | XLOC_046457 | chr9 | - | 135772570 | 135772928 | 135772570 | 135772732 | 135772809 | 77 | 135772928 | 3764 | 16,3 | 61,57 | 30,17 | 58,57 | 226 | 149 | 0.003204065 | 0.029622336 | 0.147,0.034 | 0.254,0.164 | -0.119 |
| 13152 | XLOC_018790 | chr17 | - | 80018576 | 80018830 | 80018576 | 80018659 | 80018736 | 77 | 80018830 | 13152 | 4,7 | 6,4 | 46,62 | 2,0 | 226 | 149 | 1.59E-08 | 1.42E-06 | 0.305,0.536 | 0.938,1.0 | -0.548 |
| 13190 | XLOC_034753 | chr5 | + | 149747251 | 149747480 | 149747251 | 149747329 | 149747406 | 77 | 149747480 | 13190 | 7,2 | 8,0 | 8,8 | 0,0 | 226 | 149 | 0.000580889 | 0.008090486 | 0.366,1.0 | 1.0,1.0 | -0.317 |
| 14440 | XLOC_018386 | chr17 | - | 7224938 | 7225329 | 7224938 | 7225106 | 7225183 | 77 | 7225329 | 14440 | 20,19 | 104,113 | 59,41 | 103,116 | 226 | 149 | 6.43E-05 | 0.001458729 | 0.113,0.1 | 0.274,0.189 | -0.125 |
| 10097 | XLOC_007211 | chr11 | + | 71940967 | 71941312 | 71940967 | 71941063 | 71941141 | 78 | 71941312 | 10097 | 4,6 | 0,5 | 8,9 | 0,0 | 227 | 149 | 0.00188332 | 0.019935885 | 1.0,0.441 | 1.0,1.0 | -0.279 |
| 2892 | XLOC_016217 | chr16 | + | 2284859 | 2285144 | 2284859 | 2285002 | 2285081 | 79 | 2285144 | 2892 | 32,31 | 169,182 | 59,84 | 181,142 | 228 | 149 | 2.04E-05 | 0.000583276 | 0.11,0.1 | 0.176,0.279 | -0.122 |
| 6126 | XLOC_044184 | chr8 | + | 145693063 | 145693370 | 145693063 | 145693162 | 145693241 | 79 | 145693370 | 6126 | 11,22 | 126,91 | 28,33 | 70,74 | 228 | 149 | 8.96E-05 | 0.001902785 | 0.054,0.136 | 0.207,0.226 | -0.121 |
| 14657 | XLOC_006953 | chr11 | + | 828574 | 828963 | 828574 | 828772 | 828852 | 79 | 828963 | 14657 | 0,1 | 20,25 | 4,3 | 7,15 | 228 | 149 | 0.000413862 | 0.006300927 | 0.0,0.025 | 0.272,0.116 | -0.181 |
| 556 | XLOC_030002 | chr3 | - | 123674689 | 123674962 | 123674689 | 123674812 | 123674892 | 80 | 123674962 | 556 | 79,86 | 46,69 | 168,110 | 47,42 | 229 | 149 | 0.000214937 | 0.003835212 | 0.528,0.448 | 0.699,0.63 | -0.176 |
| 1060 | XLOC_021627 | chr19 | - | 19767468 | 19767725 | 19767468 | 19767575 | 19767655 | 80 | 19767725 | 1060 | 109,78 | 66,55 | 197,228 | 91,74 | 229 | 149 | 0.006383917 | 0.04947289 | 0.518,0.48 | 0.585,0.667 | -0.127 |
| 3737 | XLOC_046490 | chr9 | - | 139905424 | 139905752 | 139905424 | 139905557 | 139905637 | 80 | 139905752 | 3737 | 43,65 | 211,175 | 118,106 | 131,146 | 229 | 149 | 2.05E-09 | 2.49E-07 | 0.117,0.195 | 0.37,0.321 | -0.19 |
| 4311 | XLOC_041495 | chr7 | - | 44609407 | 44609832 | 44609407 | 44609502 | 44609582 | 80 | 44609832 | 4311 | 11,13 | 11,8 | 28,18 | 6,0 | 229 | 149 | 0.000477482 | 0.006986662 | 0.394,0.514 | 0.752,1.0 | -0.422 |
| 1779 | XLOC_000300 | chr1 | + | 53343353 | 53344440 | 53343353 | 53343515 | 53343596 | 81 | 53344440 | 1779 | 14,16 | 36,37 | 64,55 | 56,74 | 230 | 149 | 0.00271298 | 0.026145685 | 0.201,0.219 | 0.425,0.325 | -0.165 |
| 4220 | XLOC_045898 | chr9 | + | 35660561 | 35661014 | 35660561 | 35660855 | 35660936 | 81 | 35661014 | 4220 | 250,186 | 69,53 | 519,435 | 64,73 | 230 | 149 | 9.91E-05 | 0.002055803 | 0.701,0.695 | 0.84,0.794 | -0.119 |
| 1451 | XLOC_023617 | chr2 | + | 109367719 | 109368159 | 109367719 | 109367901 | 109367983 | 82 | 109368159 | 1451 | 6,14 | 181,158 | 34,40 | 163,139 | 231 | 149 | 2.85E-08 | 2.40E-06 | 0.021,0.054 | 0.119,0.157 | -0.101 |
| 1774 | XLOC_018149 | chr17 | + | 48452545 | 48453554 | 48452545 | 48453344 | 48453426 | 82 | 48453554 | 1774 | 15,17 | 55,96 | 47,35 | 67,65 | 231 | 149 | 1.68E-05 | 0.000493829 | 0.15,0.103 | 0.312,0.258 | -0.159 |
| 2660 | XLOC_011393 | chr13 | + | 115037658 | 115038417 | 115037658 | 115037956 | 115038038 | 82 | 115038417 | 2660 | 240,189 | 144,141 | 259,274 | 100,110 | 231 | 149 | 0.000417365 | 0.00634637 | 0.518,0.464 | 0.626,0.616 | -0.13 |
| 5705 | XLOC_028552 | chr22 | + | 50960771 | 50961002 | 50960771 | 50960844 | 50960926 | 82 | 50961002 | 5705 | 3,4 | 9,29 | 24,11 | 12,21 | 231 | 149 | 0.001161597 | 0.013714868 | 0.177,0.082 | 0.563,0.253 | -0.278 |
| 428 | XLOC_034609 | chr5 | + | 81600433 | 81601300 | 81600433 | 81600669 | 81600752 | 83 | 81601300 | 428 | 7,3 | 3,10 | 3,4 | 0,0 | 232 | 149 | 3.41E-07 | 1.97E-05 | 0.6,0.162 | 1.0,1.0 | -0.619 |
| 353 | XLOC_035168 | chr5 | - | 178029485 | 178030935 | 178029485 | 178030758 | 178030844 | 86 | 178030935 | 353 | 4,5 | 84,53 | 26,27 | 31,58 | 235 | 149 | 3.66E-10 | 5.56E-08 | 0.029,0.056 | 0.347,0.228 | -0.245 |
| 10550 | XLOC_028486 | chr22 | + | 39066888 | 39067297 | 39066888 | 39066994 | 39067080 | 86 | 39067297 | 10550 | 3,9 | 3,1 | 3,10 | 0,0 | 235 | 149 | 0.001915458 | 0.020204989 | 0.388,0.851 | 1.0,1.0 | -0.381 |
| 13221 | XLOC_016216 | chr16 | + | 2257035 | 2257408 | 2257035 | 2257111 | 2257197 | 86 | 2257408 | 13221 | 1,0 | 13,20 | 6,4 | 16,16 | 235 | 149 | 0.000303979 | 0.004975467 | 0.047,0.0 | 0.192,0.137 | -0.141 |
| 13813 | XLOC_021582 | chr19 | - | 14521027 | 14521417 | 14521027 | 14521146 | 14521232 | 86 | 14521417 | 13813 | 7,12 | 12,26 | 31,40 | 20,21 | 235 | 149 | 0.000224344 | 0.003951507 | 0.27,0.226 | 0.496,0.547 | -0.274 |
| 1193 | XLOC_024434 | chr2 | - | 230656581 | 230656942 | 230656581 | 230656781 | 230656868 | 87 | 230656942 | 1193 | 1,13 | 295,232 | 70,98 | 248,279 | 236 | 149 | 0 | 0 | 0.002,0.034 | 0.151,0.182 | -0.148 |
| 2509 | XLOC_007251 | chr11 | + | 93462779 | 93463135 | 93462779 | 93462947 | 93463035 | 88 | 93463135 | 2509 | 11,17 | 132,156 | 42,55 | 149,116 | 237 | 149 | 2.88E-08 | 2.40E-06 | 0.05,0.064 | 0.151,0.23 | -0.134 |
| 7042 | XLOC_029383 | chr3 | + | 40502898 | 40503152 | 40502898 | 40503010 | 40503098 | 88 | 40503152 | 7042 | 3,8 | 4,1 | 12,6 | 0,1 | 237 | 149 | 0.001727975 | 0.018712887 | 0.32,0.834 | 1.0,0.79 | -0.318 |
| 14289 | XLOC_000967 | chr1 | - | 27682878 | 27683175 | 27682878 | 27682991 | 27683080 | 89 | 27683175 | 14289 | 0,1 | 37,40 | 10,17 | 26,39 | 238 | 149 | 2.61E-09 | 3.09E-07 | 0.0,0.015 | 0.194,0.214 | -0.197 |
| 10085 | XLOC_013517 | chr14 | - | 96798641 | 96798989 | 96798641 | 96798747 | 96798837 | 90 | 96798989 | 10085 | 0,0 | 16,13 | 3,3 | 19,7 | 239 | 149 | 0.002239458 | 0.022608002 | 0.0,0.0 | 0.09,0.211 | -0.15 |
| 14721 | XLOC_007127 | chr11 | + | 63970483 | 63970700 | 63970483 | 63970506 | 63970596 | 90 | 63970700 | 14721 | 5,16 | 5,0 | 3,3 | 0,0 | 239 | 149 | 0.00393877 | 0.03462244 | 0.384,1.0 | 1.0,1.0 | -0.308 |
| 15925 | XLOC_001222 | chr1 | - | 109823391 | 109823873 | 109823391 | 109823668 | 109823758 | 90 | 109823873 | 15925 | 136,111 | 105,123 | 155,136 | 58,48 | 239 | 149 | 5.69E-07 | 3.01E-05 | 0.447,0.36 | 0.625,0.639 | -0.229 |
| 8724 | XLOC_021139 | chr19 | + | 36104769 | 36105036 | 36104769 | 36104802 | 36104896 | 94 | 36105036 | 8724 | 2,5 | 4,1 | 8,7 | 0,0 | 243 | 149 | 2.72E-05 | 0.00074809 | 0.235,0.754 | 1.0,1.0 | -0.506 |
| 9385 | XLOC_026746 | chr20 | + | 31015933 | 31016225 | 31015933 | 31016032 | 31016127 | 95 | 31016225 | 9385 | 31,42 | 4,5 | 39,37 | 2,0 | 244 | 149 | 0.004654716 | 0.039468 | 0.826,0.837 | 0.923,1.0 | -0.13 |
| 9975 | XLOC_007774 | chr11 | - | 134026646 | 134026990 | 134026646 | 134026817 | 134026912 | 95 | 134026990 | 9975 | 35,23 | 13,9 | 25,29 | 2,4 | 244 | 149 | 0.004400346 | 0.037822859 | 0.622,0.609 | 0.884,0.816 | -0.235 |
| 9756 | XLOC_014584 | chr15 | + | 41793383 | 41793757 | 41793383 | 41793564 | 41793660 | 96 | 41793757 | 9756 | 1,1 | 0,4 | 2,6 | 0,0 | 245 | 149 | 0.001886595 | 0.019956511 | 1.0,0.132 | 1.0,1.0 | -0.434 |
| 4931 | XLOC_034487 | chr5 | + | 1802195 | 1802488 | 1802195 | 1802335 | 1802434 | 99 | 1802488 | 4931 | 6,9 | 3,2 | 8,18 | 0,0 | 248 | 149 | 9.01E-05 | 0.001903505 | 0.546,0.73 | 1.0,1.0 | -0.362 |
| 15689 | XLOC_000426 | chr1 | + | 113217176 | 113217541 | 113217176 | 113217351 | 113217453 | 102 | 113217541 | 15689 | 23,52 | 77,77 | 38,41 | 36,38 | 251 | 149 | 0.002458306 | 0.024295557 | 0.151,0.286 | 0.385,0.39 | -0.169 |
| 8819 | XLOC_016786 | chr16 | - | 69402278 | 69402498 | 69402278 | 69402385 | 69402489 | 104 | 69402498 | 8819 | 2,0 | 2,6 | 19,11 | 12,5 | 253 | 149 | 0.001411653 | 0.015917608 | 0.371,0.0 | 0.483,0.564 | -0.338 |
| 14229 | XLOC_018153 | chr17 | + | 48827861 | 48828213 | 48827861 | 48828002 | 48828107 | 105 | 48828213 | 14229 | 915,899 | 219,188 | 11,811,140 | 58,62 | 254 | 149 | 0 | 0 | 0.71,0.737 | 0.923,0.915 | -0.196 |
| 7976 | XLOC_034626 | chr5 | + | 96092881 | 96093198 | 96092881 | 96093040 | 96093147 | 107 | 96093198 | 7976 | 10,5 | 6,2 | 10,8 | 0,0 | 256 | 149 | 1.19E-06 | 5.62E-05 | 0.492,0.593 | 1.0,1.0 | -0.458 |
| 5266 | XLOC_029841 | chr3 | - | 49153246 | 49153595 | 49153246 | 49153371 | 49153479 | 108 | 49153595 | 5266 | 8,3 | 40,40 | 21,11 | 46,35 | 257 | 149 | 0.002881118 | 0.027376988 | 0.104,0.042 | 0.209,0.154 | -0.108 |
| 11082 | XLOC_046244 | chr9 | - | 35812893 | 35813335 | 35812893 | 35813153 | 35813262 | 109 | 35813335 | 11082 | 9,2 | 3,7 | 4,4 | 1,0 | 258 | 149 | 0.00059843 | 0.008296397 | 0.634,0.142 | 0.698,1.0 | -0.461 |
| 15204 | XLOC_023671 | chr2 | + | 135920148 | 135920421 | 135920148 | 135920208 | 135920317 | 109 | 135920421 | 15204 | 9,11 | 3,4 | 15,13 | 0,2 | 258 | 149 | 0.005385784 | 0.043957116 | 0.634,0.614 | 1.0,0.79 | -0.271 |
| 3991 | XLOC_044466 | chr8 | - | 145737063 | 145737445 | 145737063 | 145737172 | 145737284 | 112 | 145737445 | 3991 | 6,6 | 6,4 | 22,23 | 4,3 | 261 | 149 | 0.002924851 | 0.027569316 | 0.363,0.461 | 0.758,0.814 | -0.374 |
| 12922 | XLOC_018124 | chr17 | + | 45696374 | 45696688 | 45696374 | 45696530 | 45696643 | 113 | 45696688 | 12922 | 64,61 | 21,23 | 46,44 | 0,9 | 262 | 149 | 0.002052757 | 0.021207126 | 0.634,0.601 | 1.0,0.735 | -0.25 |
| 13452 | XLOC_017993 | chr17 | + | 27896335 | 27896643 | 27896335 | 27896377 | 27896491 | 114 | 27896643 | 13452 | 8,15 | 3,1 | 6,10 | 0,0 | 263 | 149 | 0.005922452 | 0.046986036 | 0.602,0.895 | 1.0,1.0 | -0.252 |
| 15273 | XLOC_007445 | chr11 | - | 18320167 | 18320517 | 18320167 | 18320224 | 18320338 | 114 | 18320517 | 15273 | 4,2 | 4,0 | 4,2 | 0,0 | 263 | 149 | 0.005301958 | 0.043343504 | 0.362,1.0 | 1.0,1.0 | -0.319 |
| 1105 | XLOC_021261 | chr19 | + | 45567269 | 45567888 | 45567269 | 45567485 | 45567600 | 115 | 45567888 | 1105 | 46,56 | 117,123 | 125,103 | 92,131 | 264 | 149 | 3.99E-05 | 0.001015057 | 0.182,0.204 | 0.434,0.307 | -0.177 |
| 7542 | XLOC_038282 | chr6 | - | 43536217 | 43536474 | 43536217 | 43536332 | 43536447 | 115 | 43536474 | 7542 | 0,2 | 1,2 | 6,1 | 0,0 | 264 | 149 | 0.00010575 | 0.002174812 | 0.0,0.361 | 1.0,1.0 | -0.82 |

| | | | | | | | | | | | | | | | | | | | | | | |
|---|---|---|---|---|---|---|---|---|---|---|---|---|---|---|---|---|---|---|---|---|---|---|
| 11805 | XLOC_008857 | chr12 | + | 44164840 | 44165168 | 44164840 | 44164907 | 44165022 | 115 | 44165168 | 11805 | 4,1 | 2,1 | 8,3 | 0,0 | 264 | 149 | 0.006028777 | 0.047645284 | 0.53,0.361 | 1.0,1.0 | -0.554 |
| 2931 | XLOC_023538 | chr2 | + | 74717151 | 74717600 | 74717151 | 74717254 | 74717370 | 116 | 74717600 | 2931 | 25,20 | 66,51 | 48,31 | 51,49 | 265 | 149 | 0.004707318 | 0.03982423 | 0.176,0.181 | 0.346,0.262 | -0.126 |
| 1934 | XLOC_028602 | chr22 | - | 24180897 | 24181318 | 24180897 | 24180963 | 24181081 | 118 | 24181318 | 1934 | 20,11 | 26,27 | 61,33 | 36,15 | 267 | 149 | 0.000167636 | 0.003179792 | 0.3,0.185 | 0.486,0.551 | -0.276 |
| 7992 | XLOC_018673 | chr17 | - | 62499920 | 62500205 | 62499920 | 62499986 | 62500104 | 118 | 62500205 | 7992 | 19,14 | 1,5 | 15,12 | 1,0 | 267 | 149 | 0.003386189 | 0.030888447 | 0.914,0.61 | 0.893,1.0 | -0.184 |
| 10448 | XLOC_011394 | chr13 | + | 115051703 | 115052104 | 115051703 | 115051875 | 115051993 | 118 | 115052104 | 10448 | 88,99 | 204,203 | 114,118 | 133,137 | 267 | 149 | 8.95E-05 | 0.001902785 | 0.194,0.214 | 0.324,0.325 | -0.12 |
| 15257 | XLOC_047875 | chrX | + | 152959359 | 152959713 | 152959359 | 152959472 | 152959591 | 119 | 152959713 | 15257 | 21,19 | 10,7 | 27,37 | 3,3 | 268 | 149 | 0.001156733 | 0.013678916 | 0.539,0.601 | 0.833,0.873 | -0.283 |
| 14017 | XLOC_000373 | chr1 | + | 93586105 | 93586451 | 93586105 | 93586229 | 93586351 | 122 | 93586451 | 14017 | 23,22 | 5,3 | 5,9 | 1,0 | 271 | 149 | 0.005782123 | 0.046140423 | 0.717,0.801 | 0.733,1.0 | -0.108 |
| 14549 | XLOC_004988 | chr10 | + | 97816649 | 97817004 | 97816649 | 97816757 | 97816880 | 123 | 97817004 | 14549 | 2,0 | 7,8 | 6,4 | 8,6 | 272 | 149 | 0.002580707 | 0.025125565 | 0.135,0.0 | 0.291,0.268 | -0.212 |
| 13620 | XLOC_018545 | chr17 | - | 40139608 | 40139928 | 40139608 | 40139712 | 40139836 | 124 | 40139928 | 13620 | 6,8 | 11,6 | 25,8 | 5,1 | 273 | 149 | 0.000463905 | 0.006861421 | 0.229,0.421 | 0.732,0.814 | -0.448 |
| 15690 | XLOC_000426 | chr1 | + | 113217318 | 113217541 | 113217318 | 113217351 | 113217475 | 124 | 113217541 | 15690 | 11,7 | 8,12 | 11,18 | 0,3 | 273 | 149 | 6.49E-05 | 0.001470498 | 0.429,0.241 | 1.0,0.766 | -0.548 |
| 14675 | XLOC_007153 | chr11 | + | 65338052 | 65338441 | 65338052 | 65338231 | 65338356 | 125 | 65338441 | 14675 | 1,0 | 4,8 | 7,2 | 14,6 | 274 | 149 | 0.001765024 | 0.018950384 | 0.12,0.0 | 0.214,0.153 | -0.123 |
| 9268 | XLOC_037650 | chr6 | + | 13601065 | 13601353 | 13601065 | 13601181 | 13601312 | 131 | 13601353 | 9268 | 7,2 | 3,2 | 15,14 | 1,0 | 280 | 149 | 0.000131825 | 0.002590726 | 0.554,0.347 | 0.889,1.0 | -0.494 |
| 9623 | XLOC_047718 | chrX | + | 53113616 | 53114048 | 53113616 | 53113804 | 53113936 | 132 | 53114048 | 9623 | 26,16 | 43,37 | 13,21 | 6,8 | 281 | 149 | 2.77E-05 | 0.000759497 | 0.243,0.187 | 0.535,0.582 | -0.344 |
| 4365 | XLOC_041366 | chr7 | + | 157046360 | 157046836 | 157046360 | 157046513 | 157046647 | 134 | 157046836 | 4365 | 0,2 | 1,4 | 9,2 | 0,0 | 283 | 149 | 5.08E-09 | 5.46E-07 | 0.0,0.208 | 1.0,1.0 | -0.896 |
| 5672 | XLOC_028587 | chr22 | - | 21983589 | 21984050 | 21983589 | 21983696 | 21983831 | 135 | 21984050 | 5672 | 80,43 | 48,39 | 129,133 | 31,52 | 284 | 149 | 0.000893509 | 0.011303753 | 0.466,0.366 | 0.686,0.573 | -0.213 |
| 12026 | XLOC_000229 | chr1 | + | 40214378 | 40214760 | 40214378 | 40214439 | 40214574 | 135 | 40214760 | 12026 | 1,2 | 6,4 | 6,2 | 1,0 | 284 | 149 | 5.31E-06 | 0.000194341 | 0.08,0.208 | 0.759,1.0 | -0.735 |
| 12027 | XLOC_021106 | chr19 | + | 33073100 | 33073364 | 33073100 | 33073138 | 33073275 | 137 | 33073364 | 12027 | 16,23 | 4,2 | 27,18 | 1,0 | 286 | 149 | 0.001172158 | 0.013817865 | 0.676,0.857 | 0.934,1.0 | -0.201 |
| 7752 | XLOC_034882 | chr5 | - | 40833878 | 40834367 | 40833878 | 40834143 | 40834282 | 139 | 40834367 | 7752 | 8,9 | 9,0 | 30,27 | 0,0 | 288 | 149 | 0.000681711 | 0.009188442 | 0.315,1.0 | 1.0,1.0 | -0.343 |
| 11335 | XLOC_009489 | chr12 | - | 95450972 | 95451419 | 95450972 | 95451188 | 95451328 | 140 | 95451419 | 11335 | 0,5 | 9,15 | 15,11 | 12,21 | 289 | 149 | 0.005621654 | 0.045340978 | 0.0,0.147 | 0.392,0.213 | -0.229 |
| 12943 | XLOC_028747 | chr22 | - | 50720358 | 50720742 | 50720358 | 50720471 | 50720612 | 141 | 50720742 | 12943 | 6,3 | 2,2 | 1,11 | 0,0 | 290 | 149 | 0.000280981 | 0.004672266 | 0.607,0.435 | 1.0,1.0 | -0.479 |
| 7751 | XLOC_034882 | chr5 | - | 40833878 | 40834367 | 40833878 | 40834139 | 40834282 | 143 | 40834367 | 7751 | 8,9 | 30,24 | 30,27 | 39,26 | 292 | 149 | 0.001284459 | 0.014824947 | 0.12,0.161 | 0.282,0.346 | -0.173 |
| 4345 | XLOC_041420 | chr7 | - | 21945580 | 21946074 | 21945580 | 21945762 | 21945906 | 144 | 21946074 | 4345 | 0,4 | 4,3 | 5,8 | 0,1 | 293 | 149 | 8.42E-06 | 0.000283243 | 0.0,0.404 | 1.0,0.803 | -0.7 |
| 10324 | XLOC_021503 | chr19 | - | 6730997 | 6731245 | 6730997 | 6731065 | 6731209 | 144 | 6731245 | 10324 | 24,43 | 5,1 | 62,61 | 1,0 | 293 | 149 | 0.001838997 | 0.019563081 | 0.709,0.956 | 0.969,1.0 | -0.152 |
| 2576 | XLOC_014712 | chr15 | + | 75646993 | 75648036 | 75646993 | 75647159 | 75647304 | 145 | 75648036 | 2576 | 7,5 | 38,40 | 35,32 | 31,45 | 294 | 149 | 1.35E-07 | 8.84E-06 | 0.085,0.06 | 0.364,0.265 | -0.242 |
| 6046 | XLOC_026812 | chr20 | + | 44441758 | 44442103 | 44441758 | 44441929 | 44442075 | 146 | 44442103 | 6046 | 25,22 | 20,20 | 56,31 | 5,6 | 295 | 149 | 4.96E-07 | 2.67E-05 | 0.387,0.357 | 0.85,0.723 | -0.414 |
| 4503 | XLOC_047701 | chrX | + | 48433948 | 48434397 | 48433948 | 48434055 | 48434202 | 147 | 48434397 | 4503 | 92,71 | 28,36 | 257,229 | 34,19 | 296 | 149 | 5.80E-07 | 3.05E-05 | 0.623,0.498 | 0.792,0.858 | -0.264 |
| 12217 | XLOC_007722 | chr11 | - | 116732917 | 116733293 | 116732917 | 116733043 | 116733190 | 147 | 116733293 | 12217 | 4,2 | 4,0 | 4,1 | 0,0 | 296 | 149 | 0.004739864 | 0.039905334 | 0.335,1.0 | 1.0,1.0 | -0.333 |
| 12950 | XLOC_028747 | chr22 | - | 50721432 | 50721874 | 50721432 | 50721612 | 50721765 | 153 | 50721874 | 12950 | 20,24 | 1,5 | 16,44 | 0,0 | 302 | 149 | 0.000144326 | 0.002801237 | 0.908,0.703 | 1.0,1.0 | -0.195 |
| 7728 | XLOC_028532 | chr22 | + | 45717927 | 45718307 | 45717927 | 45718096 | 45718251 | 155 | 45718307 | 7728 | 0,2 | 5,6 | 3,2 | 2,0 | 304 | 149 | 0.002370221 | 0.023705363 | 0.0,0.14 | 0.424,1.0 | -0.642 |
| 14149 | XLOC_029987 | chr3 | - | 121386907 | 121387275 | 121386907 | 121387005 | 121387160 | 155 | 121387275 | 14149 | 0,0 | 10,4 | 3,2 | 1,2 | 304 | 149 | 0.000128343 | 0.002532667 | 0.0,0.0 | 0.595,0.329 | -0.462 |
| 8788 | XLOC_016862 | chr16 | - | 89808927 | 89809346 | 89808927 | 89809051 | 89809207 | 156 | 89809346 | 8788 | 10,9 | 1,4 | 9,11 | 1,0 | 305 | 149 | 0.002221287 | 0.022500071 | 0.83,0.524 | 0.815,1.0 | -0.23 |
| 12347 | XLOC_043957 | chr8 | + | 21937922 | 21938136 | 21937922 | 21937971 | 21938128 | 157 | 21938136 | 12347 | 32,23 | 12,5 | 10,8 | 0,0 | 306 | 149 | 7.80E-06 | 0.00026501 | 0.565,0.691 | 1.0,1.0 | -0.372 |
| 7536 | XLOC_047975 | chrX | - | 57004724 | 57005093 | 57004724 | 57004854 | 57005012 | 158 | 57005093 | 7536 | 0,0 | 3,1 | 9,4 | 2,0 | 307 | 149 | 1.79E-08 | 1.57E-06 | 0.0,0.0 | 0.686,1.0 | -0.843 |
| 15064 | XLOC_018794 | chr17 | - | 80203851 | 80204341 | 80203851 | 80203915 | 80204073 | 158 | 80204341 | 15064 | 105,139 | 83,87 | 194,181 | 37,46 | 307 | 149 | 2.71E-10 | 4.24E-08 | 0.38,0.437 | 0.718,0.656 | -0.279 |
| 8695 | XLOC_047679 | chrX | + | 41064555 | 41064936 | 41064555 | 41064746 | 41064905 | 159 | 41064936 | 8695 | 0,1 | 2,1 | 1,1 | 0,0 | 308 | 149 | 1.07E-06 | 5.13E-05 | 0.0,0.326 | 1.0,1.0 | -0.837 |
| 8792 | XLOC_016862 | chr16 | - | 89851261 | 89851567 | 89851261 | 89851372 | 89851531 | 159 | 89851567 | 8792 | 0,7 | 4,1 | 5,4 | 0,0 | 308 | 149 | 1.15E-05 | 0.000371005 | 0.0,0.772 | 1.0,1.0 | -0.614 |
| 10092 | XLOC_007211 | chr11 | + | 71939770 | 71940080 | 71939770 | 71939891 | 71940051 | 160 | 71940080 | 10092 | 87,107 | 74,67 | 130,132 | 63,42 | 309 | 149 | 0.004293973 | 0.037094418 | 0.362,0.435 | 0.499,0.602 | -0.152 |
| 7929 | XLOC_005345 | chr10 | - | 101945270 | 101945645 | 101945270 | 101945394 | 101945555 | 161 | 101945645 | 7929 | 2,4 | 2,4 | 8,11 | 1,0 | 310 | 149 | 1.65E-05 | 0.000487948 | 0.325,0.325 | 0.794,1.0 | -0.572 |
| 10020 | XLOC_007716 | chr11 | - | 113678672 | 113679159 | 113678672 | 113678858 | 113679019 | 161 | 113679159 | 10020 | 0,1 | 1,3 | 6,3 | 0,0 | 310 | 149 | 4.38E-09 | 4.84E-07 | 0.0,0.138 | 1.0,1.0 | -0.931 |
| 4583 | XLOC_038232 | chr6 | - | 33545311 | 33545685 | 33545311 | 33545412 | 33545574 | 162 | 33545685 | 4583 | 0,0 | 1,6 | 4,2 | 2,0 | 311 | 149 | 5.82E-06 | 0.000210474 | 0.0,0.0 | 0.489,1.0 | -0.744 |
| 12612 | XLOC_007214 | chr11 | + | 73074791 | 73075284 | 73074791 | 73075047 | 73075216 | 169 | 73075284 | 12612 | 13,13 | 3,1 | 10,7 | 0,0 | 318 | 149 | 0.00446648 | 0.038260136 | 0.67,0.859 | 1.0,1.0 | -0.236 |
| 3177 | XLOC_026791 | chr20 | + | 39985679 | 39986087 | 39985679 | 39985828 | 39986000 | 172 | 39986087 | 3177 | 28,38 | 29,19 | 61,50 | 2,7 | 321 | 149 | 2.56E-07 | 1.56E-05 | 0.309,0.481 | 0.934,0.768 | -0.456 |
| 12227 | XLOC_021877 | chr19 | - | 55558472 | 55558856 | 55558472 | 55558579 | 55558754 | 175 | 55558856 | 12227 | 25,31 | 4,4 | 53,42 | 0,0 | 324 | 149 | 1.50E-05 | 0.000450719 | 0.742,0.781 | 1.0,1.0 | -0.238 |
| 3898 | XLOC_046087 | chr9 | + | 131831006 | 131831408 | 131831006 | 131831178 | 131831354 | 176 | 131831408 | 3898 | 4,8 | 4,0 | 2,3 | 0,0 | 325 | 149 | 0.006130751 | 0.048130878 | 0.314,1.0 | 1.0,1.0 | -0.343 |
| 10093 | XLOC_007211 | chr11 | + | 71939770 | 71940080 | 71939770 | 71939891 | 71940067 | 176 | 71940080 | 10093 | 61,76 | 46,72 | 116,112 | 32,37 | 325 | 149 | 4.90E-07 | 2.67E-05 | 0.378,0.326 | 0.624,0.581 | -0.251 |
| 3549 | XLOC_048083 | chrX | - | 130928021 | 130928494 | 130928021 | 130928143 | 130928324 | 181 | 130928494 | 3549 | 1,0 | 8,3 | 23,19 | 16,15 | 330 | 149 | 2.89E-06 | 0.000116877 | 0.053,0.0 | 0.394,0.364 | -0.352 |
| 6598 | XLOC_023898 | chr2 | + | 234356503 | 234356827 | 234356503 | 234356551 | 234356732 | 181 | 234356827 | 6598 | 5,14 | 67,68 | 18,40 | 36,17 | 330 | 149 | 1.41E-05 | 0.000429121 | 0.033,0.085 | 0.184,0.515 | -0.291 |
| 15296 | XLOC_001334 | chr1 | - | 153913379 | 153913870 | 153913379 | 153913524 | 153913706 | 182 | 153913870 | 15296 | 32,33 | 66,60 | 67,73 | 65,46 | 331 | 149 | 0.000114746 | 0.002329292 | 0.179,0.198 | 0.317,0.417 | -0.178 |
| 4935 | XLOC_034481 | chr5 | + | 647919 | 648338 | 647919 | 648031 | 648214 | 183 | 648338 | 4935 | 12,10 | 18,17 | 13,10 | 2,5 | 332 | 149 | 0.00263591 | 0.025596748 | 0.23,0.209 | 0.745,0.473 | -0.389 |
| 12050 | XLOC_038395 | chr6 | - | 109766973 | 109767615 | 109766973 | 109767155 | 109767338 | 183 | 109767615 | 12050 | 0,1 | 3,1 | 2,1 | 0,0 | 332 | 149 | 4.59E-09 | 5.04E-07 | 0.0,0.31 | 1.0,1.0 | -0.845 |
| 6115 | XLOC_026682 | chr20 | + | 2636783 | 2637195 | 2636783 | 2636860 | 2637046 | 186 | 2637195 | 6115 | 159,136 | 157,144 | 313,362 | 196,202 | 335 | 149 | 0.000128469 | 0.002532667 | 0.311,0.296 | 0.415,0.444 | -0.126 |
| 7500 | XLOC_016577 | chr16 | - | 2162788 | 2163285 | 2162788 | 2162964 | 2163157 | 193 | 2163285 | 7500 | 8,2 | 21,13 | 18,12 | 13,18 | 342 | 149 | 0.005425628 | 0.044253352 | 0.142,0.063 | 0.376,0.225 | -0.198 |
| 10492 | XLOC_044467 | chr8 | - | 145753407 | 145753819 | 145753407 | 145753456 | 145753651 | 195 | 145753819 | 10492 | 38,38 | 4,3 | 68,68 | 0,1 | 344 | 149 | 0.000391229 | 0.006029578 | 0.804,0.846 | 1.0,0.967 | -0.159 |
| 9808 | XLOC_018520 | chr17 | - | 37009274 | 37009967 | 37009274 | 37009358 | 37009554 | 196 | 37009967 | 9808 | 23,24 | 4,2 | 62,45 | 0,1 | 345 | 149 | 0.000498082 | 0.007224832 | 0.713,0.838 | 1.0,0.951 | -0.2 |
| 7501 | XLOC_016577 | chr16 | - | 2162788 | 2163285 | 2162788 | 2162964 | 2163161 | 197 | 2163285 | 7501 | 8,3 | 53,45 | 22,16 | 58,38 | 346 | 149 | 0.000215923 | 0.003848235 | 0.061,0.028 | 0.14,0.153 | -0.102 |
| 11093 | XLOC_013429 | chr14 | - | 68059031 | 68059435 | 68059031 | 68059153 | 68059351 | 198 | 68059435 | 11093 | 0,0 | 2,4 | 11,2 | 6,5 | 347 | 149 | 4.07E-05 | 0.001029275 | 0.0,0.0 | 0.44,0.147 | -0.293 |
| 2130 | XLOC_007597 | chr11 | - | 66133596 | 66134035 | 66133596 | 66133702 | 66133901 | 199 | 66134035 | 2130 | 78,77 | 106,130 | 122,106 | 102,75 | 348 | 149 | 0.000104229 | 0.002150628 | 0.24,0.202 | 0.339,0.377 | -0.137 |
| 445 | XLOC_034514 | chr5 | + | 35709049 | 35710594 | 35709049 | 35709223 | 35709426 | 203 | 35710594 | 445 | 17,23 | 2,4 | 12,18 | 0,0 | 352 | 149 | 0.000178437 | 0.003309556 | 0.783,0.709 | 1.0,1.0 | -0.254 |
| 2199 | XLOC_009410 | chr12 | - | 56720096 | 56720743 | 56720096 | 56720193 | 56720400 | 207 | 56720743 | 2199 | 27,38 | 92,80 | 60,63 | 71,84 | 356 | 149 | 0.000380221 | 0.005888823 | 0.109,0.166 | 0.261,0.239 | -0.112 |
| 15749 | XLOC_007017 | chr11 | + | 18418026 | 18418515 | 18418026 | 18418157 | 18418365 | 208 | 18418515 | 15749 | 14,17 | 13,11 | 16,8 | 3,1 | 357 | 149 | 0.003150057 | 0.029295865 | 0.31,0.392 | 0.69,0.77 | -0.379 |
| 457 | XLOC_041145 | chr7 | + | 75044869 | 75045469 | 75044869 | 75045048 | 75045258 | 210 | 75045469 | 457 | 211,241 | 53,34 | 340,367 | 19,12 | 359 | 149 | 6.88E-09 | 7.04E-07 | 0.623,0.746 | 0.881,0.927 | -0.22 |
| 12541 | XLOC_000835 | chr1 | - | 1647784 | 1648250 | 1647784 | 1647917 | 1648127 | 210 | 1648250 | 12541 | 8,4 | 3,1 | 7,11 | 1,0 | 359 | 149 | 0.002159978 | 0.022033115 | 0.525,0.624 | 0.744,1.0 | -0.297 |
| 3788 | XLOC_046425 | chr9 | - | 130690343 | 130690762 | 130690343 | 130690498 | 130690711 | 213 | 130690762 | 3788 | 0,0 | 2,1 | 4,1 | 6,5 | 362 | 149 | 0.006244736 | 0.048709679 | 0.0,0.0 | 0.215,0.076 | -0.145 |
| 3376 | XLOC_009413 | chr12 | - | 56814746 | 56815277 | 56814746 | 56814918 | 56815134 | 216 | 56815277 | 3376 | 108,103 | 399,382 | 328,324 | 394,337 | 365 | 149 | 0 | 0 | 0.1,0.099 | 0.254,0.282 | -0.169 |
| 6795 | XLOC_004772 | chr10 | + | 13167417 | 13167836 | 13167417 | 13167567 | 13167784 | 217 | 13167836 | 6795 | 7,6 | 1,3 | 6,10 | 0,1 | 366 | 149 | 0.003395139 | 0.030926268 | 0.74,0.449 | 1.0,0.803 | -0.307 |

| | | | | | | | | | | | | | | | | | | | | | | |
|---|---|---|---|---|---|---|---|---|---|---|---|---|---|---|---|---|---|---|---|---|---|---|
| 1177 | XLOC_020248 | chr18 | - | 60985314 | 60987270 | 60985314 | 60986185 | 60986406 | 221 | 60987270 | 1177 | 18,11 | 31,23 | 40,55 | 10,25 | 370 | 149 | 9.38E-07 | 4.54E-05 | 0.19,0.161 | 0.617,0.47 | -0.368 |
| 15070 | XLOC_020938 | chr19 | + | 7599778 | 7600088 | 7599778 | 7599788 | 7600012 | 224 | 7600088 | 15070 | 0,2 | 3,0 | 6,4 | 1,0 | 373 | 149 | 0.003349595 | 0.030666227 | 0.0,1.0 | 0.706,1.0 | -0.353 |
| 23 | XLOC_047880 | chrX | + | 153641543 | 153641904 | 153641543 | 153641589 | 153641818 | 229 | 153641904 | 23 | 34,38 | 98,73 | 115,89 | 141,97 | 378 | 149 | 0.000202668 | 0.0036597 | 0.12,0.17 | 0.243,0.266 | -0.109 |
| 2899 | XLOC_014590 | chr15 | + | 42128672 | 42129138 | 42128672 | 42128749 | 42128978 | 229 | 42129138 | 2899 | 10,29 | 16,20 | 38,53 | 16,14 | 378 | 149 | 0.004075813 | 0.035561706 | 0.198,0.364 | 0.484,0.599 | -0.26 |
| 7873 | XLOC_018791 | chr17 | - | 80047125 | 80047602 | 80047125 | 80047277 | 80047507 | 230 | 80047602 | 7873 | 14,8 | 0,7 | 14,15 | 0,0 | 379 | 149 | 0.001064078 | 0.012897549 | 1.0,0.31 | 1.0,1.0 | -0.345 |
| 3986 | XLOC_045902 | chr9 | + | 35800018 | 35800480 | 35800018 | 35800154 | 35800385 | 231 | 35800480 | 3986 | 0,1 | 6,4 | 5,3 | 5,15 | 380 | 149 | 0.005662861 | 0.045599975 | 0.0,0.089 | 0.282,0.073 | -0.133 |
| 262 | XLOC_038373 | chr6 | - | 99845590 | 99849506 | 99845590 | 99848187 | 99848419 | 232 | 99849506 | 262 | 364,303 | 141,143 | 337,330 | 52,92 | 381 | 149 | 0.000440597 | 0.006594491 | 0.502,0.453 | 0.717,0.584 | -0.173 |
| 2749 | XLOC_000004 | chr1 | + | 897008 | 898297 | 897008 | 897851 | 898083 | 232 | 898297 | 2749 | 18,11 | 20,27 | 50,69 | 29,27 | 381 | 149 | 0.00011585 | 0.002345372 | 0.26,0.137 | 0.403,0.5 | -0.253 |
| 5465 | XLOC_029402 | chr3 | + | 45043029 | 45043506 | 45043029 | 45043100 | 45043333 | 233 | 45043506 | 5465 | 8,6 | 4,0 | 4,9 | 0,0 | 382 | 149 | 0.004632813 | 0.039371061 | 0.438,1.0 | 1.0,1.0 | -0.281 |
| 10508 | XLOC_027816 | chr21 | - | 46274786 | 46275171 | 46274786 | 46274891 | 46275124 | 233 | 46275171 | 10508 | 13,16 | 5,1 | 7,8 | 0,0 | 382 | 149 | 6.96E-05 | 0.001561627 | 0.504,0.862 | 1.0,1.0 | -0.317 |
| 2890 | XLOC_016217 | chr16 | + | 2283011 | 2283991 | 2283011 | 2283648 | 2283882 | 234 | 2283991 | 2890 | 14,32 | 72,69 | 39,73 | 72,67 | 383 | 149 | 0.000667884 | 0.009060008 | 0.07,0.153 | 0.174,0.298 | -0.124 |
| 526 | XLOC_044238 | chr8 | - | 21963963 | 21965857 | 21963963 | 21965406 | 21965643 | 237 | 21965857 | 526 | 4,8 | 9,20 | 25,26 | 24,15 | 386 | 149 | 0.002694867 | 0.026020549 | 0.146,0.134 | 0.287,0.401 | -0.204 |
| 6489 | XLOC_014964 | chr15 | - | 68503600 | 68504079 | 68503600 | 68503656 | 68503893 | 237 | 68504079 | 6489 | 31,23 | 15,12 | 58,63 | 5,12 | 386 | 149 | 0.000637402 | 0.008708263 | 0.444,0.425 | 0.817,0.67 | -0.309 |
| 6690 | XLOC_023576 | chr2 | + | 96688421 | 96689121 | 96688421 | 96688501 | 96688739 | 238 | 96689121 | 6690 | 1,6 | 0,4 | 3,4 | 0,0 | 387 | 149 | 0.003045875 | 0.028492571 | 1.0,0.366 | 1.0,1.0 | -0.317 |
| 430 | XLOC_034597 | chr5 | + | 78908274 | 78908898 | 78908274 | 78908592 | 78908833 | 241 | 78908898 | 430 | 157,210 | 21,15 | 146,131 | 3,7 | 390 | 149 | 0.003155119 | 0.029295865 | 0.741,0.842 | 0.949,0.877 | -0.122 |
| 10847 | XLOC_011260 | chr13 | + | 20244979 | 20245429 | 20244979 | 20245104 | 20245345 | 241 | 20245429 | 10847 | 16,19 | 4,2 | 23,30 | 0,1 | 390 | 149 | 0.000468368 | 0.006902234 | 0.604,0.784 | 1.0,0.92 | -0.266 |
| 11603 | XLOC_009165 | chr12 | + | 123465366 | 123465816 | 123465366 | 123465434 | 123465675 | 241 | 123465816 | 11603 | 118,124 | 26,24 | 280,252 | 21,28 | 390 | 149 | 0.000698501 | 0.009347729 | 0.634,0.664 | 0.836,0.775 | -0.156 |
| 13562 | XLOC_029295 | chr3 | + | 5236470 | 5236966 | 5236470 | 5236650 | 5236893 | 243 | 5236966 | 13562 | 1,2 | 3,5 | 11,5 | 3,1 | 392 | 149 | 0.000495076 | 0.007188159 | 0.112,0.132 | 0.582,0.655 | -0.497 |
| 7897 | XLOC_013249 | chr14 | + | 96991646 | 96992096 | 96991646 | 96991728 | 96991974 | 246 | 96992096 | 7897 | 0,1 | 2,1 | 3,3 | 0,0 | 395 | 149 | 3.77E-07 | 2.16E-05 | 0.0,0.274 | 1.0,1.0 | -0.863 |
| 4940 | XLOC_038429 | chr6 | - | 131201283 | 131201629 | 131201283 | 131201346 | 131201593 | 247 | 131201629 | 4940 | 16,18 | 4,1 | 14,6 | 0,0 | 396 | 149 | 0.0003089 | 0.005045025 | 0.601,0.871 | 1.0,1.0 | -0.264 |
| 11223 | XLOC_018789 | chr17 | - | 80008537 | 80008892 | 80008537 | 80008640 | 80008887 | 247 | 80008892 | 11223 | 30,12 | 2,3 | 31,34 | 0,0 | 396 | 149 | 0.00047319 | 0.006963821 | 0.849,0.601 | 1.0,1.0 | -0.275 |
| 1582 | XLOC_045894 | chr9 | + | 34660500 | 34660933 | 34660500 | 34660597 | 34660850 | 253 | 34660933 | 1582 | 13,11 | 7,1 | 14,12 | 1,0 | 402 | 149 | 0.000664811 | 0.009041666 | 0.408,0.803 | 0.838,1.0 | -0.314 |
| 2888 | XLOC_016217 | chr16 | + | 2283011 | 2283648 | 2283011 | 2283184 | 2283438 | 254 | 2283648 | 2888 | 20,34 | 87,99 | 58,78 | 107,88 | 403 | 149 | 4.31E-05 | 0.001077557 | 0.078,0.113 | 0.167,0.247 | -0.112 |
| 837 | XLOC_004889 | chr10 | + | 75560397 | 75560953 | 75560397 | 75560517 | 75560774 | 257 | 75560953 | 837 | 17,36 | 81,44 | 55,68 | 52,53 | 406 | 149 | 0.000319489 | 0.005161928 | 0.072,0.231 | 0.28,0.32 | -0.149 |
| 5726 | XLOC_028548 | chr22 | + | 50354965 | 50355420 | 50354965 | 50355016 | 50355275 | 259 | 50355420 | 5726 | 45,20 | 7,0 | 33,45 | 0,0 | 408 | 149 | 0.000252934 | 0.004363107 | 0.701,1.0 | 1.0,1.0 | -0.149 |
| 491 | XLOC_032542 | chr4 | - | 2243246 | 2243898 | 2243246 | 2243517 | 2243777 | 260 | 2243898 | 491 | 74,72 | 16,20 | 141,110 | 6,6 | 409 | 149 | 6.09E-07 | 3.17E-05 | 0.628,0.567 | 0.895,0.87 | -0.285 |
| 12013 | XLOC_021828 | chr19 | - | 50364705 | 50365128 | 50364705 | 50364767 | 50365028 | 261 | 50365128 | 12013 | 82,79 | 9,3 | 55,44 | 1,0 | 410 | 149 | 0.000359688 | 0.005675864 | 0.768,0.905 | 0.952,1.0 | -0.139 |
| 1427 | XLOC_018607 | chr17 | - | 45915017 | 45915816 | 45915017 | 45915368 | 45915635 | 267 | 45915816 | 1427 | 54,89 | 133,152 | 75,67 | 56,54 | 416 | 149 | 5.96E-07 | 3.11E-05 | 0.127,0.173 | 0.324,0.308 | -0.166 |
| 6327 | XLOC_016250 | chr16 | + | 3335719 | 3336149 | 3335719 | 3335754 | 3336022 | 268 | 3336149 | 6327 | 23,28 | 52,63 | 26,51 | 33,31 | 417 | 149 | 0.001083394 | 0.013026704 | 0.136,0.137 | 0.22,0.37 | -0.158 |
| 13374 | XLOC_030032 | chr3 | - | 129115882 | 129116378 | 129115882 | 129115983 | 129116253 | 270 | 129116378 | 13374 | 24,17 | 8,6 | 25,31 | 0,2 | 419 | 149 | 3.39E-05 | 0.000895878 | 0.516,0.502 | 1.0,0.846 | -0.414 |
| 2527 | XLOC_007071 | chr11 | + | 47182677 | 47184358 | 47182677 | 47182741 | 47183014 | 273 | 47184358 | 2527 | 36,64 | 72,86 | 67,77 | 40,59 | 422 | 149 | 4.27E-05 | 0.001071261 | 0.15,0.208 | 0.372,0.315 | -0.165 |
| 1320 | XLOC_018401 | chr17 | - | 7982758 | 7983259 | 7982758 | 7982842 | 7983122 | 280 | 7983259 | 1320 | 5,13 | 0,9 | 7,10 | 1,0 | 429 | 149 | 0.006319276 | 0.049118899 | 1.0,0.334 | 0.709,1.0 | -0.188 |
| 15746 | XLOC_047699 | chrX | + | 48380267 | 48380640 | 48380267 | 48380295 | 48380580 | 285 | 48380640 | 15746 | 10,10 | 7,4 | 25,27 | 2,4 | 434 | 149 | 0.003996114 | 0.03495032 | 0.329,0.462 | 0.811,0.699 | -0.359 |
| 3159 | XLOC_018763 | chr17 | - | 78187602 | 78188127 | 78187602 | 78187684 | 78187970 | 286 | 78188127 | 3159 | 22,50 | 125,118 | 87,112 | 71,84 | 435 | 149 | 4.52E-14 | 1.79E-11 | 0.057,0.127 | 0.296,0.314 | -0.213 |
| 15882 | XLOC_000712 | chr1 | + | 212529967 | 212530386 | 212529967 | 212530018 | 212530304 | 286 | 212530386 | 15882 | 8,8 | 2,5 | 14,13 | 0,2 | 435 | 149 | 0.005199306 | 0.042759957 | 0.578,0.354 | 1.0,0.69 | -0.379 |
| 11117 | XLOC_009338 | chr12 | - | 49233216 | 49233692 | 49233216 | 49233334 | 49233626 | 292 | 49233692 | 11117 | 0,0 | 3,1 | 1,6 | 10,3 | 441 | 149 | 0.001654911 | 0.018077827 | 0.0,0.0 | 0.033,0.403 | -0.218 |
| 13915 | XLOC_046078 | chr9 | + | 131389278 | 131389777 | 131389278 | 131389369 | 131389664 | 295 | 131389777 | 13915 | 0,1 | 2,0 | 3,2 | 0,0 | 444 | 149 | 0.000363764 | 0.005717595 | 0.0,1.0 | 1.0,1.0 | -0.5 |
| 4499 | XLOC_044293 | chr8 | - | 48846193 | 48846650 | 48846193 | 48846224 | 48846524 | 300 | 48846650 | 4499 | 4,11 | 2,8 | 11,9 | 0,0 | 449 | 149 | 2.78E-08 | 2.38E-06 | 0.399,0.313 | 1.0,1.0 | -0.644 |
| 3030 | XLOC_016775 | chr16 | - | 67863667 | 67864451 | 67863667 | 67863991 | 67864292 | 301 | 67864451 | 3030 | 251,197 | 42,44 | 267,247 | 24,28 | 450 | 149 | 0.002055827 | 0.021224263 | 0.664,0.597 | 0.786,0.745 | -0.135 |
| 10175 | XLOC_013336 | chr14 | - | 24910879 | 24911378 | 24910879 | 24911001 | 24911303 | 302 | 24911378 | 10175 | 30,47 | 14,19 | 41,36 | 3,1 | 451 | 149 | 1.79E-06 | 7.89E-05 | 0.415,0.45 | 0.819,0.922 | -0.438 |
| 2198 | XLOC_009410 | chr12 | - | 56716398 | 56716993 | 56716398 | 56716542 | 56716845 | 303 | 56716993 | 2198 | 238,186 | 78,59 | 313,281 | 68,49 | 452 | 149 | 0.00383442 | 0.033956493 | 0.501,0.51 | 0.603,0.654 | -0.123 |
| 132 | XLOC_044027 | chr8 | + | 48641494 | 48642027 | 48641494 | 48641664 | 48641973 | 309 | 48642027 | 132 | 25,28 | 5,8 | 13,21 | 0,0 | 458 | 149 | 1.10E-06 | 5.22E-05 | 0.619,0.532 | 1.0,1.0 | -0.424 |
| 13566 | XLOC_021518 | chr19 | - | 8654335 | 8654875 | 8654335 | 8654458 | 8654772 | 314 | 8654875 | 13566 | 8,12 | 2,3 | 10,10 | 1,0 | 463 | 149 | 0.001306755 | 0.014993295 | 0.563,0.563 | 0.763,1.0 | -0.319 |
| 14923 | XLOC_018785 | chr17 | - | 79894623 | 79895123 | 79894623 | 79894737 | 79895055 | 318 | 79895123 | 14923 | 100,100 | 0,13 | 135,156 | 1,0 | 467 | 149 | 0.000119626 | 0.002405628 | 1.0,0.711 | 0.977,1.0 | -0.133 |
| 11684 | XLOC_028526 | chr22 | + | 44330485 | 44330964 | 44330485 | 44330546 | 44330865 | 319 | 44330964 | 11684 | 15,26 | 3,1 | 19,34 | 0,0 | 468 | 149 | 0.005053433 | 0.041834749 | 0.614,0.892 | 1.0,1.0 | -0.247 |
| 1952 | XLOC_044467 | chr8 | - | 145750199 | 145753171 | 145750199 | 145752023 | 145752343 | 320 | 145753171 | 1952 | 206,234 | 43,51 | 429,466 | 35,38 | 469 | 149 | 5.80E-07 | 3.05E-05 | 0.603,0.593 | 0.796,0.796 | -0.198 |
| 5239 | XLOC_029869 | chr3 | - | 52029057 | 52029479 | 52029057 | 52029122 | 52029442 | 320 | 52029479 | 5239 | 9,4 | 5,1 | 6,8 | 0,1 | 469 | 149 | 0.00069128 | 0.009267594 | 0.364,0.56 | 1.0,0.718 | -0.397 |
| 2231 | XLOC_009247 | chr12 | - | 9573326 | 9574486 | 9573326 | 9574087 | 9574413 | 326 | 9574486 | 2231 | 190,118 | 61,38 | 303,318 | 67,54 | 475 | 149 | 0.006040391 | 0.047645284 | 0.494,0.493 | 0.587,0.649 | -0.124 |
| 3969 | XLOC_045897 | chr9 | + | 35606833 | 35607406 | 35606833 | 35606980 | 35607308 | 328 | 35607406 | 3969 | 10,18 | 4,0 | 3,16 | 0,0 | 477 | 149 | 0.005836483 | 0.04645114 | 0.438,1.0 | 1.0,1.0 | -0.281 |
| 2838 | XLOC_009549 | chr12 | - | 112277460 | 112278281 | 112277460 | 112277807 | 112278136 | 329 | 112278281 | 2838 | 246,208 | 43,35 | 433,401 | 22,18 | 478 | 149 | 5.40E-09 | 5.72E-07 | 0.641,0.649 | 0.86,0.874 | -0.222 |
| 3405 | XLOC_021451 | chr19 | - | 1609289 | 1611848 | 1609289 | 1611368 | 1611702 | 334 | 1611848 | 3405 | 486,609 | 101,98 | 900,838 | 96,47 | 483 | 149 | 3.45E-06 | 0.000133758 | 0.597,0.657 | 0.743,0.846 | -0.167 |
| 3374 | XLOC_017907 | chr17 | + | 7358602 | 7359260 | 7358602 | 7358775 | 7359112 | 337 | 7359260 | 3374 | 17,24 | 6,6 | 16,37 | 0,1 | 486 | 149 | 3.10E-06 | 0.000123622 | 0.465,0.551 | 1.0,0.919 | -0.452 |
| 2894 | XLOC_016217 | chr16 | + | 2287767 | 2289507 | 2287767 | 2288019 | 2288362 | 343 | 2289507 | 2894 | 3,4 | 16,11 | 9,17 | 5,8 | 492 | 149 | 2.87E-05 | 0.000782193 | 0.054,0.099 | 0.353,0.392 | -0.296 |
| 3333 | XLOC_000851 | chr1 | - | 3547163 | 3548221 | 3547163 | 3547681 | 3548029 | 348 | 3548221 | 3333 | 37,21 | 29,24 | 76,48 | 24,23 | 497 | 149 | 0.001973841 | 0.020729023 | 0.277,0.208 | 0.487,0.385 | -0.194 |
| 8515 | XLOC_017884 | chr17 | + | 5284683 | 5285164 | 5284683 | 5284800 | 5285148 | 348 | 5285164 | 8515 | 24,21 | 3,2 | 29,29 | 1,0 | 497 | 149 | 0.002011847 | 0.020946998 | 0.706,0.759 | 0.897,1.0 | -0.216 |
| 4092 | XLOC_046494 | chr9 | - | 140006141 | 140006621 | 140006141 | 140006206 | 140006563 | 357 | 140006621 | 4092 | 70,71 | 17,35 | 69,83 | 6,1 | 506 | 149 | 3.13E-06 | 0.000123873 | 0.548,0.374 | 0.772,0.961 | -0.406 |
| 2582 | XLOC_006946 | chr11 | + | 581491 | 582081 | 581491 | 581597 | 581964 | 367 | 582081 | 2582 | 18,5 | 6,0 | 27,18 | 0,1 | 516 | 149 | 0.003800596 | 0.03378757 | 0.464,1.0 | 1.0,0.839 | -0.188 |
| 501 | XLOC_046480 | chr9 | - | 139326275 | 139327527 | 139326275 | 139327038 | 139327407 | 369 | 139327527 | 501 | 24,13 | 26,13 | 38,17 | 16,4 | 518 | 149 | 0.004153821 | 0.036033318 | 0.21,0.223 | 0.406,0.55 | -0.262 |
| 2589 | XLOC_000006 | chr1 | + | 982199 | 982834 | 982199 | 982337 | 982706 | 369 | 982834 | 2589 | 85,88 | 302,291 | 254,297 | 254,282 | 518 | 149 | 0 | 0 | 0.075,0.08 | 0.223,0.233 | -0.151 |
| 11793 | XLOC_000548 | chr1 | + | 155295205 | 155295701 | 155295205 | 155295281 | 155295654 | 373 | 155295701 | 11793 | 23,17 | 5,1 | 26,7 | 0,1 | 522 | 149 | 0.003473505 | 0.031456029 | 0.568,0.829 | 1.0,0.666 | -0.135 |
| 996 | XLOC_026793 | chr20 | + | 42158923 | 42159674 | 42158923 | 42159059 | 42159437 | 378 | 42159674 | 996 | 20,13 | 2,2 | 15,14 | 0,0 | 527 | 149 | 0.001559373 | 0.01720916 | 0.739,0.648 | 1.0,1.0 | -0.306 |
| 2671 | XLOC_001160 | chr1 | - | 78412214 | 78414985 | 78412214 | 78414459 | 78414839 | 380 | 78414985 | 2671 | 672,575 | 833,725 | 928,849 | 529,549 | 529 | 149 | 2.11E-10 | 3.41E-08 | 0.185,0.183 | 0.331,0.303 | -0.133 |
| 1482 | XLOC_016684 | chr16 | - | 28890054 | 28891076 | 28890054 | 28890425 | 28890809 | 384 | 28891076 | 1482 | 26,13 | 3,5 | 10,20 | 0,1 | 533 | 149 | 0.000550044 | 0.007786903 | 0.708,0.421 | 1.0,0.848 | -0.359 |
| 128 | XLOC_044112 | chr8 | + | 101225317 | 101226146 | 101225317 | 101225656 | 101226046 | 390 | 101226146 | 128 | 0,0 | 13,13 | 6,2 | 19,4 | 539 | 149 | 0.00149377 | 0.016607017 | 0.0,0.0 | 0.08,0.121 | -0.101 |
| 15728 | XLOC_009117 | chr12 | + | 113725967 | 113726652 | 113725967 | 113726039 | 113726430 | 391 | 113726652 | 15728 | 56,50 | 5,0 | 66,53 | 0,0 | 540 | 149 | 0.005024864 | 0.041713027 | 0.756,1.0 | 1.0,1.0 | -0.122 |

| | | | | | | | | | | | | | | | | | | | | | | |
|---|---|---|---|---|---|---|---|---|---|---|---|---|---|---|---|---|---|---|---|---|---|---|
| 2740 | XLOC_005357 | chr10 | - | 103344358 | 103345235 | 103344358 | 103344676 | 103345072 | 396 | 103345235 | 2740 | 39,33 | 60,54 | 56,62 | 45,34 | 545 | 149 | 0.000223377 | 0.003939084 | 0.151,0.143 | 0.254,0.333 | -0.146 |
| 6963 | XLOC_016235 | chr16 | + | 2818505 | 2819285 | 2818505 | 2818600 | 2818997 | 397 | 2819285 | 6963 | 355,308 | 59,82 | 650,487 | 38,32 | 546 | 149 | 3.44E-09 | 3.91E-07 | 0.621,0.506 | 0.824,0.806 | -0.251 |
| 228 | XLOC_041046 | chr7 | + | 27138692 | 27140058 | 27138692 | 27138985 | 27139397 | 412 | 27140058 | 228 | 22,21 | 33,26 | 42,50 | 17,18 | 561 | 149 | 8.89E-06 | 0.000295782 | 0.15,0.177 | 0.396,0.425 | -0.247 |
| 125 | XLOC_044167 | chr8 | + | 144462234 | 144462953 | 144462234 | 144462345 | 144462758 | 413 | 144462953 | 125 | 234,215 | 38,25 | 215,252 | 12,8 | 562 | 149 | 8.58E-06 | 0.000288207 | 0.62,0.695 | 0.826,0.893 | -0.202 |
| 7557 | XLOC_024366 | chr2 | - | 201778017 | 201778715 | 201778017 | 201778136 | 201778550 | 414 | 201778715 | 7557 | 14,38 | 6,0 | 78,87 | 0,0 | 563 | 149 | 0.001080935 | 0.013007544 | 0.382,1.0 | 1.0,1.0 | -0.309 |
| 1119 | XLOC_044280 | chr8 | - | 38125888 | 38126508 | 38125888 | 38125979 | 38126399 | 420 | 38126508 | 1119 | 37,61 | 64,54 | 64,57 | 46,26 | 569 | 149 | 0.00404822 | 0.035341452 | 0.131,0.228 | 0.267,0.365 | -0.137 |
| 2602 | XLOC_007095 | chr11 | + | 60702723 | 60704627 | 60702723 | 60702915 | 60703335 | 420 | 60704627 | 2602 | 74,89 | 150,133 | 98,76 | 83,61 | 569 | 149 | 3.69E-05 | 0.000954575 | 0.114,0.149 | 0.236,0.246 | -0.109 |
| 15850 | XLOC_000295 | chr1 | + | 52221991 | 52222487 | 52221991 | 52222030 | 52222452 | 422 | 52222487 | 15850 | 8,12 | 7,2 | 10,21 | 1,0 | 571 | 149 | 9.83E-05 | 0.002042848 | 0.23,0.61 | 0.723,1.0 | -0.441 |
| 12307 | XLOC_000147 | chr1 | + | 27087346 | 27087964 | 27087346 | 27087446 | 27087874 | 428 | 27087964 | 12307 | 15,12 | 4,1 | 7,18 | 0,0 | 577 | 149 | 2.93E-05 | 0.000794017 | 0.492,0.756 | 1.0,1.0 | -0.376 |
| 466 | XLOC_035162 | chr5 | - | 176958282 | 176959201 | 176958282 | 176958522 | 176958953 | 431 | 176959201 | 466 | 25,46 | 24,37 | 60,45 | 27,15 | 580 | 149 | 0.004170182 | 0.036133566 | 0.211,0.242 | 0.363,0.435 | -0.173 |
| 2261 | XLOC_014987 | chr15 | - | 75197004 | 75197572 | 75197004 | 75197053 | 75197494 | 441 | 75197572 | 2261 | 48,56 | 82,80 | 88,67 | 57,50 | 590 | 149 | 5.62E-05 | 0.001317054 | 0.129,0.15 | 0.281,0.253 | -0.128 |
| 8423 | XLOC_017964 | chr17 | + | 19283094 | 19283814 | 19283094 | 19283260 | 19283707 | 447 | 19283814 | 8423 | 14,13 | 0,4 | 8,11 | 0,0 | 596 | 149 | 0.004361866 | 0.037556488 | 1.0,0.448 | 1.0,1.0 | -0.276 |
| 10120 | XLOC_013470 | chr14 | - | 77270128 | 77270758 | 77270128 | 77270255 | 77270717 | 462 | 77270758 | 10120 | 5,6 | 2,2 | 8,19 | 1,0 | 611 | 149 | 0.000317932 | 0.005155618 | 0.379,0.422 | 0.661,1.0 | -0.43 |
| 940 | XLOC_023708 | chr2 | + | 170550968 | 170551757 | 170550968 | 170551251 | 170551724 | 473 | 170551757 | 940 | 3,5 | 4,7 | 19,7 | 4,2 | 622 | 149 | 0.001873827 | 0.019849366 | 0.152,0.146 | 0.532,0.456 | -0.345 |
| 492 | XLOC_032538 | chr4 | - | 1719242 | 1720353 | 1719242 | 1719402 | 1719878 | 476 | 1720353 | 492 | 82,64 | 64,46 | 89,67 | 26,15 | 625 | 149 | 6.06E-06 | 0.00021695 | 0.234,0.249 | 0.449,0.516 | -0.241 |
| 103 | XLOC_044433 | chr8 | - | 144363051 | 144363914 | 144363051 | 144363167 | 144363651 | 484 | 144363914 | 103 | 92,90 | 59,70 | 151,157 | 28,28 | 633 | 149 | 2.56E-11 | 5.58E-09 | 0.268,0.232 | 0.559,0.569 | -0.314 |
| 13736 | XLOC_014788 | chr15 | + | 99500289 | 99507746 | 99500289 | 99500705 | 99501191 | 486 | 99507746 | 13736 | 81,89 | 10,15 | 119,91 | 5,4 | 635 | 149 | 0.000975772 | 0.012050538 | 0.655,0.582 | 0.848,0.842 | -0.226 |
| 6591 | XLOC_023924 | chr2 | - | 38828 | 42305 | 38828 | 41627 | 42117 | 490 | 42305 | 6591 | 5,4 | 3,2 | 1,9 | 0,0 | 639 | 149 | 9.32E-07 | 4.52E-05 | 0.28,0.318 | 1.0,1.0 | -0.701 |
| 7636 | XLOC_020063 | chr18 | + | 21406665 | 21407431 | 21406665 | 21406781 | 21407277 | 496 | 21407431 | 7636 | 17,18 | 3,6 | 25,34 | 0,0 | 645 | 149 | 5.33E-08 | 4.09E-06 | 0.567,0.409 | 1.0,1.0 | -0.512 |
| 3186 | XLOC_007240 | chr11 | + | 82783664 | 82808540 | 82783664 | 82783846 | 82784344 | 498 | 82808540 | 3186 | 70,72 | 95,101 | 101,68 | 53,65 | 647 | 149 | 0.001863462 | 0.019767415 | 0.145,0.141 | 0.305,0.194 | -0.107 |
| 7 | XLOC_048071 | chrX | - | 122756955 | 122757822 | 122756955 | 122757134 | 122757637 | 503 | 122757822 | 7 | 227,233 | 164,183 | 302,292 | 117,140 | 652 | 149 | 0.000101474 | 0.002099556 | 0.24,0.225 | 0.371,0.323 | -0.114 |
| 297 | XLOC_038044 | chr6 | + | 157800198 | 157809091 | 157800198 | 157803298 | 157803814 | 516 | 157809091 | 297 | 44,70 | 5,0 | 74,41 | 0,0 | 665 | 149 | 0.003382954 | 0.030877668 | 0.663,1.0 | 1.0,1.0 | -0.168 |
| 6067 | XLOC_026782 | chr20 | + | 37075330 | 37076266 | 37075330 | 37075591 | 37076108 | 517 | 37076266 | 6067 | 30,21 | 3,2 | 44,34 | 0,0 | 666 | 149 | 0.000194958 | 0.003547609 | 0.691,0.701 | 1.0,1.0 | -0.304 |
| 3202 | XLOC_032226 | chr4 | + | 1243043 | 1246399 | 1243043 | 1244161 | 1244683 | 522 | 1246399 | 3202 | 102,124 | 26,21 | 256,249 | 12,8 | 671 | 149 | 1.14E-08 | 1.06E-06 | 0.466,0.567 | 0.826,0.874 | -0.334 |
| 3329 | XLOC_000753 | chr1 | + | 226032198 | 226033232 | 226032198 | 226032324 | 226032846 | 522 | 226033232 | 3329 | 153,171 | 200,188 | 429,382 | 181,190 | 671 | 149 | 4.60E-12 | 1.17E-09 | 0.145,0.168 | 0.345,0.309 | -0.17 |
| 5827 | XLOC_027790 | chr21 | - | 38444733 | 38445459 | 38444733 | 38444837 | 38445360 | 523 | 38445459 | 5827 | 42,33 | 6,11 | 48,25 | 0,1 | 672 | 149 | 2.85E-06 | 0.000116072 | 0.608,0.399 | 1.0,0.847 | -0.42 |
| 4454 | XLOC_041046 | chr7 | + | 27138692 | 27140058 | 27138692 | 27138858 | 27139397 | 539 | 27140058 | 4454 | 14,13 | 4,4 | 34,33 | 0,0 | 688 | 149 | 4.07E-08 | 3.29E-06 | 0.431,0.413 | 1.0,1.0 | -0.578 |
| 2260 | XLOC_014987 | chr15 | - | 75190467 | 75195124 | 75190467 | 75194404 | 75194944 | 540 | 75195124 | 2260 | 319,293 | 18,39 | 369,404 | 15,15 | 689 | 149 | 0.005072071 | 0.041919834 | 0.793,0.619 | 0.842,0.853 | -0.141 |
| 176 | XLOC_041435 | chr7 | - | 27168126 | 27181704 | 27168126 | 27169190 | 27169736 | 546 | 27181704 | 176 | 0,0 | 6,3 | 2,7 | 2,3 | 695 | 149 | 4.23E-05 | 0.001062518 | 0.0,0.0 | 0.177,0.333 | -0.255 |
| 4 | XLOC_048096 | chrX | - | 134184969 | 134186258 | 134184969 | 134185398 | 134185946 | 548 | 134186258 | 4 | 596,645 | 422,374 | 10,911,051 | 201,184 | 697 | 149 | 0 | 0 | 0.232,0.269 | 0.537,0.55 | -0.293 |
| 385 | XLOC_034900 | chr5 | - | 54518003 | 54518851 | 54518003 | 54518227 | 54518778 | 551 | 54518851 | 385 | 15,20 | 15,11 | 47,48 | 7,14 | 700 | 149 | 0.001978217 | 0.020750582 | 0.176,0.279 | 0.588,0.422 | -0.277 |
| 9264 | XLOC_016339 | chr16 | + | 28847253 | 28848525 | 28847253 | 28847497 | 28848048 | 551 | 28848525 | 9264 | 369,327 | 27,15 | 333,380 | 5,5 | 700 | 149 | 6.71E-06 | 0.000235865 | 0.744,0.823 | 0.934,0.942 | -0.154 |
| 9061 | XLOC_021326 | chr19 | + | 50381754 | 50382717 | 50381754 | 50381829 | 50382382 | 553 | 50382717 | 9061 | 47,50 | 3,2 | 27,14 | 0,0 | 702 | 149 | 0.001960661 | 0.020638389 | 0.769,0.841 | 1.0,1.0 | -0.195 |
| 3167 | XLOC_017914 | chr17 | + | 7761923 | 7768020 | 7761923 | 7762142 | 7762699 | 557 | 7768020 | 3167 | 61,78 | 84,91 | 102,124 | 65,39 | 706 | 149 | 3.16E-05 | 0.000838972 | 0.133,0.153 | 0.249,0.402 | -0.182 |
| 1771 | XLOC_000505 | chr1 | + | 150965676 | 150969350 | 150965676 | 150967175 | 150967735 | 560 | 150969350 | 1771 | 50,31 | 16,20 | 68,45 | 5,0 | 709 | 149 | 3.26E-06 | 0.000127805 | 0.396,0.246 | 0.741,1.0 | -0.55 |
| 11858 | XLOC_015047 | chr15 | - | 89876326 | 89878071 | 89876326 | 89877144 | 89877705 | 561 | 89878071 | 11858 | 46,29 | 3,5 | 52,48 | 0,0 | 710 | 149 | 3.22E-06 | 0.000126606 | 0.763,0.549 | 1.0,1.0 | -0.344 |
| 2757 | XLOC_014776 | chr15 | + | 91504254 | 91506740 | 91504254 | 91504284 | 91504847 | 563 | 91506740 | 2757 | 686,724 | 351,322 | 1,044,938 | 327,242 | 712 | 149 | 1.87E-05 | 0.000541046 | 0.29,0.32 | 0.401,0.448 | -0.119 |
| 2125 | XLOC_028670 | chr22 | - | 38694099 | 38694939 | 38694099 | 38694209 | 38694790 | 581 | 38694939 | 2125 | 13,171,464 | 133,159 | 13,081,277 | 30,44 | 730 | 149 | 2.22E-16 | 1.34E-13 | 0.669,0.653 | 0.899,0.856 | -0.216 |
| 2940 | XLOC_000699 | chr1 | + | 207940357 | 207941168 | 207940357 | 207940540 | 207941123 | 583 | 207941168 | 2940 | 12,181,293 | 279,363 | 28,362,421 | 313,282 | 732 | 149 | 6.08E-12 | 1.52E-09 | 0.471,0.42 | 0.648,0.636 | -0.197 |
| 15292 | XLOC_001334 | chr1 | - | 153905958 | 153906778 | 153905958 | 153906023 | 153906610 | 587 | 153906778 | 15292 | 109,88 | 8,0 | 154,136 | 1,0 | 736 | 149 | 0.000792704 | 0.010305837 | 0.734,1.0 | 0.969,1.0 | -0.117 |
| 9230 | XLOC_016177 | chr16 | + | 456714 | 457560 | 456714 | 456833 | 457424 | 591 | 457560 | 9230 | 10,25 | 1,6 | 24,26 | 0,0 | 740 | 149 | 5.45E-07 | 2.91E-05 | 0.668,0.456 | 1.0,1.0 | -0.438 |
| 2853 | XLOC_020911 | chr19 | + | 4501202 | 4502209 | 4501202 | 4501329 | 4501925 | 596 | 4502209 | 2853 | 50,62 | 42,48 | 93,100 | 36,49 | 745 | 149 | 0.004646214 | 0.039440383 | 0.192,0.205 | 0.341,0.29 | -0.117 |
| 1509 | XLOC_016525 | chr16 | + | 88740011 | 88741311 | 88740011 | 88740167 | 88740771 | 604 | 88741311 | 1509 | 6,9 | 4,0 | 11,10 | 0,0 | 753 | 149 | 0.002743528 | 0.026365349 | 0.229,1.0 | 1.0,1.0 | -0.385 |
| 355 | XLOC_035165 | chr5 | - | 177651642 | 177652430 | 177651642 | 177651730 | 177652355 | 625 | 177652430 | 355 | 62,60 | 102,74 | 109,52 | 53,52 | 774 | 149 | 0.001127896 | 0.013422314 | 0.105,0.135 | 0.284,0.161 | -0.102 |
| 16230 | XLOC_009618 | chr12 | - | 133403153 | 133405283 | 133403153 | 133403553 | 133404192 | 639 | 133405283 | 16230 | 33,37 | 16,8 | 25,16 | 2,0 | 788 | 149 | 0.000974543 | 0.012045252 | 0.281,0.467 | 0.703,1.0 | -0.477 |
| 12428 | XLOC_007509 | chr11 | - | 47598962 | 47600541 | 47598962 | 47599580 | 47600236 | 656 | 47600541 | 12428 | 5,4 | 1,1 | 8,6 | 1,0 | 805 | 149 | 0.003227742 | 0.029786313 | 0.481,0.425 | 0.597,1.0 | -0.346 |
| 3031 | XLOC_016775 | chr16 | - | 67863667 | 67865261 | 67863667 | 67864451 | 67865118 | 667 | 67865261 | 3031 | 77,72 | 88,104 | 147,192 | 56,71 | 816 | 149 | 4.30E-12 | 1.12E-09 | 0.138,0.112 | 0.324,0.331 | -0.203 |
| 8006 | XLOC_018641 | chr17 | - | 55943834 | 55944776 | 55943834 | 55943905 | 55944589 | 684 | 55944776 | 8006 | 22,29 | 6,0 | 13,21 | 0,0 | 833 | 149 | 0.00087872 | 0.011173042 | 0.396,1.0 | 1.0,1.0 | -0.302 |
| 3454 | XLOC_005363 | chr10 | - | 103866339 | 103868934 | 103866339 | 103868080 | 103868785 | 705 | 103868934 | 3454 | 619,607 | 81,57 | 766,775 | 53,51 | 854 | 149 | 0.003736763 | 0.033338308 | 0.571,0.65 | 0.716,0.726 | -0.11 |
| 609 | XLOC_029512 | chr3 | + | 101571934 | 101573527 | 101571934 | 101572707 | 101573421 | 714 | 101573527 | 609 | 14,16 | 25,37 | 44,19 | 17,23 | 863 | 149 | 0.001716746 | 0.018644978 | 0.088,0.069 | 0.309,0.125 | -0.139 |
| 9983 | XLOC_041043 | chr7 | + | 26241280 | 26242229 | 26241280 | 26241446 | 26242160 | 714 | 26242229 | 9983 | 89,65 | 1,4 | 149,144 | 0,1 | 863 | 149 | 0.005068043 | 0.041919834 | 0.939,0.737 | 1.0,0.961 | -0.142 |
| 3004 | XLOC_000907 | chr1 | - | 16944679 | 16946519 | 16944679 | 16945598 | 16946339 | 741 | 16946519 | 3004 | 16,35 | 50,38 | 55,47 | 18,26 | 890 | 149 | 1.07E-06 | 5.13E-05 | 0.051,0.134 | 0.338,0.232 | -0.193 |
| 2291 | XLOC_046145 | chr9 | + | 139745206 | 139746391 | 139745206 | 139745557 | 139746308 | 751 | 139746391 | 2291 | 13,5 | 11,6 | 55,41 | 16,10 | 900 | 149 | 0.001075093 | 0.012957966 | 0.164,0.121 | 0.363,0.404 | -0.241 |
| 2318 | XLOC_021921 | chr19 | - | 59080882 | 59081894 | 59080882 | 59080953 | 59081710 | 757 | 59081894 | 2318 | 109,106 | 96,113 | 226,175 | 102,84 | 906 | 149 | 6.86E-06 | 0.000239691 | 0.157,0.134 | 0.267,0.255 | -0.115 |
| 7328 | XLOC_021165 | chr19 | + | 37341336 | 37342806 | 37341336 | 37341870 | 37342629 | 759 | 37342806 | 7328 | 2,3 | 2,2 | 1,5 | 0,0 | 908 | 149 | 2.68E-07 | 1.59E-05 | 0.141,0.198 | 1.0,1.0 | -0.831 |
| 8009 | XLOC_018641 | chr17 | - | 55943834 | 55944776 | 55943834 | 55943905 | 55944698 | 793 | 55944776 | 8009 | 22,29 | 0,6 | 14,22 | 0,0 | 942 | 149 | 0.000632536 | 0.008661183 | 1.0,0.433 | 1.0,1.0 | -0.283 |
| 2645 | XLOC_016664 | chr16 | - | 21829976 | 21831764 | 21829976 | 21830802 | 21831615 | 813 | 21831764 | 2645 | 27,16 | 6,4 | 38,27 | 2,1 | 962 | 149 | 0.004463428 | 0.038255777 | 0.411,0.383 | 0.746,0.807 | -0.379 |
| 590 | XLOC_029664 | chr3 | + | 180326488 | 180327791 | 180326488 | 180326598 | 180327417 | 819 | 180327791 | 590 | 445,508 | 46,37 | 462,477 | 24,21 | 968 | 149 | 0.005550649 | 0.044912785 | 0.598,0.679 | 0.748,0.778 | -0.124 |
| 5215 | XLOC_029956 | chr3 | - | 108300954 | 108301930 | 108300954 | 108300985 | 108301823 | 838 | 108301930 | 5215 | 87,61 | 2,3 | 56,47 | 0,0 | 987 | 149 | 0.001699631 | 0.018472433 | 0.868,0.754 | 1.0,1.0 | -0.189 |
| 16086 | XLOC_034614 | chr5 | + | 87565061 | 87578683 | 87565061 | 87565504 | 87566342 | 838 | 87578683 | 16086 | 4,11 | 2,1 | 11,11 | 0,0 | 987 | 149 | 0.001376608 | 0.015616088 | 0.232,0.624 | 1.0,1.0 | -0.572 |
| 364 | XLOC_035102 | chr5 | - | 148871471 | 148876423 | 148871471 | 148873184 | 148874036 | 852 | 148876423 | 364 | 343,334 | 16,8 | 310,362 | 3,4 | 1001 | 149 | 0.001648036 | 0.018015807 | 0.761,0.861 | 0.939,0.931 | -0.124 |
| 503 | XLOC_032343 | chr4 | + | 68586139 | 68588555 | 68586139 | 68586339 | 68587194 | 855 | 68588555 | 503 | 37,34 | 1,9 | 71,35 | 1,0 | 1004 | 149 | 0.000226751 | 0.003984564 | 0.846,0.359 | 0.913,1.0 | -0.354 |
| 40 | XLOC_046502 | chr9 | - | 140469203 | 140470619 | 140469203 | 140469298 | 140470160 | 862 | 140470619 | 40 | 280,259 | 33,45 | 360,352 | 21,16 | 1011 | 149 | 1.44E-05 | 0.000435211 | 0.556,0.459 | 0.716,0.764 | -0.232 |
| 9438 | XLOC_014944 | chr15 | - | 65113565 | 65114590 | 65113565 | 65113596 | 65114464 | 868 | 65114590 | 9438 | 50,61 | 4,3 | 74,65 | 0,1 | 1017 | 149 | 0.00031807 | 0.005155618 | 0.647,0.749 | 1.0,0.905 | -0.255 |
| 522 | XLOC_037917 | chr6 | + | 88049833 | 88054575 | 88049833 | 88050055 | 88050943 | 888 | 88054575 | 522 | 63,68 | 7,2 | 89,65 | 0,0 | 1037 | 149 | 5.23E-06 | 0.000191994 | 0.564,0.83 | 1.0,1.0 | -0.303 |

| | | | | | | | | | | | | | | | | | | | | | | |
|---|---|---|---|---|---|---|---|---|---|---|---|---|---|---|---|---|---|---|---|---|---|---|
| 1583 | XLOC_016758 | chr16 | - | 67212101 | 67213465 | 67212101 | 67212407 | 67213304 | 897 | 67213465 | 1583 | 195,240 | 7,4 | 216,218 | 0,1 | 1046 | 149 | 0.000103605 | 0.002140687 | 0.799,0.895 | 1.0,0.969 | -0.137 |
| 2131 | XLOC_007597 | chr11 | - | 66133901 | 66135019 | 66133901 | 66134035 | 66134934 | 899 | 66135019 | 2131 | 268,266 | 143,125 | 384,306 | 108,85 | 1048 | 149 | 8.54E-05 | 0.001833049 | 0.21,0.232 | 0.336,0.339 | -0.117 |
| 833 | XLOC_029685 | chr3 | + | 184556501 | 184557540 | 184556501 | 184556534 | 184557485 | 951 | 184557540 | 833 | 20,14 | 16,9 | 66,47 | 16,13 | 1100 | 149 | 0.001607869 | 0.017666597 | 0.145,0.174 | 0.358,0.329 | -0.184 |
| 822 | XLOC_018271 | chr17 | + | 78084739 | 78085899 | 78084739 | 78084824 | 78085781 | 957 | 78085899 | 822 | 233,248 | 155,163 | 479,479 | 175,156 | 1106 | 149 | 2.63E-06 | 0.00010833 | 0.168,0.17 | 0.269,0.293 | -0.112 |
| 2659 | XLOC_011393 | chr13 | + | 115007595 | 115008823 | 115007595 | 115007755 | 115008731 | 976 | 115008823 | 2659 | 514,552 | 258,297 | 842,700 | 169,161 | 1125 | 149 | 4.12E-12 | 1.09E-09 | 0.209,0.198 | 0.398,0.365 | -0.178 |
| 1978 | XLOC_009388 | chr12 | - | 54104919 | 54106990 | 54104919 | 54105905 | 54106883 | 978 | 54106990 | 1978 | 432,431 | 116,127 | 716,697 | 41,59 | 1127 | 149 | 6.66E-16 | 3.85E-13 | 0.33,0.31 | 0.698,0.61 | -0.334 |
| 2160 | XLOC_021499 | chr19 | - | 6413123 | 6415312 | 6413123 | 6414131 | 6415151 | 1020 | 6415312 | 2160 | 19,221,731 | 61,95 | 39,984,001 | 80,90 | 1169 | 149 | 8.29E-05 | 0.001795144 | 0.801,0.699 | 0.864,0.85 | -0.107 |
| 2603 | XLOC_041436 | chr7 | - | 27201389 | 27204586 | 27201389 | 27203460 | 27204496 | 1036 | 27204586 | 2603 | 90,120 | 43,39 | 183,144 | 35,26 | 1185 | 149 | 0.000556342 | 0.007858199 | 0.208,0.279 | 0.397,0.411 | -0.161 |
| 2571 | XLOC_014712 | chr15 | + | 75644942 | 75646207 | 75644942 | 75645042 | 75646079 | 1037 | 75646207 | 2571 | 106,106 | 8,5 | 177,142 | 2,3 | 1186 | 149 | 0.004394625 | 0.037795279 | 0.625,0.727 | 0.917,0.856 | -0.211 |
| 3157 | XLOC_018763 | chr17 | - | 78183459 | 78186073 | 78183459 | 78184810 | 78185869 | 1059 | 78186073 | 3157 | 382,428 | 59,60 | 426,378 | 36,21 | 1208 | 149 | 0.000380922 | 0.005888823 | 0.444,0.468 | 0.593,0.689 | -0.185 |
| 1490 | XLOC_035181 | chr5 | - | 180272366 | 180278493 | 180272366 | 180275082 | 180276153 | 1071 | 180278493 | 1490 | 561,535 | 21,21 | 712,678 | 4,6 | 1220 | 149 | 1.25E-07 | 8.30E-06 | 0.765,0.757 | 0.956,0.932 | -0.183 |
| 2015 | XLOC_011561 | chr13 | - | 113978518 | 113980161 | 113978518 | 113978892 | 113979975 | 1083 | 113980161 | 2015 | 103,114 | 80,101 | 225,249 | 75,67 | 1232 | 149 | 1.62E-09 | 2.01E-07 | 0.135,0.12 | 0.266,0.31 | -0.161 |
| 14173 | XLOC_007142 | chr11 | + | 64852586 | 64854080 | 64852586 | 64852786 | 64853886 | 1100 | 64854080 | 14173 | 25,16 | 2,3 | 18,39 | 1,0 | 1249 | 149 | 0.000854974 | 0.010949508 | 0.599,0.389 | 0.682,1.0 | -0.347 |
| 2771 | XLOC_001135 | chr1 | - | 59154709 | 59155921 | 59154709 | 59154788 | 59155897 | 1109 | 59155921 | 2771 | 152,203 | 35,55 | 406,327 | 40,35 | 1258 | 149 | 3.56E-06 | 0.000137281 | 0.34,0.304 | 0.546,0.525 | -0.214 |
| 206 | XLOC_041217 | chr7 | + | 99950995 | 99952863 | 99950995 | 99951635 | 99952765 | 1130 | 99952863 | 206 | 107,130 | 126,102 | 343,268 | 76,77 | 1279 | 149 | 1.11E-16 | 7.26E-14 | 0.09,0.129 | 0.345,0.288 | -0.207 |
| 528 | XLOC_032657 | chr4 | - | 83343838 | 83347282 | 83343838 | 83346036 | 83347189 | 1153 | 83347282 | 528 | 26,172,600 | 623,596 | 34,953,592 | 500,457 | 1302 | 149 | 6.72E-07 | 3.39E-05 | 0.325,0.333 | 0.444,0.474 | -0.13 |
| 175 | XLOC_041438 | chr7 | - | 27228813 | 27238061 | 27228813 | 27233177 | 27234404 | 1227 | 27238061 | 175 | 21,20 | 4,0 | 14,16 | 0,0 | 1376 | 149 | 0.002497984 | 0.024542568 | 0.362,1.0 | 1.0,1.0 | -0.319 |
| 917 | XLOC_009559 | chr12 | - | 115117309 | 115118951 | 115117309 | 115117456 | 115118683 | 1227 | 115118951 | 917 | 231,207 | 116,117 | 487,535 | 158,138 | 1376 | 149 | 2.35E-05 | 0.000658335 | 0.177,0.161 | 0.25,0.296 | -0.104 |
| 6995 | XLOC_020849 | chr19 | + | 798014 | 799443 | 798014 | 798171 | 799412 | 1241 | 799443 | 6995 | 10,13 | 2,1 | 24,13 | 0,0 | 1390 | 149 | 0.002074555 | 0.021358975 | 0.349,0.582 | 1.0,1.0 | -0.534 |
| 7984 | XLOC_001539 | chr1 | - | 226109903 | 226111471 | 226109903 | 226110083 | 226111336 | 1253 | 226111471 | 7984 | 31,47 | 6,0 | 57,70 | 0,0 | 1402 | 149 | 0.000779767 | 0.010181646 | 0.354,1.0 | 1.0,1.0 | -0.323 |
| 9696 | XLOC_014760 | chr15 | + | 89824995 | 89826481 | 89824995 | 89825066 | 89826362 | 1296 | 89826481 | 9696 | 67,73 | 2,6 | 98,82 | 0,0 | 1445 | 149 | 2.70E-06 | 0.000110681 | 0.775,0.556 | 1.0,1.0 | -0.335 |
| 2388 | XLOC_005079 | chr10 | + | 124187791 | 124193261 | 124187791 | 124187832 | 124189139 | 1307 | 124193261 | 2388 | 38,64 | 14,16 | 138,122 | 8,13 | 1456 | 149 | 0.000174454 | 0.003267916 | 0.217,0.29 | 0.638,0.49 | -0.311 |
| 147 | XLOC_041749 | chr7 | - | 141356343 | 141364040 | 141356343 | 141362651 | 141363971 | 1320 | 141364040 | 147 | 236,176 | 67,83 | 204,230 | 45,43 | 1469 | 149 | 0.002047788 | 0.021199463 | 0.263,0.177 | 0.315,0.352 | -0.114 |
| 8555 | XLOC_021853 | chr19 | - | 52566421 | 52570863 | 52566421 | 52566425 | 52567746 | 1321 | 52570863 | 8555 | 8,17 | 2,1 | 15,20 | 0,0 | 1470 | 149 | 0.00200704 | 0.020930873 | 0.288,0.633 | 1.0,1.0 | -0.539 |
| 5470 | XLOC_029391 | chr3 | + | 42702976 | 42704619 | 42702976 | 42703124 | 42704465 | 1341 | 42704619 | 5470 | 1,0 | 0,2 | 7,5 | 0,0 | 1490 | 149 | 1.13E-05 | 0.000363808 | 1.0,0.0 | 1.0,1.0 | -0.5 |
| 5124 | XLOC_041515 | chr7 | - | 64465336 | 64467093 | 64465336 | 64465572 | 64466918 | 1346 | 64467093 | 5124 | 2,9 | 0,3 | 10,9 | 0,0 | 1495 | 149 | 0.002457601 | 0.024295557 | 1.0,0.23 | 1.0,1.0 | -0.385 |
| 11413 | XLOC_030086 | chr3 | - | 156868070 | 156869719 | 156868070 | 156868170 | 156869517 | 1347 | 156869719 | 11413 | 160,173 | 8,1 | 307,261 | 0,0 | 1496 | 149 | 2.15E-05 | 0.000610852 | 0.666,0.945 | 1.0,1.0 | -0.195 |
| 8873 | XLOC_016718 | chr16 | - | 46989257 | 46991132 | 46989257 | 46989534 | 46990918 | 1384 | 46991132 | 8873 | 311,267 | 10,15 | 379,387 | 0,2 | 1533 | 149 | 8.25E-09 | 8.11E-07 | 0.751,0.634 | 1.0,0.95 | -0.282 |
| 2178 | XLOC_009511 | chr12 | - | 104166091 | 104171859 | 104166091 | 104169408 | 104170821 | 1413 | 104171859 | 2178 | 139,162 | 3,2 | 129,111 | 0,0 | 1562 | 149 | 0.003165715 | 0.029339913 | 0.815,0.885 | 1.0,1.0 | -0.15 |
| 2522 | XLOC_013112 | chr14 | + | 50790662 | 50798961 | 50790662 | 50790878 | 50792326 | 1448 | 50798961 | 2522 | 84,68 | 3,2 | 86,111 | 0,0 | 1597 | 149 | 0.000366821 | 0.005729728 | 0.723,0.76 | 1.0,1.0 | -0.258 |
| 15758 | XLOC_000882 | chr1 | - | 10516563 | 10523266 | 10516563 | 10521759 | 10523221 | 1462 | 10523266 | 15758 | 158,171 | 6,0 | 215,231 | 0,0 | 1611 | 149 | 0.000787926 | 0.010261457 | 0.709,1.0 | 1.0,1.0 | -0.145 |
| 3419 | XLOC_004807 | chr10 | + | 30726104 | 30728203 | 30726104 | 30726335 | 30727844 | 1509 | 30728203 | 3419 | 52,52 | 64,77 | 46,68 | 29,13 | 1658 | 149 | 4.07E-05 | 0.001029275 | 0.068,0.057 | 0.125,0.32 | -0.16 |
| 5867 | XLOC_034684 | chr5 | + | 134124202 | 134125912 | 134124202 | 134124281 | 134125795 | 1514 | 134125912 | 5867 | 31,24 | 17,16 | 45,40 | 12,4 | 1663 | 149 | 0.004816185 | 0.040359362 | 0.14,0.118 | 0.251,0.473 | -0.233 |
| 52 | XLOC_046427 | chr9 | - | 130823531 | 130826179 | 130823531 | 130824105 | 130825652 | 1547 | 130826179 | 52 | 72,127 | 34,51 | 255,206 | 35,46 | 1696 | 149 | 6.74E-05 | 0.001521194 | 0.157,0.18 | 0.39,0.282 | -0.167 |
| 3288 | XLOC_028580 | chr22 | - | 20386516 | 20388425 | 20386516 | 20386675 | 20388299 | 1624 | 20388425 | 3288 | 21,13 | 2,1 | 60,42 | 0,0 | 1773 | 149 | 0.000260125 | 0.004446362 | 0.469,0.522 | 1.0,1.0 | -0.504 |
| 1034 | XLOC_023585 | chr2 | + | 97858717 | 97860491 | 97858717 | 97858790 | 97860462 | 1672 | 97860491 | 1034 | 8,7 | 11,18 | 11,12 | 7,4 | 1821 | 149 | 0.004769617 | 0.04008077 | 0.056,0.031 | 0.114,0.197 | -0.112 |
| 3548 | XLOC_048083 | chrX | - | 130919580 | 130928143 | 130919580 | 130926262 | 130928021 | 1759 | 130928143 | 3548 | 0,4 | 3,2 | 12,16 | 1,0 | 1908 | 149 | 7.76E-06 | 0.000264201 | 0.0,0.135 | 0.484,1.0 | -0.674 |
| 2099 | XLOC_030015 | chr3 | - | 126178495 | 126181063 | 126178495 | 126178580 | 126180377 | 1797 | 126181063 | 2099 | 699,639 | 57,36 | 831,684 | 23,11 | 1946 | 149 | 2.35E-06 | 9.85E-05 | 0.484,0.576 | 0.734,0.826 | -0.25 |
| 263 | XLOC_038373 | chr6 | - | 99850415 | 99852578 | 99850415 | 99850586 | 99852478 | 1892 | 99852578 | 263 | 24,452,523 | 71,43 | 21,982,111 | 25,20 | 2041 | 149 | 0.000208763 | 0.003742804 | 0.715,0.811 | 0.865,0.885 | -0.112 |
| 15005 | XLOC_021620 | chr19 | - | 19102148 | 19104549 | 19102148 | 19102551 | 19104446 | 1895 | 19104549 | 15005 | 172,157 | 5,1 | 192,153 | 0,0 | 2044 | 149 | 0.000159147 | 0.003030244 | 0.715,0.92 | 1.0,1.0 | -0.182 |
| 2236 | XLOC_009218 | chr12 | - | 6571410 | 6574107 | 6571410 | 6572135 | 6574055 | 1920 | 6574107 | 2236 | 399,434 | 4,5 | 289,253 | 0,0 | 2069 | 149 | 0.00021668 | 0.003855951 | 0.878,0.862 | 1.0,1.0 | -0.13 |
| 454 | XLOC_041145 | chr7 | + | 75042066 | 75044301 | 75042066 | 75042210 | 75044162 | 1952 | 75044301 | 454 | 475,513 | 52,53 | 658,649 | 27,17 | 2101 | 149 | 1.13E-07 | 7.68E-06 | 0.393,0.407 | 0.633,0.73 | -0.281 |
| 4409 | XLOC_041489 | chr7 | - | 44116107 | 44118410 | 44116107 | 44116354 | 44118338 | 1984 | 44118410 | 4409 | 6,9 | 4,1 | 19,14 | 0,0 | 2133 | 149 | 7.01E-10 | 1.01E-07 | 0.095,0.386 | 1.0,1.0 | -0.76 |
| 15570 | XLOC_001218 | chr1 | - | 109468392 | 109477564 | 109468392 | 109475123 | 109477241 | 2118 | 109477564 | 15570 | 75,53 | 2,9 | 73,71 | 0,0 | 2267 | 149 | 6.17E-07 | 3.18E-05 | 0.711,0.279 | 1.0,1.0 | -0.505 |
| 913 | XLOC_023912 | chr2 | + | 241500746 | 241503936 | 241500746 | 241500940 | 241503090 | 2150 | 241503936 | 913 | 66,71 | 3,1 | 86,84 | 0,0 | 2299 | 149 | 0.001482182 | 0.016518693 | 0.588,0.821 | 1.0,1.0 | -0.296 |
| 14038 | XLOC_021000 | chr19 | + | 13261200 | 13265742 | 13261200 | 13261507 | 13263757 | 2250 | 13265742 | 14038 | 124,109 | 0,5 | 174,133 | 0,0 | 2399 | 149 | 0.001178263 | 0.01385726 | 1.0,0.575 | 1.0,1.0 | -0.213 |
| 14902 | XLOC_001012 | chr1 | - | 35643589 | 35652723 | 35643589 | 35650194 | 35652547 | 2353 | 35652723 | 14902 | 116,112 | 21,22 | 151,169 | 2,6 | 2502 | 149 | 4.21E-07 | 2.37E-05 | 0.248,0.233 | 0.818,0.627 | -0.482 |
| 524 | XLOC_037689 | chr6 | + | 26428870 | 26432480 | 26428870 | 26429013 | 26431499 | 2486 | 26432480 | 524 | 128,140 | 2,5 | 155,157 | 0,1 | 2635 | 149 | 0.0003595 | 0.005675864 | 0.784,0.613 | 1.0,0.899 | -0.251 |
| 3980 | XLOC_046188 | chr9 | - | 14677060 | 14680160 | 14677060 | 14677498 | 14680030 | 2532 | 14680160 | 3980 | 3,5 | 0,4 | 2,2 | 0,0 | 2681 | 149 | 0.00012026 | 0.002411943 | 1.0,0.065 | 1.0,1.0 | -0.468 |
| 1232 | XLOC_000461 | chr1 | + | 145281954 | 145293580 | 145281954 | 145290511 | 145293113 | 2602 | 145293580 | 1232 | 381,433 | 24,17 | 945,734 | 9,19 | 2751 | 149 | 0.000896484 | 0.011331863 | 0.462,0.58 | 0.85,0.677 | -0.243 |
| 13255 | XLOC_001307 | chr1 | - | 150944087 | 150947366 | 150944087 | 150944437 | 150947094 | 2657 | 150947366 | 13255 | 1,1 | 0,2 | 10,2 | 0,0 | 2806 | 149 | 1.17E-05 | 0.000374604 | 1.0,0.026 | 1.0,1.0 | -0.487 |
| 64 | XLOC_046278 | chr9 | - | 74975543 | 74978522 | 74975543 | 74975703 | 74978385 | 2682 | 74978522 | 64 | 642,573 | 203,227 | 714,678 | 102,116 | 2831 | 149 | 1.63E-09 | 2.01E-07 | 0.143,0.117 | 0.269,0.235 | -0.122 |
| 12413 | XLOC_020127 | chr18 | + | 59942606 | 59945686 | 59942606 | 59942706 | 59945448 | 2742 | 59945686 | 12413 | 2,4 | 2,0 | 2,6 | 0,0 | 2891 | 149 | 0.00057956 | 0.008079468 | 0.049,1.0 | 1.0,1.0 | -0.476 |
| 1247 | XLOC_013404 | chr14 | - | 59937884 | 59942724 | 59937884 | 59939808 | 59942586 | 2778 | 59942724 | 1247 | 240,255 | 43,53 | 438,469 | 42,44 | 2927 | 149 | 0.000122114 | 0.002429626 | 0.221,0.197 | 0.347,0.352 | -0.14 |
| 1190 | XLOC_020232 | chr18 | - | 47794348 | 47799108 | 47794348 | 47796188 | 47799046 | 2858 | 47799108 | 1190 | 766,806 | 47,29 | 10,481,086 | 19,25 | 3007 | 149 | 0.000654448 | 0.008916852 | 0.447,0.579 | 0.732,0.683 | -0.195 |
| 14772 | XLOC_045867 | chr9 | + | 6493825 | 6497360 | 6493825 | 6493932 | 6496959 | 3027 | 6497360 | 14772 | 150,131 | 5,7 | 236,195 | 4,0 | 3176 | 149 | 0.004560712 | 0.038934297 | 0.585,0.468 | 0.735,1.0 | -0.341 |
| 13616 | XLOC_018612 | chr17 | - | 46629388 | 46632980 | 46629388 | 46629849 | 46632892 | 3043 | 46632980 | 13616 | 2,1 | 0,2 | 6,2 | 0,0 | 3192 | 149 | 3.11E-05 | 0.00082753 | 1.0,0.023 | 1.0,1.0 | -0.489 |
| 2657 | XLOC_041202 | chr7 | + | 99074025 | 99085371 | 99074025 | 99074103 | 99077283 | 3180 | 99085371 | 2657 | 300,273 | 37,31 | 338,265 | 5,5 | 3329 | 149 | 4.69E-11 | 9.30E-09 | 0.266,0.283 | 0.752,0.703 | -0.453 |
| 642 | XLOC_029298 | chr3 | + | 9471510 | 9475634 | 9471510 | 9471735 | 9475053 | 3318 | 9475634 | 642 | 172,179 | 30,35 | 324,316 | 9,12 | 3467 | 149 | 3.73E-11 | 7.79E-09 | 0.198,0.18 | 0.607,0.531 | -0.38 |
| 5838 | XLOC_027757 | chr21 | - | 33735464 | 33739064 | 33735464 | 33735638 | 33738962 | 3324 | 33739064 | 5838 | 88,91 | 4,0 | 150,134 | 0,0 | 3473 | 149 | 0.003509129 | 0.031759517 | 0.486,1.0 | 1.0,1.0 | -0.257 |
| 5837 | XLOC_027757 | chr21 | - | 33735464 | 33739064 | 33735464 | 33735581 | 33738926 | 3345 | 33739064 | 5837 | 82,94 | 6,0 | 129,130 | 0,0 | 3494 | 149 | 0.000645484 | 0.008810676 | 0.368,1.0 | 1.0,1.0 | -0.316 |
| 14486 | XLOC_009610 | chr12 | - | 125267228 | 125271049 | 125267228 | 125267357 | 125270986 | 3629 | 125271049 | 14486 | 14,581,497 | 36,53 | 14,061,302 | 25,20 | 3778 | 149 | 0.005946249 | 0.047125124 | 0.615,0.527 | 0.689,0.72 | -0.133 |
| 2911 | XLOC_000386 | chr1 | + | 100609624 | 100613882 | 100609624 | 100609699 | 100613449 | 3750 | 100613882 | 2911 | 819,706 | 36,40 | 16,831,432 | 17,16 | 3899 | 149 | 2.45E-12 | 6.83E-10 | 0.465,0.403 | 0.791,0.774 | -0.348 |
| 3346 | XLOC_032661 | chr4 | - | 83816844 | 83821376 | 83816844 | 83816927 | 83821229 | 4302 | 83821376 | 3346 | 902,805 | 26,35 | 903,927 | 17,11 | 4451 | 149 | 0.001588753 | 0.017482093 | 0.537,0.435 | 0.64,0.738 | -0.203 |
| 641 | XLOC_029298 | chr3 | + | 9470506 | 9475246 | 9470506 | 9470693 | 9475053 | 4360 | 9475246 | 641 | 229,242 | 19,24 | 397,383 | 14,8 | 4509 | 149 | 9.00E-05 | 0.001903505 | 0.285,0.25 | 0.484,0.613 | -0.281 |

| | | | | | | | | | | | | | | | | | | | | | | |
|---|---|---|---|---|---|---|---|---|---|---|---|---|---|---|---|---|---|---|---|---|---|---|
| 2121 | XLOC_011529 | chr13 | - | 79884552 | 79894845 | 79884552 | 79889128 | 79893669 | 4541 | 79894845 | 2121 | 10,991,077 | 12,22 | 13,811,396 | 1,4 | 4690 | 149 | 7.56E-08 | 5.45E-06 | 0.744,0.609 | 0.978,0.917 | -0.271 |
| 3006 | XLOC_000907 | chr1 | - | 16944679 | 16952500 | 16944679 | 16946519 | 16952269 | 5750 | 16952500 | 3006 | 266,244 | 41,30 | 495,519 | 39,29 | 5899 | 149 | 0.000362258 | 0.005701256 | 0.141,0.17 | 0.243,0.311 | -0.122 |
| 2919 | XLOC_034863 | chr5 | - | 34159948 | 34176062 | 34159948 | 34169258 | 34175549 | 6291 | 34176062 | 2919 | 281,218 | 17,18 | 391,344 | 8,6 | 6440 | 149 | 3.18E-05 | 0.000843346 | 0.277,0.219 | 0.531,0.57 | -0.302 |
| 595 | XLOC_029597 | chr3 | + | 140681968 | 140689835 | 140681968 | 140682069 | 140689768 | 7699 | 140689835 | 595 | 26,602,571 | 163,161 | 41,243,705 | 66,97 | 7848 | 149 | 1.44E-09 | 1.84E-07 | 0.237,0.233 | 0.543,0.42 | -0.247 |
| 10933 | XLOC_014966 | chr15 | - | 70358352 | 70366946 | 70358352 | 70358554 | 70366841 | 8287 | 70366946 | 10933 | 280,255 | 8,6 | 277,274 | 0,1 | 8436 | 149 | 2.35E-06 | 9.85E-05 | 0.382,0.429 | 1.0,0.829 | -0.509 |
